# Proteome-Wide Discovery of Degradable Proteins Using Bifunctional Molecules

**DOI:** 10.1101/2025.03.21.644652

**Authors:** Ines Forrest, Louis P. Conway, Appaso M. Jadhav, Clara Gathmann, Tzu-Yuan Chiu, Christian M. Chaheine, Michelle Estrada, Anurupa Shrestha, Justin M. Reitsma, Scott E. Warder, Anil Vasudevan, Shaun M. McLoughlin, Christopher G. Parker

**Affiliations:** The Scripps Research Institute – La Jolla, CA; Skaggs Graduate School of Chemical and Biological Sciences – La Jolla, CA; Technology and Therapeutic Platforms, Discovery Research. AbbVie Inc – North Chicago, IL

**Keywords:** targeted protein degradation, PROTACs, chemical probes, chemical proteomics, induced proximity, mass spectrometry

## Abstract

Targeted protein degradation (TPD) is an emergent therapeutic strategy with the potential to circumvent challenges associated with targets unamenable to conventional pharmacological inhibition. Among TPD approaches, Proteolysis Targeting Chimeras (PROTACs) have shown marked advancement with numerous candidates in clinical development. Despite their potential, most PROTACs utilize advanced small molecule inhibitors, inherently limiting the scope of this approach. More generally, the fraction of the proteome tractable to PROTAC-type strategies is unknown. Here, we describe a chemical proteomic strategy for the agnostic discovery of degradable human proteins in cells using a new class of bifunctional degrader molecules called “AgnoTACs”. Proteome-wide screening of 72 AgnoTACs in human cells uncovered downregulation events spanning >50 functionally and structurally diverse proteins, most of which lack chemical probes. Our findings highlight the potential of function-biased chemical libraries coupled with proteomic profiling to discover degrader starting points as well as furnish a blueprint for expanding our understanding of the chemically degradable proteome.

## INTRODUCTION

The human genome encodes more than 20,000 proteins, yet only a small fraction has useful chemical probes available to investigate their function^1^. Conventional chemical probe and drug discovery efforts have focused mainly on the identification of active/functional site inhibitors. Although this approach is highly successful for target protein classes such as enzymes and receptors, it is inevitably limited to proteins possessing functionally well-defined, druggable sites^2^. By harnessing the cell’s natural degradation machinery to eliminate proteins, targeted protein degradation (TPD) methods, such as proteolysis targeting chimeras (PROTACs), offer a compelling alternative^3, 4^. PROTACs are heterobifunctional molecules comprising a target binding moiety (TBM) linked to an E3 ligase binding moiety (EBM), which induce proximity between an E3 ligase and protein of interest (POI), facilitating directed ubiquitination of POIs leading to their subsequent degradation^5, 6^. PROTACs possess distinct advantages over conventional small molecule inhibition, including i) a catalytic mechanism of action (MoA), ii) the potential for increased selectivity owing to mandatory ternary complex formation, iii) the ability to engage a target protein potentially anywhere on its surface rather than being limited to functional sites, and iv) the ability to disrupt multiple protein functions (e.g., scaffolding roles)^7, 8^. Not surprisingly, PROTACs have now been developed for a multitude of disease areas and have shown promising preliminary results in clinical trials ^5, 9, 10^. However, despite their potential, the vast majority of PROTACs utilize pre-existing small molecule inhibitors as TBMs, including all disclosed structures of those currently in clinical development. For instance, ARV-471, an estrogen receptor (ER) degrader, is based on the known ER antagonist Lasofoxifene^11^. Similarly, ARV-110 which degrades the androgen receptor (AR) is derived from enzalutamide AR inhibitor scaffolds^12^, and KT-333 that targets STAT3-dependent pathologies, originates from SI-109, a STAT3 SH2 domain inhibitor^13^. This reliance inherently restricts PROTAC scope to targets possessing well-characterized, advanced ligands.

In this vein, powerful ‘binding-first’ screening approaches, including DNA-encoded libraries (DELs) and chemical proteomics, have demonstrated potential to furnish small molecules that can be co-opted for TPD. For example, DEL-based and other high-throughput synthesis-coupled screens have recently been deployed for the discovery of molecular glues^14-20^. Chemical proteomic screening has broadened the available repertoire of E3 ligase components^21-27^, enabled the development of PROTACs using covalent ligands as TBMs^28^, and have enabled the discovery of molecular glues^29-31^. Despite the potential of these approaches, to date, they have largely been limited to the targeting pre-defined proteins, the identification of IMiD-induced CRBN neosubstrates, or for the discovery of new covalent E3 recruiting ligands. More generally, the operational path to convert ‘binders’ into functional compounds is often not straightforward. Critical challenges include pre-requisite knowledge of a given pocket to accommodate modified ligands for proximity induction with concomitant effector proteins. Additionally, in the context of TPD, ligand binding itself may influence protein stability through a variety of mechanisms, potentially complicating differentiation of binders and designed degraders^32, 33 34^. Finally, since many of these screening approaches are often target specific, they require target-dependent readouts and available ligands that can accommodate further synthetic derivatization.

Inspired by these concepts, we report a chemical proteomic strategy that integrates target-agnostic libraries of small molecules that are functionally biased towards protein degradation with quantitative proteomics to discover degradable proteins *de novo*. With this goal in mind, we designed a series of 72 target-<u>agno</u>stic PRO<u>TACs</u>, or AgnoTACs, composed of Cereblon (CRBN) ligase ligands^10^, chemically linked to structurally diverse, drug-like small molecules. Proteome-wide profiling of this library in human cancer cells revealed that AgnoTACs can mediate the degradation of 50+ proteins, including enzymes, transcription factors, adaptor proteins, transporters, regulatory proteins and protein complexes. Mechanistic investigations demonstrated that AgnoTAC degradation events often proceed through expected CRBN- and proteasome-mediated pathways, though alternative degradation pathways were also encountered. We observed that chemical features of AgnoTACs, including linker length and composition, not only impact apparent potency of degradation for given targets but also proteome-wide selectivity. Further, preliminary studies highlight the potential of AgnoTACs to serve as ‘pathfinder’ chemical tools for functional investigations of proteins. These results underline how proteome-wide profiling of target-agnostic, yet function-focused chemical libraries can expedite the discovery of small molecule degraders for a broad spectrum of human proteins.

## RESULTS

### Proteome-Wide Profiling of AgnoTACs in Human Cells

Recent investigations of PROTACs based upon promiscuous kinase and HDAC inhibitors have provided valuable insights into principles governing TPD^35-37^. Specifically, these studies revealed that key factors traditionally associated with successful small-molecule inhibition, such as high target affinity and cellular target engagement (TE), do not reliably predict degradation efficiency or selectivity. Motivated by these findings, we rationalized that PROTAC libraries composed of relatively low molecular weight, drug-like ligands lacking pre-defined targets could serve as starting points to achieve degradation of proteins.

Towards this end, we designed and synthesized a library of 72 AgnoTAC molecules composed of thalidomide-based CRBN ligands conjugated to 18 structurally diverse small molecules possessing drug-like properties through four different linkers (**Fig. 1a, Fig. S1, Fig. S2**). ^38^ AgnoTACs were first evaluated for their impact on cell viability in human MDA-MB-231 triple-negative breast cancer cells, as highly cytotoxic compounds could impact protein abundances through indirect mechanisms. Overall, we observed minimal viability effects for most library members (**Fig. S3**).

**Fig. 1.**
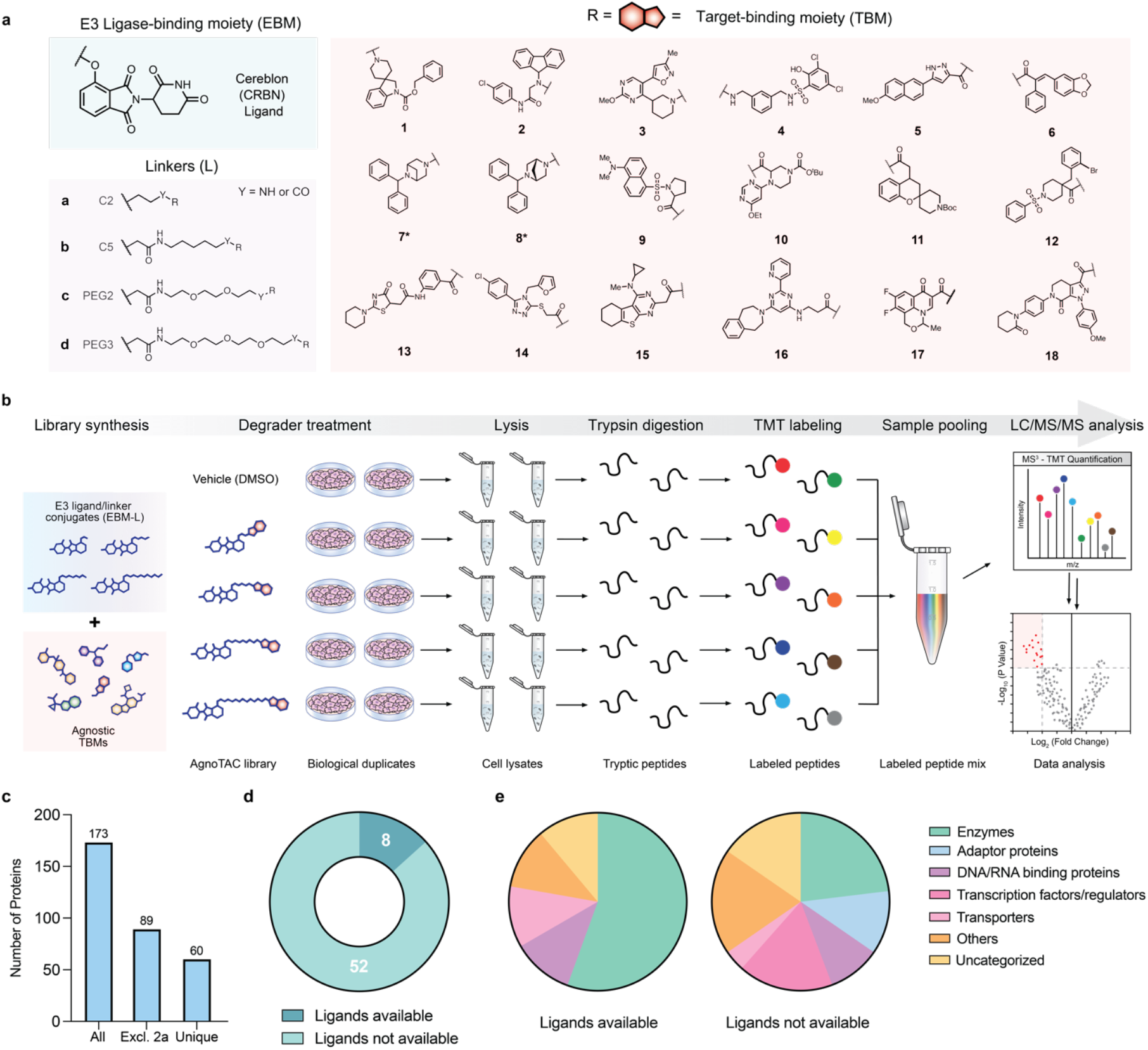
Chemical proteomic strategy to interrogate the degradable proteome. (**a**) Chemical structures of 72 AgnoTACs designed with CRBN E3 ligand, PEG or aliphatic linkers, and 18 diverse target-binding headgroups. The asterisk denotes an aryl ether linker instead of an oxyacetamide group. (**b**) Chemical proteomic workflow to identify degraded proteins. Cells are incubated with AgnoTAC library members (18 h treatment), followed by lysis, tryptic digestion, labeling with tandem mass tags (TMT), and protein identification and quantitation by LC-MS analysis (**Supplemental Dataset 1**). (**c**) Number of downregulated targets, defined as proteins displaying >2-fold decrease in MS^3^ abundance relative to DMSO and p-value < 0.05 across duplicated experiments, (excl. = excluding). (**d, e**) Categorization of AgnoTAC-downregulated targets based on existence of ligands in DrugBank and protein class distribution (associated data can be found in **Supplemental Dataset 2**).

We next employed quantitative proteomics to identify potential targets by measuring changes in protein abundance in response to treatment with each AgnoTAC. Given that library members exhibited minimal cytotoxicity, we sought to maximize the discovery of potential degradable proteins. F01-F09 AgnoTACs, which predominantly feature fragment-like head groups were screened at 100 µM, while F10-F18 degraders, containing higher molecular weight head groups, were tested at 50 µM. Adopting a TMT-based proteomics workflow^39^ (**Fig. 1b**), we quantified 8037 unique proteins, among which 173 showed substantial decreases in abundance (>2-fold relative to DMSO, and *p*-value < 0.05 across duplicated experiments, **Fig. 1c, Fig. S4, Supplemental Dataset 1**). We noted that **2a** downregulated a relatively large number of targets (i.e., ∼85), which we attributed to potential secondary effects emanating from its apparent cytotoxicity (**Fig. S3b**). To mitigate the inclusion of false positives (e.g., indirect abundance changes related to cytotoxicity) we excluded this AgnoTAC from our analysis, resulting in 60 uniquely downregulated targets (**Fig. 1c, Fig. S5a, Supplemental Dataset 1**). Despite the structurally simple TBMs, we observed each AgnoTAC to be remarkably selective, with the vast majority inducing downregulation of a single protein (**Fig. S5a, Supplemental Dataset 1**). Additionally, we identified cores (e.g., F02, F07, F08, F12, F15) which induce broader proteomic effects, while some were found to minimally affect protein abundances (e.g., F04, F09, F13, F14, F16-F18). We observed that a significant portion of targets fall out of traditional “druggable” classes, including adaptor proteins and transcription factors (**Fig. 1d, e**) with only a small fraction (∼13%) possessing known functional ligands (largely enzymes) as estimated by their presence or absence in the DrugBank database (**Fig. 1d, e, Fig. S5b, c**). In contrast, the larger subset of targets (∼87%) lacking reported bioactive ligands showed a broader functional distribution, with a reduced fractional representation of enzymes counterbalanced by expanded coverage of DNA/RNA binding proteins, transcription factors/regulators, and functionally uncategorized proteins (**Fig. 1d, e, Fig. S5b, c)**.

### Validation of Targeted Degradation of Identified Proteins

We recognize that changes in protein abundance upon AgnoTAC treatment may not necessarily be reflective of expected TPD mechanisms. We first determined half-maximal degradation concentration (DC_50_) ranges of 12.5 – 28.5 µM and maximal degradation (D_max_) ranges of 54-74% for a select set of AgnoTAC-target pairs, including: **1c**-UFD1, **2d**-PLOD2, **7b\***-CHCHD2 and **10d**-BRD2 (**Fig. 2**). We next sought to assess whether targets were being downregulated through a Cullin Ring ligase (CRL)-dependent mechanism^40^. MDA-MB-231 cells were treated with increasing concentrations of AgnoTACs, MG132 proteasome inhibitor (10 µM), or MLN4924 neddylation inhibitor (1 µM) to inhibit CRL activity^41, 42^, and protein abundance was monitored for a subset of identified downregulated targets. Co-incubation of **1c** and MG132 or **1c** and MLN4924 resulted in marked rescue of the ubiquitin recognition factor UFD1, an adaptor protein involved the ubiquitin-dependent proteolytic pathway^43^, suggesting that **1c** promotes proteasomal degradation of UFD1 via the action of a CRL in a dose-dependent fashion (**Fig. 2a-c**). We observed similar results upon cell treatment with AgnoTACs **2d, 7b\*** and **10d** degrading the PLOD2 lysyl hydroxylase^44^ (**Fig. 2d-f**), Parkinson’s disease associated transcription factor CHCHD2^45^ (**Fig. 2g-i**), and the chromatin reader protein BRD2^46^ (**Fig. 2j-l**) respectively. We observed that for a subset of targets, such as CHCHD2, protein levels were influenced by MG132 treatment alone, potentially convoluting interpretation of rescue experiments. In such instances, we relied on protein rescue through neddylation inhibition, which did not affect protein levels, to confirm degradation mechanism. In addition to high-confidence targets, we also observed proteins that fall outside our strict significance cut-offs but display trends of decreased protein abundances. For instance, Sirtuin 1 deacetylase (SIRT1)^47^ exhibited trends of downregulation upon treatment with compound **5a** but did not meet the statistical thresholds in our chemical proteomic analysis (**Fig. S6a, b**). We confirmed its degradation via the proteasome in a dose-dependent manner (**Fig. S6c, d**), suggesting these proteins could be *bona fide* AgnoTAC targets, and should be evaluated on a case-by-case basis. Collectively, this data supports that AgnoTACs promote dose-dependent CRL-mediated proteasomal degradation for a wide variety of proteins.

**Fig. 2.**
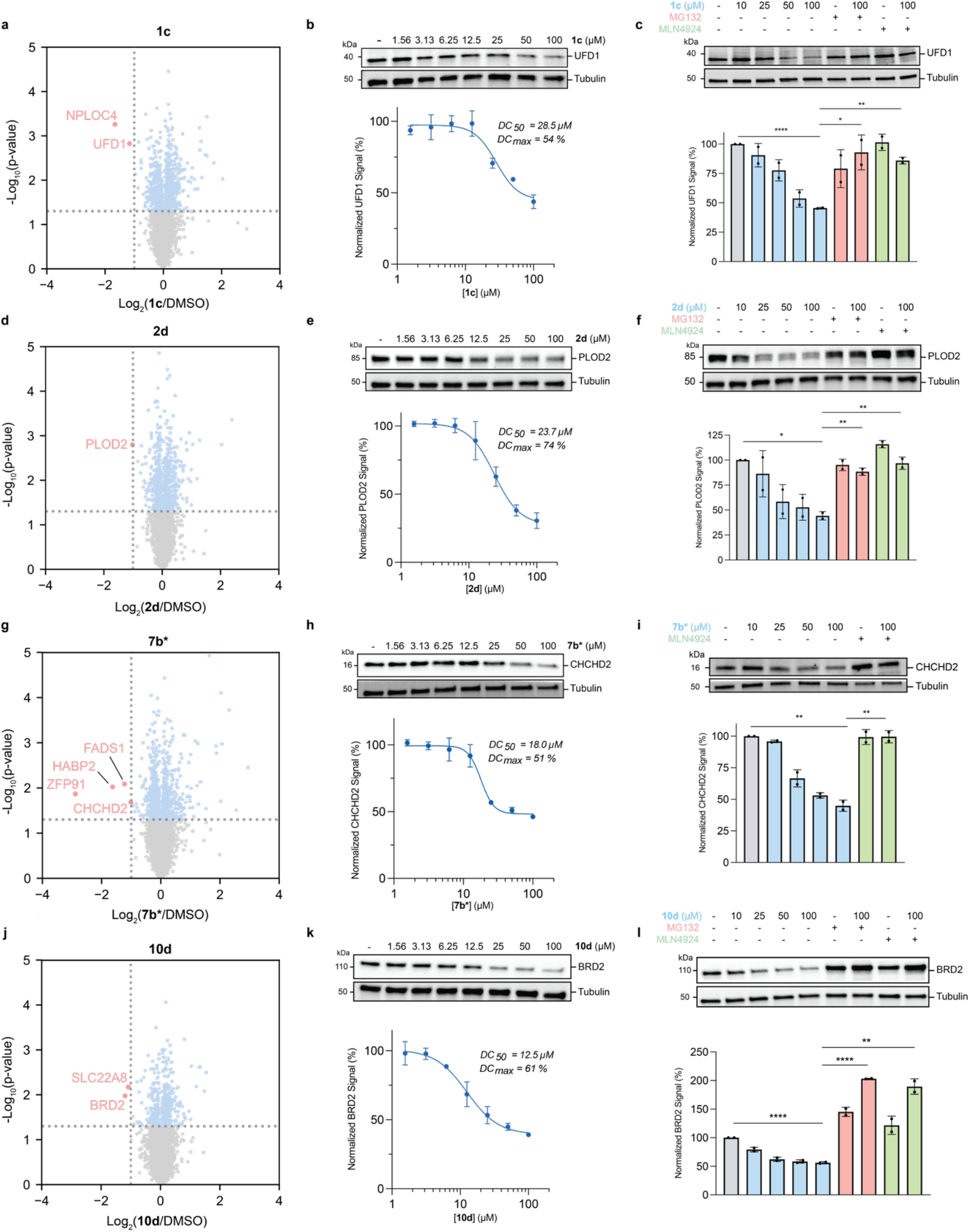
Targets degraded by AgnoTACs. Representative validation of select targets. MDA-MB-231 cells were treated with various AgnoTACs for 18 h, including (**a-c**) **1c**, (**d-f**) **2d**, (**g-i**) **7c\***, and (**j-l**) **10d**. Scatter plots depict relative fold change in protein abundance following treatment of MDA-MB-231 cells with probes. The x-axis represents log_2_(fold change) in protein expression. The y-axis represents statistical significance (p-value). Downregulated proteins are highlighted in red. Dashed lines represent thresholds (fold change >2, p-value <0.05). Data represents n = 2 biologically independent experiments. Associated proteomics data is provided in **supplemental dataset 1**. Validation by immunoblot analysis was performed by pre-incubating cells with MG132 (10 µM) or MLN4924 (1 µM) for 2 h, and subsequently co-incubating with probes for 18 h.

In addition to the expected canonical mechanism described above, we also observed intriguing instances where AgnoTACs induced proteasomal degradation of targets independently of CRL activity. While CRBN is typically associated with Cullin-mediated ubiquitination, these findings suggest the existence of alternative degradation pathways mediated by AgnoTACs. For example, degradation of the oncogenic transcription factor ETS-1^48^ by **5a** (**Fig. 3a-c**) and the functionally uncharacterized transmembrane protein TMEM205^49^ by **8d\*** (**Fig. 3d-f**) were dependent upon proteasomal activity but were surprisingly unaffected by co-treatment with MLN4924. We speculate that such degradation events could result from indirect effects, protein destabilization or alternative yet-to-be-characterized mechanisms. Nonetheless, these findings emphasize the need for thorough validation experiments to fully account for distinct mechanisms driving small molecule-mediated protein degradation.

**Fig. 3.**
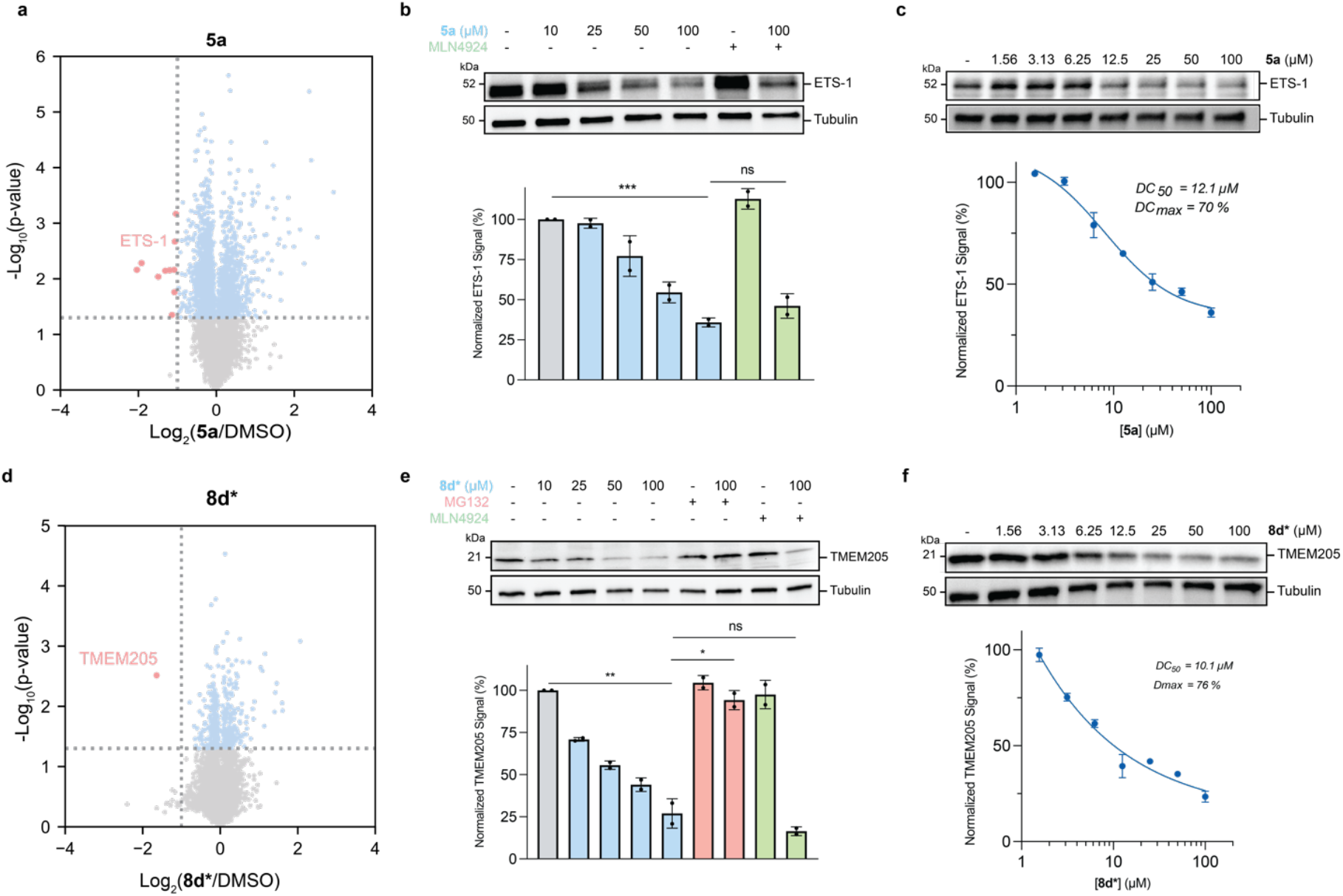
Non-canonical AgnoTAC-induced degradation events. (**a**) Volcano plot depicting global protein abundance changes upon **5a** treatment of MDA-MB-231 cells for 18 h. Vertical dashed line represents a fold change (FC) threshold of 2, while horizontal dashed lines denote a p-value threshold of 0.05. Statistical significance was assessed using a two-tailed Student’s t-test. Red dots indicate targets that meet both significance thresholds. (**b-f**) Dose-dependent proteasome-mediated and non-neddylation-dependent degradation of (**b, c**) ETS-1 transcription factor by **5a**, and (**d-f**) TMEM205 transmembrane protein by **8d\***. For rescue experiments, MDA-MB-231 cells are pre-treated with MG132 (10 µM) or MLN4924 (1 µM) and subsequently co-incubated with probes for 18 h. Data are represented as mean ± SD (n = 2 biological replicates). Statistical significance was determined using a two-tailed Student’s t-test (* *p* <0.05; ** *p* <0.01; *** *p* <0.001; ns = not significant).

As noted above, **2a** significantly impacted cell viability and our first pass proteomics profiling revealed a substantial number of proteins with altered abundance (**Fig. 4a**). Interestingly, most downregulated targets were mitochondrial proteins and components of the oxidative phosphorylation (OXPHOS)^50^ pathway (**Fig. 4b, Fig. S7**). Notably, co-incubation with MLN4924 substantially rescued **2a**-induced cytotoxicity (**Fig. 4c**), suggesting toxicity may be partly driven by an AgnoTAC-mediated degradation event. To investigate further, we monitored proteome-wide changes over 2, 4, 6, and 12 hours, revealing mitochondrial carrier homolog 2 (MTCH2) protein, an outer mitochondrial membrane insertase^51^ as the protein first affected by compound **2a**. MTCH2 was the only downregulated protein detected within the first 2 hours of treatment and continued to be reduced at all subsequent time points (**Fig. 4d**). Rescue experiments confirmed that MTCH2 degradation occurred dose-dependently and via proteasome and neddylation-mediated pathways (**Fig. 4e**). Recent reports have implicated MTCH2 in maintaining mitochondrial homeostasis and regulating apoptosis^52, 53^, suggesting that its degradation by **2a** may drive the observed cytotoxic effects. However, further investigation is needed to clarify whether MTCH2 degradation is the primary mechanism of action or if other mitochondrial disturbances also contribute to **2a**’s cytotoxicity.

**Fig. 4.**
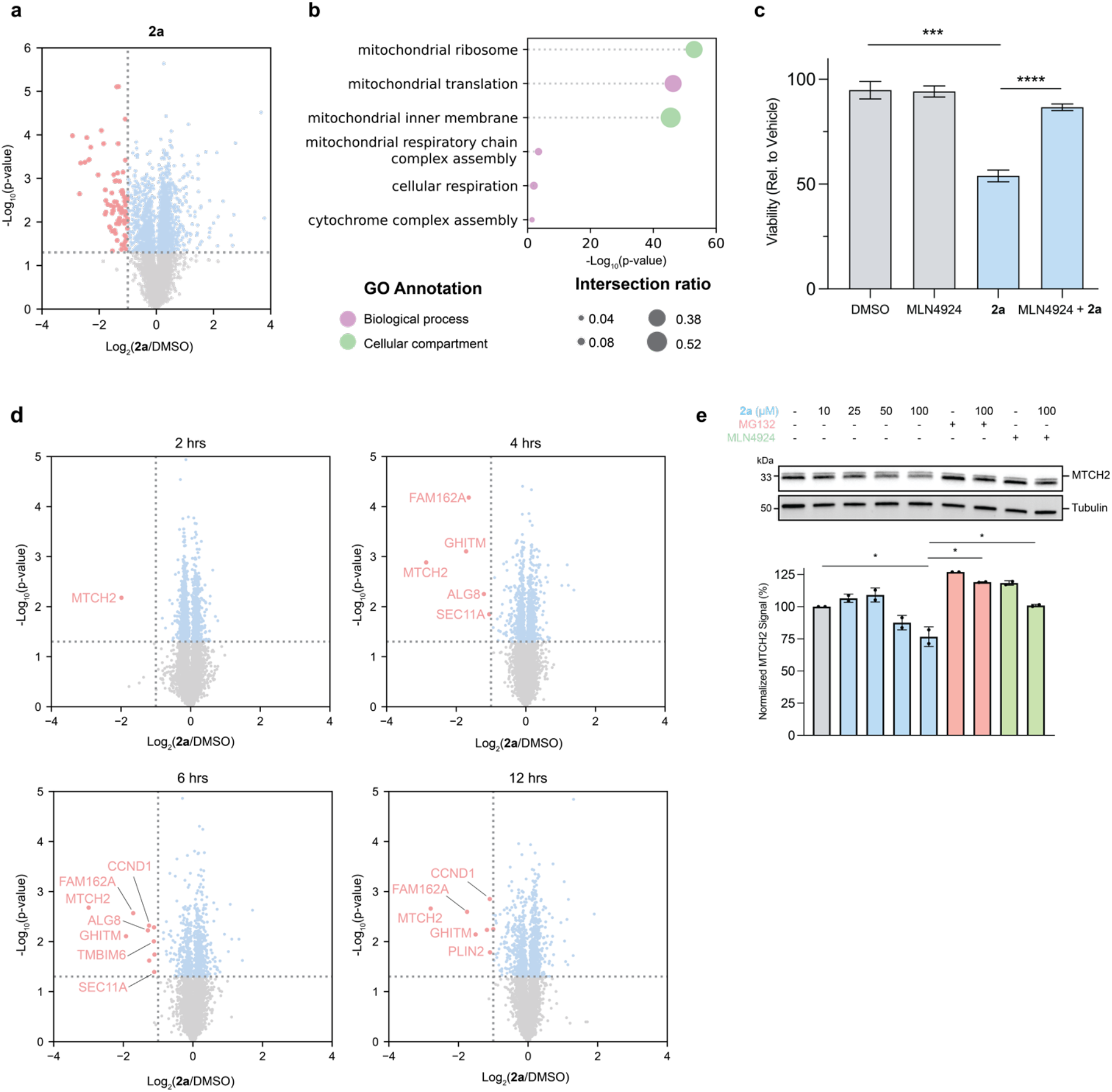
Characterization of 2a proteomic effects. (**a**) Volcano plot of protein abundances following 18 h treatment of MDA-MB-231 cells with **2a**. (**b**) Top Gene Ontology (GO) driver terms categorized by biological process and cellular compartments. The adjusted p-value is calculated with Fisher’s exact test. Intersection ratio represents proportion of targets within a given category relative to the total number of targets in the query. (**c**) Cell viability assay of MDA-MB-231 cells treated for 18 h with DMSO (vehicle), MLN4924 (1 µM), **2a** (50 µM) or a combination of MLN4924 and **2a**. Results depict the relative luminescence percentage, normalized to vehicle. Error bars represent mean ± SD from n = 3 independent experiments. Statistical significance was determined using a two-tailed Student’s t-test (* p <0.05, ***p <0.001; **** p <0.0001). (**d**) Volcano plots of protein abundances at 2, 4, 6 and 12 h following **2a** treatment. Downregulated targets are highlighted in red. (**e**) Immunoblot and statistical analyses confirming proteasome- and CRBN-mediated degradation of MTCH2 following 2 h pre-incubation with MG132 (10 µM) or MLN4924 (1 µM), and subsequent 6 h co-treatment with **2a**.

### Impact of AgnoTAC Modifications on Target Degradation

It is well-established that changes to linker length and composition can impact the potency and selectivity of bifunctional molecules^54-56^. We therefore wondered how variations in linker design affect the proteome-wide degradation profiles across all scaffolds. Analysis of our chemical proteomic data revealed that each linker series within a scaffold generally downregulated the same number of targets (∼25), with the exception of PEG2, which had 12 targets (**Fig. 5a, Fig. S8**). Moreover, unique targets across all scaffolds were overwhelmingly downregulated by a single linker type (**Fig. 5b**), indicating that linker structure plays a critical role in AgnoTAC selectivity. For example, though AgnoTACs featuring scaffold F05 led to the degradation of several proteins (**Fig. 5c**), we observe that degradation of Cyclin A2 (CCNA2)^57^ and ETS-1 is mediated preferentially by AgnoTAC **5a**, which possess a C2 linker (**Fig. 5d-e**). Additionally, in some cases, linker composition appears to be more a critical variable over length. For instance, BRD2 degradation is primarily observed with PEG linker-containing AgnoTACs based on scaffold F10, whereas aliphatic linkers had minimal effect on BRD2 abundance (**Fig. 5f, g)**. This observation was notably carried over to other BET family proteins (e.g., BRD3 and BRD4, **Fig. S9a-e**). Additional examples of linker-dependent degradation profiles include UFD1 and NPLOC4 (nuclear protein localization 4, NPL4) both degraded equipotently by **1c** (**Fig. S9f-h**) and TMEM205 degraded by **8d**\* (**Fig. S9i, j**).

**Fig. 5.**
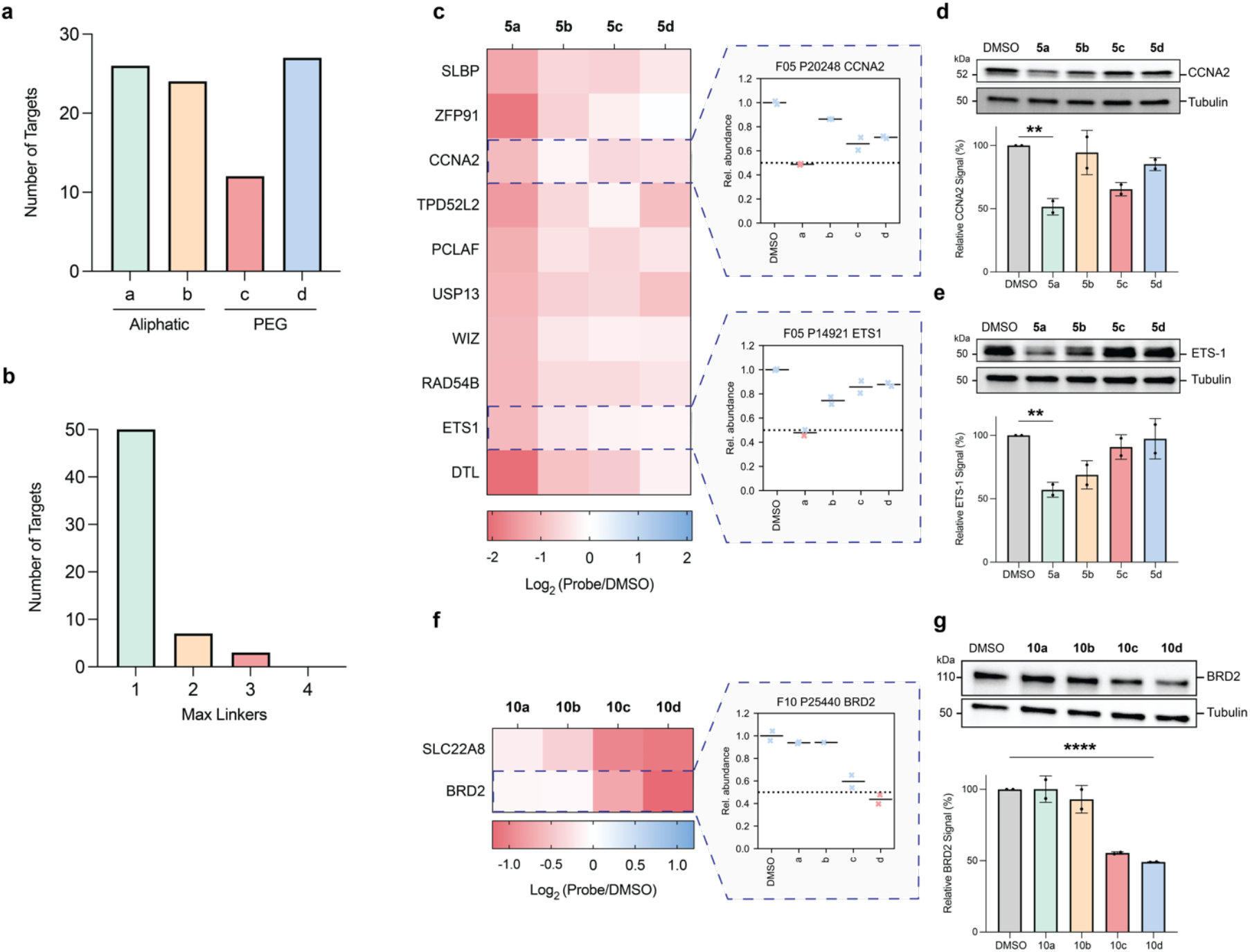
Linker composition influences degradation profiles. (**a**) Total number of targets downregulated classified by linker type (aliphatic or PEG). (**b**) Number of unique downregulated proteins targeted by 1, 2, 3 or all 4 linker types across scaffolds. (**c-g**) Protein abundance heatmaps and quantitative analysis of the mean abundance of select targets across different linker types, including (**c-e**) CCNA2 and ETS-1 in response to 100 µM treatments, and (**f, g**) BRD2 in response to 50 µM AgnoTAC treatments. Dashed line represents downregulation threshold. Bar plots represent me*an* ± SD (n = 2 biological replicates). Statistical significance was determined using a two-tailed Student’s t-test (** p <0.01; **** p <0.0001).

Previous studies have shown that thalidomide-based PROTACs can induce degradation of immunomodulatory (IMiD) targets through molecular glue (MGD) mechanisms^58-60^. Structural studies of CRBN ternary complexes with IMiDs have revealed a recurring motif among various neosubstrates, including CK1α, GSPT1 and IKFZ proteins^61, 62^. This motif is characterized by a β-hairpin loop containing a key glycine (G) residue, facilitating interactions with the CRBN/MGD binding interface^30, 63^. Given the prevalence of β-hairpin G-loops in the human proteome^64^, we wondered whether additional AgnoTAC targets possess this motif. Using a known G-loop structure as a query, we identified nine proteins amongst AgnoTAC targets containing at least one β-hairpin loop (**Fig. S10a**), raising the possibility of being potential IMiD off-targets. Among 72 AgnoTAC profiles, we identified 17 that degrade established IMiD glue-based off-targets, including GSPT1, GSPT2 and ZFP91 (**Fig. S5a, Fig. S10b**). Notably, these degradation events occurred with both aryl ether and oxyacetamide linkages used in our studies, though IMiD off-targets were often not shared across all headgroups with the same linkages (**Fig. S10c, d**). We noted even subtle structural modifications, such as altering the connectivity of the bridging C-C bond in the benzhydrylpiperazine group between **7b\*** and **8b\*** (C5 linker) resulted in distinct effects on GSPT1 levels, with **8b\*** causing more pronounced downregulation (**Fig. S10e-h**).

Collectively, these findings underscore the significant influence of linker design on target degradation patterns and the importance of considering multiple linker structures, as well as highlights the challenges in extrapolating generalizable linker design principles when examining a broad range of targets.

### Mechanistic investigation of degraded targets

Though we have demonstrated, in several examples, that AgnoTAC-dependent protein degradation is mediated through Cullin and proteasome pathways, we sought to further characterize these degradation events. To this end, we conducted a series of experiments to further delineate the mechanism of **10d**-induced BRD2 proteasomal degradation (**Fig. 2j-l**). We first assessed degradation kinetics, where we observed BRD2 degradation occurs as early as two hours following **10d** treatment (**Fig. S11a**). We next assessed E3 ligase-binding dependence by synthesizing a structurally similar negative control **10d-neg** in which the glutarimide moiety of the thalidomide derivative is capped with a methyl group (**Fig. 6a**), preventing CRBN recruitment^65^. As expected, **10d-neg** failed to induce BRD2 degradation, confirming that E3 ligase engagement is needed for **10d**’s activity (**Fig. 6b**). We further evaluated BRD2 depletion using a HiBiT assay in HEK293T BRD2-HiBiT cell line (**Fig. S11b**), which confirmed that **10d** most potently degraded BRD2, as observed in our proteomic studies. Finally, co-treatment of **10d** and JQ1, a bromodomain-targeting inhibitor^66^, rescued BRD2 levels, suggesting that **10d** binds BRD2 competitively with JQ1 and co-incubation disrupts ternary complex formation to prevent degradation (**Fig. 6c, d**). We generated analogous negative control AgnoTACs for several additional targets displaying Cullin-dependent degradation, including CHCHD2 (**Fig. S11c, d**), SIRT1 (**Fig. S11e, f**) and PLOD2 (**Fig. 6e, f**), confirming degradation dependence on CRBN. Perhaps not surprisingly, for targets undergoing Cullin-independent degradation, both the parent as well as the methyl-capped thalidomide control displayed similar profiles, confirming, in those instances, that degradation occurs via alternative mechanisms (**Fig. S11g-k**).

**Fig. 6.**
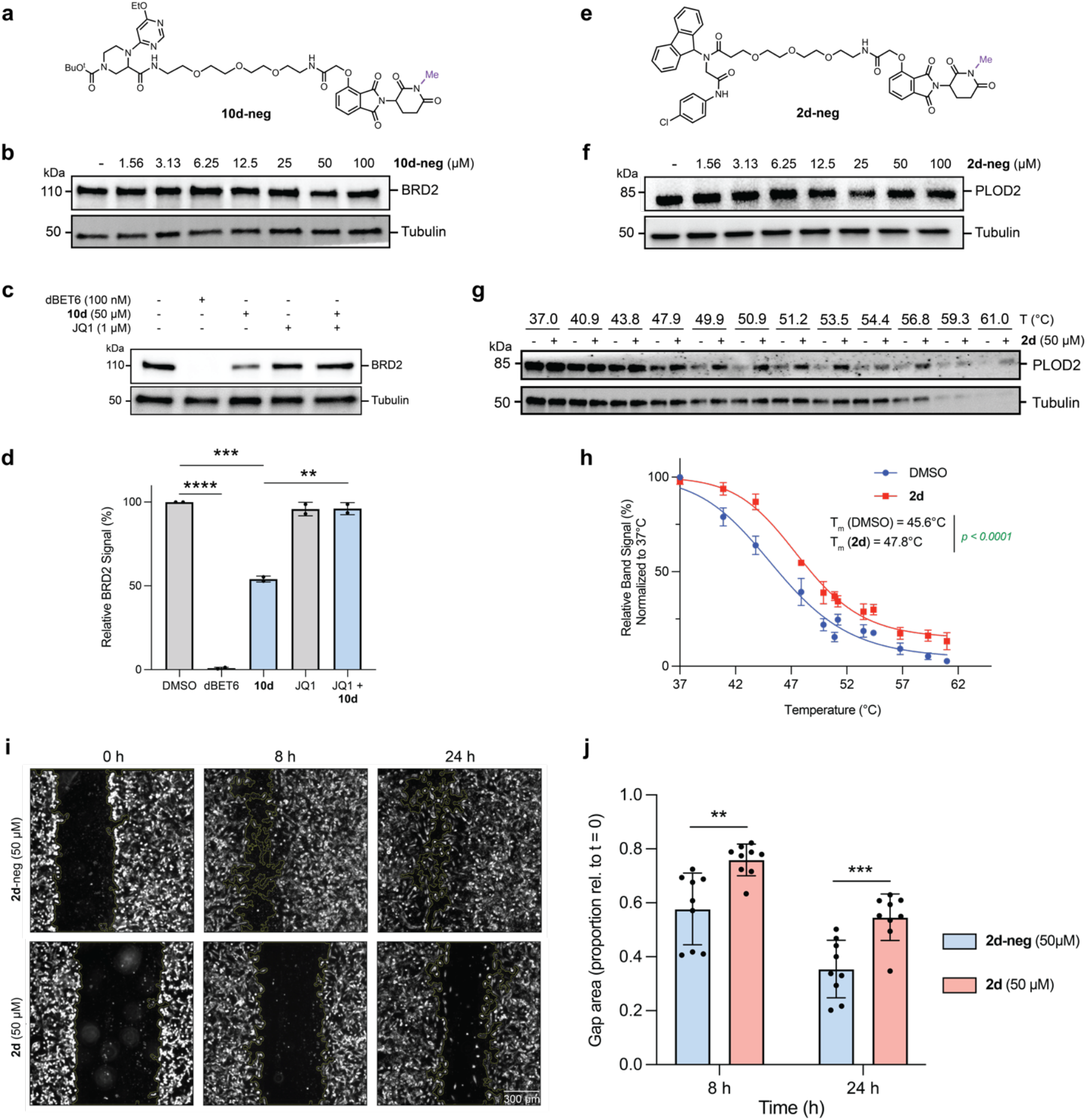
Mechanistic characterization of AgnoTAC-induced degradation events. (**a**) Chemical structure of negative control **10d-neg** and (**b**) immunoblot analysis of dose-response at indicated concentrations following 18 h treatment of MDA-MB-231 cells. (**c, d**) Immunoblot and quantitative analysis of BRD2 degradation after 18 h treatment with dBET6 (100 nM), JQ1 (1 µM) and **10d** (50 µM). JQ1-treated cells were pre-treated with inhibitor for 1 h. Bar plot data represents mean ± SD (n = 2 biological replicates), statistical significance was determined using a two-tailed Student’s tailed t-test (** *p* <0.01; *** *p* <0.001; **** *p* <0.0001). (**e**) Chemical structure of negative control **2d-neg** and (**f**) immunoblot analysis of PLOD2 with **2d** dose-response at indicated concentrations following 18 h treatment. (**g, h**) CETSA for PLOD2 upon MDA-MB-231 treatment with **2d** for 30 min. Protein levels were assessed by immunoblot and quantified to generate thermal stabilization curves. Data points represent mean ± SD, n = 2 independent experiments. Statistical significance is indicated with *p*-values. (**i**) Representative images of **2d** and **2d-neg** effects on MDA-MB-231 migration measured by a wound healing assay and (**j**) corresponding statistical significance analysis for n = 9 biological replicates.

Among the validated targets, we observed one of the highest levels of degradation for PLOD2, mediated by **2d** (DC_50_ = 23.7 µM and D_max_ = 74%, **Fig. 2d-f**). Depletion of PLOD2 was observable as early as 6 h and continued until 24 h after treatment (**Fig. S11l**). To further verify that PLOD2 degradation was mediated by binding, we performed cellular thermal shift assay (CETSA) experiments, where we observed marked stabilization of PLOD2 upon treatment with **2d** (**Fig. 6g, h**) over an applied temperature gradient. PLOD2 catalyzes lysyl hydroxylation on the Gly-X-Y motif of collagen peptides and is required for the formation of stabilized collagen crosslinks^67^. Given these functions, it is perhaps not surprising that PLOD2 is critical for epithelial mesenchymal transition (EMT), migration, and metastasis of various cancers^68, 69^, as well as tissue repair and fibrosis^70^, though no selective inhibitors have been reported. To assess whether targeted degradation of PLOD2 would disrupt cellular migration, we treated MDA-MB-231 cells with **2d** and observed a substantial decrease in migration rates, whereas **2d-neg** had minimal effects (**Fig. 6i, j**). Taken together, these data suggest that AgnoTACs directly engage and degrade targets through a CRBN-mediated mechanism and can serve as chemical probes to investigate protein functions.

As noted above, AgnoTAC **1c** induces linker-dependent, selective and equipotent downregulation of UFD1 and NPL4 (**Fig. 2a, Fig. S9f-h**). UFD1 (ubiquitin fusion degradation 1) and NPL4 (nuclear protein localization 4) are adaptor proteins that complex with AAA ATPase p97 (also known as VCP), mediating various p97 functions in protein ubiquitination and degradation^43, 71, 72^, endoplasmic reticulum-associated degradation (ERAD)^73-75^, and transport processes^76, 77^. Previous studies demonstrated that genetic ablation of either UFD1 or NPL4 destabilizes the VCP-UFD1-NPL4 complex, leading to proteasomal degradation of the other component, releasing free VCP^78^. Interestingly, we observe no changes in VCP abundance (**Supplemental dataset 1**), prompting us to speculate that AgnoTAC **1c** either directly targets UFD1 and NPL4 for degradation, or contributes to the destabilization of this complex resulting in subsequent degradation. As noted previously, we found UFD1 to be degraded by **1c** via a CRBN-mediated pathway (**Fig 2c**). Though degradation of NPL4 by **1c** appeared to progress equipotently (**Fig. 7a, b**) and with the same kinetics as UFD1 (**Fig, S11m**), its levels were restored upon proteasome inhibition, but not by neddylation inhibition (**Fig. 7c, d**). Notably, UFD1 and NPL4 degradation was recapitulated across multiple human cell lines derived from diverse tissues (**Fig. S12**), suggesting that these effects are not restricted to a specific model system. In addition, the negative control analog **1c-neg (Fig. 7e)**, had no effect on either UFD1 or NPL4 levels (**Fig. 7f**), implying that the degradation of both proteins is dependent on CRBN and is likely not the result of **1c** binding alone. To further assess whether UFD1 or NPL4 is the direct binding target of **1c**, we conducted CETSA at shorter incubation periods where protein loss was not observed (**Fig. 7g**). Treatment of MDA-MB-231 cells with **1c** resulted in increased thermal stability for UFD1 (**Fig. 7h**), whereas no stabilization was observed for NPL4 (**Fig. 7i**), suggesting direct engagement of UFD1 by **1c**. Collectively these results indicate that **1c** likely binds UFD1, resulting in its CRBN-mediated proteasomal degradation, and that this binding and recruitment of CRBN leads to subsequent proteasomal degradation of NPL4 via endogenous pathways.

**Fig. 7.**
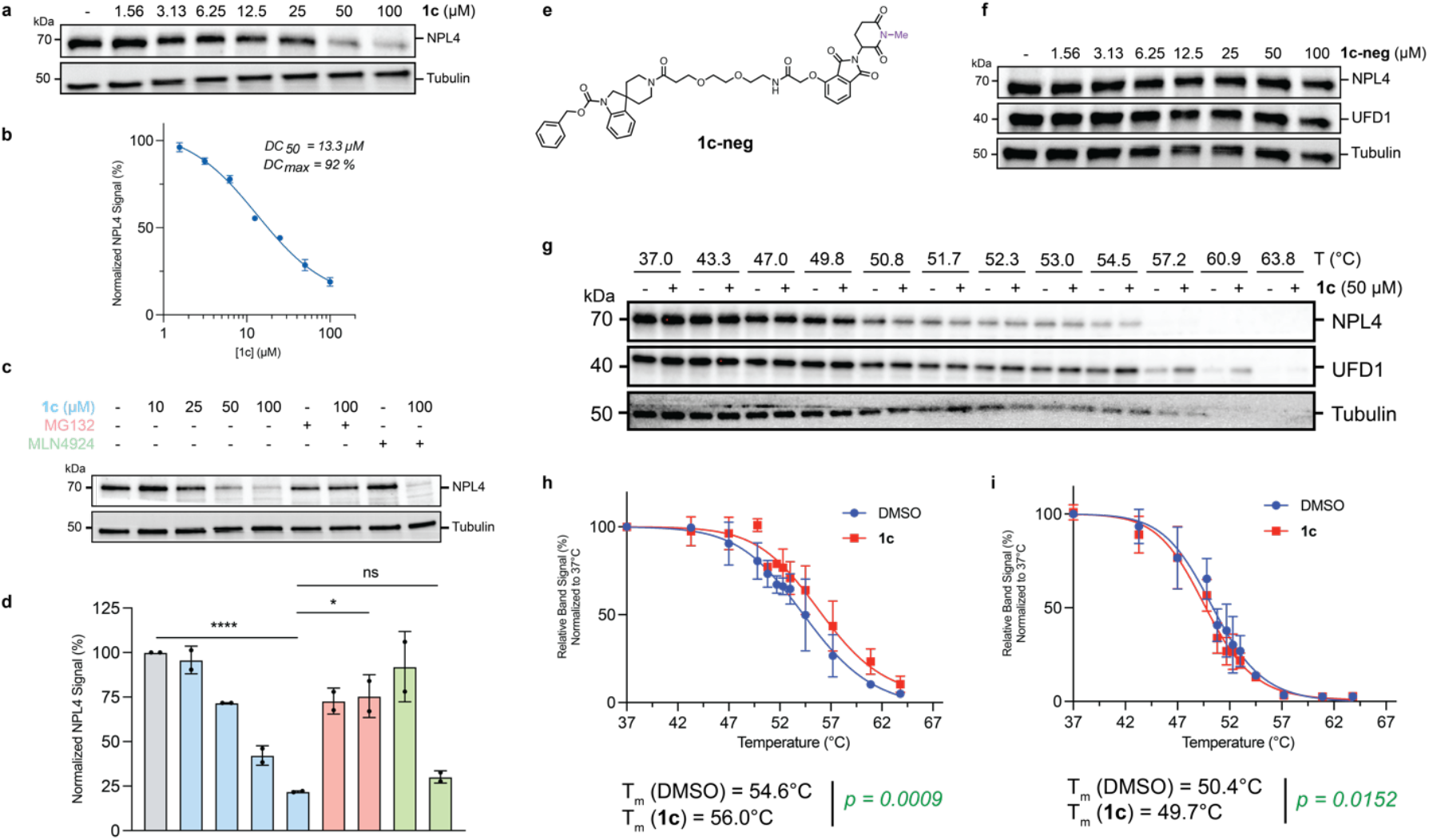
UFD1/NPL4 adaptor protein complex degradation mechanism. (**a-d**) Immunoblot analyses of NPL4 in MDA-MB-231 cells treated with various concentrations of **1c**, MG132 (10 µM) or MLN4924 (1 µM). Bar graph displays normalized NPL4 signal intensities, highlighting significant reductions and protein level rescues upon **1c** treatment (mean ± SD, n = 2 independent experiments, * *p* <0.05; ** *p* <0.01; *** *p* <0.001; **** *p* <0.0001; ns = not significant). (**e**) Chemical structure of **1c-neg** control, followed by (**f**) immunoblot analysis of NPL4 and UFD1 levels after **1c-neg** treatment at various concentrations. (**g**) CETSA for NPL4 and UFD1 upon **1c** treatment for 30 min. Protein levels were assessed by immunoblot. (**h, i**) Thermal stabilization curves for (**h**) UFD1 and (**i**) NPL4 in the presence of **1c** compared to DMSO control. Data points represent mean ± SD, n = 2 independent experiments. Statistical significance is indicated with *p*-values.

## CONCLUSION

Chemical proteomics is firmly established as a powerful strategy for global ligand and target discovery in native systems. Founded on the use of active site-directed probes as reporters for enzyme function and pocket occupancy (i.e., ABPP),^79-83^ these methods are now routinely deployed to map covalent and non-covalent small molecule interactions proteome-wide, independent of protein function or class^84^. Despite numerous instances of chemical proteomic methods being utilized to develop functional small molecules against diverse targets^85-94^, conventionally such methods report binding, rather than functional outcomes. More generally, understanding how small molecule binding might relate to downstream consequences, and how binders can be converted into functional compounds, is often target-specific. To begin addressing this gap, recent efforts have aimed to read out proteome-wide functional changes, such as perturbations to protein abundance^34^ and complex formation^95^, in response to small molecule treatment. However, in these examples, functionally unbiased chemical libraries are utilized and necessitate substantial follow-up investigation to pinpoint the mechanism of observed functional outcomes. Considering these challenges, we sought a chemical proteomic strategy for the systematic and global discovery of small molecules that impose specific functional consequences. This effort was driven primarily by the lack of such methods to discover small molecule protein degraders, beyond targets with established ligands. We hypothesized that integrating function-specific library design principles with quantitative proteomics would enable prospective discovery of small molecules that are tuned to degrade proteins through a pre-defined mechanism.

To establish a proof-of-concept, we employed straightforward coupling chemistry to generate a library of PROTACs aimed at agnostically identifying degradable proteins via CRBN-mediated proteasomal degradation. Given the library’s small size relative to conventional high-throughput screening (HTS), we prioritized structurally diverse, drug-like TBMs (**Figs. S1, S2**) to maximize exploration of chemical space, which we suspect contributed to the successful identification of novel degraders. While the importance of linkers in achieving productive ternary complex formation for specific targets is well-established, our findings emphasize their crucial roles in *de novo* target discovery using agnostic screens. From these efforts, we identified 50+ unique downregulation events, many of which were validated to proceed via the intended degradation mechanism. Notably, we identified, to our knowledge, first-in-class chemical probes for several proteins, including UFD1/NPL4, CHCHD2, and PLOD2, resulting in their targeted degradation. We generally observed DC_50_ values ranged from 10-30 µM with maximum degradation typically between 40-70%, indicating that TBMs likely possess lower affinities for their targets, unoptimized linkers, and/or targets are poor substrates for CRBN. Consequently, we did not observe characteristic autoinhibition at higher AgnoTAC concentrations^96^. Despite these modest potencies, we observed surprising proteome-wide selectivity, indicating that AgnoTACs could serve as suitable probes to investigate protein function, (e.g., PLOD2, **Fig. 6i, j**), as well as valuable starting points for further optimization.

A key takeaway from our study is the importance of rigorous validation to confirm AgnoTAC-mediated degradation proceeds through the intended mechanism. Employing negative control probes and direct target engagement assays is essential to differentiate genuine degradation events from indirect effects. For example, in the case of UFD1/NPL4, our data suggests that UFD1 is the direct target of **1c**, while NPL4 degradation likely occurs through a bystander mechanism resulting from destabilization of the UFD1/NPL4 complex and subsequent proteasomal clearance, a pathway that, to our knowledge, has yet to be annotated. Additionally, we identified proteasomal degradation events that appeared to be CRBN-independent (e.g., ETS-1, TMEM205). In such cases, we speculate that direct binding of the AgnoTAC leads to protein destabilization, or perhaps downstream consequences of degradation events not captured in our proteomic studies. In some instances, we expect AgnoTACs to target essential proteins or disrupt key cellular processes (e.g., transcription, cell cycle), leading to broader proteomic changes and presenting additional challenges in elucidating the primary target and mechanism. This was evidenced with **2a**, which reduced cell viability and mediated downregulation of 85 proteins. We demonstrated that **2a** viability effects were, in part, CRBN-dependent and converged on MTCH2 as a primary degradation target. In this vein, we see potential in applying AgnoTAC libraries in phenotypic screens where, akin to other chemical proteomic libraries^86, 88, 97, 98^, hit compounds could directly serve as tools for target identification.^99^

Building on the evolving concepts of chemical proteomic library design^100-104^, future AgnoTAC libraries incorporating sp^3^-rich, stereochemically defined and densely functionalized cores should enable deeper exploration of the proteome while expediting the discovery of authentic, molecular recognition-driven degradation events. To aide deconvolution efforts, integration of profiling methods that provide mechanistic information, for example, MS-based ubiquitinomics^31^ or pulse labeling experiments^105^, should streamline the identification of AgnoTAC direct targets. Projecting forward, we believe that global profiling of target-agnostic, functionally defined libraries will fill gaps in small molecule-probe development by enabling the concatenated discovery of functional compounds and corresponding targets. In this context, this strategy extends TPD beyond targets with well-established ligands and we expect will furnish valuable starting points for the development of advanced small molecule degraders for biologically and therapeutically compelling proteins.

## Supporting information

Supplemental Chemistry Information

## SUPPORTING INFORMATION

Supplementary Figures S1-S13, detailed procedures for all experiments, specifics for reagents and instruments used can be found in Supplemental Information document.

Supplemental Document 1 – Synthetic methods and compound characterization Supplemental Dataset 1 – Proteomics data

Supplemental Dataset 2 – Figure data

## COMPETING INTERESTS

I.F., L.P.C. and C.G.P. are inventors on a patent application (WO2022187650) submitted by The Scripps Research Institute that covers heterobifunctional molecules for targeted protein degradation. M.E., A.S., J.M.R, S.E.W., A.V and S.M.M are employees of AbbVie. All other authors declare no competing interests.

## ACKNOWLEDGEMENTS

This work was funded, in part, by AbbVie Inc. and Inception Therapeutics. I.F. was supported by a Ford Foundation Predoctoral Fellowship.

## AUTHOR CONTRIBUTIONS

I.F., L.P.C. and C.G.P. conceived this study, designed experiments and interpreted results. I.F. prepared figures. I.F. and C.G.P. wrote the manuscript. I.F., A.M.J., C.G., C.M.C., M.E. and A.S. synthesized chemical probes and characterized new compounds. I.F. and L.P.C. performed chemoproteomic experiments, analyzed MS data, performed validation experiments and functional characterization of compounds. T-Y. C. and C. G. assisted with validation experiments. M.E., A.S., J.M.R, S.E.W., A.V and S.M.M. provided reagents and assistance with experimental design. All authors assisted with editing of the manuscript.

