## Supplemental Information for "Proteome-Wide Discovery of Degradable Proteins Using Bifunctional Molecules"

#### Table of content

**Supplementary Figures**
Figure S1 – Chemical structures of AgnoTAC degrader library
Figure S2 – Chemoinformatic analysis of the AgnoTAC library
Figure S3 – Evaluation of AgnoTAC effects on cell viability
Figure S4 – Proteomic profiles of AgnoTAC library
Figure S5 – Analysis of detected and downregulated targets
Figure S6 – Additional target degraded by AgnoTAC **5a**
Figure S7 – Protein cluster analysis of 2a-treated samples
Figure S8 – Linker composition influences target degradation, related to Fig. 5
Figure S9 – Additional linker effects, related to Fig. 5.
Figure S10 – Analysis of molecular glue off-targets
Figure S11 – Additional mechanistic characterization of AgnoTAC targets
Figure S12 – AgnoTAC **1c** degrades UFD1/NPL4 in several cell lines
Figure S13 – Compiled uncropped Western blots

**Supplementary Tables**
Table S1 – Reagents, kits, assays, antibodies and softwares used in this study.
Supplemental Dataset 1 – Proteomics data
Supplemental Dataset 2 – Figure data

**Biological Methods**

**References**

### SUPPLEMENTARY FIGURES

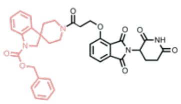

**1a**

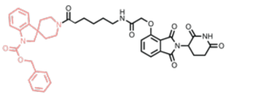

**1b**

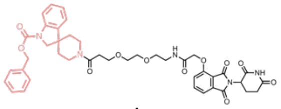

**1c**

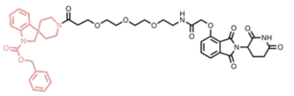

**1d**

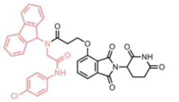

**2a**

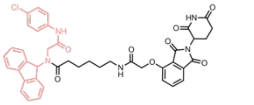

**2b**

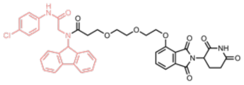

**2c**

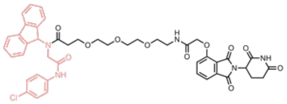

**2d**

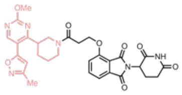

**3a**

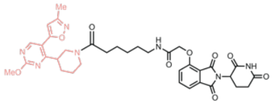

**3b**

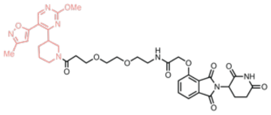

**3c**

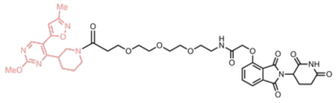

**3d**

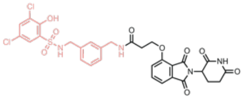

**4a**

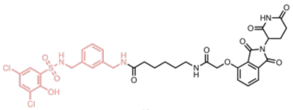

**4b**

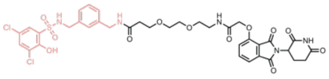

**4c**

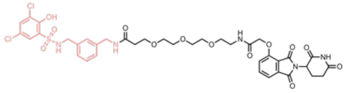

**4d**

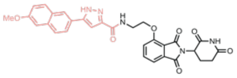

**5a**

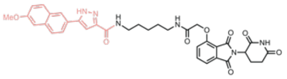

**5b**

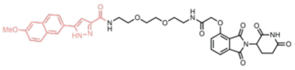

**5c**

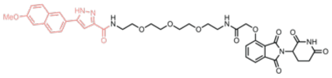

**5d**

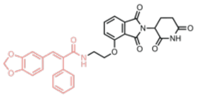

**6a**

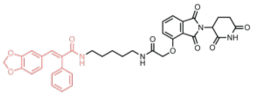

**6b**

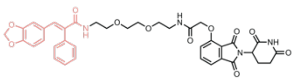

**6c**

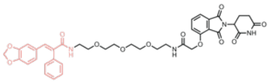

**6d**

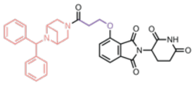

**7a***

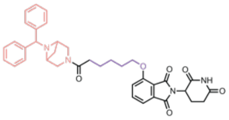

**7b***

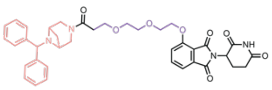

**7c***

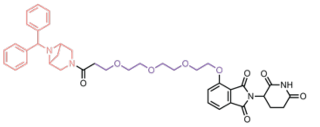

**7d***

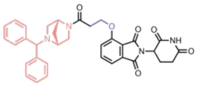

**8a***

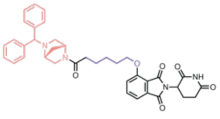

**8b***

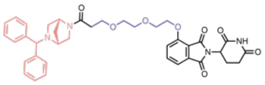

**8c***

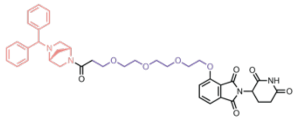

**8d***

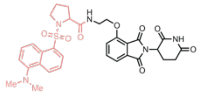

**9a**

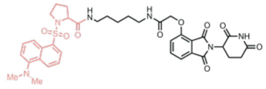

**9b**

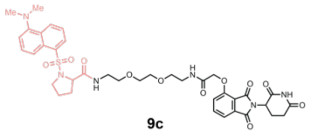

**9c**

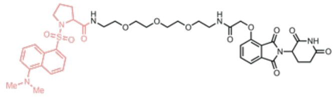

**9d**

**Fig. S1| Chemical structures of AgnoTAC degrader library** **(continued on next page)**

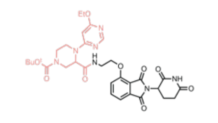

**10a**

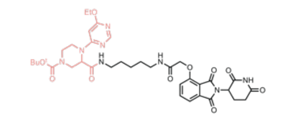

**10b**

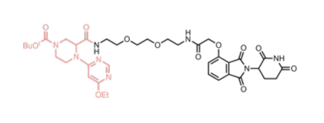

**10c**

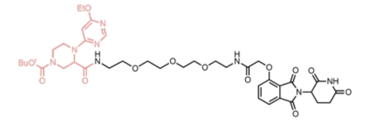

**10d**

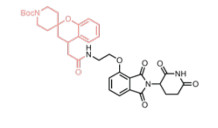

**11a**

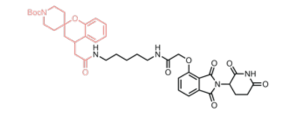

**11b**

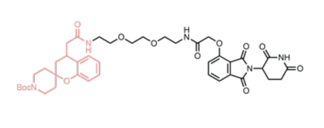

**11c**

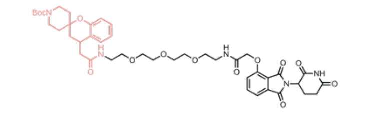

**11d**

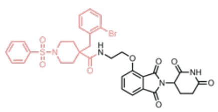

**12a**

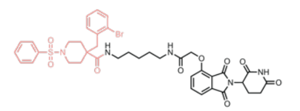

**12b**

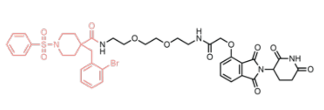

**12c**

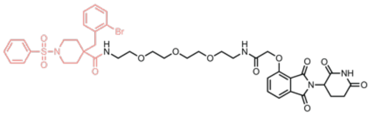

**12d**

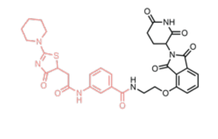

**13a**

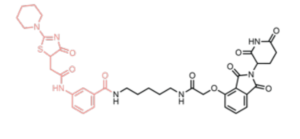

**13b**

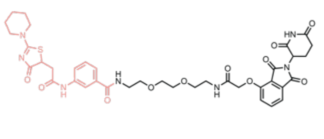

**13c**

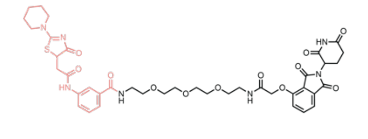

**13d**

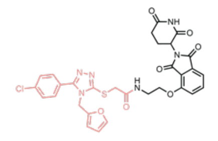

**14a**

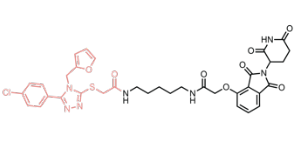

**14b**

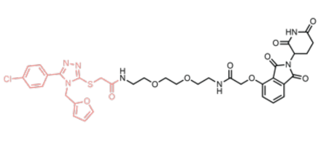

**14c**

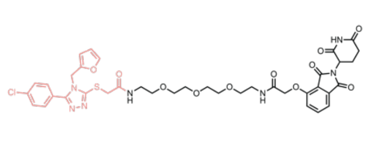

**14d**

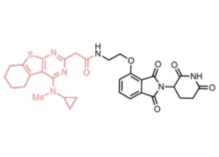

**15a**

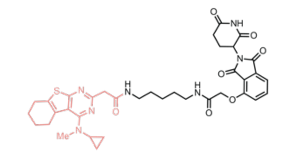

**15b**

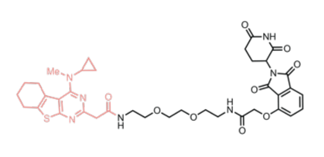

**15c**

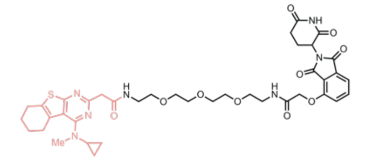

**15d**

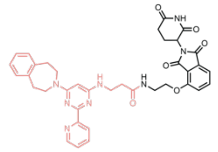

**16a**

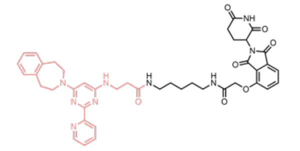

**16b**

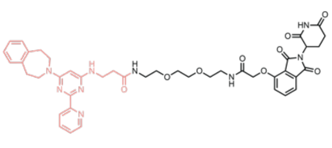

**16c**

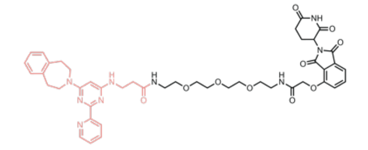

**16d**

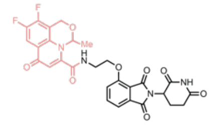

**17a**

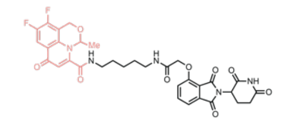

**17b**

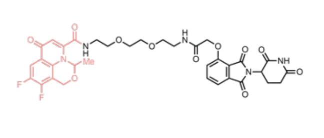

**17c**

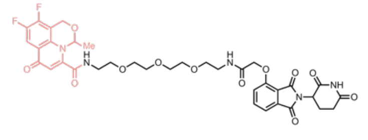

**17d**

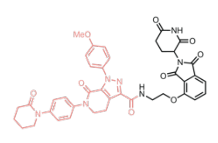

**18a**

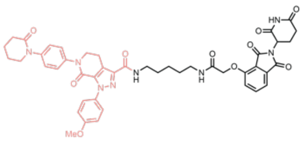

**18b**

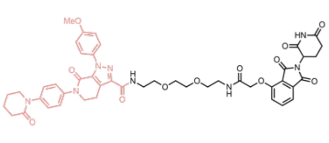

**18c**

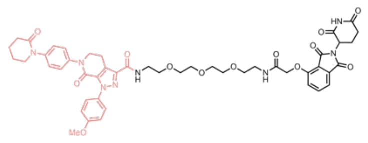

**18d**

**Fig. S1| Chemical structures of AgnoTAC degrader library.**

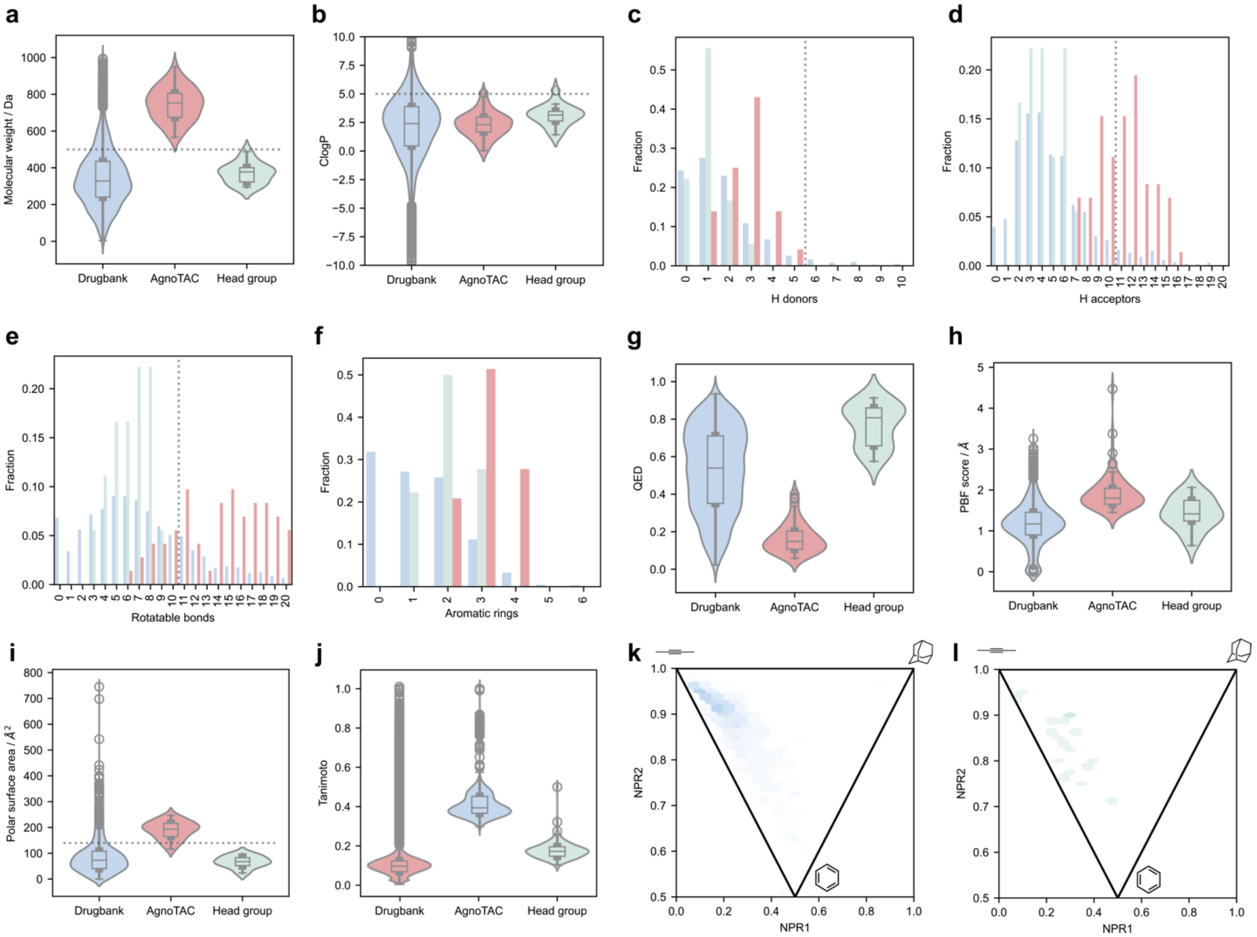


**Fig. S2| Chemoinformatic analysis of AgnoTAC library.** Overview of the molecular properties of AgnoTAC headgroups (green) and degraders (red) compared to DrugBank ligands (blue): (**a**) molecular weight; (**b**) calculated LogP (cLogP); (**c**) number of H bond donors and (**d**) H bond acceptors; (**e**) number of rotatable bonds; (**f**) number of aromatic rings; (**g**) quantitative estimate of drug likeness (QED); (**h**) characterization of compound $sp^3$ character using plane of best fit (PBF); (**i**) polar surface area distribution; (**j**) Tanimoto coefficient: similarity analysis measuring chemical diversity. (**k, l**) distributions of normalized principal moments of inertia ratios (NPR) depicting molecular shape of (**k**) DrugBank ligands compared to (**l**) AgnoTAC headgroups.

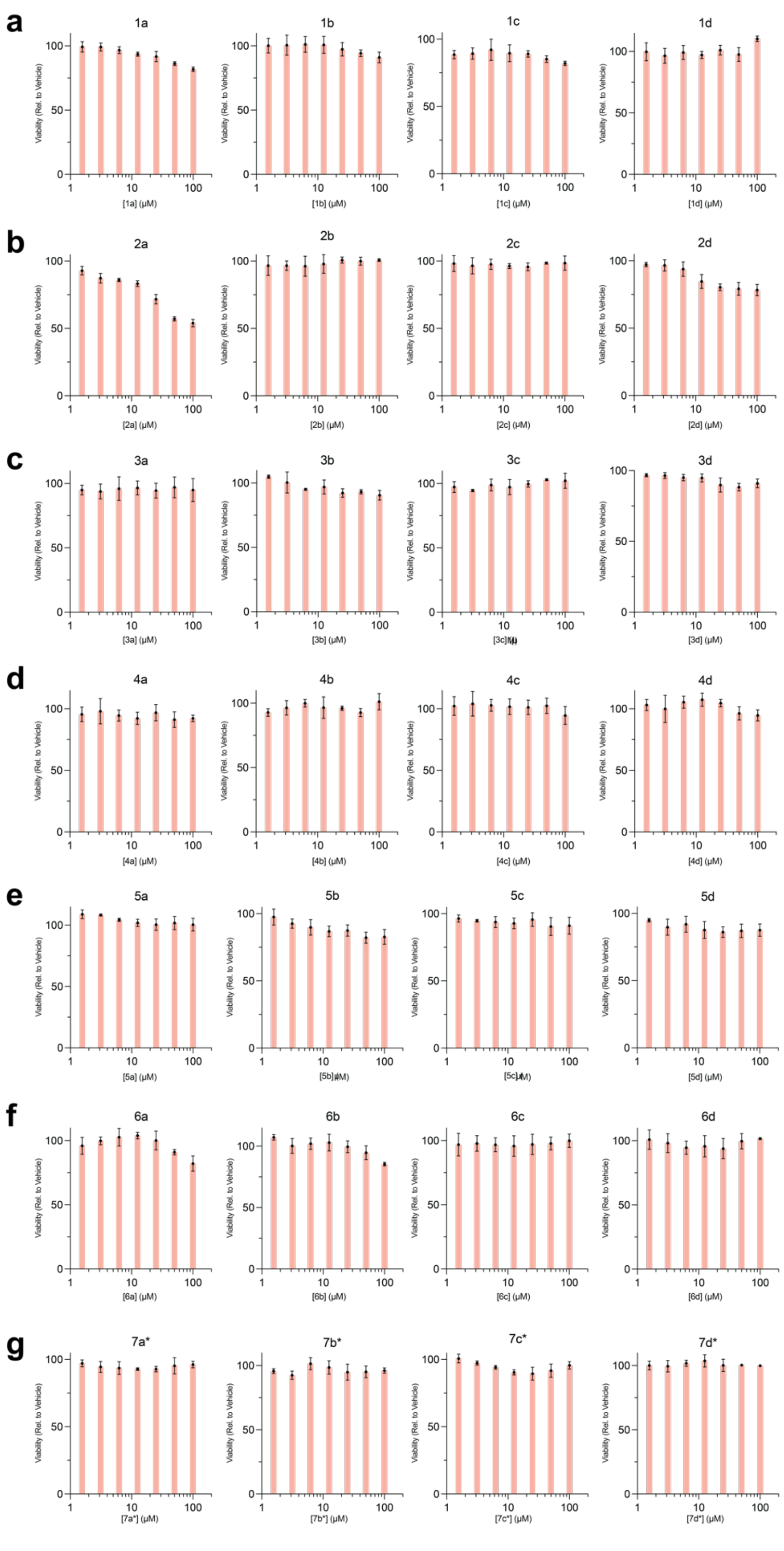


**Fig. S3| Evaluation of AgnoTAC effects on cell viability** **(continued on next page)**

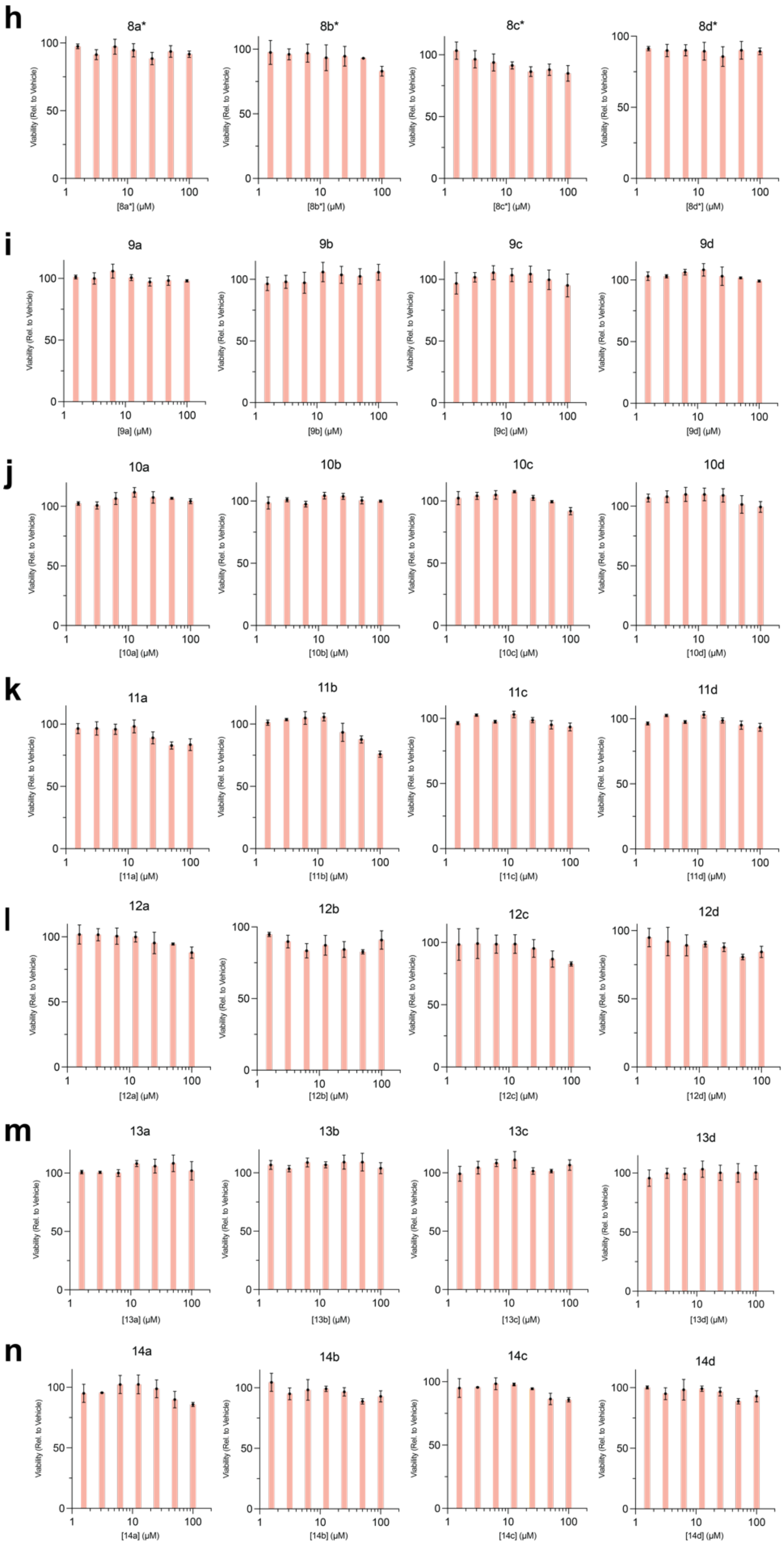


**Fig. S3| Evaluation of AgnoTAC effects on cell viability** (continued on next page)

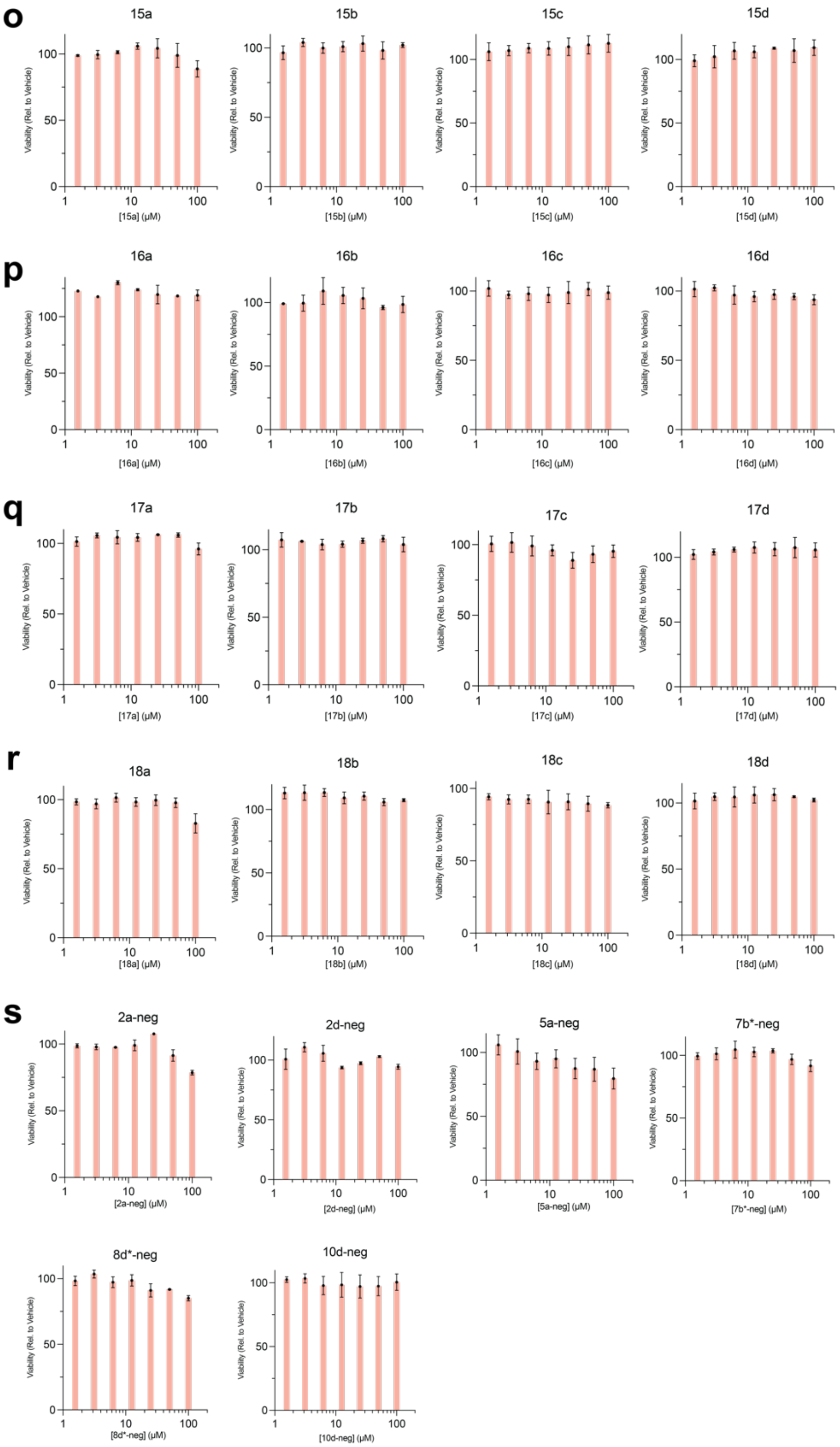


**Fig. S3| Evaluation of AgnoTAC effects on cell viability.** Cell viability was assessed using Cell Titer-Glo luminescence assay following MDA-MB-231 cell treatment with AgnoTACs at varying concentrations for 18 h. Results depict the relative luminescence percentage, normalized to DMSO vehicle. Error bars represent mean ± standard deviation (SD) from n = 3 independent experiments.

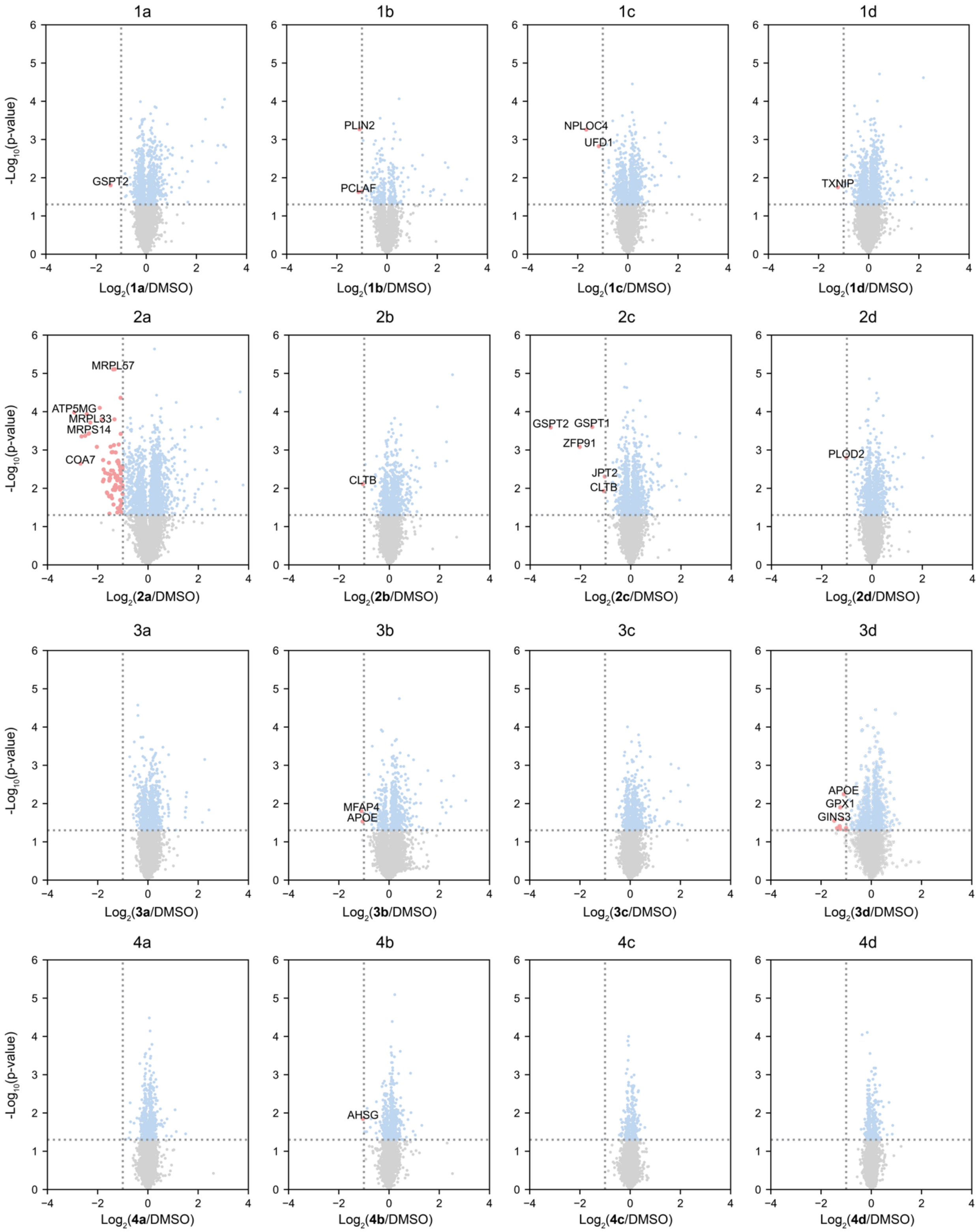


**Fig. S4| Proteomic profiles of AgnoTAC library (continued on next page)**

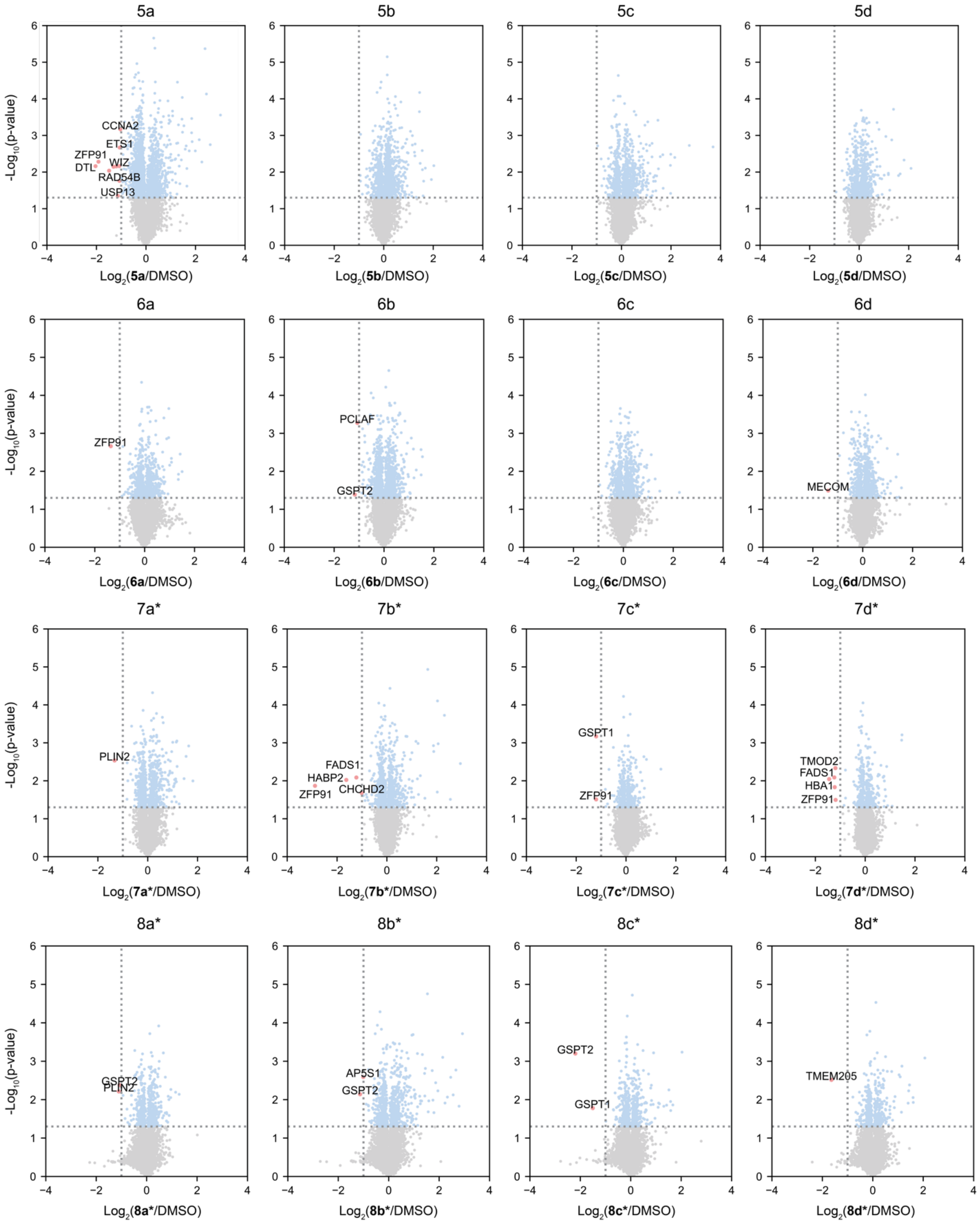


**Fig. S4| Proteomic profiles of AgnoTAC library** **(continued on next page)**

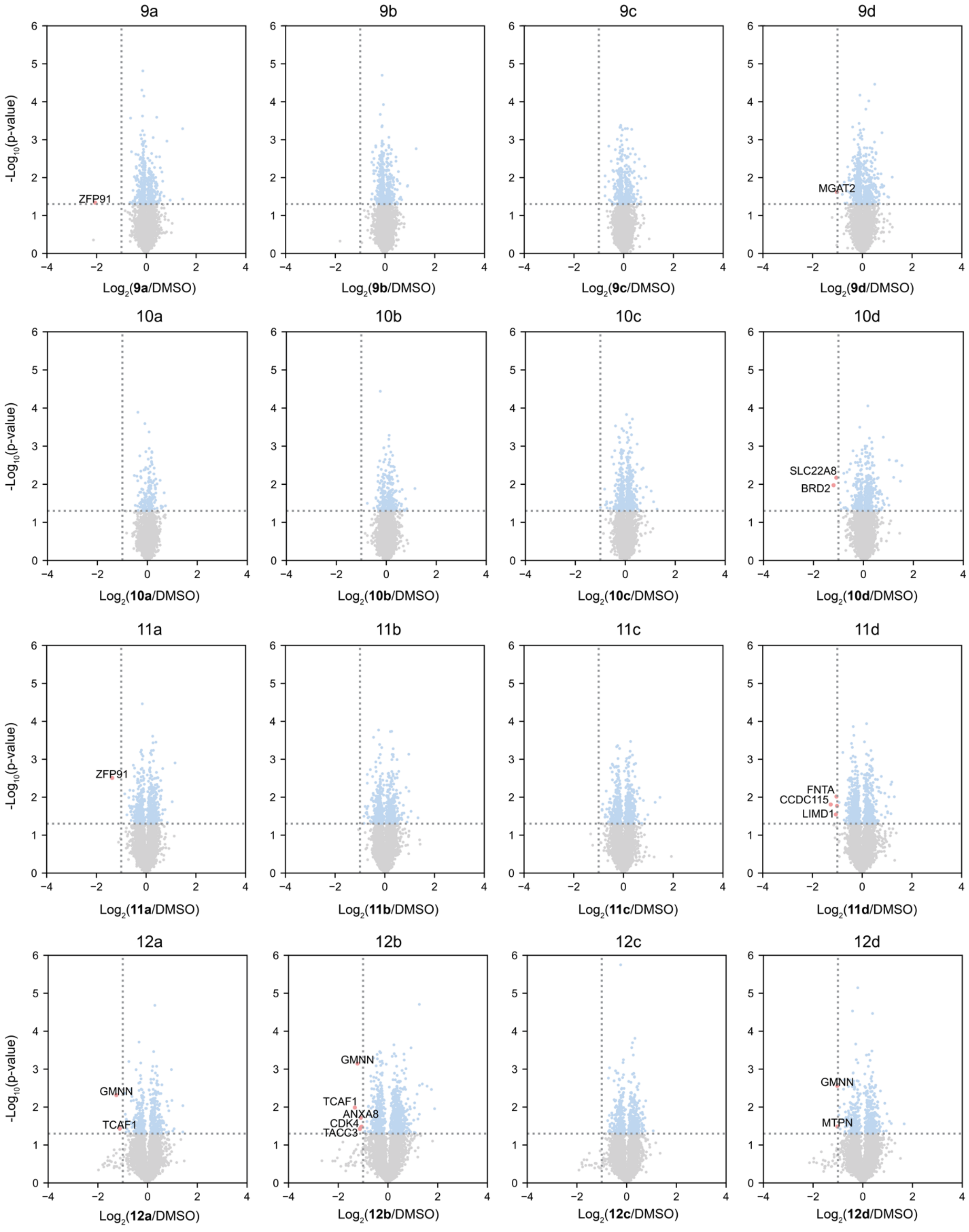


**Fig. S4| Proteomic profiles of AgnoTAC library** **(continued on next page)**

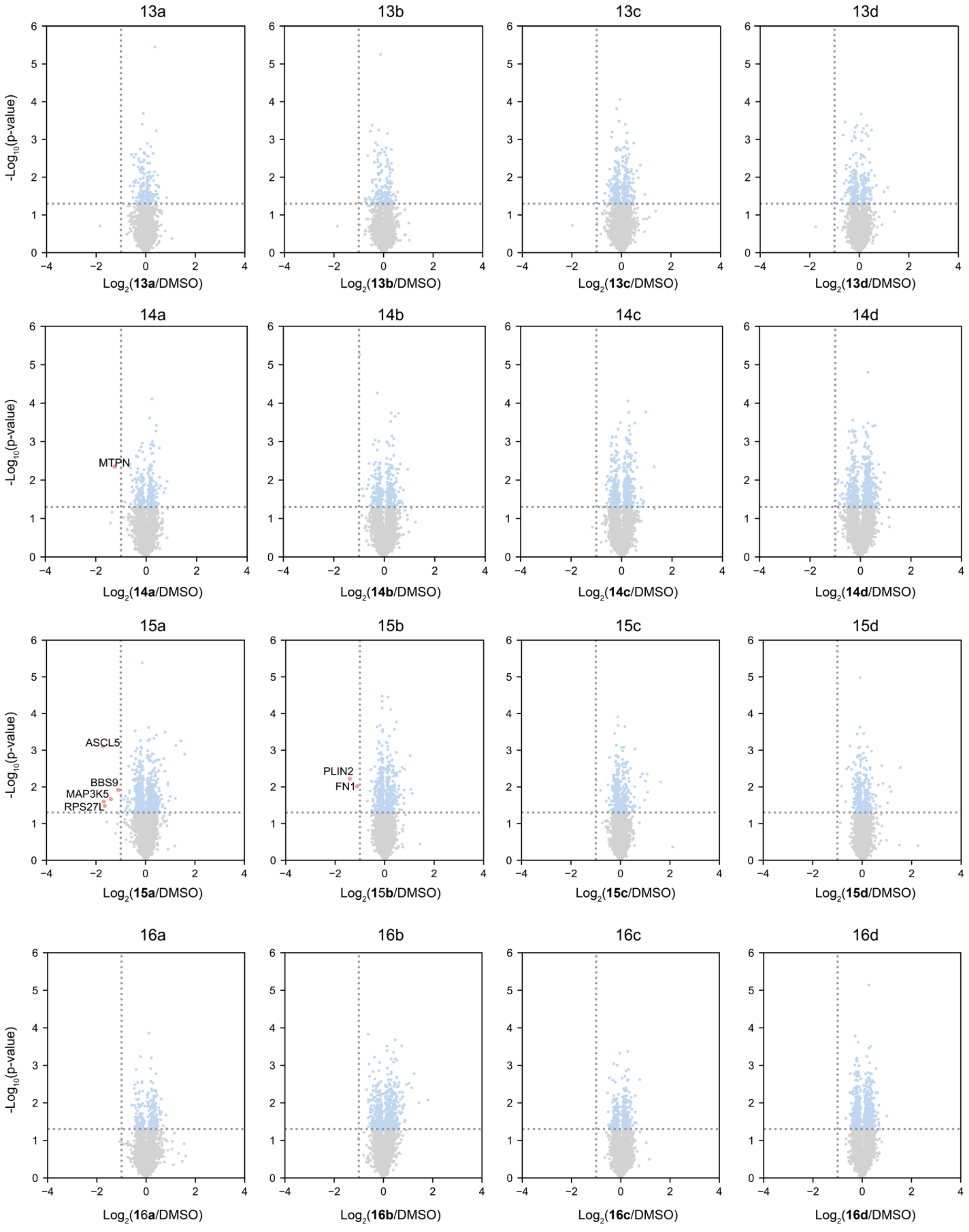


**Fig. S4| Proteomic profiles of AgnoTAC library** **(continued on next page)**

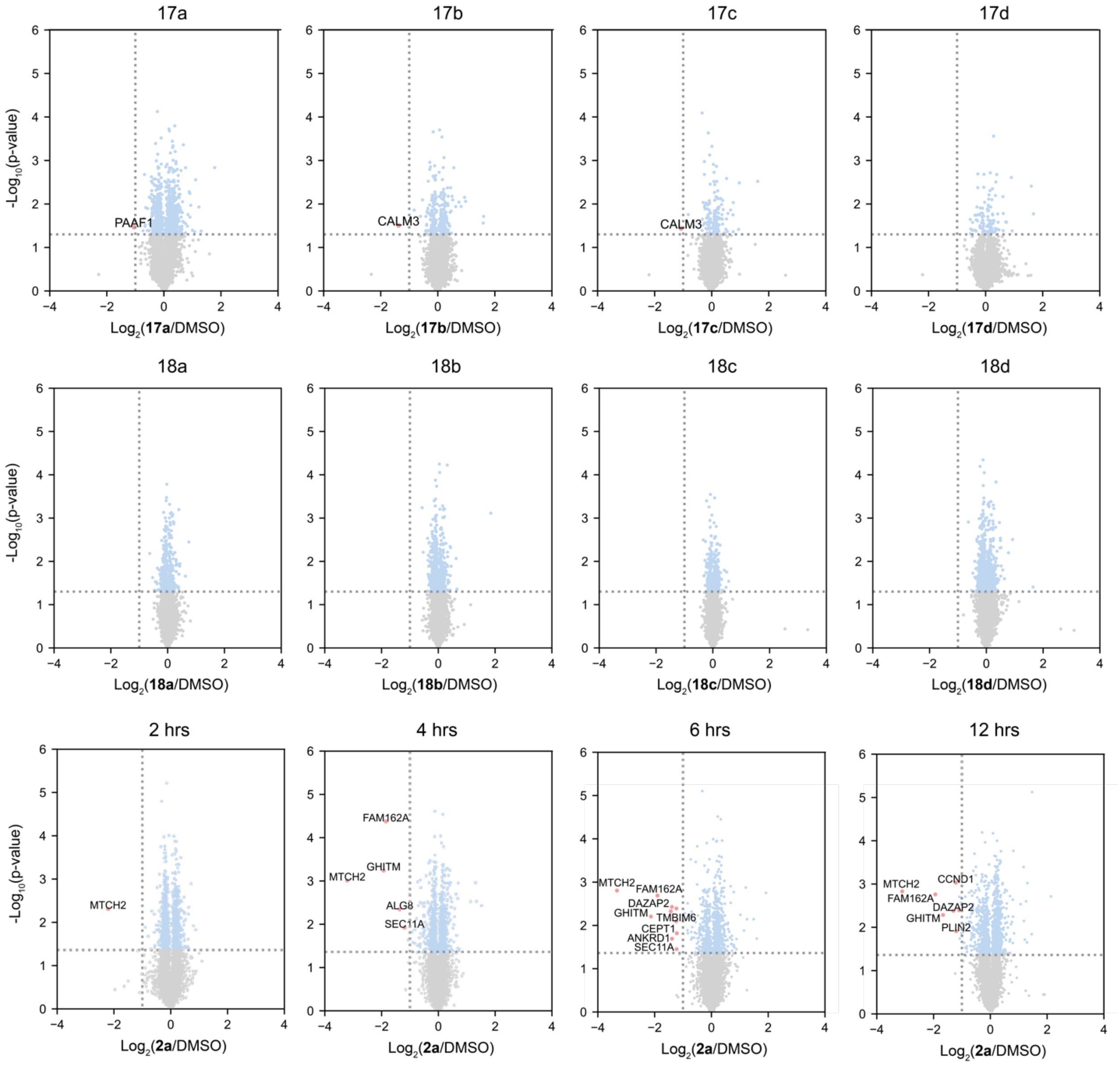


**Fig. S4| Proteomic profiles of AgnoTAC library.** Volcano plots of proteomic changes upon AgnoTAC treatment (18h). Detailed data sets and full list of downregulated targets can be found in **supplemental dataset 1**.

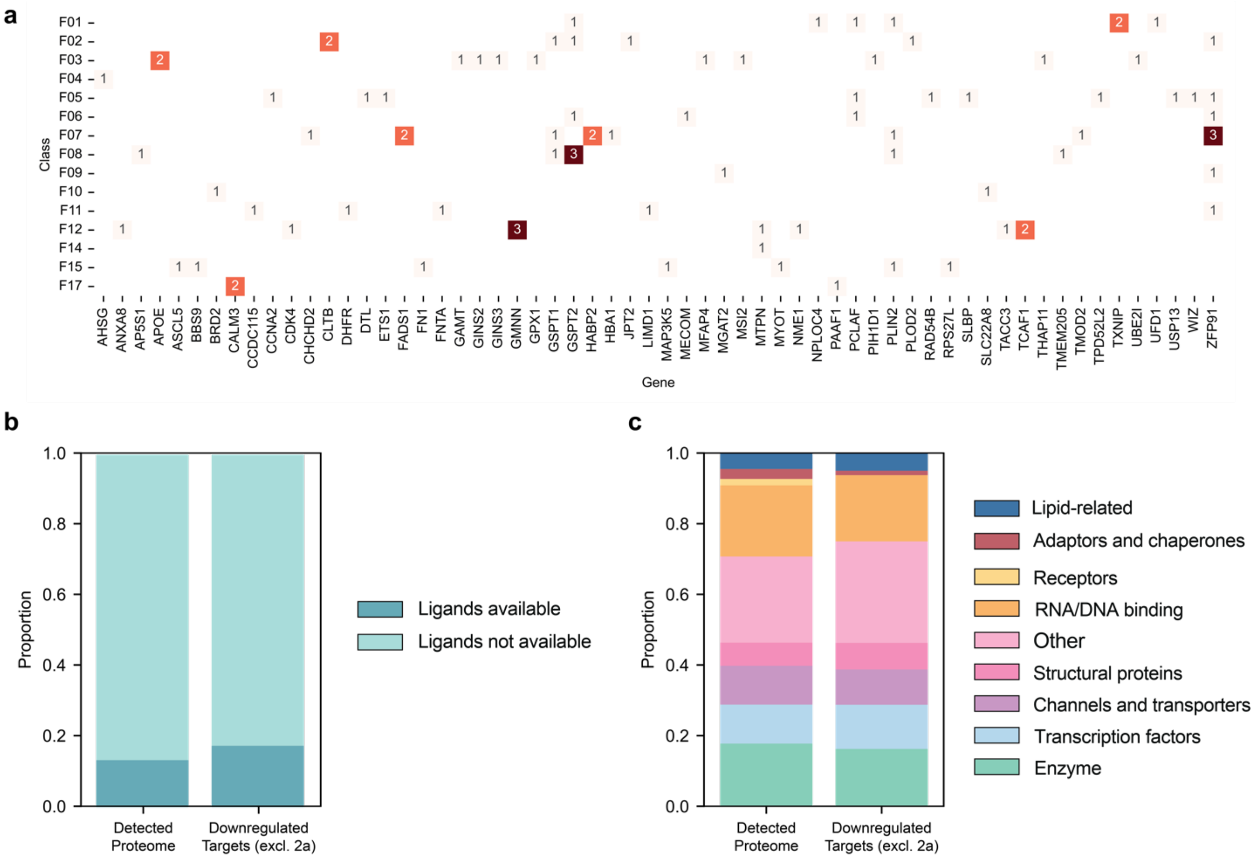


**Fig. S5| Analysis of detected and downregulated targets.** (**a**) Heatmap of unique targets significantly downregulated across headgroups. (**b, c**) Bar graphs comparing the proportion of downregulated targets (excluding cytotoxic AgnoTAC **2a**) to all detected proteins in MDA-MB-231 cells classified by (**b**) the availability of chemical probes as assigned by the DrugBank database and (**c**) protein functional classes as assigned by UniProt. The "Other" category refers to motor, metal-ion binding, carbohydrate-binding, protein-binding, heme-related, fatty acid metabolism-related proteins and uncharacterized proteins.

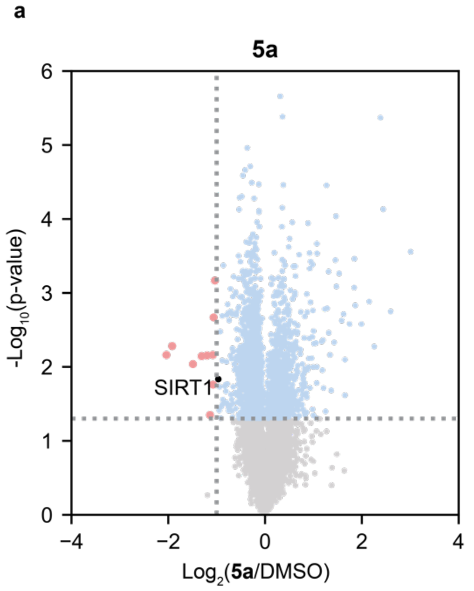


b

| Protein | SIRT1 |
|---|---|
| Fold Change (Ratio) | 0.51 |
| *p*-Value | 0.015 |
| Unique Peptides | 3 |
| PSMs | 4 |

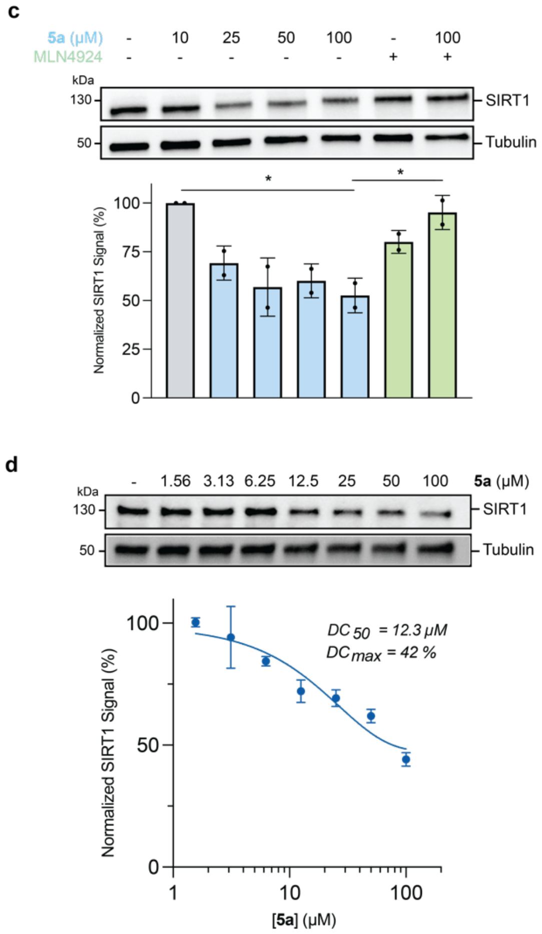


**Fig. S6| Additional target degraded by AgnoTAC 5a.** (**a**) Volcano plot depicting global protein abundance changes upon **5a** treatment of MDA-MB-231 cells for 18 h. SIRT1 downregulation is highlighted by a black dot. (**b**) Fold change, p-value, unique peptide and peptide-spectrum matches (PSMs) for SIRT1 deacetylase. (**c, d**) Dose-dependent, proteasome, and neddylation-mediated degradation of SIRT1.

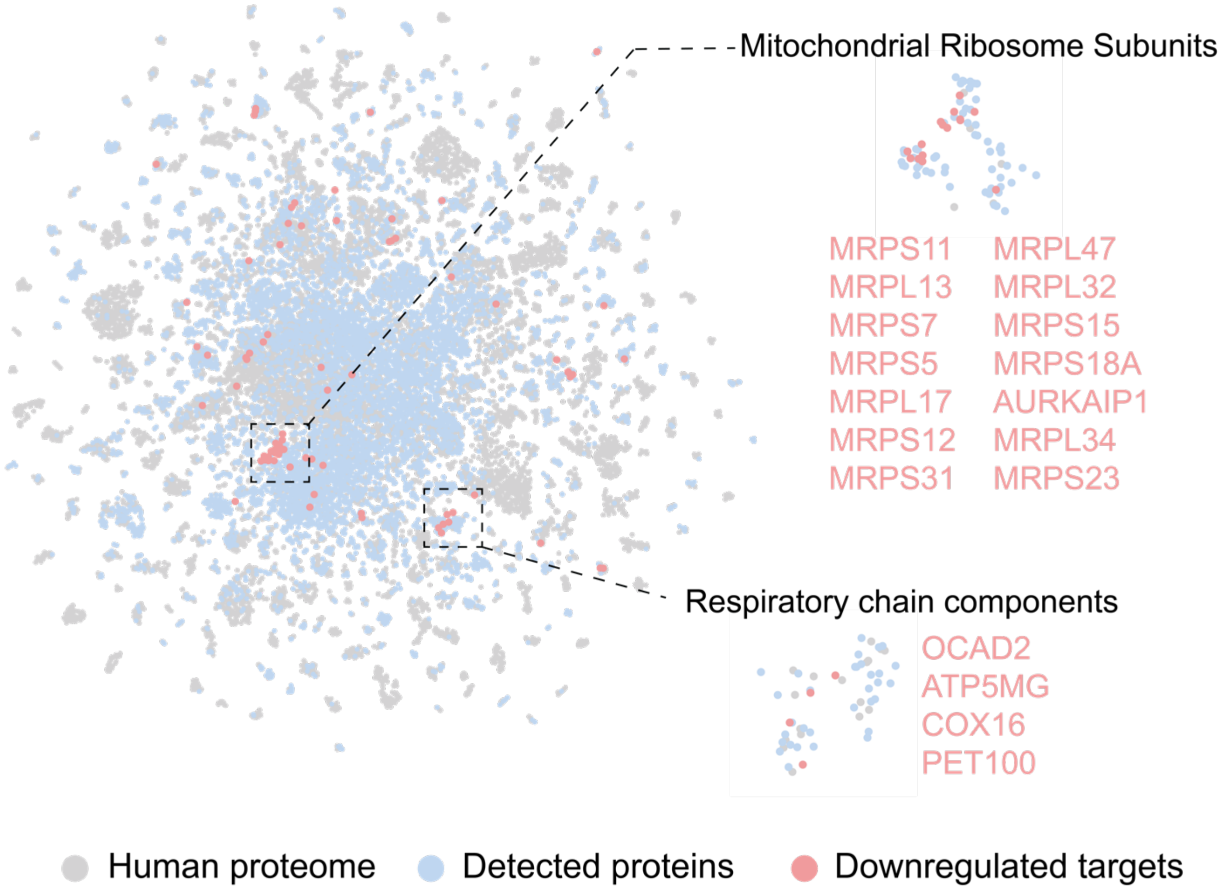


**Fig. S7| Protein cluster analysis of 2a-treated samples**. 2D UMAP projection illustrating clustered proteins following MDA-MB-231 cell treatment with 2a for 18 h. Gray dots represent all human proteins, blue dots represent detected proteins, and red dots highlight downregulated targets. Dashed insets emphasize protein clusters.

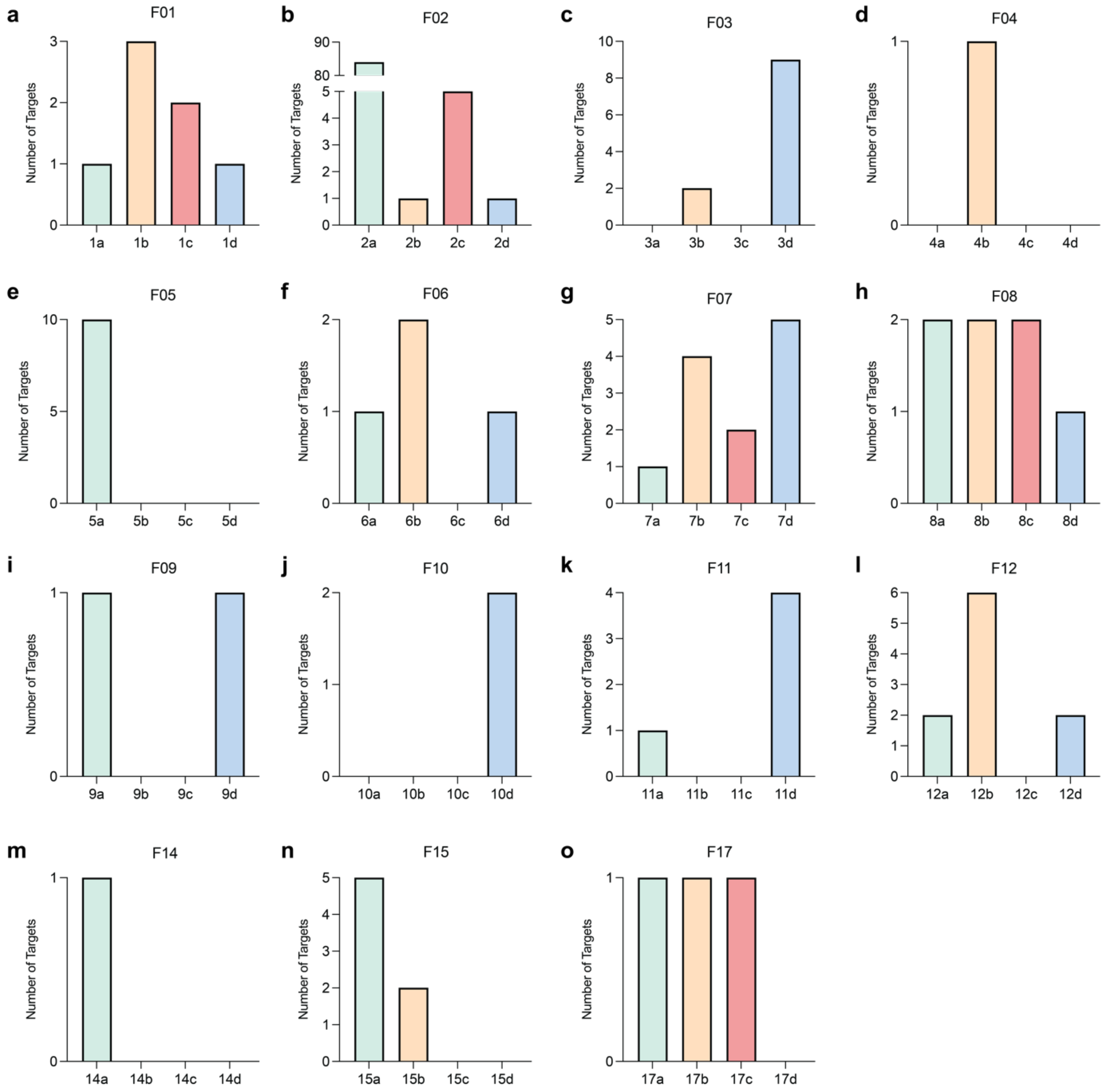


**Fig. S8| Linker composition influences degradation profiles, related to Fig. 5.** (**a-o**) Bar plots depicting total number of targets downregulated by each probe across all 18 headgroups. AgnoTACs corresponding to headgroups not displayed (i.e., F13, F16 and F18) did not significantly downregulate any target. Summary of results (excluding AgnoTAC 2a) is provided in **Fig. 3a.** Associated data can be found in **supplemental dataset 2**.

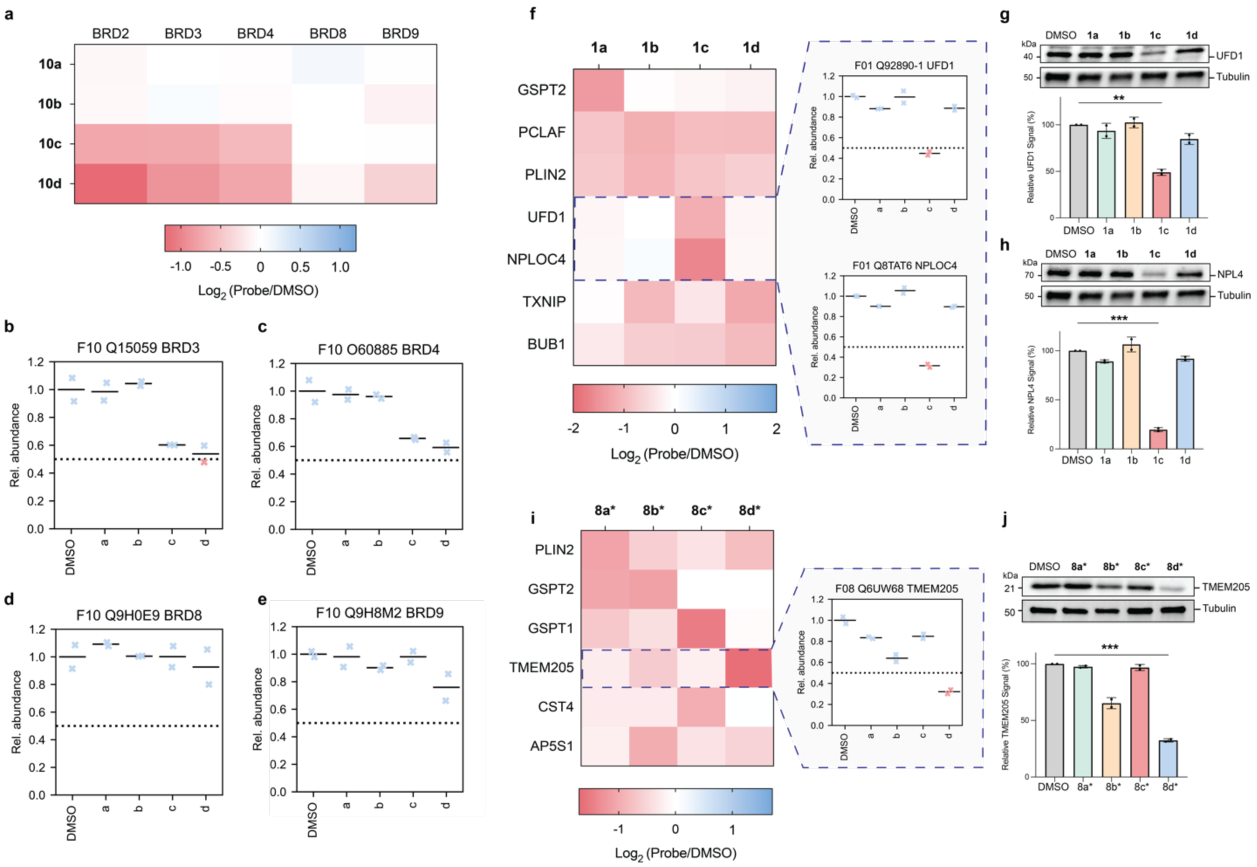


**Fig. S9| Additional linker effects, related to Fig. 5.** (**a**) Heatmap of additional bromodomain proteins identified in F10 AgnoTAC-treated samples. (**b-e**) Mean proteomic abundance ratios for (**b**) BRD3, (**c**) BRD4, (**d**) BRD8 and (**e**) BRD9 proteins. Both BRD3 and BRD4 are downregulated by F10 PEG-based linker AgnoTACs (**10c** and **10d**), following the same trends as BRD2 (**Fig. 3f, g**). (**f-j**) Additional degradability heatmaps and abundance ratios highlighting linker-dependent changes in proteomic profiles for (**f-h**) UFD1/NPLOC4, and (**i, j**) TMEM205. Immunoblots were obtained after 18 h AgnoTAC treatments. Error bars in bar plots represent mean ± SD (n = 2 biological replicates).

**a**

| Gene | Accession | AA_Start | AA_End | Turn Query | Peptide | Turn Overlap | Control Align | Query Align | Overlap Align | RMSD |
|---|---|---|---|---|---|---|---|---|---|---|
| **ZFP91** | Q96JP5 | 400 | 409 | LQCEICGFT | LQCEICGFTC | *..**. | FQCNQCGASF | LQCEICGFTC | .**..**... | 0.192818418 |
| **ZFP91** | Q96JP5 | 430 | 439 | FSCNICGKK | FSCNICGKKF | **.**. | FQCNQCGASF | FSCNICGKKF | *.**.**..* | 0.55828917 |
| **GSPT2** | Q8IYD1 | 548 | 593 | LVDKKSGEK | CIEEVEITALISLVDKKSGEKSKTRPRFVKQDQVCIARLRTAGTI | ....*. | ------------FQCNQCGASF------------------------ | CIEEVEITALISLVDKKSGEKSKTRPRFVKQDQVCIARLRTAGTI | ......*... | 0.443048328 |
| **MECOM** | Q03112 | 348 | 357 | HACPECGKT | HACPECGKTF | *..**. | FQCNQCGASF | HACPECGKTF | ..*..**..* | 0.372687995 |
| **MECOM** | Q03112 | 405 | 414 | IKCKDCGQM | IKCKDCGQMF | *..**. | FQCNQCGASF | IKCKDCGQMF | ..*..**..* | 0.60982269 |
| **MECOM** | Q03112 | 912 | 921 | YTCRYCGKI | YTCRYCGKIF | *..**. | FQCNQCGASF | YTCRYCGKIF | ..*..**..* | 0.435923904 |
| **GSPT1** | P15170 | 419 | 442 | LVDKKSGEK | CIEEVEITALICLVDKKSGEKSKT | ....*. | ------------FQCNQCGASF-- | CIEEVEITALICLVDKKSGEKSKT | ......*.. | 0.422952771 |
| **CDK4** | P11802 | 18 | 40 | ARDPHSGHF | GTVYKARDPHSGHFVALKSVRVP | ....*. | -----FQCNQCGASF-------- | GTVYKARDPHSGHFVALKSVRVP | ......*... | 0.737835705 |
| **FN1** | P02751 | 96 | 107 | CFDKYTGNT | TCFDKYTGNTYR | ....*. | -FQCNQCGASF- | TCFDKYTGNTYR | ......*... | 0.959801316 |
| **FN1** | P02751 | 560 | 571 | CQDSETGTF | QCQDSETGTFYQ | ....*. | -FQCNQCGASF- | QCQDSETGTFYQ | .*....*... | 0.718473136 |
| **PAAF1** | Q9BRP4 | 116 | 130 | IWQASNGEL | MKIWQASNGELRRVL | ....*. | --FQCNQCGASF--- | MKIWQASNGELRRVL | ......*.. | 0.382800102 |
| **MAP3K5** | Q99683 | 809 | 820 | LINTYSGVL | VLINTYSGVLKI | ....*. | -FQCNQCGASF- | VLINTYSGVLKI | ......*... | 0.534017503 |
| **WIZ** | O95785 | 304 | 313 | LACGECGWA | LACGECGWAF | *..**. | FQCNQCGASF | LACGECGWAF | ..*..**..* | 0.494413674 |
| **WIZ** | O95785 | 416 | 425 | SACVFCGFP | SACVFCGFPA | *..**. | FQCNQCGASF | SACVFCGFPA | ..*..**.. | 0.304864138 |
| **WIZ** | O95785 | 870 | 879 | TTCEVCGAC | TTCEVCGACF | *..*** | FQCNQCGASF | TTCEVCGACF | ..*..***.* | 0.403276116 |
| **WIZ** | O95785 | 1043 | 1052 | IRCEFCGEF | IRCEFCGEFF | *..**. | FQCNQCGASF---------- | IRCEFCGEFFIRCEFCGEFF | ..*..**..* | 0.516328692 |
| **WIZ** | O95785 | 1227 | 1236 | IRCEFCGEF | IRCEFCGEFF | *..**. | FQCNQCGASF---------- | IRCEFCGEFFIRCEFCGEFF | ..*..**..* | 0.516328692 |

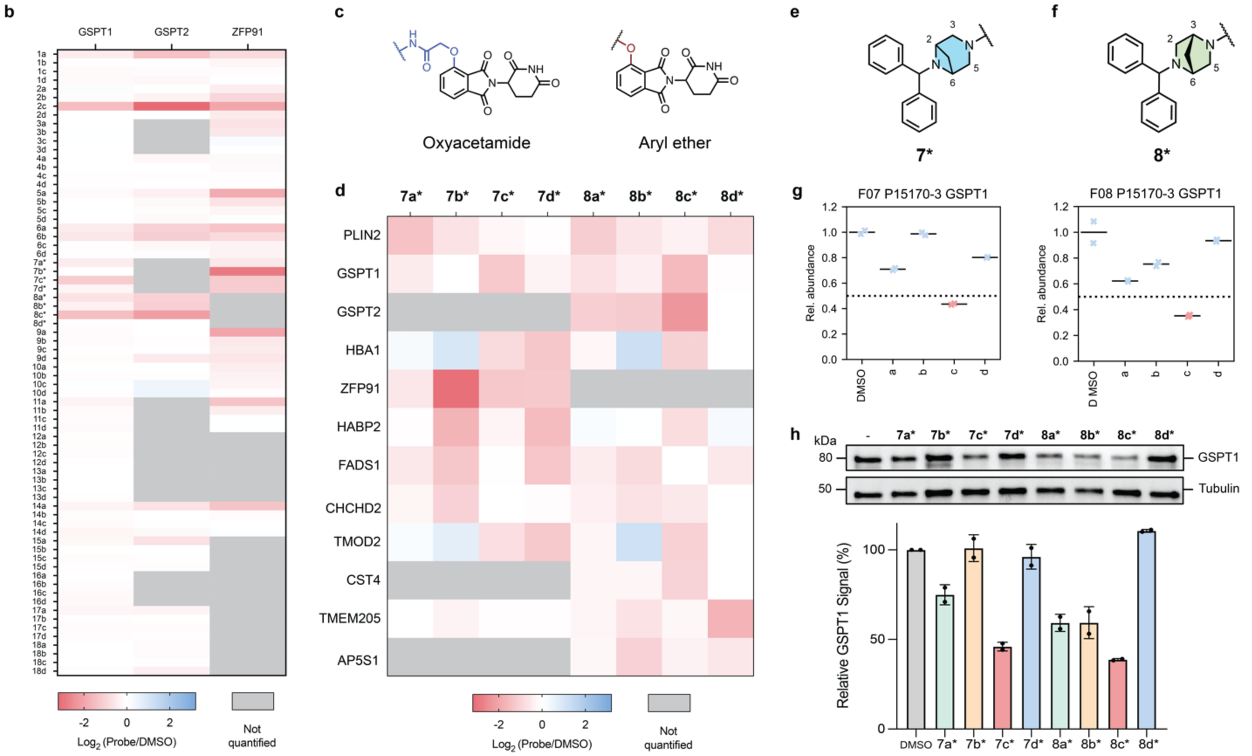


**Fig. S10| Analysis of molecular glue off-targets.** (**a**) Heatmap depicting changes in the abundances of known molecular glue targets, including GSPT1, GSPT2 and ZFP91, across all 72 AgnoTACs. (**b**) Structural comparison of 4-oxyacetamide and 4-hydroxythalidomide (aryl ether) moieties. (**c**) Heatmap displaying $\log_2$(fold-change) values of proteins targeted by structurally similar **F7***- and **F8***-based AgnoTAC analogs. (**d, e**) Chemical structures of **F7*** and **F8*** headgroups. (**f**) Comparison of GSPT1 mean proteomic ratios following treatment with **F7*** and **F8***. (**g**) Immunoblot validation of GSPT1 downregulation by **F7***- and **F8***. Error bars in bar plot represent mean ± SD (n = 2 biological replicates). All experiments were performed in MDA-MB-231 cells treated for 18 h. (**h**) Table of AgnoTAC targets possessing motifs that resemble β-hairpin G-loop structures.

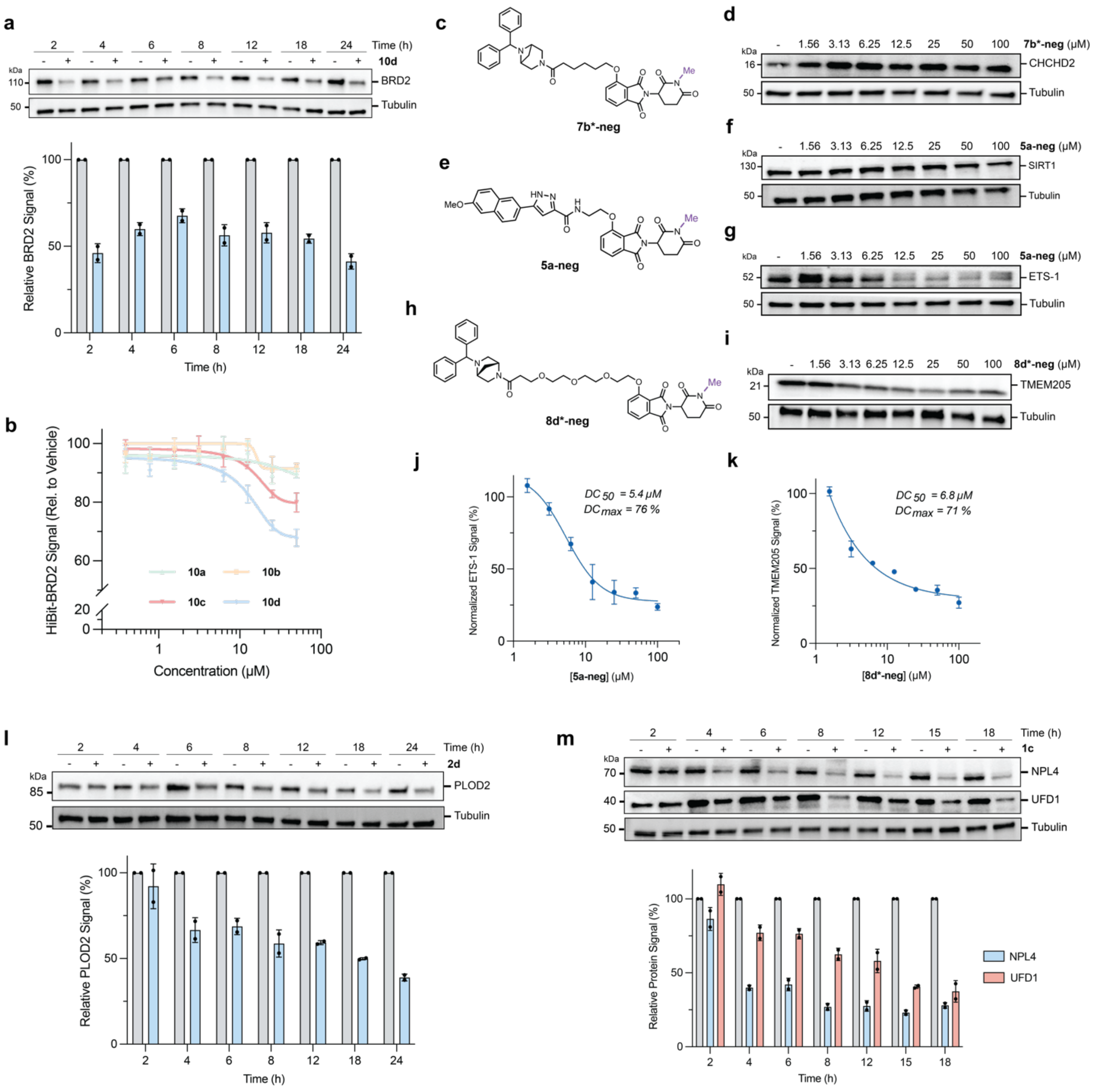


**Fig. S11| Additional mechanistic characterization of AgnoTAC targets.** (**a**) Time-course analysis of BRD2 degradation following treatment of MDA-MB-231 cells with 50 µM **10d** at indicated times and quantitative analysis (normalized to loading control). (**b**) BRD2 degradation assessed by HiBiT assay. HEK293t cells containing endogenously tagged HiBiT-BRD2 were treated with 50 µM of **10a**, **10b**, **10c** or **10d**. Data was normalized to DMSO control. Errors bars are expressed as mean ± SD (n = 3 biological replicates). (**c-i**) Chemical structures of additional negative controls and corresponding immunoblot dose-response analyses for (**c, d**) CHCHD2 upon **7b*-neg** treatment, (**e-g**) SIRT1 and ETS-1 following **5a-neg** treatment, and (**h, i**) TMEM205 levels after **8d*-neg** treatment. (**j, k**) Dose-dependent depletion of (**j**) ETS-1 by **5a-neg** and (**k**) TMEM205 by **8d*-neg**, highlighting Cullin-independent degradation mechanism. All shown results were obtained after 18 h treatments of MDA-MB-231 cells at indicated concentrations. Dose response curves for ETS-1 and TMEM205 are representative of n = 2 biological replicates. (**l, m**) Time-course analyses of (**l**) PLOD2 degradation by **2d** (50 µM) and (**m**) NPL4/UFD1 degradation by **1c** (40 µM) at indicated times in MDA-MB-231 cells.

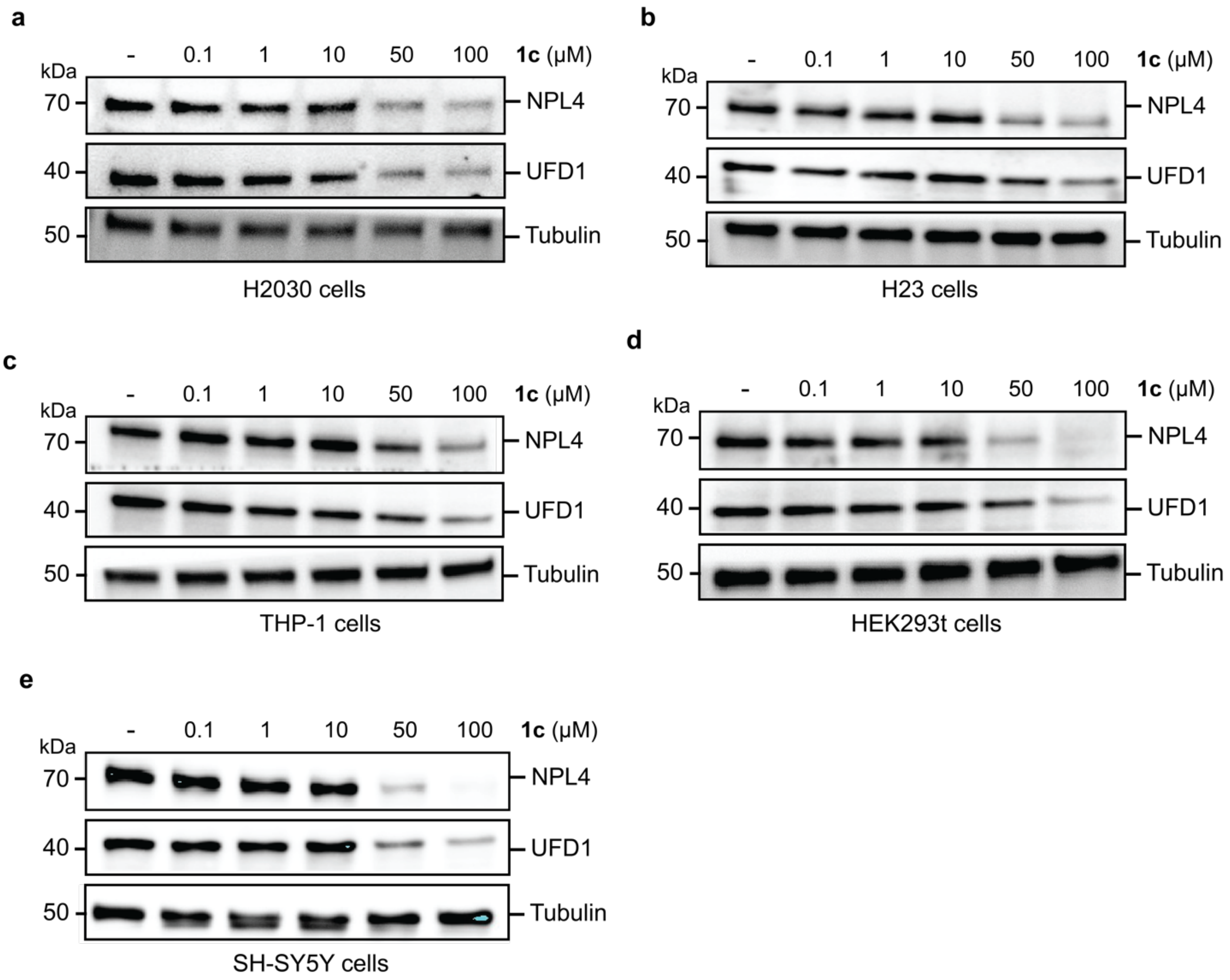


**Fig. S12| AgnoTAC 1c degrades UFD1/NPL4 in several cell lines.** Immunoblot analysis of NPL4 and UFD1 downregulation in (**a**) H2030 and (**b**) H23 non-small lung cancer cells, (**c**) THP-1 leukemia monocytes, (**d**) HEK293t kidney epithelial cells, and (**e**) SH-SY5Y neuroblastoma cells upon **1c** treatment at indicated concentrations for 6 h.

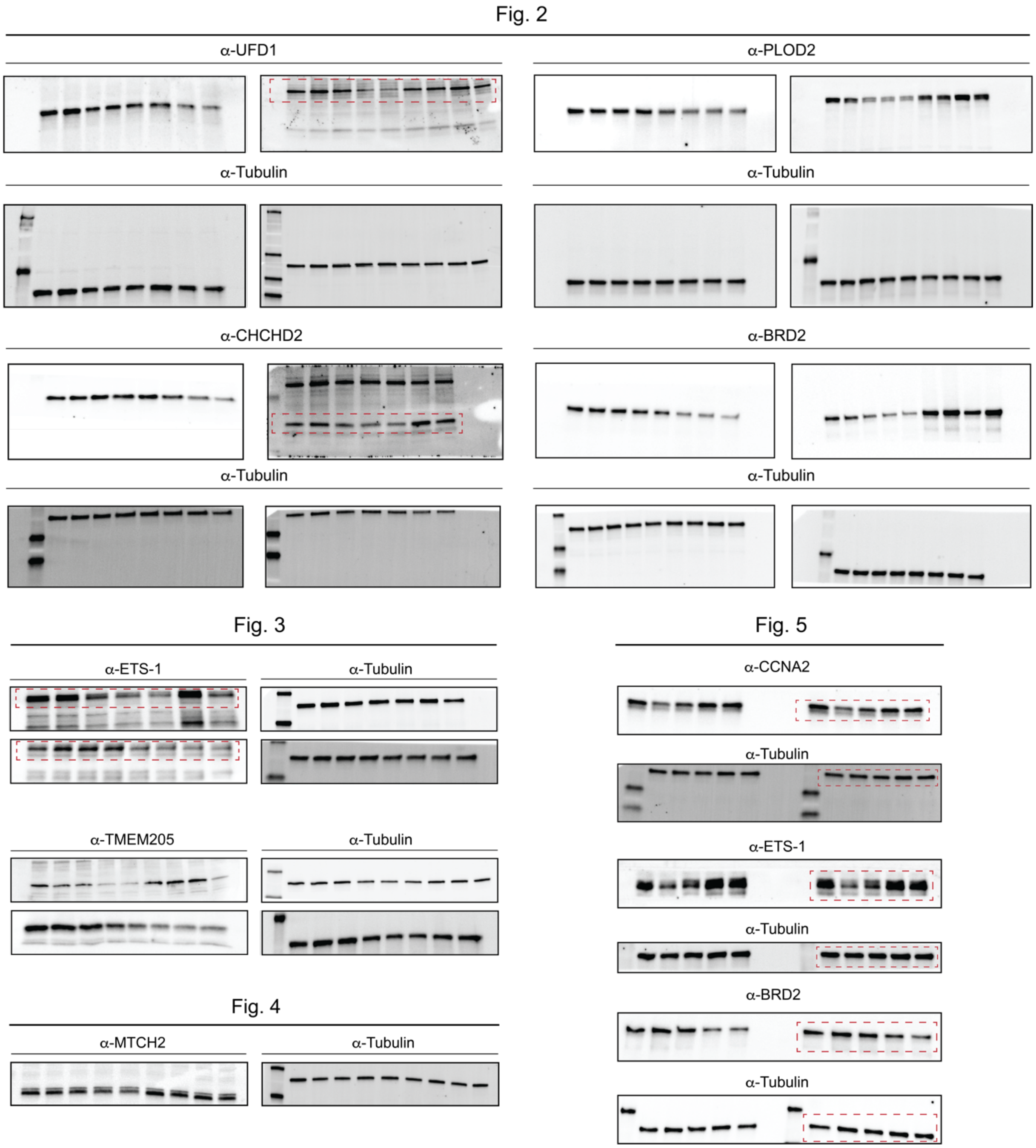


**Fig. S13| Compiled uncropped Western blots (continued on next page)**

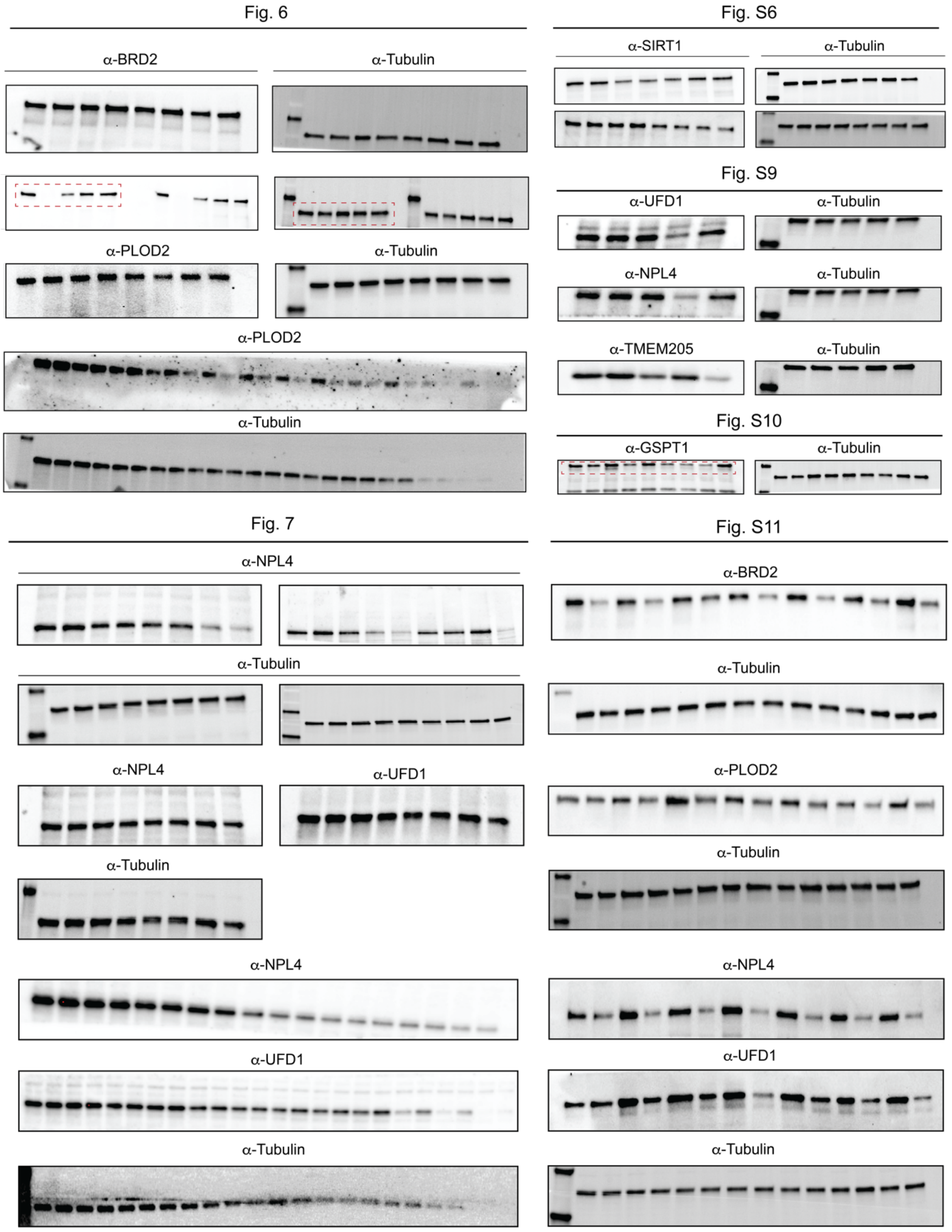


**Fig. S13| Compiled uncropped Western blots (continued on next page)**

#### Fig. S11 (continued)

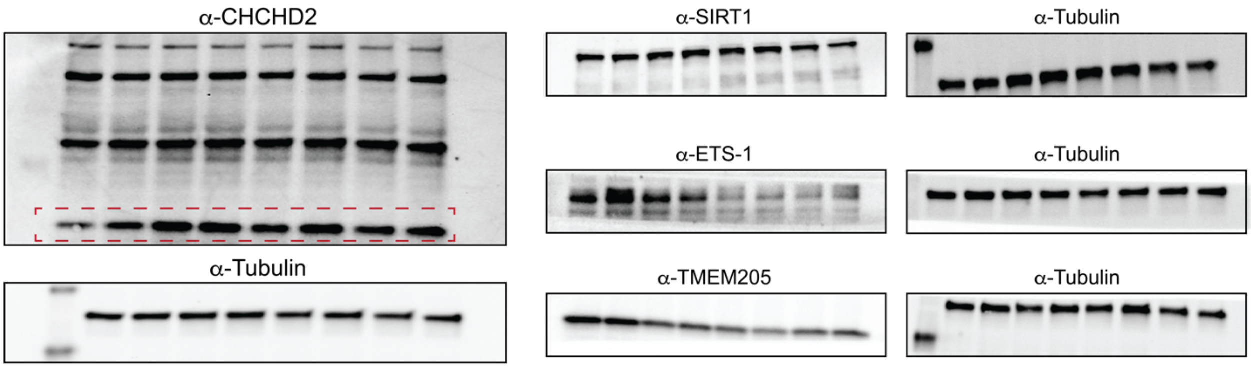


#### Fig. S12

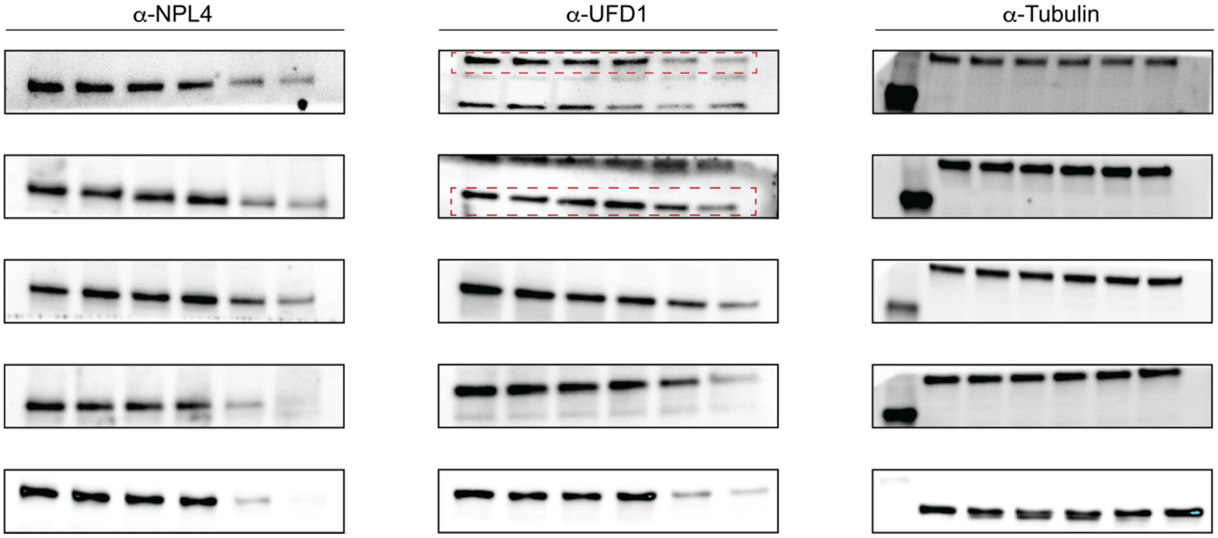


**Fig. S13| Compiled uncropped Western blots.** Red insets indicate correct bands.

#### SUPPLEMENTARY TABLES

| REAGENT | SOURCE | IDENTIFIER |
|---|---|---|
| **ANTIBODIES** | | |
| Rabbit anti-UFD1 (1:1,000 dilution), | Cell Signaling Technology | 13789S |
| Rabbit anti-PLOD2 (1:1,000 dilution) | Cell Signaling Technology | 44709S |
| Rabbit anti-CHCHD2 (1:2,000 dilution) | Proteintech | 19424-1-AP |
| Rabbit anti-BRD2 D89B4 clone (1:1,000 dilution) | Cell Signaling Technology | 5848S |
| Rabbit anti-SIRT1 D1D7 clone (1:1,000 dilution) | Cell Signaling Technology | 9475S |
| Rabbit anti-ETS-1 D8O8A clone (1:1,000 dilution) | Cell Signaling Technology | 14069S |
| Rabbit anti-TMEM205 (1:1,000 dilution) | Invitrogen | PA5-59599 |
| Rabbit anti-CCNA2 E6D1J clone (1:1,000 dilution) | Cell Signaling Technology | 67955S |
| Rabbit anti-MTCH2 (1:10,000 dilution) | Proteintech | 16888-1-AP |
| Rabbit anti-NPL4 (1:1,000 dilution) | Cell Signaling Technology | 13489S |
| Rabbit anti-GSPT1 (1:2,000 dilution) | Proteintech | 10763-1-AP |
| Rabbit anti-HA Tag C29F4 clone (1:2,000 dilution) | Cell Signaling Technology | 3724S |
| Rabbit anti-Ubiquitin E4I2J clone (1:2,000 dilution) | Cell Signaling Technology | 43124S |
| Anti-tubulin hFAB rhodamine IgG (1:10,000 dilution) | Bio-Rad | 12004165 |
| Goat anti-Rabbit IgG, HRP (1:10,000 dilution) | Thermo Fisher Scientific | 31460 |
| **EXPERIMENTAL MODELS: CELL LINES** | | |
| MDA-MB-231 | ATCC | HTB-26 |
| HEK293t | ATCC | CRL-3216 |
| SH-SY5Y | ATCC | CRL--2266 |
| NCI-H2030 | ATCC | CRL-5914 |
| NCI-H23 | ATCC | CRL-5800 |
| THP-1 | ATCC | TIB-202 |
| **COMMERICAL ASSAYS, KITS AND REAGENTS** | | |
| CellTiter-Glo Luminescent Cell Viability Assay | Promega | G7570 |
| Nano-Glo HiBiT Lytic Detection System | Promega | N3030 |
| DC Protein Assay | Bio-Rad | 5000112 |
| Pierce High pH Reversed-Phase Fractionation Kit | Thermo Fisher Scientific | 84868 |
| Pierce Quantitative Fluorometric Peptide Assay Kit | Thermo Fisher Scientific | 23290 |
| TMT10plexTM Isobaric label reagent set | Thermo Fisher Scientific | 90110 |
| Sequencing Grade Modified Trypsin | Promega | V5111 |
| Endoproteinase LysC | New England Biolabs | P8109 |
| Halt Protease Inhibitor | Thermo Fisher Scientific | 78438 |
| ProteaseMAX™ Surfactant, Trypsin Enhancer | Promega | V2072 |
| Pierce C18 Spin Columns | Thermo Fisher Scientific | 89873 |
| Iodoacetamide (IAA) | Sigma-Aldrich | 16125-10G |
| Tetraethylammonium bromide (TEAB) | Thermo Scientific | 9C114 |
| Trifluoroacetic acid (TFA) | Thermo Scientific | 85183 |
| Formic acid (FA) | Fisher | A117-50 |
| 50% wt. $H_2O$ Hydroxylamine | Sigma-Aldrich | 467804 |
| Urea, Molecular Biology Grade | Sigma-Aldrich | U5128 |
| Tris(2-carboxyethyl)phosphine (TCEP) | Research Products International | T26500-10 |
| Potassium carbonate ($K_2CO_3$) | Sigma-Aldrich | 209619 |
| Methanol, Optima LC-MS Grade | Fisher Scientific | A456-4 |
| Chloroform, HPLC-Grade | Fisher Scientific | C607-4 |
| Acetonitrile, Optima LC-MS Grade | Fisher Scientific | A955-4 |
| Water, Optima LC-MS Grade | Fisher Scientific | W64 |
| Calcium chloride ($CaCl_2$), ACS Grade | Sigma-Aldrich | 2082904X |
| 4X Laemmli SDS Reducing Sample Buffer | BioWorld | 10570020 |

| | | |
|---|---|---|
| 4-20% Mini-PROTEAN TGX Precast Gels, 10 wells | Bio-Rad | 4561093 |
| 4-20% Criterion TGX Precast Midi gels, 26 wells | Bio-Rad | 5671095 |
| Dulbecco's phosphate-buffered saline (DBPS) | Corning | 21-031-CV |
| Tween-20 | Millipore Sigma | 655205 |
| Trans-Blot Turbo RTA transfer Kit, LF, PVDF | Bio-Rad | 1704275 |
| (S)-MG132 | MedChem Express | HY-13259 |
| Pevonedistat (MLN4924) | MedChem Express | HY-70062 |
| Anti-HA Magnetic Beads | MedChem Express | HY-K0201 |
| Lipofectamine™ 3000 Transfection Reagent | Thermo Fisher Scientific | L3000015 |
| UltraPure 1M Tris-HCl Buffer, pH 7.5 | Invitrogen | 15567027 |
| ZymoPURE Plasmid Miniprep Kit | Zymo Research | D4020 |
| Miller's LB Broth | Corning | 46-050-CM |
| Carbenicillin, Disodium Salt, 5 Grams | Research Products International | C46000-5.0 |
| Dulbecco's Modified Eagle Medium (DMEM) | Gibco | 11995-065 |
| Roswell Park Memorial Institute (RPMI) Medium | Gibco | 21870-076 |
| Opti-MEM™ I Reduced Serum Medium | Gibco | 31985070 |
| Fetal Bovine Serum (FBS) | Omega Scientific | FB-01 |
| L-Glutamine 200 mM solution | Corning | 25-005-CI |
| Penicillin/Streptomycin 100X solution | Corning | 30-002-CI |
| **SOFTWARES AND ALGORITHMS** | | |
| Proteome Discoverer (Version 3.0) | Thermo Fisher Scientific | - |
| Prism (Version 10.1.1) | GraphPad | - |
| Xcalibur (Version 4.1.50) | Thermo Fisher Scientific | - |
| ImageLab (Version 6.1.0) | Bio-Rad Laboratories | - |
| ChemDraw (Version 22.2.0) | Revvity Signals | - |
| MestReNova (Version 14.0.0) | Mestrelab Research | - |
| Excel (Version 16.93.1) | Microsoft | - |
| Python (Version 3.11.3 ) | Python Software Foundation | - |
| Illustrator 2025 (Version 29.1) | Adobe | - |

**Table S1| Reagents, kits, assays, antibodies and softwares used in this study.**

#### BIOLOGICAL METHODS

##### Cell Culture

MDA-MB-231 cells (ATCC) and HEK293t cells (ATCC) were maintained in DMEM supplemented with 10% (v/v) fetal bovine serum (FBS), 1% (v/v) penicillin/streptomycin, and 2mM glutamine. SH-SY5Y cells (gift from the Hyeryun Choe lab) were maintained in DMEM supplemented with 20% (v/v) fetal bovine serum (FBS), 1% (v/v) penicillin/streptomycin, and 2mM glutamine. NCI-H2030 cells (ATCC), NCI-H23 cells (ATCC) and THP-1 cells (ATCC) were maintained in RPMI 1640 supplemented with 10% (v/v) fetal bovine serum (FBS), 1% (v/v) penicillin/streptomycin, and 2mM glutamine. All cells were grown at 37°C in a humidified 5% $CO_2$ atmosphere.

##### Live Cell Treatment

Cells were grown to 80-95% confluency in appropriate medium. The growth medium was aspirated, and the cells were incubated with media containing compounds at indicated concentrations and time points. The cultures were scraped, washed with cold DPBS, and collected into 15 mL centrifuge tubes, then transferred to 1.5 mL Eppendorf tubes. The cell suspensions were centrifuged (1,400 g, 3 min) and the pellets were stored at -80 °C until the next stage of processing.

##### Quantitative Proteomics Sample Preparation

Cell pellets were resuspended in 350 µL DPBS containing 1 x Halt protease inhibitor cocktail, and lysed by sonication (15 ms on, 40 ms off, 15% amplitude, 1 s total). Protein concentrations were normalized (2 mg/mL in 100 µL with cold DPBS) using the Lowry Protein Assay (Pierce). Urea (48 mg) was weighed for each sample in 2 mL LoBind Eppendorf tubes and cell lysates (100 µL = 200 µg) were added (final urea concentration = 8 M). Pellets were resuspended in 50 µL of a freshly prepared 1:1 solution of TCEP (200 mM in DPBS) and $K_2CO_3$ (600 mM in DPBS) and incubated for 30 min at 37 °C) while shaking. After reaction, 70 µL of a solution of freshly prepared iodoacetamide (70 µL, 400 mM in DPBS) was added and incubated for 30 min at room temperature while protected from light. After reaction, 1.8 mL of cold 3:1:2 MeOH/$CHCl_3$/$H_2O$ solution was added to each tube, and the samples were centrifuged (10,000 x *g,* 10 min, 4 °C), forming a disc. The supernatant was carefully removed, MeOH (600 µL) was added, samples were centrifuged as previously described and the supernatant was removed. Cell pellets were resuspended in TEAB (160 µL, 100 mM, pH 8.5) and sonicated as described previously. Endoproteinase LysC (20 µL, 1/2 vial, 10 µg dissolved in 220 µL of 100 mM TEAB pH 8.5) was added to each and incubated at 37°C for 2 h with shaking. Sequencing-grade modified porcine trypsin (20 µL, 1 vial, 20 µg dissolved in 220 µL of 100 mM TEAB pH 8.5), a solution of Protease Max (2 µL, 1% (w/v) in 100 mM TEAB pH 8.5) and a solution of $CaCl_2$ (2 µL, 100 mM) were added to the samples and incubated at 37 °C overnight (14 h) with shaking. The digest was separated by centrifugation (12,000 *x g*, 10 min, 4 °C). Peptide concentration was determined using Pierce Quantitative Fluorometric Peptide Assay Kit (Thermo Fisher Scientific, 23290) according to manufacturer's instructions. MS-grade acetonitrile (12.5 µL) was added to each tube, and samples were labeled with respective TMT 10 plex or 16-plex isotope (8 µL, 20 µg/ µL) for 1 h with occasional vortexing at RT. To quench, hydroxylamine (3 µL, 5% *v/v*) was added to each sample, vortexed, and incubated for 15 min at RT. Formic acid (5 µL) was added to each tube for acidification, and samples were dried under vacuum centrifugation. The samples were combined by redissolving the contents of one tube in a solution of trifluoroacetic acid (TFA, 400 µL, 0.1% in water) and transferred into each sample until all samples were redissolved. The stepwise process was repeated with formic acid (Buffer A, 200 µL, 0.1% in water) for a final volume of 600 µL. The samples were fractionated using a fractionation kit (Pierce high pH Reversed-Phase Fractionation Kit, Thermo Fisher Scientific 84868) according to manufacturer's instructions. The peptide fractions were eluted from the spin column with consecutive solutions of 0.1% triethylamine combined with MeCN (5-50% MeCN). The fractions were combined pairwise (fraction 1 and fraction 10, fraction 2 and fraction 11, etc.), dried via vacuum centrifugation, and stored at -80 °C until ready for mass spectrometer injection.

#### LC-MS Analysis of TMT-Labeled Samples

TMT labeled samples were redissolved in MS buffer A (65 µL, 0.1% formic acid in water). 10 µL of each sample was loaded onto an Acclaim PepMap 100 precolumn (75 µm x 2 mm) and eluted on an Acclaim PepMap RSLC analytical column (75 µm x 15 cm) using the UltiMate 3000 RSLCnano system (Thermo Fisher Scientific). Buffer A was prepared as described above and buffer B (0.1% formic acid in MeCN) were used in a 220 min gradient (flow rate 0.3 mL min, 35 °C) of 2 % buffer B for 10 min, followed by an incremental increase to 30 % buffer B over 192 min, 60 % buffer B for 5 min, 60-95 % buffer B for 1 min, hold at 95 % buffer B for 5 min, followed by descent to 2% buffer B for 1 min followed by re-equilibration at 2 % for 6 min. The elutions were analyzed with a Thermo Fisher Scientific Orbitrap Fusion Lumos mass spectrometer with a cycle time of 3 s and nano-LC electrospray ionization source applied voltage of 2.0 kV. $MS^1$ spectra were recorded at a resolution of 120,000 with an automatic gain control (AGC) value of $1x10^6$ ions, maximum injection time of 50 ms (dynamic exclusion enabled, repeat count 1, duration 20 s). The scan range was specified from 375 to 1,500 m/z. Peptide fragmentation $MS^2$ spectra was recorded via collision-induced diffusion (CID) and quadrupole ion trap analysis (AGC $1.8x10^4$, 30 % collision energy, maximum inject time 120 ms, isolation window 1.6). MS3 spectra were generated by high-energy collision-induced dissociation (HCD) with collision energy of 65 %. Precursor selection included up to 10 $MS^2$ ions for the $MS^3$ spectrum.

#### Proteomics Data Processing

Raw MS-data analysis was performed using processing software Proteome Discoverer 3.0 (Thermo Fisher Scientific). Peptide sequences were identified by matching proteome databases with experimental fragmentation patterns via the SEQUEST HT algorithm. Fragment tolerances were set to 0.6 Da, and MS1 precursor mass tolerances set to 10 ppm with one missed cleavage site allowed. Trypsin specificity was designated with a maximum of one missed cleavage. Variable amino acid modifications included methionine oxidation (M, +15.994915) and cysteine carbamidomethylation (C, +57.02146), while TMT-tags (K and N-termini, +229.1629 for 10-plex) were specified as fixed modifications. Spectra were searched against the *Homo Sapiens* proteome database (74,782 sequences) using a false discovery rate (FDR) of 1 % (Percolator). $MS^3$ peptide quantitation of TMT reporter ions was performed with a mass tolerance of 20 ppm. Relative TMT ratios obtained by Proteome Discoverer were transformed with $\log_2(x)$, and statistical significance (p-values) were calculated via Student's two-tailed t-tests across two biological replicates (significance threshold was set at $p < 0.05$).

#### Cell Viability Assays

Cells were seeded in white-opaque 96-well plates in full growth media at a density of 5,000 cells/well (50 µL) and were allowed to grow for 24 h at 37°C in a humidified 5% $CO_2$ atmosphere. The cells were then treated with compounds in triplicate and incubated at 37°C in a humidified 5% $CO_2$ atmosphere for 18 hours. Cell viability was determined using the luciferase-based Cell Titer-Glo Luminescent Cell Viability Assay (Promega) following manufacturer's guidelines. Data represents the average and standard deviation of triplicates in measured luminescence.

#### Proteasome and Neddylation Inhibition Experiments

To evaluate the dependence of AgnoTAC-induced protein degradation on the proteasome and neddylation pathways, cells were pre-treated with 10 µM (S)-MG132 (proteasome inhibitor) or 1 µM MLN4924 (NEDD8-activating enzyme inhibitor) for 2 hrs. Following pre-incubation, media was aspirated, and cells were co-treated with inhibitors and the AgnoTAC at indicated time points and concentrations. Cell lysates were then collected and further analyzed by Western blotting.

#### Immunoblots (Western blots)

After cells were harvested, cell pellets were resuspended in 100 µL DPBS containing 1 x Halt protease inhibitor cocktail, and lysed by sonication (15 ms on, 40 ms off, 15% amplitude, 1 s total). Protein concentrations were normalized (2 mg/mL in 100 µL with cold DPBS) as previously described. 4X SDS gel loading buffer (33 µL) was added to the solution of protein lysate and the resulting mixture was heated at 95 °C for 10 min. Proteins (15 µg total protein loaded per gel lane) were resolved by SDS-PAGE (10 % acrylamide) made in-house, and transferred to PVDF membrane (0.2 µM, 1620177, Bio-Rad). The membrane was blocked with 5% BSA (w/v) in Tris-buffered saline with Tween (TBST) buffer (0.1% Tween 20, 20 mM Tris-HCl 7.6, 150 mM NaCl) at room temperature for 1 h. The antibody was diluted with fresh 5% BSA in TBST buffer (dilutions were performed following manufacturer's guidelines) and incubated with membrane overnight at 4 °C. Membrane was washed three times with TBST buffer, left 5 minutes between each wash on a rocker and then incubated with secondary antibody in 5% dry milk in TBST at room temperature for 2 h on a rocker. Membrane was washed three times with TBST buffer and visualized by in-gel fluorescence on a Bio-Rad ChemiDoc MP Imaging System. The images were processed using Image Lab (version 6.1.0) software.

##### HiBiT assay
HEK293t cells expressing HiBiT-BRD2 were generated as previously reported[1]. Degradation of HiBiT-BRD2 was quantified using the Nano-Glo HiBiT Lytic Detection System (Promega, #N3030) according to manufacturer protocol. Briefly, $2*10^3$ cells were plated in 30 µL complete medium (DMEM, 4 mM glutamine, supplemented with 10% FBS, penicillin 100 u/mL, and streptomycin 100 µg/mL) in a 384-well plate. Cells were incubated for 24 hrs prior to the experiment (37 °C/5% $CO_2$). Compounds (20 mM stock in DMSO) were diluted to 3x stocks into complete medium and serially diluted in medium maintaining a constant DMSO concentration (1.5%). 15 µL of these solutions were then added to cells in duplicate and incubated for 18 hrs (37 °C/5% $CO_2$). After equilibration at room temperature, 40 µL of Nano-Glo HiBiT lysis buffer freshly supplemented with LgBiT (1:100) and Nano-Glo Lytic substrate (1:50) were added to each well. Plate was shaken at 200 rpm for 10 min before imaging using a microplate reader (unfiltered luminescence, ClarioSTAR, BMG Labtech). Signal was normalized to control wells containing no compound as 100% HiBiT-BRD2.

##### Cellular Thermal Shift Assay (CETSA)
MDA-MB-231 cells ($1*10^7$ cells per milliliter for each condition) were centrifuged (400g, 4 min, 4°C), resuspended in growth medium supplemented with DMSO or indicated concentration of compounds, and incubated for 30 min at 37°C. Sample were then carefully washed with cold DPBS supplemented with 1x Halt protease inhibitor cocktail and compound two times (400g, 4 min, 4°C). Next, 50 µL aliquots were transferred to polymerase chain reaction (PCR) tubes and heated at different temperatures on a Bio-Rad C1000 Thermal Cycler for 3 min and further incubated for 3 min at 25 °C. Each PCR tube underwent three snap freeze–thaw cycles in liquid nitrogen. Lysates were then transferred to 1.5-ml Eppendorf tubes and centrifuged at 20,000g for 20 min at 4 °C. Lastly, 30 µl of soluble protein was transferred to 10 µl of 4× SDS sample buffer-containing PCR tubes, boiled at 95°C for 5 min, and resolved by SDS-PAGE before immunoblot analysis.

##### Wound Healing Assay
MDA-MB-231 cells were grown to ~60 % confluence in 3 × 6-well plates that were scored on the base with a razor to facilitate image alignment. The cells were treated with **2d** or **2d-neg** (50 µM) in growth medium for 18 h, then scratched with a 200 µl pipette tip. The medium was aspirated to remove debris and replaced with the appropriate solution of compound in growth medium. The cells were imaged (bright field) at 8 hrs and 24 hrs post-scratch. Images were aligned and cropped to a constant size using a custom Python script, and the wound area was quantified using ImageJ with the Wound Healing Tool macro from Montpellier Ressources Imagerie (RRID:SCR_025260; Method: variance, Variance filter radius: 10, Threshold: 200, Radius open: 4, Min. size: 10000). p-Values were calculated using the two-tailed Student's t-test. Protocol was based on ref [2].

##### Meta-Analysis of Identified Targets
Protein targets were classified as 'liganded' if they were annotated as having approved, non-nutraceutical, small molecule ligands in the DrugBank database[3] (retrieved 20230401). Proteins were classified as enzymes if they were assigned Enzyme Commission numbers in the UniProt database[4]

(retrieved 20220516), otherwise their functional classification was based on UniProt keywords adapted from Parker *et al.*[5]. Chemical space plots were generated by applying the UMAP dimension reduction algorithm (0.5.5, arXiv:1802.03426) to 2048-bit molecular fingerprints generated using RDKit .

##### Chemoinformatic Analysis

Chemoinformatic analysis was performed using Python (3.11.3) and the RDKit library (2022.09.5) to calculate chemoinformatic descriptors. Where the descriptors necessitated the calculation of 3D structures (PBF, NPR1, NPR2), a previous procedure was adapted Ogasawara *et al.*[6]. For each structure, 200 conformers were calculated using the RDKit ETKDG algorithm. These conformers were refined using the RDKit MMFF algorithm, then pruned by iteratively selecting the lowest energy conformer, calculating its RMSD from other conformers and excluding those with RMSD < 1.0 until three conformers remained. The mean of the descriptor values for these three conformers was then calculated.

##### Quantification and Statistical Analysis

All data fitting and statistical analysis were performed using GraphPad Prism version 10.1.1 for Windows and Mac, www.graphpad.com. Statistical significance was defined as $p < 0.05$ and determined by two-tailed Student's *t* tests.

##### Data Availability

The mass spectrometry proteomics data have been deposited to the ProteomeXChange Consortium via the PRIDE[7] partner repository with the dataset identifier PXD061964.
