## Supplemental Chemistry Information for "Proteome-Wide Discovery of Degradable Proteins Using Bifunctional Molecules"

#### SYNTHETIC METHODS AND COMPOUND CHARACTERIZATION

##### General Synthetic Information

All commercial reagents acquired from commercial vendors, including Sigma-Aldrich, Fisher Scientific, Combi-Blocks, MedChemExpress, eNovation Chemicals, AstaTech, Matrix Scientific, BroadPharm and AmBeed were used without further purification. Distilled water was used for all water necessities in synthetic procedures (e.g., reagent, solvent, work-up). Anhydrous solvents were purchased from Sigma-Aldrich in Sure/Seal™ formulations. All reactions were monitored by thin-layer chromatography (TLC, Merck silica gel 60 F-254 plates) and directly visualized with UV light (254 nm) or stained either with para-anisaldehyde (2.5% p-anisaldehyde, 1% AcOH, 3.5% $H_2SO_4$ (conc.) in 95% EtOH), iodine, ninhydrin (0.3% ninhydrin (w/v), 97:3 EtOH-AcOH) or KMnO4 (1.5g of KMnO4, 10g $K_2CO_3$, and 1.25mL 10% NaOH in 200mL water), followed by gentle heating. Reaction purification was carried out using Flash chromatography (230 – 400 mesh silica gel), and Biotage® (manually packed columns of 5G, 10G or 25G of silica) or preparative thin layer chromatography (EMD Millipore silica gel coated (250 µm) F254 glass plates or Analtech glass backed plates (1000-2000 µm thickness). $^1$H- and $^{13}$C-NMR spectra were acquired in the indicated solvents and recorded on a Bruker AVANCE NEO 400 MHz instrument with SampleXpress, Bruker AVANCE NEO 500 MHz NMR instrument equipped with 5mm BBFO SP probe, and SampleCase-24 sample changer, and Bruker AVANCE III HD 600 instrument equipped with a 5mm CPDCH CryoProbe. Data was collected at ambient temperature unless otherwise stated using standard pulse methods as supplied by Bruker software. Coupling constants are quoted to the nearest 0.1 Hz and multiplicities are given by the following abbreviations and combinations: m (multiplet), s (singlet), d (doublet), dd (doublet of doublets), ddd (doublet of doublet of doublets), t (triplet), td (triplet of doublets), tt (triplet of triplets), q (quartet), br (broad). All NMR data was processed in MestReNova v14.0.0. Chemical shifts for proton and carbon resonances are reported in parts per million (ppm) on the δ scale relative to the residual protons of the deuterated solvent of relevance. Mass spectrometry data were collected on a Thermo Scientific ISQ single-quadrupole instrument (ESI; low resolution), Agilent 6125 and 6135 single-quadrupole instruments (ESI; low resolution), and Agilent 6230 single-quadrupole TOF (ESI-TOF; high resolution HRMS).

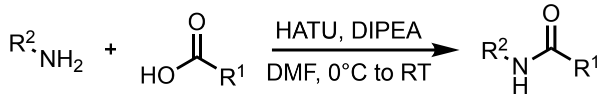


**General Procedure 1A: Synthesis of amides using HATU as coupling reagent (in vial/flask)**

To a solution of amine (1.1 eq.), carboxylic acid (1.0 eq.) and HATU (1.5 eq.) in dry DMF at 0°C was added DIPEA (3.0 eq.). The reaction mixture was stirred for 2 - 18 h and then quenched with DI $H_2O$ and diluted with Ethyl acetate (EtOAc). The aqueous layer was extracted 3x with EtOAc, and the combined organic layers were washed 2x with $H_2O$, 2x with a saturated aqueous solution of $NH_4Cl$ and 2x with a saturated aqueous solution of NaCl before being dried over anhydrous $Na_2SO_4$ and filtered. Volatiles were removed by rotary evaporation and the crude product was purified by automated column chromatography, pTLC, or organic solvent washes (i.e., the organic solvent was added into the vial and mixed with the solid decanting).

**General Procedure 1B: Synthesis of amides using HATU as coupling reagent (high-throughput)**

In a 96 well plate, stock solutions of amine (HCl or TFA salt, 0.42 M, 1.05 eq.), carboxylic acid (0.40 M, 1.0 eq.) and HATU (0.42 M, 1.05 eq.) were combined. A stock solution of DIPEA (1.4 M, 3.5 eq.) in dry DMF was then added and the reaction mixture was shaken at RT for 6 h. The reaction was diluted with 2x MeCN before purification by preparative HPLC (MeCN/$H_2O$ gradient, 0.1% FA), followed by pTLC if necessary.

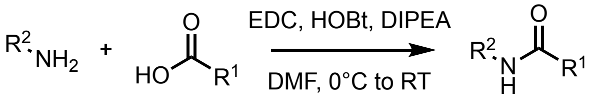


**General Procedure 2: Synthesis of amides using EDC as coupling reagent**

To a solution of amine (1.1 eq.), carboxylic acid (1.0 eq.), EDC (1.5 eq.) and HOBt (1.5 eq.) in dry $CH_2Cl_2$ (DCM) at 0°C was added DIPEA (3.0 eq.). The reaction mixture was stirred overnight, then quenched with deionized $H_2O$. The aqueous layer was extracted 3x with $CH_2Cl_2$, and the combined organic layers were washed 2x with $H_2O$, 2x with a saturated aqueous solution of $NH_4Cl$ and 2x with a saturated aqueous solution of NaCl before being dried over anhydrous $Na_2SO_4$ and filtered. Volatiles were removed by rotary evaporation and the crude product was purified by automated column chromatography, pTLC, or organic solvent washes (i.e., the organic solvent was added into the vial and mixed with the solid decanting).

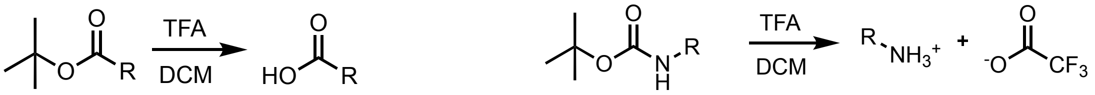


**General Procedure 3: *Tert*-butyl ester deprotection of carboxylic acids and Boc deprotection of amines using TFA**

To a solution of t-Butyl protected acid (left, 1.0 eq.) or Boc protected amine (right, 1.0 eq.) in dry $CH_2Cl_2$ (DCM) was added a solution of TFA (20-50%). The reaction mixture was stirred at RT for 0.5 - 12 h. The crude product was concentrated through a nitrogen flow, diluted in $CH_2Cl_2$ and evaporated again. No purification was needed.

**General Procedure 4: Methylated negative control**

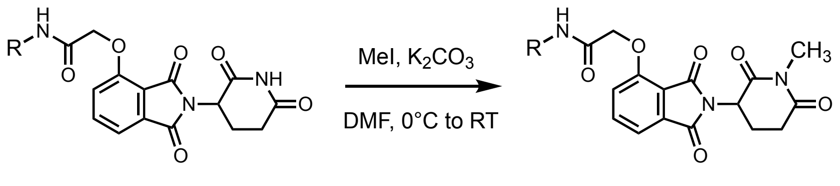


To a solution of selected bifunctional molecule (1.0 eq) in DMF (1 mL) at 0°C, was added potassium carbonate ($K_2CO_3$, 1.2 eq) and methyl iodide (MeI, 1.2 eq). The reaction mixture was stirred at RT for 0.5 – 1 h, then quenched with cold with $H_2O$ and extracted with EtoAC (2 × 50 mL). The combined organic extracts were dried over anhydrous $Na_2SO_4$, filtered, and concentrated under reduced pressure. The crude product was purified by pTLC or Biotage® using a 0–6% methanol gradient in $CH_2Cl_2$ (DCM) to afford the corresponding methylated negative control.

### Compound Characterization

#### Thalidomide-linker intermediates

Thalidomide-O-C2-acid, thalidomide-O-C2-amine, thalidomide-O-acetamido-C5-acid, thalidomide-O-acetamido-C5-amine, thalidomide-O-C5-acid, thalidomide-O-acetamido-PEG2-C2-acid, thalidomide-O-acetamido-PEG2-C2-amine, thalidomide-O-PEG2-acid, thalidomide-O-acetamido-PEG3-C2-acid, thalidomide-O-acetamido-PEG3-C2-amine and thalidomide-O-PEG3-C2-acid starting materials were purchased from BroadPharm and Tenova Pharmaceuticals Inc. and used as received without purification.

#### Headgroup intermediates

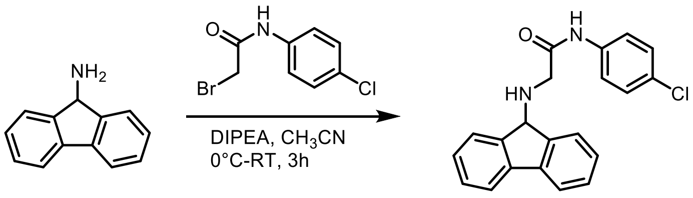


**F2**

**Fragment F2: 2-((9H-fluoren-9-yl)amino)-N-(4-chlorophenyl)acetamide (F2)**

To a solution of 9H-Fluoren-9-amine hydrochloride 88 mg (40.4 µmol, 1 eq) and 2-bromo-N-(4-chlorophenyl)acetamide 100 mg (40.4 µmol, 1 eq) in acetonitrile (5 mL), DIPEA (120.4 µmol, 3 eq) were added at 0°C. The reaction mixture was stirred at RT for 3 h. The reaction mixture was then diluted with cold water and extracted with ethyl acetate. The organic layer was dried over anhydrous $Na_2SO_4$, filtered, and concentrated under reduced pressure. The crude product was purified by Biotage® using a 0–60% ethyl acetate in hexane gradient to afford the desired product (**F2**) as a white powder (125 mg, 35.3 µmol, 65%). $^1H$ NMR (600 MHz, $CDCl_3$) δ 9.42 (s, 1H), 7.74 (d, J = 7.6 Hz, 2H), 7.65 (d, J = 7.4 Hz, 2H), 7.56 – 7.50 (m, 2H), 7.46 – 7.42 (m, 2H), 7.36 – 7.34 (m, 2H), 7.32 – 7.29 (m, 2H), 5.08 (s, 1H), 3.24 (s, 2H). $^{13}C$ NMR (151 MHz, CDCl3) δ 169.60, 143.83, 140.96, 136.07, 129.17, 129.11, 129.07, 128.88, 127.71, 124.65, 120.67, 120.29, 63.17, 48.50. LC-MS (ESI+) calc'd for $C_{21}H_{17}ClN_2O10$: 348.10, found $[M+H]^+$ 349.10.

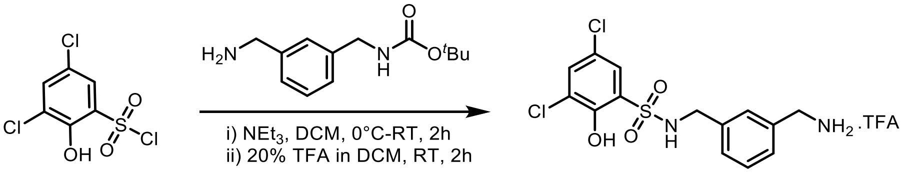


**F4**

**Fragment F4**: **N-(3-(aminomethyl)benzyl)-3,5-dichloro-2-hydroxybenzenesulfonamide (F4)**

To a solution of tert-Butyl 3-(aminomethyl)benzylcarbamate 300 mg (1.27 µmol, 1 eq) and 3,5-Dichloro-2-hydroxybenzenesulfonyl chloride 332 mg (1.27 µmol, 1 eq) in dichloromethane (5 mL), triethylamine 257 mg (2.54 µmol, 2 eq) were added at 0°C. The reaction mixture was stirred at RT for 2 h. The reaction mixture was then diluted with cold water and extracted with dichloromethane. The organic layer was dried over anhydrous $Na_2SO_4$, filtered, and concentrated under reduced pressure. The crude product was purified by Biotage® using a 0–80% ethyl acetate in hexane gradient to afford the Boc intermediate, which was subjected to deprotection as described in general procedure 3 to obtain N-(3-(aminomethyl)benzyl)-3,5-dichloro-2-hydroxybenzenesulfonamide (**F4**) as TFA salt which was used without further purification. (470 mg, 72 %) $^1H$ NMR (600 MHz, DMSO-d6) δ 10.97 (s, 1H), 8.24 – 8.19 (m, 2H), 7.82 – 7.72 (m, 1H), 7.53 (d, *J* = 2.6 Hz, 1H), 7.33 – 7.28 (m, 3H), 4.13 (s, 2H), 3.97 (s, 2H). LC-MS (ESI+) calc'd for $C_{14}H_{14}Cl_2N_2O_3S$: 360.01, found $[M+H]^+$ 361.01.

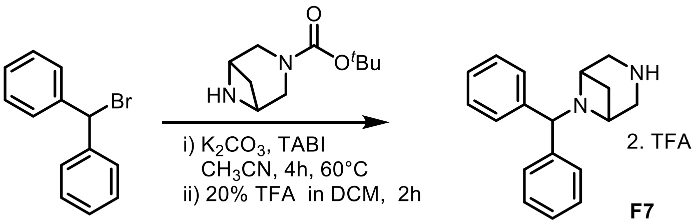


**F7**

**Fragment F7**: **2-benzhydryl-2,5-diazabicyclo[2.2.1]heptane (F7)**

To a stirred solution of bromodiphenylmethane 100 mg (0.40 µmol, 1 eq) in acetonitrile (5 mL) was added tert-Butyl 3,6-diazabicyclo[3.1.1]heptane-3-carboxylate 88 mg (0.44 µmol, 1.2 eq), $K_2CO_3$ 167 mg (1.21 µmol, 3 eq) and TABI 15 mg (0.04 µmol, 0.1 eq). The mixture was stirred for 4h at 60 °C, then cooled to room temperature, diluted with water, and extracted in ethyl acetate (50 mL X2). The combined extract was dried over $Na_2SO_4$, filtered, and concentrated. The crude product was purified by Biotage® using a 0–50% Ethyl acetate in Hexane gradient to afford tert-butyl-5-benzhydryl-2,5-diazabicyclo[2.2.1]heptane-2-carboxylate intermediate was subjected to deprotection as described in general procedure 3 to obtain 2-benzhydryl-2,5-diazabicyclo[2.2.1]heptane (**F7**) as TFA salt which was used without further purification. (150 mg, 67 %) $^1$H NMR (600 MHz, $CD_3OD$) δ 7.65 (d, $J$ = 7.7 Hz, 4H), 7.44 (d, $J$ = 7.6 Hz, 4H), 7.38 (d, $J$ = 7.1 Hz, 2H), 5.88 (s, 1H), 4.18 – 4.12 (m, 2H), 4.10 – 4.02 (m, 2H), 3.74 (s, 2H), 3.61 – 3.38 (m, 1H), 2.18 (s, 1H). LC-MS (ESI+) calc'd for $C_{18}H_{20}N_2$: 264.1, found $[M+H]^+$ 265.1.

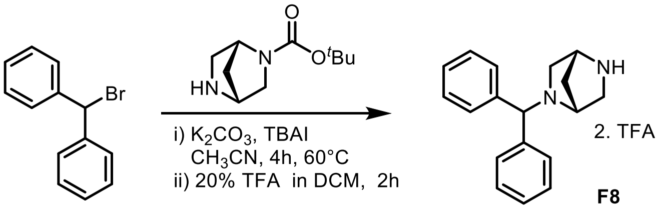


**F8**

**Fragment F8**: **(1R,4R)-2-benzhydryl-2,5-diazabicyclo[2.2.1]heptane (F8)**

To a stirred solution of bromodiphenylmethane 100 mg (0.40 µmol, 1 eq) in acetonitrile (5 mL) was added (*1R*, *4R*)-diazabicyclo[2,2,1]-heptane 88 mg (0.44 µmol, 1.2 eq), $K_2CO_3$ 167 mg (1.21 µmol, 3 eq) and TABI 15 mg (0.04 µmol, 0.1 eq). The mixture was stirred for 4h at 60 °C, then cooled to room temperature, diluted with water, and extracted in ethyl acetate (50 mL x 2). The combined extract was dried over $Na_2SO_4$, filtered, and concentrated. The crude product was purified by Biotage® using a 0–50% Ethyl acetate in Hexane gradient to afford the tert-butyl (*1R,4R*)-5-benzhydryl-2,5-diazabicyclo[2.2.1]heptane-2-carboxylate intermediate which was subjected for deprotection as a general procedure 3 to obtain (1R,4R)-2-benzhydryl-2,5-diazabicyclo[2.2.1]heptane **(F8)** as TFA salt which was used without further purification. (112 mg, 56 %) $^1$H NMR (600 MHz, $CD_3OD$) δ 7.61 – 7.51 (m, 4H), 7.38 – 7.32 (m, 4H), 7.32 – 7.23 (m, 2H), 4.95 (s, 1H), 4.32 (s, 1H), 3.89 – 3.59 (m, 2H), 3.22 – 3.18 (m, 1H), 3.02 – 2.98 (m, 2H), 2.45 – 2.43 (m, 1H), 1.91 – 1.89 (m, 1H). LC-MS (ESI+) calc'd for $C_{18}H_{20}N_2$: 264.1, found $[M+H]^+$ 265.1.

#### Final Compounds

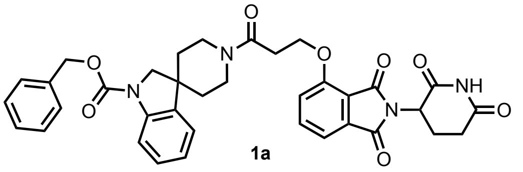

**1a**

**Benzyl 1'-(3-((2-(2,6-dioxopiperidin-3-yl)-1,3-dioxoisoindolin-4-yl)oxy)propanoyl)spiro[indoline-3,4'-piperidine]-1-carboxylate (1a)**

**General Procedure 1A.** Reaction scale: Thalidomide-O-C2-acid 19 mg (1 eq, 55.7 µmol) and Benzyl spiro[indoline-3,4'-piperidine]-1-carboxylate 20 mg (1 eq, 55.7 µmol). Purified by pTLC (6% MeOH/DCM) to afford **1a** as an off-white solid (8.0 mg, 43 %). $^{1}$H NMR (600 MHz, $CDCl_3$) δ 8.65 (s, 1H), 7.93 (s, 1H), 7.74 – 7.64 (m, 2H), 7.51 – 7.44 (m, 6H), 7.46 – 7.40 (m, 3H), 7.36 – 7.30 (m, 2H), 7.24 – 7.16 (m, 2H), 7.16 – 7.12 (m, 2H), 7.06 – 6.98 (m, 2H), 5.42 – 5.21 (m, 4H), 4.96 (s, 2H), 4.70 – 4.64 (m, 2H), 4.62 – 4.54 (m, 4H), 4.06 – 3.96 (m, 5H), 3.48 (s, 1H), 3.25 (s, 1H), 3.08 – 2.98 (s, 2H), 2.83 – 2.68 (m, 6H), 2.10 (s, 2H), 1.97 (s, 1H), 1.93 – 1.77 (m, 6H), 1.73 – 1.71 (m, 4H). $^{13}$C NMR (151 MHz, $CDCl_3$) δ 167.01, 165.79, 156.23, 136.72, 133.73, 128.68, 128.39, 128.31, 123.05, 119.16, 117.11, 116.15, 49.16, 39.00, 37.20, 36.33, 31.40. (*Note: rotameric mixture observed*) HR-MS (ESI+) calc'd for $C_{36}H_{34}N_4O_8$: 650.2377, found $[M+H]^+$ 651.2448.

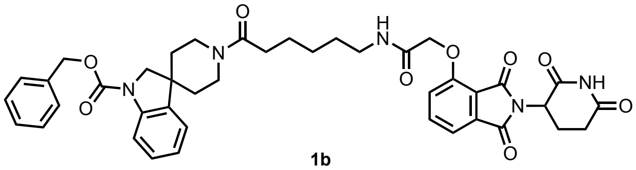

**1b**

**benzyl 1'-(6-(2-((2-(2,6-dioxopiperidin-3-yl)-1,3-dioxoisoindolin-4 yl)oxy) acetamido) hexanoyl) spiro[indoline-3,4'-piperidine]-1-carboxylate (1b)**

**General Procedure 1A**. Reaction scale: Thalidomide-O-acetamido-C5-acid 19 mg (1eq, 41.7 µmol) and Benzyl spiro[indoline-3,4'-piperidine]-1-carboxylate 15 mg (1 eq, 41.7 µmol). Purified by pTLC (6% MeOH/DCM) to afford **1b** as an off-white solid (7.0 mg, 41 %). $^{1}$H NMR (600 MHz, CDCl3) δ 9.69 (s, 1H), 7.92 (s, 1H), 7.81 – 7.72 (m, 2H), 7.70 – 7.62 (m, 2H), 7.61 – 7.52 (m, 2H), 7.51 – 7.34 (m, 10H), 7.27 – 7.19 (m, 3H), 7.14 – 7.06 (m, 2H), 7.02 (s, 2H), 5.36 – 5.28 (m, 4H), 5.07 – 4.92 (m, 2H), 4.77 – 4.56 (m, 6H), 3.98 – 3.92 (m, 4H), 3.88 (d, J = 13.9 Hz, 2H), 3.44 (dd, J = 13.4, 6.7 Hz, 2H), 3.40 – 3.28 (m, 2H), 3.24 – 3.13 (m, 2H), 2.94 – 2.85 (m, 2H), 2.84 – 2.76 (m, 4H), 2.74 – 2.68 (m, 2H), 2.46 – 2.39 (m, 4H), 2.23 – 2.12 (m, 2H), 1.88 – 1.78 (m, 4H), 1.78 – 1.60 (m, 12H), 1.48 – 1.39 (m, 6H). $^{13}$C NMR (151 MHz, $CDCl_3$) δ 171.49, 168.56, 166.95, 166.62, 166.31, 154.79, 137.10, 133.54, 128.67, 128.37, 128.29, 123.12, 117.59, 68.63, 55.44, 49.34, 39.20, 39.04, 38.90, 33.22, 31.41, 28.88, 26.76, 25.34, 22.82. (*Note: rotameric mixture observed*) HR-MS (ESI+) calc'd for $C_{41}H_{43}N_5O_9$: 749.3177, found $[M+H]^+$ 750.3142.

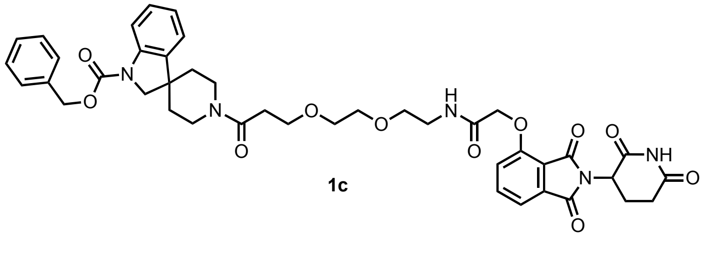

**1c**

**Benzyl 1-(3-(2-(2-(2-((2-(2,6-dioxopiperidin-3-yl)-1,3-dioxoisoindolin-4-yl)oxy)acetamido)ethoxy) ethoxy)propanoyl)spiro[indoline-3,4'-piperidine]-1'-carboxylate (1c)**

**General Procedure 1A**. Reaction scale: Thalidomide-O-acetamido-PEG2-C2-acid 20 mg (1 eq, 31.7 µmol) and 1'Benzyl spiro[indoline-3,4'-piperidine]-1-carboxylate 14 mg (1.1 eq, 34.9 µmol). Purified by pTLC (7% MeOH/DCM) to afford **1c** as an off-white solid (12.0 mg, 42%). $^1$H NMR (600 MHz, $CDCl_3$) δ 9.64 (s, 2H), 7.98 – 7.96 (m, 1H), 7.84 – 7.72 (m, 3H), 7.68 – 7.55 (m, 2H), 7.50 – 7.44 (m, 6H), 7.37 – 7.21 (m, 3H), 7.19 – 7.13 (m, 2H), 7.11 – 6.99 (m, 2H), 6.70 – 6.68 (m, 1H), 5.42 – 5.34 (m, 4H), 5.17 – 4.92 (m, 2H), 4.76 – 4.72 (m, 5H), 4.09 – 3.83 (m, 8H), 3.83 – 3.67 (m, 8H), 3.66 – 3.64 (m, 2H), 3.26 – 3.24 (m, 3H), 3.03 – 2.59 (m, 10H), 2.23 – 2.19 (m, 2H), 2.10 – 2.06 (m, 3H), 1.90 – 1.86 (m, 3H), 1.78 – 1.74 (m, 3H), 1.58 – 1.38 (m, 8H). $^{13}$C NMR (151 MHz, $CDCl_3$) δ 171.60, 169.68, 168.50, 166.87, 166.62, 165.72, 154.18, 153.45, 152.59, 141.42, 140.21, 138.27, 137.47, 136.88, 136.03, 133.45, 128.51, 128.32, 128.21, 128.11, 122.95, 122.42, 119.15, 117.70, 117.03, 114.79, 70.10, 70.07, 69.32, 67.84, 67.54, 67.14, 67.03, 56.86, 55.17, 49.15, 43.19, 42.78, 42.48, 41.82, 38.83, 37.00, 36.11, 33.30, 31.31, 29.53, 22.54, 18.38, 16.97, 12.45. (*Note: rotameric mixture observed*) HR-MS (ESI+) calc'd for $C_{42}H_{45}N_5O_{11}$: 795.3116, found $[M+H]^+$ 796.3183.

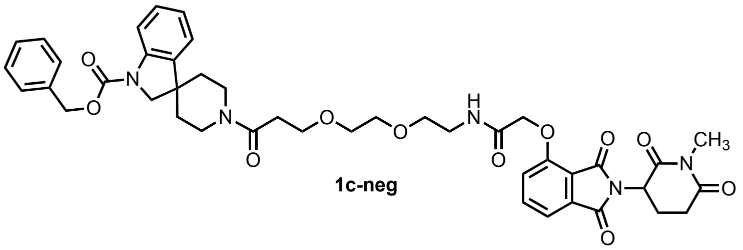

**1c-neg**

**Benzyl 1-(3-(2-(2-(2-((2-(1-methyl-2,6-dioxopiperidin-3-yl)-1,3-dioxoisoindolin-4-yl)oxy)acetamido) ethoxy)ethoxy)propanoyl)spiro[indoline-3,4'-piperidine]-1'-carboxylate (1c-neg)**

**General Procedure 4**. Reaction scale: 10 mg (1 eq, 13.3 µmol) **1c** and methyl iodide 6 mg (2.0 eq, 26.6 µmol). Purified by pTLC (6% MeOH/DCM) to afford **1c-neg** as a sticky solid (3 mg, 35%). $^1$H NMR (600 MHz, $CDCl_3$) δ 7.90 (s, 1H), 7.72 (dd, *J* = 8.3, 7.4 Hz, 1H), 7.63 – 7.58 (m, 1H), 7.53 (d, *J* = 7.3 Hz, 1H), 7.43 (d, *J* = 7.4 Hz, 2H), 7.43 – 7.38 (m, 2H), 7.37 – 7.31 (m, 1H), 7.22 (s, 1H), 7.18 (d, *J* = 8.4 Hz, 1H), 7.08 (d, *J* = 7.4 Hz, 1H), 7.00 (d, *J* = 7.7 Hz, 1H), 5.46 – 5.14 (m, 3H), 5.05 – 4.95 (m, 1H), 4.64 (s, 3H), 3.93 – 3.87 (m, 3H), 3.85 – 3.75 (m, 2H), 3.67 – 3.61 (m, 5H), 3.60 – 3.56 (s, 2H), 3.20 (s, 3H), 3.17 – 3.14 (m, 1H), 3.03 – 2.94 (m, 1H), 2.86 – 2.76 (m, 2H), 2.69 – 2.63 (m, 3H), 2.14 – 2.08 (m, 1H), 1.82 – 1.78 (m, 2H), 1.75 – 1.64 (m, 4H). $^{13}$C NMR (151 MHz, $CDCl_3$) δ 171.22, 171.20, 169.37, 168.82, 168.79, 167.02, 166.84, 166.82, 165.97, 165.95, 154.47, 154.45, 136.97, 136.95, 133.69, 133.67, 128.68, 128.66, 128.51, 128.37, 128.29, 128.27, 123.09, 122.51, 119.44, 118.04, 117.24, 115.02, 77.35, 77.14, 76.93, 70.39, 70.38, 70.32, 70.30, 69.57, 69.55, 67.97, 67.42, 67.20, 57.02, 55.11, 50.02, 50.01, 43.13, 42.82, 38.99, 38.98, 38.78, 37.21, 36.31, 33.56, 31.86, 31.84, 29.69, 27.26, 27.25, 21.94, 21.93, 18.59, 17.25, 12.41 (*Note: rotameric mixture observed*) HR-MS (ESI+) calc'd for $C_{43}H_{47}N_5O_{11}$: 809.3272, found $[M+H]^+$ 810.3352.

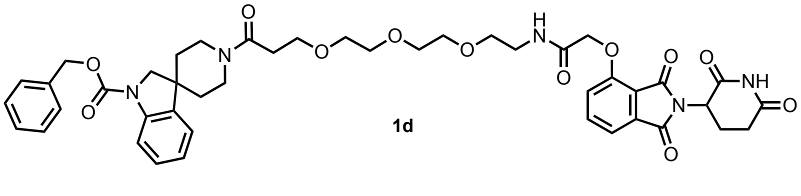

**1d**

**Benzyl 1'-(1-((2-(2,6-dioxopiperidin-3-yl)-1,3-dioxoisoindolin-4-yl)oxy)-2-oxo-6,9,12-trioxa-3-azapentadecan-15-oyl)spiro[indoline-3,4'-piperidine]-1-carboxylate (1d)**

**General Procedure 1A**. Reaction scale: Thalidomide-O-acetamido-PEG3-C2-acid 20 mg (1 eq, 31.7 µmol) and 1'Benzyl spiro[indoline-3,4'-piperidine]-1-carboxylate 13 mg (1.1 eq, 34.9 µmol). Purified by pTLC (7% MeOH/DCM) to afford **1d** as an off-white solid (5.0 mg, 38%). $^1$H NMR (600 MHz, $CDCl_3$) δ 9.29 (s, 2H),

7.92 – 7.89 (m, 1H), 7.77 – 7.69 (m, 2H), 7.70 – 7.64 (m, 2H), 7.57 – 7.52 (m, 2H), 7.47 – 7.42 (m, 4H), 7.42 – 7.38 (m, 5H), 7.38 – 7.34 (m, 3H), 7.23 – 7.17 (m, 5H), 7.13 – 7.06 (m, 2H), 7.04 – 6.98 (m, 3H), 5.36 – 5.28 (m, 5H), 5.02 – 4.98 (m, 2H), 4.69 – 4.64 (m, 7H), 3.96 – 3.86 (m, 7H), 3.89 – 3.82 (m, 5H), 3.68 – 3.64 (m, 16H), 3.61 – 3.54 (m, 5H), 3.19 – 3.15 (m, 2H), 2.93 – 2.85 (m, 2H), 2.85 – 2.65 (m, 12H), 2.16 – 2.09 (m, 3H), 1.85 – 1.79 (m, 5H), 1.72 – 1.67 (m, 5H). $^{13}$C NMR (151 MHz, $CDCl_3$) δ 171.41, 171.35, 169.51, 168.39, 168.35, 166.90, 166.71, 165.83, 154.47, 136.96, 133.66, 128.66, 128.48, 128.36, 128.27, 123.08, 122.54, 119.39, 118.01, 117.24, 115.02, 77.33, 77.11, 76.90, 70.42, 70.37, 70.30, 69.46, 67.92, 67.30, 67.19, 57.03, 49.34, 42.88, 42.67, 39.09, 38.87, 37.25, 36.33, 33.50, 31.48, 29.69, 29.36, 22.69, 22.66, 14.14. (*Note: rotameric mixture observed*) HR-MS (ESI+) calc'd for $C_{44}H_{49}N_5O_{12}$: 839.3378, found $[M+H]^+$ 840.3449.

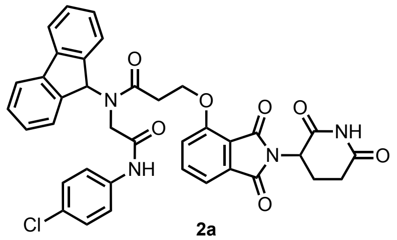

**2a**

**N-(2-((4-chlorophenyl)amino)-2-oxoethyl)-3-((2-(2,6-dioxopiperidin-3-yl)-1,3 dioxoisoindolin-4-yl)oxy)-N-(9H-fluoren-9-yl)propenamide (2a)**

**General Procedure 1A**. Reaction scale: Thalidomide-O-C2-acid 20 mg (1 eq, 57.3 μmol) and **F2** 20 mg (57.3 μmol, 1 eq). Purified by pTLC (6% MeOH/DCM) to afford **2a** as a sticky solid (8.0 mg, 42 %). $^1$H NMR (600 MHz, $CDCl_3$) δ 9.62 (s, 1H), 8.14 – 8.12 (m, 1H), 7.84 – 7.68 (m, 4H), 7.63 (s, 2H), 7.56 – 7.47 (m, 2H), 7.42 – 7.39 (m, 2H), 7.21 – 7.18 (m, 3H), 6.73 – 6.70 (m, 1H), 6.40 – 6.36 (m, 1H), 6.13 – 6.11 (m, 1H), 5.30 (s, 2H), 5.17 (s, 1H), 4.85 – 4.76 (m, 1H), 3.42 (t, J = 5.7 Hz, 2H), 3.16 – 2.98 (m, 4H), 2.93 – 2.83 (m, 1H), 2.86 – 2.76 (m, 1H), 2.75 – 2.67 (m, 1H), 2.66 – 2.68 (m, 2H). $^{13}$C NMR (151 MHz, $CDCl_3$) δ 170.76, 170.64, 170.05, 167.70, 167.64, 166.36, 166.19, 164.92, 164.75, 163.33, 146.84, 146.40, 141.12, 136.47, 136.29, 136.20, 134.33, 133.26, 133.16, 128.93, 128.86, 128.69, 127.57, 126.70, 125.41, 122.46, 121.71, 121.50, 120.82, 120.40, 66.66, 56.28, 53.46, 49.40, 49.35, 31.34, 31.29, 22.52, 22.37. (*Note: rotameric mixture observed*) HR-MS (ESI+) calc'd for $C_{37}H_{29}ClN_4O_7$: 676.1725, found $[M+H]^+$ 677.1798.

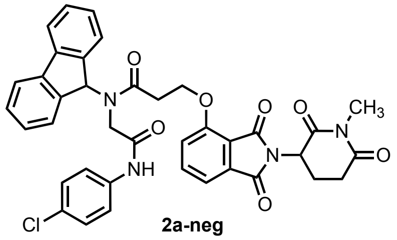

**2a-neg**

**N-(2-((4-chlorophenyl)amino)-2-oxoethyl)-N-(9H-fluoren-9-yl)-3-((2-(1-methyl-2,6-dioxopiperidin-3-yl)-1,3-dioxoisoindolin-4-yl)oxy)propenamide (2a-neg)**

**General Procedure 4**. Reaction scale: bifunctional molecule **2a** 9.6 mg (14.1 μmol, 1 eq) and methyl iodide 2 mg (28.2 μmol, 1 eq). Purified by pTLC (6% MeOH/DCM) to afford **2a-neg** as a sticky solid (3.0 mg, 32%). $^1$H NMR (600 MHz, $CDCl_3$) δ 9.61 (s, 1H), 7.84 – 7.68 (m, 6H), 7.62 (s, 3H), 7.54 – 7.52 (m, 3H), 7.45 – 7.38 (m, 4H), 7.21 – 7.17 (m, 4H), 5.17 (s, 1H), 4.82 – 4.78 (m, 1H), 3.43 (s, 2H), 3.16 – 3.13 (m, 4H), 3.10 – 3.02 (m, 4H), 2.96 – 2.88 (m, 2H), 2.65 (s, 3H), 1.97 (s, 1H). $^{13}$C NMR (151 MHz, CDCl3) δ 170.76, 170.64, 170.05, 167.70, 167.64, 166.36, 166.19, 164.92, 164.75, 163.33, 146.84, 146.40, 141.12, 136.47, 136.29, 136.20, 134.33, 133.26, 133.16, 128.93, 128.86, 128.69, 127.57, 126.70, 125.41, 122.46, 121.71, 121.50, 120.82, 120.40, 66.66, 56.28, 53.46, 49.40, 49.35, 31.34, 31.29, 22.52, 22.37. (*Note: rotameric mixture observed*) HR-MS (ESI+) calc'd for $C_{38}H_{31}ClN_4O_7$: 690.1881, found $[M+H]^+$ 691.1957.

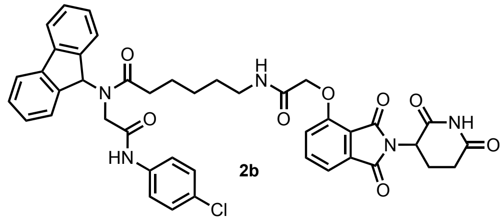

**N-(2-((4-chlorophenyl)amino)-2-oxoethyl)-6-(2-((2-(2,6-dioxopiperidin-3-yl)-1,3-dioxoisoindolin-4-yl)oxy)acetamido)-N-(9H-fluoren-9-yl)hexanamide (2b)**

**General Procedure 1A**. Reaction scale: Thalidomide-O-C5-acid 20 mg (1 eq, 44.9 µmol) and **F2** 16 mg (1.1 eq, 44.9 µmol). Purified by pTLC (6% MeOH/DCM) to afford **2b** as a sticky solid (7.0 mg, 45 %). $^1$H NMR (600 MHz, $CDCl_3$) δ 8.44 (s, 1H), 7.74 – 7.68 (m, 6H), 7.55 (s, 3H), 7.42 – 7.38 (m, 6H), 7.19 (s, 6H), 7.06 (s, 2H), 6.98 (s, 1H), 5.95 (s, 1H), 4.96 – 4.92 (m, 2H), 4.64 – 4.58 (m, 4H), 3.56 – 3.24 (m, 7H), 3.00 – 2.82 (m, 4H), 2.74 (s, 3H), 2.65 (s, 1H), 2.50 (s, 1H), 2.14 (s, 2H), 1.95 (s, 2H), 1.84 (s, 2H), 1.72 (s, 3H), 1.56 (s, 4H), 1.46 – 1.43 (m, 3H), 1.26 – 1.23 (m, 7H). $^{13}$C NMR (151 MHz, DMSO) δ 172.85, 170.14, 169.93, 166.75, 165.54, 155.07, 145.02, 140.37, 137.54, 136.96, 133.06, 128.57, 128.15, 127.33, 126.78, 125.13, 120.73, 120.39, 120.04, 116.81, 116.09, 67.63, 62.33, 48.82, 47.14, 38.74, 38.04, 30.97, 28.49, 26.65, 23.08, 22.01. (*Note: rotameric mixture observed*) HR-MS (ESI+) calc'd for $C_{42}H_{38}ClN_5O_8$: 775.2409, found $[M+H]^+$ 776.2471

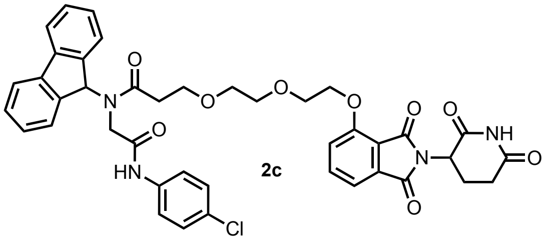

**N-(2-((4-chlorophenyl)amino)-2-oxoethyl)-3-(2-(2-((2-(2,6-dioxopiperidin-3-yl)-1,3-dioxoisoindolin-4-yl)oxy)ethoxy)ethoxy)-N-(9H-fluoren-9-yl)propenamide (2c)**

**General Procedure 1A**. Reaction scale: Thalidomide-O-PEG2-C2-acid 15 mg (34.5 µmol, 1 eq) and **F2** 12 mg (34.5 µmol, 1 eq). Purified by pTLC (7% MeOH/DCM) to afford **2c** as a sticky solid (3.0 mg, 28%). $^1$H NMR (600 MHz, $CDCl_3$) δ 8.55 (s, 1H), 8.20 – 8.17 (m, 2H), 7.91 – 7.89 (m, 1H), 7.74 – 7.68 (m, 3H), 7.66 – 7.58 (m, 5H), 7.52 – 7.46 (m, 6H), 7.44 – 7.40 (m, 5H), 7.36 – 7.30 (m, 3H), 7.22 – 7.18 (m, 7H), 7.14 – 7.10 (m, 5H), 6.96 (s, 1H), 6.09 (s, 1H), 4.89 – 8.84 (m, 1H), 4.78 (s, 1H), 4.23 – 4.17 (m, 5H), 4.07 – 4.03 (m, 4H), 4.00 – 3.96 (m, 4H), 3.91 (s, 3H), 3.89 – 3.84 (m, 5H), 3.79 – 3.74 (m, 6H), 3.46 – 3.44 (m, 4H), 3.40 – 3.34 (m, 2H), 3.21 – 3.17 (m, 3H), 2.85 – 2.82 (m, 3H), 2.79 – 2.70 (m, 2H), 2.70 – 2.61 (m, 2H), 2.06 – 2.03 (m, 4H). $^{13}$C NMR (151 MHz, $CDCl_3$) δ 174.14, 170.99, 170.80, 168.51, 168.20, 167.08, 166.84, 165.58, 156.20, 141.67, 141.20, 140.77, 140.68, 136.57, 136.44, 133.67, 129.55, 129.36, 128.94, 128.75, 128.70, 127.79, 127.65, 125.81, 125.01, 124.97, 121.32, 121.13, 120.58, 120.10, 119.12, 119.03, 117.16, 116.88, 116.12, 70.97, 70.88, 70.62, 69.17, 68.93, 67.91, 67.66, 63.25, 49.04, 48.18, 34.43, 34.29, 31.29, 31.19, 29.68, 22.55. (*Note: rotameric mixture observed*) HR-MS (ESI+) calc'd for $C_{41}H_{37}ClN_4O_9$: 764.2249, found $[M+H]^+$ 765.2313.

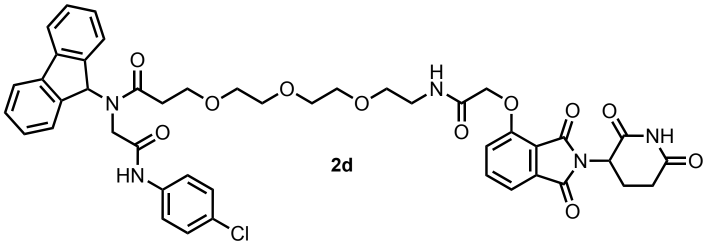

2d

**N-(2-((4-chlorophenyl)amino)-2-oxoethyl)-3-(2-(2-(2-(2-((2-(2,6-dioxopiperidin-3-yl)-1,3-dioxoisoindolin-4-yl)oxy)acetamido)ethoxy)ethoxy)ethoxy)-N-(9H-fluoren-9-yl)propenamide (2d)**

**General Procedure 1A**. Reaction scale: Thalidomide-O-acetamido-PEG3-C2-acid 20 mg (29.7 μmol, 1 eq) and **F2** 10 mg (29.7 μmol, 1 eq). Purified by pTLC (7% MeOH/DCM) to afford **2d** as a sticky solid (8mg, 40 % ). $^1$H NMR (600 MHz, $CDCl_3$) δ 9.30 (s, 1H), 9.16 (s, 1H), 8.52 (s, 1H), 8.19 (s, 1H), 7.73 – 7.70 (m, 3H), 7.69 – 7.67 (m, 1H), 7.72 – 7.64 (m, 1H), 7.67 – 7.62 (m, 3H), 7.61 (s, 1H), 7.52 – 7.48 (m, 6H), 7.44 – 7.46 (m, 2H), 7.36 – 7.30 (m, 2H), 7.28 – 7.23 (m, 6H), 7.22 – 7.19 (m, 1H), 7.18 – 7.09 (m, 6H), 6.88 (s, 1H), 6.03 (s, 1H), 4.96 – 4.94 (m, 1H), 4.93 – 4.89 (m, 1H), 4.65 – 4.50 (m, 4H), 4.07 – 3.90 (m, 4H), 3.73 – 3.68 (m, 2H), 3.64 – 3.58 (m, 7H), 3.56 – 3.52 (m, 6H), 3.51 – 3.46 (m, 2H), 3.44 – 3.40 (m, 2H), 3.38 (s, 2H), 3.22 – 3.18 (m, 2H), 3.15 – 3.12 (m, 1H), 2.80 – 2.74 (m, 4H), 2.12 – 2.08 (m, 4H), 1.46 – 1.39 (m, 4H).$^{13}$C NMR (151 MHz, $CDCl_3$) δ 174.42, 173.86, 171.57, 168.91, 167.11, 166.99, 166.89, 166.68, 166.60, 165.89, 165.86, 154.37, 154.27, 141.72, 141.60, 141.31, 141.26, 140.74, 137.03, 137.01, 136.55, 135.98, 133.52, 133.48, 129.54, 129.19, 128.98, 128.70, 128.68, 128.65, 127.85, 127.83, 127.70, 127.67, 125.85, 125.17, 121.51, 121.19, 120.52, 120.09, 120.07, 119.45, 119.34, 117.86, 117.79, 117.25, 117.20, 77.32, 77.11, 76.90, 70.64, 70.33, 70.27, 70.24, 70.22, 70.18, 70.17, 69.50, 67.81, 67.68, 67.49, 67.39, 63.20, 59.50, 55.24, 49.27, 47.95, 43.24, 39.08, 39.00, 34.35, 33.99, 31.37, 22.62, 22.59, 18.55, 17.18, 12.46. (*Note: rotameric mixture observed*) HR-MS (ESI+) calc'd for $C_{45}H_{44}ClN_5O_{11}$: 865.2726, found $[M+H]^+$ 866.2799.

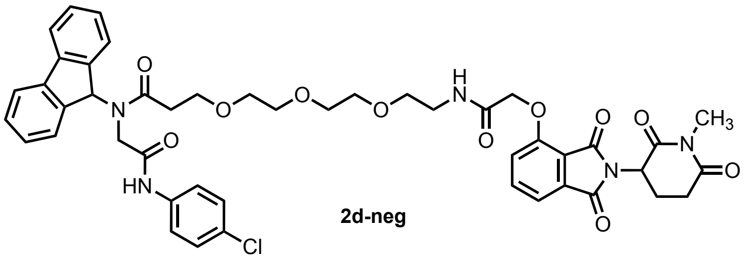

2d-neg

**N-(2-((4-chlorophenyl)amino)-2-oxoethyl)-N-(9H-fluoren-9-yl)-3-(2-(2-(2-(2-((2-(1-methyl-2,6-dioxopiperidin-3-yl)-1,3-dioxoisoindolin-4-yl)oxy)acetamido)ethoxy)ethoxy)ethoxy)propenamide (2d-neg)**

**General Procedure 4**. Reaction scale: bifunctional molecule **2d** 12 mg (13.8 μmol, 1 eq) and methyl iodide 2 mg (27.7 μmol, 2 eq). Purified by pTLC (6% MeOH/DCM) to afford **2d-neg** as a sticky solid (6.0 mg, 50 %). $^1$H NMR (600 MHz, $CDCl_3$) δ 8.62 (s, 1H), 8.02 (s, 1H), 7.79 – 7.66 (m, 6H), 7.57 – 7.50 (m, 7H), 7.47 – 7.42 (m, 2H), 7.41 – 7.36 (m 2H), 7.32 – 7.24 (m, 6H), 7.24 – 7.13 (m, 4H), 6.97 (s, 1H), 6.10 (s, 1H), 4.98 – 4.94 (m, 2H), 4.62 – 4.59 (m, 2H), 4.56 – 4.54 (m, 2H), 4.06 – 4.04 (m, 2H), 4.00 – 4.94 (m, 2H), 3.75 – 3.66 (m, 4H), 3.65 – 3.56 (m, 13H), 3.55 – 3.49 (m, 5H), 3.47 – 3.37 (m, 4H), 3.24 – 3.18 (m, 4H), 3.18 (s, 3H), 2.98 – 2.94 (m, 2H), 2.86 – 2.82 (m, 2H), 2.81 – 2.73 (m, 3H), 2.15 – 2.06 (m, 2H). $^{13}$C NMR (151 MHz, $CDCl_3$) δ 174.24, 173.80, 171.15, 171.12, 168.77, 168.70, 167.06, 166.93, 166.79, 166.75, 165.98, 154.46, 154.41, 141.92, 141.86, 141.27, 140.82, 140.75, 136.95, 136.55, 136.03, 133.63, 133.60, 129.60, 129.27, 128.95, 128.78, 127.85, 127.70, 125.94, 125.91, 125.03, 121.33, 121.18, 121.11, 120.61, 120.06, 119.45, 118.00, 117.98, 117.34, 117.30, 70.71, 70.55, 70.46, 70.38, 70.27, 70.23, 69.48, 69.38, 68.15, 67.99, 67.91, 67.65, 63.26, 59.42, 50.00, 48.16, 47.97, 39.03, 38.94, 34.45, 34.16, 31.95, 31.87, 29.72, 27.29, 22.72, 21.92. (*Note: rotameric mixture observed*) HR-MS (ESI+) calc'd for $C_{46}H_{46}ClN_5O_{11}$: 879.2882, found $[M+H]^+$ 880.2954.

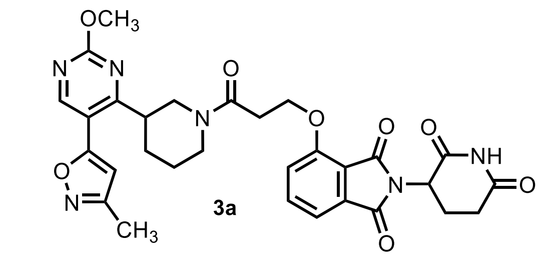

**2-(2,6-dioxopiperidin-3-yl)-4-(3-(3-(2-methoxy-5-(3-methylisoxazol-5-yl)pyrimidin-4-yl)piperidin-1-yl)-3-oxopropoxy)isoindoline-1,3-dione (3a)**

**General Procedure 1A**. Reaction scale: Thalidomide-O-C2-acid 14 mg (38.6 μmol, 1 eq) and 5-(2-methoxy-4-(piperidin-3-yl)pyrimidin-5-yl)-3-methylisoxazole 15 mg (38.6 μmol, 1 eq). Purified by pTLC (6% MeOH/DCM) to afford **3a** as a sticky solid (5.0 mg, 32 %). $^{1}$H NMR (600 MHz, $CDCl_3$) δ 8.67 (d, *J* = 1.1 Hz, 1H), 8.64 – 8.61 (m, 1H), 8.17 – 8.14 (m, 1H), 8.12 – 8.09 (m, 1H), 7.72 – 7.65 (m, 2H), 7.49 – 7.43 (m, 2H), 7.33 – 7.27 (m, 2H), 6.42 – 6.38 (m, 1H), 6.32 – 6.28 (m, 1H), 4.96 – 4.92 (m, 1H), 4.91 – 4.84 (m, 1H), 4.76 – 4.72 (m, 1H), 4.70 – 4.65 (m, 1H), 4.58 – 4.45 (m, 2H), 4.12 – 4.05 (m, 4H), 4.04 – 4.00 (m, 1H), 3.58 – 3.48 (m, 1H), 3.31 – 3.11 (m, 3H), 3.07 – 2.92 (m, 2H), 2.91 – 2.85 (m, 2H), 2.83 – 2.63 (m, 4H), 2.35 – 2.28 (m, 4H), 2.16 – 2.04 (m, 3H), 2.03 – 1.96 (m, 1H), 1.92 – 1.88 (m, 1H), 1.86 – 1.82 (m, 1H), $^{13}$C NMR (151 MHz, $CDCl_3$) δ 171.15, 170.81, 170.70, 168.31, 168.17, 166.93, 165.67, 165.53, 165.40, 165.28, 164.76, 160.43, 159.46, 159.38, 156.19, 136.64, 133.65, 133.56, 119.08, 117.01, 116.07, 115.95, 115.37, 104.53, 104.08, 65.90, 55.33, 55.25, 49.04, 48.95, 46.20, 45.67, 41.92, 40.87, 32.57, 31.33, 30.00, 25.59, 22.53, 11.42, 11.34. (*Note: rotameric mixture observed*) HR-MS (ESI+) calc'd for $C_{30}H_{30}N_6O_8$: 602.2125, found $[M+H]^+$ 603.2198.

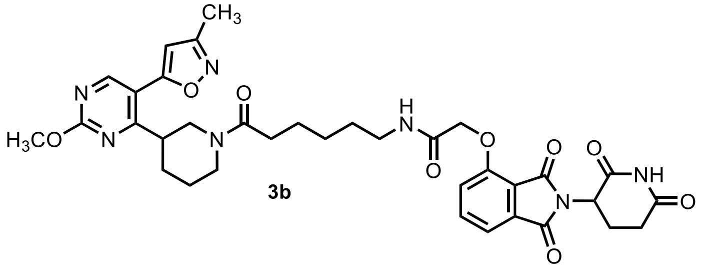

**2-((2-(2,6-dioxopiperidin-3-yl)-1,3-dioxoisoindolin-4-yl)oxy)-N-(6-(3-(2-methoxy-5-(3-methylisoxazol-5-yl)pyrimidin-4-yl)piperidin-1-yl)-6-oxohexyl)acetamide (3b)**

**General Procedure 1A**. Reaction scale: Thalidomide-O-acetamido-C5-acid 16 mg (38.6 μmol, 1 eq) and 5-(2-methoxy-4-(piperidin-3-yl)pyrimidin-5-yl)-3-methylisoxazole 15 mg (38.6 μmol, 1 eq) Purified by pTLC (6% MeOH/DCM) to afford **3b** as a sticky solid (9.0 mg, 38 %). $^{1}$H NMR (600 MHz, $CDCl_3$) δ 8.70 – 8.54 (m, 2H), 7.73 – 7.68 (m, 2H), 7.67 – 7.57 (m, 2H), 7.56 – 7.54 (m, 2H), 7.21 – 7.18 (m, 2H), 6.46 – 6.42 (m, 1H), 6.28 – 6.24 (m, 1H), 5.03 – 4.98 (m, 1H), 4.98 – 4.94 (m, 1H), 4.80 – 4.73 (m, 1H), 4.73 – 4.63 (m, 4H), 4.61 – 4.58 (m, 2H), 4.12 – 4.08 (m, 3H), 4.07 – 4.04 (m, 6H), 3.97 – 3.94 (m, 1H), 3.80 – 3.76 (m, 1H), 3.70 – 3.57 (m, 1H), 3.41 – 3.38 (m, 3H), 3.32 – 3.26 (m, 2H), 3.20 – 3.04 (m, 3H), 3.00 – 2.96 (m, 1H), 2.92 – 2.81 (m, 2H), 2.81 – 2.70 (m, 4H), 2.63 – 2.51 (m, 1H), 2.38 – 2.34 (m, 6H), 2.16 – 2.14 (m, 2H), 2.02 – 1.83 (m, 8H), 1.65 – 1.60 (m, 6H), 1.59 – 1.52 (m, 4H), 1.48 – 1.34 (m, 6H). $^{13}$C NMR (151 MHz, $CDCl_3$) δ 172.11, 172.03, 171.94, 171.43, 171.37, 171.30, 171.12, 170.95, 170.85, 170.77, 168.44, 168.36, 166.84, 166.78, 166.56, 166.19, 165.59, 165.42, 165.37, 165.23, 164.83, 164.74, 160.57, 160.43, 160.37, 159.48, 159.38, 154.70, 136.97, 133.49, 120.12, 118.38, 117.51, 117.48, 115.47, 115.39, 115.24, 104.69, 104.56, 104.01, 68.53, 60.34, 55.31, 55.19, 49.69, 49.61, 49.29, 49.26, 46.16, 46.13, 45.73, 45.63, 42.05, 41.96, 40.85, 40.80, 39.17, 39.12, 33.25, 33.18, 33.05, 33.01, 31.86, 31.34, 31.31, 30.10, 30.06, 29.87, 29.63, 28.84, 28.80, 28.77, 28.73, 26.72, 26.63, 25.69, 25.66, 25.31, 25.25, 25.17, 24.50, 22.83, 22.76, 22.62, 21.00, 14.14, 14.06, 11.44, 11.42, 11.38. (*Note: rotameric mixture observed*) HR-MS (ESI+) calc'd for $C_{35}H_{39}N_7O_9$: 701.2809, found $[M+H]^+$ 702.2878.

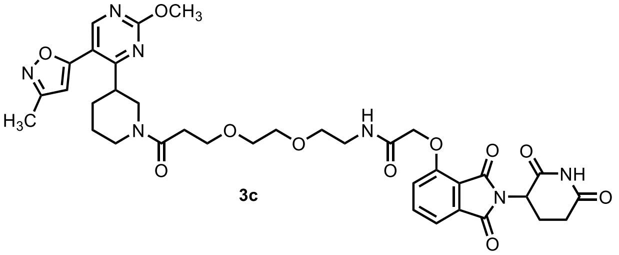

**3c**

**2-((2-(2,6-dioxopiperidin-3-yl)-1,3-dioxoisoindolin-4-yl)oxy)-N-(2-(2-(3-(3-(2-methoxy-5-(3-methylisoxazol-5-yl)pyrimidin-4-yl)piperidin-1-yl)-3-oxopropoxy)ethoxy)ethyl)acetamide (3c)**

**General Procedure 1A**. Reaction scale: Thalidomide-O-acetamido-PEG2-C2-acid 20 mg (31.7 µmol, 1 eq) and 5-(2-methoxy-4-(piperidin-3-yl)pyrimidin-5-yl)-3-methylisoxazole 14 mg (31.7 µmol, 1.1 eq) Purified by pTLC (6% MeOH/DCM) to afford **3c** as a sticky solid (5.0 mg). $^{1}$H NMR (600 MHz, $CDCl_3$) δ 8.67 – 8.64 (m, 1H), 8.63 – 8.58 (m, 1H), 7.76– 7.68 (m, 3H), 7.68 – 7.60 (m, 2H), 7.56 – 7.52 (m, 2H), 7.21 – 7.16 (m, 2H), 6.45 – 6.40 (m, 1H), 6.30 – 6.26 (m, 1H), 5.02 – 4.91 (m, 2H), 4.77 – 4.70 (m, 2H), 4.66 – 4.60 (m, 4H), 4.09 – 4.03 (m, 7H), 4.00 – 3.96 (m, 1H), 3.94 – 3.90 (m, 2H), 3.85 – 3.76 (m, 5H), 3.65 – 3.59 (m, 12H), 3.47 – 3.43 (m, 1H), 3.18 – 3.14 (m, 1H), 3.12 – 3.05 (m, 3H), 3.00 – 2.96 (m, 1H), 2.89 – 2.83 (m, 2H), 2.79 – 2.76 (m, 3H), 2.71 – 2.66 (m, 3H), 2.64 – 2.60 (m, 2H), 2.38 – 2.35 (m, 5H), 2.33 (s, 2H), 2.15 – 2.10 (m, 2H), 2.03 – 1.96 (m, 3H), 1.85 – 1.80 (m, 4H). $^{13}$C NMR (151 MHz, $CDCl_3$) δ 171.43, 171.40, 171.34, 170.82, 170.76, 169.71, 169.56, 168.37, 168.33, 166.76, 166.73, 166.72, 165.77, 165.75, 165.74, 165.63, 165.47, 165.46, 165.20, 164.83, 164.79, 160.57, 160.56, 160.47, 159.51, 159.46, 159.44, 154.33, 136.91, 136.88, 133.76, 119.02, 118.01, 117.99, 117.98, 117.20, 117.17, 115.46, 115.25, 115.24, 104.63, 104.12, 104.08, 70.31, 70.27, 70.25, 70.23, 70.14, 69.52, 69.51, 69.48, 67.66, 67.31, 67.28, 67.26, 55.38, 55.26, 49.52, 49.35, 46.14, 45.73, 45.66, 42.08, 41.87, 41.85, 40.87, 40.83, 38.97, 33.48, 33.36, 31.51, 30.29, 30.08, 30.03, 25.60, 24.52, 22.76, 11.51, 11.47, 11.44. (*Note: rotameric mixture observed*) HR-MS (ESI+) calc'd for $C_{36}H_{41}N_7O_{11}$: 747.2864, found $[M+H]^+$ 748.2924.

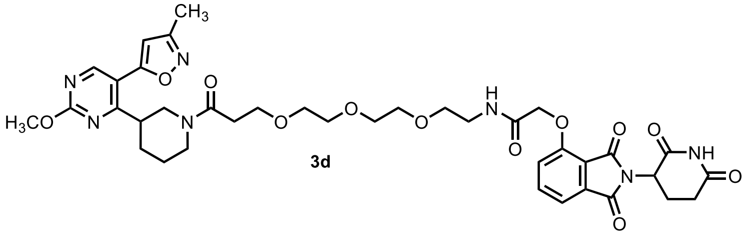

**3d**

**2-((2-(2,6-dioxopiperidin-3-yl)-1,3-dioxoisoindolin-4-yl)oxy)-N-(2-(2-(2-(3-(3-(2-methoxy-5-(3-methylisoxazol-5-yl)pyrimidin-4-yl)piperidin-1-yl)-3-oxopropoxy)ethoxy)ethoxy)ethyl)acetamide (3d)**

**General Procedure 1A**. Reaction scale: Thalidomide-O-acetamido-PEG3-C2-acid 32 mg (60.2 µmol, 1 eq) and 5-(2-methoxy-4-(piperidin-3-yl)pyrimidin-5-yl)-3-methylisoxazole 20 mg (60.2 µmol, 1 eq) Purified by pTLC (6% MeOH/DCM) to afford **3d** as a sticky solid (15.0 mg, 45%).$^{1}$H NMR (600 MHz, $CDCl_3$) δ 9.13 (s, 1H), 9.04 (s, 1H), 8.68 –8.64 (m, 1H), 8.62 – 8.58 (m, 1H), 7.74 – 7.71 (m, 2H), 7.65 – 7.62 (m, 2H), 7.54 (d, $J$ = 7.3 Hz, 2H), 7.20 – 7.17 (m, 3H), 6.46 – 6.42 (m, 1H), 6.30 – 6.28 (m, 1H), 4.96 – 4.93 (m, 2H), 4.78 – 4.58 (m, 7H), 4.10 – 4.02 (m, 7H), 3.97 – 3.94 (m, 3H), 3.81 – 3.78 (m, 5H), 3.64 – 3.56 (m, 8H), 3.46 – 3.44 (m, 1H), 3.18 – 3.06 (m, 4H), 2.96 – 2.93 (m, 2H), 2.88 – 2.84 (m, 3H), 2.82 – 2.71 (m, 5H), 2.70 – 2.65 (m, 3H), 2.65 – 2.56 (m, 3H), 2.39 – 2.34 (m, 7H), 2.18 – 2.11 (m, 3H), 2.03 – 1.98 (m, 4H), 1.86 – 1.83 (m, 4H), $^{13}$C NMR (151 MHz, $CDCl_3$) δ 171.32, 171.08, 170.88, 169.56, 168.36, 166.96, 166.75, 165.89, 165.68, 165.51, 165.29, 164.83, 160.64, 159.58, 159.48, 154.53, 137.03, 133.76, 119.42, 118.11, 117.34, 115.54, 115.34, 104.73, 104.22, 77.77, 77.34, 77.13, 76.92, 70.38, 69.58, 67.98, 67.33, 55.44, 55.32, 49.58, 49.40, 46.20, 45.75, 42.07, 41.93, 40.98, 39.18, 33.54, 33.40, 31.55, 30.38, 30.12, 29.77, 25.70, 24.62, 22.75, 11.59, (*Note: rotameric mixture observed*) HR-MS (ESI+) calc'd for $C_{38}H_{45}N_7O_{12}$: 791.3126, found $[M+H]^+$ 792.3108.

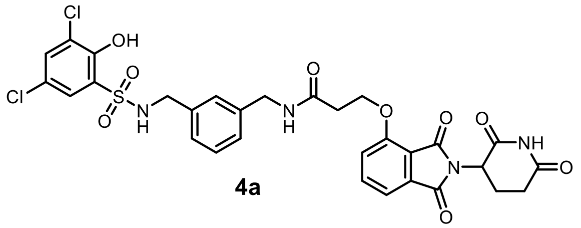

**4a**

**N-(3-(((3,5-dichloro-2-hydroxyphenyl)sulfonamido)methyl)benzyl)-3-((2-(2,6-dioxopiperidin-3-yl)-1,3-dioxoisoindolin-4-yl)oxy)propanamide (4a)**

**General Procedure 1A**. Reaction scale: thalidomide-O-C2-acid 20 mg (57.75 µmol, 1 eq) and N-(3-(aminomethyl)benzyl)-3-chloro-2-hydroxybenzenesulfonamide (**F4**) 20.85 mg (57.75 µmol, 1 eq) Purified by pTLC (6% MeOH/DCM) to afford **4a** as an off-white powder (7.4 mg, 19%). $^{1}$H NMR (500 MHz, $CDCl_3$) δ 7.70 – 7.65 (m, 2H), 7.48 – 7.44 (m, 3H), 7.32 (s, 1H), 7.23 (d, *J* = 8.2 Hz, 2H), 7.10 (s, 1H), 7.05 – 7.01 (m, 3H), 6.86 (d, *J* = 7.1 Hz, 1H), 4.95 – 4.91 (m, 1H), 4.40 (s, 2H), 4.33 (s, 2H), 4.00 – 3.96 (m, 2H), 2.80 – 2.84 (m, 4H), 2.33 – 2.29 (m, 2H). $^{13}$C NMR (151 MHz, DMSO) δ $^{13}$C NMR (151 MHz, $CDCl_3$) δ 168.70, 167.80, 166.74, 155.48, 138.88, 137.07, 133.41, 132.38, 130.88, 128.77, 128.26, 127.17, 126.89, 118.84, 117.12, 116.67, 68.16, 65.57, 49.14, 38.68, 30.32, 28.88, 23.70. HR-MS (ESI+) calc'd for $C_{30}H_{26}Cl_2N_4O_9S$: 688.0798 found $[M+H]^+$ 689.0868.

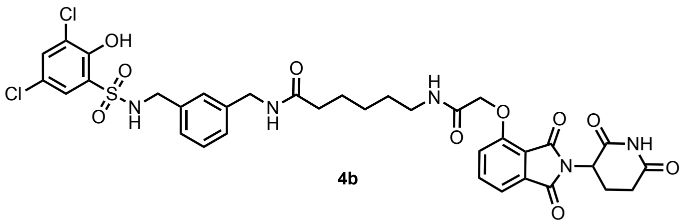

**4b**

**N-(3-(((3,5-dichloro-2-hydroxyphenyl)sulfonamido)methyl)benzyl)-6-(2-((2-(2,6-dioxopiperidin-3-yl)-1,3-dioxoisoindolin-4-yl)oxy)acetamido)hexanamide (4b)**

**General Procedure 1A**. Reaction scale: Thalidomide-O-acetamido-C5-acid 14.6 mg (32.7 µmol, 1 eq) and N-(3-(aminomethyl)benzyl)-3-chloro-2-hydroxybenzenesulfonamide (**F4**) 15.0 mg (32.7 µmol, 1 eq). Purified by pTLC (8% MeOH/DCM) to afford **4b** as an off-white powder (9.0 mg, 35%). $^{1}$H NMR (600 MHz, $CDCl_3$) δ 9.39 (s, 1H), 7.74 (d, *J* = 7.9 Hz, 1H), 7.65 – 7.62 (m, 1H), 7.56 (d, *J* = 7.3 Hz, 1H), 7.52 – 7.49 (m, 1H), 7.38 – 7.35 (m, 1H), 7.20 (d, *J* = 7.9 Hz, 1H), 7.13 – 7.10 (m, 1H), 7.04 (d, *J* = 7.6 Hz, 1H), 6.96 (d, *J* = 7.5 Hz, 1H), 6.17 (s, 1H), 4.99 – 4.96 (m, 2H), 4.69 – 4.55 (m, 2H), 4.38 – 4.33 (m, 2H), 4.15 (s, 2H), 3.44 – 3.41 (m, 1H), 3.32 – 3.27 (m, 1H), 2.89 – 2.86 (m, 1H), 2.77 – 2.74 (m, 2H), 2.31 – 2.28 (m, 2H), 2.17 – 2.14 (m, 1H), 1.72 – 1.69 (m, 2H), 1.62 – 1.59 (m, 2H), 1.44 – 1.41 (m, 2H). $^{13}$C NMR (176 MHz, $CDCl_3$) δ 174.13, 171.97, 168.85, 167.37, 166.68, 166.49, 154.83, 139.08, 137.30, 136.23, 134.04, 133.63, 128.67, 127.30, 127.26, 127.24, 127.16, 126.67, 123.98, 120.40, 118.48, 117.82, 68.61, 49.39, 47.31, 43.41, 39.03, 36.56, 31.44, 29.85, 28.75, 26.44, 25.56, 22.93. HR-MS (ESI+) calc'd for $C_{35}H_{35}Cl_2N_5O_{10}S$: 787.1482 found $[M+H]^+$ 788.1549.

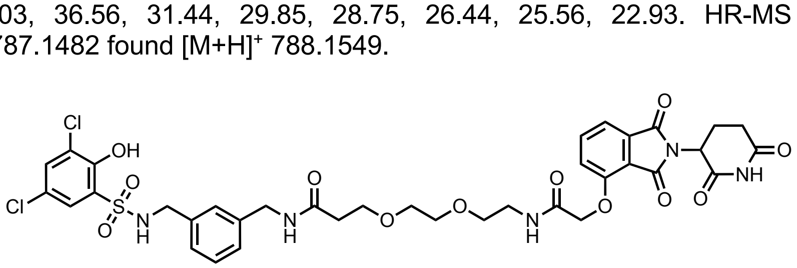

**4c**

**N-(3-(((3,5-dichloro-2-hydroxyphenyl)sulfonamido)methyl)benzyl)-3-(2-(2-(2-((2-(2,6-dioxopiperidin-3-yl)-1,3-dioxoisoindolin-4-yl)oxy)acetamido)ethoxy)ethoxy)propenamide (4c)**

**General Procedure 1A**. Reaction scale: Thalidomide-O-acetamido-PEG2-C2-acid 14 mg (28.5 µmol, 1 eq) and N-(3-(aminomethyl)benzyl)-3-chloro-2-hydroxybenzenesulfonamide (**F4**) 10.3 mg (28.5 µmol, 1 eq) Purified by pTLC (6% MeOH/DCM) to afford **4c** as a sticky solid (8.0 mg, 34%). $^{1}$H NMR (600 MHz, $CDCl_3$) δ 8.93 (s, 1H), 7.74 (d, *J* = 7.9 Hz, 1H), 7.64 – 7.61 (m, 1H), 7.56 – 7.54 (m, 2H), 7.38 – 7.36 (m, 1H), 7.21 (s, 1H), 7.18 (d, *J* = 8.4 Hz, 1H), 7.11 – 7.00 (m, 3H), 6.89 (d, *J* = 7.1 Hz, 1H), 5.02 – 4.92 (m, 1H), 4.60 (s, 2H), 4.37 (d, *J* = 6.1 Hz, 2H), 4.10 (s, 2H), 3.83 – 3.77 (m, 2H), 3.73 – 3.65 (m, 2H), 3.63 – 3.58 (m, 2H), 3.54 – 3.50 (m, 2H), 3.39 – 3.35 (m, 2H), 2.88 – 2.84 (m, 2H), 2.77 – 2.74 (m, 2H), 2.59 – 2.48 (m, 2H), 2.16 – 2.14 (m, 1H), 1.46 – 1.44 (m, 2H). $^{13}$C NMR (151 MHz, DMSO) δ 173.27, 170.58, 170.36, 167.38, 167.21, 165.91, 155.45, 139.85, 137.96, 137.40, 133.50, 133.30, 130.52, 128.50, 127.48, 126.88, 126.47, 126.42, 124.43, 121.48, 120.76, 117.19, 116.49, 70.01, 69.93, 69.28, 67.91, 67.30, 54.01, 49.26, 46.73, 42.38, 38.86, 36.57, 31.41, 22.47. HR-MS (ESI+) calc'd for $C_{36}H_{37}Cl_2N_5O_{12}S$: 833.1536, found $[M+H]^+$ 834.1595.

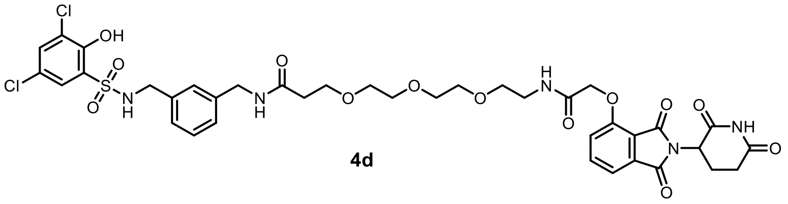

**4d**

**N-(3-(((3,5-dichloro-2-hydroxyphenyl)sulfonamido)methyl)benzyl)-3-(2-(2-(2-(2-((2-(2,6-dioxopiperidin-3-yl)-1,3-dioxoisoindolin-4-yl)oxy)acetamido)ethoxy)ethoxy)ethoxy)propenamide (4d)**

**General Procedure 1A**. Reaction scale: Thalidomide-O-acetamido-PEG3-C2-acid 9 mg (16.8 µmol, 1 eq) and N-(3-(aminomethyl)benzyl)-3-chloro-2-hydroxybenzenesulfonamide (**F4**) 8 mg (16.8 µmol, 1 eq) Purified by pTLC (6% MeOH/DCM) to afford **4d** as a brown sticky solid (4.3 mg, 30%). $^{1}$H NMR (600 MHz, $CDCl_3$) δ 8.95 (s, 1H), 7.74 (d, *J* = 7.9 Hz, 1H), 7.66 (s, 1H), 7.61 – 7.48 (m, 2H), 7.37 (d, *J* = 2.5 Hz, 1H), 7.24 – 7.15 (m, 2H), 7.05 (d, *J* = 7.2 Hz, 2H), 6.88 (d, *J* = 6.8 Hz, 1H), 4.92 – 4.89 (m, 2H), 4.62 – 4.59 (m, 2H), 4.40 – 4.36 (m, 2H), 4.10 (s, 2H), 3.83 – 3.81 (m, 2H), 3.75 – 3.38 (m, 8H), 3.16 – 3.12 (m, 1H), 2.89 – 2.82 (m, 1H), 2.79 – 2.63 (m, 2H), 2.55 (t, *J* = 5.6 Hz, 2H), 2.16 – 2.09 (m, 1H), 1.48 – 1.43 (m, 2H). $^{13}$C NMR (176 MHz, $CDCl_3$) δ 172.61, 171.57, 168.67, 167.41, 166.76, 166.16, 154.58, 139.29, 137.25, 136.12, 133.93, 133.70, 128.43, 127.48, 126.91, 126.88, 126.43, 119.73, 118.14, 117.60, 70.38, 70.24, 70.22, 70.19, 69.56, 68.03, 67.49, 49.37, 47.39, 43.19, 39.13, 37.07, 31.48, 29.85, 22.80. HR-MS (ESI+) calc'd for $C_{38}H_{41}Cl_2N_5O_{13}S$: 877.1799, found $[M+H]^+$ 878.1851.

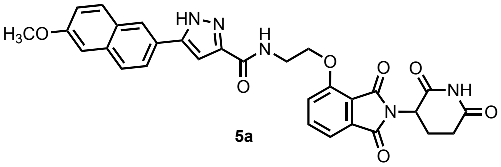

**5a**

**N-(2-((2-(2,6-dioxopiperidin-3-yl)-1,3-dioxoisoindolin-4-yl)oxy)ethyl)-5-(6-methoxynaphthalen-2-yl)-1H-pyrazole-3-carboxamide (5a)**

**General Procedure 1A**. Reaction scale: Thalidomide-O-C2-amine 30 mg (93.1 µmol, 1 eq) and 5-(6-methoxynaphthalen-2-yl)-1H-pyrazole-3-carboxylic acid 25 mg (93.1 µmol, 1.1 eq) Purified by pTLC (5% MeOH/DCM) to afford **5a** as a sticky solid (15.0 mg, 35%). $^{1}$H NMR (600 MHz, DMSO) δ 11.12 (s, 1H), 8.36 – 8.28 (m, 2H), 7.98 – 7.77 (m, 4H), 7.63 (d, *J* = 8.5 Hz, 2H), 7.48 (d, *J* = 7.2 Hz, 2H), 7.36 (s, 1H), 7.27 – 7.07 (m, 1H), 5.08 – 5.14 (m, 1H), 4.42 – 4.38 (m, 2H), 3.90 (s, 3H), 3.73 – 3.70 (m, 2H), 2.92 – 2.88 (m, 2H), 2.65 – 2.52 (m, 2H). $^{13}$C NMR (151 MHz, DMSO) δ 173.29, 170.44, 167.28, 165.74, 162.49, 158.22, 156.13, 148.02, 144.24, 137.54, 134.48, 133.76, 130.07, 128.85, 127.99, 124.40, 120.50, 119.91, 116.96, 116.02, 106.47, 103.08, 67.62, 55.73, 49.21, 38.08, 31.42, 22.47. HR-MS (ESI+) calc'd for $C_{30}H_{25}N_5O_7$: 567.1836, found $[M+H]^+$ 568.1828.

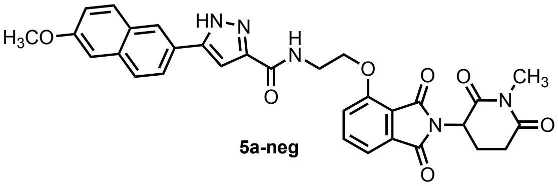

**5a-neg**

**5-(6-methoxynaphthalen-2-yl)-N-(2-((2-(1-methyl-2,6-dioxopiperidin-3-yl)-1,3-dioxoisoindolin-4-yl)oxy)ethyl)-1H-pyrazole-3-carboxamide (5a-neg)**

**General Procedure 4**. Reaction scale: Bifunctional molecule **5a** 15 mg (41.66 µmol, 1 eq) and methyl iodide 3 mg (83.3 µmol, 2 eq ). Purified by pTLC (7% MeOH/DCM) to afford **5a-neg** as a sticky solid (6.0 mg, 41%). $^{1}$H NMR (600 MHz, $CDCl_3$) δ 8.34 (d, *J* = 7.2 Hz, 1H), 7.99 (s, 1H), 7.69 (d, *J* = 8.6 Hz, 1H), 7.66 – 7.62 (m, 2H), 7.58 (d, *J* = 7.9 Hz, 1H), 7.38 (d, *J* = 7.3 Hz, 1H), 7.15 (d, *J* = 7.8 Hz, 2H), 7.12 – 7.10 (m, 1H), 7.07 – 7.05 (m, 1H), 4.90 – 4.86 (m, 1H), 4.34 – 4.30 (m, 2H), 3.89 (s, 3H), 3.74 – 3.68 (m, 2H), 3.09 (s, 3H), 2.86 – 2.75 (m, 1H), 2.64 – 2.58 (m, 2H), 2.00 – 1.83 (m, 1H). $^{13}$C NMR (151 MHz, $CDCl_3$) δ 171.14, 168.97, 166.97, 166.89, 162.48, 158.27, 156.38, 136.88, 134.45, 133.44, 129.72, 128.67, 127.62, 124.46, 124.04, 119.68, 119.51, 117.14, 116.43, 105.68,103.36, 68.79, 55.37, 49.92, 38.59, 31.72, 27.23, 21.83. HR-MS (ESI+) calc'd for $C_{31}H_{27}N_5O_7$: 581.1910, found $[M+H]^+$ 582.1984.

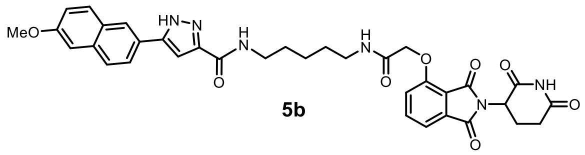

**5b**

**N-(5-(2-((2-(2,6-dioxopiperidin-3-yl)-1,3-dioxoisoindolin-4-yl)oxy)acetamido)pentyl)-5-(6-methoxynaphthalen-2-yl)-1H-pyrazole-3-carboxamide (5b)**

**General Procedure 1A**. Reaction scale: Thalidomide-O-acetamido-C5-acid 15 mg (36.02 µmol, 1 eq) and 5-(6-methoxynaphthalen-2-yl)-1H-pyrazole-3-carboxylic acid (**F5**) 10.6 mg (39.62 µmol, 1.1 eq) Purified by pTLC (8% MeOH/DCM) to afford **5b** as a white powder (8.2 mg, 34%). $^{1}$H NMR (600 MHz, $CDCl_3$) δ 12.16 (s, 1H), 7.99 (s, 1H), 7.97 – 7.91 (m, 1H), 7.84 – 7.71 (m, 3H), 7.65 (dd, *J* = 8.4, 1.8 Hz, 1H), 7.58 (d, *J* = 7.3 Hz, 1H), 7.49 (s, 1H), 7.25 (d, *J* = 8.5 Hz, 1H), 7.21 – 7.16 (m, 2H), 7.14 (d, *J* = 2.4 Hz, 1H), 5.06 (dd, *J* = 12.7, 5.3 Hz, 1H), 4.78 – 4.75 (m, 1H), 4.65 – 4.62 (m, 1H), 3.94 (s, 3H), 3.82 – 3.66 (m, 2H), 3.10 (s, 1H), 3.02 – 2.98 (m, 2H), 2.93 – 2.76 (m, 2H), 2.26 – 2.18 (m, 1H), 1.78 – 1.82 (m, 4H), 1.51 – 1.46 (m, 2H). $^{13}$C NMR (151 MHz, $CDCl_3$) δ 173.69, 169.28, 167.05, 166.64, 166.40, 161.57, 158.58, 155.42, 147.59, 145.09, 137.30, 134.86, 133.51, 129.95, 128.87, 128.06, 124.44, 123.78, 123.74, 121.32, 119.99, 118.76, 118.04, 105.92, 103.81, 69.56, 55.53, 49.33, 39.37, 39.18, 31.51, 29.85, 29.12, 28.93, 23.83. HR-MS (ESI+) calc'd for $C_{35}H_{34}N_6O_8$: 666.2438, found $[M+H]^+$ 667.2512.

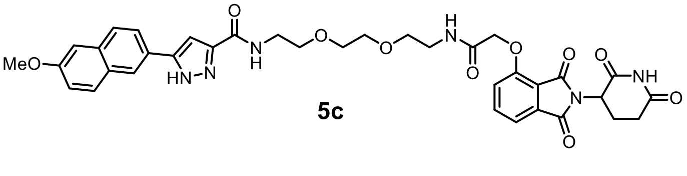

**5c**

**N-(2-(2-(2-(2-((2-(2,6-dioxopiperidin-3-yl)-1,3-dioxoisoindolin-4-yl)oxy)acetamido)ethoxy)ethoxy)ethyl)-5-(6-methoxynaphthalen-2-yl)-1H-pyrazole-3-carboxamide (5c)**

**General Procedure 1A**. Reaction scale: Thalidomide-O-acetamido-PEG2-C2-amine 15 mg (32.44 µmol, 1 eq) and 5-(6-methoxynaphthalen-2-yl)-1H-pyrazole-3-carboxylic acid (**F5**) 9.57 mg (35.68 µmol, 1.1 eq) Purified by pTLC (8% MeOH/DCM) to afford **5c** as a white powder (5.5 mg, 24%). $^{1}$H NMR (600 MHz,

$CDCl_3$) δ 10.33 (s, 1H), 8.00 (s, 1H), 7.78 – 7.74 (m, 2H), 7.71 – 7.65 (m, 3H), 7.64 – 7.61 (m, 1H), 7.49 (d, *J* = 7.3 Hz, 1H), 7.20 – 7.07 (m, 4H), 5.03 – 4.98 (m, 1H), 4.64 – 4.65 (m, 2H), 3.93 (s, 3H), 3.70 – 3.54 (m, 11H), 2.97 – 2.88 (m, 1H), 2.88 – 2.71 (m, 2H), 2.22 – 2.08 (m, 2H). $^{13}C$ NMR (151 MHz, $CDCl_3$) δ 172.62, 169.05, 167.35, 166.79, 166.08, 161.94, 158.43, 154.58, 147.00, 145.55, 137.10, 134.68, 133.71, 129.91, 128.89, 127.87, 124.37, 124.34, 123.90, 119.82, 119.42, 118.02, 117.43, 105.90, 103.45, 70.28, 70.22, 69.83, 69.61, 68.10, 55.51, 49.36, 39.32, 31.51, 29.84, 22.81. HR-MS (ESI+) calc'd for $C_{36}H_{36}N_6O_{10}$: 712.2493, found $[M+H]^+$ 713.2557.

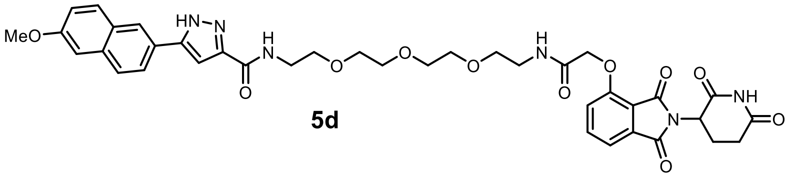


**5d**

**N-(1-((2-(2,6-dioxopiperidin-3-yl)-1,3-dioxoisoindolin-4-yl)oxy)-2-oxo-6,9,12-trioxa-3-azatetradecan-14-yl)-5-(6-methoxynaphthalen-2-yl)-1H-pyrazole-3-carboxamide (5d)**

**General Procedure 1A**. Reaction scale: Thalidomide-O-acetamido-PEG3-C2-amine 15 mg (29.6 µmol, 1 eq) and 5-(6-methoxynaphthalen-2-yl)-1H-pyrazole-3-carboxylic acid (**F5**) 8.7 mg (32.6 µmol, 1.1 eq) Purified by pTLC (8% MeOH/DCM) to afford **5d** as a white powder (13.58 mg, 62%). $^1H$ NMR (500 MHz, $CDCl_3$) δ 10.41 (s, 1H), 8.03 (d, *J* = 1.7 Hz, 1H), 7.75 (t, *J* = 8.8 Hz, 2H), 7.71 – 7.68 (m, 3H), 7.61 – 7.59 (m, 1H), 7.52 (d, *J* = 7.3 Hz, 1H), 7.19 – 7.13 (m, 2H), 7.11 (d, *J* = 8.4 Hz, 2H), 5.05 – 5.01 (m, 1H), 4.65 – 4.51 (m, 2H), 3.93 (s, 3H), 3.71 – 3.56 (m, 16H), 2.97 – 2.71 (m, 4H), 2.17 – 2.14 (m, 1H). $^{13}C$ NMR (151 MHz, $CD_3OD$_SPE) δ 174.63, 173.03, 171.48, 168.48, 167.90, 159.79, 157.60, 138.11, 136.05, 135.03, 130.70, 130.28, 128.78, 125.20, 124.94, 120.90, 120.61, 118.37, 116.95, 106.77, 103.42, 69.44, 61.55, 55.82, 54.81, 50.46, 39.60, 32.14, 23.62, 20.86, 14.45. HR-MS (ESI+) calc'd for $C_{38}H_{40}N_6O_{11}$: 756.2755, found $[M+H]^+$ 757.2830.

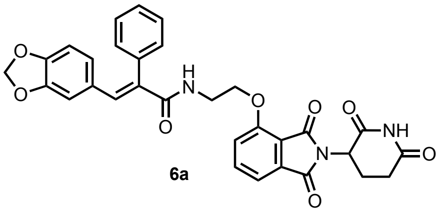


**6a**

**(E)-3-(benzo[d][1,3]dioxol-5-yl)-N-(2-((2-(2,6-dioxopiperidin-3-yl)-1,3-dioxoisoindolin-4-yl)oxy)ethyl)-2-phenylacrylamide (6a)**

**General Procedure 1A**. Reaction scale: Thalidomide-O-C2-amine 18 mg (55.9 µmol, 1 eq) and (E)-3-(benzo[d][1,3]dioxol-5-yl)-2-phenylacrylic acid 15 mg (55.9 µmol, 1 eq) Purified by pTLC (5% MeOH/DCM) to afford **6a** as an off-white solid (15.0 mg, 55%). $^1H$ NMR (600 MHz, DMSO) $^1H$ NMR (600 MHz, $CDCl_3$) δ 8.33 (s, 1H), 7.73 (s, 1H), 7.69 – 7.67 (m, 1H), 7.47 (d, *J* = 7.3 Hz, 1H), 7.42 – 7.37 (m, 3H), 7.25 (d, *J* = 8.5 Hz, 1H), 7.21 – 7.17 (m, 2H), 6.67 (dd, *J* = 8.1, 1.7 Hz, 1H), 6.62 (d, *J* = 8.2 Hz, 1H), 6.24 (d, *J* = 1.7 Hz, 1H), 6.17 (d, *J* = 5.9 Hz, 1H), 5.85 (s, 2H), 4.94 – 4.92 (m, 1H), 4.29 (t, *J* = 5.3 Hz, 2H), 3.82 – 3.63 (m, 2H), 2.95 – 2.69 (m, 4H), 2.12 – 2.10 (m, 1H).$^{13}C$ NMR (151 MHz, DMSO) δ 171.04, 168.09, 166.92, 165.52, 156.18, 148.07, 147.39, 137.06, 136.70, 135.85, 133.75, 132.24, 129.85, 129.78, 128.63, 126.49, 119.50, 117.31, 116.41, 109.36, 108.15, 101.18, 67.92, 49.13, 39.44, 31.41, 22.62. HR-MS (ESI+) calc'd for $C_{31}H_{25}N_3O_8$: 567.1642, found $[M+H]^+$ 568.1713

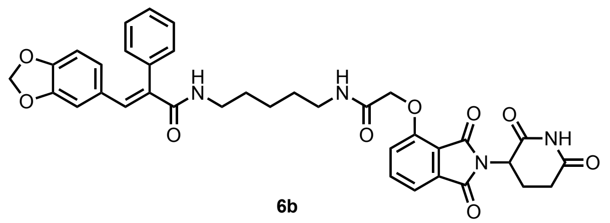

**6b**

**(E)-3-(benzo[d][1,3]dioxol-5-yl)-N-(5-(2-((2-(2,6-dioxopiperidin-3-yl)-1,3-dioxoisoindolin-4-yl)oxy)acetamido)pentyl)-2-phenylacrylamide (6b)**

**General Procedure 1A**. Reaction scale: Thalidomide-O-acetamido-C5-amine 14 mg (26.1 µmol, 1 eq) and (E)-3-(benzo[d][1,3]dioxol-5-yl)-2-phenylacrylic acid 7 mg (26.1 µmol, 1 eq). Purified by pTLC (5% MeOH/DCM) to afford **6b** as an off-white solid (14.0 mg, 64 %). $^1$H NMR (600 MHz, CDCl3) δ 9.45 (s, 1H), 7.82 – 7.71 (m, 2H), 7.66 – 7.60 (m, 1H), 7.58 (d, J = 7.3 Hz, 1H), 7.51 – 7.44 (m, 3H), 7.30 – 7.19 (m, 3H), 6.73 (dd, J = 8.1, 1.7 Hz, 1H), 6.67 (d, J = 8.2 Hz, 1H), 6.48 – 6.45 (m, 1H), 6.26 (d, J = 1.6 Hz, 1H), 5.90 (s, 2H), 5.56 – 5.48 (m, 1H), 5.12 – 5.07 (m, 1H), 4.70 – 4.66 (m, 2H), 3.78 – 3.72 (m, 2H), 3.52 – 3.46 (m, 1H), 3.42 – 3.19 (m, 5H), 3.02 (s, 1H), 3.01 – 2.75 (m, 3H), 2.27 – 2.16 (m, 1H), 1.64 – 1.58 (m, 2H), 1.55 – 1.39 (m, 8H). $^{13}$C NMR (151 MHz, $CDCl_3$) δ 171.49, 168.39, 167.53, 166.86, 166.67, 166.21, 154.57, 148.00, 137.10, 137.06, 136.20, 133.56, 132.15, 129.90, 129.80, 129.10, 128.69, 126.53, 119.91, 118.30, 117.49, 109.28, 108.22, 108.16, 101.15, 68.23, 55.64, 55.61, 49.35, 43.60, 40.21, 38.98, 31.45, 29.17, 28.72, 24.13, 22.81, 18.58, 17.17, 12.59. HR-MS (ESI$^+$) calc'd for $C_{36}H_{34}N_4O_9$: 666.6870, found [M+H]$^+$ 667.2391

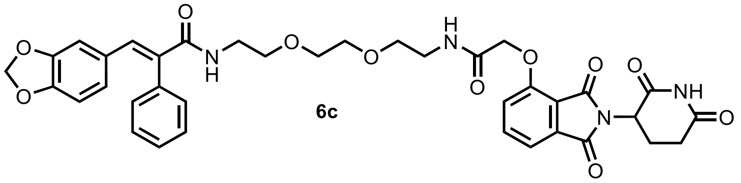

**6c**

**(E)-3-(benzo[d][1,3]dioxol-5-yl)-N-(2-(2-(2-(2-((2-(2,6-dioxopiperidin-3-yl)-1,3-dioxoisoindolin-4-yl)oxy)acetamido)ethoxy)ethoxy)ethyl)-2-phenylacrylamide (6c)**

**General Procedure 1A**. Reaction scale: Thalidomide-O-acetamido-PEG2-C2-amine 12.5 mg (25.0 µmol, 1 eq) and (E)-3-(benzo[d][1,3]dioxol-5-yl)-2-phenylacrylic acid 7 mg (25.0 µmol, 1 eq). Purified by pTLC (7% MeOH/DCM) to afford **6c** as an off-white solid (12.0 mg, 60 %).$^1$H NMR (600 MHz, $CDCl_3$) δ 8.97 (s, 1H), 7.83 – 7.75 (m, 1H), 7.70 – 7.68 (m, 1H), 7.60 (d, *J* = 7.3 Hz, 1H), 7.55 – 7.43 (m, 2H), 7.35 – 7.29 (m, 2H), 7.25 (d, *J* = 8.4 Hz, 1H), 6.73 (dd, *J* = 8.2, 1.7 Hz, 1H), 6.69 (d, *J* = 8.2 Hz, 1H), 6.28 (d, *J* = 1.6 Hz, 1H), 6.01 (d, *J* = 5.7 Hz, 1H), 5.92 (s, 1H), 5.08 – 4.95 (m, 1H), 4.71 (s, 1H), 3.78 (t, *J* = 7.4 Hz, 1H), 3.68 – 3.56 (m, 6H), 3.56 – 3.54 (m, 1H), 3.26 – 3.23 (m, 1H), 2.97 – 2.72 (m, 2H), 1.56 – 1.46 (m, 5H).$^{13}$C NMR (151 MHz, $CDCl_3$) δ 171.20, 168.22, 167.64, 166.97, 166.67, 165.89, 154.40, 148.01, 147.36, 137.01, 136.16, 133.67, 132.30, 129.97, 129.68, 129.09, 128.58, 126.52, 119.35, 118.01, 117.33, 109.27, 108.15, 101.15, 70.29, 70.00, 69.77, 69.53, 67.87, 55.62, 49.31, 43.60, 39.91, 39.08, 31.41, 22.69, 18.62, 17.24, 12.56. HR-MS (ESI$^+$) calc'd for $C_{37}H_{36}N_4O_{11}$: 712.2381, found [M+H]$^+$ 713.2449.

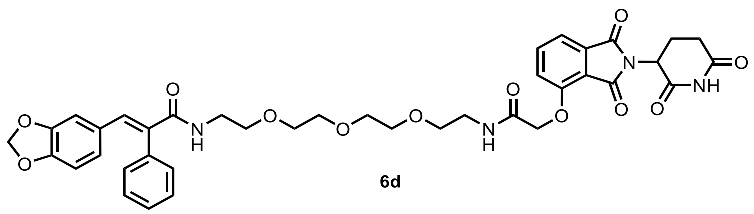

**6d**

**(E)-3-(benzo[d][1,3]dioxol-5-yl)-N-(1-((2-(2,6-dioxopiperidin-3-yl)-1,3-dioxoisoindolin-4-yl)oxy)-2-oxo-6,9,12-trioxa-3-azatetradecan-14-yl)-2-phenylacrylamide (6d)**

**General Procedure 1A.** Reaction scale: Thalidomide-O-acetamido-PEG3-C2-amine 11.5 mg (20.5 µmol, 1 eq) and (E)-3-(benzo[d][1,3]dioxol-5-yl)-2-phenylacrylic acid 6 mg (20.5 µmol, 1 eq). Purified by pTLC (7% MeOH/DCM) to afford **6d** as an off-white solid (8.0 mg, 52%).$^{1}$H NMR (600 MHz, CDCl3) δ 8.94 (s, 1H), 7.80 – 7.71 (m, 2H), 7.69 – 7.67 (m, 1H), 7.56 (d, J = 7.3 Hz, 1H), 7.49 – 7.38 (m, 3H), 7.33 – 7.24 (m, 3H), 7.22 (d, J = 8.4 Hz, 1H), 6.69 (dd, J = 8.2, 1.7 Hz, 1H), 6.64 (d, J = 8.1 Hz, 1H), 6.25 (d, J = 1.6 Hz, 1H), 6.01 (t, J = 5.2 Hz, 1H), 5.88 (s, 2H), 5.01 – 4.89 (m, 1H), 4.67 (s, 2H), 3.74 (s, 1H), 3.69 – 3.52 (m, 16H), 3.50 (q, J = 5.6 Hz, 2H), 3.20 (s, 1H), 2.92 – 2.65 (m, 3H), 2.21 – 2.09 (m, 1H), 1.54 – 1.40 (m, 7H). $^{13}$C NMR (151 MHz, CDCl3) δ 171.16, 168.21, 166.64, 165.87, 147.96, 147.36, 137.04, 136.19, 132.49, 129.96, 129.64, 129.14, 128.51, 126.44, 119.44, 118.03, 117.34, 109.29, 108.14, 70.27, 70.23, 70.21, 70.19, 69.77, 69.51, 67.94, 55.57, 49.27, 43.60, 39.87, 39.12, 31.38, 22.69, 18.72, 17.34, 12.63. HR-MS (ESI$^{+}$) calc'd for $C_{39}H_{40}N_4O_{12}$: 756.2643, found [M+H]$^{+}$ 757.2720.

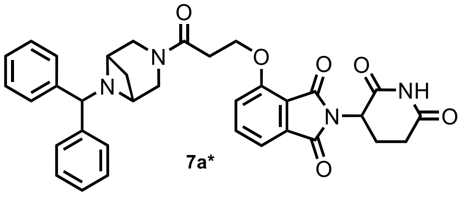

**7a***

**4-(3-(6-benzhydryl-3,6-diazabicyclo[3.1.1]heptan-3-yl)-3-oxopropoxy)-2-(2,6-dioxopiperidin-3-yl)isoindoline-1,3-dione (7a*)**

**General Procedure 1A.** Reaction scale: Thalidomide-O-C2-acid 15 mg (43.6 µmol, 1 eq) of and 6-benzhydryl-3,6-diazabicyclo[3.1.1]heptane 20.0 mg (43.6 µmol, 1. eq) Purified by pTLC (7% MeOH/DCM) to afford **7a*** as an off-white solid (16 mg, 48%).$^{1}$H NMR (500 MHz, $CDCl_3$) δ 8.10 (s, 1H), 7.78 (d, *J* = 7.3 Hz, 1H), 7.68 – 7.50 (m, 5H), 7.33 (s, 10H), 4.98 – 4.94 (m, 1H), 4.80 (s, 1H), 4.67 – 4.63 (m, 3H), 3.84 (s, 1H), 3.64 – 3.60 (m, 5H), 3.07 (s, 1H), 3.04 – 2.89 (m, 3H), 2.89 – 2.70 (m, 4H), 2.27 – 2.12 (m, 1H), 1.70 (s, 5H). $^{13}$C NMR (151 MHz, $CDCl_3$) δ 173.99, 171.10, 171.05, 168.22, 167.07, 165.74, 156.65, 136.56, 133.79, 128.72, 127.45, 127.30, 118.99, 117.11, 115.78, 69.14, 64.89, 53.47, 49.09, 44.46, 42.68, 38.63, 33.19, 31.39, 28.83, 25.71, 22.62. (*Note: rotameric mixture observed*), HR-MS (ESI$^{+}$) calc'd for $C_{34}H_{32}N_4O_6$: 592.2322, found [M+H]$^{+}$ 593.2398.

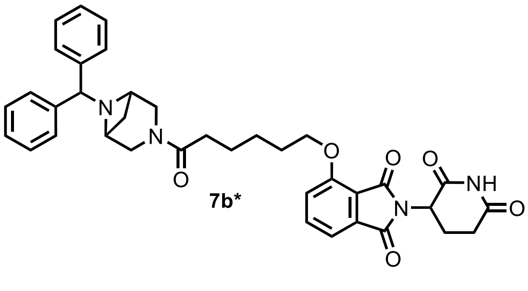

**7b***

**4-((6-(6-benzhydryl-3,6-diazabicyclo[3.1.1]heptan-3-yl)-6-oxohexyl)oxy)-2-(2,6-dioxopiperidin-3-yl)isoindoline-1,3-dione (7b*)**

**General Procedure 1A.** Reaction scale: Thalidomide-O-C5-acid 12.7 mg (32.7 µmol, 1 eq) of and 6-benzhydryl-3,6-diazabicyclo[3.1.1]heptane 15.0 mg (32.7 µmol, 1. eq). Purified by pTLC (7% MeOH/DCM) to afford **7b*** as an off-white solid (15.0 mg, 52%). $^{1}$H NMR (500 MHz, $CDCl_3$) δ 8.14 (s, 1H), 7.72 – 7.68 (m, 1H), 7.58 – 7.54 (m, 3H), 7.47 (d, *J* = 7.1 Hz, 1H), 7.31 – 7.19 (m, 7H), 4.97 – 4.94 (m, 1H), 4.75 (s, 1H), 4.26 – 4.23 (m, 2H), 3.79 – 3.60 (m, 4H), 3.61 – 3.42 (m, 2H), 3.20 – 3.14 (m, 1H), 2.95 – 2.65 (m, 4H), 2.44 – 2.40 (m, 2H), 2.12 – 2.08 (m, 1H), 1.98 – 1.96 (m, 2H), 1.86 – 1.84 (m, 3H), 1.66 – 1.62 (m, 3H), 1.55 – 1.49 (m, 3H), 1.49 – 1.38 (m, 4H). $^{13}$C NMR (151 MHz, $CDCl_3$) δ 174.11, 171.13, 168.26, 167.08, 165.75, 156.67, 136.62, 133.75, 128.89, 128.77, 127.46, 127.33, 119.05, 117.06, 115.76, 69.15, 66.52,

64.84, 55.24, 53.47, 49.07, 44.41, 43.26, 42.63, 33.17, 31.37, 29.70, 28.79, 28.77, 28.49, 25.68, 24.35, 24.33, 22.61, 18.57, 17.19, 12.50. (*Note: rotameric mixture observed*) HR-MS (ESI$^+$) calc'd for $C_{37}H_{38}N_4O_6$: 634.2791, found $[M+H]^+$ 635.2862.

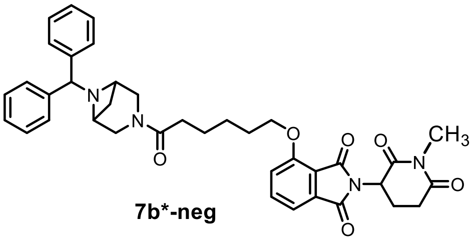


**7b*-neg**

**4-((6-(6-benzhydryl-3,6-diazabicyclo[3.1.1]heptan-3-yl)-6-oxohexyl)oxy)-2-(2,6-dioxopiperidin-3-yl)isoindoline-1,3-dione (7b*-neg)**

**General Procedure 4.** Reaction scale: bifunctional molecule **7b** 10 mg (15.8 µmol, 1.0 eq) of and 4.9 mg (31.6 µmol, 2.0 eq) Purified by pTLC (7% MeOH/DCM) to afford **7b*-neg** as an off-white solid (5 mg, 45%). $^1$H NMR (600 MHz, $CDCl_3$) δ 7.85 (s, 2H), 7.69 – 7.67 (m, 1H), 7.47 – 7.45 (m, 3H), 7.32 – 7.03 (m, 6H), 5.02 – 4.87 (m, 1H), 4.67 – 6.63 (m, 1H), 4.28 – 4.13 (m, 2H), 3.73 – 3.60 (m, 1H), 3.56 – 3.54 (m, 1H), 3.48 – 3.46 (m, 1H), 3.36 – 3.34 (m, 1H), 3.21 – 3.18 (m, 2H), 3.00 – 2.90 (m, 1H), 2.83 – 2.61 (m, 2H), 2.39 – 2.36 (m, 1H), 2.11 – 1.99 (m, 1H), 1.96 – 1.92 (m, 1H), 1.88 – 1.75 (m, 2H), 1.63 – 1.61 (m 3H). $^{13}$C NMR (151 MHz, DMSO) δ 173.00, 171.82, 169.73, 166.87, 165.36, 156.09, 142.25, 137.14, 133.25, 132.75, 129.65, 128.61, 127.21, 127.11, 119.86, 116.20, 115.22, 68.82, 49.33, 32.26, 31.14, 28.43, 26.64, 25.20, 24.08, 21.24. (*Note: rotameric mixture observed*) HR-MS (ESI$^+$) calc'd for $C_{38}H_{40}N_4O_6$: 648.2948, found $[M+H]^+$ 649.3021.

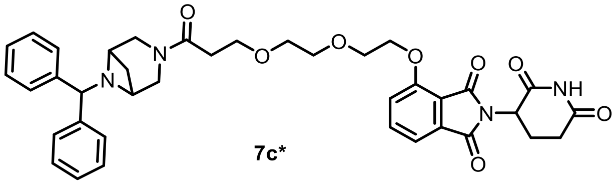


**7c***

**4-(2-(2-(3-(6-benzhydryl-3,6-diazabicyclo[3.1.1]heptan-3-yl)-3-oxopropoxy)ethoxy)ethoxy)-2-(2,6-dioxopiperidin-3-yl)isoindoline-1,3-dione (7c*)**

**General Procedure 1A.** Reaction scale: Thalidomide-O-PEG2-acid 13.2 mg (32.7 µmol, 1 eq) of and 6-benzhydryl-3,6-diazabicyclo[3.1.1]heptane 15.0 mg (32.7 µmol, 1. eq). Purified by pTLC (7% MeOH/DCM) to afford **7b*** as an off-white solid (15.0 mg, 52%). $^1$H NMR (600 MHz, CDCl3) δ 8.81 (s, 1H), 7.67 – 7.65 (d, *J* = 7.9 Hz, 1H), 7.57 – 7.49 (m, 3H), 7.42 – 7.39 (m,1H), 7.26 – 7.18 (m, 4H), 7.17 – 7.14 (m, 2H), 4.86 – 4.84 (m, 1H), 4.72 – 4.70 (m, 1H), 4.34 – 4.30 (m, 2H), 3.96 – 3.94 (m, 2H), 3.88 – 3.75 (m, 4H), 3.72 – 3.63 (m, 4H), 3.57 – 3.54 (m, 2H), 3.48 – 3.46 (m, 1H), 3.37 – 3.34 (m, 1H), 3.16 – 3.14 (m, 1H), 2.67 – 2.64 (m, 3H), 2.43 (s, 1H), 1.92 (s, 1H), 1.48 – 1.38 (m, 6H). $^{13}$C NMR (151 MHz, DMSO) δ 172.85, 171.35, 170.00, 166.86, 165.33, 155.86, 142.26, 137.04, 133.27, 128.58, 127.24, 127.07, 120.05, 116.34, 115.43, 70.21, 69.86, 68.94, 68.74, 66.72, 53.52, 48.79, 41.78, 32.94, 30.99, 22.03, 18.07, 16.74, 12.43. (*Note: rotameric mixture observed*) HR-MS (ESI$^+$) calc'd for $C_{38}H_{40}N_4O_8$: 680.2846, found $[M+H]^+$ 681.2843.

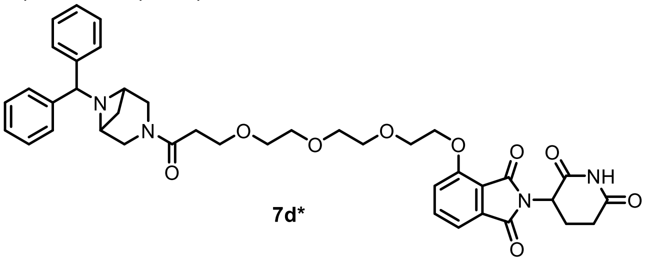


**7d***

**4-(2-(2-(2-(3-(6-benzhydryl-3,6-diazabicyclo[3.1.1]heptan-3-yl)-3-oxopropoxy)ethoxy)ethoxy)ethoxy)-2-(2,6-dioxopiperidin-3-yl)isoindoline-1,3-dione (7d*)**

**General Procedure 1A.** Reaction scale: Thalidomide-O-PEG3-C2-acid 10.0 mg (21.8 µmol, 1.0 eq) of and 6-benzhydryl-3,6-diazabicyclo[3.1.1]heptane 10.0 mg (21.8 µmol, 1.0 0eq) Purified by pTLC (7% MeOH/DCM) to afford **7d*** as an off-white solid (10 mg, 50%). $^{1}H$ NMR (600 MHz, $CDCl_3$) δ 8.62 (s, 1H), 7.65 – 7.63 (m, 1H), 7.52 – 7.50 (m, 2H), 7.43 – 7.41 (m, 1H), 7.31 – 7.24 (m, 3H), 7.24 – 7.15 (m, 2H), 4.94 – 4.92 (m, 1H), 4.69 (s, 1H), 4.35 – 4.33 (s, 1H), 3.94 – 3.92 (m, 1H), 3.87 – 3.85 (m, 2H), 3.78 – 3.74 (m, 2H), 3.73 – 3.63 (m, 5H), 3.59 – 3.57 (m, 2H), 3.45 – 3.43 (m, 1H), 3.16 – 3.14 (m, 1H), 2.86 – 2.84 (m, 1H), 2.80 – 2.68 (m, 2H), 2.65 – 2.62 (m, 1H), 2.11 – 2.07 (m, 1H), 1.52 – 1.38 (m, 5H). $^{13}C$ NMR (151 MHz, $CDCl_3$) δ 172.52, 171.21, 171.19, 168.27, 167.00, 165.83, 156.34, 156.32, 136.54, 133.71, 133.69, 128.78, 128.69, 127.40, 127.39, 127.35, 119.39, 119.35, 117.40, 117.37, 116.19, 116.18, 71.10, 70.47, 70.39, 70.37, 70.33, 69.24, 69.23, 69.15, 69.10, 67.13, 67.09, 64.66, 55.38, 49.17, 49.15, 44.47, 43.38, 42.55, 33.67, 31.41, 29.71, 28.52, 22.63, 18.58, 17.21, 12.47. (*Note: rotameric mixture observed*) HR-MS ($ESI^+$) calc'd for $C_{40}H_{44}N_4O_9$: 724.3108, found $[M+H]^+$ 725.3179.

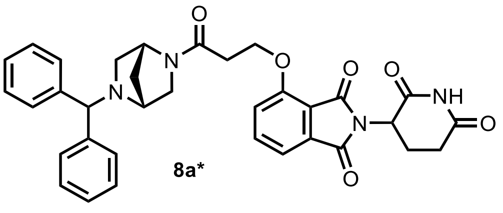

8a*

**4-(3-((1R,4R)-5-benzhydryl-2,5-diazabicyclo[2.2.1]heptan-2-yl)-3-oxopropoxy)-2-(2,6-dioxopiperidin-3-yl)isoindoline-1,3-dione (8a)**

**General Procedure 1A.** Reaction scale: Thalidomide-O-C2-acid 15 mg (43.6 µmol, 1 eq) and (1R,4R)-2-benzhydryl-2,5-diazabicyclo[2.2.1]heptane 20 mg (43.6 µmol, 1.0 eq). Purified by pTLC (7% MeOH/DCM) to afford **8a** as an off-white solid (8.0 mg, 40%). $^{1}H$ NMR (600 MHz, $CDCl_3$) δ 8.89 (s, 1H), 7.67 – 7.65 (m, 2H), 7.59 – 7.38 (m, 6H), 7.36 – 7.21 (m, 6H), 7.22 – 7.05 (m, 3H), 4.94 – 4.90 (m, 1H), 4.77 (s, 1H), 4.71 – 4.38 (m, 4H), 3.80 – 3.78 (s, 1H), 3.61 – 3.40 (m, 1H), 3.24 – 3.04 (m, 1H), 2.97 – 2.94 (s, 1H), 2.91 – 2.63 (m, 6H), 2.56 – 2.54 (m, 1H), 2.04 (s, 2H), 1.94 (s, 1H), 1.74 (s, 1H), 1.66 (s, 1H), 1.44 – 1.31 (m, 2H). $^{13}C$ NMR (151 MHz, $CDCl_3$) δ 171.57, 171.51, 168.51, 167.76, 167.52, 167.06, 167.02, 165.78, 165.68, 156.26, 156.13, 136.74, 136.71, 133.67, 133.64, 128.63, 127.74, 127.34, 127.30, 127.20, 119.24, 119.21, 117.06, 116.09, 116.06, 72.69, 71.99, 65.91, 65.83, 59.65, 59.37, 59.13, 57.99, 56.51, 55.4 (*Note: rotameric mixture observed*). HR-MS ($ESI^+$) calc'd for $C_{34}H_{32}N_4O_6$: 592.2322, found $[M+H]^+$ 593.2400.

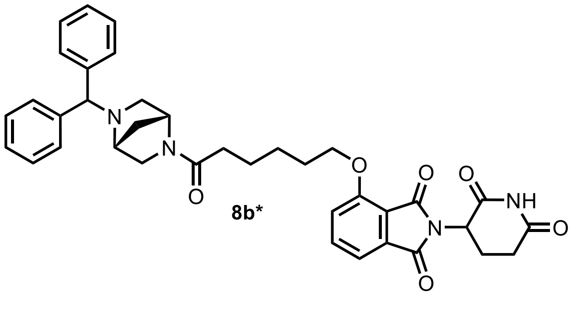

8b*

**4-((6-((1R,4R)-5-benzhydryl-2,5-diazabicyclo[2.2.1]heptan-2-yl)-6-oxohexyl)oxy)-2-(2,6-dioxopiperidin-3-yl)isoindoline-1,3-dione (8b*)**

**General Procedure 1A.** Reaction scale: Thalidomide-O-C5-acid 12.7 mg (32.7 µmol, 1.0 eq) and (1R,4R)-2-benzhydryl-2,5-diazabicyclo[2.2.1]heptane15 mg (32.7 µmol, 1.0 eq). Purified by pTLC (7% MeOH/DCM) to afford **8b** as an off-white solid (6 mg, 42%). $^{1}H$ NMR (600 MHz, $CDCl_3$) δ 8.18 (s, 1H), 7.72 – 7.63 (m,

1H), 7.51 – 7.40 (m, 4H), 7.31 – 7.24 (m, 4H), 7.24 – 7.14 (m, 2H), 4.98 – 4.94 (m, 1H), 4.76 (s, 1H), 4.63 (s, 1H), 4.59 – 4..54 (s, 1H), 4.27 (s, 1H), 4.23 – 4.20 (s, 2H), 3.81 – 3.79 (s, 1H), 3.66 – 3.64 (s, 1H), 3.52 – 3.48 (s, 1H), 3.25 – 3.22 (m, 1H), 3.19 – 3.15 (m, 1H), 3.00 – 2.92 (m, 1H), 2.92 – 2.67 (m, 3H), 2.62 – 2.57 (m, 1H), 2.47 – 2.39 (m, 1H), 2.40 – 2.33 (m, 1H), 2.33 – 2.24 (m, 1H), 2.15 – 2.05 (m, 1H), 2.06 – 1.99 (m, 1H), 1.99 – 1.88 (m, 2H), 1.79 – 1.69 (m, 3H), 1.69 – 1.45 (m, 3H). (*Note: rotameric mixture observed*) HR-MS (ESI$^+$) calc'd for $C_{37}H_{38}N_4O_6$: 634.7330, found $[M+H]^+$ 635.2867.

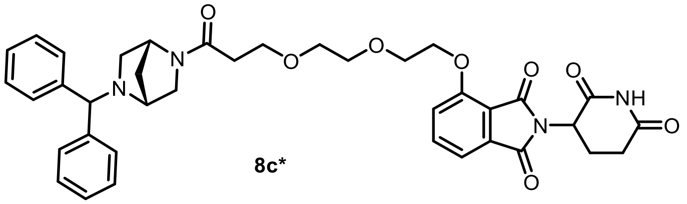

**8c***

**4-(2-(2-(3-((1R,4R)-5-benzhydryl-2,5-diazabicyclo[2.2.1]heptan-2-yl)-3-oxopropoxy)ethoxy)ethoxy)-2-(2,6-dioxopiperidin-3-yl)isoindoline-1,3-dione (8c*)**

**General Procedure 1A.** Reaction scale: Thalidomide-O-PEG2-acid 14.2 mg (32.7 µmol, 1.0 eq) and (1R,4R)-2-benzhydryl-2,5-diazabicyclo[2.2.1]heptane 15 mg (32.7 µmol, 1.0 eq). Purified by pTLC (7% MeOH/DCM) to afford **8c** as an off-white solid (11.0 mg, 38%). $^1$H NMR (600 MHz, $CDCl_3$) δ 8.80 (s, 1H), 7.66 – 7.62 (m, 2H), 7.53 – 7.34 (m, 9H), 7.31 – 7.19 (m, 9H), 7.15 – 7.12 (m, 3H), 7.06 (s, 1H), 4.92 – 4.88 (s, 2H), 4.70 (s, 1H), 4.30 – 4.26 (s, 4H), 3.92 – 3.88 (m, 2H), 3.84 – 3.74 (m, 6H), 3.69 – 3.63 (m, 6H), 3.60 – 3.56 (s, 2H), 3.24 – 3.20 (s, 1H), 3.15 – 3.12 (m, 3H), 2.85 – 2.64 (m, 6H), 2.64 – 2.40 (m, 6H), 2.07 – 2.05 (s, 2H), 1.46 – 1.31 (m, 6H). $^{13}$C NMR (151 MHz, $CDCl_3$) δ 171.55, 168.63, 167.01, 165.78, 156.34, 136.66, 133.65, 128.70, 127.75, 127.33, 119.53, 119.51, 119.48, 117.19, 117.15, 116.14, 116.12, 70.95, 70.87, 70.42, 70.37, 69.25, 69.16, 67.34, 67.17, 59.16, 55.31, 49.10, 43.32, 35.15, 34.46, 31.33, 31.31, 29.69, 22.59, 22.57, 22.55, 18.49, 17.04, 12.62. (*Note: rotameric mixture observed*) HR-MS (ESI$^+$) calc'd for $C_{38}H_{40}N_4O_8$: 680.2846, found $[M+H]^+$ 681.2841.

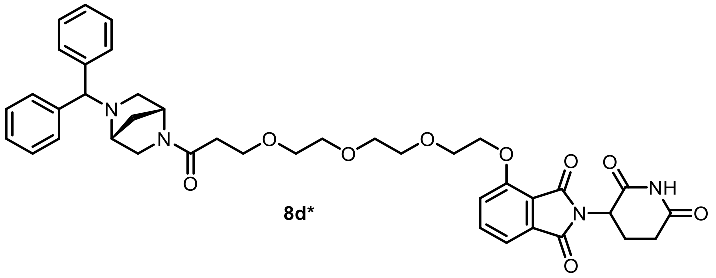

**8d***

**4-(2-(2-(2-(3-((1R,4R)-5-benzhydryl-2,5-diazabicyclo[2.2.1]heptan-2-yl)-3-oxopropoxy)ethoxy)ethoxy)ethoxy)-2-(2,6-dioxopiperidin-3-yl)isoindoline-1,3-dione (8d*)**

**General Procedure 1A.** Reaction scale: Thalidomide-O-PEG3-acid 15.2 mg (32.7 µmol, 1.0 eq) and (1R,4R)-2-benzhydryl-2,5-diazabicyclo[2.2.1]heptane 15 mg (32.7 µmol, 1.0 eq). Purified by pTLC (7% MeOH/DCM) to afford **8d*** as an off-white solid (9.0 mg, 34%). $^1$H NMR (600 MHz, CDCl3) δ 8.98 (s, 1H), 7.72 – 7.60 (m, 1H), 7.48 – 7.45 (m, 4H), 7.30 – 7.25 (m, 5H), 7.24 – 7.05 (m, 4H), 4.96 – 4.92 (m, 1H), 4.58 – 4.54 (m, 1H), 4.35 – 4.30 (m, 3H), 3.93 – 3.88 (m, 3H), 3.82 – 3.58 (m, 13H), 3.46 (s, 1H), 3.08 – 3.04 (m, 1H), 2.89 – 2.70 (m, 4H), 2.66 – 2.40 (m, 3H), 2.12 – 2.09 (m, 1H), 1.99 (s, 1H), 1.64 –1.62 (m, 1H). $^{13}$C NMR (151 MHz, $CDCl_3$) δ $^{13}$C NMR (151 MHz, $CDCl_3$) δ 171.66, 168.65, 167.02, 156.25, 136.74, 133.60, 128.69, 127.76, 127.37, 119.50, 117.36, 116.27, 70.86, 70.14, 69.15, 69.05, 67.12, 54.86, 49.18, 34.87, 31.39, 29.74, 22.61, 22.54, 18.59, 17.21. (*Note: rotameric mixture observed*) HR-MS (ESI$^+$) calc'd for $C_{40}H_{44}N_4O_9$: 724.3108, found $[M+H]^+$ 724.3108.

.

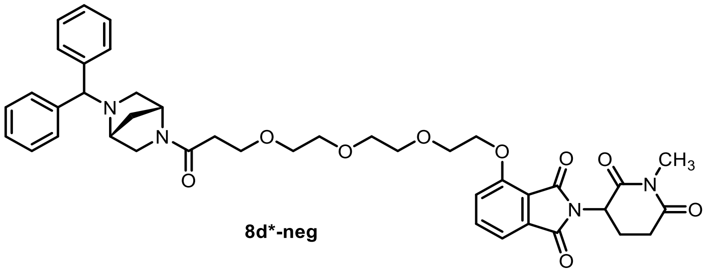

**8d*-neg**

**4-(2-(2-(2-(3-((1R,4R)-5-benzhydryl-2,5-diazabicyclo[2.2.1]heptan-2-yl)-3-oxopropoxy)ethoxy)ethoxy)ethoxy)-2-(1-methyl-2,6-dioxopiperidin-3-yl)isoindoline-1,3-dione (8d*-neg)**

**General Procedure 4**. Reaction scale: bifunctional molecule **8d*** 14.5 mg (20.0 µmol,1 eq) and methyl iodide 7 mg (50.0 µmol, 2.5 eq). Purified by pTLC (6% MeOH/DCM) to afford **8d*-neg** as a sticky solid (8.0 mg, 54% ). $^1$H NMR (600 MHz, $CDCl_3$) δ 7.68 – 7.95 (m, 1H), 7.58 – 7.37 (m, 5H), 7.37 – 7.20 (m, 5H), 7.18 – 7.15 (m, 2H), 5.02 – 4.92 (m, 1H), 4.72 (s, 1H), 4.62 (s, 1H), 4.57 (s, 1H), 4.40 – 4.29 (m, 2H), 3.96 – 3.92 (m, 2H), 3.88 – 3.81 (m, 1H), 3.81 – 3.76 (m, 2H), 3.76 – 3.74 (m, 1H), 3.71 – 3.72 (m, 3H), 3.65 – 3.56 (m, 3H), 3.50 – 3.46 (m, 1H), 3.27 – 3.09 (m, 4H), 3.01 – 2.89 (m, 2H), 2.83 – 2.69 (m, 2H), 2.67 – 2.43 (m, 3H), 2.16 – 2.02 (m, 3H), 1.73 – 1.61 (m, 2H). $^{13}$C NMR (151 MHz, $CDCl_3$) δ 171.24, 168.79, 168.73, 168.52, 167.16, 165.79, 156.40, 156.38, 143.98, 143.90, 143.13, 143.11, 136.46, 136.44, 133.79, 128.66, 128.55, 127.72, 127.65, 127.29, 127.22, 119.51, 119.41, 117.36, 116.15, 116.08, 72.74, 71.91, 71.87, 71.10, 70.56, 70.43, 69.35, 69.21, 67.36, 67.27, 59.20, 59.12, 59.06, 58.03, 56.30, 56.20, 51.14, 49.93, 49.78, 49.05, 36.37, 35.17, 34.51, 31.89, 27.30, 27.21, 21.92. (*Note: rotameric mixture observed*) HR-MS (ESI+) calc'd for $C_{41}H_{46}N_4O_9$: 738.3265, found [M+H]$^+$ 739.3343.

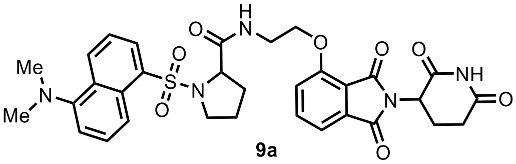

**9a**

**1-((5-(dimethylamino)naphthalen-1-yl)sulfonyl)-N-(2-((2-(2,6-dioxopiperidin-3-yl)-1,3-dioxoisoindolin-4-yl)oxy)ethyl)pyrrolidine-2-carboxamide (9a)**

**General Procedure 1B**. Reaction scale: ((5-(dimethylamino)naphthalen-1-yl)sulfonyl)proline (6.6 mg, 20 µmol, 1.0 eq.) was reacted with thalidomide-O-C2-amine (TFA salt, 9.1 mg, 21 µmol, 1.05 eq.). Purified by preparative HPLC (MeCN/$H_2O$ gradient, 0.1% FA) to afford **9a** as a white solid (9.5 mg, 14.7 µmol, 73%). $^1$H NMR (600 MHz, $CD_3OD$) (rotamers present) δ 8.58 – 8.52 (m, 1H), 8.48 – 8.43 (m, 1H), 8.24 – 8.17 (m, 1H), 7.76 (ddd, J = 8.4, 7.2, 1.0 Hz, 1H), 7.63 – 7.50 (m, 2H), 7.47 (dd, J = 7.2, 2.4 Hz, 1H), 7.40 – 7.33 (m, 1H), 7.23 – 7.16 (m, 1H), 5.19 – 5.05 (m, 1H), 4.34 – 4.24 (m, 1H), 4.16 – 3.97 (m, 2H), 3.65 – 3.57 (m, 1H), 3.57 – 3.42 (m, 2H), 3.37 – 3.32 (m, 1H), 2.92 – 2.82 (m, 1H), 2.85 (s, 6H), 2.79 – 2.63 (m, 2H), 2.17 – 2.07 (m, 1H), 2.01 – 1.82 (m, 3H), 1.77 – 1.68 (m, 1H). $^{13}$C NMR (151 MHz, $CD_3OD$) δ 174.69, 174.66, 171.40, 168.49 & 168.46 (rot.), 167.72, 157.40, 153.18, 138.08, 135.01, 134.76 & 134.68 (rot.), 132.06 & 132.04 (rot.), 131.76 & 131.74 (rot.), 131.60 & 131.48 (rot.), 131.25, 129.57, 124.47 & 124.44 (rot.), 121.13 & 121.01 (rot.), 120.56 & 120.54 (rot.), 118.63 & 118.58 (rot.), 117.08 & 117.01 (rot.), 116.52, 68.84 & 68.74 (rot.), 63.27 & 63.25 (rot.), 50.45, 50.40 & 50.39 (rot.), 45.75, 39.51, 32.57 & 32.54 (rot.), 32.15, 25.67 & 25.65 (rot.), 23.68 & 23.66 (rot.). HR-MS (ESI+) calc'd for [M+H]$^+$ $C_{32}H_{34}N_5O_8S$: 648.2123, found 648.2121 (0.4 ppm).

.

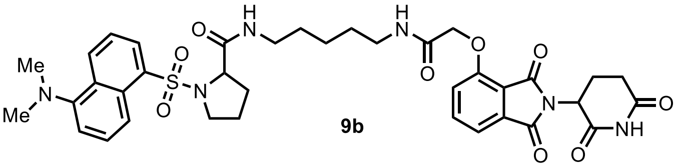

9b

**1-((5-(dimethylamino)naphthalen-1-yl)sulfonyl)-N-(5-(2-((2-(2,6-dioxopiperidin-3-yl)-1,3-dioxoisoindolin-4-yl)oxy)acetamido)pentyl)pyrrolidine-2-carboxamide (9b)**

**General Procedure 1B**. Reaction scale: ((5-(dimethylamino)naphthalen-1-yl)sulfonyl)proline (6.6 mg, 20 µmol, 1.0 eq.) was reacted with thalidomide-O-acetamido-C5-amine (TFA salt, 11.1 mg, 21 µmol, 1.05 eq.). Purified by preparative HPLC (MeCN/$H_2O$ gradient, 0.1% FA) to afford **9b** as a white solid (11.2 mg, 15.0 µmol, 75%). $^1H$ NMR (600 MHz, $CD_3OD$) δ 8.59 – 8.54 (m, 1H), 8.42 (dd, *J* = 8.7, 2.6 Hz, 1H), 8.19 – 8.08 (m, 1H), 7.64 – 7.48 (m, 3H), 7.35 – 7.20 (m, 3H), 5.12 (dd, *J* = 12.8, 5.5 Hz, 1H), 4.72 (dd, *J* = 14.4, 2.5 Hz, 1H), 4.68 (dd, *J* = 14.3, 2.5 Hz, 1H), 4.20 (dd, *J* = 8.2, 4.0 Hz, 1H), 3.58 – 3.45 (m, 1H), 3.43 – 3.26 (m, 3H), 3.18 – 3.01 (m, 2H), 2.88 (s, 6H), 2.93 – 2.83 (m, 1H), 2.81 – 2.65 (m, 2H), 2.22 – 2.10 (m, 1H), 2.00 – 1.76 (m, 3H), 1.73 – 1.51 (m, 3H), 1.49 – 1.29 (m, 4H). $^{13}C$ NMR (151 MHz, $CD_3OD$) δ 174.64, 174.27, 171.41, 169.94, 168.26, 167.71, 155.98, 153.23, 137.82, 134.64, 134.30 & 134.26 (rot.), 132.03, 131.75 & 131.63 (rot.), 131.56, 131.32, 129.57, 124.50, 121.38, 120.50, 119.21, 117.57, 116.62, 69.19, 63.31, 50.50, 50.28, 45.79, 40.20, 39.97, 32.30, 32.16, 29.94, 29.84, 25.61, 24.95, 23.68. HR-MS (ESI+) calc'd for $[M+H]^+$ $C_{37}H_{43}N_6O_9S$: 747.2807, found 747.2801 (0.9 ppm).

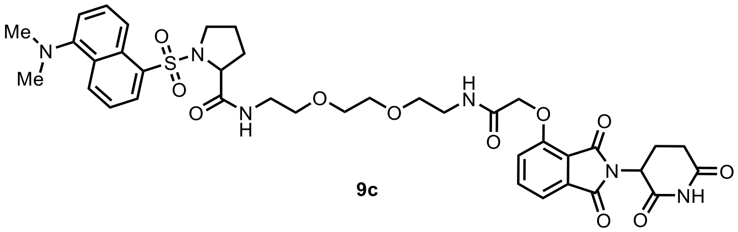

9c

**1-((5-(dimethylamino)naphthalen-1-yl)sulfonyl)-N-(2-(2-(2-(2-((2-(2,6-dioxopiperidin-3-yl)-1,3-dioxoisoindolin-4-yl)oxy)acetamido)ethoxy)ethoxy)ethyl)pyrrolidine-2-carboxamide (9c)**

**General Procedure 1B**. Reaction scale: ((5-(dimethylamino)naphthalen-1-yl)sulfonyl)proline (6.6 mg, 20 µmol, 1.0 eq.) was reacted with thalidomide-O-acetamido-PEG2-amine (HCl salt, 10.5 mg, 21 µmol, 1.05 eq.). The product was purified by preparative HPLC (MeCN/$H_2O$ gradient, 0.1% FA) followed by preparative TLC (7% MeOH/DCM) to afford **9c** as a white solid (8.2 mg, 10.3 µmol, 52%). $^1H$ NMR (400 MHz, $CDCl_3$) δ 9.13 – 8.86 (m, 1H), 8.56 (dd, *J* = 8.5, 3.2 Hz, 1H), 8.52 (dd, *J* = 8.7, 2.6 Hz, 1H), 8.31 – 8.11 (m, 1H), 7.70 (t, *J* = 7.9 Hz, 1H), 7.65 (br s, 1H), 7.62 – 7.49 (m, 3H), 7.25 – 7.21 (m, 1H), 7.21 – 7.08 (m, 2H), 4.97 (dt, *J* = 12.1, 4.9 Hz, 1H), 4.64 (s, 2H), 4.35 – 4.27 (m, 1H), 3.79 – 3.24 (m, 14H), 2.88 (d, *J* = 1.2 Hz, 6H), 2.95 – 2.70 (m, 3H), 2.20 – 2.07 (m, 2H), 1.85 – 1.71 (m, 1H), 1.68 – 1.58 (m, 2H). *Note: shifted $H_2O$ peak at 5.30 ppm integrating to 2H.* $^{13}C$ NMR (151 MHz, $CDCl_3$) δ 171.93 & 171.86 (rot.), 171.56 & 171.47 (rot.), 168.47 & 168.37 (rot.), 166.99 & 166.97 (rot.), 166.83, 165.95 & 165.92 (rot.), 154.55 & 154.54 (rot.), 151.98, 137.01, 133.81, 132.65 & 132.55 (rot.), 131.23 & 131.20 (rot.), 130.98 & 130.89 (rot.), 130.75 & 130.73 (rot.), 130.17, 128.78, 123.47 & 123.46 (rot.), 119.39 & 119.34 (rot.), 119.29, 118.17, 117.35 & 117.34 (rot.), 115.51, 70.34 & 70.30 (rot.), 70.21 & 70.16 (rot.), 69.78 & 69.67 (rot.), 69.63, 68.02 & 67.99 (rot.), 62.40 & 62.34 (rot.), 53.58, 49.41, 45.54, 39.41 & 39.37 (rot.), 39.26 & 39.20 (rot.), 31.56 & 31.53 (rot.), 30.79 & 30.72 (rot.), 24.62 & 24.58 (rot.), 22.84. HR-MS (ESI+) calc'd for $[M+H]^+$ $C_{38}H_{45}N_6O_{11}S$: 793.2862, found 793.2846 (2.1 ppm).

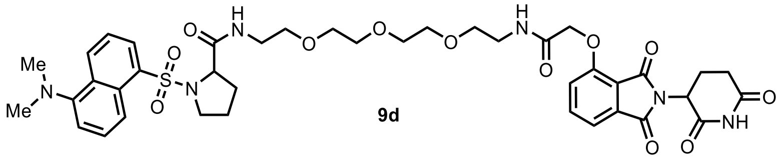

**9d**

**1-((5-(dimethylamino)naphthalen-1-yl)sulfonyl)-N-(1-((2-(2,6-dioxopiperidin-3-yl)-1,3-dioxoisoindolin-4-yl)oxy)-2-oxo-6,9,12-trioxa-3-azatetradecan-14-yl)pyrrolidine-2-carboxamide (9d)**

**General Procedure 1B**. Reaction scale: ((5-(dimethylamino)naphthalen-1-yl)sulfonyl)proline (6.6 mg, 20 µmol, 1.0 eq.) was reacted with thalidomide-O-acetamido-PEG3-amine (HCl salt, 11.4 mg, 21 µmol, 1.05 eq.). Purified by preparative HPLC (MeCN/$H_2O$ gradient, 0.1% FA) to afford **9d** as a white solid (11.0 mg, 13.1 µmol, 66%). $^1$H NMR (600 MHz, $CD_3OD$) δ 8.59 (d, *J* = 8.5 Hz, 1H), 8.47 (d, *J* = 8.7 Hz, 1H), 8.20 (dd, *J* = 7.3, 1.3 Hz, 1H), 7.74 (t, *J* = 7.9 Hz, 1H), 7.59 (t, *J* = 8.0 Hz, 2H), 7.48 (d, *J* = 7.3 Hz, 1H), 7.36 (d, *J* = 8.5 Hz, 1H), 7.27 (d, *J* = 7.5 Hz, 1H), 5.12 (dd, *J* = 12.9, 5.5 Hz, 1H), 4.73 (s, 2H), 4.32 – 4.23 (m, 1H), 3.70 – 3.59 (m, 8H), 3.58 – 3.47 (m, 5H), 3.46 – 3.35 (m, 3H), 3.29 – 3.22 (m, 1H), 3.21 – 3.12 (m, 1H), 2.87 (s, 6H), 2.91 – 2.82 (m, 1H), 2.80 – 2.67 (m, 2H), 2.19 – 2.09 (m, 1H), 1.97 – 1.81 (m, 3H), 1.76 – 1.63 (m, 1H). $^{13}$C NMR (151 MHz, $CD_3OD$) δ 174.61, 174.38, 171.35, 169.97, 168.30, 167.55, 156.09, 153.23, 138.13, 134.90, 134.63, 132.03, 131.78, 131.50, 131.33, 129.60, 124.50, 121.44, 120.61, 119.15, 117.83, 116.62, 71.60, 71.57, 71.43, 71.34, 70.30, 70.24, 69.14, 63.15, 50.53, 50.32, 45.79, 40.28, 40.22, 32.37, 32.18, 25.65, 23.67. HR-MS (ESI+) calc'd for $[M+H]^+$ $C_{40}H_{49}N_6O_{12}S$: 837.3124, found 837.3110 (1.7 ppm).

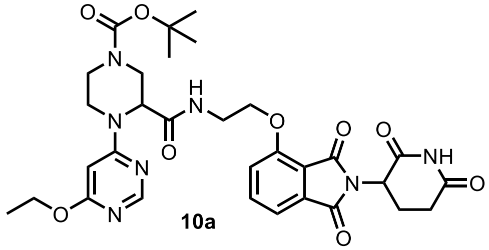

**10a**

***tert*-butyl 3-((2-((2-(2,6-dioxopiperidin-3-yl)-1,3-dioxoisoindolin-4-yl)oxy)ethyl)carbamoyl)-4-(6-ethoxypyrimidin-4-yl)piperazine-1-carboxylate (10a)**

**General Procedure 1B**. Reaction scale: 4-(*tert*-butoxycarbonyl)-1-(6-ethoxypyrimidin-4-yl)piperazine-2-carboxylic acid (7.0 mg, 20 µmol, 1.0 eq.) was reacted with thalidomide-O-C2-amine (TFA salt, 9.1 mg, 21 µmol, 1.05 eq.). Purified by preparative HPLC (MeCN/$H_2O$ gradient, 0.1% FA) to afford **10a** as a white solid (10.7 mg, 16.4 µmol, 82%). $^1$H NMR (600 MHz, $CD_3OD$) *(rotamers present)* δ 8.15 – 8.03 (br m, 1H), 7.86 – 7.70 (m, 1H), 7.47 (t, *J* = 7.6 Hz, 1H), 7.44 – 7.31 (m, 1H), 6.09 – 5.85 (m, 1H), 5.18 – 5.05 (m, 1H), 4.42 (br s, 1H), 4.21 (m, 4H), 3.88 – 3.75 (m, 2H), 3.76 – 3.57 (m, 3H), 3.56 – 3.31 (m, 3H), 2.89 (ddd, *J* = 17.6, 13.9, 5.4 Hz, 1H), 2.85 – 2.66 (m, 2H), 2.26 – 2.10 (m, 1H), 1.50 – 1.25 (m, 12H). $^{13}$C NMR (151 MHz, $CD_3OD$) *(mostly rotameric multiplets, major intensity is reported)* δ 174.63, 173.44, 171.40, 168.44, 168.16, 168.00, 165.54, 158.11, 157.45, 156.39, 138.20, 134.97, 120.73, 118.27, 117.03, 87.18, 81.58, 69.25, 63.58, 58.43, 50.60, 44.83, 43.89, 42.12, 39.77, 32.19, 28.55, 23.72, 14.80. HR-MS (ESI+) calc'd for $[M+H]^+$ $C_{31}H_{38}N_7O_9$: 652.2726, found 652.2713 (2.0 ppm).

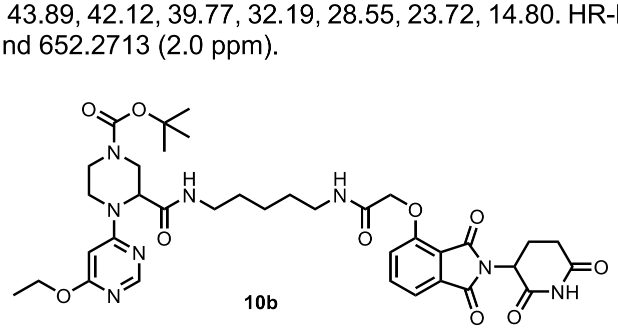

**10b**

**_tert_-butyl 3-((5-(2-((2-(2,6-dioxopiperidin-3-yl)-1,3-dioxoisoindolin-4-yl)oxy)acetamido)pentyl)carbamoyl)-4-(6-ethoxypyrimidin-4-yl)piperazine-1-carboxylate (10b)**

**General Procedure 1B**. Reaction scale: 4-(*tert*-butoxycarbonyl)-1-(6-ethoxypyrimidin-4-yl)piperazine-2-carboxylic acid (7.0 mg, 20 µmol, 1.0 eq.) was reacted with thalidomide-O-acetamido-C5-amine (TFA salt, 11.1 mg, 21 µmol, 1.05 eq.). Purified by preparative HPLC (MeCN/$H_2O$ gradient, 0.1% FA) to afford **10b** as a white solid (10.8 mg, 14.4 µmol, 72%). $^1H$ NMR (400 MHz, $CD_3OD$) δ 8.19 (br s, 1H), 7.81 (t, *J* = 7.9 Hz, 1H), 7.54 (d, *J* = 7.3 Hz, 1H), 7.43 (d, *J* = 8.4 Hz, 1H), 5.96 (br s, 1H), 5.23 – 5.09 (m, 1H), 4.77 (s, 2H), 4.40 (br s, 1H), 4.28 (q, *J* = 7.0 Hz, 2H), 3.93 – 3.76 (m, 2H), 3.71 – 3.58 (m, 1H), 3.52 – 3.31 (m, 2H), 3.31 – 3.09 (m, 4H), 2.98 – 2.83 (m, 1H), 2.82 – 2.61 (m, 2H), 2.25 – 2.07 (m, 1H), 1.67 – 1.22 (m, 18H). $^{13}C$ NMR (151 MHz, $CD_3OD$) δ 174.58, 172.83, 171.54, 171.40, 169.91, 168.30, 167.79, 165.78, 158.15, 156.31, 156.27, 138.23, 134.92, 121.77, 119.33, 117.96, 87.05, 81.57, 69.45, 63.58, 57.63 & 57.56 (rot.), 50.55, 45.43 (br), 44.17 (br), 42.25 (br), 40.47 (br), 40.02, 32.17, 30.09 (br multiplet), 29.87, 28.64, 25.10, 23.65, 14.86. HR-MS (ESI+) calc'd for $[M+H]^+$ $C_{36}H_{46}N_8O_{10}$: 751.3410, found 751.3395 (2.0 ppm).

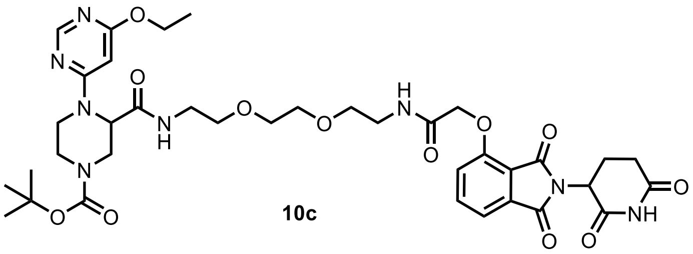

**_tert_-butyl 3-((2-(2-(2-(2-((2-(2,6-dioxopiperidin-3-yl)-1,3-dioxoisoindolin-4-yl)oxy)acetamido)ethoxy)ethoxy)ethyl)carbamoyl)-4-(6-ethoxypyrimidin-4-yl)piperazine-1-carboxylate (10c)**

**General Procedure 1B.** Reaction scale: 4-(*tert*-butoxycarbonyl)-1-(6-ethoxypyrimidin-4-yl)piperazine-2-carboxylic acid (7.0 mg, 20 µmol, 1.0 eq.) was reacted with thalidomide-O-acetamido-PEG2-amine (HCl salt, 10.5 mg, 21 µmol, 1.05 eq.). The product was purified by preparative HPLC (MeCN/$H_2O$ gradient, 0.1% FA) followed by preparative TLC (7% MeOH/DCM) to afford **10c** as a white solid (8.0 mg, 10.0 µmol, 50%). $^1H$ NMR (400 MHz, $CDCl_3$) δ 10.02 – 8.93 (m, 1H), 8.30 (s, 1H), 7.81 – 7.71 (m, 1H), 7.65 (br s, 1H), 7.55 (d, *J* = 7.3 Hz, 1H), 7.19 (dd, *J* = 8.4, 2.6 Hz, 1H), 7.08 – 6.80 (m, 1H), 5.81 (s, 1H), 4.95 (s, 1H), 4.64 (s, 2H), 4.60 – 4.45 (m, 1H), 4.40 – 4.28 (m, 2H), 3.99 – 3.72 (m, 2H), 3.70 – 3.15 (m, 16H), 2.88 – 2.74 (m, 3H), 2.28 – 2.11 (m, 1H), 1.45 (s, 9H), 1.39 – 1.30 (m, 3H). $^{13}C$ NMR (151 MHz, $CDCl_3$) δ *(3 signals at coalescence or overlapping in 173-160 ppm region)* 170.33, 166.85, 168.84, 166.65, 165.89, 157.59 (br),154.75, 154.44, 137.03, 133.68, 119.32, 118.08, 117.35, 86.44, 80.52 (br), 69.98, 69.92, 69.74, 69.50, 67.88, 62.30 (br), 56.03 (br), 49.28, 49.23, 43.47, 41.55 (br), 39.49, 39.08, 31.41, 28.35, 22.72, 14.62. HR-MS (ESI+) calc'd for $[M+H]^+$ $C_{37}H_{49}N_8O_{12}$: 797.3465, found 797.3480 (1.9 ppm).

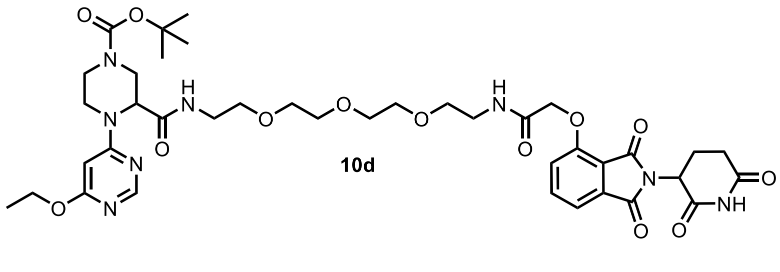

**_tert_-butyl 3-((1-((2-(2,6-dioxopiperidin-3-yl)-1,3-dioxoisoindolin-4-yl)oxy)-2-oxo-6,9,12-trioxa-3-azatetradecan-14-yl)carbamoyl)-4-(6-ethoxypyrimidin-4-yl)piperazine-1-carboxylate (10d)**

**General Procedure 1B**. Reaction scale: 4-(*tert*-butoxycarbonyl)-1-(6-ethoxypyrimidin-4-yl)piperazine-2-carboxylic acid (7.0 mg, 20 µmol, 1.0 eq.) was reacted with thalidomide-O-acetamido-PEG3-amine (HCl salt, 11.4 mg, 21 μmol, 1.05 eq.). The product was purified by preparative HPLC (MeCN/$H_2O$ gradient, 0.1% FA) followed by preparative TLC (7% MeOH/DCM) to afford **10d** as a white solid (7.6 mg, 9.0 μmol, 45%). $^1$H NMR (400 MHz, $CDCl_3$) δ 9.96 – 9.04 (m, 1H), 8.40 – 8.14 (m, 1H), 7.74 (t, *J* = 7.7 Hz, 1H), 7.70 (s, 1H), 7.55 (d, *J* = 7.3 Hz, 1H), 7.19 (d, *J* = 8.4 Hz, 1H), 7.13 – 6.95 (m, 1H), 5.80 (s, 1H), 4.95 (br s, 1H), 4.64 (s, 2H), 4.55 – 4.40 (m, 1H), 4.39 – 4.28 (m, 2H), 3.91 – 3.73 (m, 2H), 3.74 – 3.47 (m, 16H), 3.47 – 3.22 (m, 4H), 2.96 – 2.66 (m, 3H), 2.17 – 2.09 (m, 1H), 1.45 (s, 9H), 1.35 (t, *J* = 7.1 Hz, 3H). $^{13}$C NMR (151 MHz, $CDCl_3$) δ 171.74 (br), 170.42 (br), 170.12 (br), 168.46 (br), 166.85, 166.77, 165.93, 164.02 (br), 157.61 (br), 154.89 (br), 154.50, 137.12, 133.83, 119.29, 118.15, 117.43, 86.57 (br), 80.47, 70.49, 70.47, 70.38, 70.30, 70.24, 69.58, 67.87, 62.41 (br), 56.38 (br), 49.40, 43.72 (br), 43.14 (br), 41.61 (br), 39.43, 39.22, 31.55, 28.50, 22.86, 14.76. HR-MS (ESI+) calc'd for $[M+H]^+$ $C_{39}H_{53}N_8O_{13}$: 841.3727, found 841.3711 (2.0 ppm).

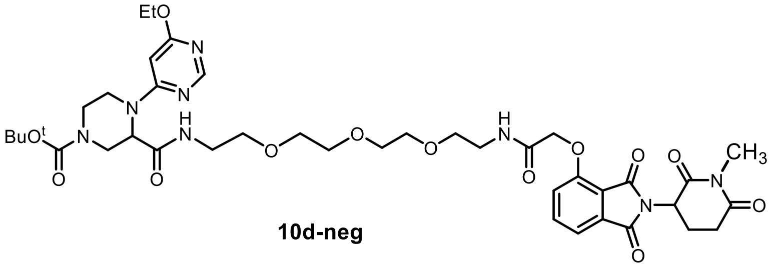

**10d-neg**

**tert-butyl 4-(6-ethoxypyrimidin-4-yl)-3-((1-((2-(1-methyl-2,6-dioxopiperidin-3-yl)-1,3-dioxoisoindolin-4-yl)oxy)-2-oxo-6,9,12-trioxa-3-azatetradecan-14-yl)carbamoyl)piperazine-1-carboxylate (10d-neg)**

**General Procedure 4.** Reaction scale: bifunctional molecule **10a** 8 mg (0.09 μmol, 1 eq) and methyl iodide 3.0 mg (0.18 μmol, 2.0 eq). Purified by pTLC (6% MeOH/DCM) to afford **10d-neg** as a sticky solid (3.0 mg, 42%). $^1$H NMR (600 MHz, $CDCl_3$) δ 8.29 (s, 1H), 7.73 (d, *J* = 7.9 Hz, 1H), 7.60 (s, 1H), 7.55 (d, *J* = 7.3 Hz, 1H), 7.20 (d, *J* = 8.4 Hz, 1H), 7.03 (s, 1H), 6.81 (s, 1H), 5.82 (s, 1H), 4.98 – 4.95 (m, 2H), 4.66 (s, 1H), 4.48 (s, 1H), 4.33 (d, *J* = 7.1 Hz, 2H), 3.85 – 3.75 (m, 2H), 3.69 – 3.45 (m, 6H), 3.35 (s, 1H), 3.20 (s, 1H), 3.04 – 2.98 (m, 2H), 2.83 – 2.78 (m, 2H), 2.10 (d, *J* = 7.7 Hz, 1H), 1.65 (s, 3H), 1.50 – 1.52 (m, 4H), 1.45( s, 9H), 1.25 (s, 3H). $^{13}$C NMR (151 MHz, $CDCl_3$) δ 171.13, 170.35, 170.00, 168.68, 167.11, 166.80, 166.01, 157.47, 154.75, 154.51, 136.99, 133.68, 119.51, 118.08, 117.35, 86.45, 80.31, 70.43, 70.26, 69.89, 69.56, 68.08, 62.26, 55.93, 50.03, 41.32, 39.40, 39.02, 31.95, 31.89, 29.72, 28.37, 27.30, 22.72, 21.94, 14.63, 14.15. HR-MS (ESI+) calc'd for $C_{40}H_{54}N_8O_{13}$: 854.9150, found $[M+H]^+$ 855.3881.

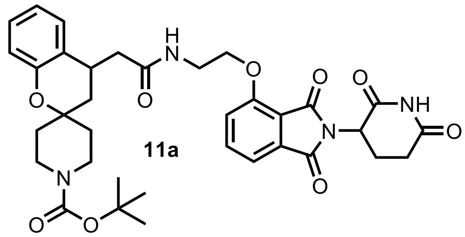

**11a**

***tert*-butyl 4-(2-((2-((2-(2,6-dioxopiperidin-3-yl)-1,3-dioxoisoindolin-4-yl)oxy)ethyl)amino)-2-oxoethyl)spiro[chromane-2,4'-piperidine]-1'-carboxylate (11a)**

**General Procedure 1B**. Reaction scale: 2-(1'-(*tert*-butoxycarbonyl)spiro[chromane-2,4'-piperidin]-4-yl)acetic acid (7.2 mg, 20 μmol, 1.0 eq.) was reacted with thalidomide-O-C2-amine (TFA salt, 9.1 mg, 21 μmol, 1.05 eq.). Purified by preparative HPLC (MeCN/$H_2O$ gradient, 0.1% FA) to afford **11a** as a white solid (10.2 mg, 15.4 μmol, 77%). $^1$H NMR (600 MHz, $CD_3OD$) (2 H's expected in water peak region) δ 7.80 (t, *J* = 7.8 Hz, 1H), 7.51 – 7.46 (m, 2H), 7.25 – 7.20 (m, 1H), 7.08 – 7.02 (m, 1H), 6.83 – 6.75 (m, 2H), 5.11 (dd, *J* = 12.9, 5.4 Hz, 1H), 4.41 – 4.24 (m, 2H), 3.84 – 3.60 (m, 4H), 3.40 – 3.32 (m, 2H), 3.02 (br s, 1H), 2.96 (ddd, *J* = 13.8, 5.0, 2.4 Hz, 1H), 2.92 – 2.82 (m, 1H), 2.78 – 2.62 (m, 2H), 2.21 (ddd, *J* = 13.3, 9.8, 2.9 Hz, 1H), 2.15 – 2.06 (m, 1H), 1.87 (ddd, *J* = 13.6, 6.3, 2.0 Hz, 1H), 1.74 (d, *J* = 13.8 Hz, 1H), 1.68 – 1.61 (m, 1H), 1.54 (ddd, *J* = 13.5, 11.7, 6.6 Hz, 1H), 1.45 (s, 9H), 1.35 (td, *J* = 12.8, 4.7 Hz, 1H). $^{13}$C NMR (151 MHz,

$CD_3OD$) δ 174.87 & 174.85 (rot.), 174.60, 171.30 & 171.26 (rot.), 168.39, 167.88, 157.65, 156.50, 154.06, 138.23, 135.12, 128.82, 128.12, 125.65, 121.65, 120.83, 118.57, 118.39, 117.03, 81.03, 73.88, 69.14, 50.47, 49.57, 42.88, 41.22 (br), 39.85, 39.64, 38.33 (br.), 32.68 (br.), 32.15, 29.59, 28.70, 23.66. HR-MS (ESI+) calc'd for $[M+H]^+$ $C_{35}H_{41}N_4O_9$: 661.2868, found 661.2863 (0.8 ppm).

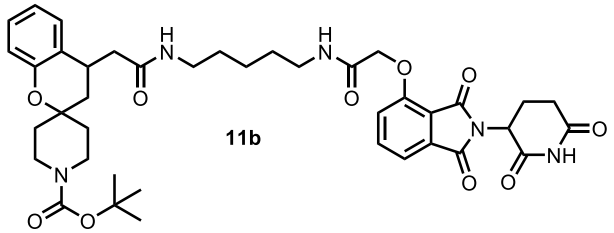

**11b**

***tert*-butyl 4-(2-((5-(2-((2-(2,6-dioxopiperidin-3-yl)-1,3-dioxoisoindolin-4-yl)oxy)acetamido)pentyl)amino)-2-oxoethyl)spiro[chromane-2,4'-piperidine]-1'-carboxylate (11b)**

**General Procedure 1B**. Reaction scale: 2-(1'-(*tert*-butoxycarbonyl)spiro[chromane-2,4'-piperidin]-4-yl)acetic acid (7.2 mg, 20 µmol, 1.0 eq.) was reacted with thalidomide-O-acetamido-C5-amine (TFA salt, 11.1 mg, 21 µmol, 1.05 eq.). Purified by preparative HPLC (MeCN/$H_2O$ gradient, 0.1% FA) to afford **11b** as a white solid (9.8 mg, 12.9 µmol, 64%). $^1H$ NMR (600 MHz, $CD_3OD$) δ 7.81 (t, *J* = 7.9 Hz, 1H), 7.54 (d, *J* = 7.3 Hz, 1H), 7.43 (d, *J* = 8.5 Hz, 1H), 7.22 (dd, *J* = 7.8, 1.6 Hz, 1H), 7.08 (t, *J* = 7.6 Hz, 1H), 6.85 (t, *J* = 7.5 Hz, 1H), 6.80 (dd, *J* = 8.1, 1.3 Hz, 1H), 5.19 – 5.12 (m, 1H), 4.76 (s, 2H), 3.86 (d, *J* = 13.4 Hz, 1H), 3.76 (d, *J* = 13.4 Hz, 1H), 3.44 – 3.27 (m, 4H), 3.27 – 3.18 (m, 2H), 3.06 (br s, 1H), 2.97 – 2.84 (m, 2H), 2.80 – 2.68 (m, 2H), 2.19 (dd, *J* = 14.1, 9.2 Hz, 1H), 2.16 – 2.11 (m, 1H), 1.93 (ddd, *J* = 13.5, 6.3, 1.7 Hz, 1H), 1.79 (d, *J* = 13.9 Hz, 1H), 1.76 – 1.70 (m, 1H), 1.69 – 1.51 (m, 6H), 1.45 (s, 9H), 1.51 – 1.39 (m, 3H). $^{13}C$ NMR (151 MHz, $CD_3OD$) δ 174.59, 174.40, 171.43, 169.89, 168.28, 167.85, 156.58, 156.24, 154.08, 138.27, 134.90, 128.82, 128.22, 125.83, 121.75, 121.67, 119.33, 118.59, 118.01, 81.09, 73.93, 69.41, 50.55, 49.57, 42.90, 41.13 (br), 40.33, 40.00, 39.79, 38.36 (br), 32.74 (br), 32.17, 29.96 & 29.94 (rot.), 29.89, 29.38 & 29.37 (rot.), 28.69, 25.20 & 25.19 (rot.), 23.66. HR-MS (ESI+) calc'd for $[M+H]^+$ $C_{40}H_{50}N_5O_{10}$: 760.3552, found 760.3566 (1.9 ppm).

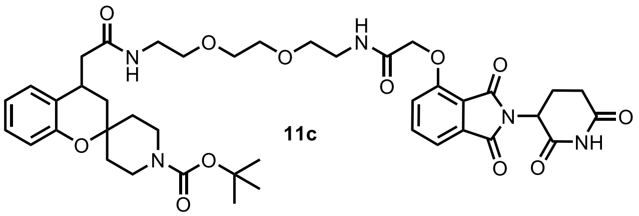

**11c**

***tert*-butyl 4-(14-((2-(2,6-dioxopiperidin-3-yl)-1,3-dioxoisoindolin-4-yl)oxy)-2,13-dioxo-6,9-dioxa-3,12-diazatetradecyl)spiro[chromane-2,4'-piperidine]-1'-carboxylate (11c)**

**General Procedure 1B**. Reaction scale: 2-(1'-(*tert*-butoxycarbonyl)spiro[chromane-2,4'-piperidin]-4-yl)acetic acid (7.2 mg, 20 µmol, 1.0 eq.) was reacted with thalidomide-O-acetamido-PEG2-amine (HCl salt, 10.5 mg, 21 µmol, 1.05 eq.). Purified by preparative HPLC (MeCN/$H_2O$ gradient, 0.1% FA) to afford **11c** as a white solid (10.8 mg, 13.4 µmol, 67%). $^1H$ NMR (600 MHz, $CD_3OD$) δ 7.79 (t, *J* = 8.1 Hz, 1H), 7.53 (d, *J* = 7.3 Hz, 1H), 7.40 (d, *J* = 8.5 Hz, 1H), 7.21 (d, *J* = 7.8 Hz, 1H), 7.07 (t, *J* = 7.6 Hz, 1H), 6.86 – 6.82 (m, 1H), 6.79 (d, *J* = 8.5 Hz, 1H), 5.16 – 5.10 (m, 1H), 4.73 (s, 2H), 3.86 (d, *J* = 13.4 Hz, 1H), 3.76 (d, *J* = 13.3 Hz, 1H), 3.68 – 3.61 (m, 6H), 3.55 (t, *J* = 5.4 Hz, 2H), 3.51 (t, *J* = 5.4 Hz, 2H), 3.38 (t, *J* = 5.4 Hz, 2H), 3.38 – 3.31 (m, 2H), 3.06 (s, 1H), 2.92 – 2.83 (m, 2H), 2.79 – 2.67 (m, 2H), 2.20 (dd, *J* = 14.1, 9.4 Hz, 1H), 2.17 – 2.11 (m, 1H), 1.94 (dd, *J* = 13.5, 6.2 Hz, 1H), 1.78 (d, *J* = 13.9 Hz, 1H), 1.71 (d, *J* = 13.7 Hz, 1H), 1.64 (td, *J* = 12.6, 4.7 Hz, 1H), 1.53 (dd, *J* = 13.6, 11.7 Hz, 1H), 1.49 – 1.42 (m, 1H), 1.45 (s, 9H). $^{13}C$ NMR (151

MHz, $CD_3OD$) δ 174.58 (2 C's), 171.35, 169.98, 168.28, 167.68, 156.56, 156.14, 154.07, 138.23, 134.93, 128.81, 128.20, 125.81, 121.69, 121.55, 119.19, 118.59, 117.92, 81.07, 73.93, 71.48, 71.32, 70.59, 70.39, 69.18, 50.54, 49.57, 42.79, 41.13 (br), 40.44, 40.16, 39.71, 38.37 (br), 32.78 (br), 32.17, 29.35, 28.69, 23.66. HR-MS (ESI+) calc'd for $[M+H]^+$ $C_{41}H_{52}N_5O_{12}$: 806.3607, found 806.3629 (2.8 ppm).

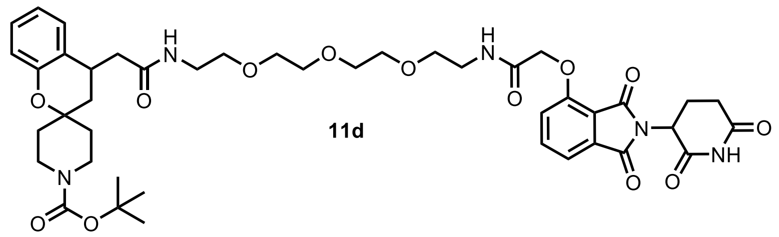

11d

**_tert_-butyl 4-(17-((2-(2,6-dioxopiperidin-3-yl)-1,3-dioxoisoindolin-4-yl)oxy)-2,16-dioxo-6,9,12-trioxa-3,15-diazaheptadecyl)spiro[chromane-2,4'-piperidine]-1'-carboxylate (11d)**

**General Procedure 1B.** Reaction scale: 2-(1'-(*tert*-butoxycarbonyl)spiro[chromane-2,4'-piperidin]-4-yl)acetic acid (7.2 mg, 20 μmol, 1.0 eq.) was reacted with thalidomide-O-acetamido-PEG3-amine (HCl salt, 11.4 mg, 21 μmol, 1.05 eq.). Purified by preparative HPLC (MeCN/$H_2O$ gradient, 0.1% FA) to afford **11d** as a white solid (11.1 mg, 13.1 μmol, 65%). $^1H$ NMR (600 MHz, $CD_3OD$) δ 7.80 (t, *J* = 7.9 Hz, 1H), 7.53 (d, *J* = 7.3 Hz, 1H), 7.41 (d, *J* = 8.5 Hz, 1H), 7.22 (d, *J* = 7.8 Hz, 1H), 7.06 (t, *J* = 7.8 Hz, 1H), 6.84 (t, *J* = 7.5 Hz, 1H), 6.81 – 6.76 (m, 1H), 5.12 (dd, *J* = 12.9, 5.2 Hz, 1H), 4.74 (s, 2H), 3.86 (d, *J* = 13.4 Hz, 1H), 3.76 (d, *J* = 13.5 Hz, 1H), 3.68 – 3.57 (m, 10H), 3.54 (t, *J* = 5.4 Hz, 2H), 3.48 (t, *J* = 5.7 Hz, 2H), 3.39 (t, *J* = 5.4 Hz, 2H), 3.37 – 3.32 (m, 2H), 3.06 (s, 1H), 2.95 – 2.83 (m, 2H), 2.81 – 2.67 (m, 2H), 2.22 (dd, *J* = 14.1, 9.3 Hz, 1H), 2.18 – 2.10 (m, 1H), 1.94 (dd, *J* = 13.5, 6.3 Hz, 1H), 1.79 (d, *J* = 13.9 Hz, 1H), 1.75 – 1.69 (m, 1H), 1.65 (td, *J* = 12.7, 4.8 Hz, 1H), 1.55 (t, *J* = 12.6 Hz, 1H), 1.50 – 1.41 (m, 1H), 1.45 (s, 9H). $^{13}C$ NMR (151 MHz, $CD_3OD$) δ 174.56 (2 Cs), 171.32, 169.92, 168.28, 167.60, 156.55, 156.14, 154.07, 138.22, 134.95, 128.81, 128.21, 125.82, 121.69, 121.53, 119.18, 118.59, 117.90, 81.06, 73.94, 71.65, 71.57, 71.38, 71.25, 70.54, 70.30, 69.15, 50.54, 49.57, 42.78, 41.03 (br), 40.40, 40.15, 39.72, 38.38 (br), 32.79 (br), 32.18, 29.37, 28.69, 23.66. HR-MS (ESI+) calc'd for $[M+H]^+$ $C_{43}H_{56}N_5O_{13}$: 850.3869, found 850.3862 (0.9 ppm).

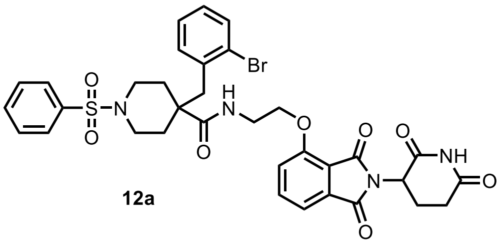

12a

**4-(2-bromobenzyl)-N-(2-((2-(2,6-dioxopiperidin-3-yl)-1,3-dioxoisoindolin-4-yl)oxy)ethyl)-1-(phenylsulfonyl)piperidine-4-carboxamide (12a)**

**General Procedure 1B**. Reaction scale: 4-(2-bromobenzyl)-1-(phenylsulfonyl)piperidine-4-carboxylic acid (8.8 mg, 20 μmol, 1.0 eq.) was reacted with thalidomide-O-C2-amine (TFA salt, 9.1 mg, 21 μmol, 1.05 eq.). Purified by preparative HPLC (MeCN/$H_2O$ gradient, 0.1% FA) to afford **12a** as a white solid (10.0 mg, 13.6 μmol, 68%). $^1H$ NMR (600 MHz, $CD_3OD$) δ 7.82 (dd, *J* = 8.5, 7.3 Hz, 1H), 7.66 – 7.61 (m, 2H), 7.54 (d, *J* = 7.2 Hz, 1H), 7.44 – 7.30 (m, 5H), 7.13 (dd, *J* = 7.7, 1.9 Hz, 1H), 7.09 (td, *J* = 7.4, 1.3 Hz, 1H), 6.96 (ddd, *J* = 7.9, 7.1, 1.9 Hz, 1H), 5.13 (dd, *J* = 12.7, 5.5 Hz, 1H), 4.03 (t, *J* = 5.1 Hz, 2H), 3.68 – 3.56 (m, 2H), 3.53 – 3.41 (m, 2H), 2.98 (s, 2H), 2.89 (ddd, *J* = 17.1, 13.7, 5.3 Hz, 1H), 2.82 – 2.66 (m, 2H), 2.35 – 2.22 (m, 4H), 2.19 – 2.12 (m, 1H), 1.77 – 1.67 (m, 2H). $^{13}C$ NMR (151 MHz, $CD_3OD$) δ 175.44, 174.63, 171.27, 168.41, 168.06, 157.47, 138.34, 137.35, 137.19, 135.03, 134.02, 133.84, 133.36, 130.04, 129.60, 128.60, 128.24, 126.61, 120.73, 118.26, 117.10, 68.97, 50.54, 47.79, 45.97, 45.06 (2 Cs), 39.95, 33.97, 33.90, 32.16, 23.72. HR-MS (ESI+) calc'd for $[M+H]^+$ $C_{34}H_{34}BrN_4O_8S$: 661.2868, found 661.2863 (0.4 ppm).

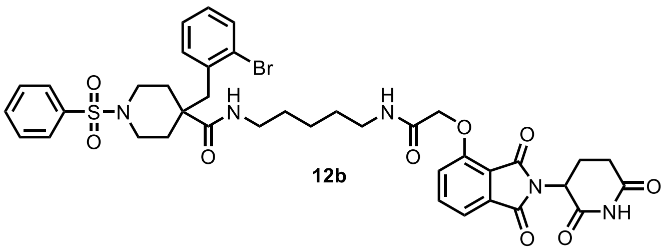

**12b**

**4-(2-bromobenzyl)-N-(5-(2-((2-(2,6-dioxopiperidin-3-yl)-1,3-dioxoisoindolin-4-yl)oxy)acetamido)pentyl)-1-(phenylsulfonyl)piperidine-4-carboxamide (12b)**

**General Procedure 1B**. Reaction scale: 4-(2-bromobenzyl)-1-(phenylsulfonyl)piperidine-4-carboxylic acid (8.8 mg, 20 µmol, 1.0 eq.) was reacted with thalidomide-O-acetamido-C5-amine (TFA salt, 11.1 mg, 21 µmol, 1.05 eq.). Purified by preparative HPLC (MeCN/$H_2O$ gradient, 0.1% FA) to afford **12b** as a white solid (7.8 mg, 9.3 µmol, 47%). $^1H$ NMR (600 MHz, $CD_3OD$) δ 7.81 (dd, *J* = 8.5, 7.3 Hz, 1H), 7.75 – 7.67 (m, 2H), 7.66 – 7.61 (m, 1H), 7.60 – 7.53 (m, 3H), 7.51 (dd, *J* = 8.4, 1.3 Hz, 1H), 7.43 (d, *J* = 8.4 Hz, 1H), 7.21 (td, *J* = 7.4, 1.3 Hz, 1H), 7.13 – 7.04 (m, 2H), 5.17 (dd, *J* = 12.8, 5.5 Hz, 1H), 4.76 (s, 2H), 3.63 – 3.53 (m, 2H), 3.29 – 3.22 (m, 2H), 3.11 – 2.98 (m, 2H), 2.93 (s, 2H), 2.87 (ddd, *J* = 18.1, 14.4, 5.4 Hz, 1H), 2.80 – 2.67 (m, 2H), 2.29 (t, *J* = 12.1 Hz, 2H), 2.22 – 2.11 (m, 3H), 1.64 (ddd, *J* = 14.9, 12.7, 4.1 Hz, 2H), 1.50 (p, *J* = 7.0 Hz, 2H), 1.31 – 1.24 (m, 2H), 1.23 – 1.15 (m, 2H). $^{13}C$ NMR (151 MHz, $CD_3OD$) δ 175.18, 174.58, 171.48, 169.88, 168.28, 167.88, 156.18, 138.31, 137.44, 136.98, 134.89, 134.23, 134.09, 133.38, 130.33, 129.67, 128.80, 128.28, 126.66, 121.77, 119.33, 118.03, 69.40, 50.55, 47.76, 45.78, 45.08, 45.06, 40.50, 39.99, 33.57 (3 Cs), 32.16, 29.88, 29.85, 25.21, 23.66. HR-MS (ESI+) calc'd for $[M+H]^+$ $C_{39}H_{43}BrN_5O_9S$: 836.1959, found 836.1974 (1.8 ppm).

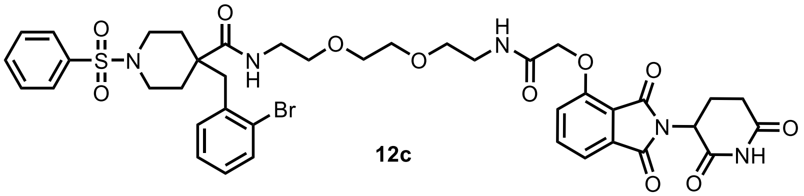

**12c**

**4-(2-bromobenzyl)-N-(2-(2-(2-(2-((2-(2,6-dioxopiperidin-3-yl)-1,3-dioxoisoindolin-4-yl)oxy)acetamido)ethoxy)ethoxy)ethyl)-1-(phenylsulfonyl)piperidine-4-carboxamide (12c)**

**General Procedure 1B**. Reaction scale: 4-(2-bromobenzyl)-1-(phenylsulfonyl)piperidine-4-carboxylic acid (8.8 mg, 20 µmol, 1.0 eq.) was reacted with thalidomide-O-acetamido-PEG2-amine (HCl salt, 10.5 mg, 21 µmol, 1.05 eq.). Purified by preparative HPLC (MeCN/$H_2O$ gradient, 0.1% FA) to afford **12c** as a white solid (14.1 mg, 16.0 µmol, 61%). $^1H$ NMR (400 MHz, $CD_3OD$) δ 7.79 (dd, *J* = 8.4, 7.3 Hz, 1H), 7.73 – 7.64 (m, 2H), 7.65 – 7.59 (m, 1H), 7.59 – 7.46 (m, 4H), 7.40 (d, *J* = 8.4 Hz, 1H), 7.21 (td, *J* = 7.4, 1.3 Hz, 1H), 7.11 (dd, *J* = 7.8, 1.8 Hz, 1H), 7.07 (td, *J* = 7.6, 1.8 Hz, 1H), 5.14 (dd, *J* = 12.6, 5.5 Hz, 1H), 4.74 (s, 2H), 3.77 – 3.52 (m, 6H), 3.53 – 3.42 (m, 4H), 3.30 – 3.28 (m, 2H), 3.21 (q, *J* = 5.3 Hz, 2H), 2.93 (s, 2H), 2.92 – 2.82 (m, 1H), 2.80 – 2.64 (m, 2H), 2.35 (td, *J* = 12.4, 2.2 Hz, 2H), 2.18 – 2.08 (m, 3H), 1.71 – 1.56 (m, 2H). $^{13}C$ NMR (151 MHz, $CD_3OD$) δ 174.05, 173.19, 169.99, 168.62, 166.89, 166.27, 154.69, 136.83, 136.12, 135.99, 133.51, 132.72, 132.65, 132.04, 128.96, 128.27, 127.32, 126.95, 125.25, 120.16, 117.79, 116.52, 70.04, 69.77, 68.91 (2 Cs), 67.77, 49.14, 46.38, 44.28, 43.56 (2 Cs), 39.11, 38.81, 32.24 (2 Cs), 30.78, 22.28. HR-MS (ESI+) calc'd for $[M+H]^+$ $C_{40}H_{45}BrN_5O_{11}S$: 882.2014, found 882.2009 (0.6 ppm).

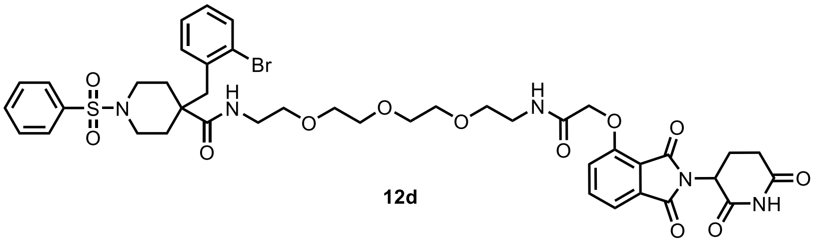

**12d**

**4-(2-bromobenzyl)-N-(1-((2-(2,6-dioxopiperidin-3-yl)-1,3-dioxoisoindolin-4-yl)oxy)-2-oxo-6,9,12-trioxa-3-azatetradecan-14-yl)-1-(phenylsulfonyl)piperidine-4-carboxamide (12d)**

**General Procedure 1B.** Reaction scale: 4-(2-bromobenzyl)-1-(phenylsulfonyl)piperidine-4-carboxylic acid (8.8 mg, 20 µmol, 1.0 eq.) was reacted with thalidomide-O-acetamido-PEG3 amine (HCl salt, 11.4 mg, 21 µmol, 1.05 eq.). Purified by preparative HPLC (MeCN/$H_2O$ gradient, 0.1% FA) to afford **12d** as a white solid (11.2 mg, 12.1 µmol, 61%). $^1$H NMR (400 MHz, $CD_3OD$) δ 7.79 (dd, *J* = 8.4, 7.3 Hz, 1H), 7.73 – 7.64 (m, 2H), 7.65 – 7.59 (m, 1H), 7.59 – 7.46 (m, 4H), 7.40 (d, *J* = 8.4 Hz, 1H), 7.21 (td, *J* = 7.4, 1.3 Hz, 1H), 7.11 (dd, *J* = 7.8, 1.8 Hz, 1H), 7.07 (td, *J* = 7.6, 1.8 Hz, 1H), 5.14 (dd, *J* = 12.6, 5.5 Hz, 1H), 4.74 (s, 2H), 3.77 – 3.52 (m, 6H), 3.53 – 3.42 (m, 4H), 3.30 – 3.28 (m, 2H), 3.21 (q, *J* = 5.3 Hz, 2H), 2.93 (s, 2H), 2.92 – 2.82 (m, 1H), 2.80 – 2.64 (m, 2H), 2.35 (td, *J* = 12.4, 2.2 Hz, 2H), 2.18 – 2.08 (m, 3H), 1.71 – 1.56 (m, 2H). $^{13}$C NMR (151 MHz, $CD_3OD$) δ 174.03, 173.18, 169.96, 168.57, 166.91, 166.20, 154.72, 136.81, 136.01 (2 Cs), 133.54, 132.72, 132.67, 132.03, 128.96, 128.28, 127.36, 126.94, 125.25, 120.11, 117.80, 116.50, 70.20, 70.16, 70.04, 69.73, 68.91, 68.85, 67.75, 49.14, 46.39, 44.33, 43.60 (2 Cs), 39.09, 38.81, 32.30, 32.28, 30.78, 22.27. HR-MS (ESI+) calc'd for $[M+H]^+$ $C_{42}H_{49}BrN_5O_{12}S$: 926.2276, found 926.2282 (0.7 ppm).

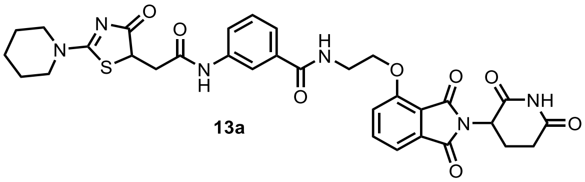

**13a**

**N-(2-((2-(2,6-dioxopiperidin-3-yl)-1,3-dioxoisoindolin-4-yl)oxy)ethyl)-3-(2-(4-oxo-2-(piperidin-1-yl)-4,5-dihydrothiazol-5-yl)acetamido)benzamide (13a)**

**General Procedure 1B**. Reaction scale: 3-(2-(4-oxo-2-(piperidin-1-yl)-4,5-dihydrothiazol-5-yl)acetamido)benzoic acid (7.2 mg, 20 µmol, 1.0 eq.) was reacted with thalidomide-O-C2-amine (TFA salt, 9.1 mg, 21 µmol, 1.05 eq.). Purified by preparative HPLC (MeCN/$H_2O$ gradient, 0.1% FA) to afford **13a** as a white solid (11.8 mg, 17.9 µmol, 89%). $^1$H NMR (600 MHz, $CD_3OD$) δ 8.00 (dt, *J* = 3.8, 1.9 Hz, 1H), 7.76 (dd, *J* = 8.5, 7.3 Hz, 1H), 7.71 – 7.65 (m, 1H), 7.53 (dt, *J* = 7.8, 1.3 Hz, 1H), 7.48 (d, *J* = 8.5 Hz, 1H), 7.45 (d, *J* = 7.3 Hz, 1H), 7.39 (t, *J* = 7.9 Hz, 1H), 5.11 (ddd, *J* = 12.7, 5.5, 1.5 Hz, 1H), 4.56 (dd, *J* = 10.8, 3.6 Hz, 1H), 4.41 (t, *J* = 5.3 Hz, 2H), 3.94 – 3.80 (m, 2H), 3.83 (t, *J* = 5.3 Hz, 2H), 3.59 – 3.47 (m, 2H), 3.38 (dd, *J* = 16.6, 3.6 Hz, 1H), 2.91 – 2.78 (m, 2H), 2.77 – 2.64 (m, 2H), 2.18 – 2.08 (m, 1H), 1.85 – 1.58 (m, 6H). $^{13}$C NMR (151 MHz, $CD_3OD$) δ 191.39, 181.77, 174.60, 171.48, 170.70, 170.17, 168.45, 167.99, 157.65, 140.00, 138.16, 136.36, 135.05, 130.12, 124.02, 123.83, 121.05, 120.03, 118.41, 117.02, 69.34, 53.15, 51.42, 50.75, 50.47, 41.07, 40.40, 32.16, 26.98, 26.58, 24.79, 23.64. HR-MS (ESI+) calc'd for $[M+H]^+$ $C_{32}H_{33}N_6O_8S$: 661.2075, found 661.2078 (0.5 ppm).

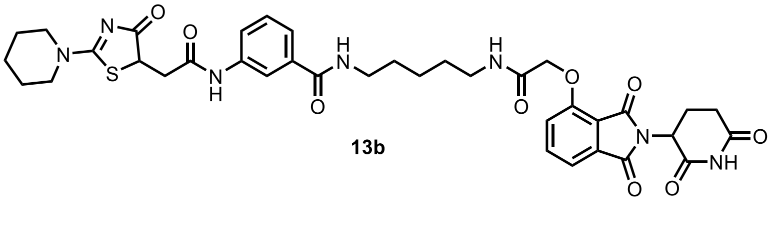

**13b**

**N-(5-(2-((2-(2,6-dioxopiperidin-3-yl)-1,3-dioxoisoindolin-4-yl)oxy)acetamido)pentyl)-3-(2-(4-oxo-2-(piperidin-1-yl)-4,5-dihydrothiazol-5-yl)acetamido)benzamide (13b)**

**General Procedure 1B**. Reaction scale: 3-(2-(4-oxo-2-(piperidin-1-yl)-4,5-dihydrothiazol-5-yl)acetamido)benzoic acid (7.2 mg, 20 µmol, 1.0 eq.) was reacted with thalidomide-O-acetamido-C5-amine (TFA salt, 11.1 mg, 21 µmol, 1.05 eq.). Purified by preparative HPLC (MeCN/$H_2O$ gradient, 0.1% FA) to afford **13b** as a white solid (12.7 mg, 16.7 µmol, 84%). $^1$H NMR (600 MHz, $CD_3OD$) δ 7.97 (t, *J* = 1.9 Hz, 1H), 7.77 (t, *J* = 7.9 Hz, 1H), 7.60 – 7.53 (m, 1H), 7.53 – 7.43 (m, 2H), 7.38 (d, *J* = 8.4 Hz, 1H), 7.32 (td, *J* = 7.9, 3.1 Hz, 1H), 5.13 (dd, *J* = 12.9, 5.5 Hz, 1H), 4.73 (s, 2H), 4.57 (dd, *J* = 10.8, 3.7 Hz, 1H), 4.01 – 3.81 (m, 2H), 3.61 – 3.51 (m, 2H), 3.48 – 3.33 (m, 5H), 3.00 – 2.63 (m, 4H), 2.14 (td, *J* = 7.1, 3.4 Hz, 1H), 1.87 – 1.60 (m, 10H), 1.48 (p, *J* = 7.8 Hz, 2H). $^{13}$C NMR (151 MHz, DMSO-$d_6$) δ 187.65, 179.26, 172.82, 169.93, 168.84, 166.76, 166.67 & 166.59 (rot.), 166.04 & 165.96 (rot.), 165.52, 155.07, 138.85, 136.95, 135.51, 133.04, 128.64, 121.70, 121.51, 120.37, 118.38, 116.80, 116.05, 67.60, 54.95, 51.65, 49.27, 48.80, 48.77, 38.31, 38.18, 30.96, 30.73, 28.77 & 28.74 & 28.73 (rot.), 25.58, 25.10, 23.78, 23.35, 22.01. HR-MS (ESI+) calc'd for $[M+H]^+$ $C_{37}H_{42}N_7O_{11}S$: 760.2759, found 760.2756 (0.4 ppm).

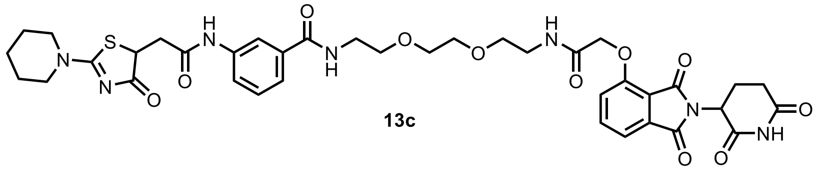

**13c**

**N-(2-(2-(2-(2-((2-(2,6-dioxopiperidin-3-yl)-1,3-dioxoisoindolin-4-yl)oxy)acetamido)ethoxy)ethoxy)ethyl)-3-(2-(4-oxo-2-(piperidin-1-yl)-4,5-dihydrothiazol-5-yl)acetamido)benzamide (13c)**

**General Procedure 1B**. Reaction scale: 3-(2-(4-oxo-2-(piperidin-1-yl)-4,5-dihydrothiazol-5-yl)acetamido)benzoic acid (7.2 mg, 20 µmol, 1.0 eq.) was reacted with thalidomide-O-acetamido-PEG2-amine (HCl salt, 10.5 mg, 21 µmol, 1.05 eq.). Purified by preparative HPLC (MeCN/$H_2O$ gradient, 0.1% FA) to afford **13c** as a white solid (10.6 mg, 13.2 µmol, 66%). $^1$H NMR (600 MHz, DMSO-$d_6$) δ 11.12 (s, 1H), 10.31 (s, 1H), 8.54 – 8.42 (m, 1H), 8.02 (t, *J* = 1.9 Hz, 1H), 8.01 (s, 1H), 7.80 (dd, *J* = 8.5, 7.3 Hz, 1H), 7.70 (d, *J* = 7.7 Hz, 1H), 7.49 (t, *J* = 7.5 Hz, 2H), 7.41 – 7.34 (m, 2H), 5.11 (dd, *J* = 12.9, 5.4 Hz, 1H), 4.78 (s, 2H), 4.44 (dd, *J* = 11.2, 3.5 Hz, 1H), 3.84 (dt, *J* = 11.5, 5.5 Hz, 1H), 3.77 (dt, *J* = 12.5, 5.6 Hz, 1H), 3.58 – 3.49 (m, 6H), 3.46 (t, *J* = 5.9 Hz, 4H), 3.39 (q, *J* = 5.9 Hz, 2H), 3.36 – 3.25 (m, 9H), 2.89 (ddd, *J* = 16.9, 13.9, 5.5 Hz, 1H), 2.72 (dd, *J* = 16.6, 11.2 Hz, 1H), 2.66 – 2.51 (m, 2H), 2.04 (ddd, *J* = 14.2, 6.3, 3.9 Hz, 1H), 1.76 – 1.49 (m, 4H). $^{13}$C NMR (151 MHz, DMSO-$d_6$) (*1 C expected under solvent*) δ 187.65, 179.27, 172.82, 169.92, 168.86, 166.92, 166.76, 166.20, 165.45, 155.00, 138.90, 136.94, 135.21, 133.04, 128.68, 121.72, 121.64, 120.31, 118.40, 116.74, 116.04, 69.61, 69.57, 68.87, 68.85, 67.46, 54.95, 51.65, 49.27, 48.80, 48.77, 38.42, 30.96, 25.58, 25.10, 23.35, 22.01. HR-MS (ESI+) calc'd for $[M+H]^+$ $C_{38}H_{44}N_7O_{11}S$: 806.2814, found 806.2832 (2.3 ppm).

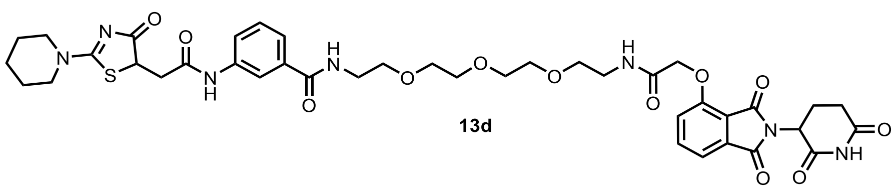

**13d**

**N-(1-((2-(2,6-dioxopiperidin-3-yl)-1,3-dioxoisoindolin-4-yl)oxy)-2-oxo-6,9,12-trioxa-3-azatetradecan-14-yl)-3-(2-(4-oxo-2-(piperidin-1-yl)-4,5-dihydrothiazol-5-yl)acetamido)benzamide (13d)**

**General Procedure 1B.** Reaction scale: 3-(2-(4-oxo-2-(piperidin-1-yl)-4,5-dihydrothiazol-5-yl)acetamido)benzoic acid (7.2 mg, 20 µmol, 1.0 eq.) was reacted with thalidomide-O-acetamido-PEG3-

amine (HCl salt, 11.4 mg, 21 μmol, 1.05 eq.). Purified by preparative HPLC (MeCN/$H_2O$ gradient, 0.1% FA) to afford **13d** as a white solid (11.7 mg, 13.8 μmol, 69%). $^1$H NMR (600 MHz, $CD_3OD$) δ 8.00 (t, *J* = 1.9 Hz, 1H), 7.78 (t, *J* = 7.9 Hz, 1H), 7.65 – 7.61 (m, 1H), 7.53 – 7.48 (m, 2H), 7.42 – 7.34 (m, 2H), 5.11 (ddd, *J* = 12.8, 5.6, 1.7 Hz, 1H), 4.73 (s, 2H), 4.56 (ddd, *J* = 10.9, 3.6, 1.3 Hz, 1H), 3.95 – 3.82 (m, 2H), 3.68 – 3.51 (m, 16H), 3.46 (t, *J* = 5.4 Hz, 2H), 3.38 (dd, *J* = 16.6, 3.2 Hz, 1H), 2.94 – 2.66 (m, 5H), 2.20 – 2.09 (m, 1H), 1.79 – 1.60 (m, 6H). $^{13}$C NMR (151 MHz, $CD_3OD$) δ 191.35, 181.77, 174.61, 171.41, 170.70, 169.93, 169.91, 168.29, 167.56, 156.11, 139.98, 138.21, 136.54, 134.90, 130.10, 123.82 (3 Cs)z, 121.46, 119.87, 119.10, 117.86, 71.59, 71.57, 71.36, 71.33, 70.51, 70.27, 69.10, 53.14, 51.42, 50.75, 50.55, 41.09, 41.01, 40.19, 32.19, 26.99, 26.59, 24.80, 23.67. HR-MS (ESI+) calc'd for $[M+H]^+$ $C_{40}H_{48}N_7O_{12}S$: 850.3076, found 850.3073 (0.4 ppm).

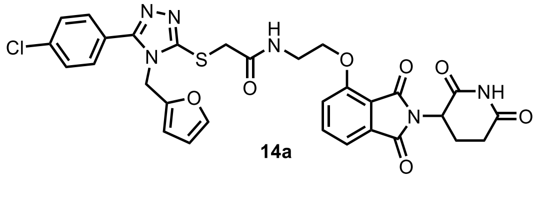

14a

**2-((5-(4-chlorophenyl)-4-(furan-2-ylmethyl)-4H-1,2,4-triazol-3-yl)thio)-N-(2-((2-(2,6-dioxopiperidin-3-yl)-1,3-dioxoisoindolin-4-yl)oxy)ethyl)acetamide (14a)**

**General Procedure 1B**. Reaction scale: 2-((5-(4-chlorophenyl)-4-(furan-2-ylmethyl)-4H-1,2,4-triazol-3-yl)thio)acetic acid (7.0 mg, 20 μmol, 1.0 eq.) was reacted with thalidomide-O-C2-amine (TFA salt, 9.1 mg, 21 μmol, 1.05 eq.). Purified by preparative HPLC (MeCN/$H_2O$ gradient, 0.1% FA) to afford **14a** as a white solid (10.3 mg, 15.9 μmol, 79%). $^1$H NMR (600 MHz, $CD_3OD$) δ 7.73 (dd, *J* = 8.5, 7.3 Hz, 1H), 7.62 – 7.56 (m, 2H), 7.56 – 7.49 (m, 2H), 7.47 – 7.37 (m, 3H), 6.34 (dd, *J* = 3.3, 1.8 Hz, 1H), 6.30 (dd, *J* = 3.3, 0.8 Hz, 1H), 5.28 (s, 2H), 5.07 (dd, *J* = 12.7, 5.5 Hz, 1H), 4.25 (t, *J* = 5.2 Hz, 2H), 3.94 (s, 2H), 3.76 – 3.59 (m, 2H), 2.85 (ddd, *J* = 17.3, 13.8, 5.4 Hz, 1H), 2.73 (ddd, *J* = 17.3, 4.5, 2.5 Hz, 1H), 2.67 (qd, *J* = 12.8, 4.5 Hz, 1H), 2.08 (dtd, *J* = 13.1, 5.3, 2.4 Hz, 1H). $^{13}$C NMR (151 MHz, $CD_3OD$) δ 174.60, 171.37, 170.21, 168.40, 167.84, 157.52, 156.55, 153.09, 149.00, 144.84, 138.14, 138.03, 135.03, 131.56, 130.39, 126.19, 120.99, 118.35, 117.00, 111.72, 111.05, 69.32, 50.45, 42.96, 40.19, 37.86, 32.15, 23.63. HR-MS (ESI+) calc'd for $[M+H]^+$ $C_{30}H_{26}ClN_6O_7S$: 649.1267, found 649.1264 (0.5 ppm).

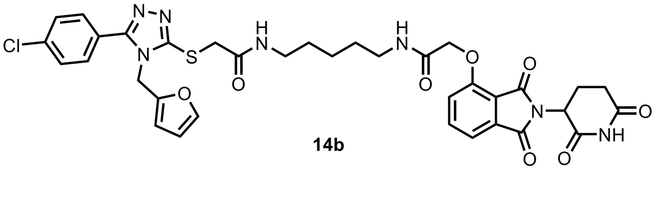

14b

**2-((5-(4-chlorophenyl)-4-(furan-2-ylmethyl)-4H-1,2,4-triazol-3-yl)thio)-N-(5-(2-((2-(2,6-dioxopiperidin-3-yl)-1,3-dioxoisoindolin-4-yl)oxy)acetamido)pentyl)acetamide (14b)**

**General Procedure 1B**. Reaction scale: Reaction scale: 2-((5-(4-chlorophenyl)-4-(furan-2-ylmethyl)-4H-1,2,4-triazol-3-yl)thio)acetic acid (7.0 mg, 20 μmol, 1.0 eq.) was reacted with thalidomide-O-acetamido-C5-amine (TFA salt, 11.1 mg, 21 μmol, 1.05 eq.). Purified by preparative HPLC (MeCN/$H_2O$ gradient, 0.1% FA) to afford **14b** as a white solid (11.1 mg, 14.8 μmol, 74%). $^1$H NMR (400 MHz, DMSO-$d_6$) δ 11.13 (s, 1H), 8.24 (t, *J* = 5.6 Hz, 1H), 7.97 (t, *J* = 5.7 Hz, 1H), 7.81 (dd, *J* = 8.5, 7.3 Hz, 1H), 7.70 (d, *J* = 8.5 Hz, 2H), 7.62 (d, *J* = 8.5 Hz, 3H), 7.49 (d, *J* = 7.2 Hz, 1H), 7.38 (d, *J* = 8.5 Hz, 1H), 6.41 (dd, *J* = 3.3, 1.9 Hz, 1H), 6.32 (d, *J* = 3.3 Hz, 1H), 5.26 (s, 2H), 5.12 (dd, *J* = 12.9, 5.4 Hz, 1H), 4.77 (s, 2H), 3.93 (s, 2H), 3.12 (q, *J* = 6.6 Hz, 2H), 3.05 (q, *J* = 6.6 Hz, 2H), 2.90 (ddd, *J* = 17.4, 14.0, 5.4 Hz, 1H), 2.69 – 2.49 (m, 2H), 2.10 – 1.98 (m, 1H), 1.48 – 1.33 (m, 4H), 1.30 – 1.20 (m, 2H). $^{13}$C NMR (151 MHz, DMSO-$d_6$) δ 172.83, 169.93, 166.77, 166.67, 166.47, 165.53, 155.08, 154.19, 151.22, 147.83, 143.72, 136.96, 135.07, 133.04, 130.31, 129.12, 125.72, 120.38, 116.80, 116.05, 110.79, 109.59, 67.61, 48.81, 41.36, 38.92, 38.25, 36.88, 30.96, 28.66, 28.57, 23.60, 22.01. HR-MS (ESI+) calc'd for $[M+H]^+$ $C_{35}H_{35}ClN_7O_8S$: 748.1951, found 748.1947 (0.6 ppm).

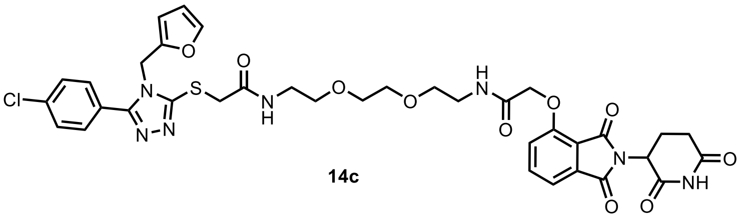

**14c**

**2-((5-(4-chlorophenyl)-4-(furan-2-ylmethyl)-4H-1,2,4-triazol-3-yl)thio)-N-(2-(2-(2-(2-((2-(2,6-dioxopiperidin-3-yl)-1,3-dioxoisoindolin-4-yl)oxy)acetamido)ethoxy)ethoxy)ethyl)acetamide (14c)**

**General Procedure 1B**. Reaction scale: 2-((5-(4-chlorophenyl)-4-(furan-2-ylmethyl)-4H-1,2,4-triazol-3-yl)thio)acetic acid (7.0 mg, 20 µmol, 1.0 eq.) was reacted with thalidomide-O-acetamido-PEG2-amine (HCl salt, 10.5 mg, 21 µmol, 1.05 eq.). Purified by preparative HPLC (MeCN/$H_2O$ gradient, 0.1% FA) to afford **14c** as a white solid (8.2 mg, 10.3 µmol, 52%). $^1H$ NMR (400 MHz, $CD_3OD$) δ 7.79 (dd, *J* = 8.5, 7.3 Hz, 1H), 7.69 – 7.64 (m, 2H), 7.60 – 7.54 (m, 2H), 7.52 (d, *J* = 7.3 Hz, 1H), 7.46 (dd, *J* = 1.9, 0.8 Hz, 1H), 7.42 (d, *J* = 8.5 Hz, 1H), 6.36 (dd, *J* = 3.3, 1.9 Hz, 1H), 6.31 (dd, *J* = 3.3, 0.8 Hz, 1H), 5.28 (s, 2H), 5.12 (dd, *J* = 12.5, 5.5 Hz, 1H), 4.76 (s, 2H), 3.91 (s, 2H), 3.64 – 3.58 (m, 6H), 3.50 (m, 4H), 3.35 (t, *J* = 5.4 Hz, 2H), 2.88 (ddd, *J* = 17.8, 14.2, 5.2 Hz, 1H), 2.79 – 2.64 (m, 2H), 2.22 – 2.07 (m, 1H). $^{13}C$ NMR (151 MHz, DMSO-$d_6$) (*1 C expected under solvent*) δ 173.28, 170.37, 167.38, 167.27, 167.21, 165.92, 155.45, 154.64, 151.63, 148.28, 144.17, 137.41, 135.52, 133.50, 130.75, 129.57, 126.18, 120.78, 117.20, 116.51, 111.24, 110.02, 70.02 (2 Cs), 69.34, 69.29, 67.92, 49.26, 41.81, 38.86 & 38.75 (rot), 37.27, 31.41, 22.46. HR-MS (ESI+) calc'd for $[M+H]^+$ $C_{36}H_{37}ClN_7O_{10}S$: 794.2006, found 794.2027 (2.7 ppm).

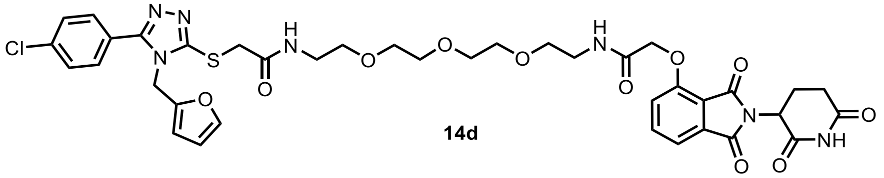

**14d**

**2-((5-(4-chlorophenyl)-4-(furan-2-ylmethyl)-4H-1,2,4-triazol-3-yl)thio)-N-(1-((2-(2,6-dioxopiperidin-3-yl)-1,3-dioxoisoindolin-4-yl)oxy)-2-oxo-6,9,12-trioxa-3-azatetradecan-14-yl)acetamide (14d)**

**General Procedure 1B.** Reaction scale: 2-((5-(4-chlorophenyl)-4-(furan-2-ylmethyl)-4H-1,2,4-triazol-3-yl)thio)acetic acid (7.0 mg, 20 µmol, 1.0 eq.) was reacted with thalidomide-O-acetamido-PEG3-amine (HCl salt, 11.4 mg, 21 µmol, 1.05 eq.). Purified by preparative HPLC (MeCN/$H_2O$ gradient, 0.1% FA) to afford **14d** as a white solid (11.3 mg, 13.5 µmol, 67%). $^1H$ NMR (400 MHz, $CD_3OD$) δ 7.79 (dd, *J* = 8.5, 7.3 Hz, 1H), 7.72 – 7.61 (m, 2H), 7.61 – 7.55 (m, 2H), 7.52 (d, *J* = 7.3 Hz, 1H), 7.46 (dd, *J* = 1.9, 0.8 Hz, 1H), 7.42 (d, *J* = 8.4 Hz, 1H), 6.36 (dd, *J* = 3.3, 1.9 Hz, 1H), 6.31 (dd, *J* = 3.3, 0.8 Hz, 1H), 5.28 (s, 2H), 5.13 (dd, *J* = 12.6, 5.5 Hz, 1H), 4.76 (s, 2H), 3.93 (s, 2H), 3.62 (s, 4H), 3.61 – 3.53 (m, 6H), 3.54 – 3.46 (m, 4H), 3.36 (t, *J* = 5.4 Hz, 2H), 2.88 (ddd, *J* = 17.7, 14.3, 5.1 Hz, 1H), 2.81 – 2.65 (m, 2H), 2.14 (dtd, *J* = 12.9, 4.9, 2.1 Hz, 1H). $^{13}C$ NMR (151 MHz, $CD_3OD$) δ 174.61, 171.36, 169.96, 169.90, 168.30, 167.59, 156.63, 156.16, 153.27, 149.04, 144.86, 138.20, 138.08, 134.93, 131.73, 130.42, 126.33, 121.52, 119.17, 117.89, 111.74, 111.04, 71.58, 71.54, 71.36, 71.27, 70.32, 70.29, 69.18, 50.55, 49.43, 42.93, 40.87, 40.19, 38.05, 32.18, 23.66. HR-MS (ESI+) calc'd for $[M+H]^+$ $C_{38}H_{41}ClN_7O_{11}S$: 838.2268, found 838.2261 (0.9 ppm).

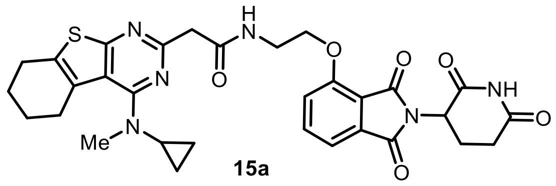

**15a**

**2-(4-(cyclopropyl(methyl)amino)-5,6,7,8-tetrahydrobenzo[4,5]thieno[2,3-d]pyrimidin-2-yl)-N-(2-((2-(2,6-dioxopiperidin-3-yl)-1,3-dioxoisoindolin-4-yl)oxy)ethyl)acetamide (15a)**

**General Procedure 1A.** Reaction scale: thalidomide-O-C2-amine 15 mg (47.2 µmol, 1.0 eq) and 2-(4-(cyclopropyl(methyl)amino)-5,6,7,8-tetrahydrobenzo[4,5]thieno[2,3-d]pyrimidin-2-yl)acetic acid 15 mg (47.2 µmol,1.0 eq). Purified by pTLC (6% MeOH/DCM) to afford **15a** as a yellow sticky solid (8 mg, 28%). $^{1}$H NMR (600 MHz, CDCl3) δ 8.49 (s, 2H), 8.15 (s, 1H), 7.67 – 7.64 (m, 1H), 7.43 (d, J = 7.3 Hz, 1H), 7.30 (d, J = 8.5 Hz, 1H), 4.98 – 4.92 (m, 1H), 4.32 (t, J = 5.5 Hz, 2H), 3.97 (s, 2H), 3.80 – 3.72 (m, J = 5.5 Hz, 2H), 3.70 – 3.62 (m, 1H), 3.21 (s, 2H), 3.16 – 3.03 (m, 2H), 2.86 – 2.78 (m, 4H), 2.19 – 2.08 (m, 1H), 1.94 – 1.87 (m, 2H), 1.80 – 1.73 (m, 2H), 1.56 – 1.50 (m, 2H), 1.44 – 1.42 (m, 1H), 1.27 – 1.23 (m, 2H), 0.90 – 0.87 (m, 2H). $^{13}$C NMR (151 MHz, $CDCl_3$) δ 171.13, 170.35, 170.00, 168.68, 167.11, 166.80, 166.01, 157.47, 154.75, 154.51, 136.99, 133.68, 119.51, 118.08, 117.35, 86.45, 80.31, 70.43, 70.26, 69.89, 69.56, 68.08, 62.26, 55.93, 50.03, 41.32, 39.40, 39.02, 31.95, 31.89, 29.72, 28.37, 27.30, 22.72, 21.94, 14.63, 14.15. HR-MS (ESI+) calc'd for $C_{31}H_{32}N_6O_6S$: 616.2103, found $[M+H]^+$ 617.2184.

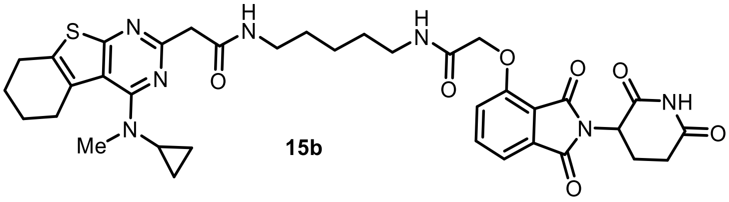

**15b**

**2-(4-(cyclopropyl(methyl)amino)-5,6,7,8-tetrahydrobenzo[4,5]thieno[2,3-d]pyrimidin-2-yl)-N-(5-(2-((2-(2,6-dioxopiperidin-3-yl)-1,3-dioxoisoindolin-4-yl)oxy)acetamido) pentyl) acetamide (15b)**

**General Procedure 1A.** Reaction scale: thalidomide-O-acetamido-C5-amine 19 mg (37.8 µmol, 1.0 eq) and 2-(4-(cyclopropyl(methyl)amino)-5,6,7,8-tetrahydrobenzo[4,5]thieno[2,3-d]pyrimidin-2-yl)acetic acid 12 mg (37.8 µmol,1.0 eq). Purified by pTLC (7% MeOH/DCM) **15b** as a yellow sticky solid (6.8 mg, 23 %). $^{1}$H NMR (600 MHz, $CDCl_3$) δ 9.27 (s, 1H), 8.32 (s, 1H), 7.80 (s, 1H), 7.76 – 7.69 (m, 1H), 7.61 – 7.58 (m, 1H), 7.53 (d, *J* = 7.4 Hz, 1H), 7.22 – 7.18 (m, 1H), 5.02 – 4.98 (m, 1H), 4.70 – 4.59 (m, 2H), 3.91 (s, 2H), 3.72 – 3.67 (m, 2H), 3.48 – 3.44 (m, 1H), 3.32 – 3.28 (m, 3H), 3.24 – 3.19 (m, 2H), 3.17 – 3.14 (m, 2H), 3.04 – 2.98 (m, 1H), 2.92 – 2.74 (m, 4H), 2.19 – 2.17 (m, 2H), 1.97 – 1.88 (m, 2H), 1.78 – 1.74 (m, 2H), 1.61 – 1.57 (m, 4H), 1.49 – 1.46 (m, 3H), 1.43 – 1.39 (m, 3H), 0.92 (s, 1H). $^{13}$C NMR (151 MHz, DMSO) δ 172.82, 169.92, 167.49, 167.43, 166.77, 166.66, 165.53, 160.25, 156.29, 155.09, 136.96, 133.04, 132.84, 128.43, 120.38, 118.01, 117.96, 116.79, 116.05, 67.61, 53.50, 48.81, 44.67, 41.76, 40.73, 38.65, 38.26, 35.17, 30.96, 28.73, 26.78, 25.20, 23.66, 22.55, 22.28, 22.02, 18.06, 16.74, 12.43, 8.62. HR-MS (ESI+) calc'd for $C_{36}H_{41}N_7O_7S$: 715.8260, found $[M+H]^+$ 716.2861.

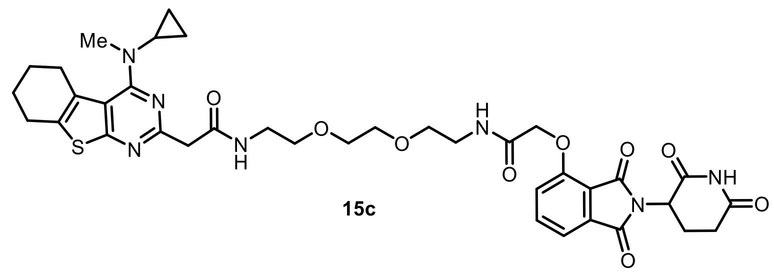

**15c**

**2-(4-(cyclopropyl(methyl)amino)-5,6,7,8-tetrahydrobenzo[4,5]thieno[2,3-d]pyrimidin-2-yl)-N-(2-(2-(2-(2-((2-(2,6-dioxopiperidin-3-yl)-1,3-dioxoisoindolin-4-yl)oxy)acetamido)ethoxy)ethoxy)ethyl)acetamide (15c)**

**General Procedure 1A.** Reaction scale: Thalidomide-O-acetamido-PEG2-C2-amine 19 mg (37.8 µmol, 1.0 eq) and 2-(4-(cyclopropyl(methyl)amino)-5,6,7,8-tetrahydrobenzo[4,5]thieno[2,3-d]pyrimidin-2-yl)acetic acid 12 mg (37.8 µmol,1.0 eq). Purified by pTLC (7% MeOH/DCM) **15c** as a yellow sticky solid (4 mg, 22 %). $^{1}$H NMR (600 MHz, CDCl3) δ 9.13 (s, 1H), 8.32 (s, 1H), 7.72 (d, *J* = 7.3 Hz, 1H), 7.64 –7.61 (m, 1H), 7.53 (d, *J* = 7.3 Hz, 1H), 7.18 (d, *J* = 8.4 Hz, 1H), 5.01 – 4.90 (m, 1H), 4.64 (s, 2H), 3.86 (s, 2H), 3.64 – 3.53 (m, 9H), 3.47 (d, *J* = 5.7 Hz, 2H), 3.08 (s, 3H), 2.87 – 2.84 (m 6H), 1.94 – 1.90 (m, 2H), 1.76 –1.72 (m, 2H), 0.82 – 0.80 (m, 1H). $^{13}$C NMR (151 MHz, $CDCl_3$) δ 171.39, 169.23, 168.38, 167.01, 166.81, 165.95, 154.52, 137.07, 133.83, 127.84, 119.37, 119.27, 118.16, 117.39, 70.51, 70.18, 70.06, 69.66, 67.98, 53.57, 49.44, 42.09, 39.46, 39.20, 34.80, 31.58, 27.14, 25.92, 23.10, 22.96, 22.81, 8.52. HR-MS (ESI+) calc'd for $C_{37}H_{43}N_7O_9S$: 761.2843, found $[M+H]^+$ 762.2902.

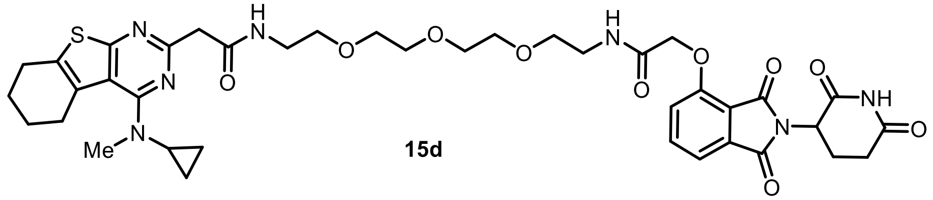

**15d**

**2-(4-(cyclopropyl(methyl)amino)-5,6,7,8-tetrahydrobenzo[4,5]thieno[2,3-d]pyrimidin-2-yl)-N-(1-((2-(2,6-dioxopiperidin-3-yl)-1,3-dioxoisoindolin-4-yl)oxy)-2-oxo-6,9,12-trioxa-3-azatetradecan-14-yl)acetamide (15d)**

**General Procedure 1A.** Reaction scale**:** Thalidomide-O-acetamido-PEG3-C2-amine 20 mg (37.8 µmol, 1.0 eq) and 2-(4-(cyclopropyl(methyl)amino)-5,6,7,8-tetrahydrobenzo[4,5]thieno[2,3-d]pyrimidin-2-yl)acetic acid 12 mg (37.8 µmol,1.0 eq). Purified by pTLC (7% MeOH/DCM) **15d** as a yellow sticky solid (10 mg, 34 %). $^{1}$H NMR (600 MHz, $CDCl_3$) δ 9.01 (s, 1H), 8.20 (t, *J* = 5.8 Hz, 1H), 7.73 (d, *J* = 7.3 Hz, 1H), 7.66 (t, *J* = 5.6 Hz, 1H), 7.54 (d, *J* = 7.3 Hz, 1H), 7.19 (d, *J* = 8.4 Hz, 1H), 4.98 – 4.94 (m, 1H), 4.65 (s, 2H), 3.86 (s, 2H), 3.67 – 3.56 (m, 10 H), 3.48 (d, *J* = 5.4 Hz, 2H), 3.11 (s, 3H), 2.90 – 2.81 (m, 4H), 2.81 – 2.71 (m, 2H), 1.95 – 1.87 (m, 2H), 1.78 – 1.74 (m, 2H), 0.82 (d, *J* = 6.6 Hz, 1H). $^{13}$C NMR (151 MHz, DMSO) δ 172.81, 169.90, 168.14, 166.92, 166.75, 165.46, 160.73, 157.46, 155.00, 136.96, 133.05, 132.43, 128.05, 120.33, 118.31, 116.75, 116.05, 69.76, 69.63, 69.13, 68.84, 67.49, 48.81, 45.44, 40.90, 40.43, 38.78, 38.42, 34.86, 30.97, 26.56, 25.23, 22.58, 22.34, 22.02, 8.28. HR-MS (ESI+) calc'd for $C_{39}H_{47}N_7O_{10}S$: 805.3105, found $[M+H]^+$ 806.3179.

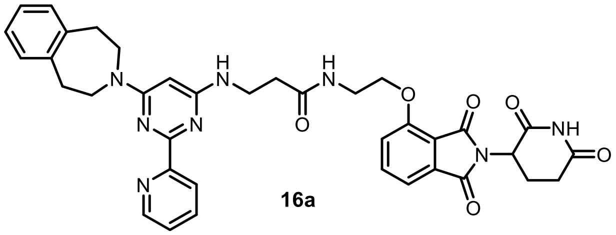

**16a**

**N-(2-((2-(2,6-dioxopiperidin-3-yl)-1,3-dioxoisoindolin-4-yl)oxy)ethyl)-3-((2-(pyridin-2-yl)-6-(1,2,4,5-tetrahydro-3H-benzo[d]azepin-3-yl)pyrimidin-4-yl)amino)propenamide (16a)**

**General Procedure 1A.** Reaction scale: Thalidomide-O-C2-amine 9 mg (25.6 μmol, 1.0 eq) and 3-((2-(pyridin-2-yl)-6-(1,2,4,5-tetrahydro-3H-benzo[d]azepin-3-yl)pyrimidin-4-yl)amino)propanoic acid 10 mg (25.6 μmol,1.0 eq). Purified by pTLC (7% MeOH/DCM) to afford **16a** as a yellow sticky solid (4 mg, 22%). $^{1}$H NMR (600 MHz, $CDCl_3$) δ 8.7 (d, *J* = 7.2 Hz, 1H), 8.33 (d, *J* = 7.8 Hz, 1H), 7.86 – 7.78 (m, 1H), 7.61 – 7.57 (m, 2H), 7.42 – 7.39 (m, 2H), 7.21 – 7.11 (m, 5H), 5.62 (s, 1H), 5.00 – 4.94 (m, 1H), 4.28 – 4.24 (m, 1H), 4.15 – 4.10 (m, 1H), 3.97– 3.91 (m, 4H), 3.77 – 3.74 (m, 1H), 3.68 – 3.63 (m, 2H), 3.57 (s, 1H), 3.06 – 2.99 (m, 4H), 2.88 – 2.81 (m, 1H), 2.78 – 2.75 (m, 2H), 2.65 – 2.61 (m, 2H). $^{13}$C NMR (151 MHz, DMSO) δ 172.82, 171.01, 169.96, 166.78, 165.27, 160.96, 155.68, 149.04, 140.38, 137.03, 133.25, 129.88, 126.33, 123.16, 120.06, 116.49, 115.57, 67.46, 54.96, 48.76, 48.62, 47.13, 40.43, 38.04, 35.74, 30.97, 22.04. HR-MS (ESI+) calc'd for $C_{37}H_{36}N_8O_6$: 688.7450, found $[M+H]^+$ 689.2818.

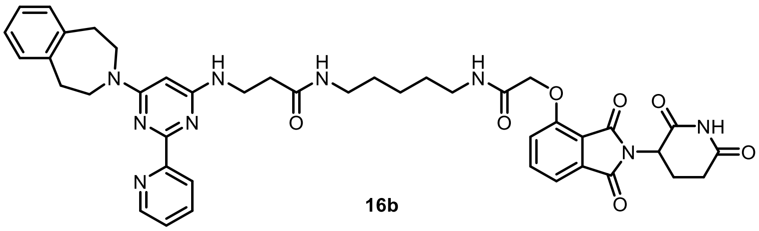

**16b**

**N-(5-(2-((2-(2,6-dioxopiperidin-3-yl)-1,3-dioxoisoindolin-4-yl)oxy)acetamido)pentyl)-3-((2-(pyridin-2-yl)-6-(1,2,4,5-tetrahydro-3H-benzo[d]azepin-3-yl)pyrimidin-4-yl)amino) propenamide (16b)**

**General Procedure 2.** Reaction scale: Thalidomide-O-acetamido-C5-amine 13 mg (25.6 μmol, 1.0 eq) and 3-((2-(pyridin-2-yl)-6-(1,2,4,5-tetrahydro-3H-benzo[d]azepin-3-yl)pyrimidin-4-yl)amino)propanoic acid 10 mg (25.6 μmol,1.0 eq). Purified by pTLC (7% MeOH/DCM) **16b** as a yellow sticky solid (4 mg, 22 %). $^{1}$H NMR (400 MHz, $CDCl_3$) δ 8.76 – 8.74 (m, 1H), 8.36 (d, *J* = 7.8 Hz, 1H), 7.85 (d, *J* = 7.0 Hz, 1H), 7.77 – 7.67 (m, 1H), 7.58 (s, 1H), 7.54 – 7.48 (m, 1H), 7.47 – 7.44 (m, 1H), 7.19 – 7.11 (m, 5H), 5.65 (s, 1H), 5.05 – 5.01 (m, 1H), 4.63 – 4.61 (m, 2H), 3.96 – 3.92 (m, 4H), 3.70 – 3.55 (m, 2H), 3.26 – 3.24 (m, 3H), 3.07 – 2.95 (m, 4H), 2.91 – 2.73 (m, 3H), 2.72 – 2.53 (m, 3H), 2.15 – 2.11 (m, 1H), 1.56 – 1.52 (m, 2H), 1.44 – 1.39 (m, 2H), $^{13}$C NMR (151 MHz, DMSO) δ 172.84, 170.50, 169.93, 166.77, 166.66, 165.54, 155.06, 148.99, 140.87, 136.96, 133.04, 129.81, 126.22, 123.18, 120.39, 116.82, 116.08, 67.63, 48.82, 46.69, 38.48, 38.29, 35.97, 30.98, 28.81, 28.73, 23.75, 22.03. HR-MS (ESI+) calc'd for $C_{42}H_{45}N_9O_7$: 787.8780, found $[M+H]^+$ 788.3516.

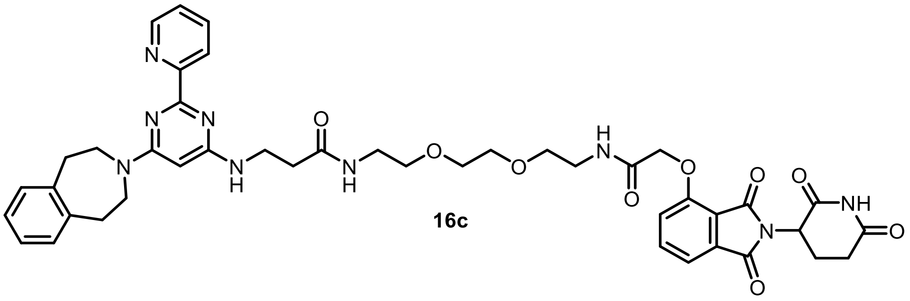

**16c**

**N-(2-(2-(2-(2-((2-(2,6-dioxopiperidin-3-yl)-1,3-dioxoisoindolin-4-yl)oxy)acetamido) ethoxy) ethoxy)ethyl)-3-((2-(pyridin-2-yl)-6-(1,2,4,5-tetrahydro-3H-benzo[d]azepin-3-yl)pyrimidin-4-yl)amino)propenamide (16c)**

**General Procedure 2.** Reaction scale: Thalidomide-O-acetamido-PEG2-C2-amine 17 mg (33.9 μmol, 1.0 eq), and 3-((2-(pyridin-2-yl)-6-(1,2,4,5-tetrahydro-3H-benzo[d]azepin-3-yl)pyrimidin-4-yl)amino)propanoic acid 12 mg (30.8 μmol,1.0 eq). Purified by pTLC (7% MeOH/DCM) **16c** as a yellow sticky solid (4 mg, 23%). $^{1}$H NMR (600 MHz, $CD_3OD$_SPE) δ 8.74 – 8.67 (m, 1H), 8.47 – 8..45 (m, 2H), 8.27 (s, 1H), 8.02 (dd, *J* = 7.8, 1.7 Hz, 1H), 7.72 (d, *J* = 7.3 Hz, 1H), 7.63 – 7.57 (m, 1H), 7.45 (d, *J* = 7.3 Hz, 1H), 7.33 (d, *J* = 8.4 Hz, 1H), 7.19 – 7.03 (m, 3H), 5.12 – 5.09 (m, 1H), 4.67 (s, 2H), 4.58 (s, 4H), 3.97 (s, 4H), 3.67 (t, *J* = 6.4 Hz, 2H), 3.61 – 3.54 (m, 4H), 3.50 – 3.46 (m, 2H), 3.34 (t, *J* = 5.4 Hz, 1H), 3.06 – 3.00 (m, 2H), 2.91 – 2.80 (m, 1H), 2.78 – 2.63 (m, 2H), 2.56 (t, *J* = 6.4 Hz, 1H), 2.13 – 2.09 (m, 2H). $^{13}$C NMR (151 MHz, DMSO) δ 172.84, 170.64, 169.92, 166.94, 166.76, 165.47, 161.19, 154.98, 149.04, 140.55, 136.97, 133.04, 129.86, 126.29, 123.18, 120.33, 116.76, 116.08, 69.57, 69.13, 68.83, 67.48, 48.82, 46.92, 38.60, 38.40, 35.82, 30.97, 22.02. HR-MS (ESI+) calc'd for $C_{43}H_{47}N_9O_9$: 833.9030, found $[M+H]^+$ 834.3573.

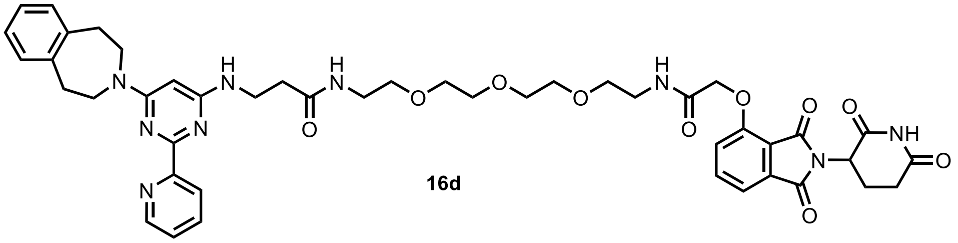

**16d**

**N-(1-((2-(2,6-dioxopiperidin-3-yl)-1,3-dioxoisoindolin-4-yl)oxy)-2-oxo-6,9,12-trioxa-3-azatetradecan-14-yl)-3-((2-(pyridin-2-yl)-6-(1,2,4,5-tetrahydro-3H-benzo[d]azepin-3-yl)pyrimidin-4-yl)amino)propenamide (16d)**

**General Procedure 2.** Reaction scale: thalidomide-O-acetamido-PEG3-C2-amine 18.4 mg (33.9 μmol, 1.0 eq), and 3-((2-(pyridin-2-yl)-6-(1,2,4,5-tetrahydro-3H-benzo[d]azepin-3-yl)pyrimidin-4-yl)amino)propanoic acid 12 mg (30.8 μmol,1.0 eq). Purified by pTLC (7% MeOH/DCM) to get **16d** as a yellow sticky solid (3.5 mg, 21%). $^{1}$H NMR (600 MHz, $CDCl_3$) δ 8.68 (d, *J* = 7.2 Hz, 1H), 8.45 – 8.42 (m, 2H), 8.05 – 7.93 (m, 1H), 7.75 – 7.73 (m, 1H), 7.56 – 7.53 (m, 1H), 7.51 – 7.47 (m, 1H), 7.35 – 7.29 (m, 1H), 7.22 – 6.98 (m, 3H), 5.81 (d, *J* = 6.3 Hz, 1H), 5.54 – 5.39 (m, 1H), 5.17 – 5.02 (m, 2H), 4.68 (t, *J* = 7.7 Hz, 2H), 3.95 (s, 2H), 3.68 (t, *J* = 7.2 Hz, 2H), 3.64 – 3.51 (m, 4H), 3.49 – 3.45 (m, 2H), 3.39 – 3.26 (m, 8H), 3.04 – 3.01 (m, 2H), 2.92 – 2.61 (m, 4H), 2.58 – 2.54 (m, 2H), 2.15 – 2.09 (m, 2H). $^{13}$C NMR (151 MHz, DMSO) δ 173.28, 171.30, 170.38, 167.39, 167.22, 165.93, 161.99, 155.45, 149.47, 141.36, 137.43, 137.09, 133.51, 130.26, 126.66, 124.76, 123.63, 120.80, 117.22, 116.53, 69.58, 69.28, 67.95, 49.28, 47.11, 40.89, 39.03, 38.87, 37.81, 36.44, 31.43, 22.48. HR-MS (ESI+) calc'd for $C_{45}H_{51}N_9O_{10}$: 877.3759, found $[M+H]^+$ 878.3839.

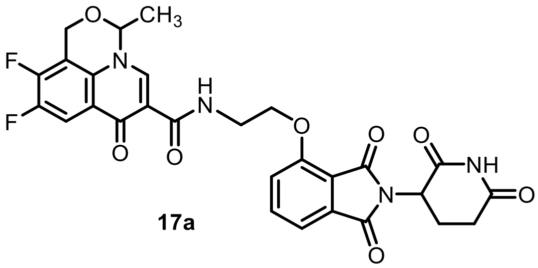

**17a**

**N-(2-((2-(2,6-dioxopiperidin-3-yl)-1,3-dioxoisoindolin-4-yl)oxy)ethyl)-9,10-difluoro-3-methyl-7-oxo-1H,3H,7H-[1,3]oxazino[5,4,3-ij]quinoline-6-carboxamide (17a)**

**General Procedure 2.** Reaction scale: Thalidomide-O-C2-amine 22 mg (71.1 µmol, 1.0 eq) and 9,10-difluoro-3-methyl-7-oxo-1H,3H,7H-[1,3]oxazino[5,4,3-ij]quinoline-6-carboxylic acid 20 mg (71.1 µmol,1.0 eq). Purified by pTLC (7% MeOH/DCM) to afford **17a** as an off-white solid (15.0 mg, 54%). $^{1}$H NMR (600 MHz, DMSO) δ 11.13 (s, 1H), 10.04 (s, 1H), 8.90 (s, 1H), 7.81 (d, J = 7.9 Hz, 1H), 7.72 (d, J = 7.6 Hz, 1H), 7.64 – 7.61 (m, 2H), 7.46 (d, J = 7.0 Hz, 1H), 5.13 – 5.07 (m, 1H), 4.97 – 4.90 (m, 1H), 4.66 – 4.63 (m, 1H), 4.48 – 4.44 (m, 1H), 4.38 (t, J = 5.8 Hz, 2H), 3.83 – 3.76 (m, 2H), 2.90 – 2.88 (m, 2H), 2.63 – 2.47 (m, 4H), 2.08 – 2.02 (m, 1H), 1.46 – 1.42 (m, 3H).$^{13}$C NMR (151 MHz, DMSO) δ 174.37, 173.29, 170.44, 167.27, 165.66, 164.69, 156.17, 149.62, 149.55, 147.99, 147.92, 146.20, 142.30, 142.18, 140.63, 140.52, 137.51, 135.84, 135.76, 133.75, 125.21, 123.27, 123.23, 120.57, 116.95, 116.04, 111.21, 104.08, 103.95, 69.34, 68.29, 54.63, 49.21, 31.42, 22.48, 18.21. HR-MS (ESI+) calc'd for $C_{28}H_{22}F_2N_4O_8$: 580.1406, found $[M+H]^+$ 581.1485.

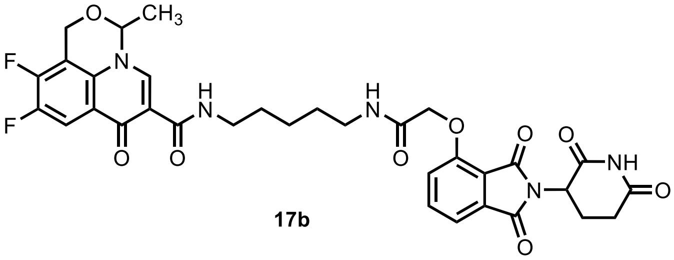

**17b**

**N-(5-(2-((2-(2,6-dioxopiperidin-3-yl)-1,3-dioxoisoindolin-4-yl)oxy)acetamido)pentyl)-9,10-difluoro-3-methyl-7-oxo-1H,3H,7H-[1,3]oxazino[5,4,3-ij]quinoline-6-carboxamide (17b)**

**General Procedure 1A.** Reaction scale: Thalidomide-O-acetamido-C5-amine 28 mg (53.3 µmol, 1.0 eq) and 9,10-difluoro-3-methyl-7-oxo-1H,3H,7H-[1,3]oxazino[5,4,3-ij]quinoline-6-carboxylic acid 15 mg (53.3 µmol,1.0 eq). Purified by pTLC (7% MeOH/DCM) to afford **17b** as an off-white solid (15.0 mg, 45%). $^{1}$H NMR (600 MHz, CDCl3) δ 10.10 (s, 1H), 9.89 (s, 1H), 8.80 (s, 1H), 7.85 – 7.82 (m, 1H), 7.73 (s, 1H), 7.71 – 7.63 (m, 1H), 7.53 (dd, J = 7.3, 2.2 Hz, 1H), 7.22 – 7.18 (m, 1H), 5.09 – 4.93 (m, 1H), 4.74 – 4.53 (m, 3H), 4.50 – 4.48 (m, 1H), 4.42 – 4.38 (m, 1H), 3.56 – 3.54 (m, 2H), 3.42 – 3.38 (m, 1H), 3.26 – 3.24 (m, 1H), 2.92 – 2.71 (m, 3H), 2.17 – 2.14 (m, 1H), 1.71 – 1.55 (m, 6H), 1.54 – 1.38 (m, 4H). $^{13}$C NMR (151 MHz, DMSO) δ 174.48, 173.29, 170.37, 167.20, 167.14, 165.99, 164.05, 155.45, 149.58, 149.50, 147.94, 147.87, 145.97, 142.24, 142.13, 140.58, 140.47, 137.39, 135.78, 135.70, 133.44, 125.20, 125.18, 123.21, 123.17, 120.84, 117.27, 116.48, 111.53, 104.03, 103.90, 69.34, 68.11, 54.60, 54.03, 49.27, 42.29, 38.75, 38.65, 31.42, 29.31, 29.06, 24.21, 22.45, 18.53, 18.19, 17.18, 12.95. HR-MS (ESI+) calc'd for $C_{33}H_{31}F_2N_5O_9$: 679.2090, found $[M+H]^+$ 680.2144.

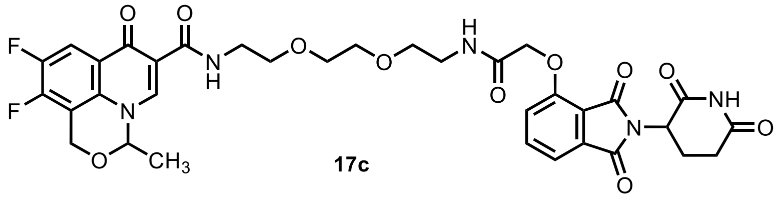

**17c**

**N-(2-(2-(2-(2-((2-(2,6-dioxopiperidin-3-yl)-1,3-dioxoisoindolin-4-yl)oxy)acetamido)ethoxy)ethoxy) ethyl)-9,10-difluoro-3-methyl-7-oxo-1H,3H,7H-[1,3]oxazino[5,4,3-ij]quinoline-6-carboxamide (17c)**

**General Procedure 1A.** Reaction scale: Thalidomide-O-acetamido-PEG2-C2-amine 18 mg (36.0 µmol, 1.0 eq) and 9,10-difluoro-3-methyl-7-oxo-1H,3H,7H-[1,3]oxazino[5,4,3-ij]quinoline-6-carboxylic acid 10.6 mg (37.9 µmol,1.0 eq). Purified by pTLC (7% MeOH/DCM) to afford **17c** as an off-white solid (7.0 mg, 38%). $^1$H NMR (600 MHz, $CDCl_3$) $^1$H NMR (600 MHz, $CDCl_3$) δ 9.97 (s, 1H), 8.73 (s, 1H), 7.77 – 7.74 (m, 1H), 7.71 – 7.55 (m, 2H), 7.46 – 7.44 (m, 1H), 7.13 – 7.11 (m, 1H), 4.94 – 4.92 (m, 1H), 4.65 – 4.51 (m, 2H), 4.50 – 4.46 (m, 1H), 4.42 – 4.38 (m, 1H), 4.32 – 4.28 (m, 1H), 3.68 – 3.47 (m, 10H), 2.87 – 2.68 (m, 2H), 2.07 – 2.05 (m, 1H), 1.52 – 1.48 (m, 4H), 1.48 – 1.36 (m, 1H). $^{13}$C NMR (151 MHz, DMSO) δ 174.41, 173.27, 170.35, 167.33, 167.20, 165.88, 164.21, 155.38, 149.59, 149.51, 147.95, 147.88, 146.16, 146.06, 142.26, 142.14, 140.60, 140.48, 137.37, 135.79, 135.72, 133.45, 125.19, 123.27, 123.22, 120.73, 117.15, 116.44, 111.42, 104.07, 103.94, 70.17, 70.12, 69.74, 69.36, 67.91, 54.59, 49.26, 38.99, 38.93, 31.41, 22.45, 18.20, HR-MS (ESI+) calc'd for $C_{34}H_{33}F_2N_5O_{11}$: 725.2145, found $[M+H]^+$ 726.2235.

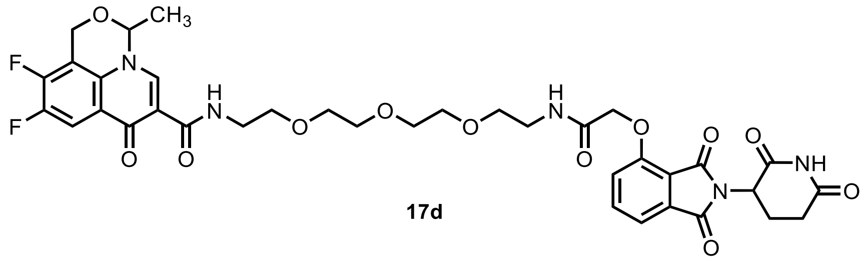

**17d**

**N-(1-((2-(2,6-dioxopiperidin-3-yl)-1,3-dioxoisoindolin-4-yl)oxy)-2-oxo-6,9,12-trioxa-3-azatetradecan-14-yl)-9,10-difluoro-3-methyl-7-oxo-1H,3H,7H-[1,3]oxazino[5,4,3-ij]quinoline-6-carboxamide (17d)**

**General Procedure 1A.** Reaction scale: Thalidomide-O-acetamido-PEG3-C2-amine 20 mg (36.9 µmol, 1.0 eq) and 9,10-difluoro-3-methyl-7-oxo-1H,3H,7H-[1,3]oxazino[5,4,3-ij]quinoline-6-carboxylic acid 11 mg (38.7 µmol,1.0 eq). Purified by pTLC (7% MeOH/DCM) to afford **17d** as an off-white solid (12.0 mg, 45%). $^1$H NMR (600 MHz, $CD_3OD$) δ 10.24 (s, 1H), 8.77 (s, 1H), 8.54 (s, 1H), 7.83 – 7.64 (m, 2H), 7.52 – 7.44 (m, 1H), 7.44 – 7.33 (m, 1H), 5.19 – 5.05 (m, 2H), 4.75 – 4.73 (m, 4H), 4.58 – 4.54 (m, 2H), 4.45 (m, 1H), 3.71 – 3.54 (m, 10H), 3.49 – 3.38 (m, 4H), 2.89 – 2.86 (m, 1H), 2.79 – 2.61 (m, 2H), 2.18 – 2.07 (m, 1H), 1.54 (d, $J$ = 6.9 Hz, 3H). $^{13}$C NMR (151 MHz, DMSO) δ 174.40, 173.28, 170.36, 167.35, 167.20, 165.91, 164.23, 155.40, 149.59, 149.52, 147.96, 147.89, 146.06, 142.26, 142.15, 140.60, 140.49, 137.40, 135.80, 135.73, 133.46, 125.19, 120.74, 117.17, 116.48, 111.41, 104.05, 103.92, 70.25, 70.17, 70.10, 69.72, 69.33, 69.26, 67.90, 54.60, 49.26, 38.99, 38.88, 31.40, 22.46, 18.19. HR-MS (ESI+) calc'd for $C_{36}H_{37}F_2N_5O_{12}$: 769.2407, found $[M+H]^+$ 770.2470.

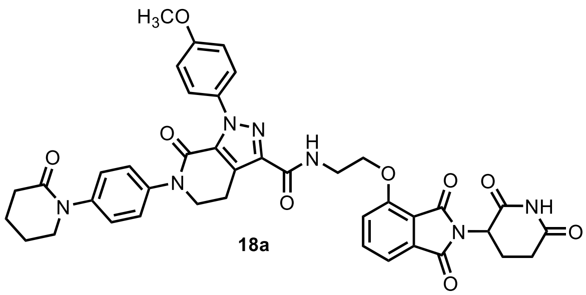

**18a**

**N-(2-((2-(2,6-dioxopiperidin-3-yl)-1,3-dioxoisoindolin-4-yl)oxy)ethyl)-1-(4-methoxyphenyl)-7-oxo-6-(4-(2-oxopiperidin-1-yl)phenyl)-4,5,6,7-tetrahydro-1H-pyrazolo[3,4-c]pyridine-3-carboxamide (18a)**

**General Procedure 1A.** Reaction scale: Thalidomide-O-C2-amine 10.3 mg (32.6 µmol, 1.0 eq) and 1-(4-methoxyphenyl)-7-oxo-6-(4-(2-oxopiperidin-1-yl)phenyl)-4,5,6,7-tetrahydro-1H-pyrazolo[3,4-c]pyridine-3-carboxylic acid 15 mg (32.6 µmol,1.0 eq). Purified by pTLC (7% MeOH/DCM) to afford **18a** as an off-white solid (7.0 mg, 38%). $^{1}H$ NMR (600 MHz, $CDCl_3$) δ 8.28 (s, 1H), 7.73 – 7.61 (m, 2H), 7.49 – 7.46 (m, 3H), 7.35 – 7.30 (m, 2H), 7.28 (d, *J* = 8.5 Hz, 1H), 7.25 – 7.21 (m, 2H), 6.95 – 6.83 (m, 2H), 4.90 – 4.87 (m, 1H), 4.37 (t, *J* = 5.4 Hz, 2H), 4.14 – 4.02 (m, 2H), 3.93 –3.89 (m, 2H), 3.80 (s, 3H), 3.67 – 3.55 (m, 2H), 3.35 (t, *J* = 6.7 Hz, 2H), 3.12 – 3.08 (m, 2H), 2.85 – 2.79 (m, 1H), 2.78 – 2.62 (m, 2H), 2.54 (t, *J* = 6.1 Hz, 2H), 2.07 – 1.99 (m, 1H), 1.94 – 1.92 (m, 3H), 1.83 (s, 2H), 1.43 – 1.31 (m, 4H). HR-MS (ESI+) calc'd for $C_{40}H_{37}N_7O_9$; 759.7760, $^{13}C$ NMR (151 MHz, DMSO) δ 173.29, 170.42, 169.35, 167.28, 165.73, 162.01, 159.64, 157.04, 156.09, 141.85, 141.52, 140.23, 137.52, 133.74, 133.50, 132.93, 127.34, 126.81, 126.47, 125.58, 120.49, 116.97, 116.01, 113.86, 67.42, 55.94, 54.02, 51.40, 51.31, 49.21, 42.28, 37.99, 33.05, 31.41, 23.46, 22.47, 21.42, 21.37, 18.53, 17.18, 12.94. HR-MS (ESI+) calc'd for $C_{40}H_{37}N_7F_2N_5O_{12}$: 759.2653, found $[M+H]^+$ 760.2727.

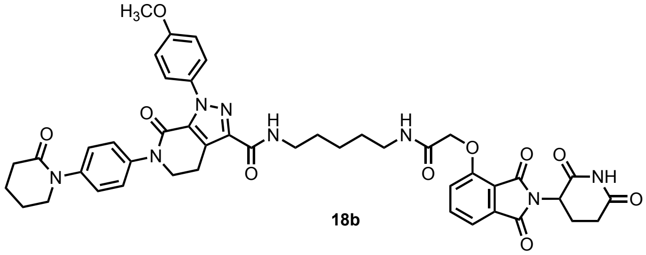

**18b**

**N-(5-(2-((2-(2,6-dioxopiperidin-3-yl)-1,3-dioxoisoindolin-4-yl)oxy)acetamido)pentyl)-1-(4-methoxyphenyl)-7-oxo-6-(4-(2-oxopiperidin-1-yl)phenyl)-4,5,6,7-tetrahydro-1H-pyrazolo[3,4-c]pyridine-3-carboxamide (18b)**

**General Procedure 1A.** Reaction scale: Thalidomide-O-acetamido-C5-amine 17 mg (32.5 µmol, 1.0 eq) and 1-(4-methoxyphenyl)-7-oxo-6-(4-(2-oxopiperidin-1-yl)phenyl)-4,5,6,7-tetrahydro-1H-pyrazolo[3,4-c]pyridine-3-carboxylic acid 15 mg (32.5 µmol,1.0 eq). Purified by pTLC (7% MeOH/DCM) to afford **18b** as an off-white solid (14 mg, 42%). 1H NMR (600 MHz, CDCl3) δ 9.29 (s, 1H), 7.69 – 7.64 (m, 1H), 7.52 – 7.48 (m, 1H), 7.48 (d, J = 7.3 Hz, 1H), 7.40 – 7.35 (m, 2H), 7.29 – 7.25 (m, 2H), 7.20 (s, 1H), 7.20 – 7.15 (m, 2H), 7.12 (d, J = 8.4 Hz, 1H), 7.03 – 7.00 (m, 1H), 6.88 – 6.82 (m, 1H), 4.98 – 4.88 (m, 1H), 4.59 – 4.57 (m, 2H), 4.09 – 4.00 (m, 2H), 3.74 (s, 2H), 3.60 – 3.48 (m, 2H), 3.44 – 3.42 (m, 2H), 3.39 – 3.18 (m, 3H), 3.10 – 3.03 (m, 1H), 2.84 – 2.65 (m, 2H), 2.48 (t, J = 6.1 Hz, 2H), 2.12 – 2.00 (m, 1H), 1.98 (s, 1H), 1.92 – 1.73 (m, 4H), 1.65 – 1.53 (m, 2H), 1.48 – 1.42 (m, 2H), 1.39 – 1.27 (m, 2H), 1.19 (t, J = 7.2 Hz, 1H). $^{13}C$ NMR (151 MHz, DMSO) δ 173.28, 170.38, 169.34, 167.22, 167.12, 166.00, 161.58, 159.59, 157.08, 155.52, 141.96, 141.83, 140.26, 137.42, 133.49, 133.40, 132.99, 127.36, 126.80, 126.46, 125.41, 120.84, 117.26, 116.51, 113.83, 68.09, 55.94, 54.02, 51.40, 51.31, 49.27, 42.28, 38.77, 33.05, 31.41, 29.37, 29.18, 24.22, 23.46, 22.46, 21.43, 21.37, 18.53, 17.18, 12.94. HR-MS (ESI+) calc'd for $C_{45}H_{46}N_8O_{10}$: 858.9090, found [M+H]+ 859.3403.

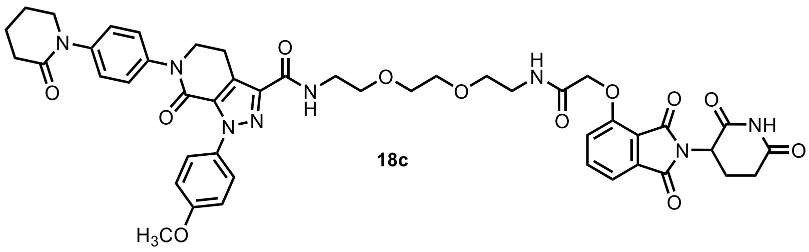

**18c**

**N-(2-(2-(2-(2-((2-(2,6-dioxopiperidin-3-yl)-1,3-dioxoisoindolin-4-yl)oxy)acetamido)ethoxy)ethoxy)ethyl)-1-(4-methoxyphenyl)-7-oxo-6-(4-(2-oxopiperidin-1-yl)phenyl)-4,5,6,7-tetrahydro-1H-pyrazolo[3,4-c]pyridine-3-carboxamide (18c)**

**General Procedure 1A.** Reaction scale: Thalidomide-O-acetamido-PEG2-C2-amine 13 mg (26.0 μmol, 1.0 eq) and 1-(4-methoxyphenyl)-7-oxo-6-(4-(2-oxopiperidin-1-yl)phenyl)-4,5,6,7-tetrahydro-1H-pyrazolo[3,4-c]pyridine-3-carboxylic acid 13 mg (26.0 μmol,1.0 eq). Purified by pTLC (7% MeOH/DCM) to afford **18c** as an off-white solid (11 mg, 48%). $^{1}$H NMR (600 MHz, $CDCl_3$) δ 8.91 (s, 1H), 7.69 – 7.62 (m, 1H), 7.56 – 7.51 (m, 1H), 7.47 (d, *J* = 7.3 Hz, 1H), 7.38 (d, *J* = 7.2, Hz, 2H), 7.28 – 7.23 (m, 2H), 7.19 – 7.15 (m, 1H), 7.10 (d, *J* = 8.4 Hz, 1H), 6.87 – 6.83 (m, 1H), 4.86 – 4.81 (m, 2H), 4.56 (s, 1H), 4.04 – 4.01 (m, 2H), 3.74 (s, 2H), 3.63 – 3.51 (m, 10H), 3.50 – 3.42 (m, 2H), 3.31 – 3.23 (m, 2H), 2.78 – 2.74 (m, 2H), 2.73 – 2.65 (m, 2H), 2.65 – 2.54 (m, 1H), 2.52 – 2.43 (m, 1H), 2.06 – 1.98 (m, 1H), 1.89 – 1.82 (m, 3H), 1.42 –1.34 (m, 2H), 1.32 –1.26 (m, 2H). $^{13}$C NMR (151 MHz, DMSO) δ 173.28, 170.37, 169.34, 167.36, 167.22, 165.92, 161.70, 159.61, 157.04, 155.42, 141.83, 141.66, 140.24, 137.41, 133.47, 132.94, 127.32, 126.80, 126.46, 125.46, 120.75, 117.18, 116.49, 113.85, 70.08, 69.94, 69.30, 69.25, 67.91, 55.93, 53.98, 51.38, 51.30, 49.27, 38.87, 38.62, 33.05, 31.41, 23.46, 22.46, 21.41, 21.37, 18.51, 17.18. HR-MS (ESI+) calc'd for $C_{46}H_{48}N_8O_{12}$: 904.9340, found $[M+H]^+$ 905.3461.

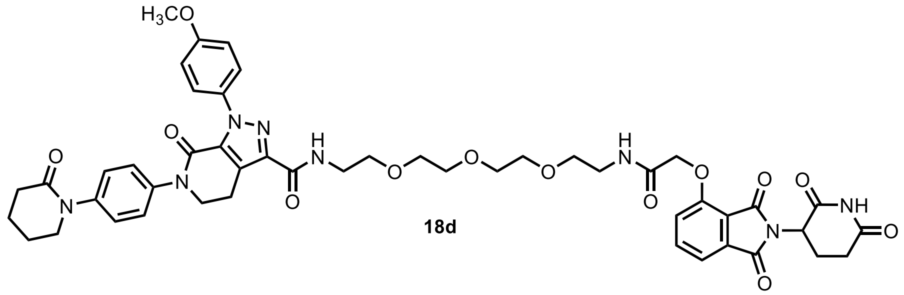

**18d**

**N-(1-((2-(2,6-dioxopiperidin-3-yl)-1,3-dioxoisoindolin-4-yl)oxy)-2-oxo-6,9,12-trioxa-3-azatetradecan-14-yl)-1-(4-methoxyphenyl)-7-oxo-6-(4-(2-oxopiperidin-1-yl)phenyl)-4,5,6,7-tetrahydro-1H-pyrazolo[3,4-c]pyridine-3-carboxamide (18d)**

**General Procedure 1A.** Reaction scale: Thalidomide-O-acetamido-PEG3-C2-amine 16 mg (29.5 μmol, 1.0 eq) and 1-(4-methoxyphenyl)-7-oxo-6-(4-(2-oxopiperidin-1-yl)phenyl)-4,5,6,7-tetrahydro-1H-pyrazolo[3,4-c]pyridine-3-carboxylic acid 13 mg (30.1 μmol,1.0 eq). Purified by pTLC (7% MeOH/DCM) to afford **18d** as an off-white solid (16 mg, 52%). $^{1}$H NMR (600 MHz, $CDCl_3$) δ 9.09 (s, 1H), 7.74 – 7.72 (m, 1H), 7.63 – 7.61 (m, 1H), 7.52 (d, *J* = 7.3 Hz, 1H), 7.50 – 7.46 (m, 2H), 7.46 – 7.42 (m, 1H), 7.32 (d, *J* = 8.7 Hz, 2H), 7.24 (d, *J* = 8.7 Hz, 2H), 7.17 (d, *J* = 8.4 Hz, 1H), 6.96 – 6.91 (m 2H), 4.86 – 4.80 (m, 1H), 4.62 (s, 2H), 4.14 – 4.01 (m, 2H), 3.80 (s, 3H), 3.73 – 3.62 (m, 9H), 3.62 – 3.54 (m, 7H), 3.53 – 3.44 (m, 1H), 3.36 – 3.32 (m, 2H), 2.78 – 2.63 (m, 2H), 2.58 – 2.53(m, 2H), 2.05 – 1.97 (m, 1H), 1.96 – 1.91 (m, 4H), 1.87 (s, 1H), 1.44 – 1.35 (m, 1H). $^{13}$C NMR (151 MHz, DMSO) δ 173.28, 170.37, 169.36, 167.37, 167.22, 165.92, 161.70, 159.61, 157.05, 155.42, 141.83, 141.67, 140.24, 137.43, 133.47, 132.95, 127.32, 126.80, 126.47, 125.47, 120.77, 117.19, 116.52, 113.85, 70.21, 70.08, 69.97, 69.27, 69.23, 67.92, 55.93, 51.38, 51.31, 49.27, 38.87, 38.62, 33.05, 31.41, 23.45, 22.45, 21.41, 21.36. HR-MS (ESI$^+$) calc'd for $C_{48}H_{52}N_8O_{13}$: 948.9870, found $[M+H]^+$ 949.3722.

### NMR Spectra

#### Headgroup intermediates

$^{1}H$ NMR **F2**

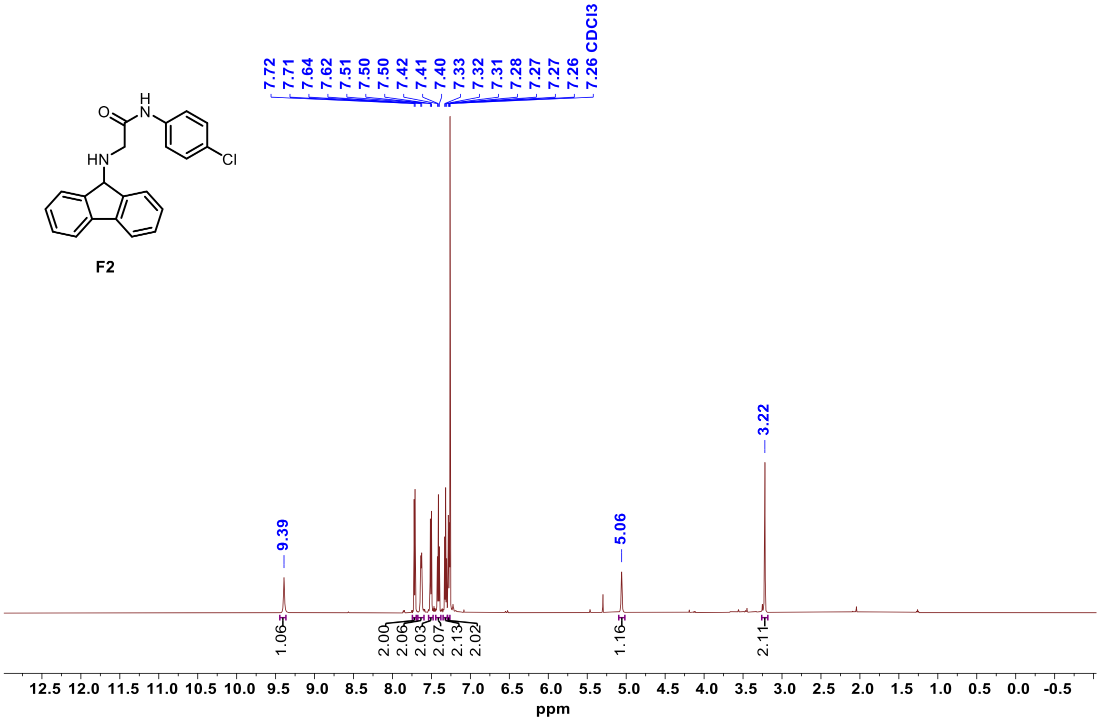


$^{13}C$ NMR **F2**

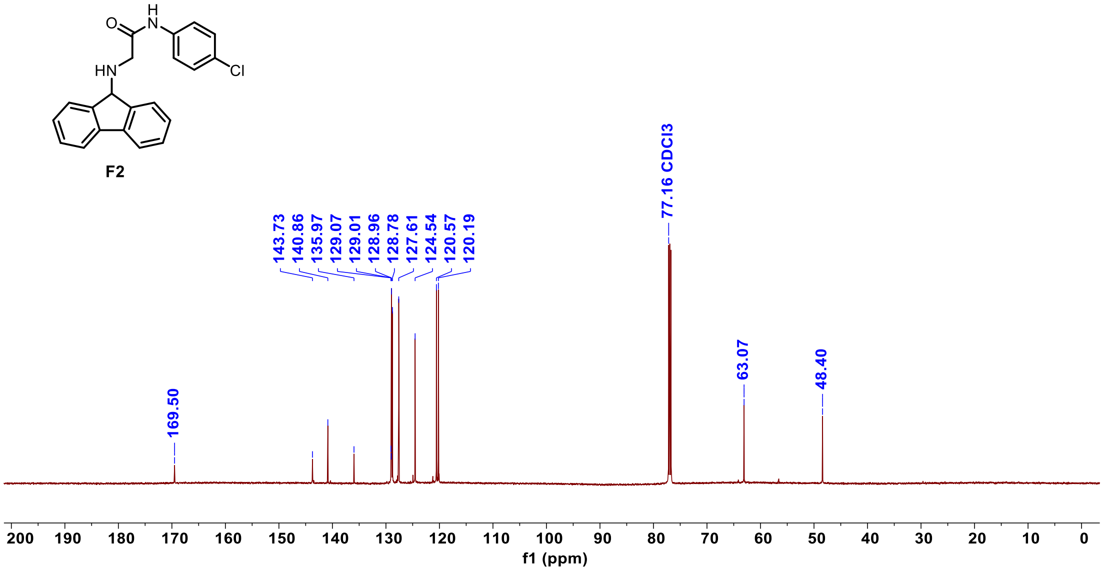

<sup></sup>

#### [1]H NMR **F4**

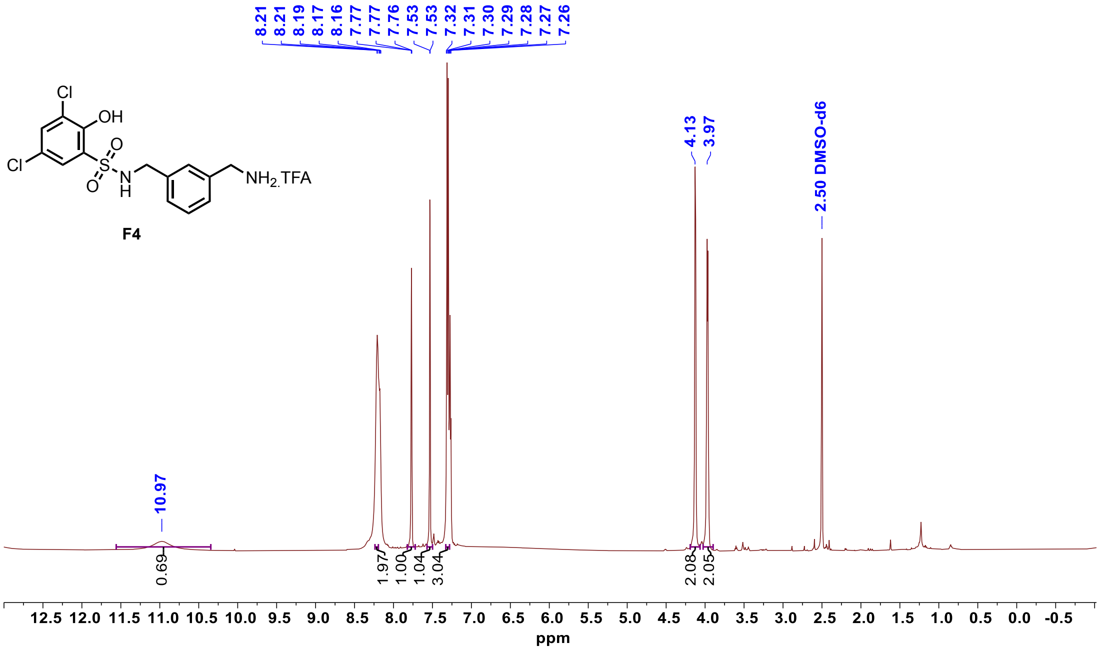


#### [1]H NMR **F7**

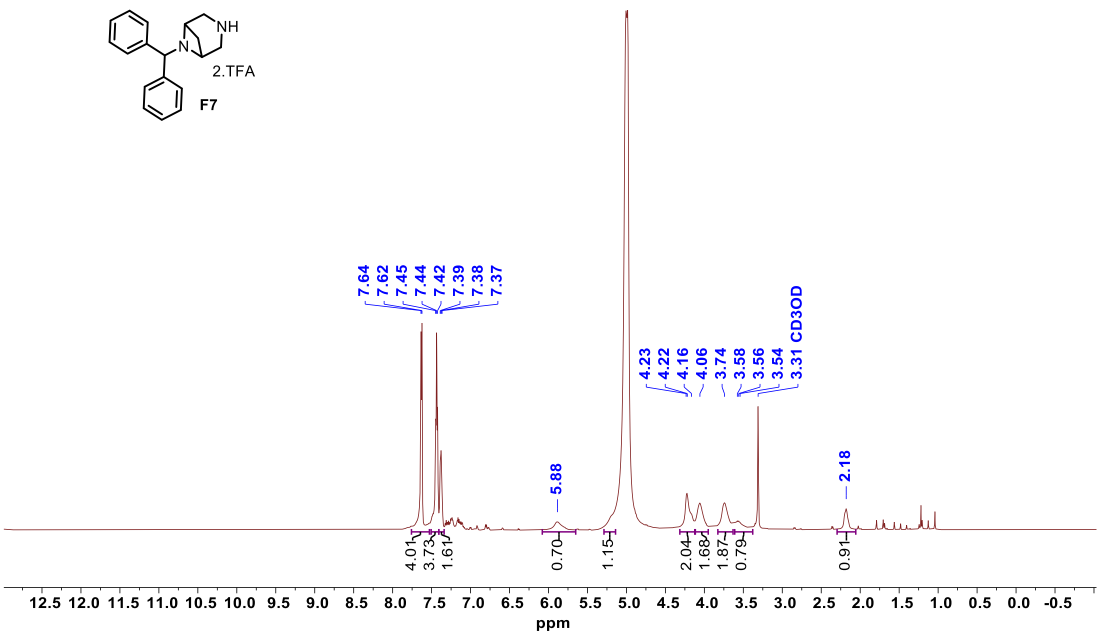

$^{1}$H NMR **F8**

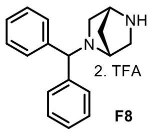
NH
N
2. TFA

**F8**

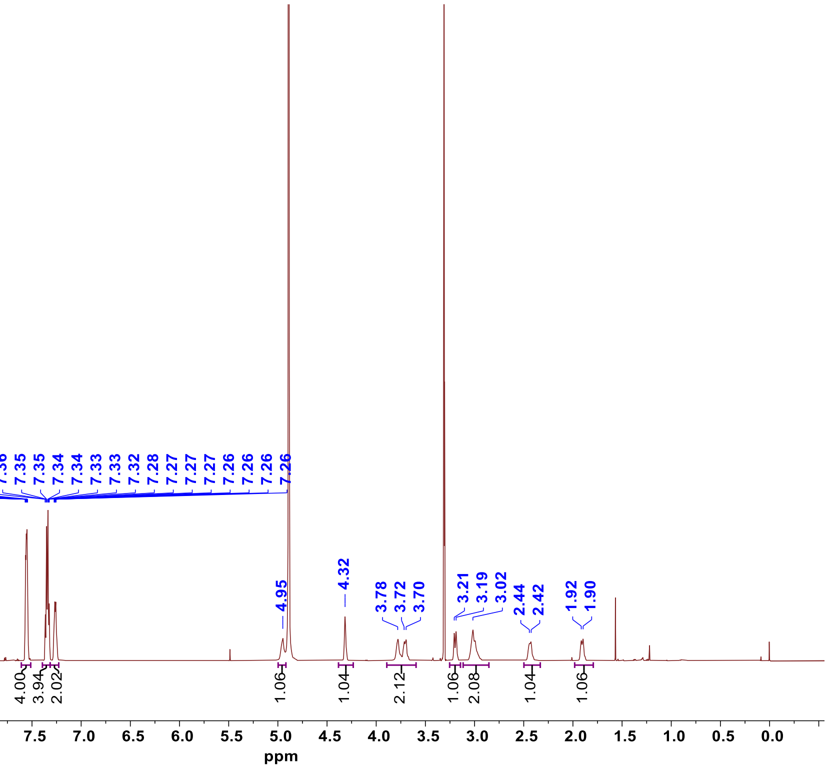
7.57
7.57
7.56
7.56
7.56
7.55
7.55
7.54
7.37
7.36
7.35
7.35
7.34
7.34
7.33
7.33
7.32
7.28
7.27
7.27
7.27
7.26
7.26
7.26
7.26
4.95
4.32
3.78
3.72
3.70
3.21
3.19
3.02
2.44
2.42
1.92
1.90
4.00
3.94
2.02
1.06
1.04
2.12
1.06
2.08
1.04
1.06
10.0
9.5
9.0
8.5
8.0
7.5
7.0
6.5
6.0
5.5
5.0
4.5
4.0
3.5
3.0
2.5
2.0
1.5
1.0
0.5
0.0
ppm

#### Final Compounds

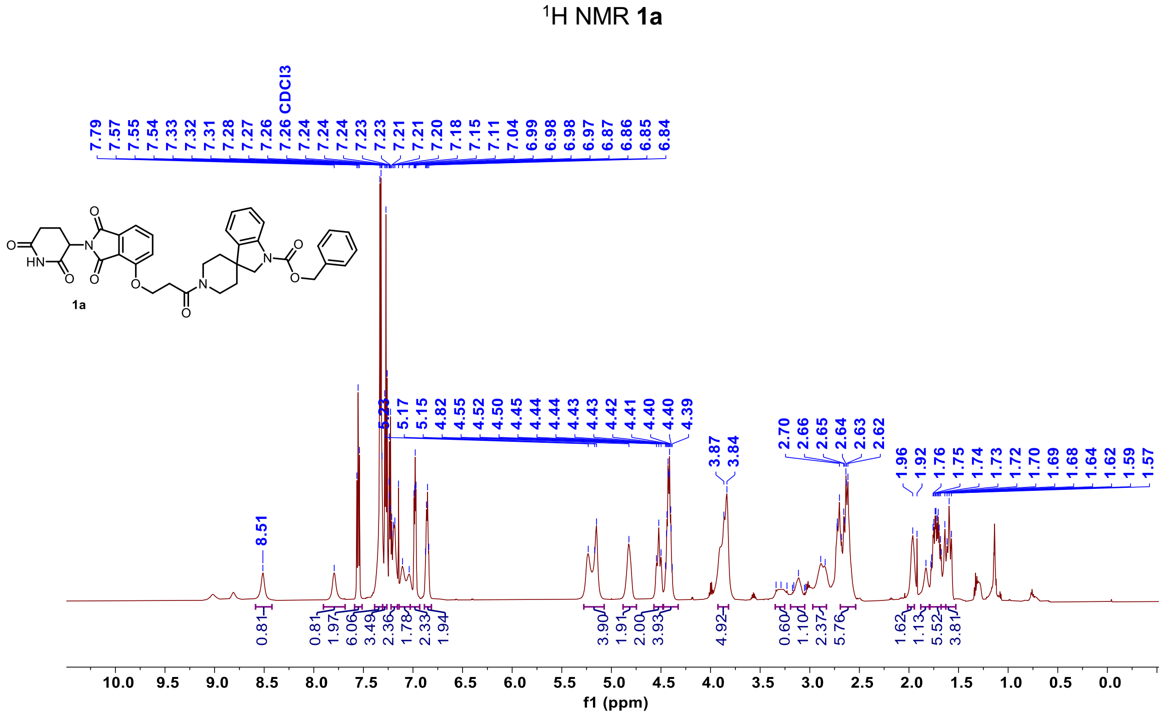


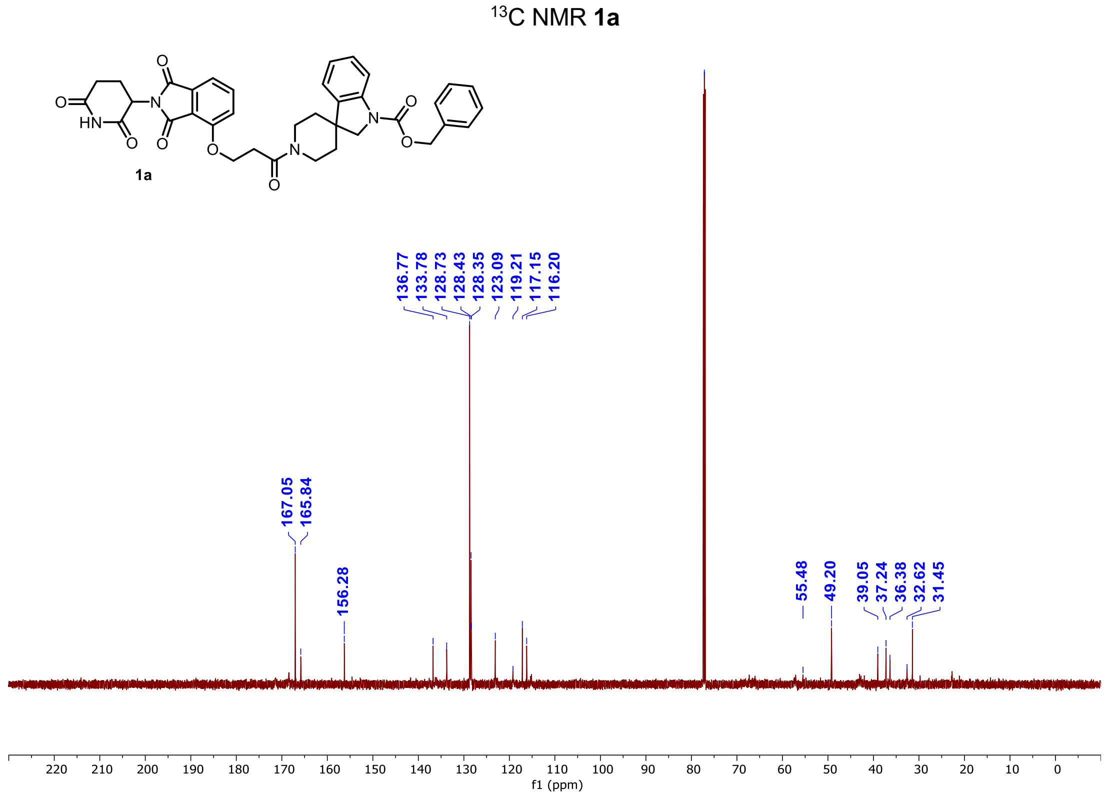

[1]H NMR **1b**

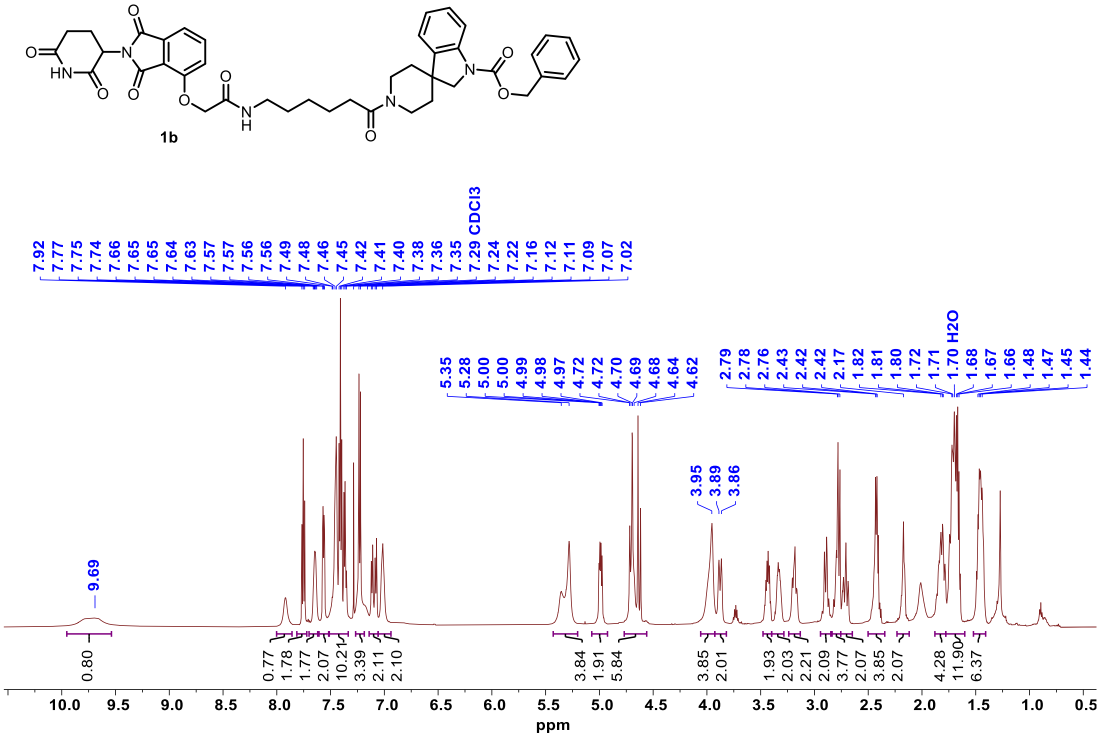


[13]C NMR **1b**

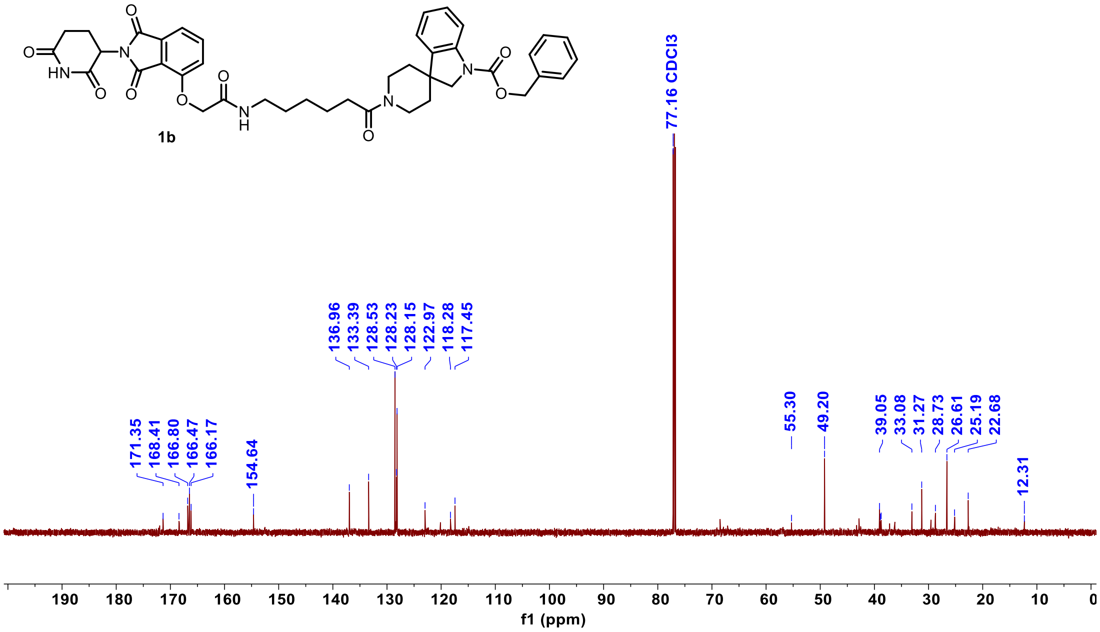

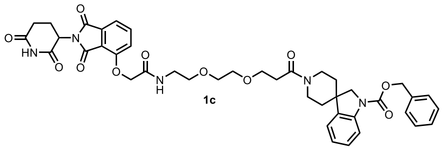


$^{1}$H NMR **1c**

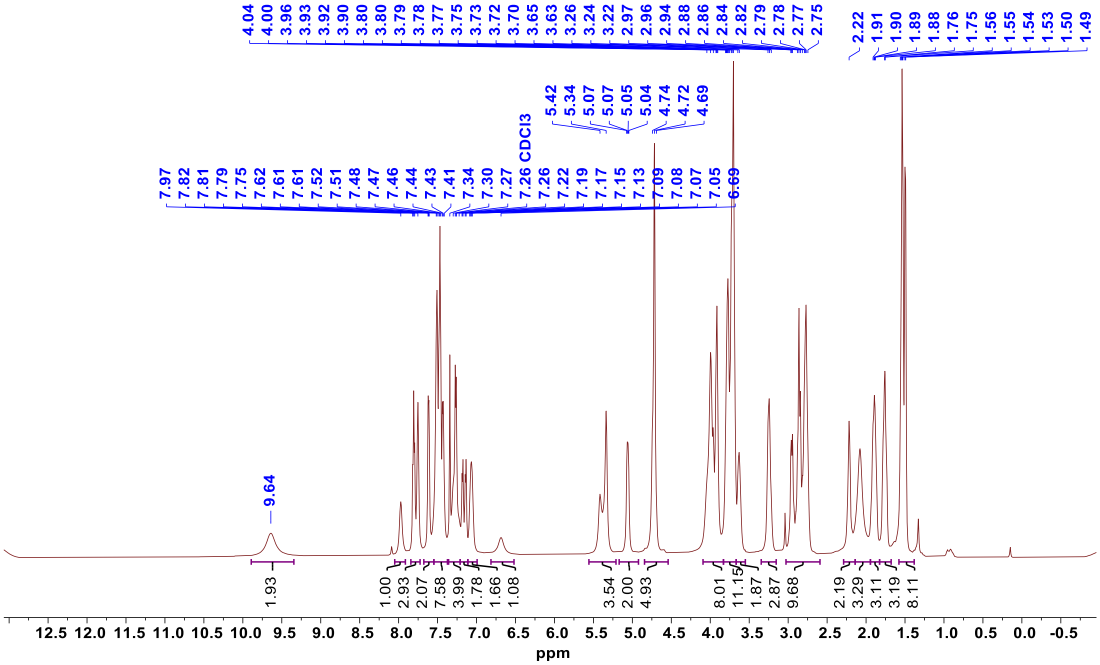


$^{13}$C NMR **1c**

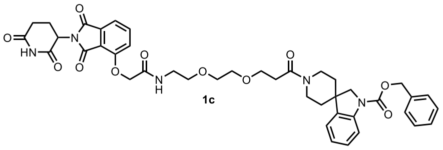


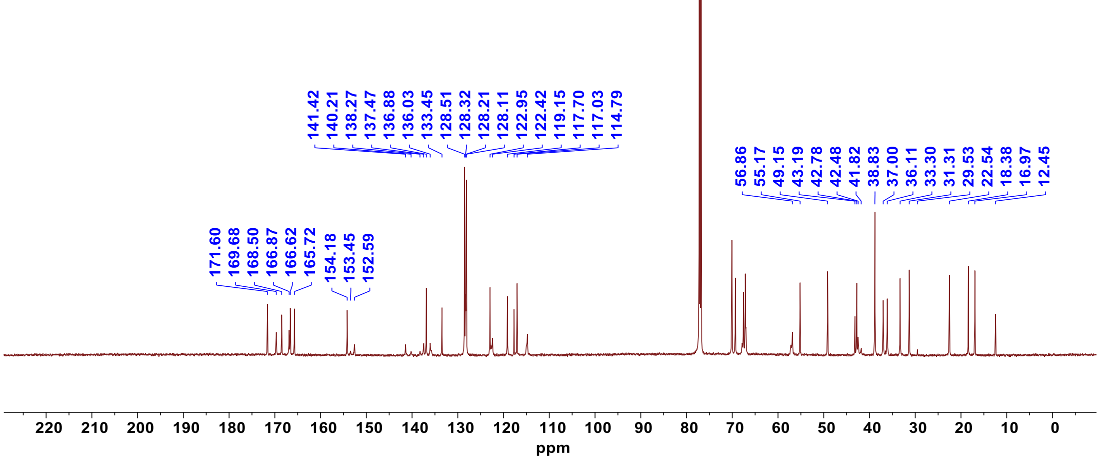

#### $^{1}$H NMR **1c-neg**

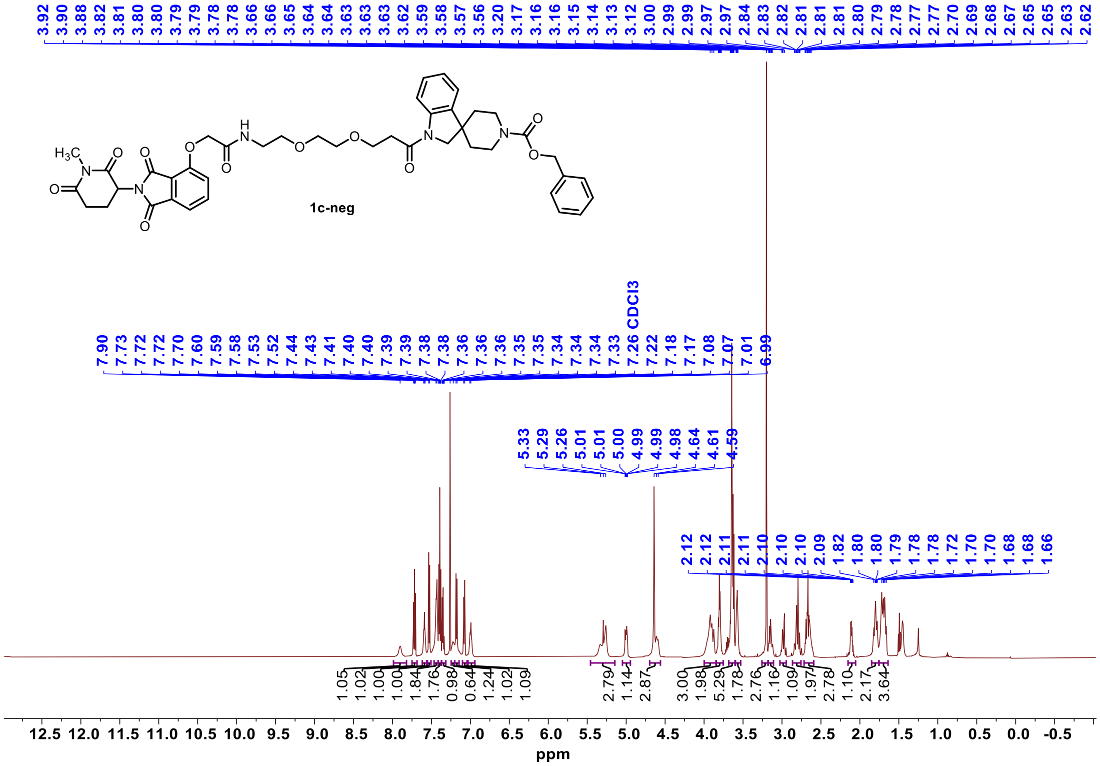


#### $^{13}$C NMR **1c-neg**

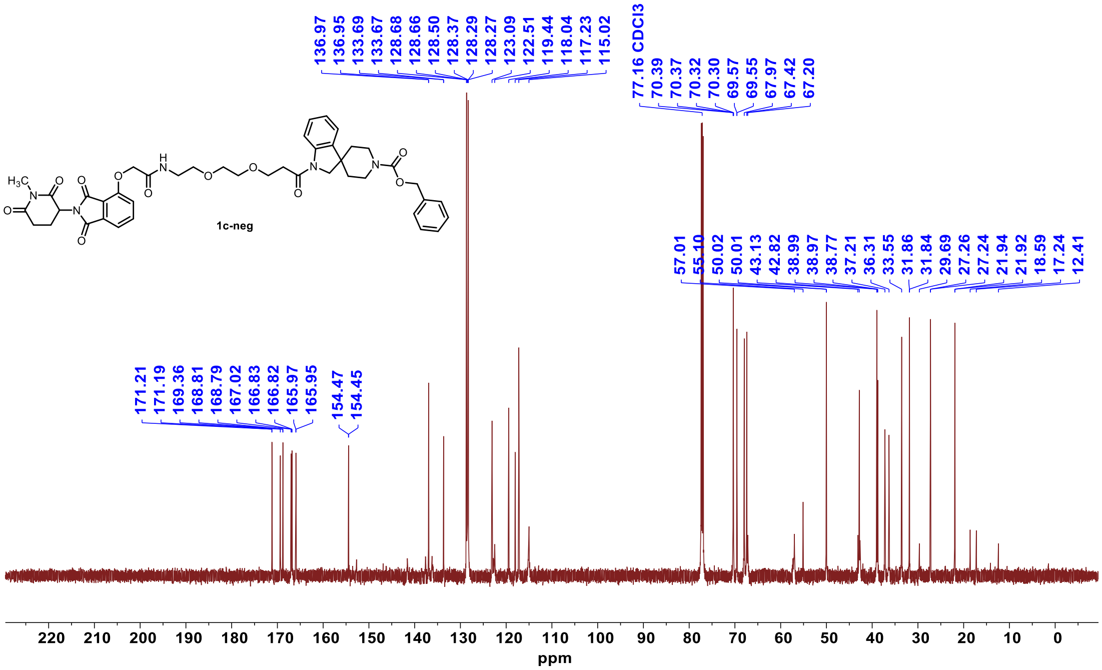

$^{1}$H NMR **1d**

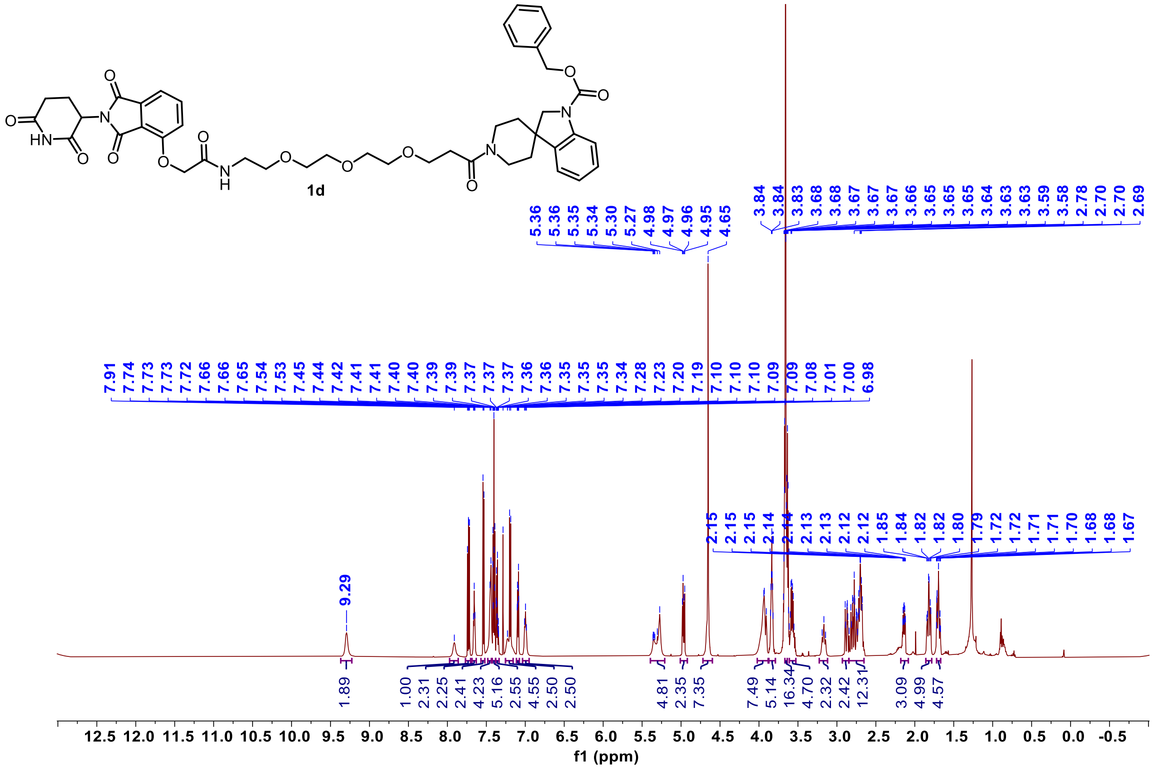

$^{13}$C NMR **1d**

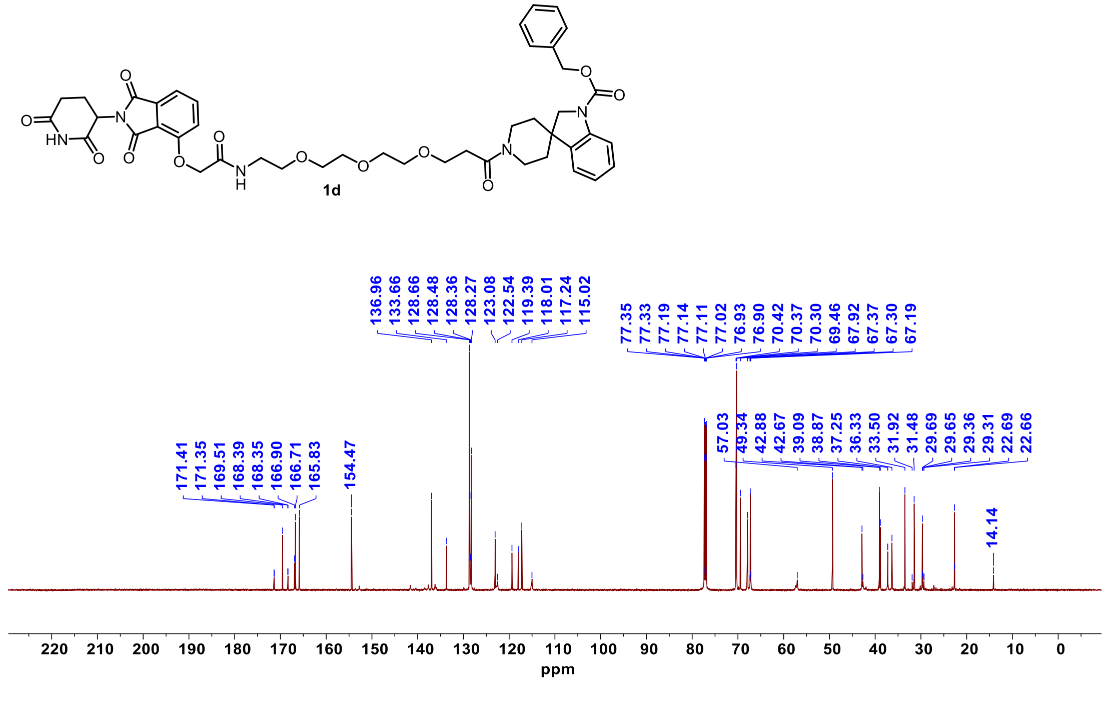

### $^{1}$H NMR **2a**

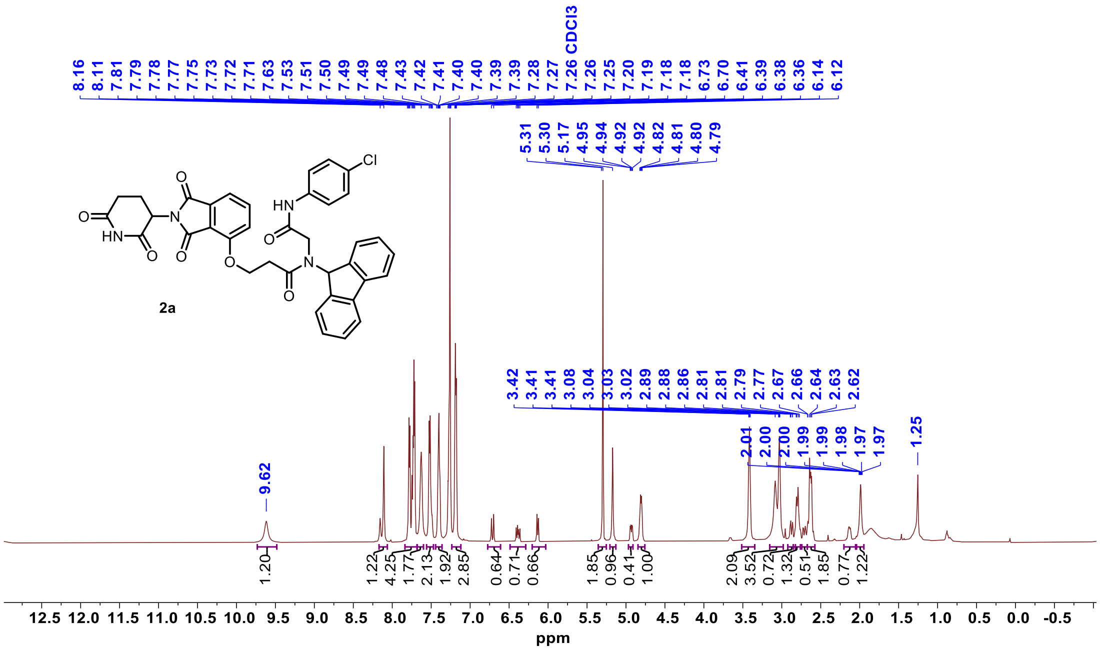


### $^{13}$C NMR **2a**

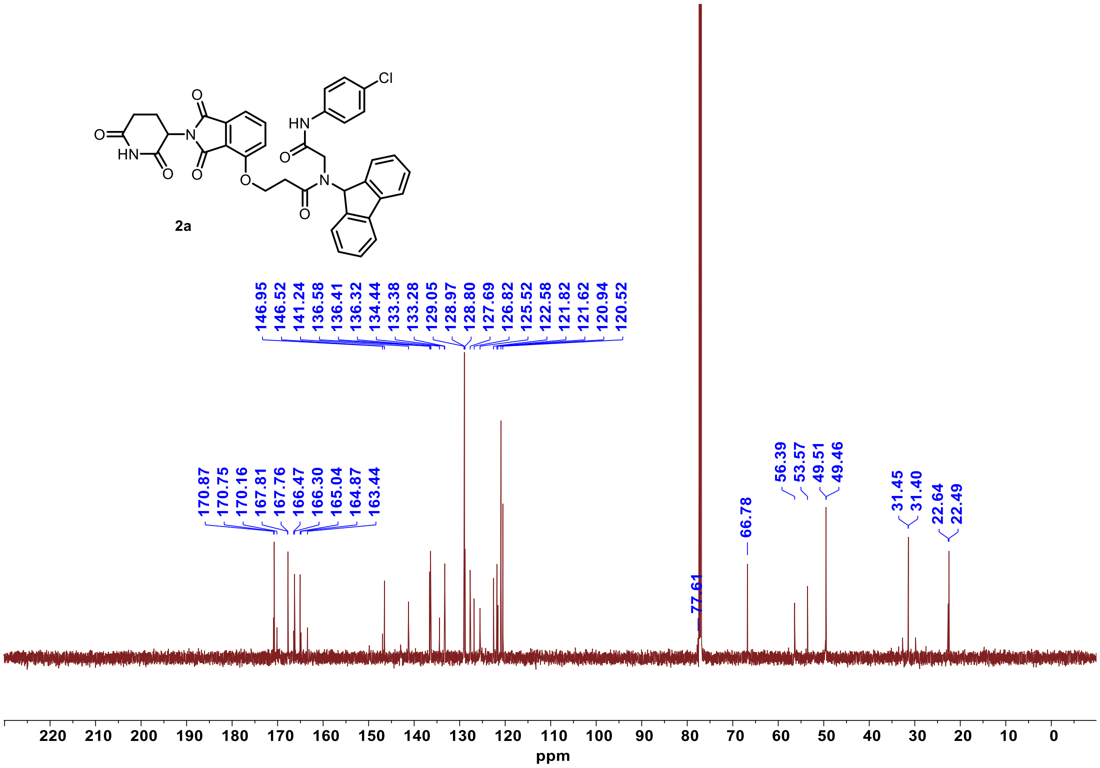

[1]H NMR **2a-neg**

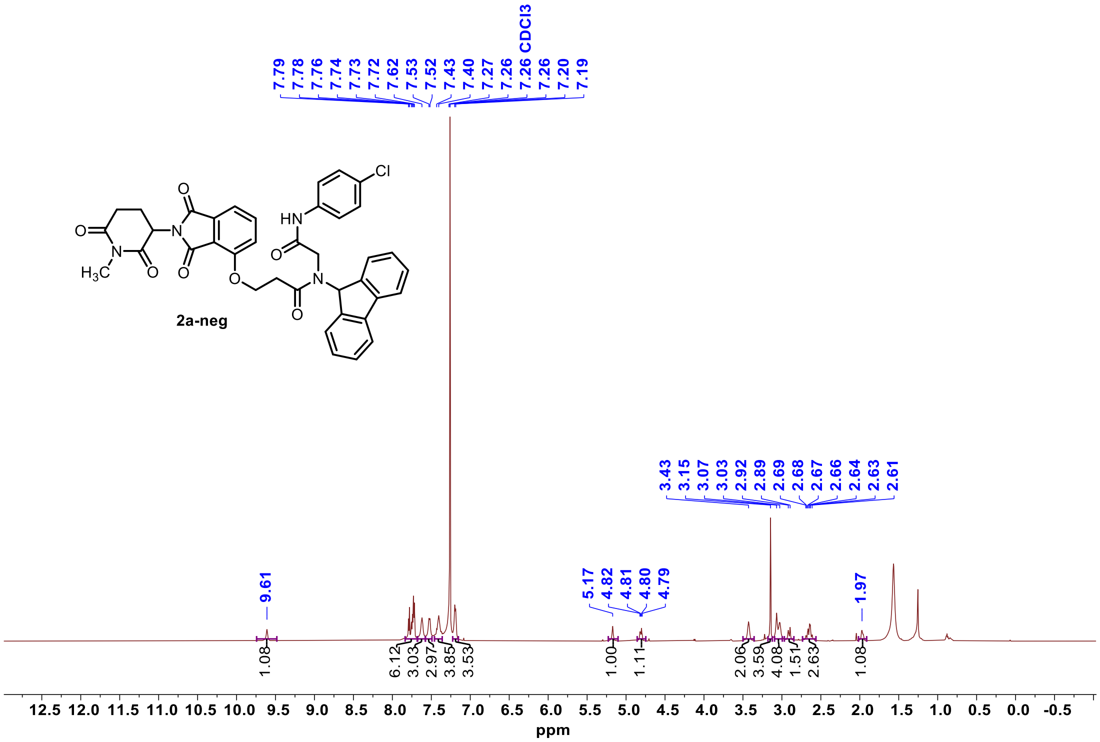


[13]C NMR **2a-neg**

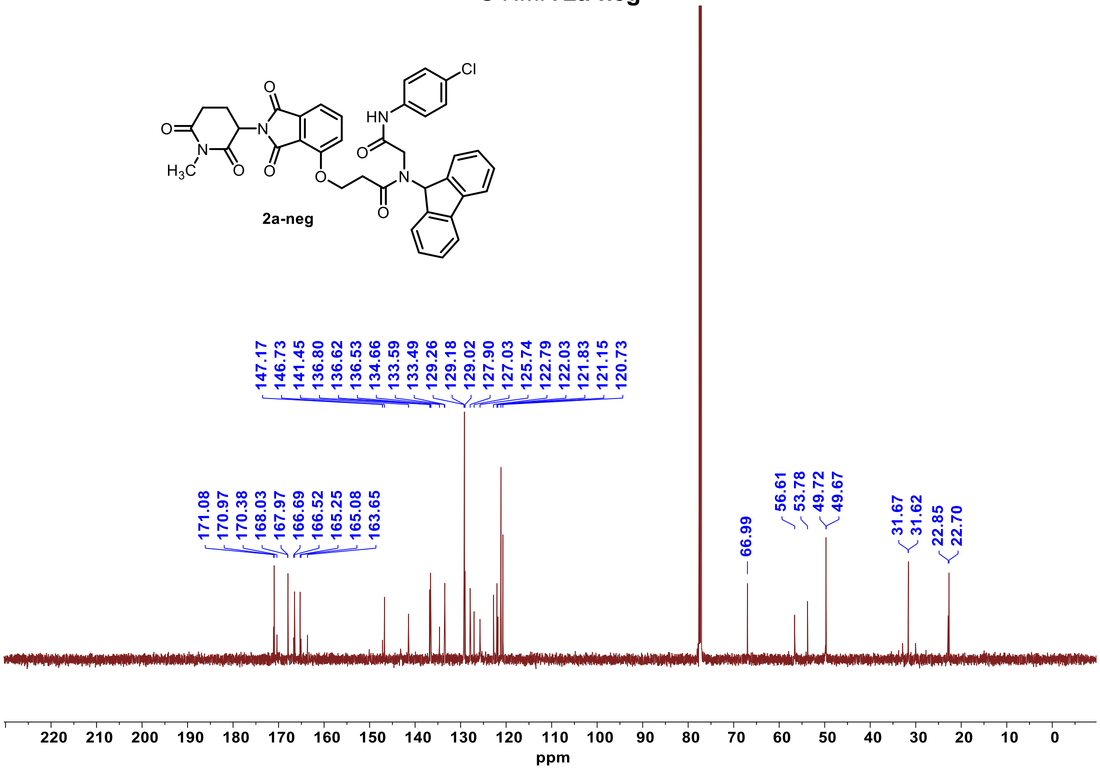

[1]H NMR **2b**

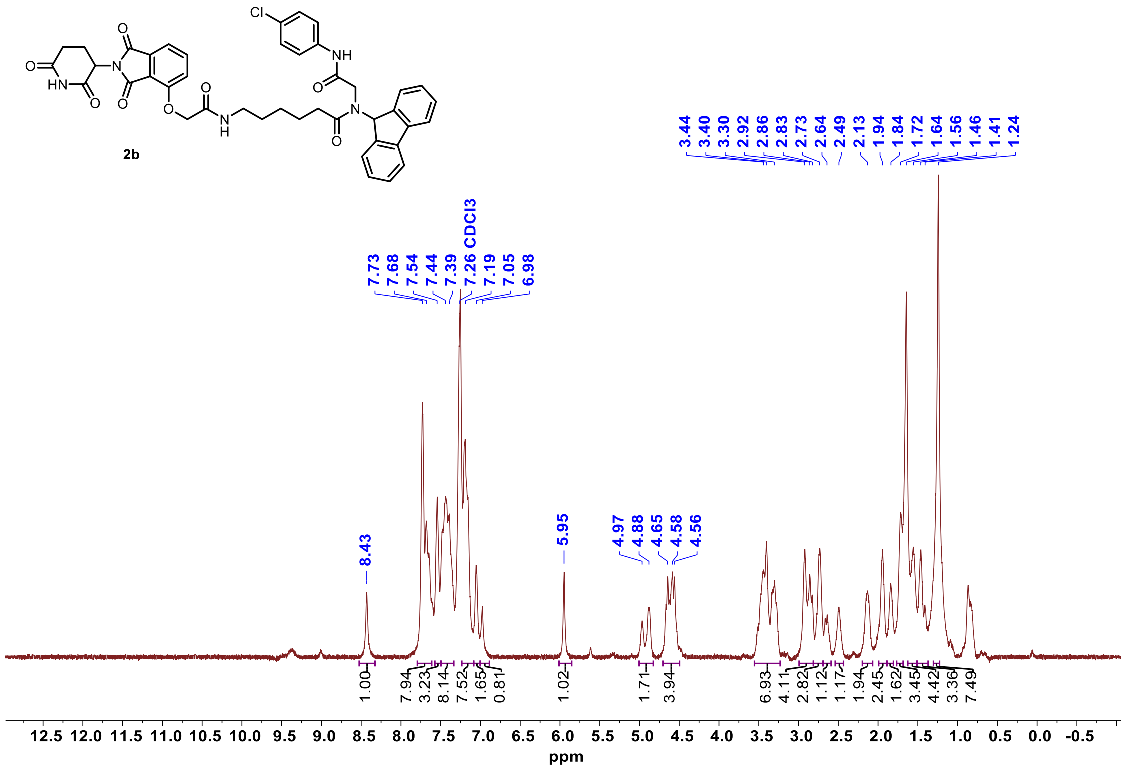


[13]C NMR **2b**

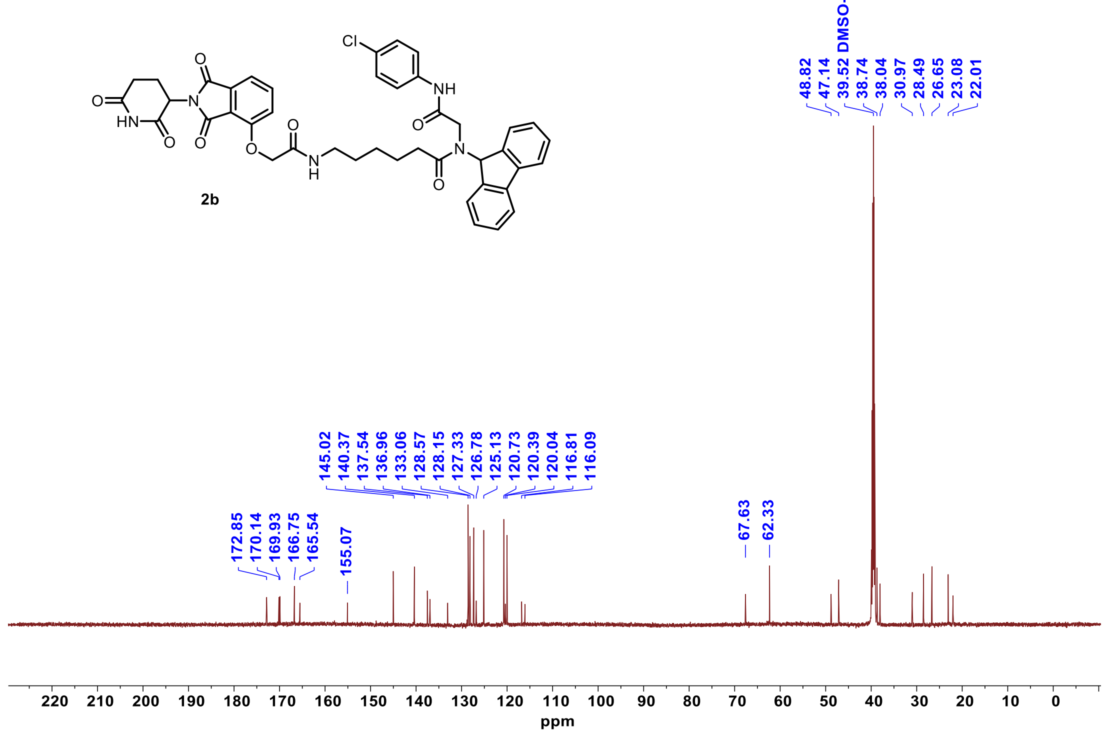

[1]H NMR **2c**

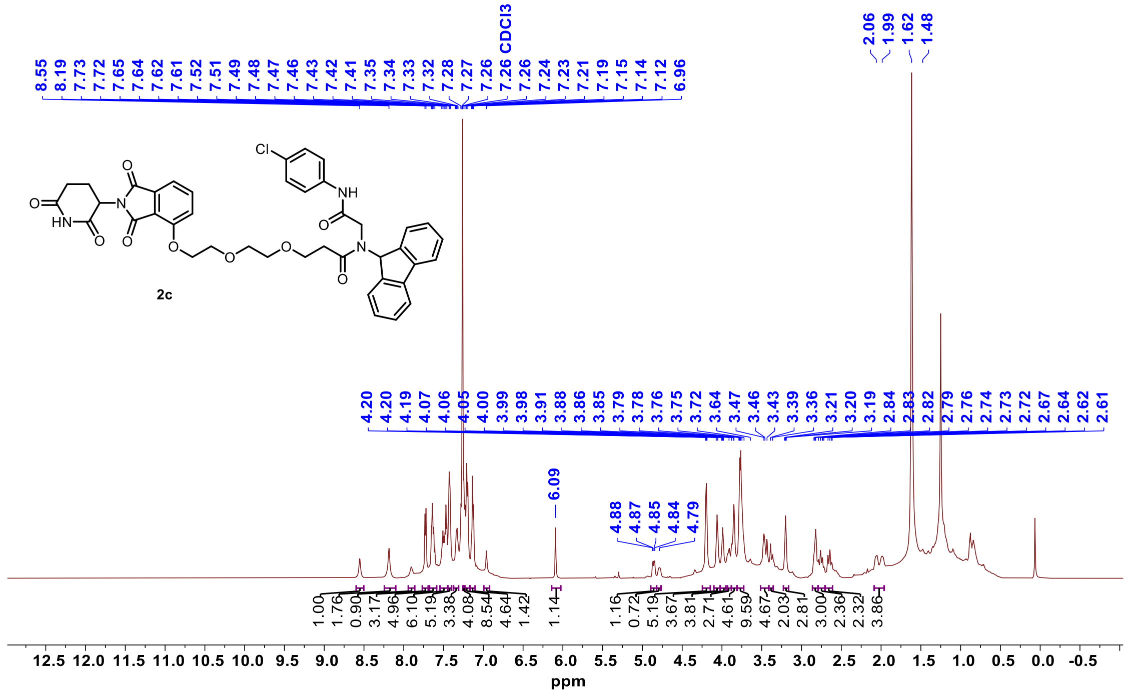


[13]C NMR **2c**

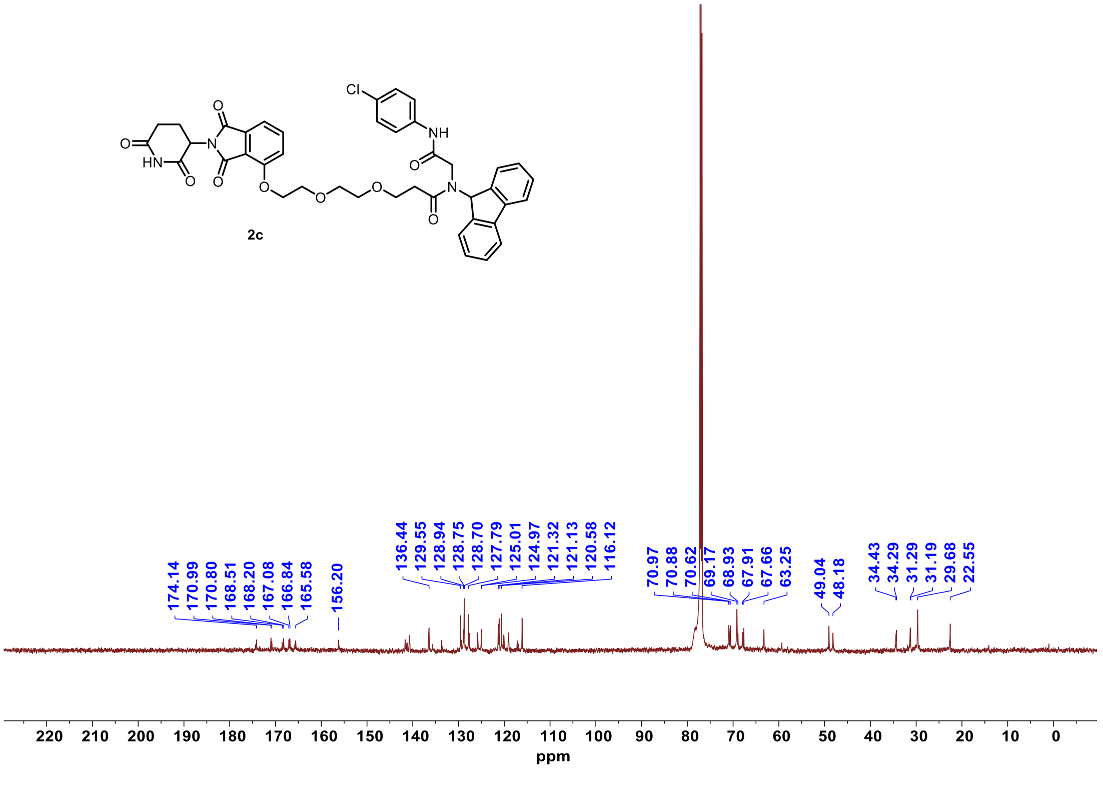

[1]H NMR **2d**

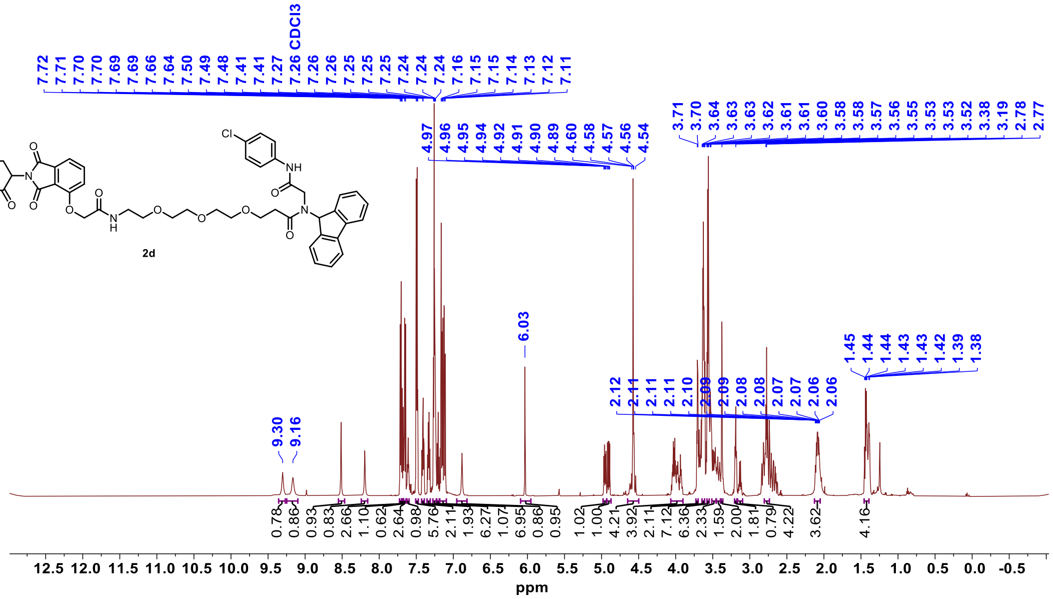


[13]C NMR **2d**

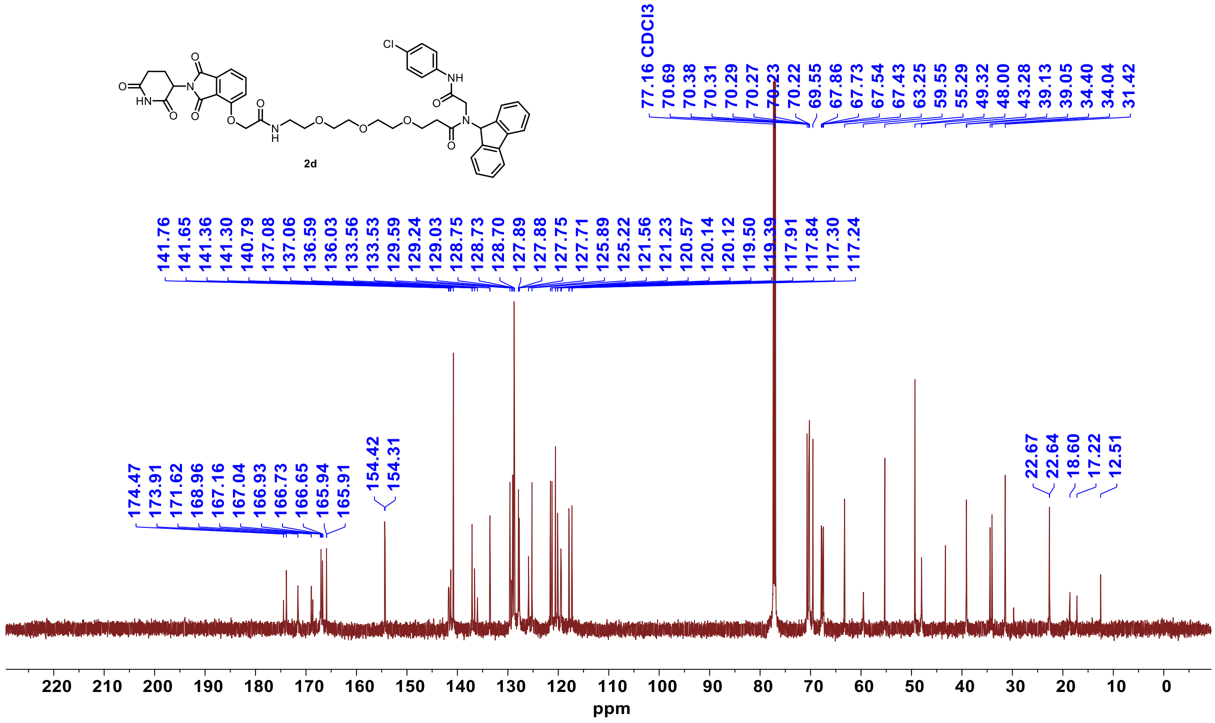

$^{1}$H NMR **2d-neg**

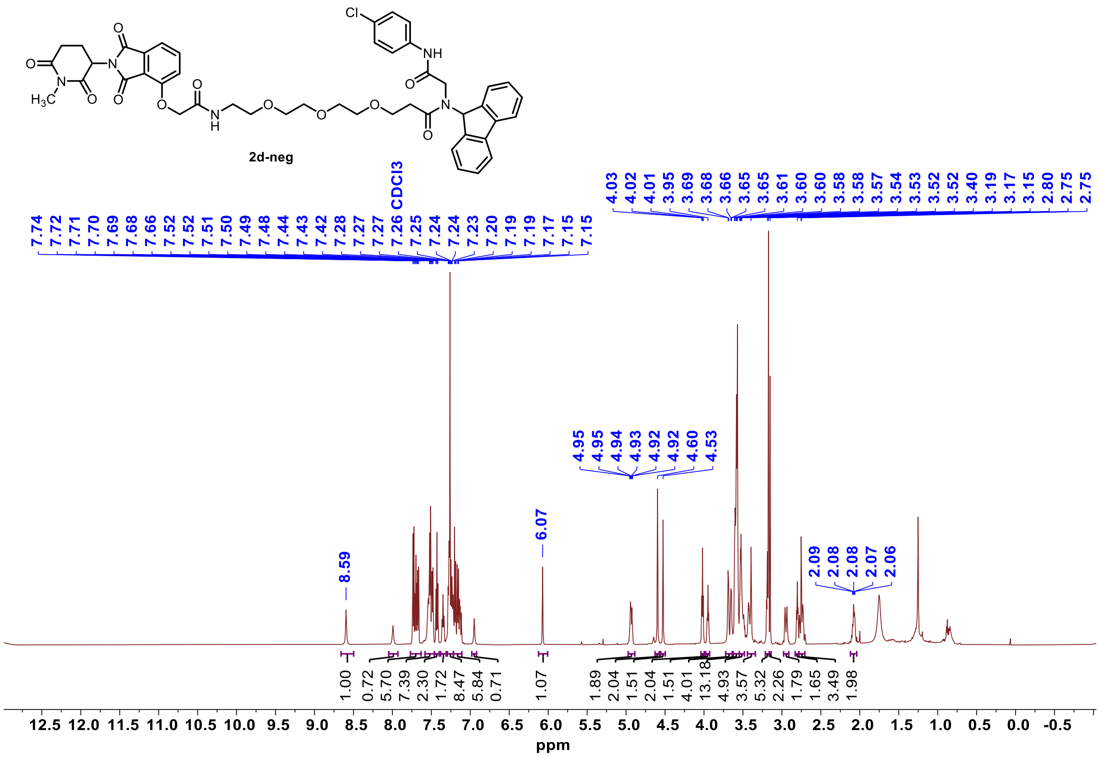


$^{13}$C NMR **2d-neg**

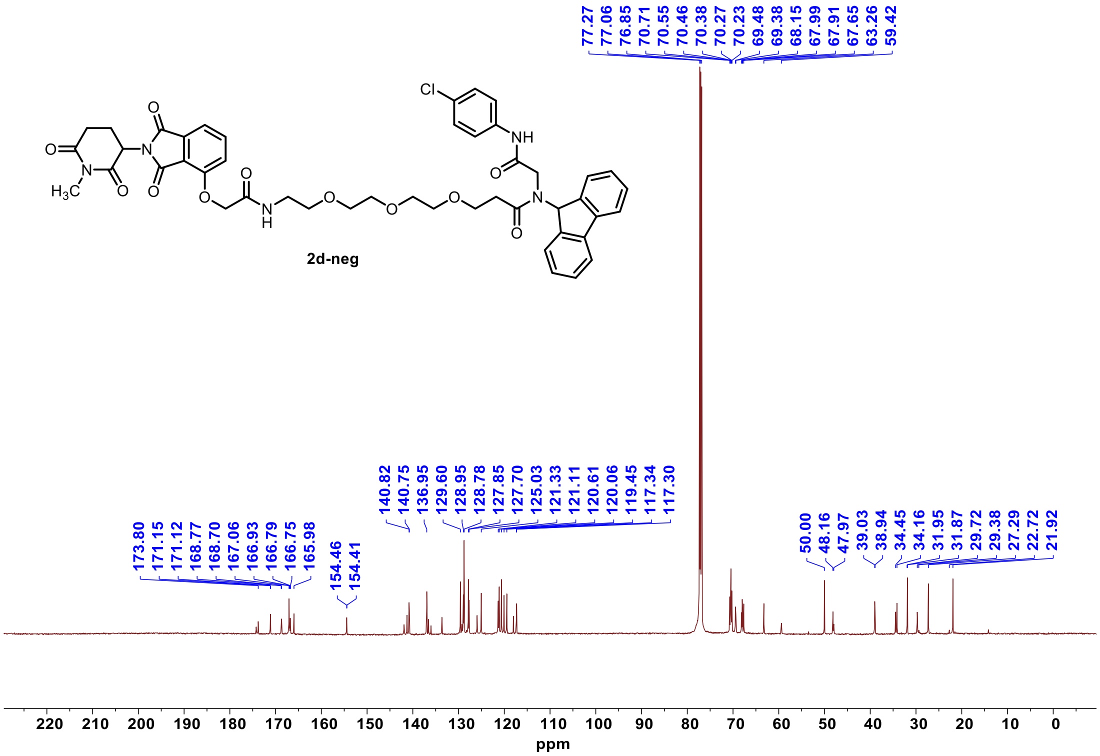

#### $^{1}$H NMR **3a**

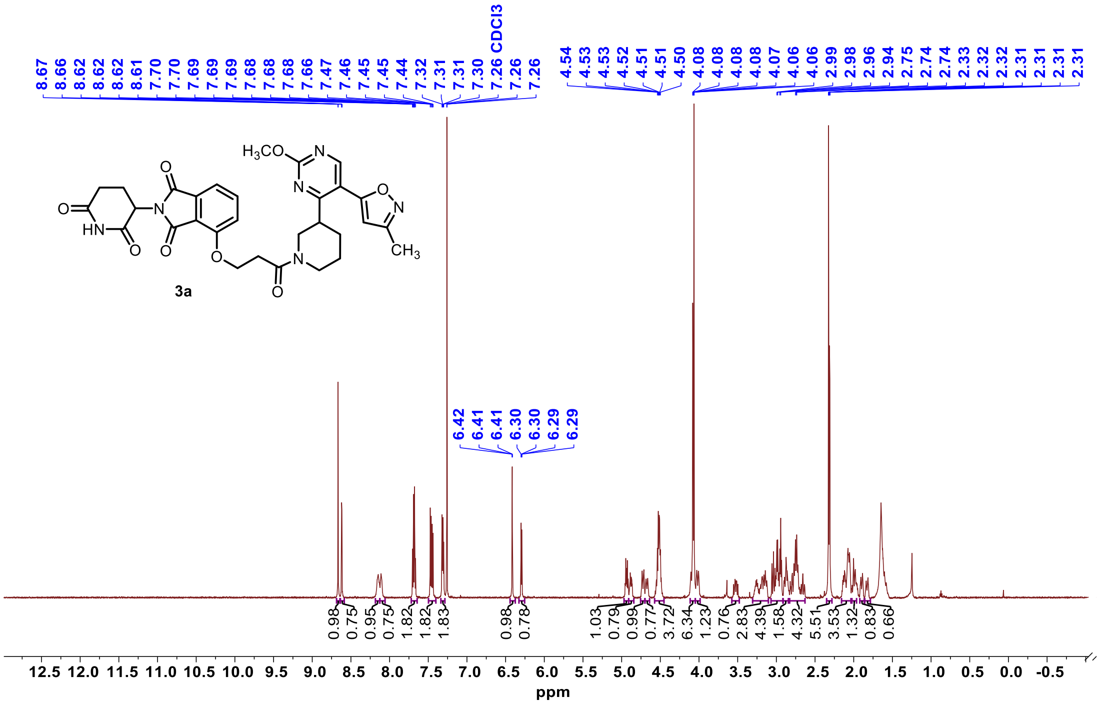

3a

#### $^{13}$C NMR **3a**

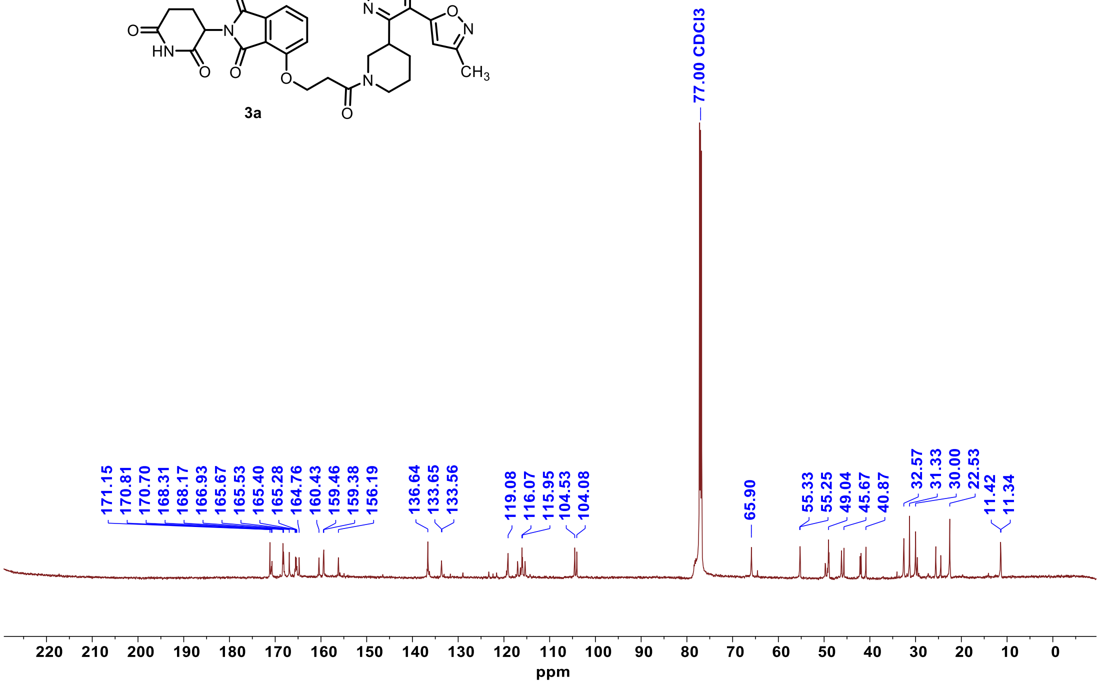

3a

#### $^{1}H$ NMR **3b**

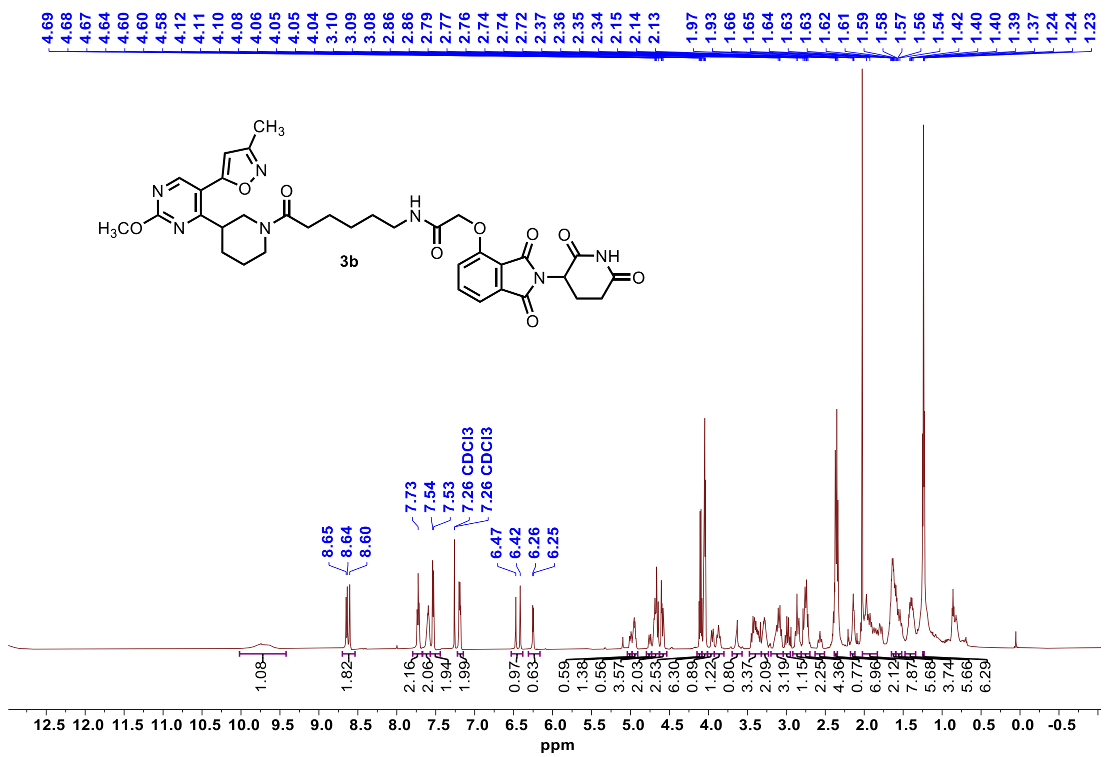


#### $^{13}C$ NMR **3b**

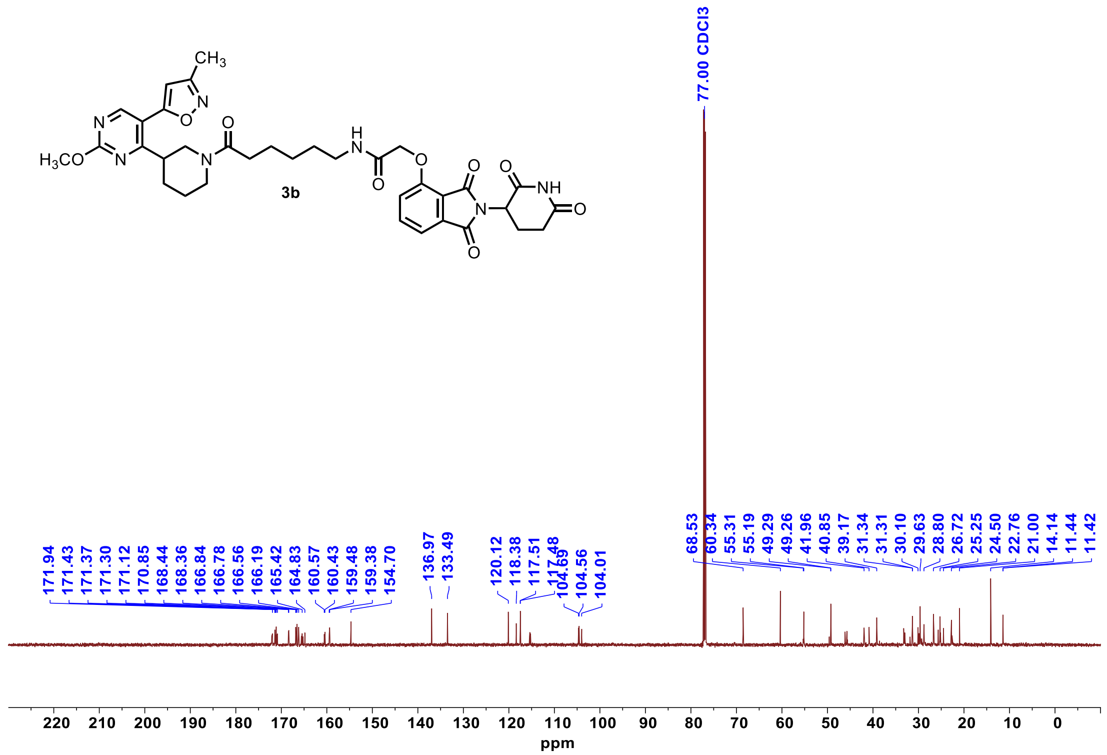

[1]H NMR **3c**

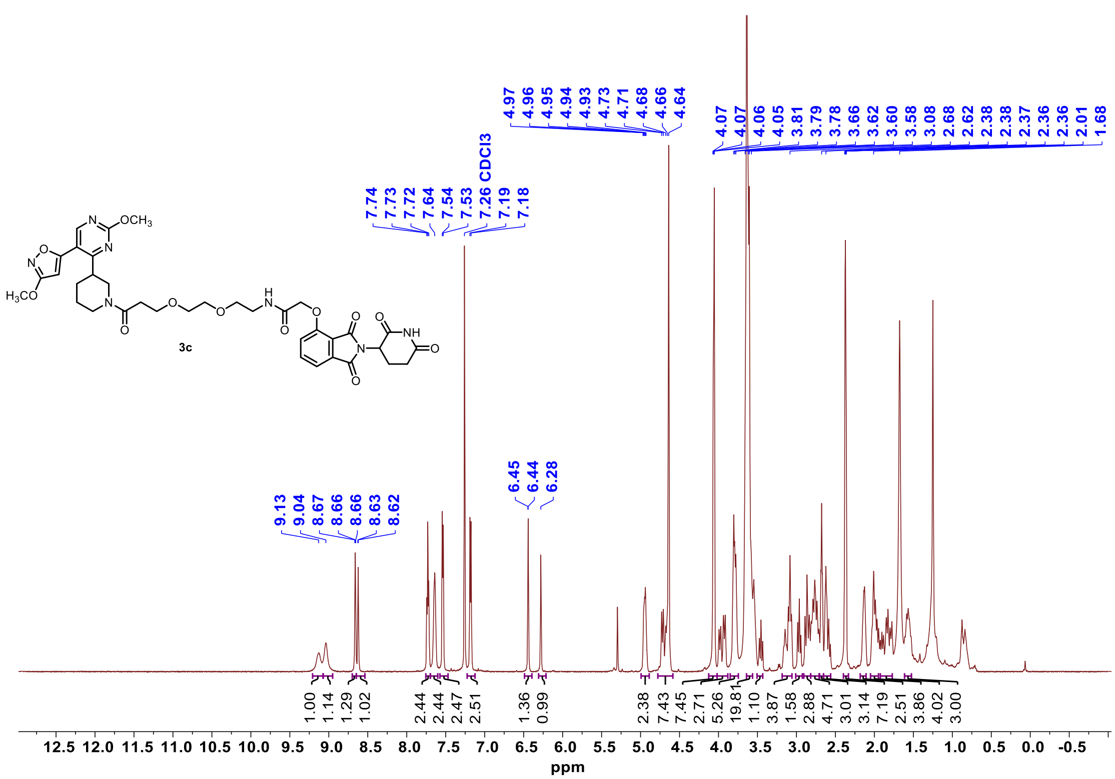


[13]C NMR **3c**

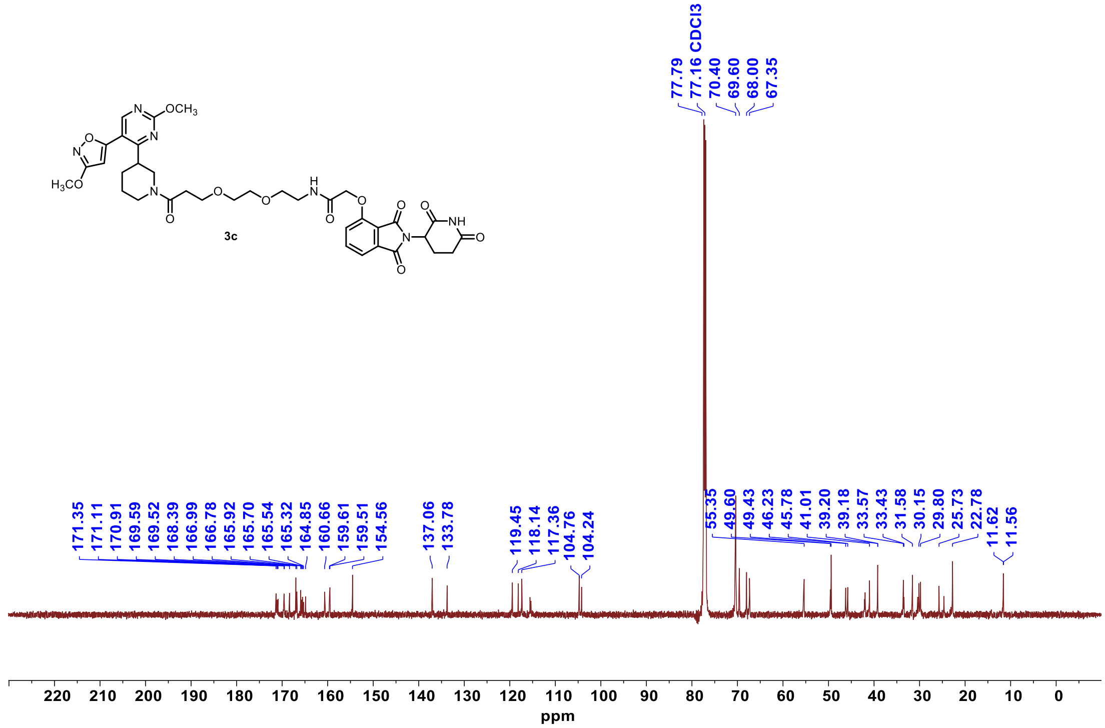

### $^{1}$H NMR **3d**

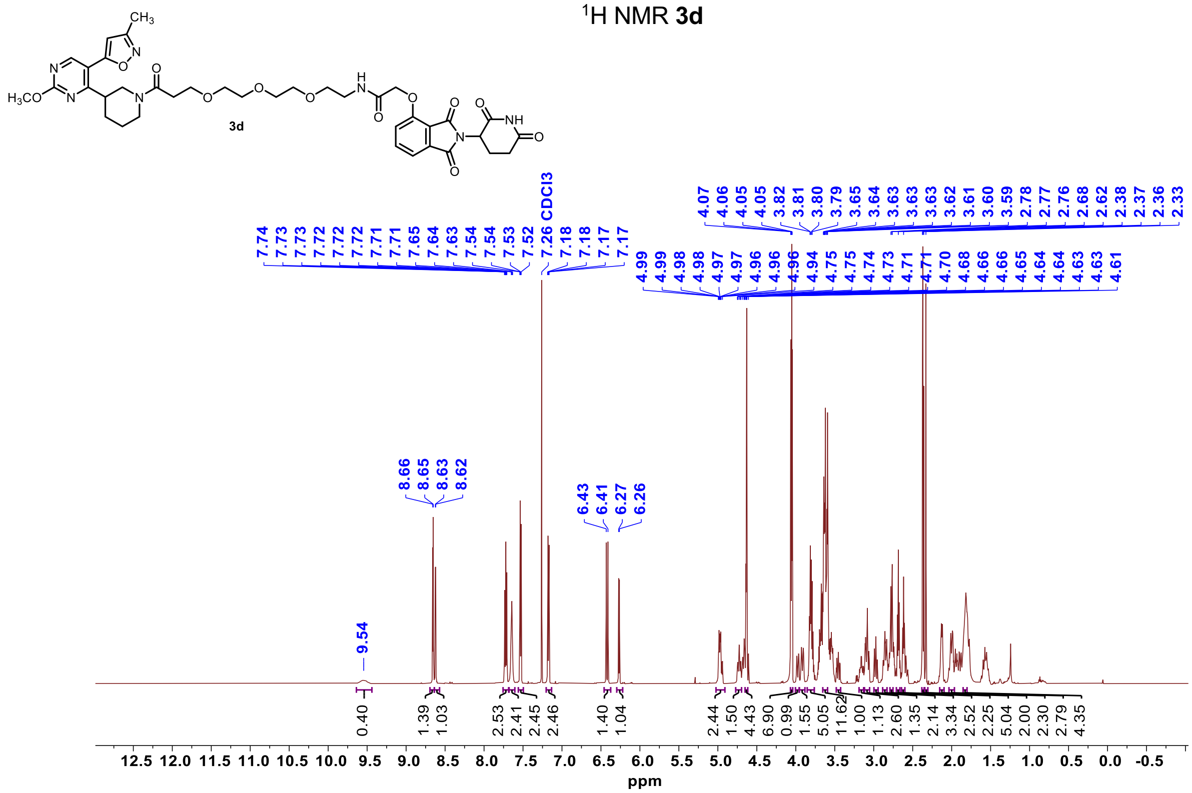


### $^{13}$C NMR **3d**

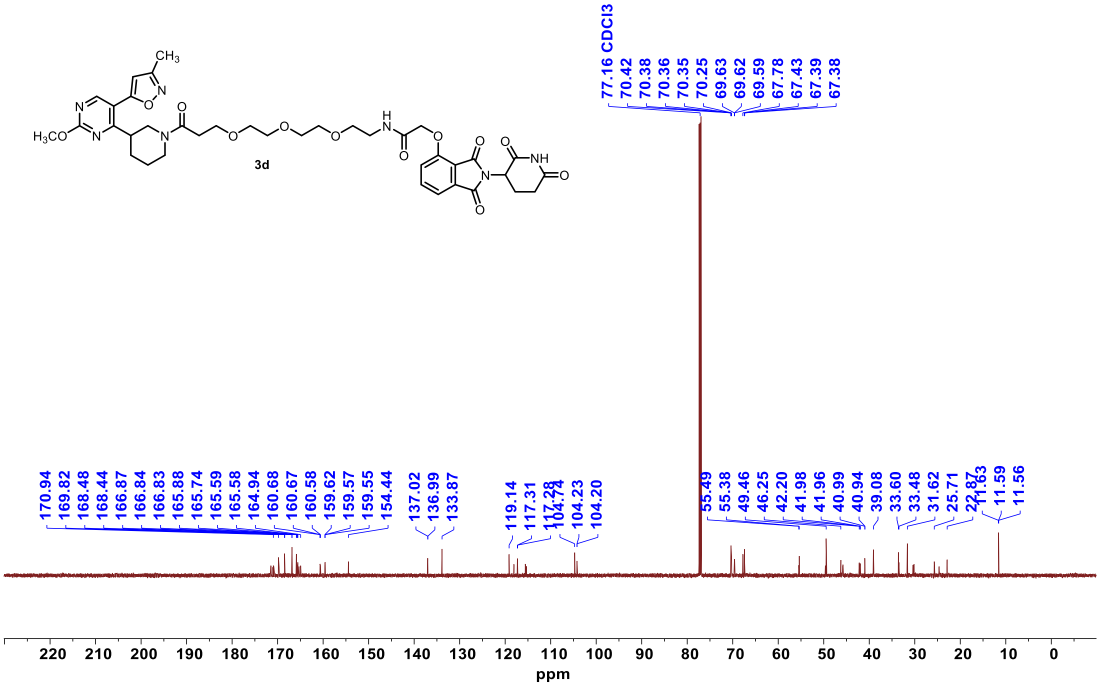

#### $^{1}$H-NMR-**4a**

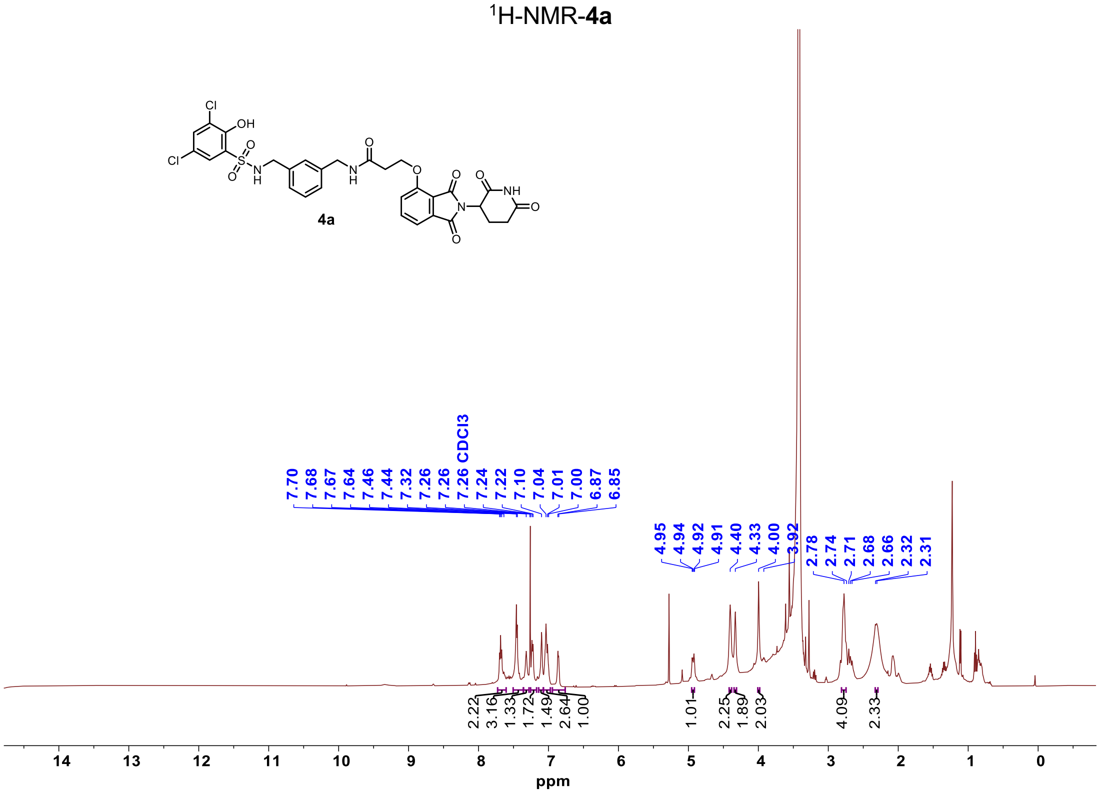


#### $^{13}$C-NMR-**4a**

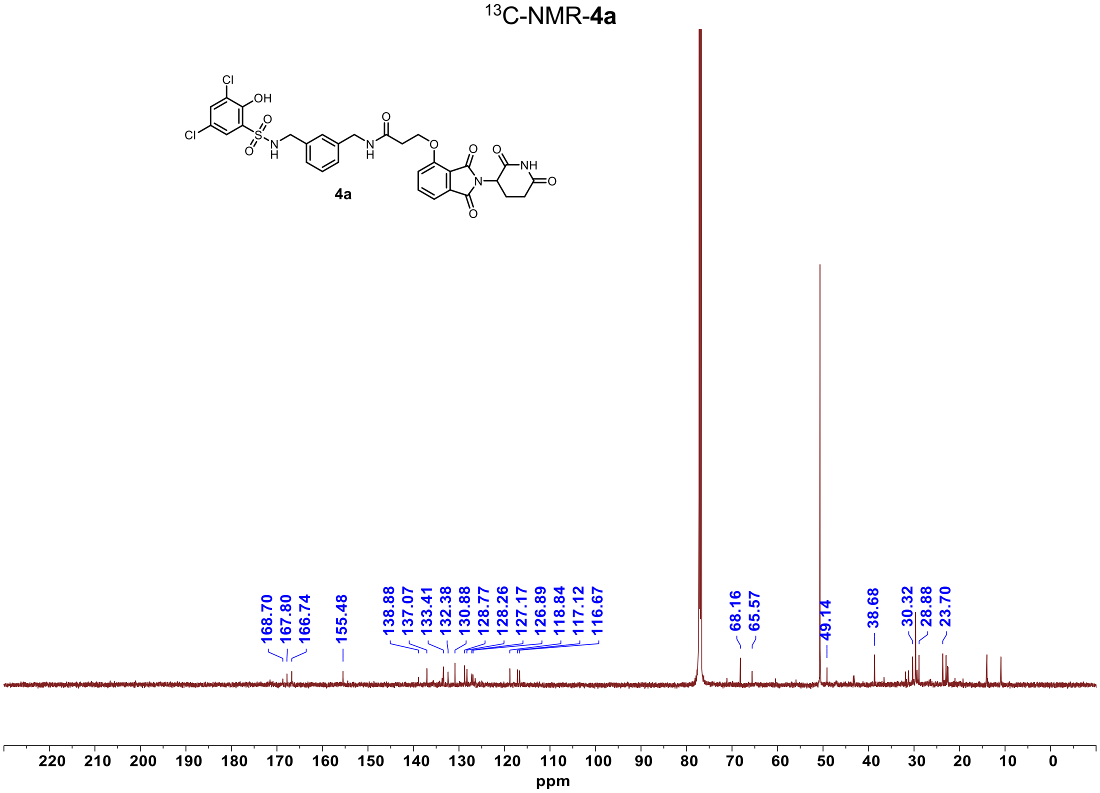

$^{1}$H-NMR-**4b**

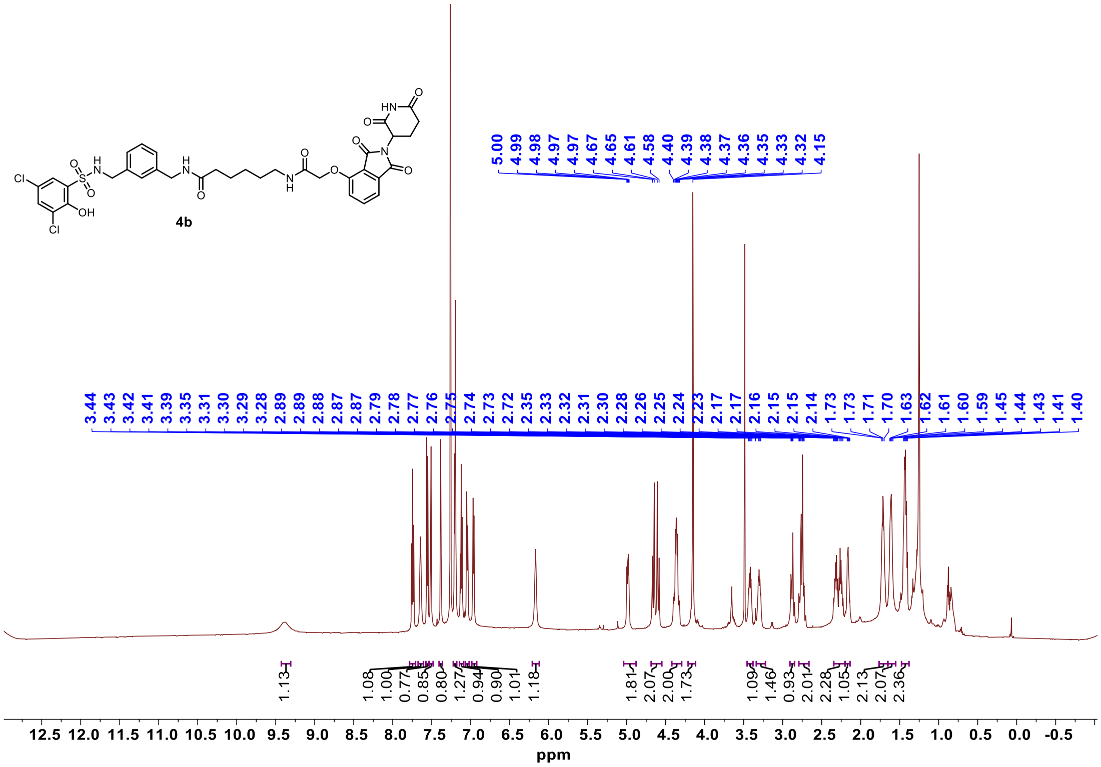

$^{13}$C-NMR-**4b**

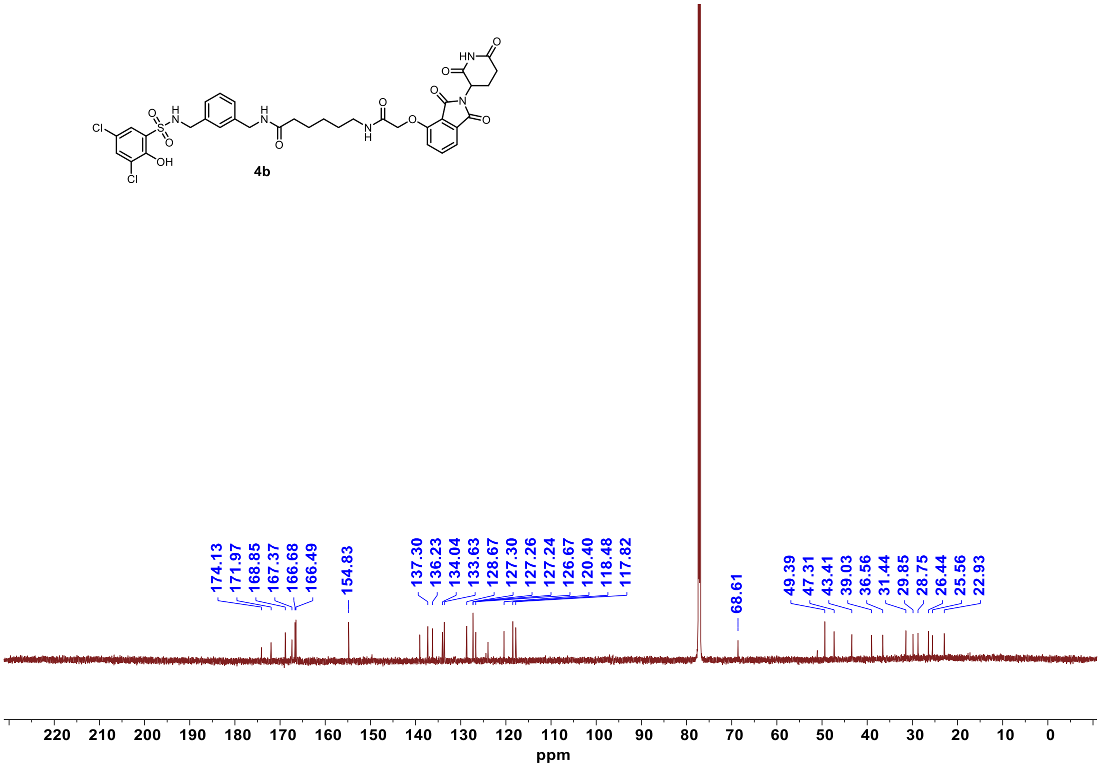

$^{1}H$ NMR **4c**

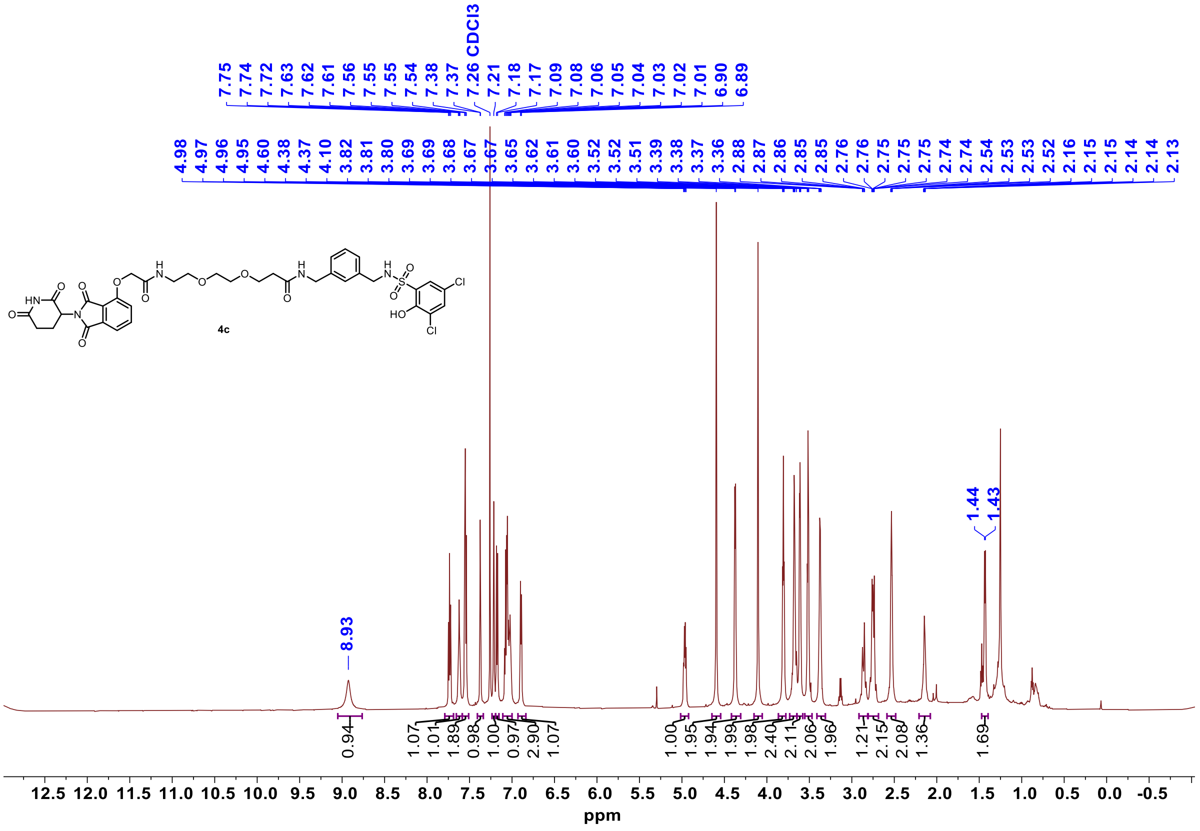


$^{13}C$ NMR **4c**

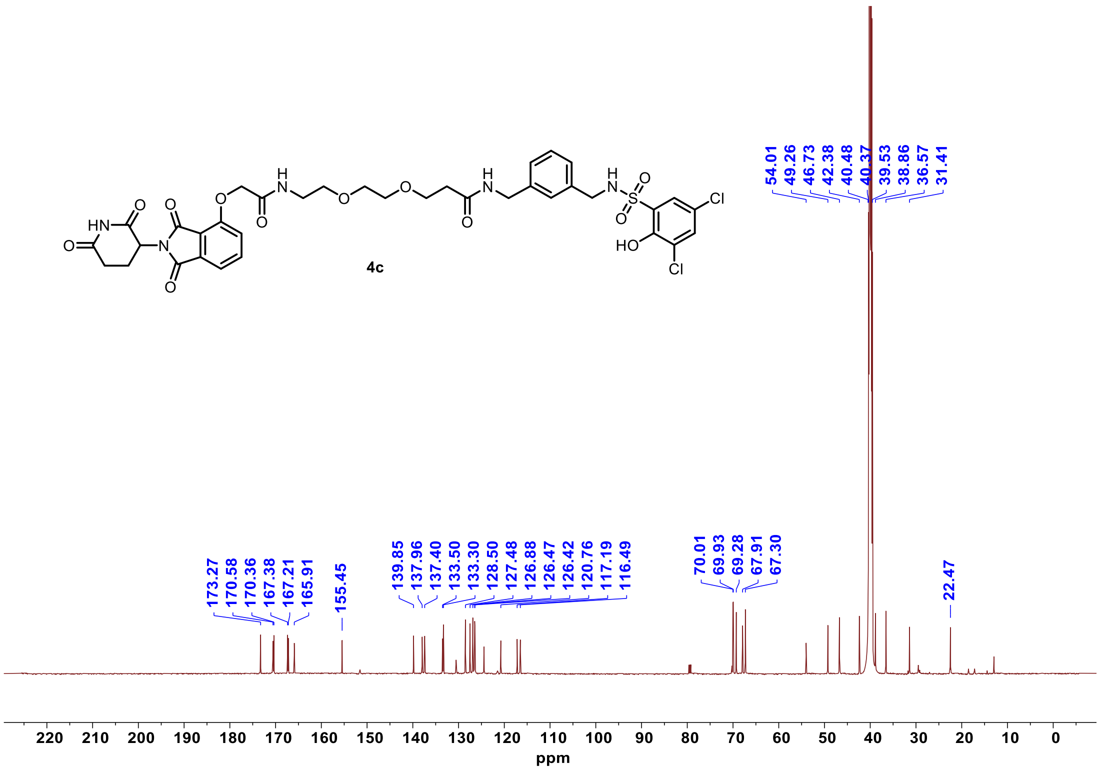

[1]H-NMR-**4d**

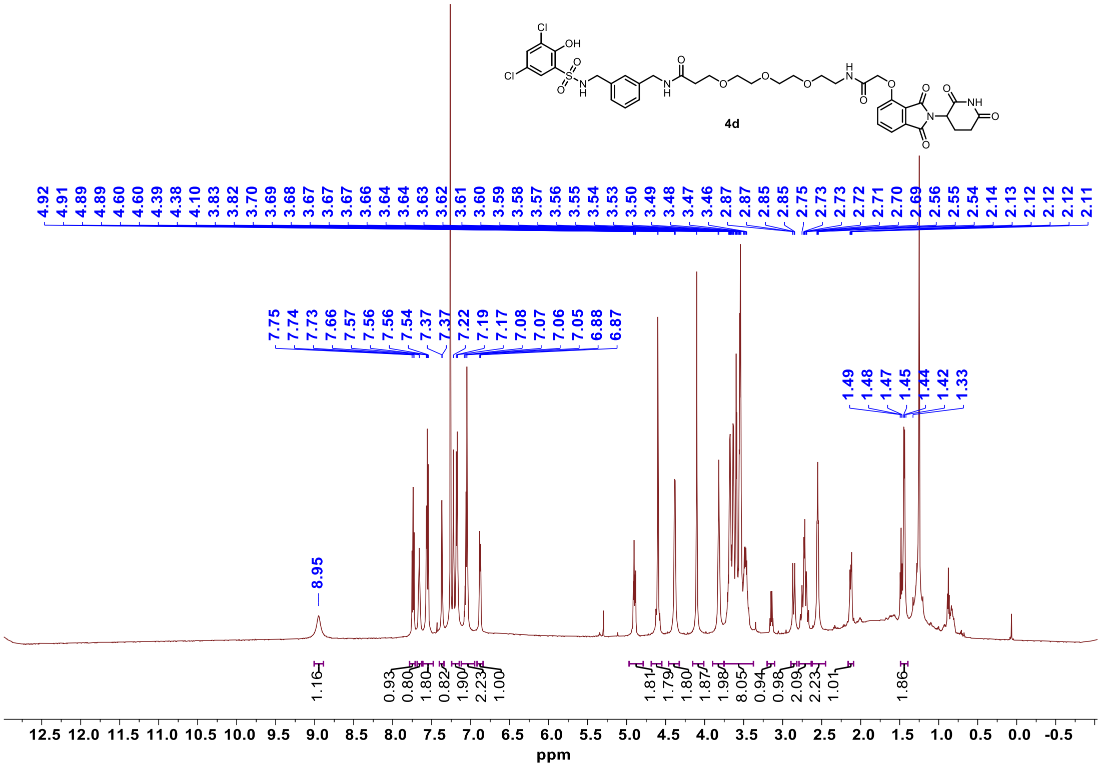


[13]C-NMR-**4d**

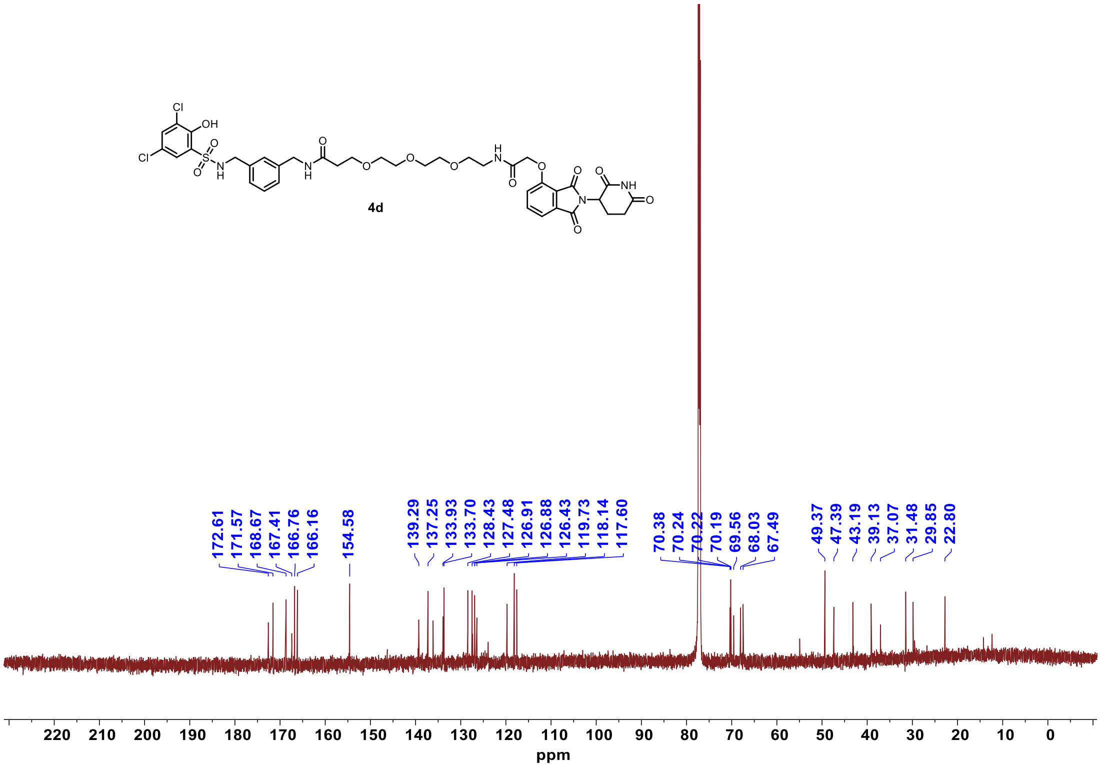

$^{1}$H NMR **5a**

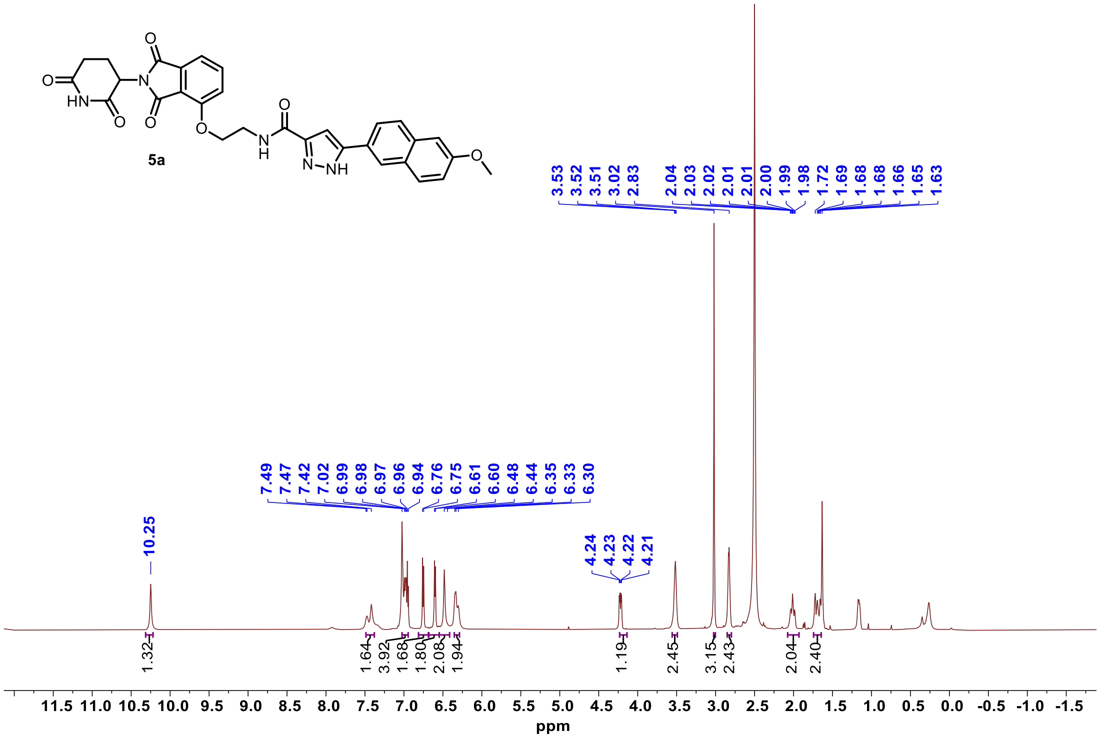


$^{13}$C NMR **5a**

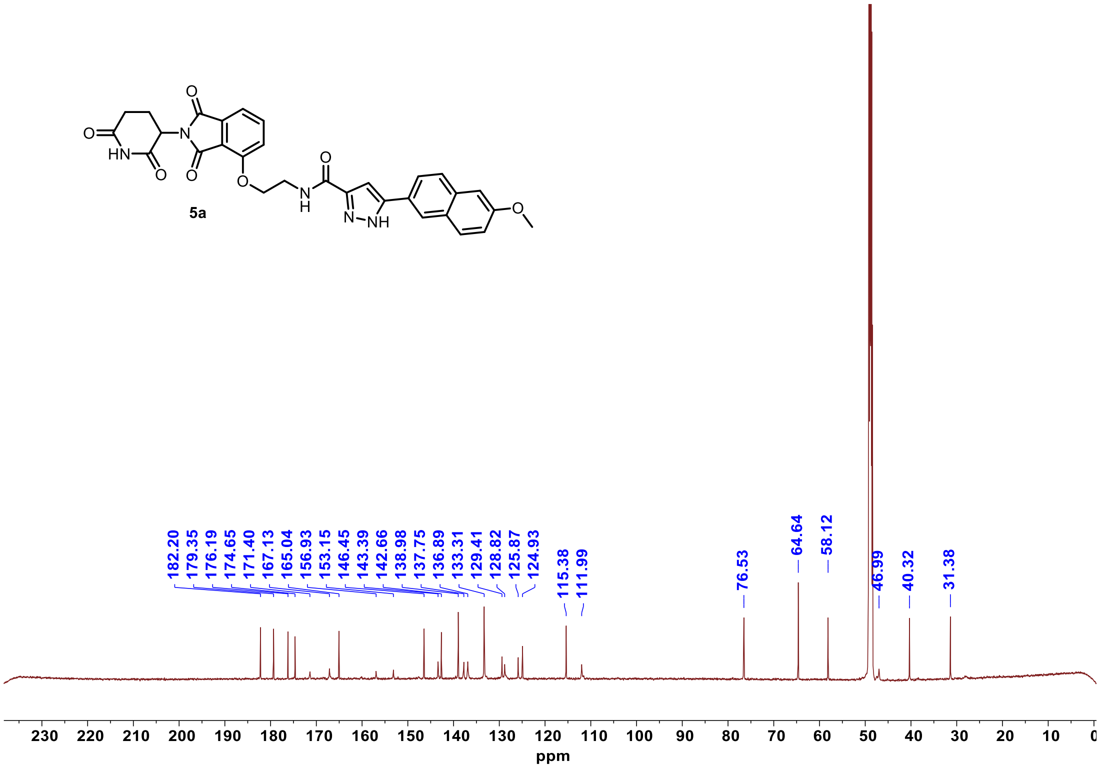

#### $^{1}$H NMR **5a-neg**

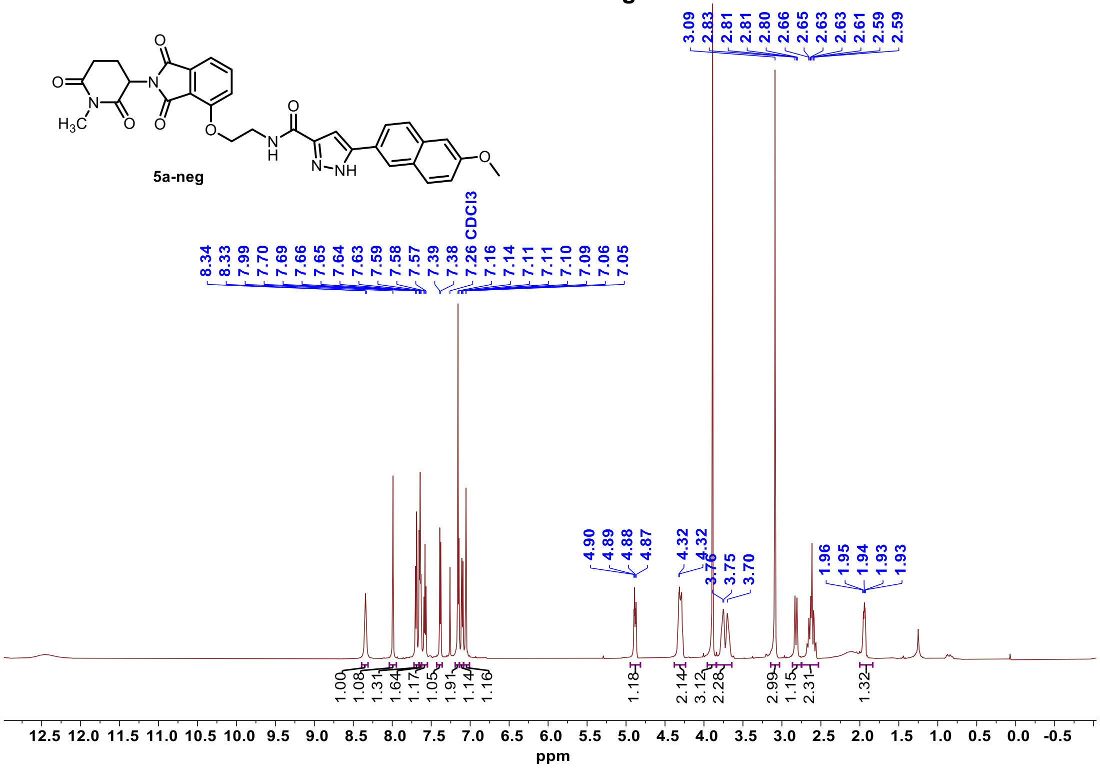


#### $^{13}$C NMR **5a-neg**

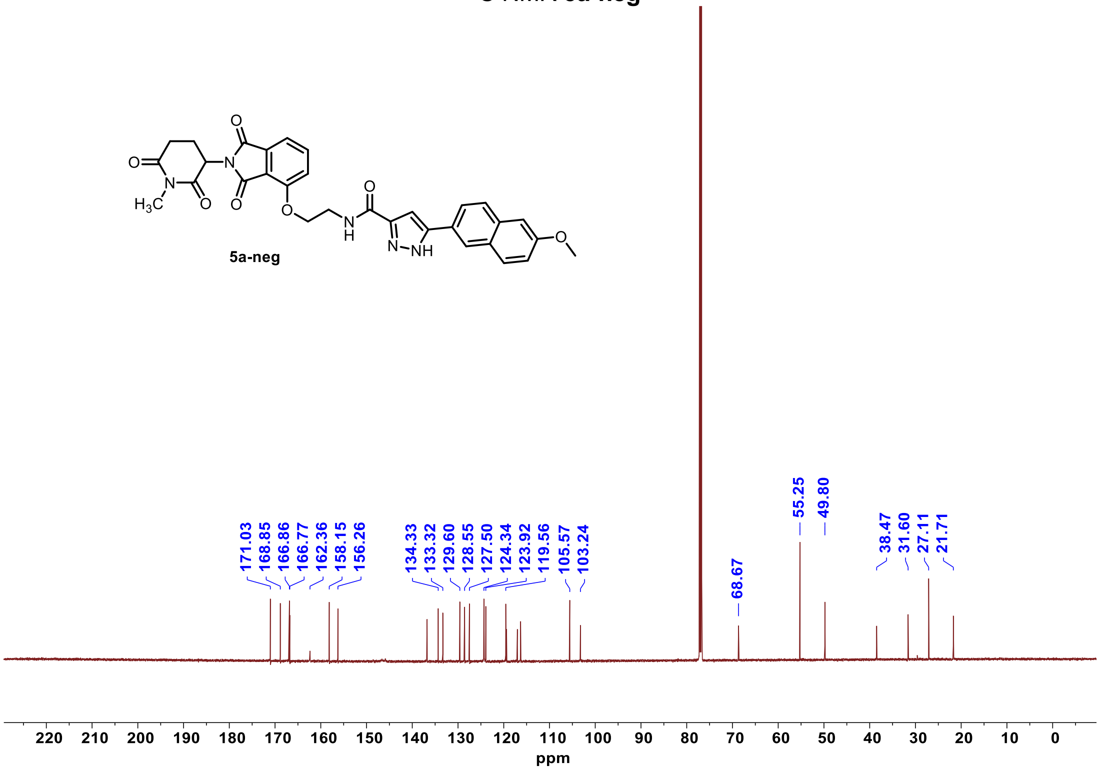

$^{1}$H-NMR-**5b**

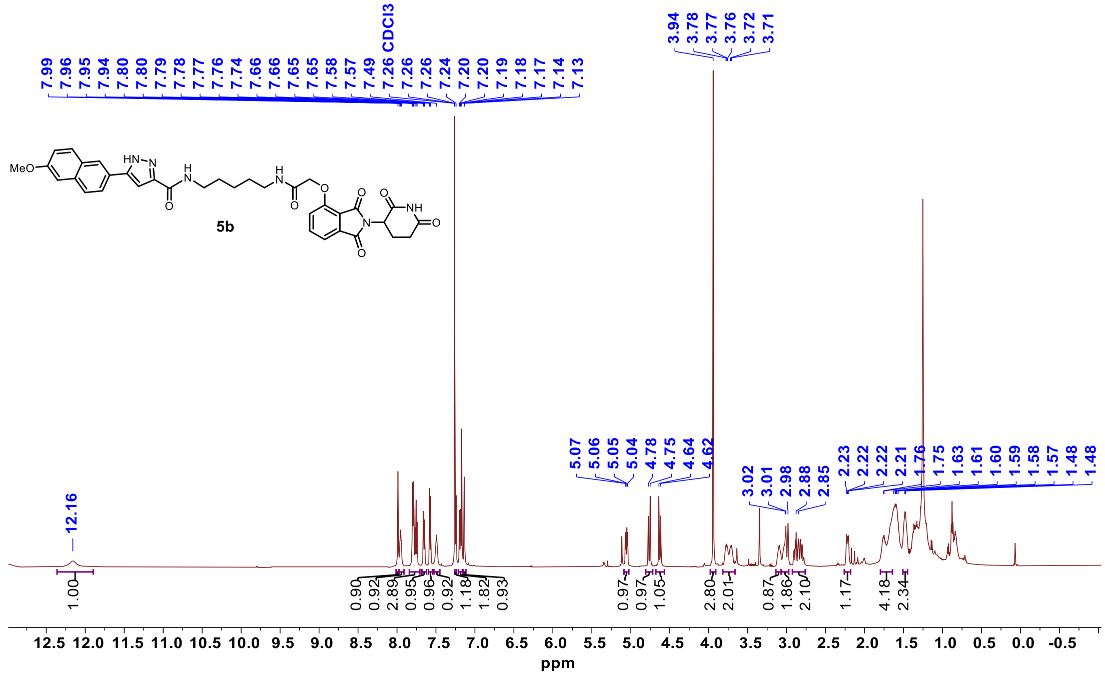


$^{13}$C-NMR-**5b**

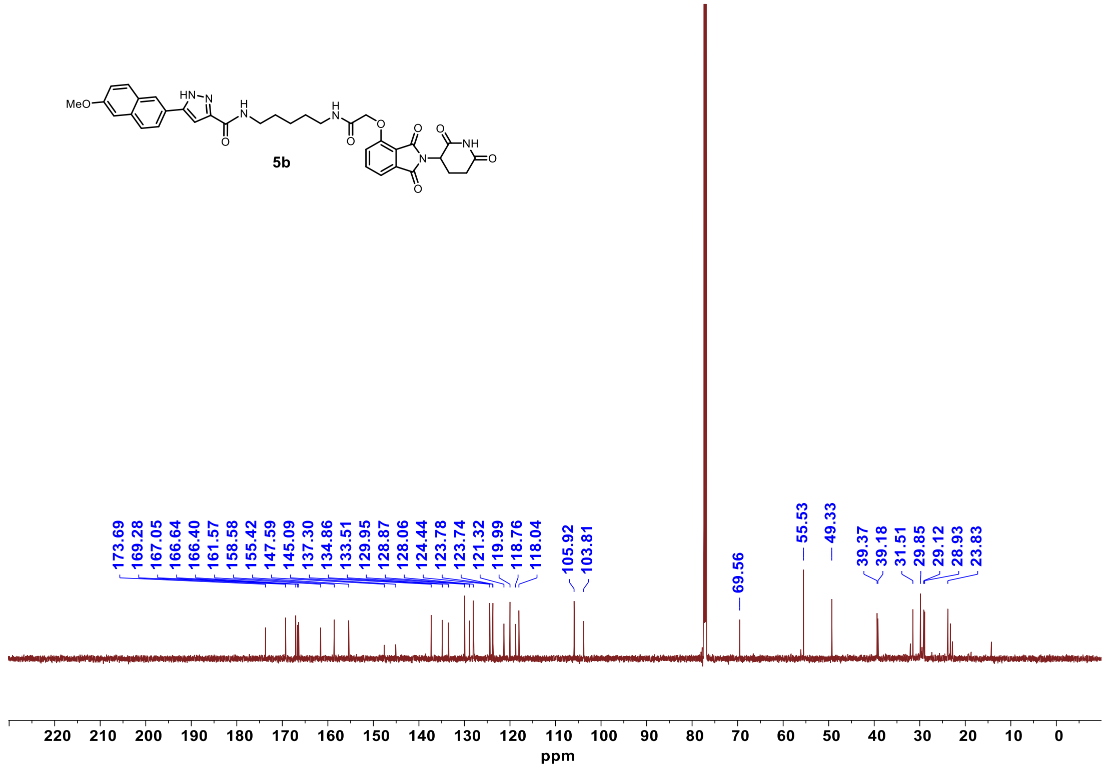

[1]H-NMR-**5c**

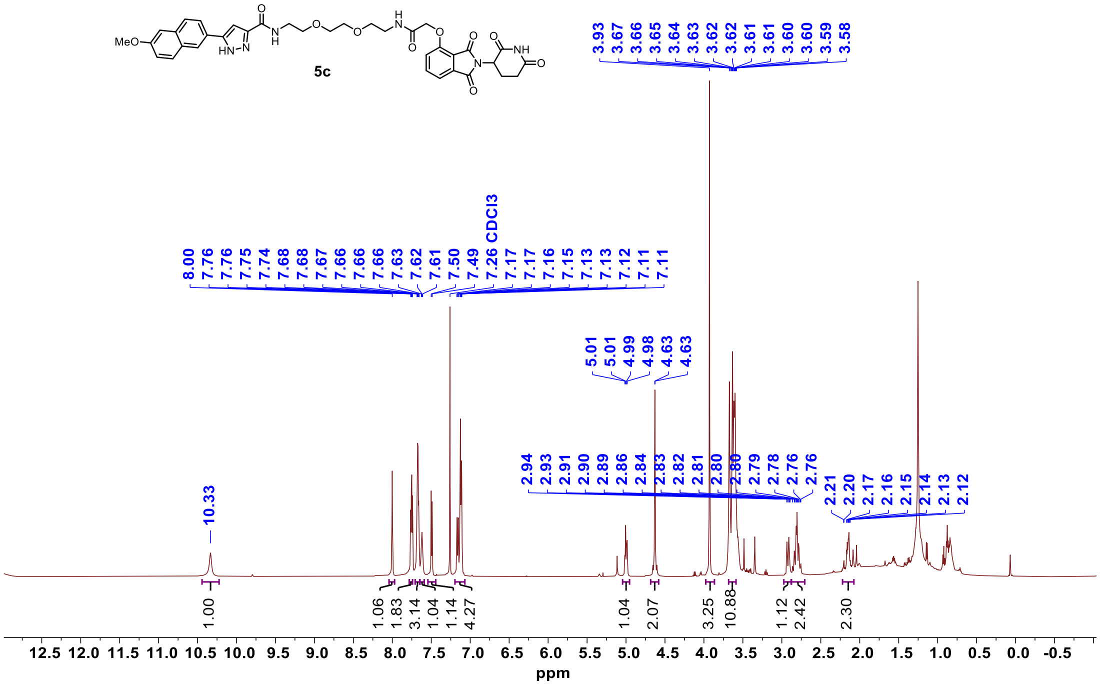


[13]C-NMR-**5c**

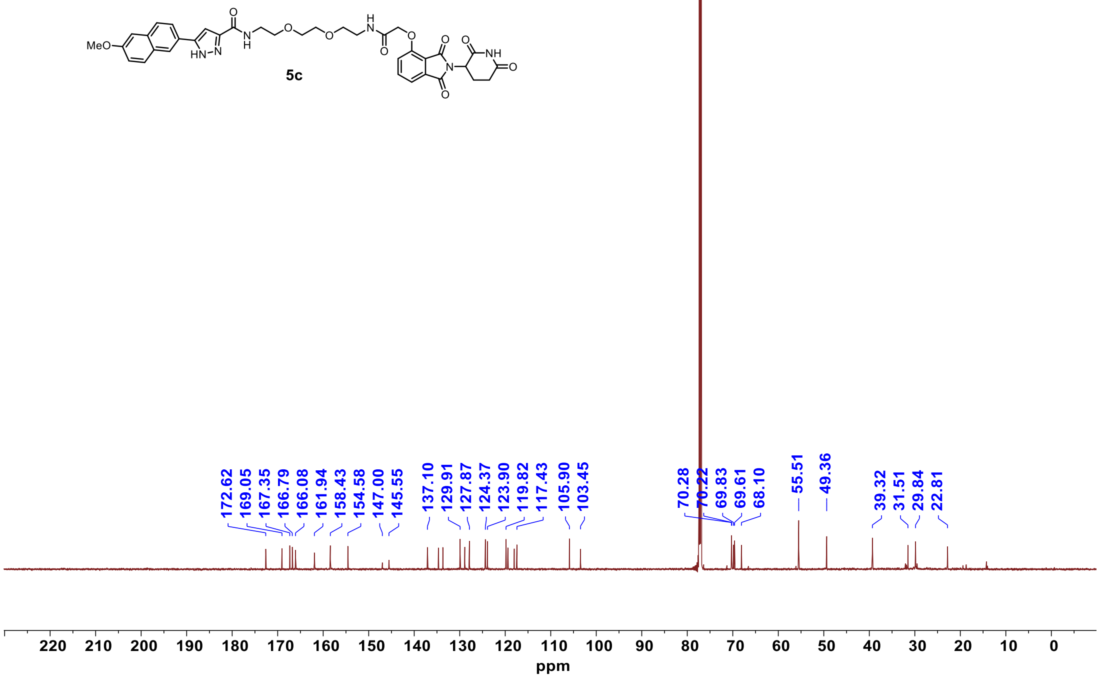

$^{1}$H-NMR-**5d**

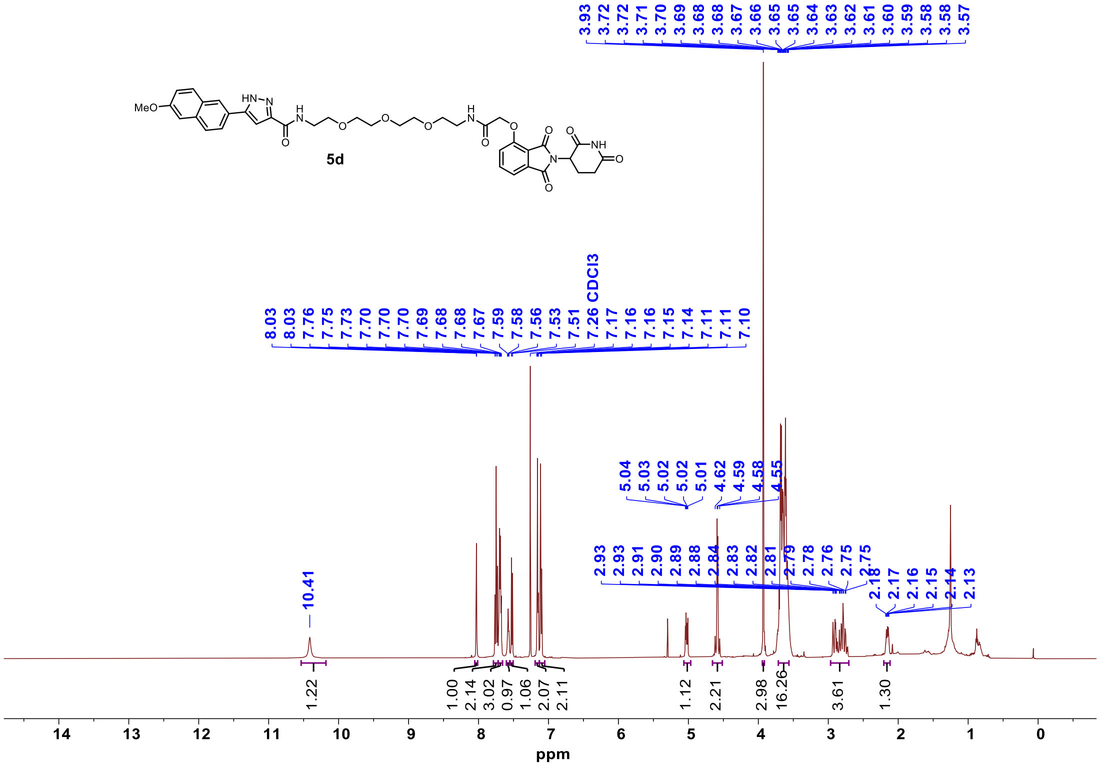


$^{13}$C-NMR-**5d**

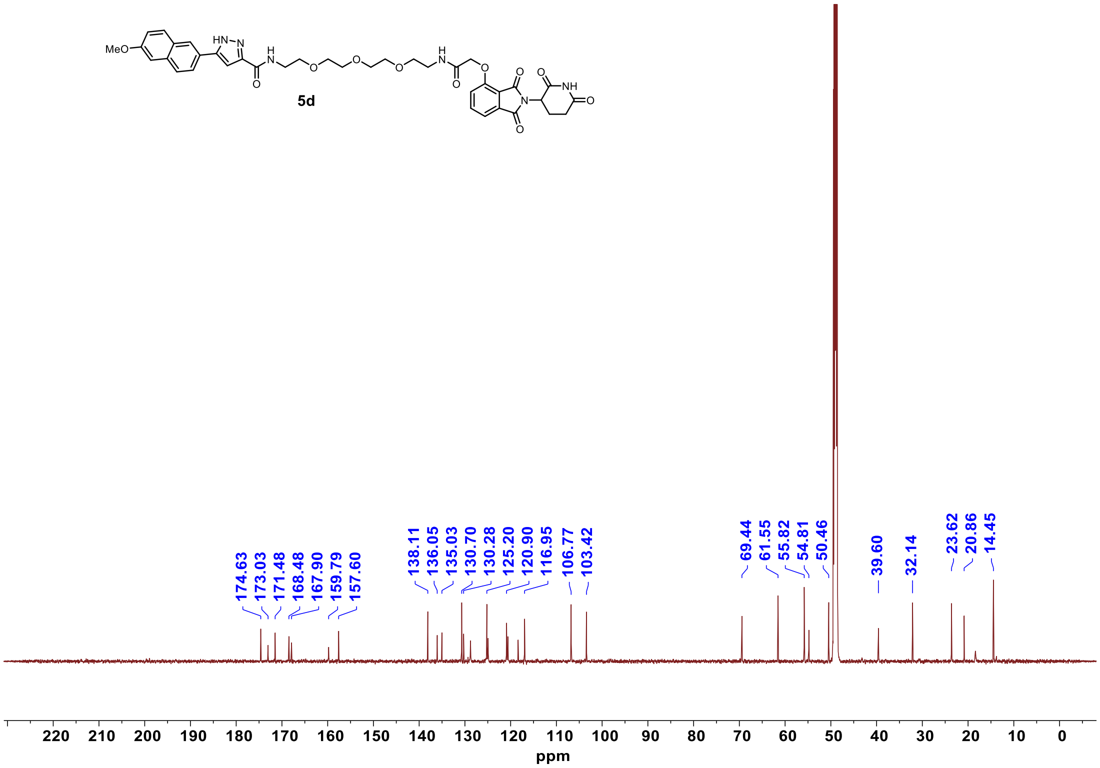

[1]H NMR **6a**

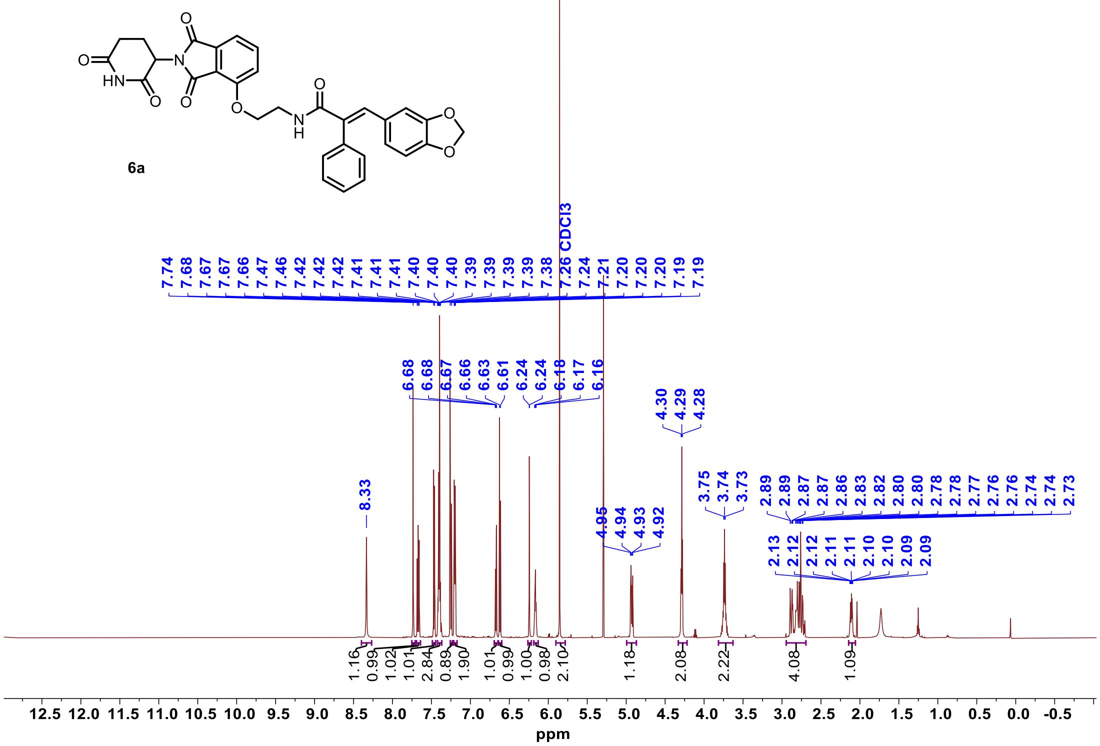


[13]C NMR **6a**

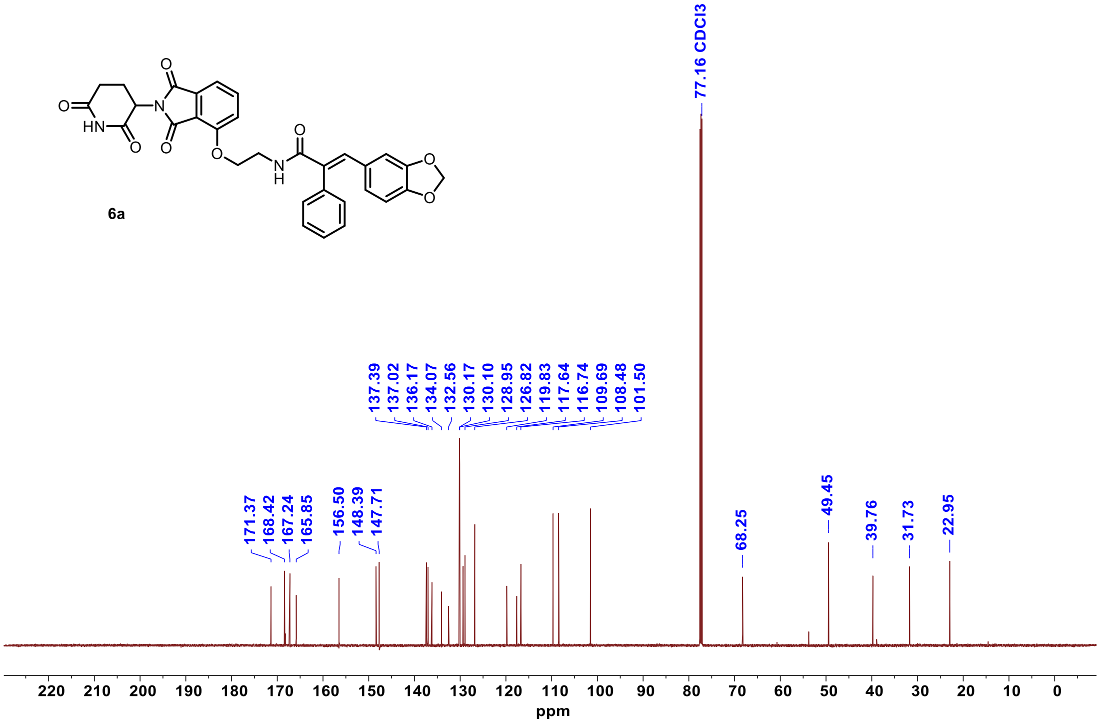

#### $^{1}$H NMR **6b**

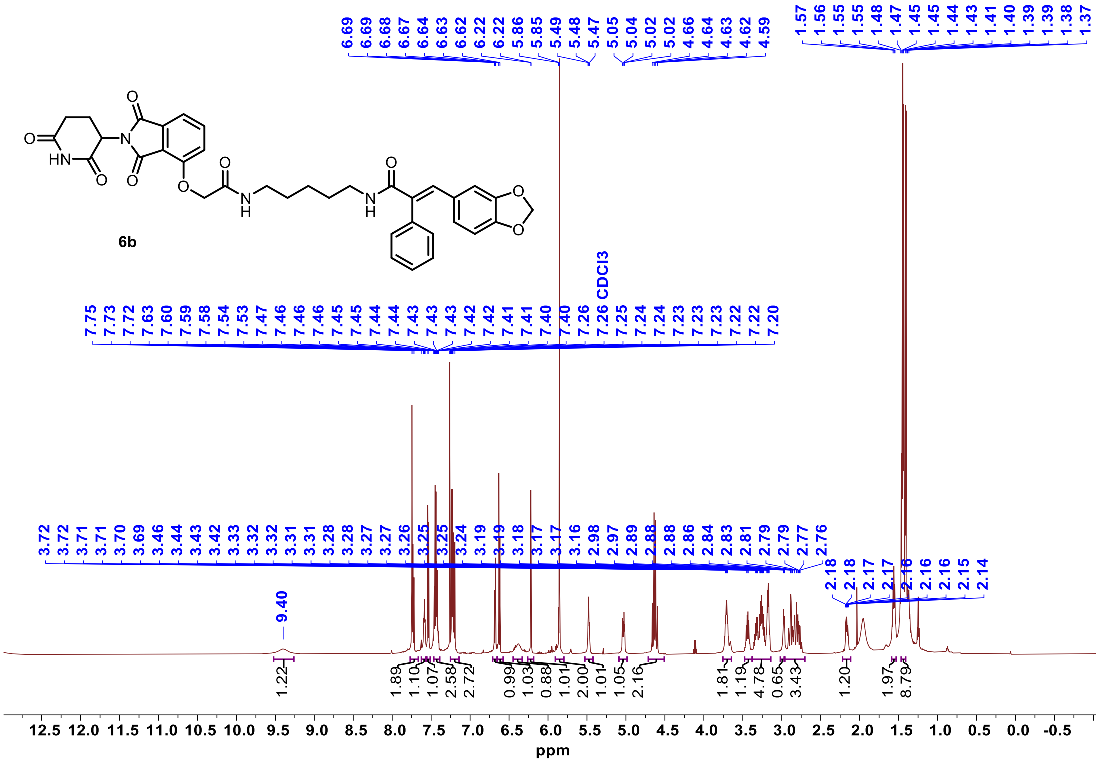


#### $^{13}$C NMR **6b**

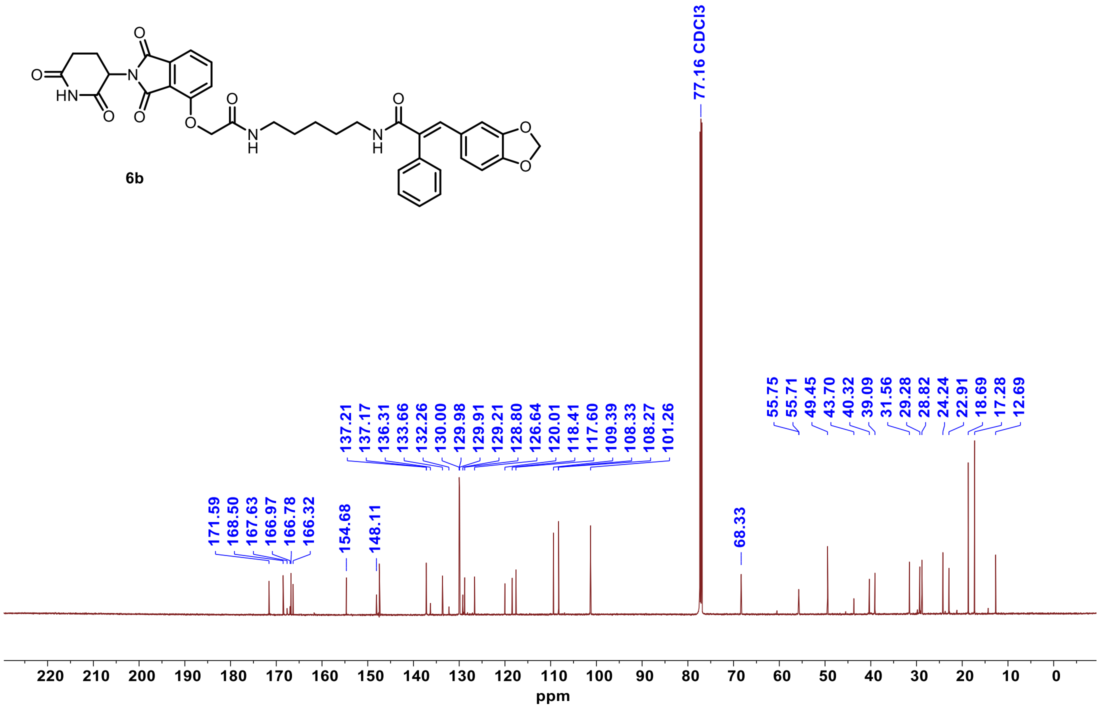

$^{1}$H NMR **6c**

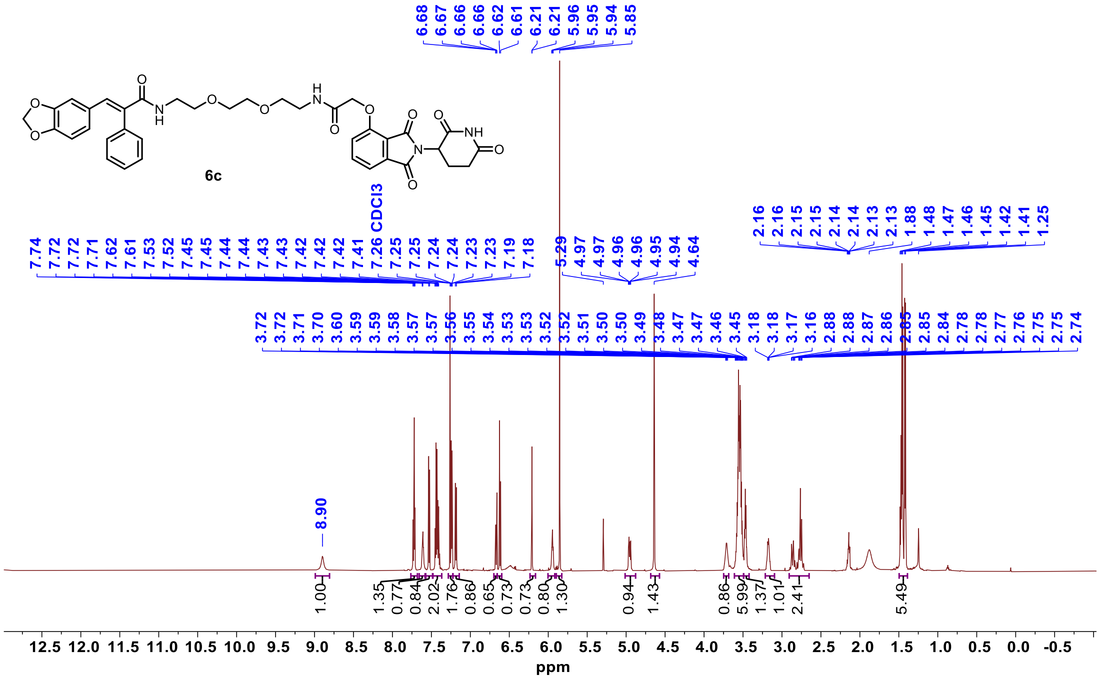


$^{13}$C NMR **6c**

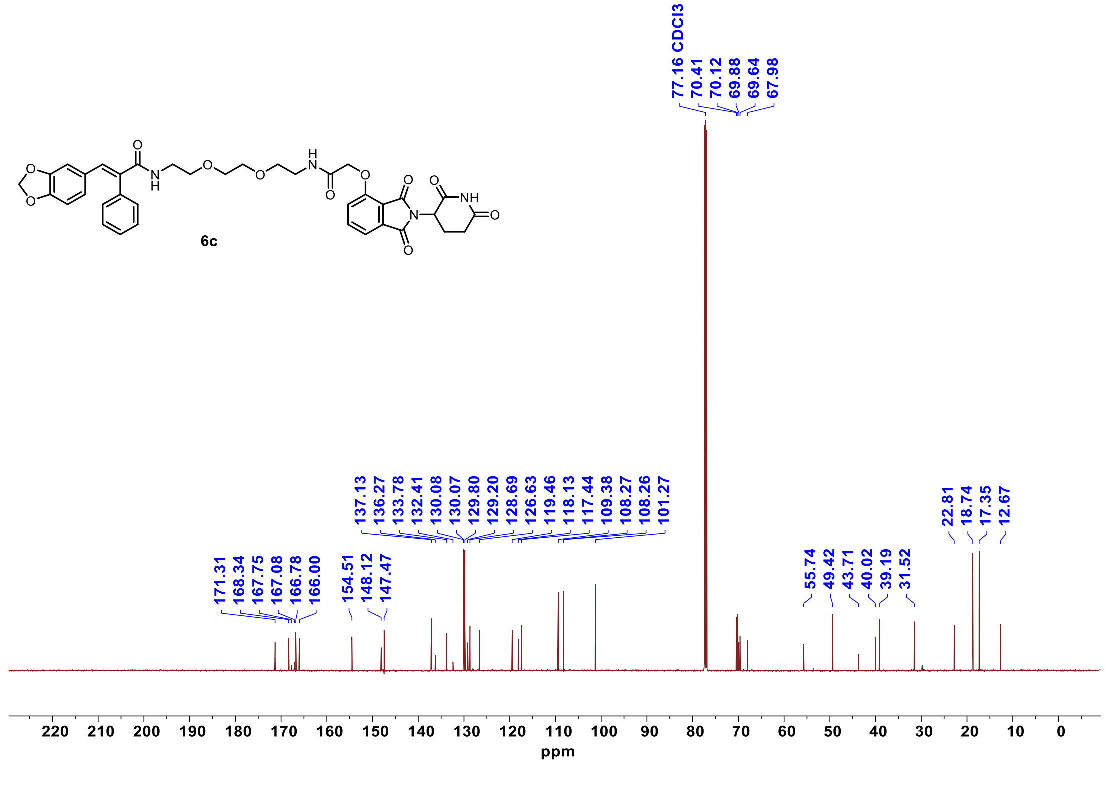

<sup></sup>

$^1$H NMR **6d**

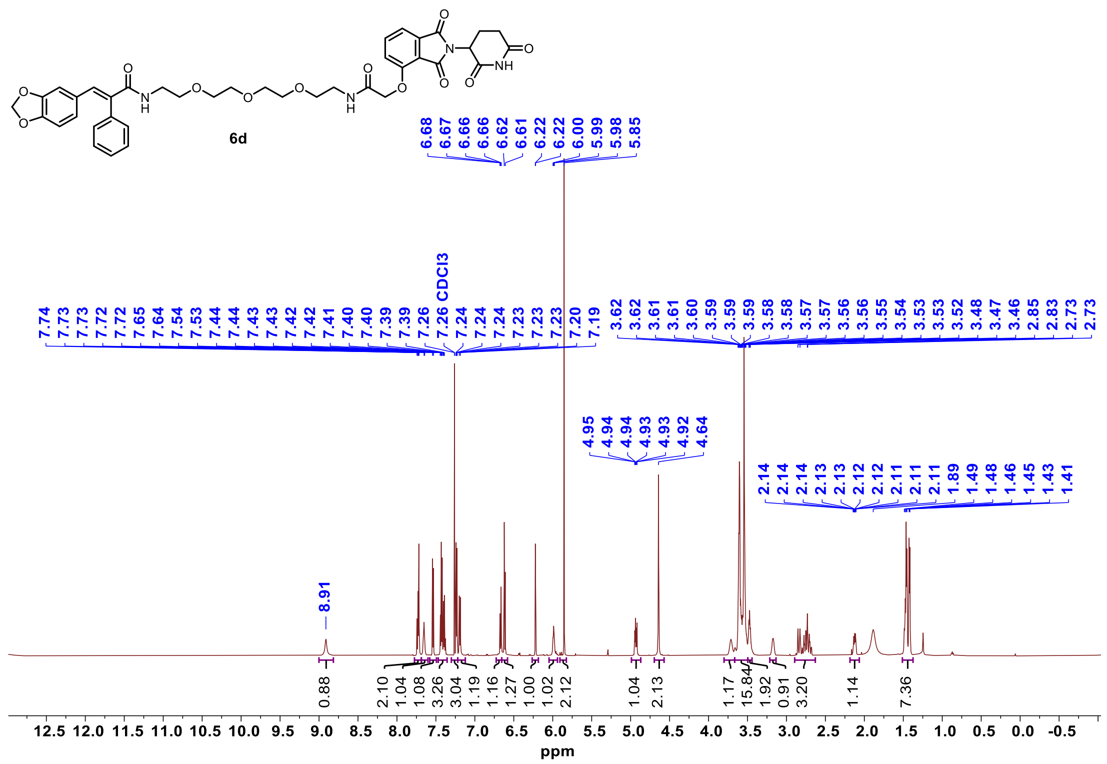


$^{13}$C NMR **6d**

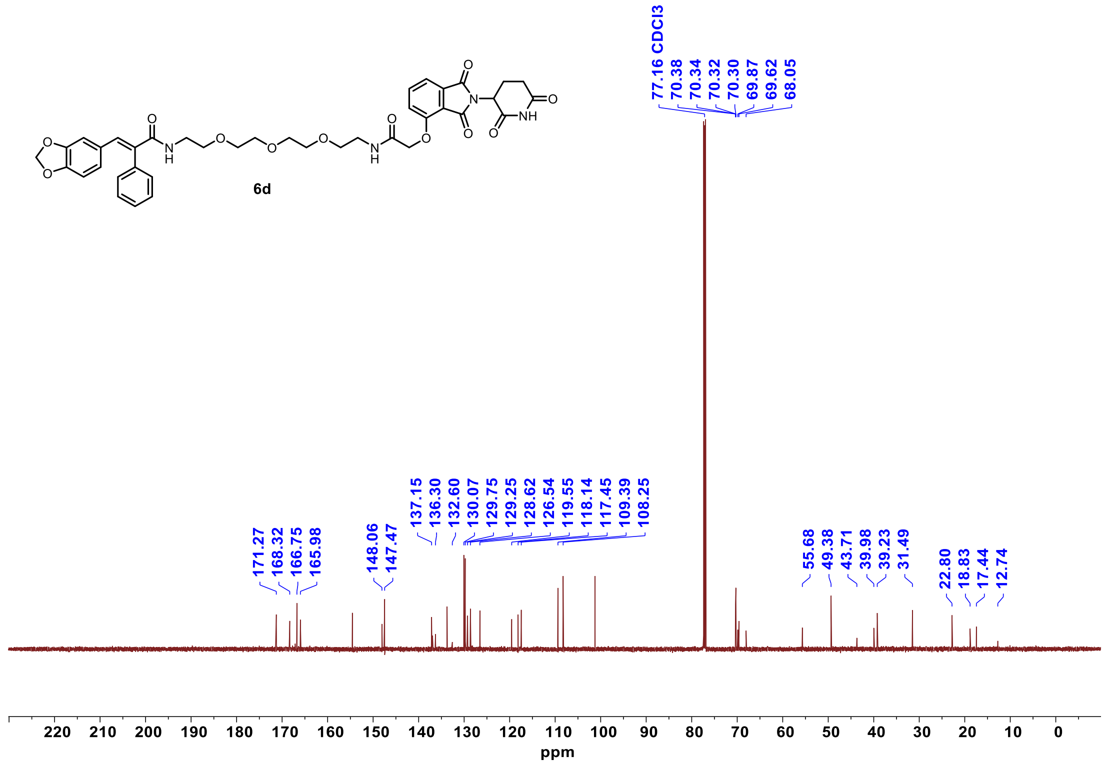

$^{1}$H NMR **7a**

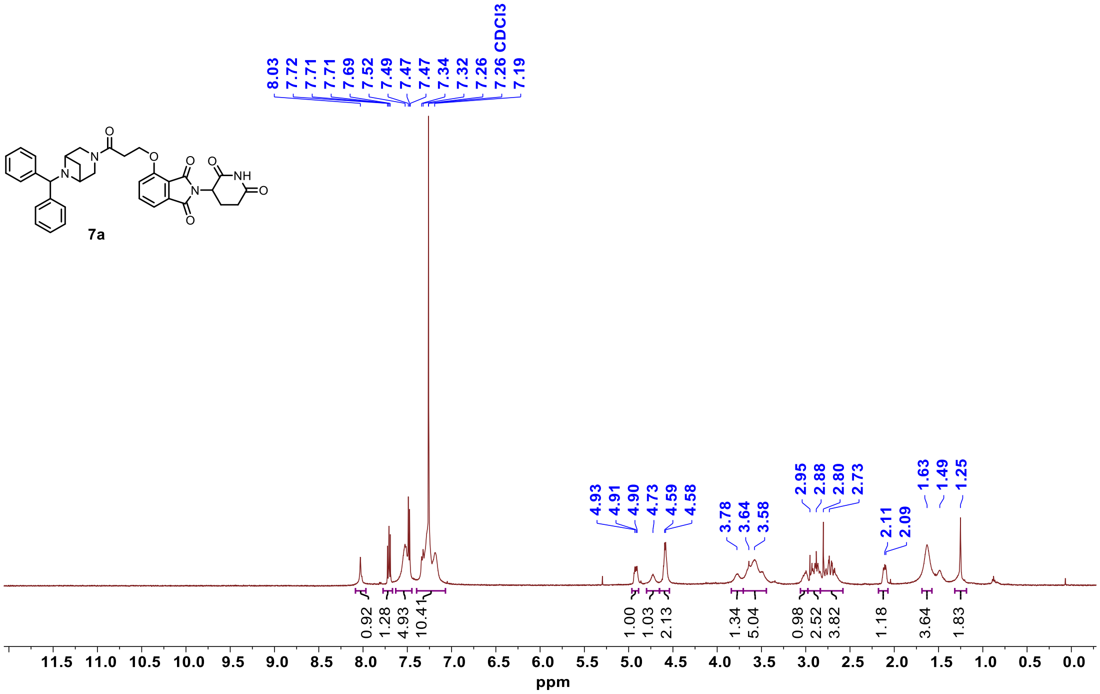


$^{13}$C NMR **7a**

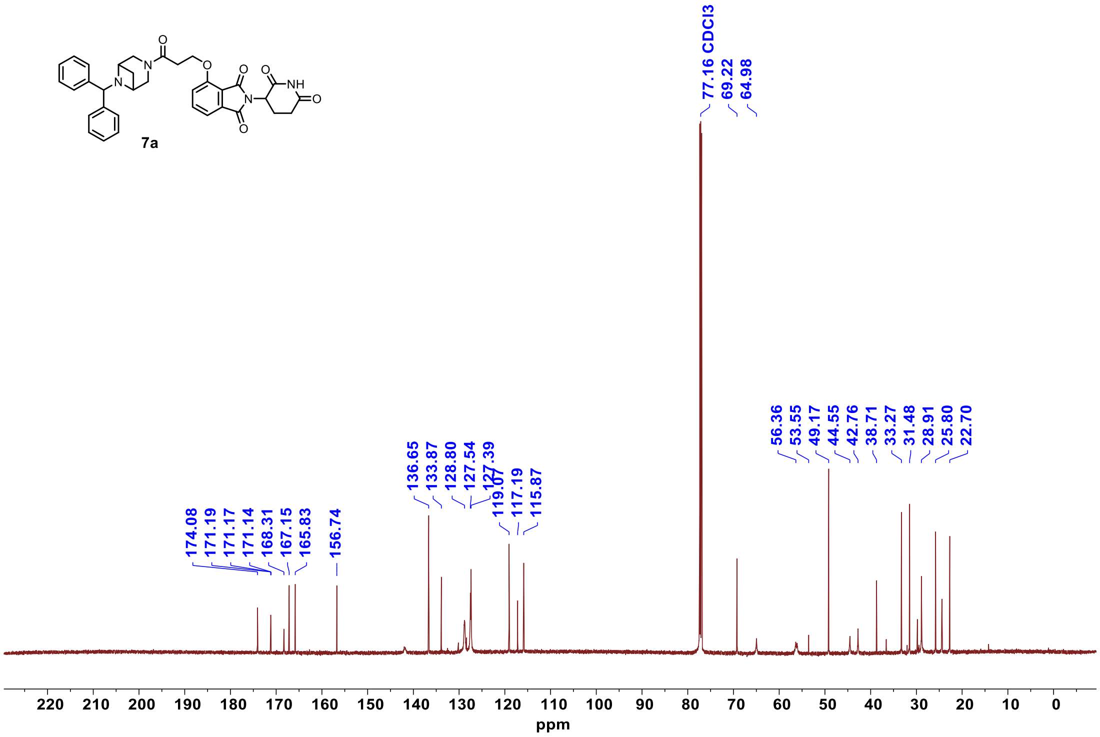

$^{1}$H NMR **7b**

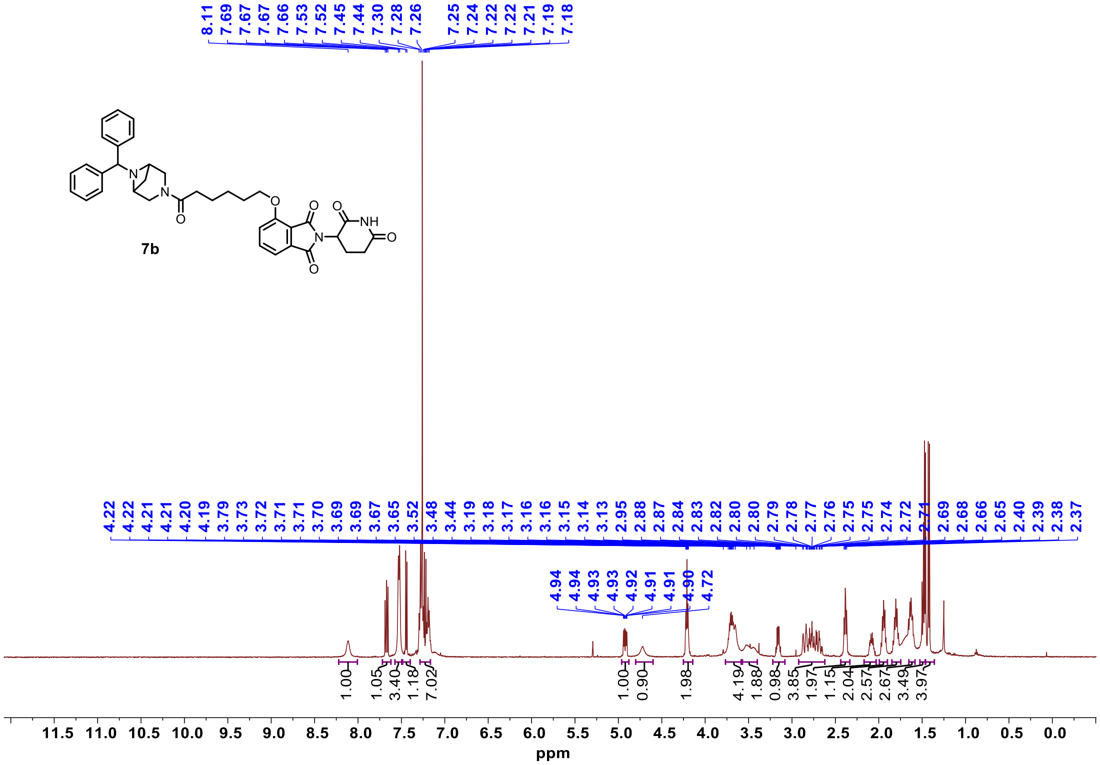

$^{13}$C NMR **7b**

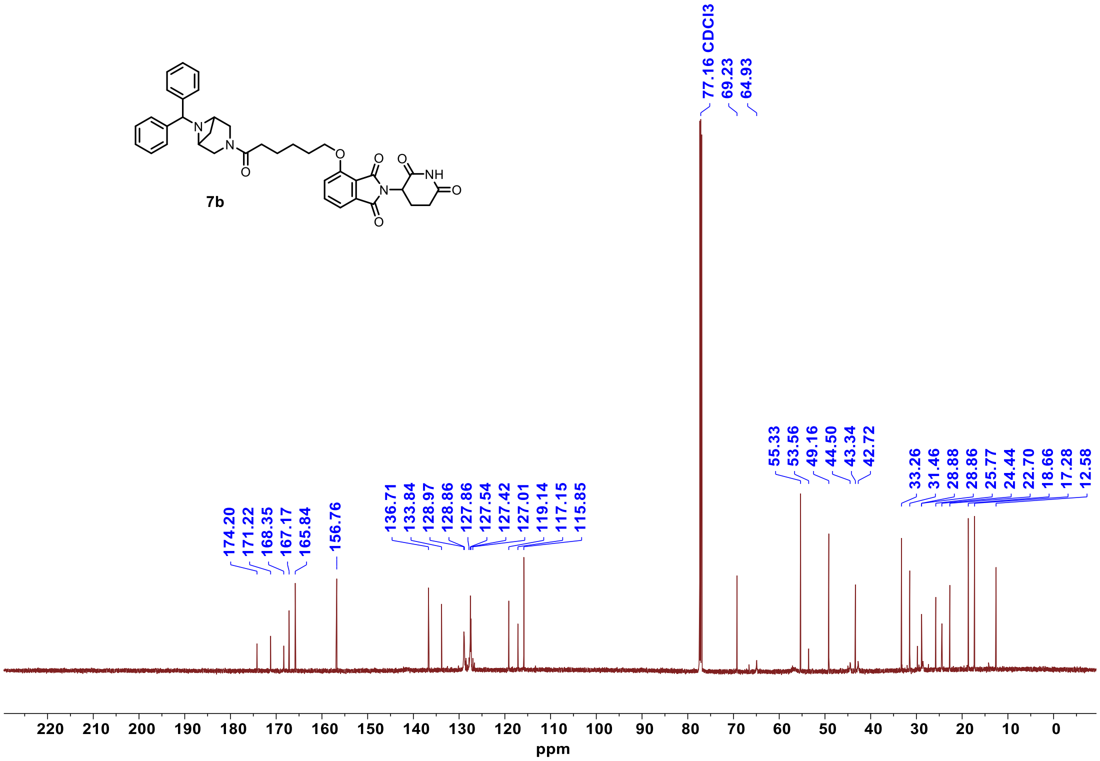

$^{1}$H NMR **7b-neg**

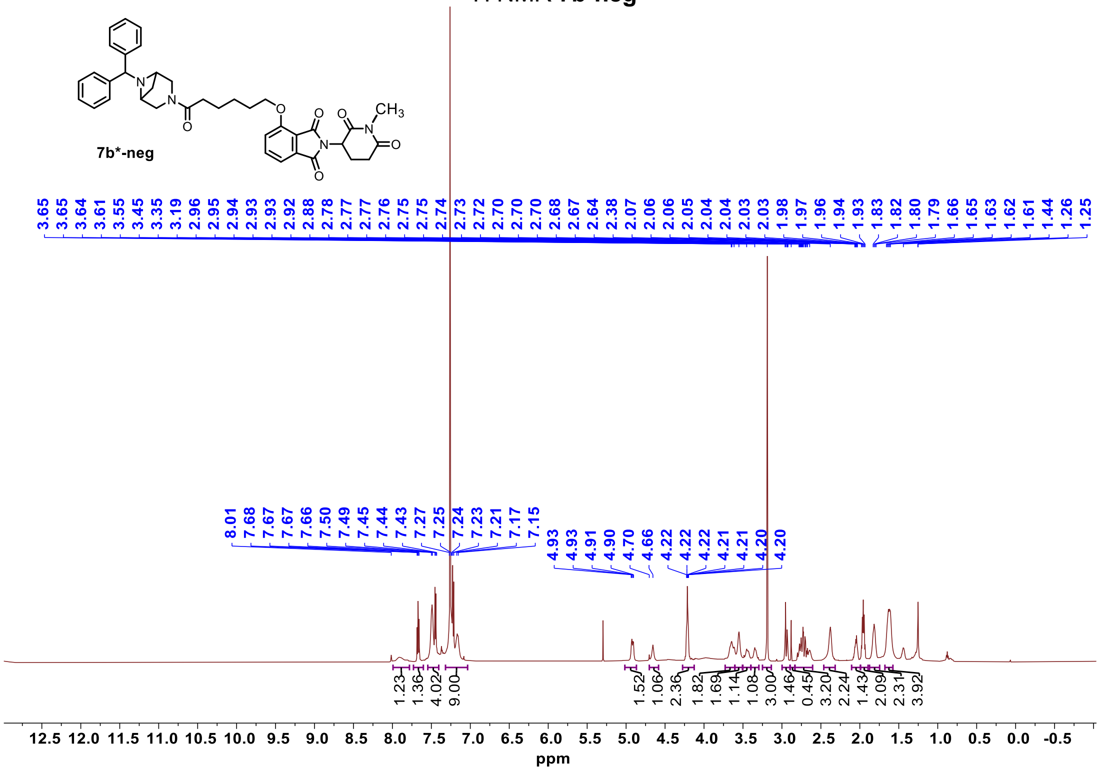


$^{13}$C NMR **7b-neg**

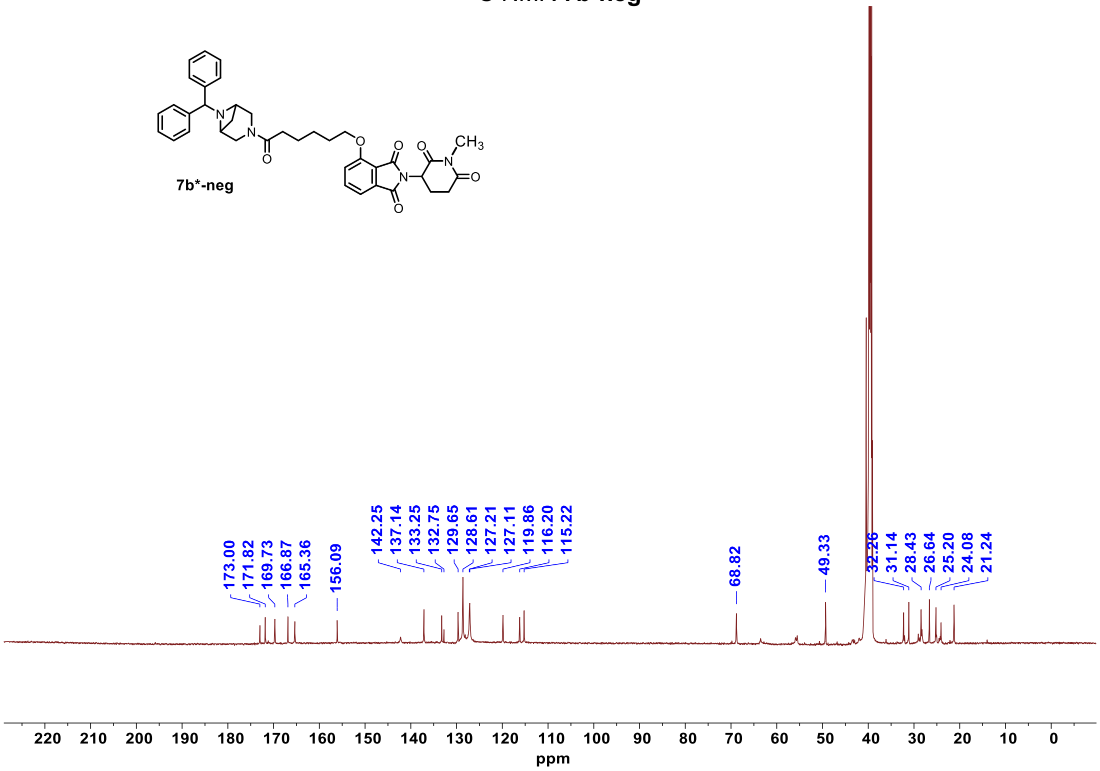

[1]H NMR **7c**

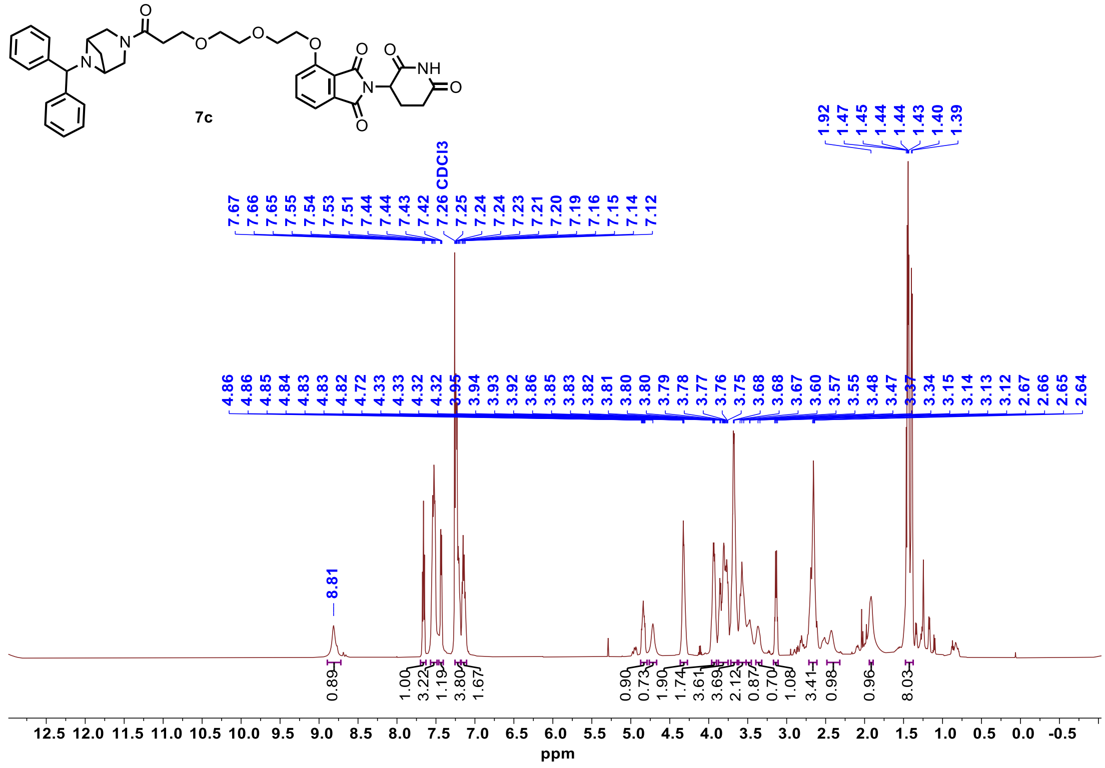


[13]C NMR **7c**

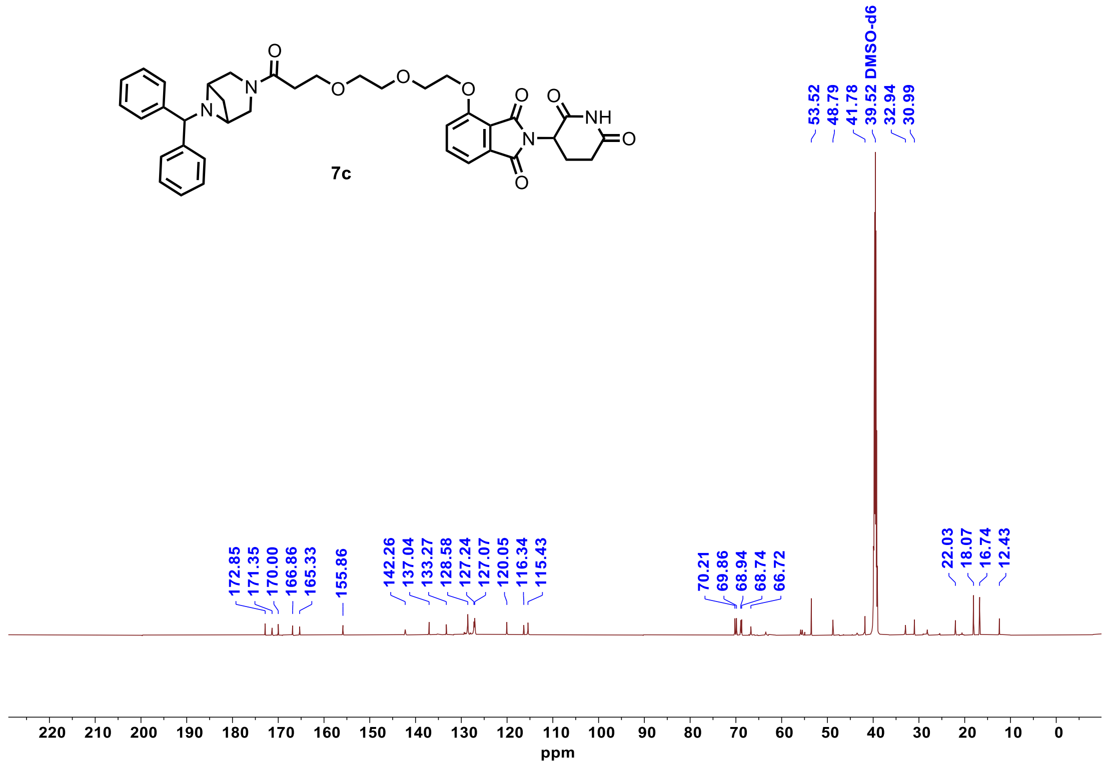

$^{1}$H NMR **7d**

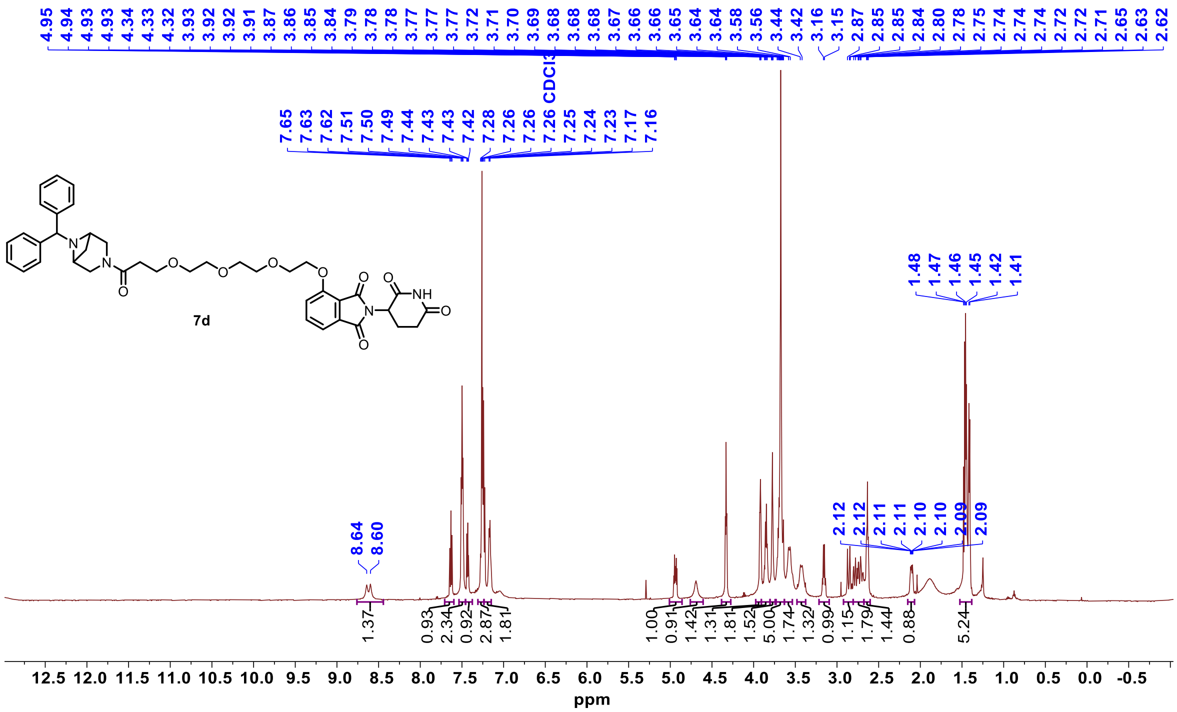


$^{13}$C NMR **7d**

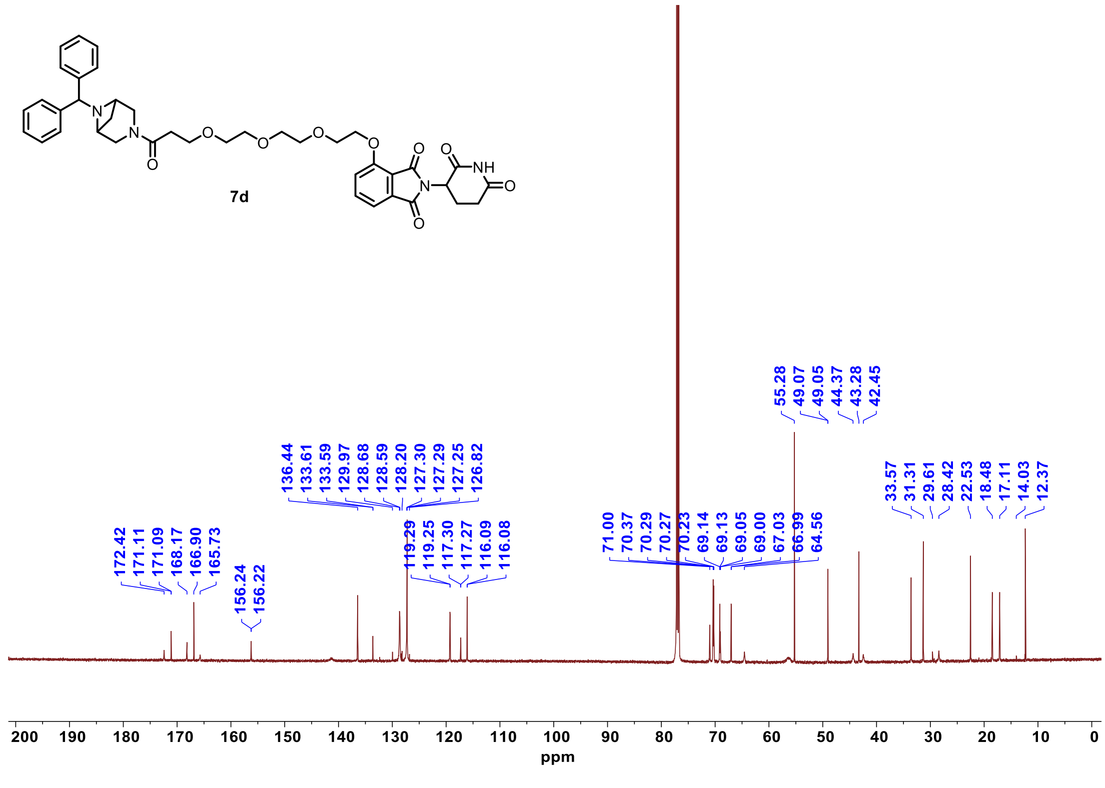

$^{1}$H NMR **8a**

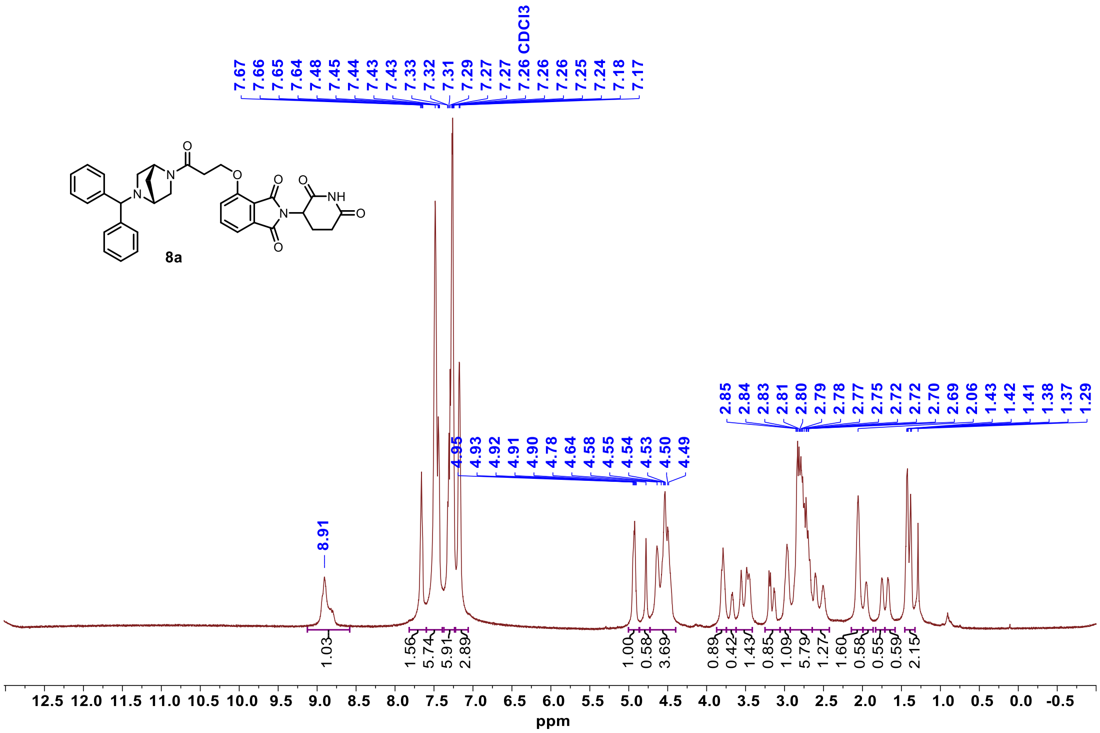


$^{13}$C NMR **8a**

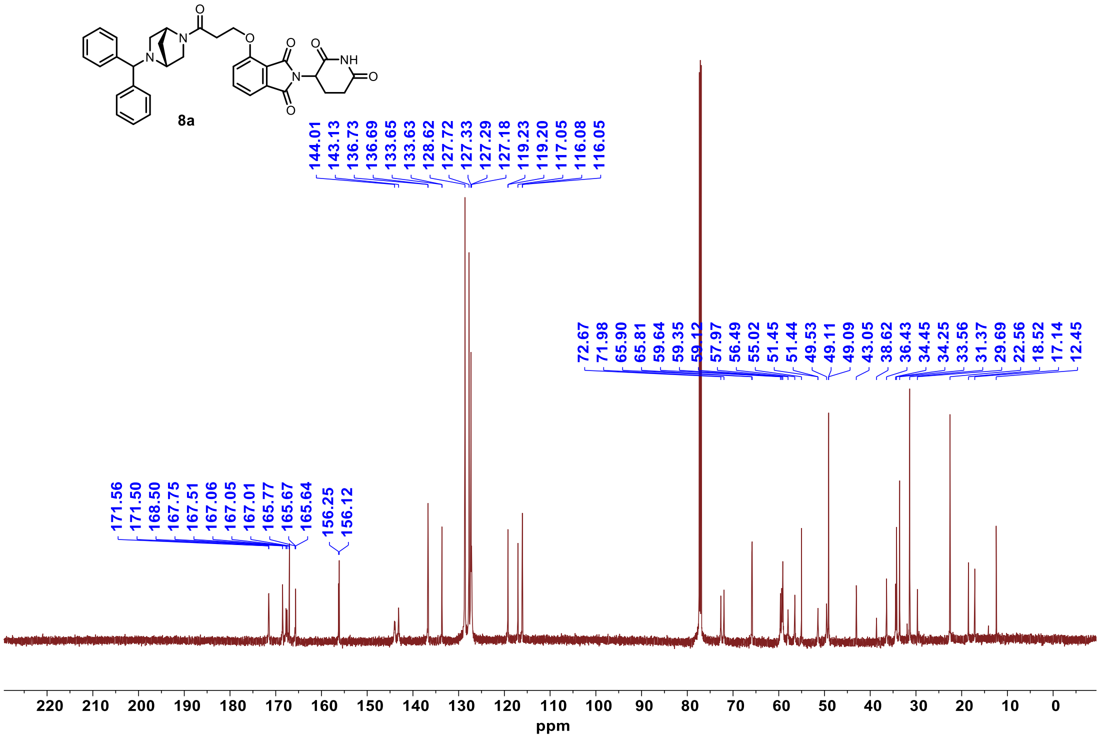

$^{1}$H NMR **8b**

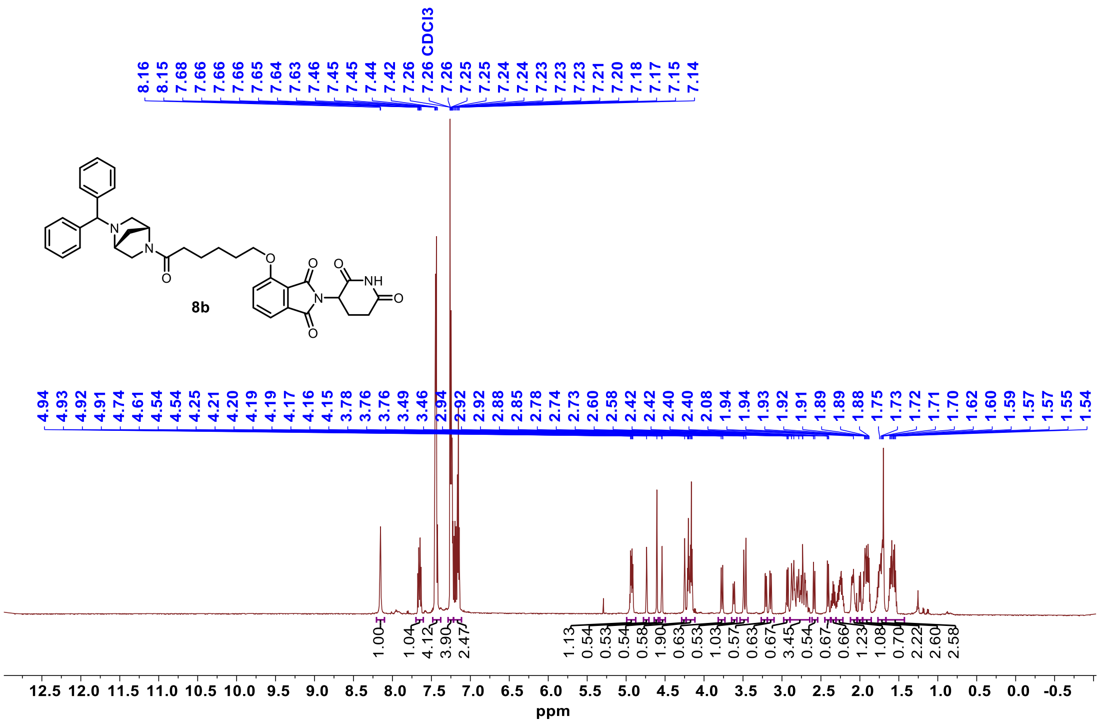


$^{13}$C NMR **8b**

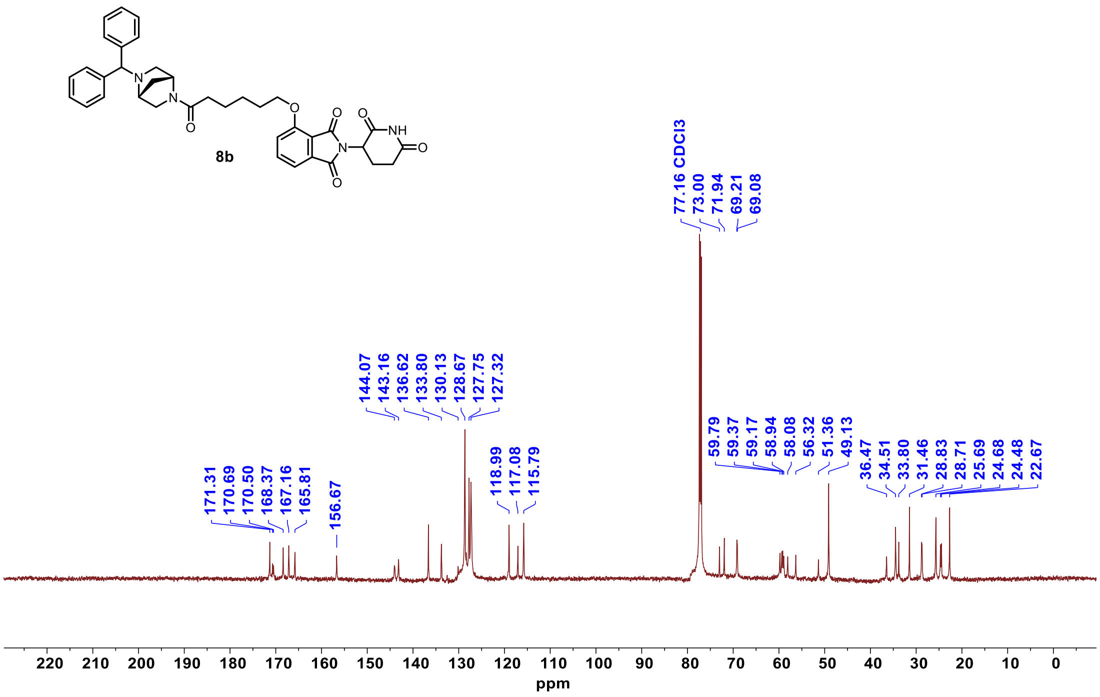

#### $^{1}$H NMR **8c**

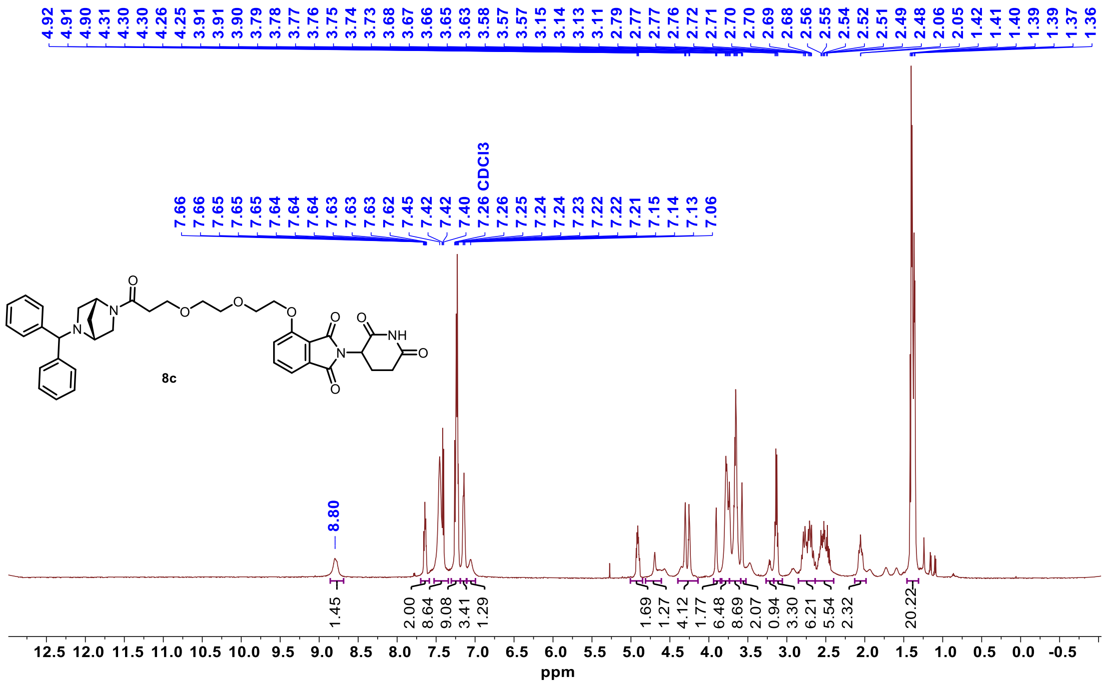


#### $^{13}$C NMR **8c**

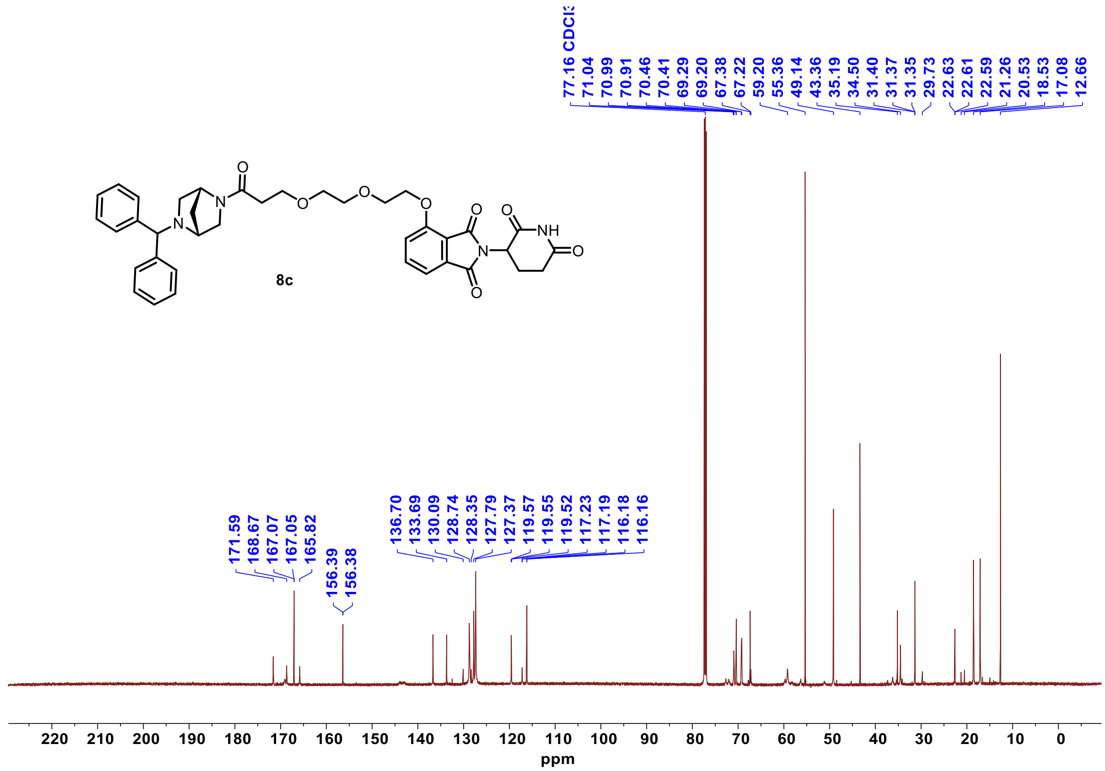

$^{1}H$ NMR **8d**

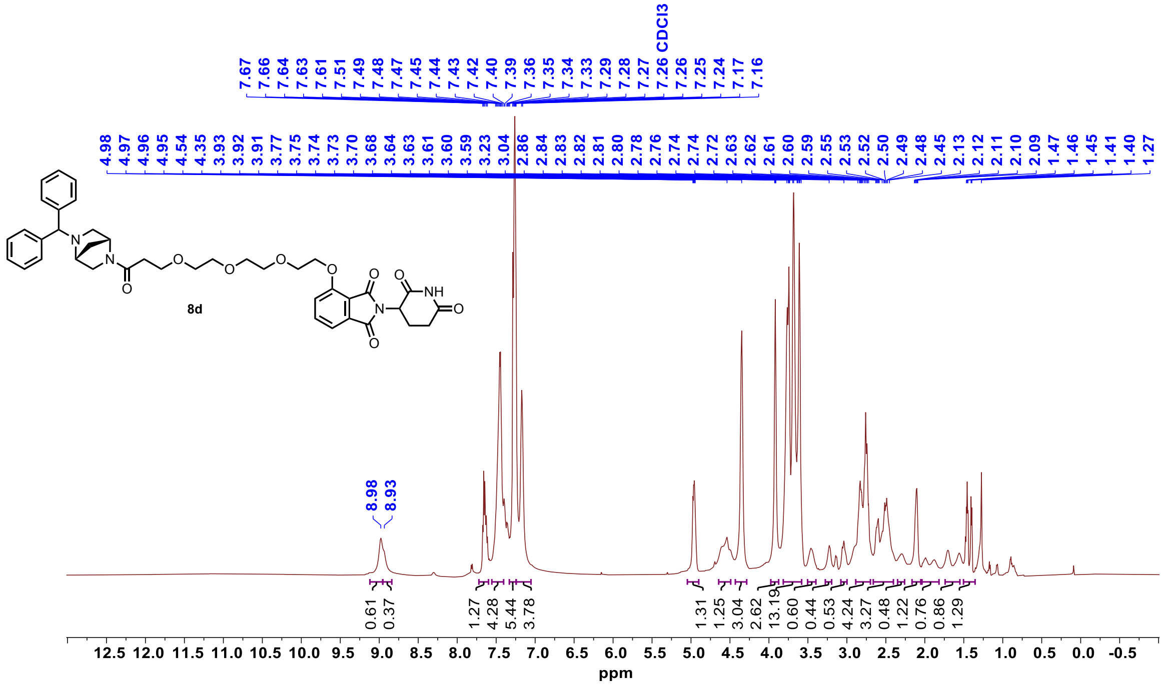


$^{13}C$ NMR **8d**

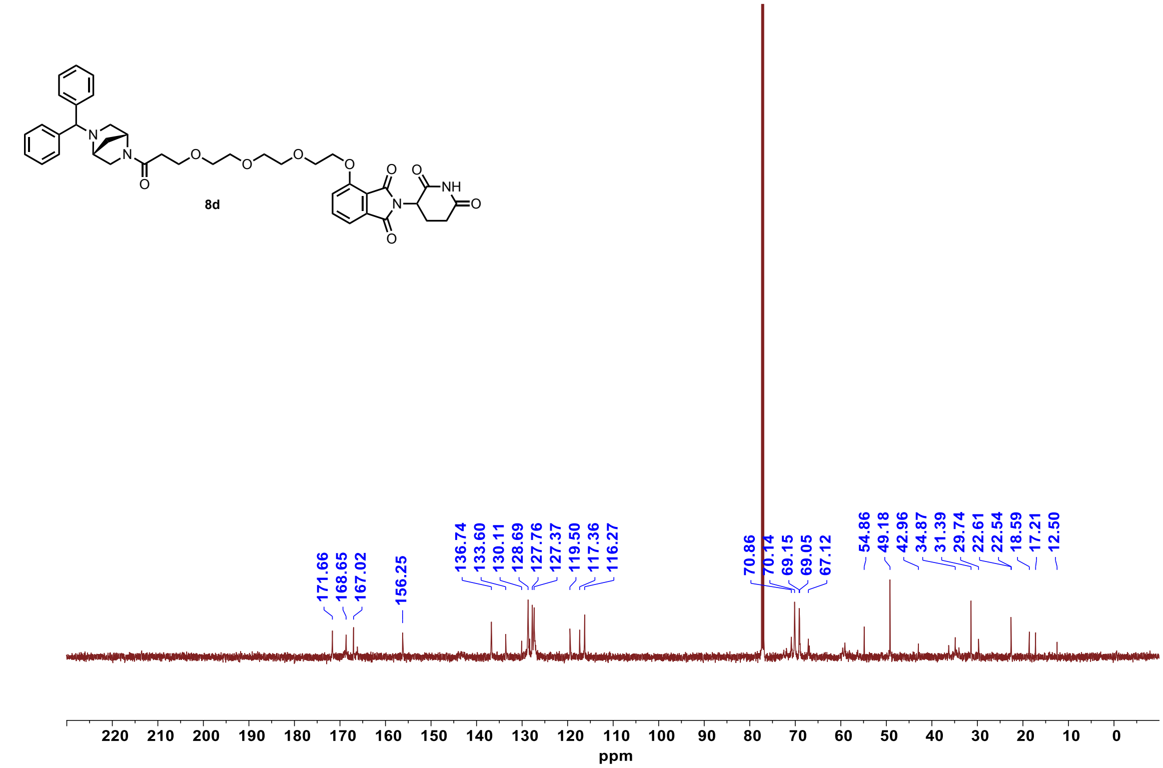

#### [1]H NMR **8d*-neg**

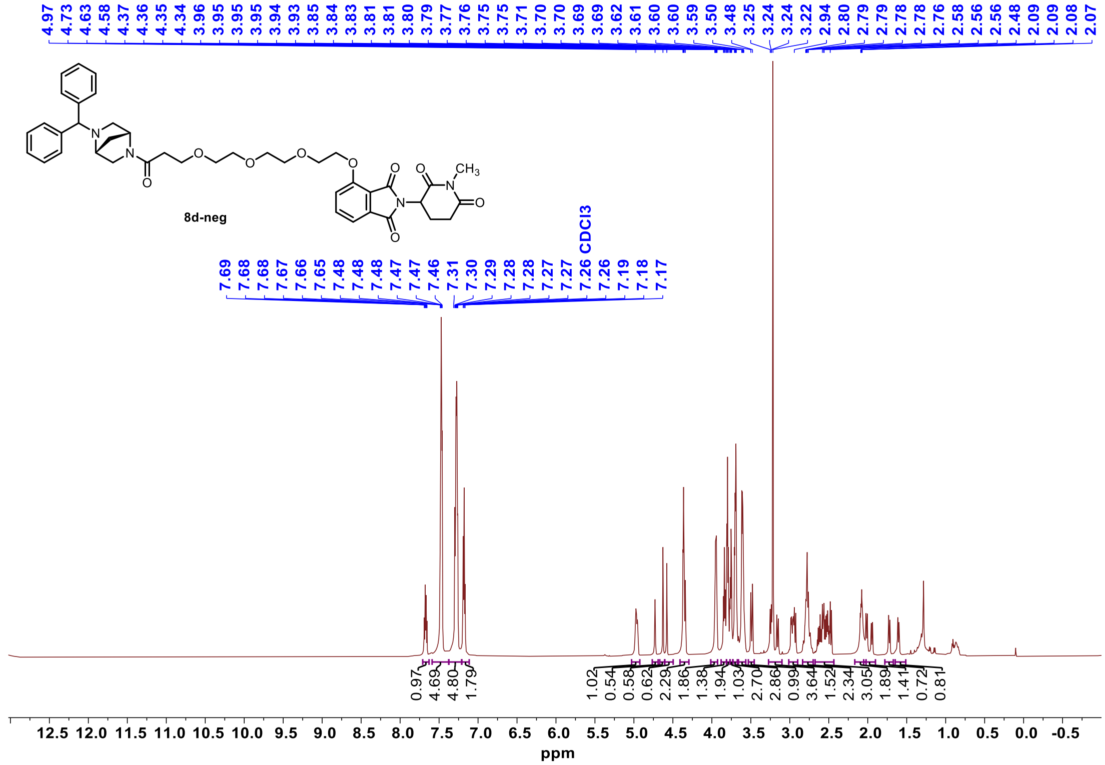


#### [13]C NMR **8d*-neg**

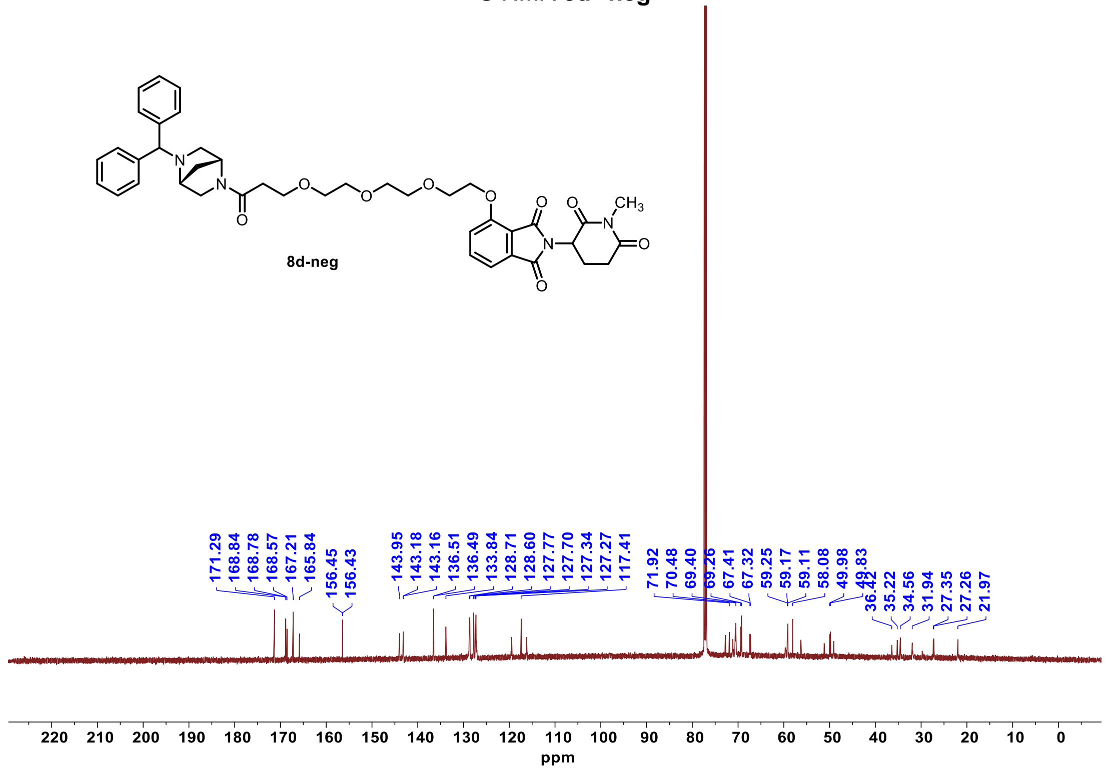

$^{1}H$ NMR **9a**

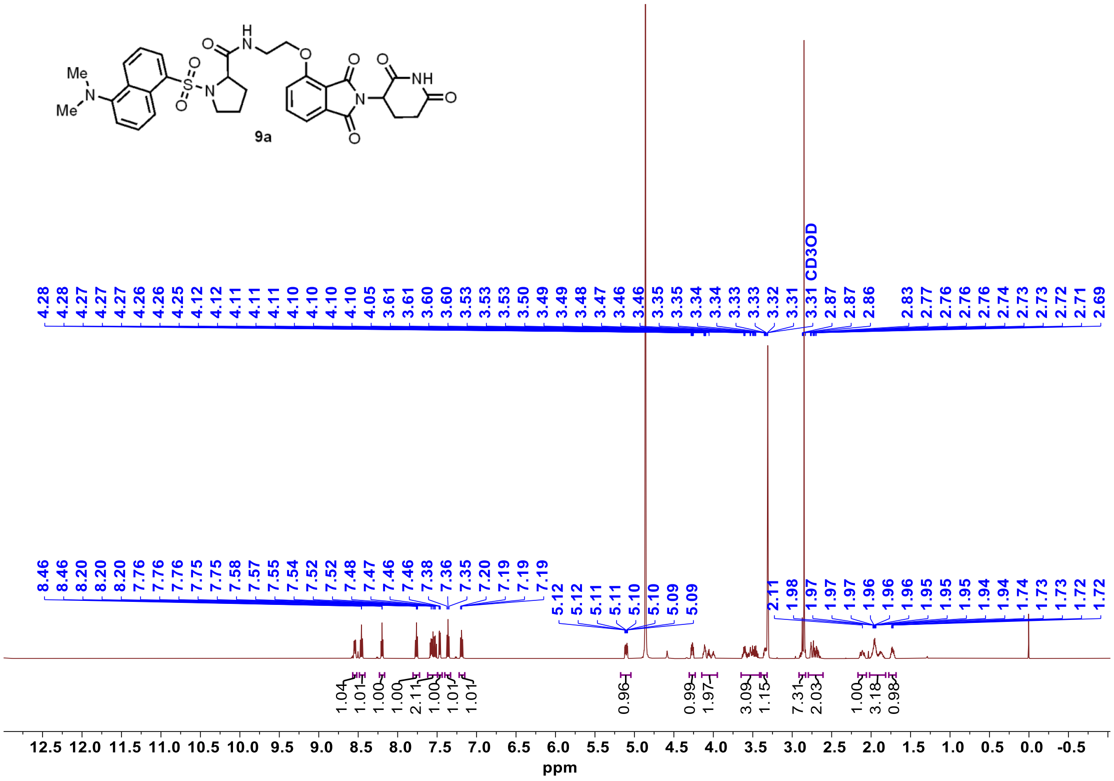


$^{13}C$ NMR **9a**

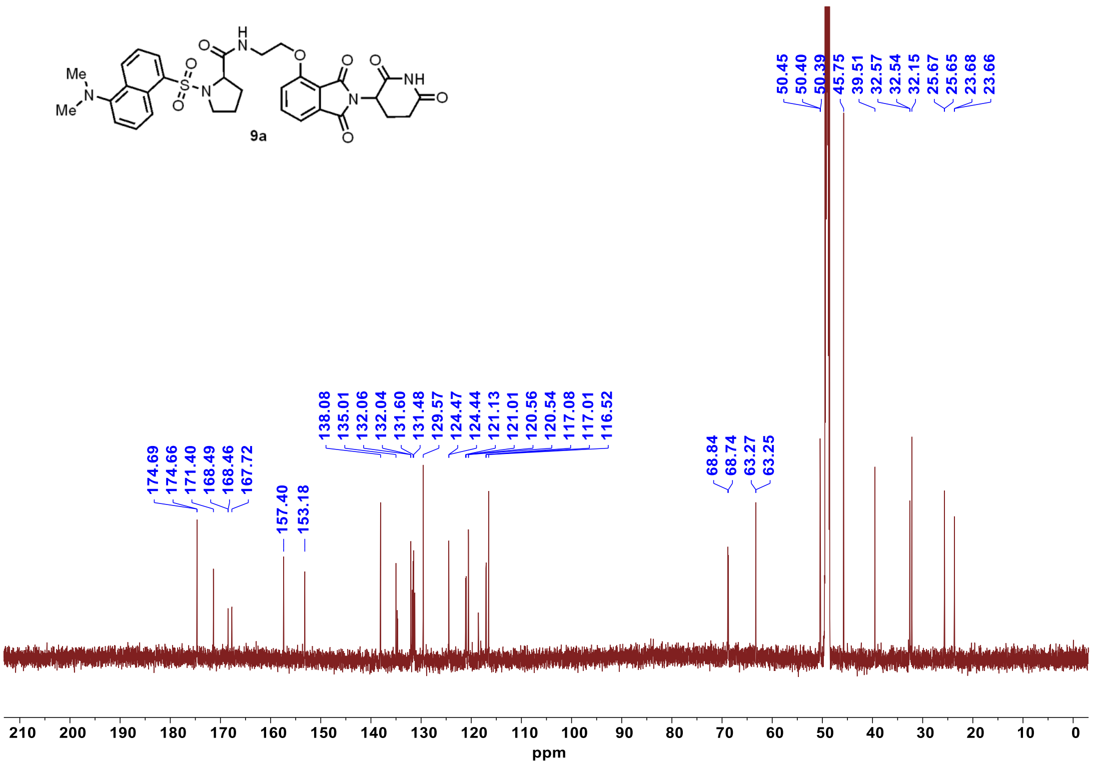

$^{1}$H NMR **9b**

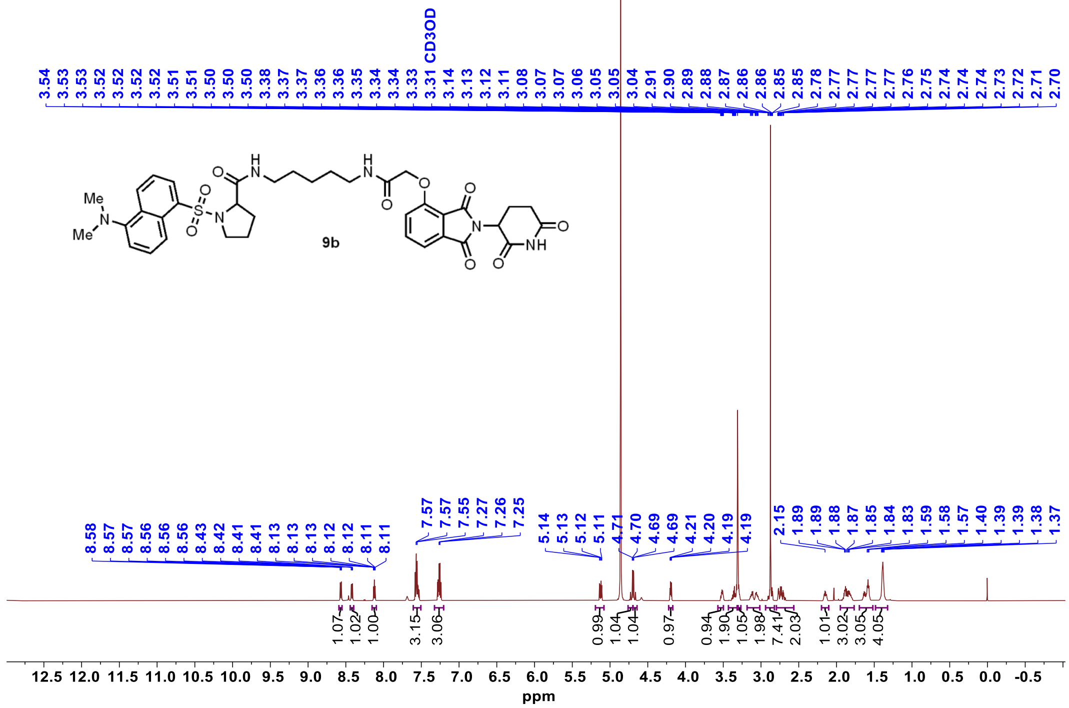


$^{13}$C NMR **9b**

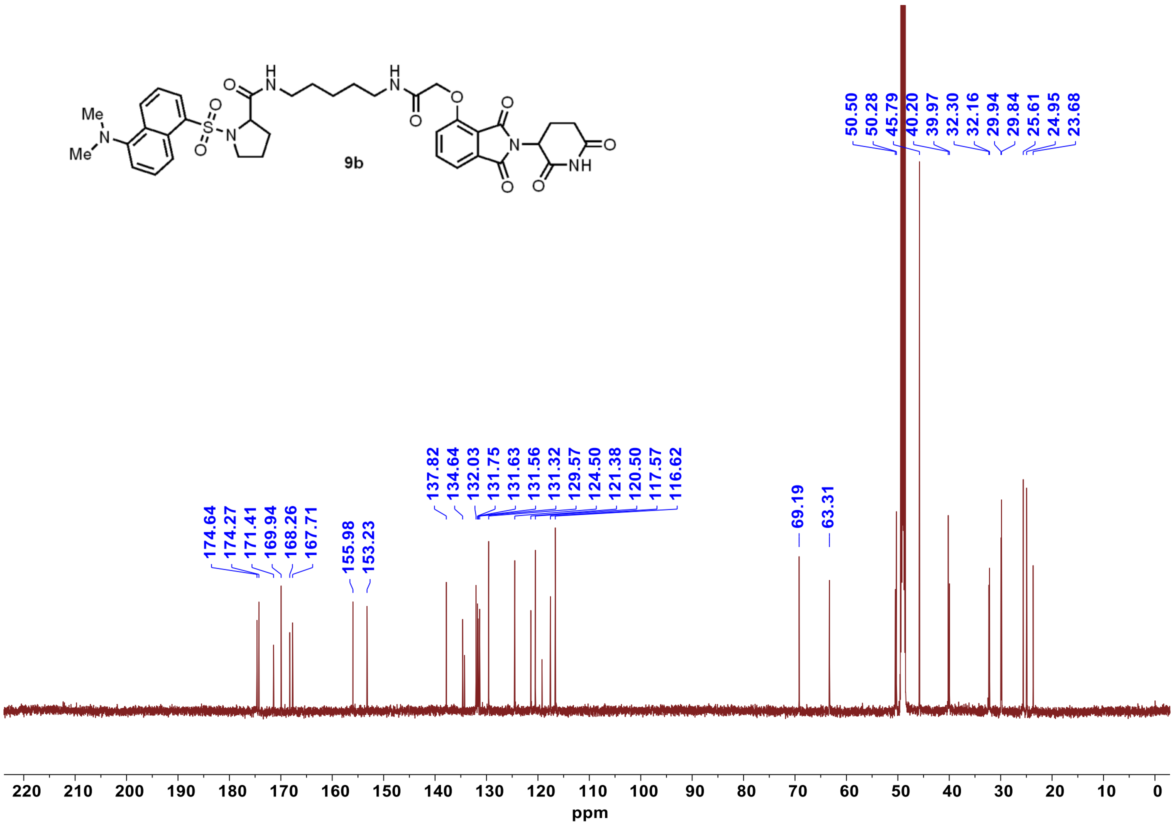

[1]H NMR **9c**

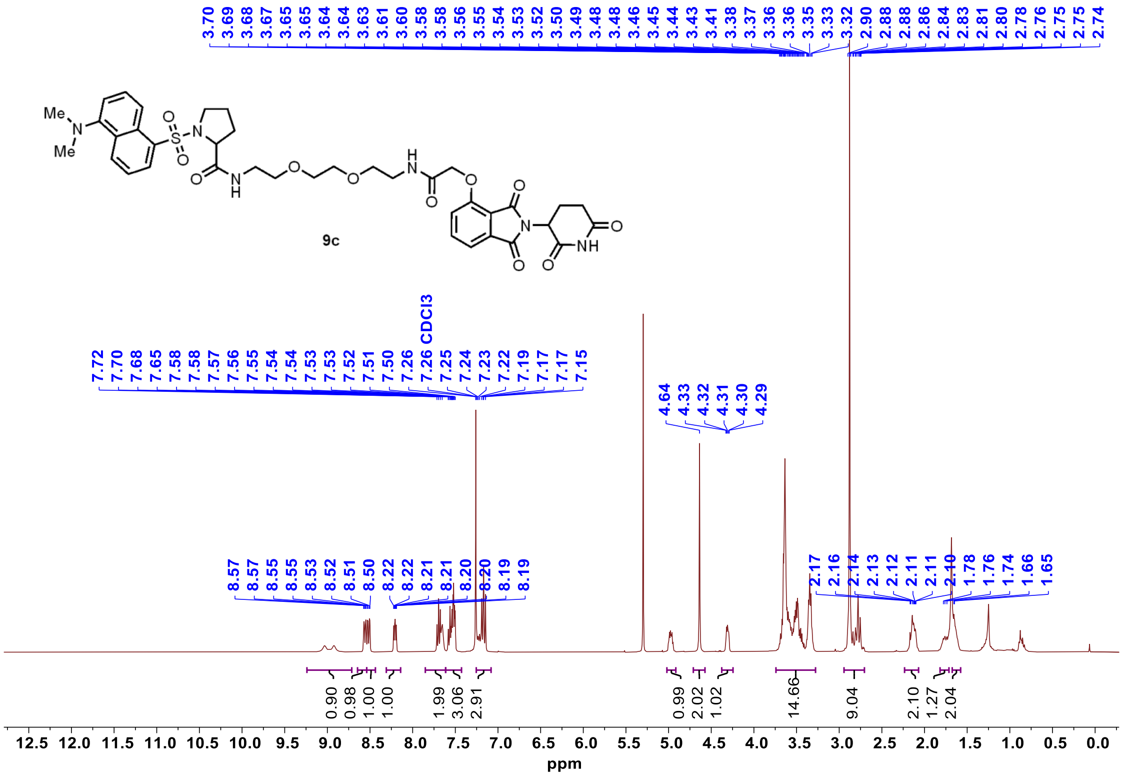


[13]C NMR **9c**

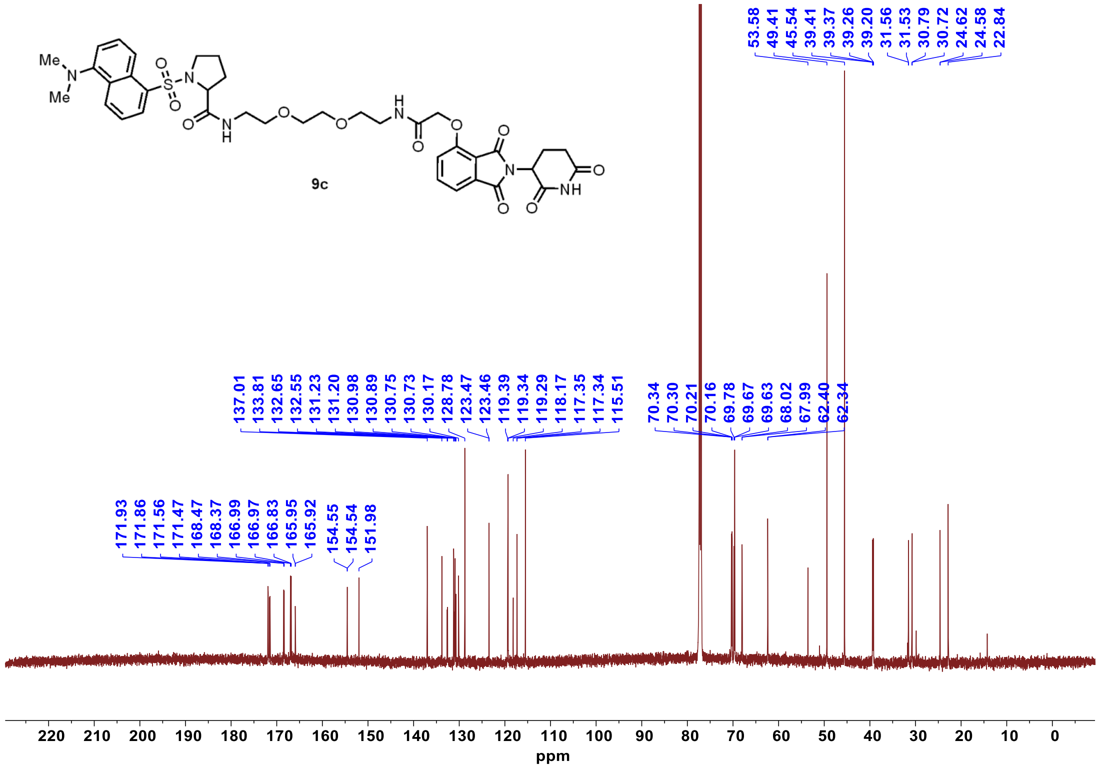

$^{1}$H NMR **9d**

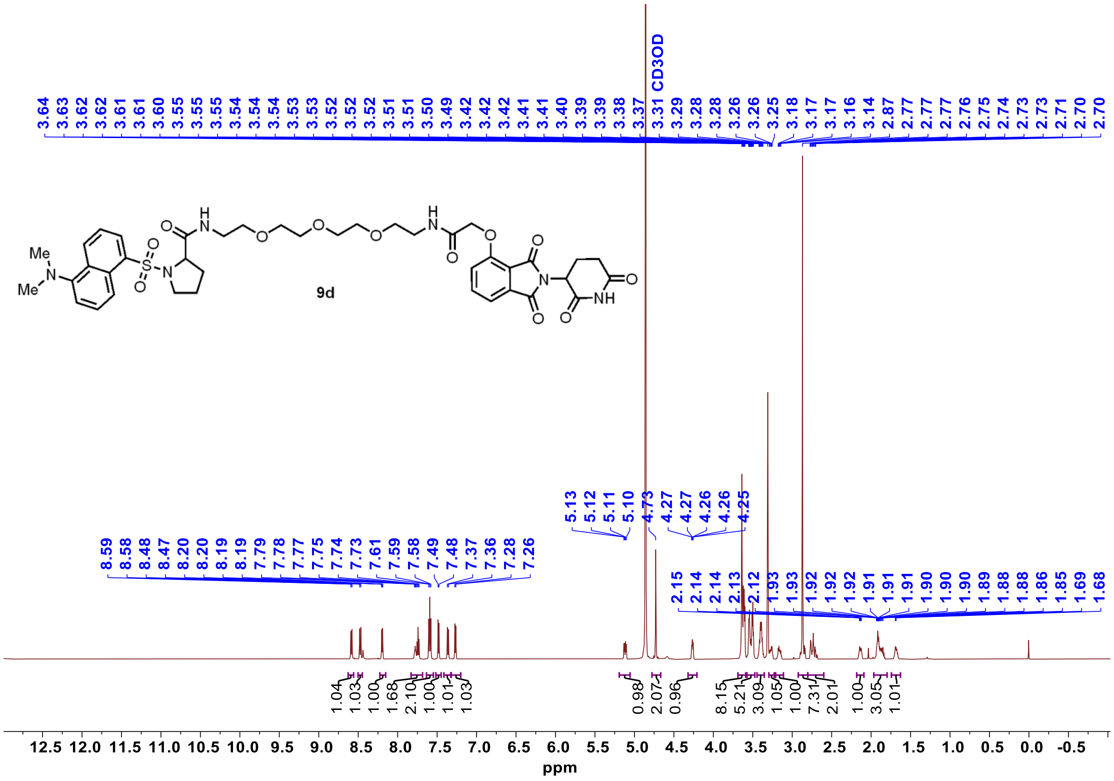


$^{13}$C NMR **9d**

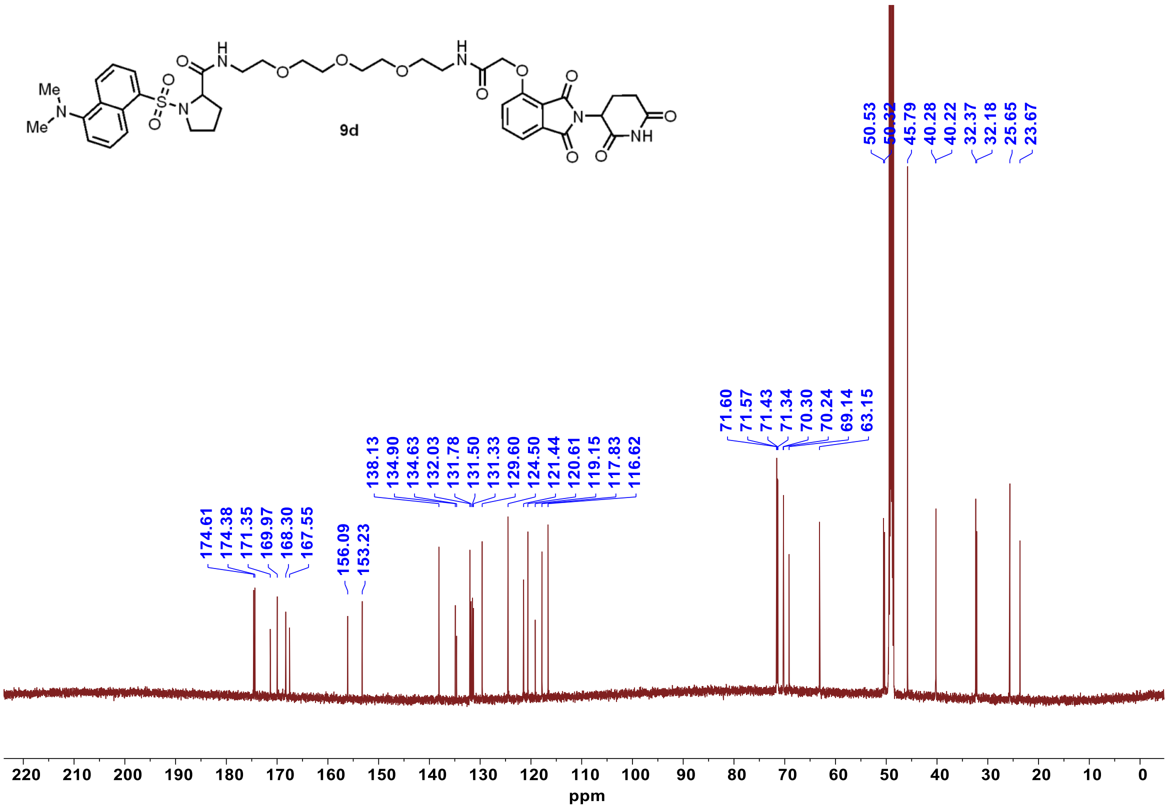

$^1$H NMR **10a**

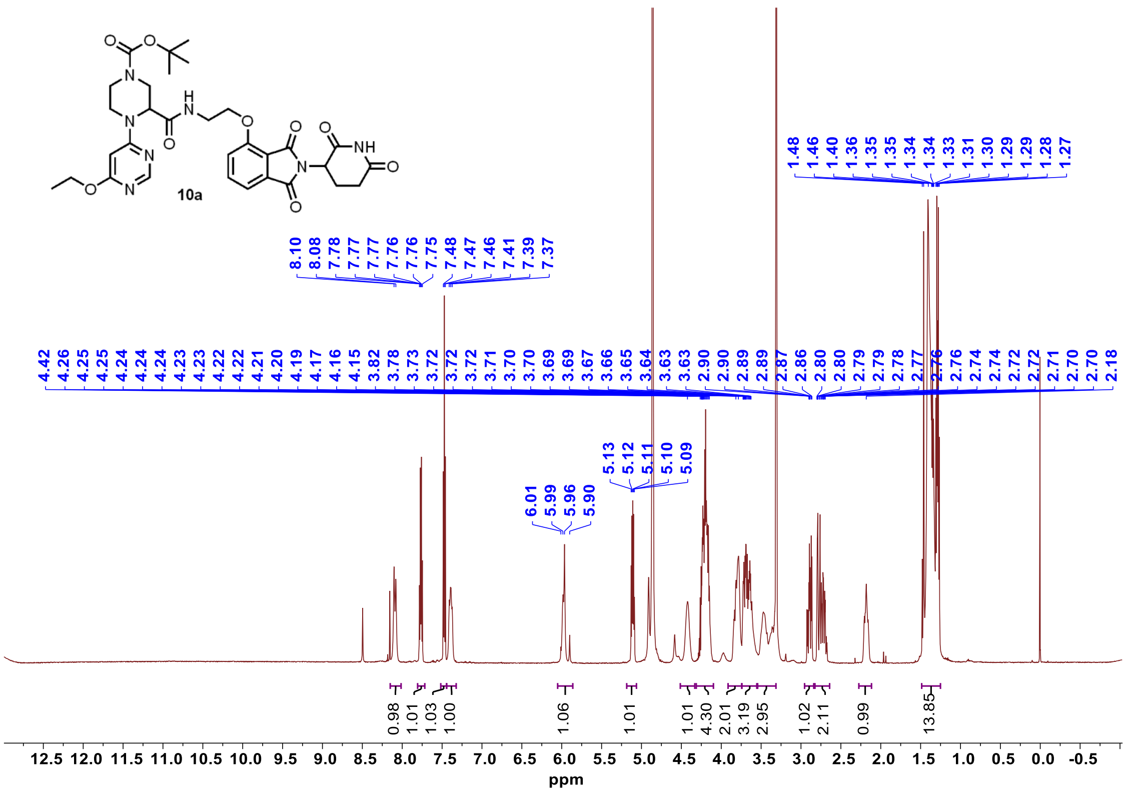


$^{13}$C NMR **10a**

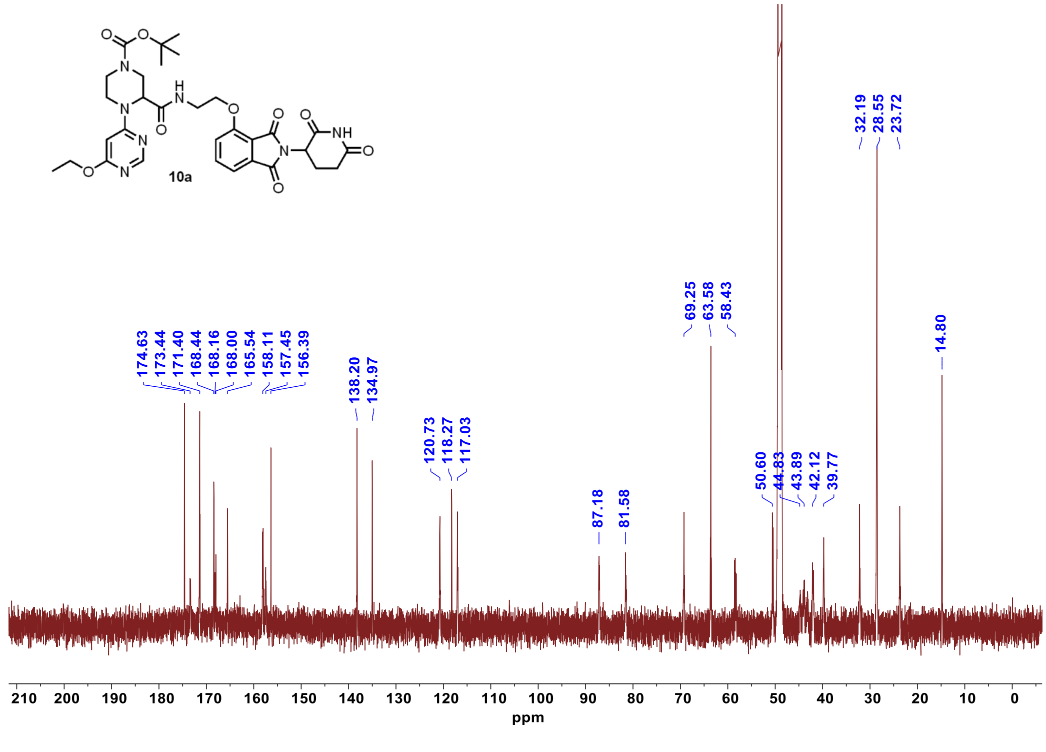

<sup></sup>
#### $^{1}$H NMR **10b**

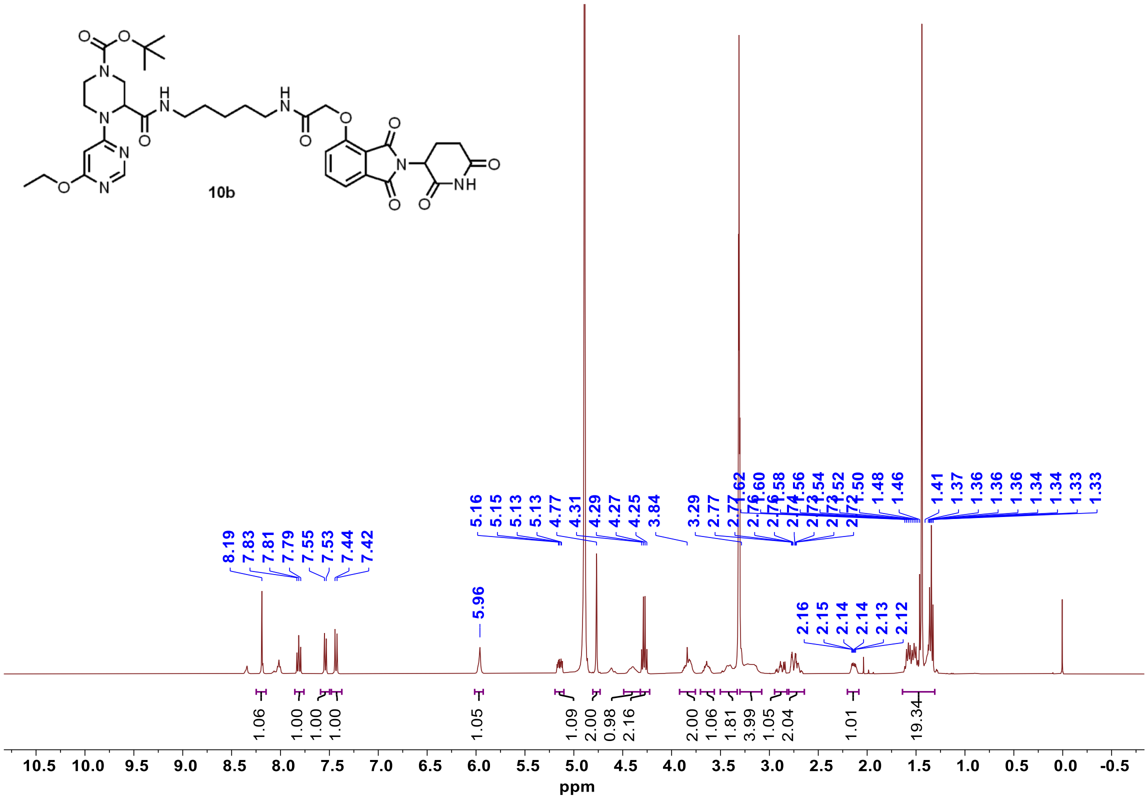


#### $^{13}$C NMR **10b**

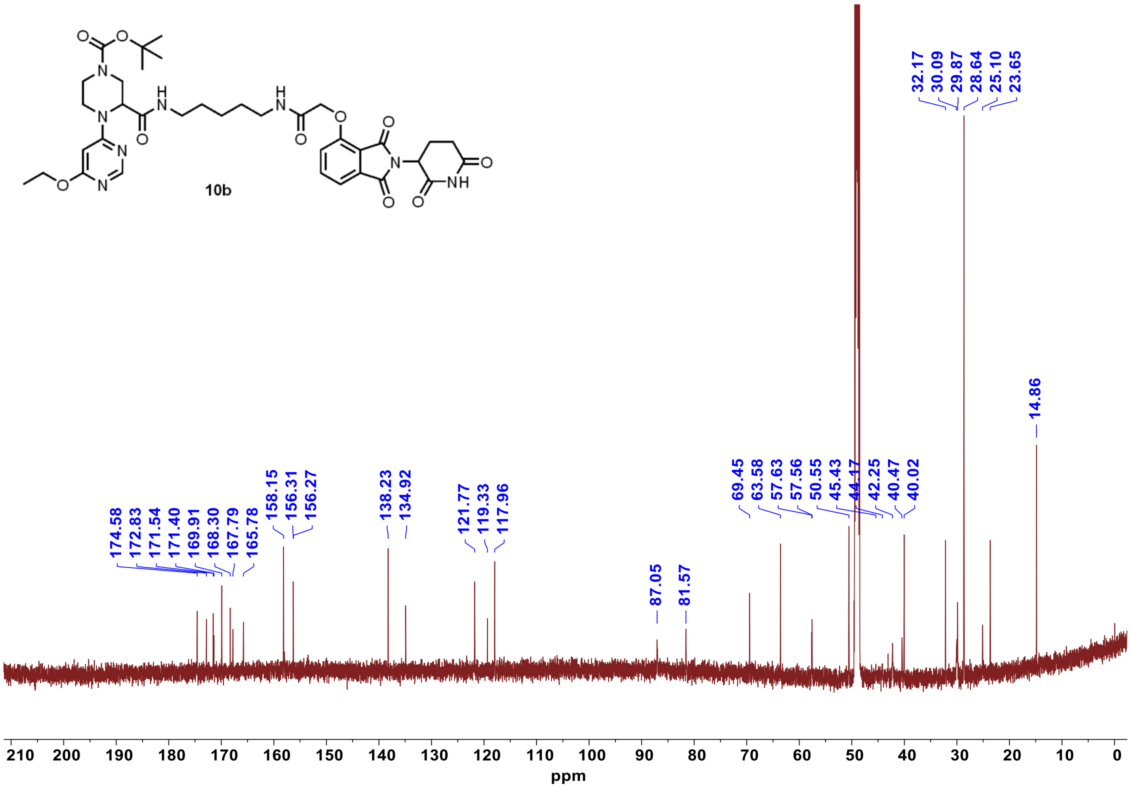

[1]H NMR **10c**

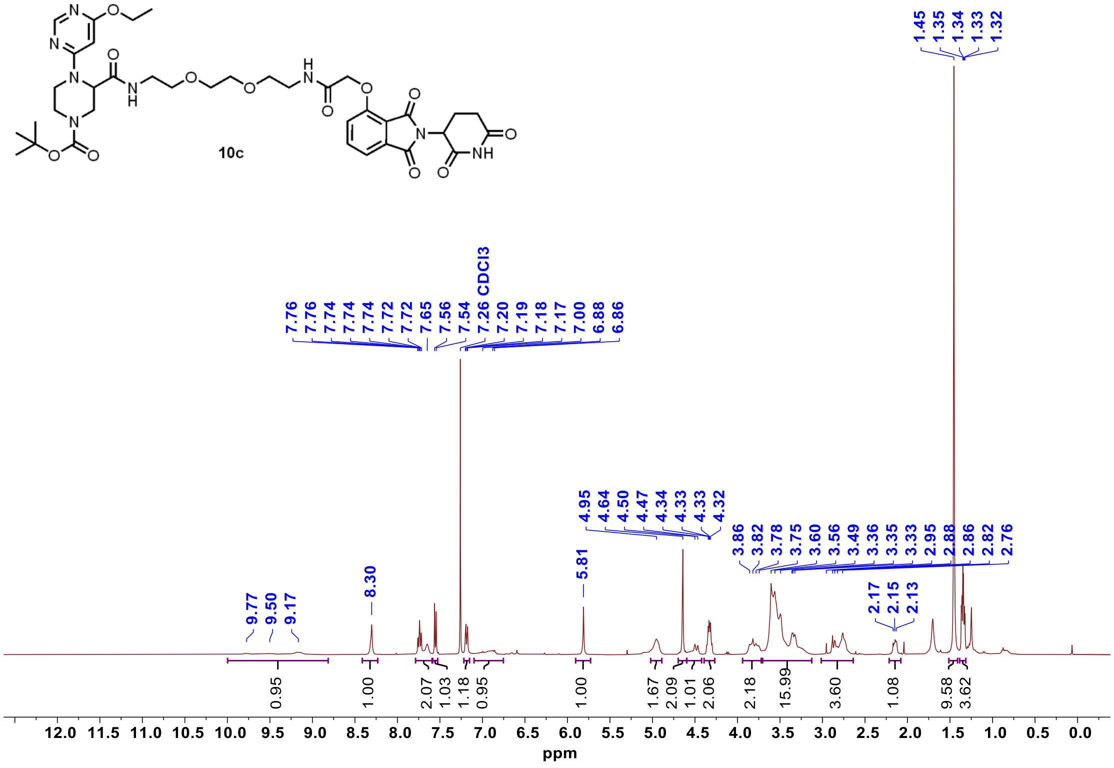


[13]C NMR **10c**

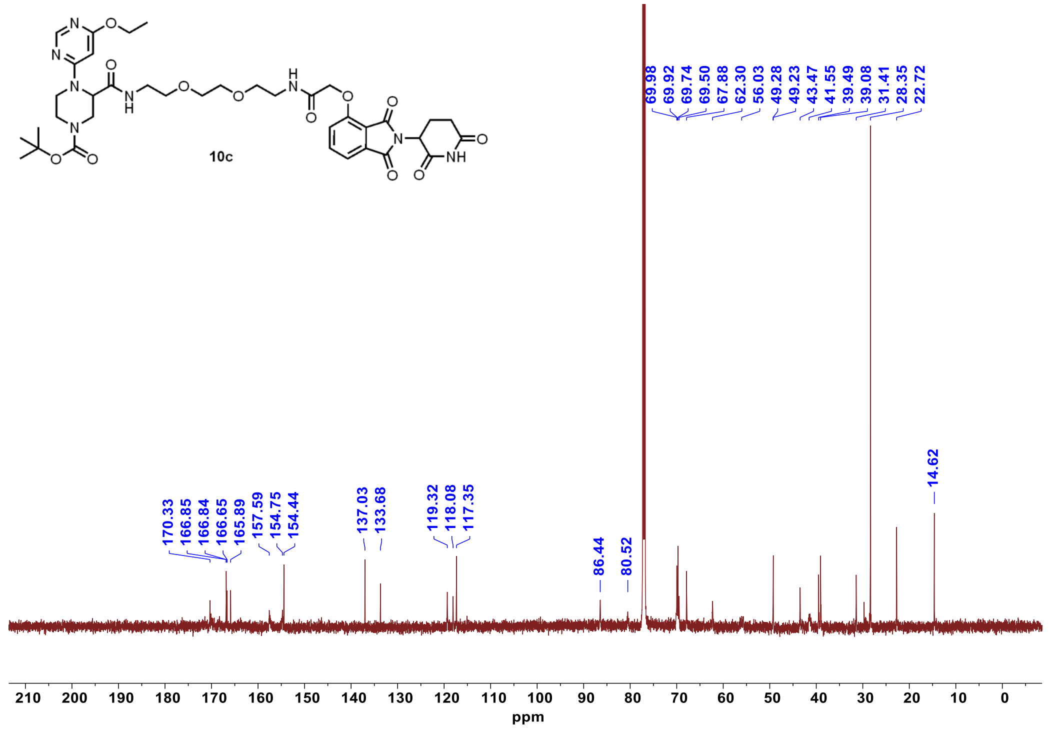

$^{1}$H NMR **10d**

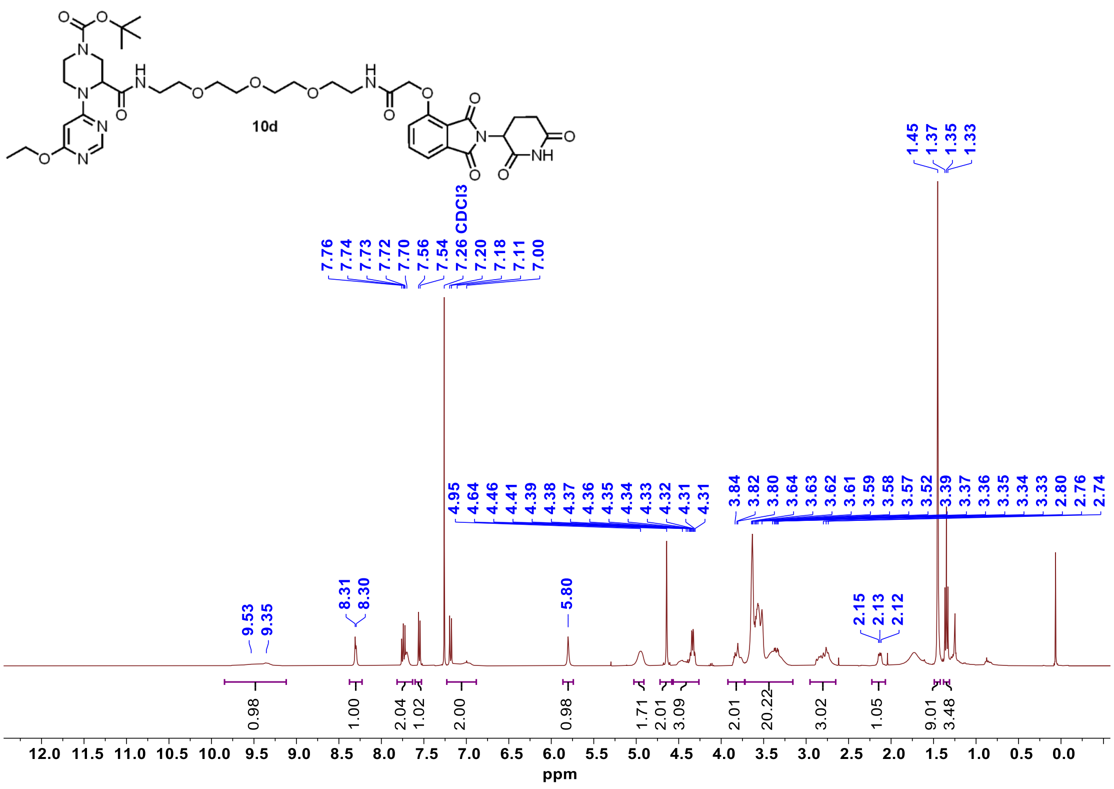


$^{13}$C NMR **10d**

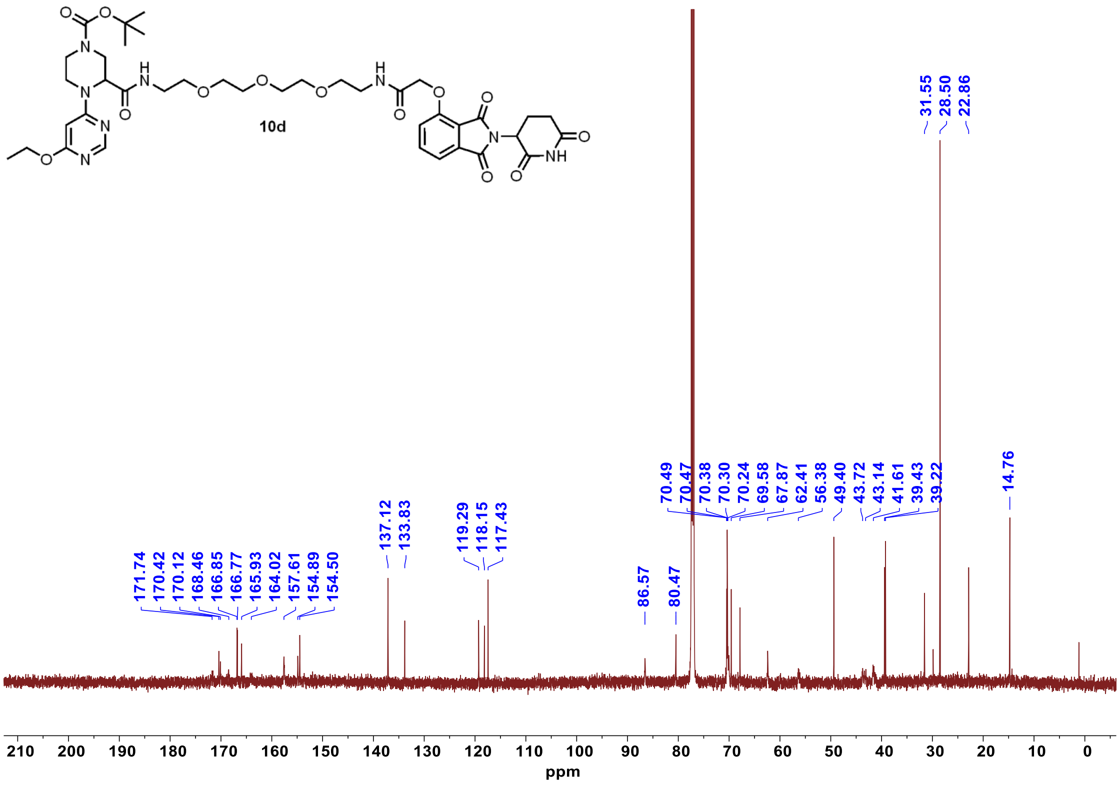

#### $^{1}H$ NMR **10d-neg**

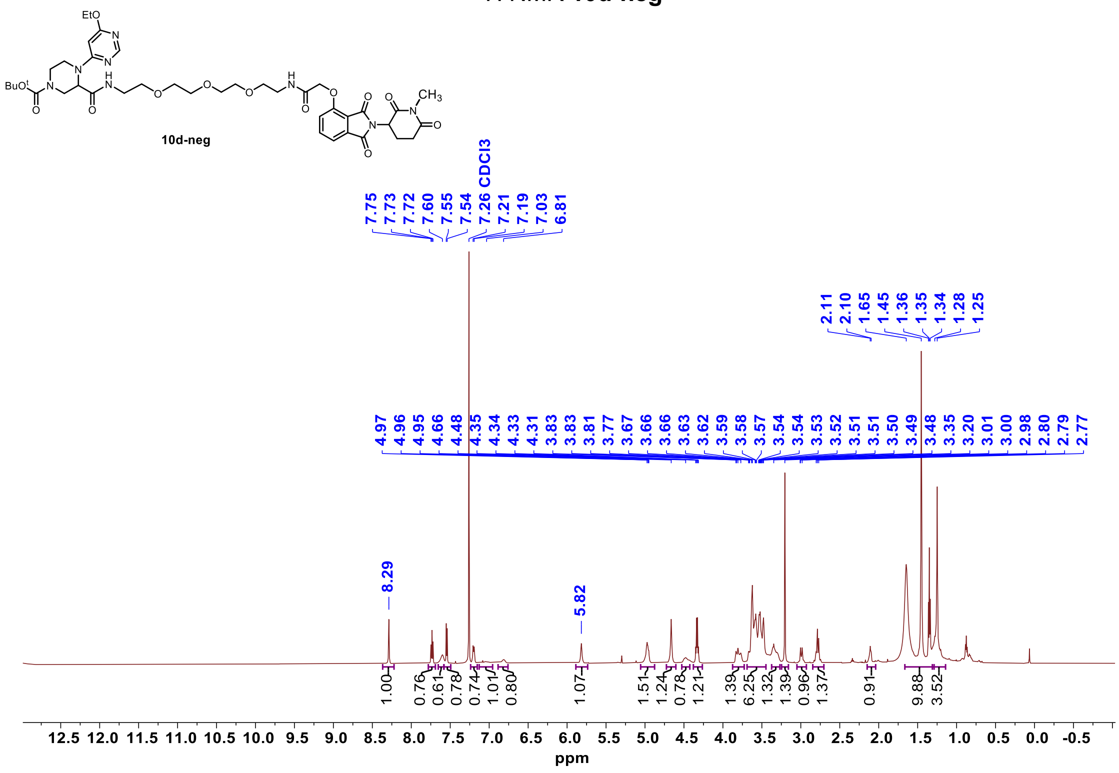


#### $^{13}C$ NMR **10d-neg**

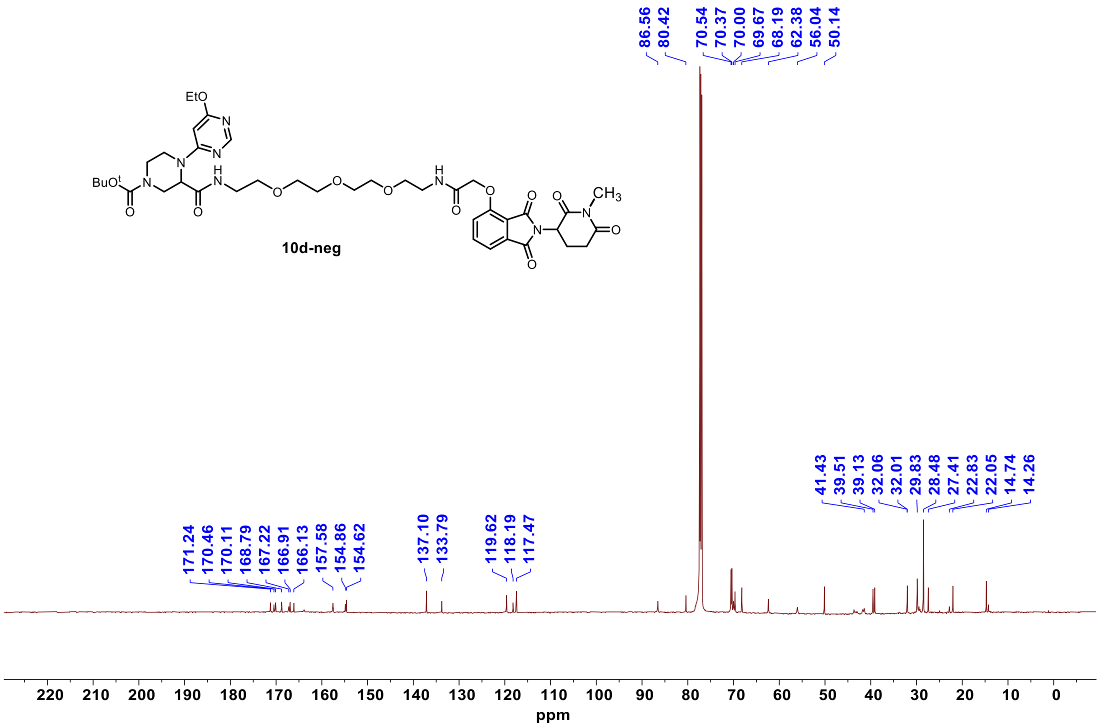

$^{1}$H NMR **11a**

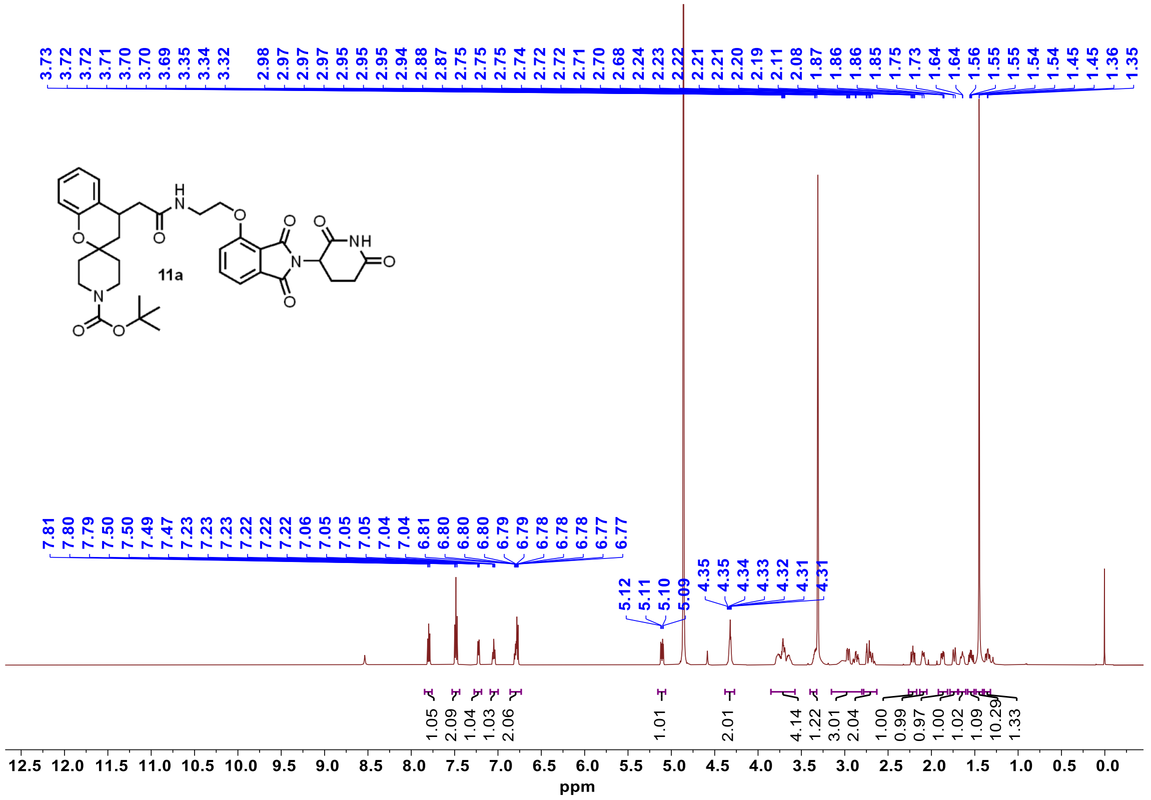


$^{13}$C NMR **11a**

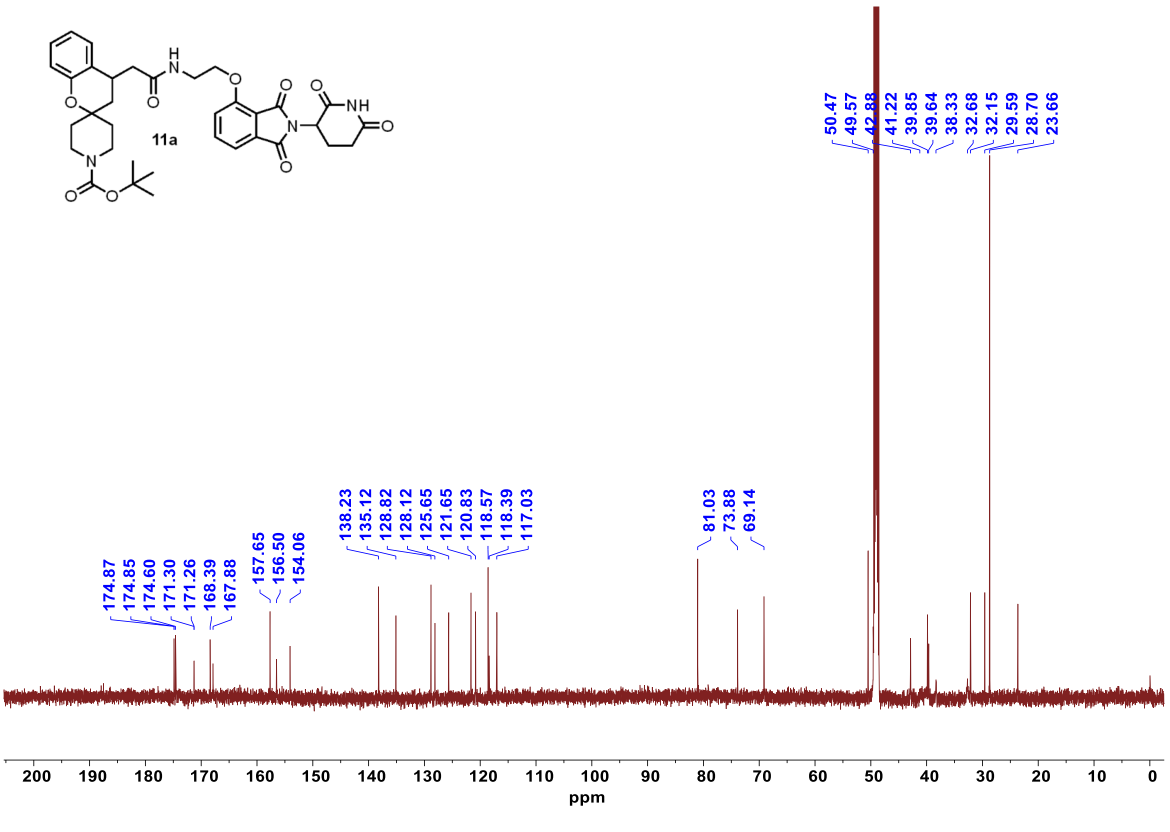

[1]H NMR **11b**

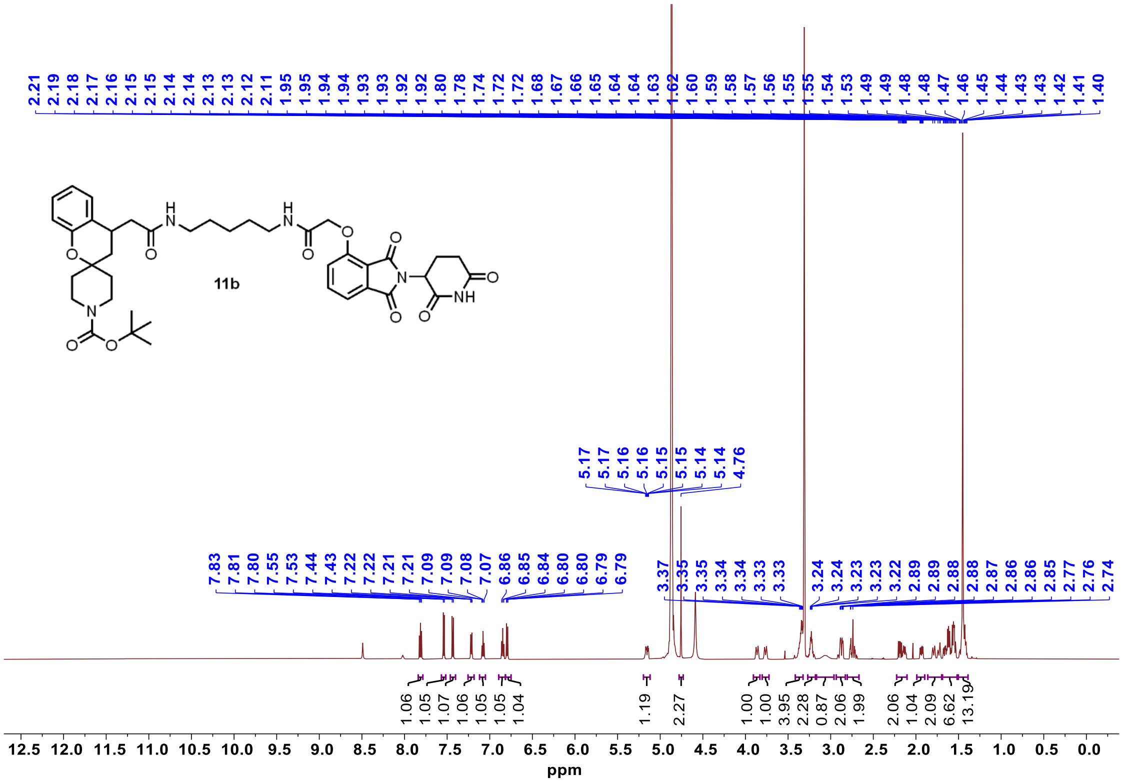


[13]C NMR **11b**

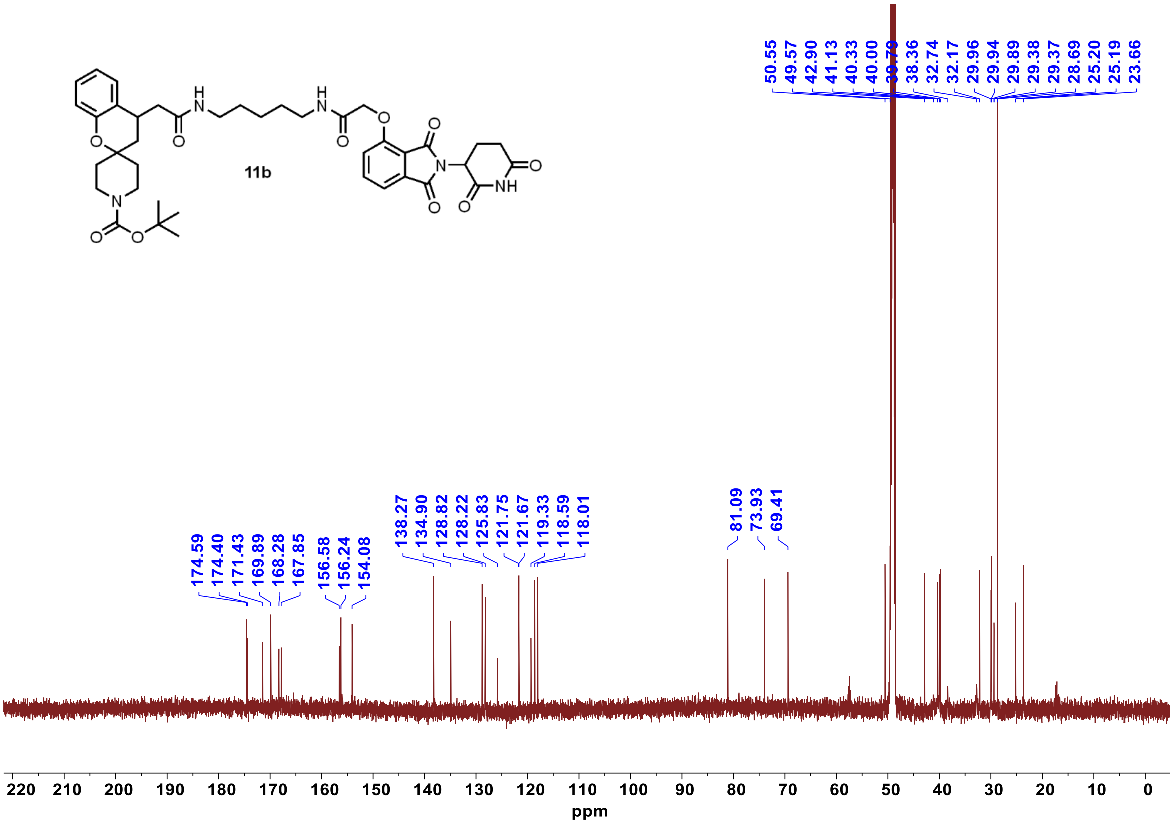

$^{1}H$ NMR **11c**

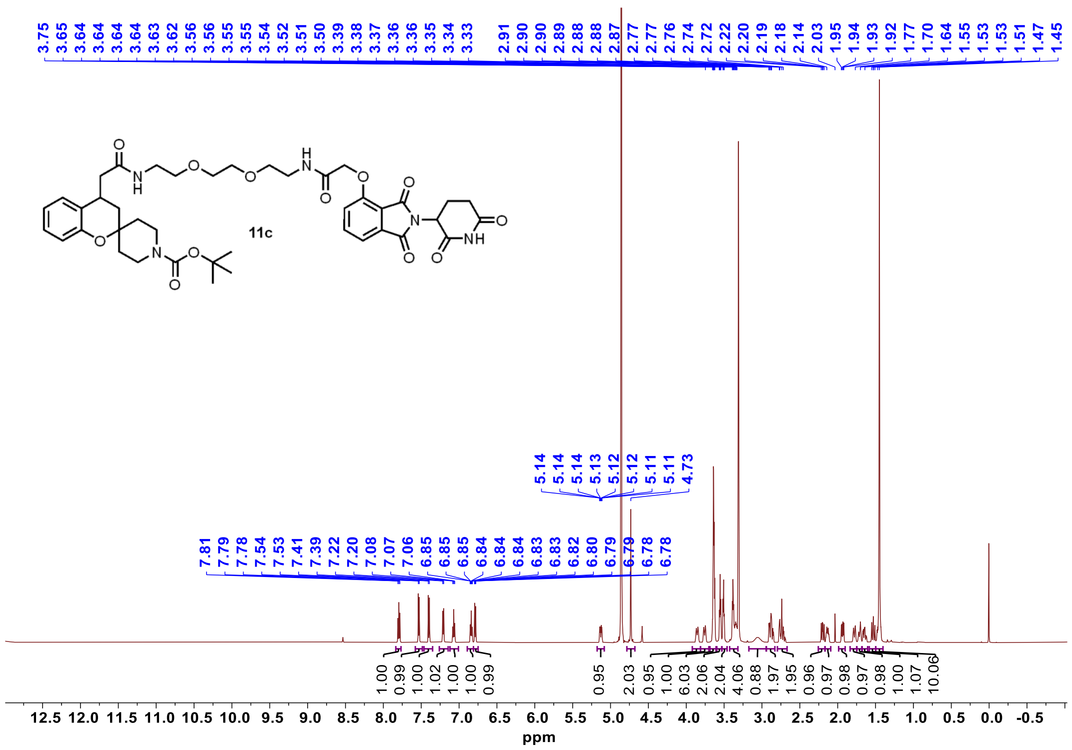


$^{13}C$ NMR **11c**

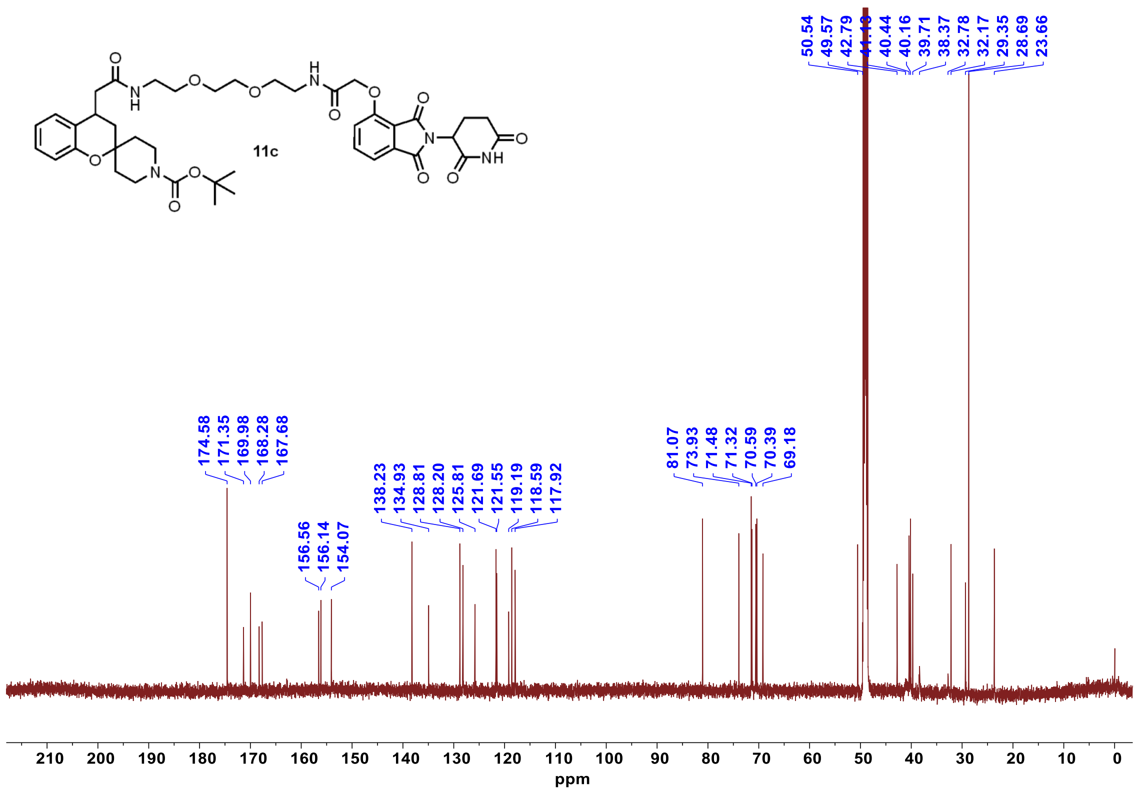

$^{1}$H NMR **11d**

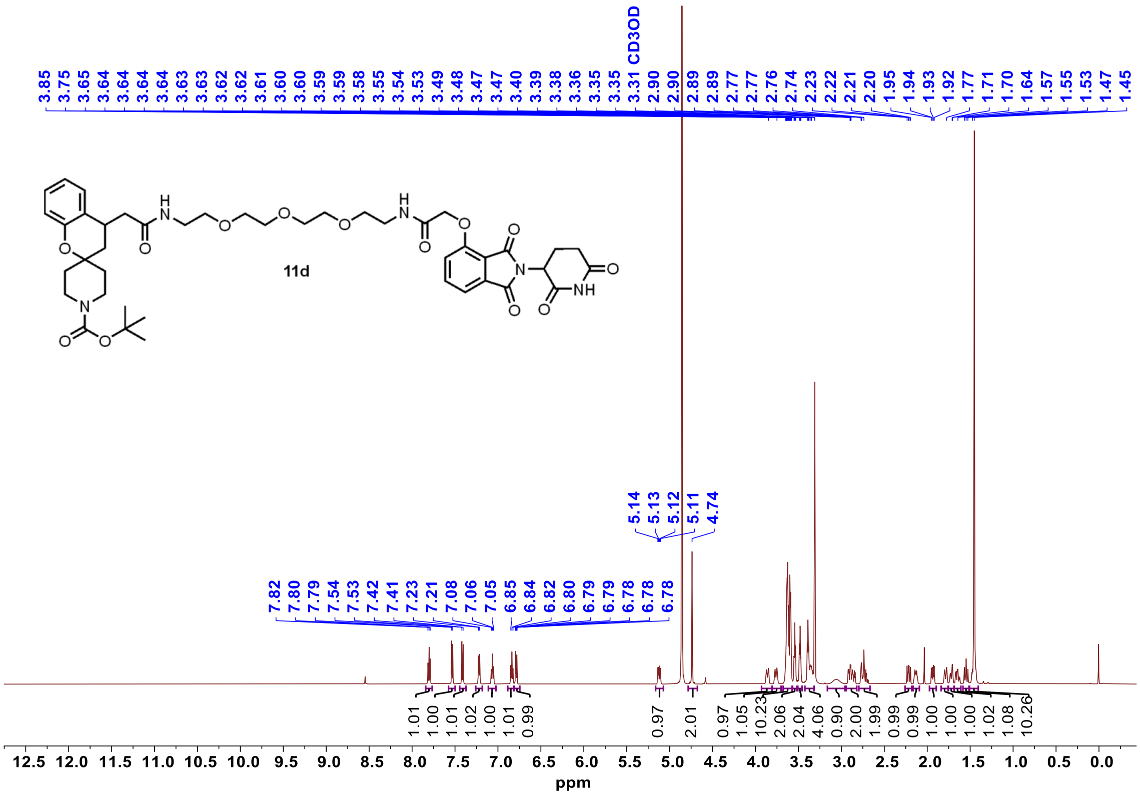


$^{13}$C NMR **11d**

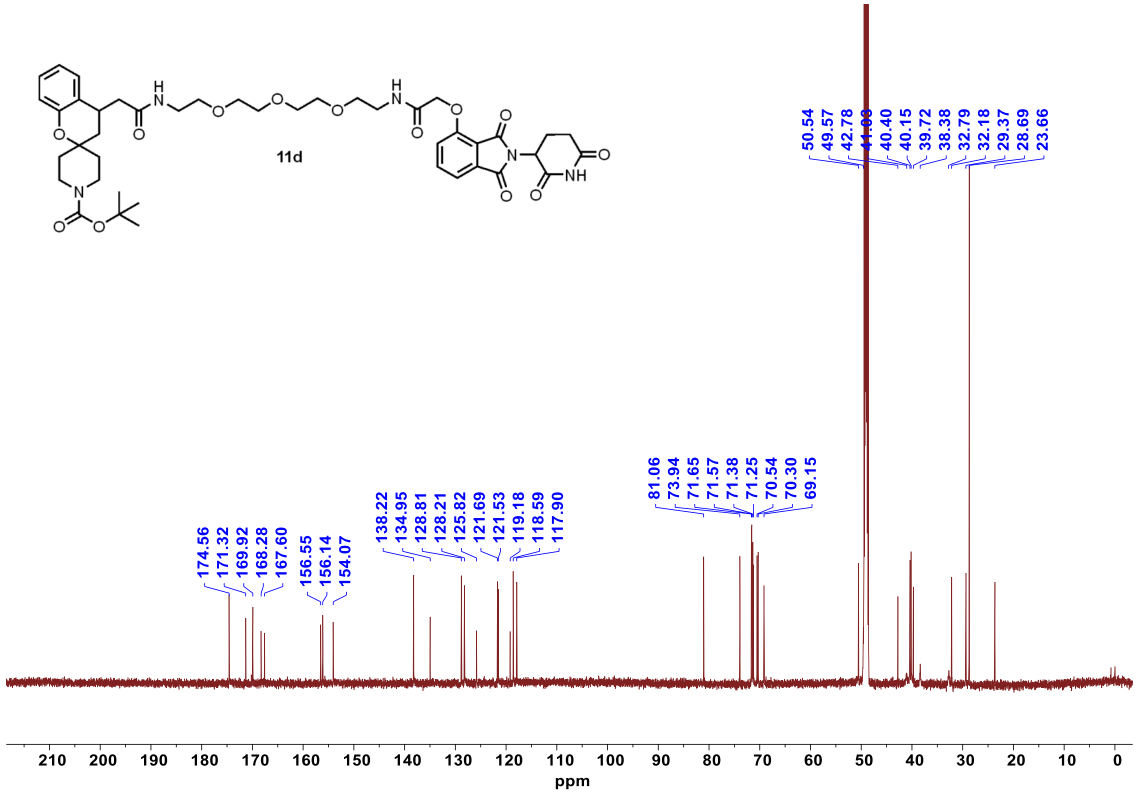

$^{1}$H NMR **12a**

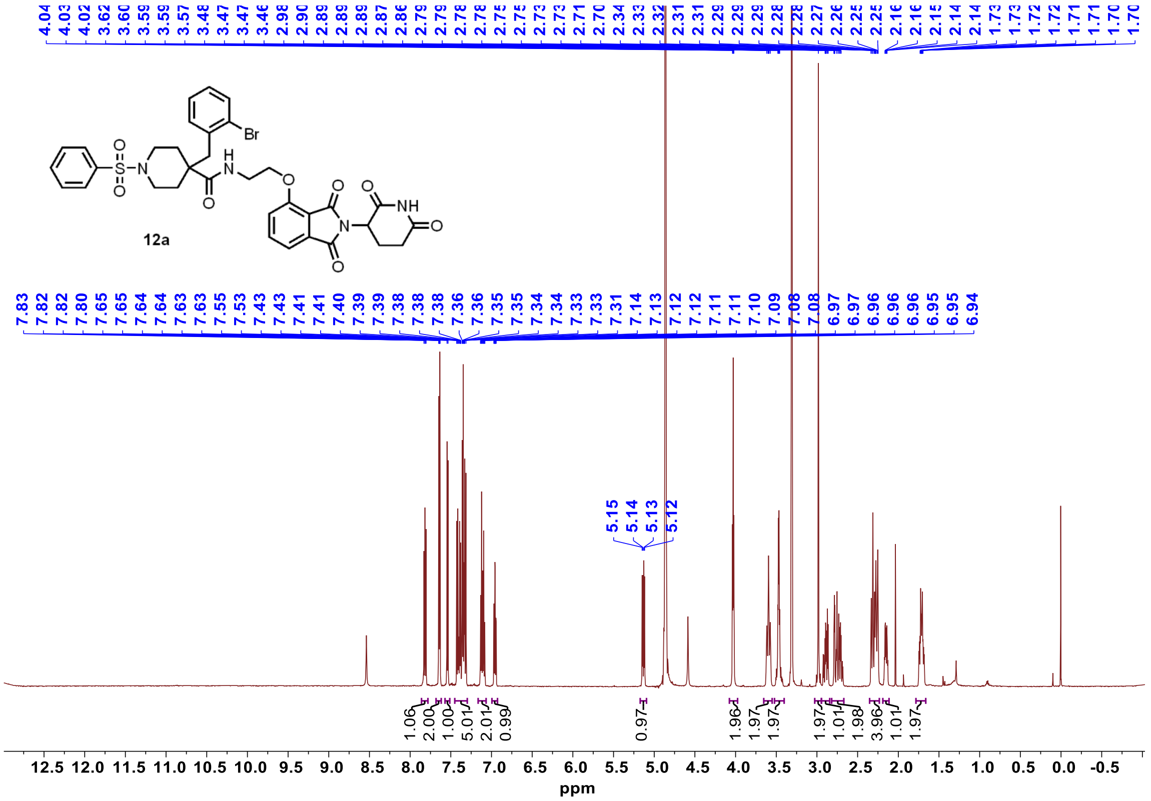


$^{13}$C NMR **12a**

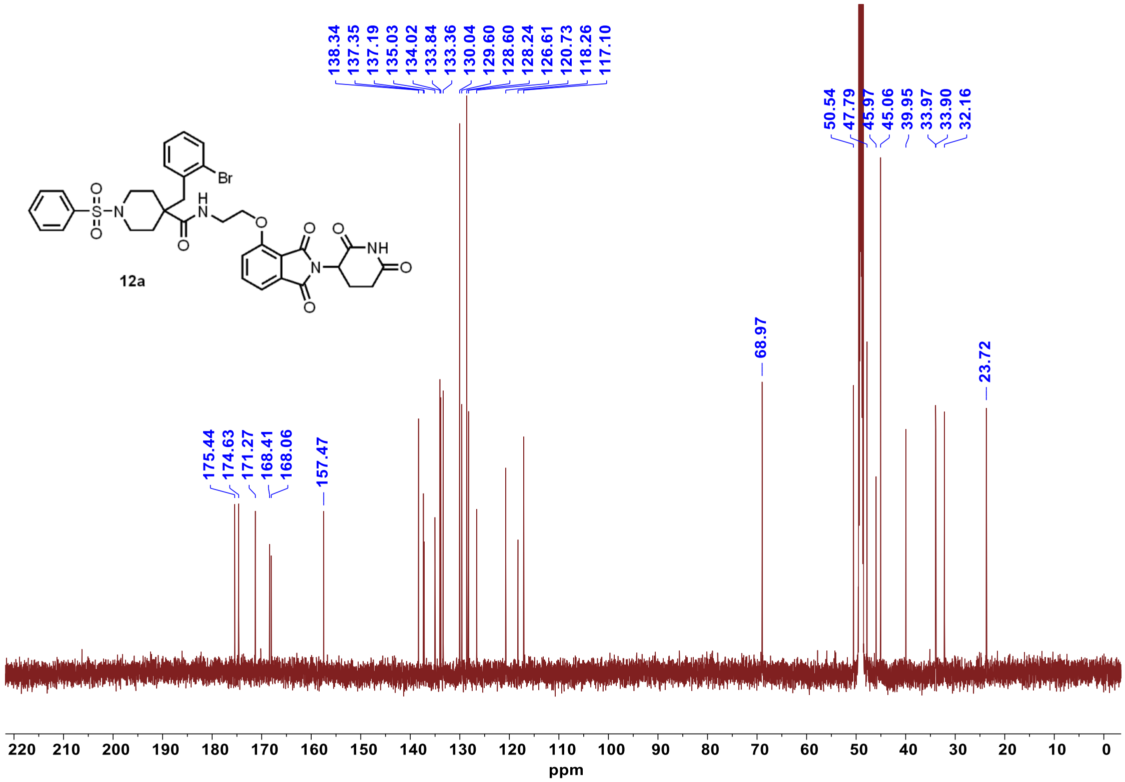

$^{1}$H NMR **12b**

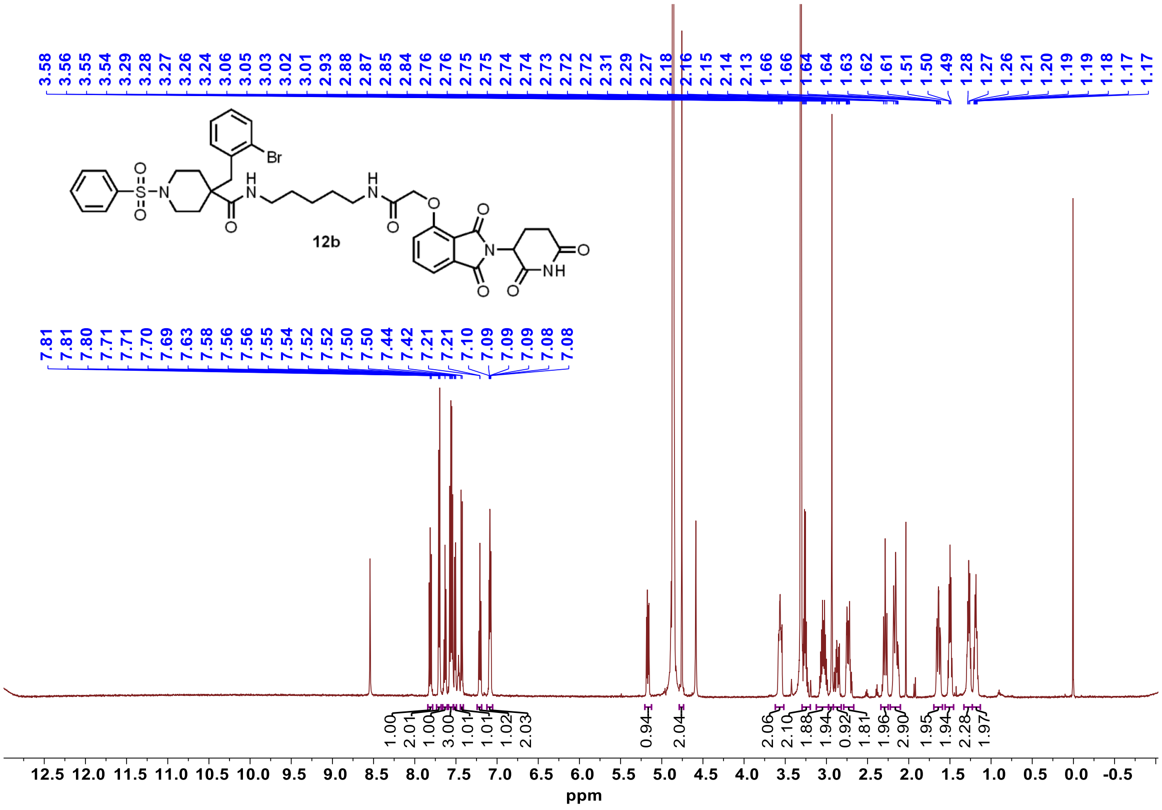


$^{13}$C NMR **12b**

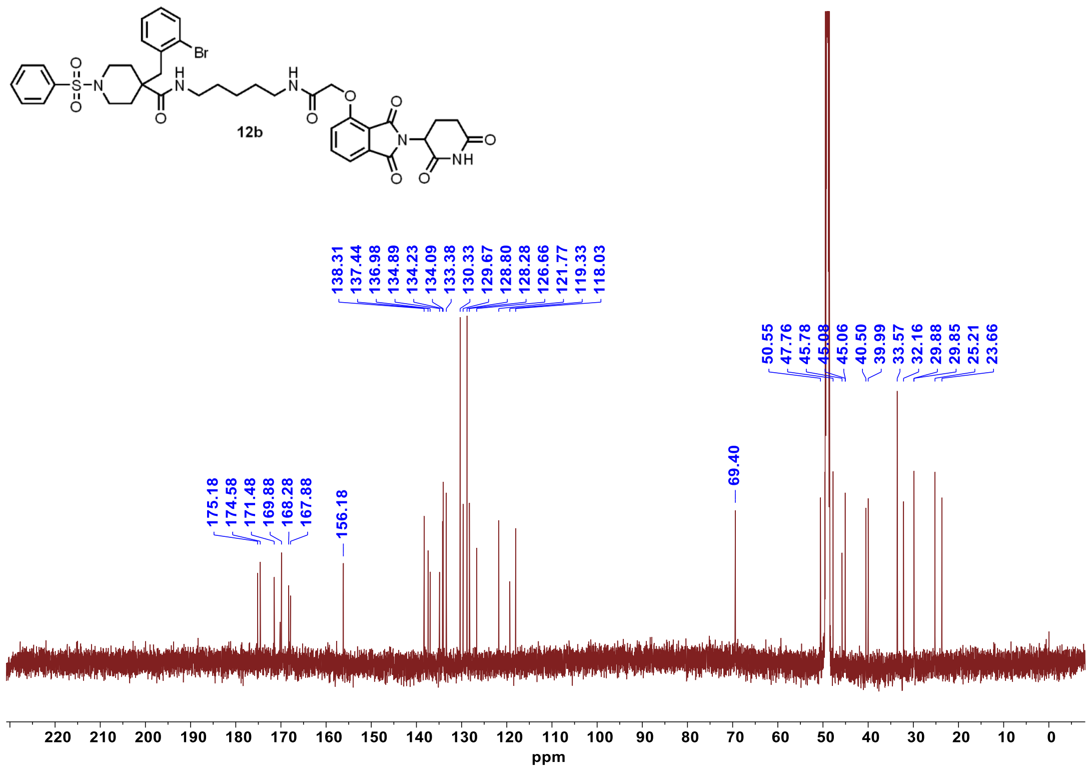

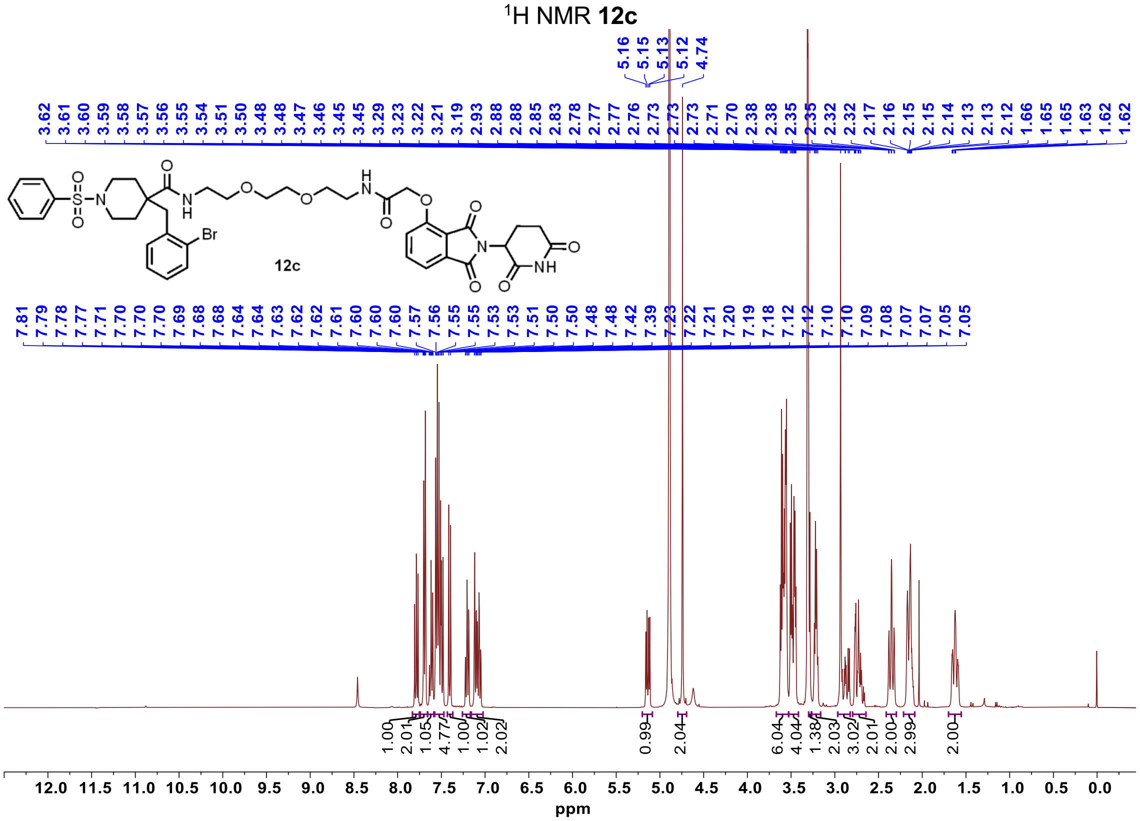

[1]H NMR 12c
3.62 3.61 3.60 3.59 3.58 3.57 3.56 3.55 3.54 3.51 3.50 3.48 3.48 3.47 3.46 3.45 3.45 3.29 3.23 3.22 3.21 3.19 2.93 2.88 2.88 2.85 2.83 2.78 2.77 2.77 2.76 2.73 2.73 2.73 2.71 2.70 2.38 2.38 2.35 2.35 2.32 2.32 2.17 2.16 2.15 2.15 2.14 2.13 2.13 2.12 1.66 1.65 1.63 1.62 1.62
5.16 5.15 5.13 5.12 4.74
7.81 7.79 7.78 7.77 7.71 7.70 7.70 7.69 7.68 7.68 7.64 7.64 7.63 7.62 7.62 7.61 7.60 7.60 7.60 7.57 7.56 7.55 7.55 7.53 7.53 7.51 7.50 7.50 7.48 7.48 7.42 7.39 7.23 7.22 7.21 7.20 7.19 7.18 7.12 7.12 7.10 7.10 7.09 7.08 7.07 7.07 7.05 7.05
12c
Br
1.00 2.01 1.05 4.77 1.00 1.02 2.02 0.99 2.04 6.04 4.04 1.38 2.03 3.02 2.01 2.00 2.99 2.00
12.0 11.5 11.0 10.5 10.0 9.5 9.0 8.5 8.0 7.5 7.0 6.5 6.0 5.5 5.0 4.5 4.0 3.5 3.0 2.5 2.0 1.5 1.0 0.5 0.0
ppm


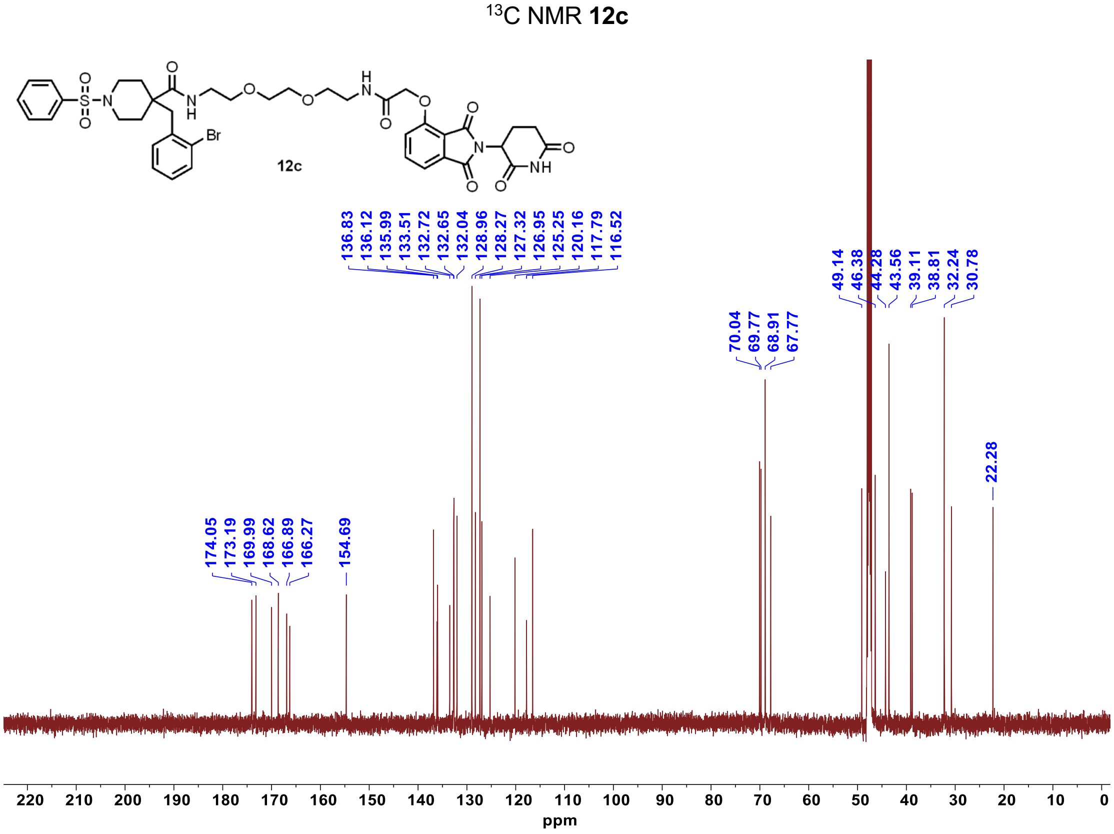

[13]C NMR 12c
12c
Br
136.83 136.12 135.99 133.51 132.72 132.65 132.04 128.96 128.27 127.32 126.95 125.25 120.16 117.79 116.52
49.14 46.38 44.28 43.56 39.11 38.81 32.24 30.78
70.04 69.77 68.91 67.77
174.05 173.19 169.99 168.62 166.89 166.27
154.69
22.28
220 210 200 190 180 170 160 150 140 130 120 110 100 90 80 70 60 50 40 30 20 10 0
ppm

$^{1}H$ NMR **12d**

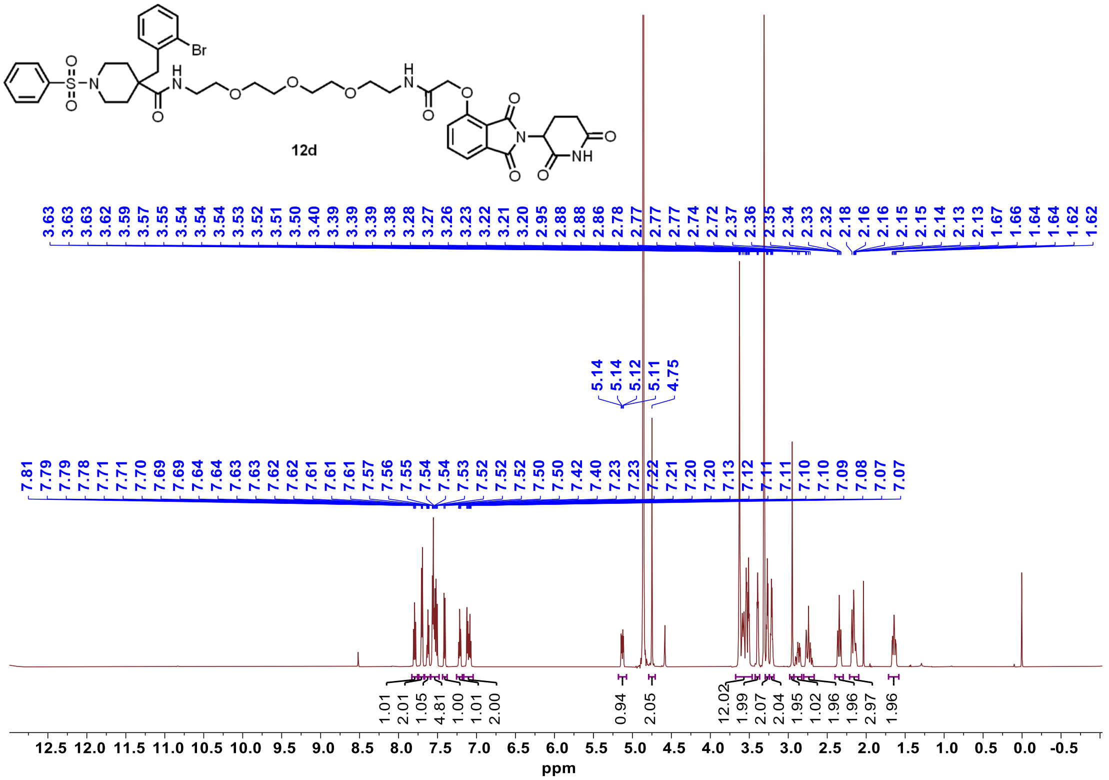


$^{13}C$ NMR **12d**

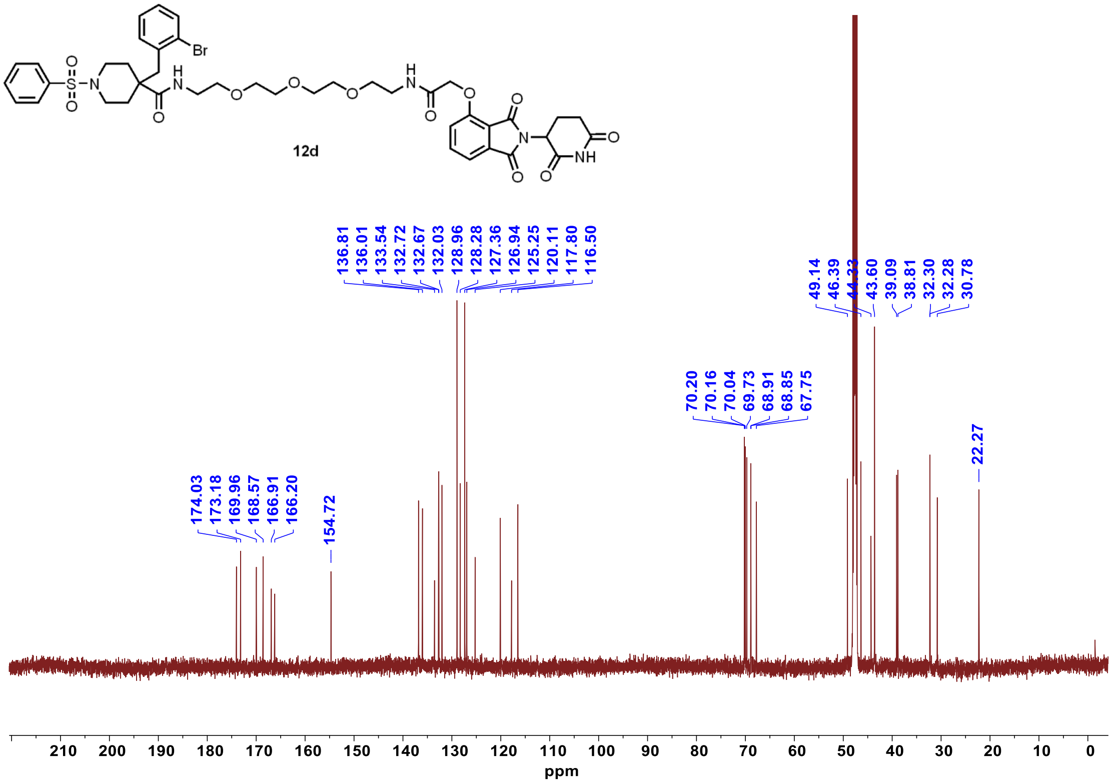

$^{1}$H NMR **13a**

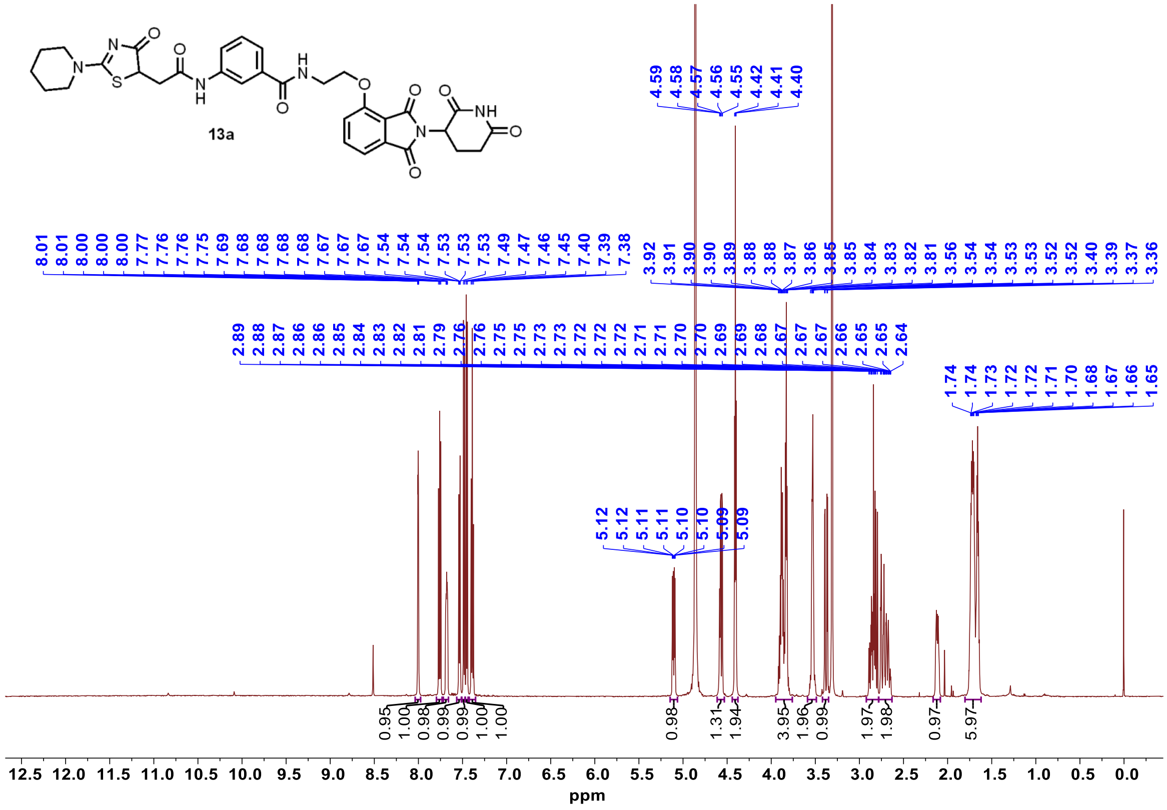

$^{13}$C NMR **13a**

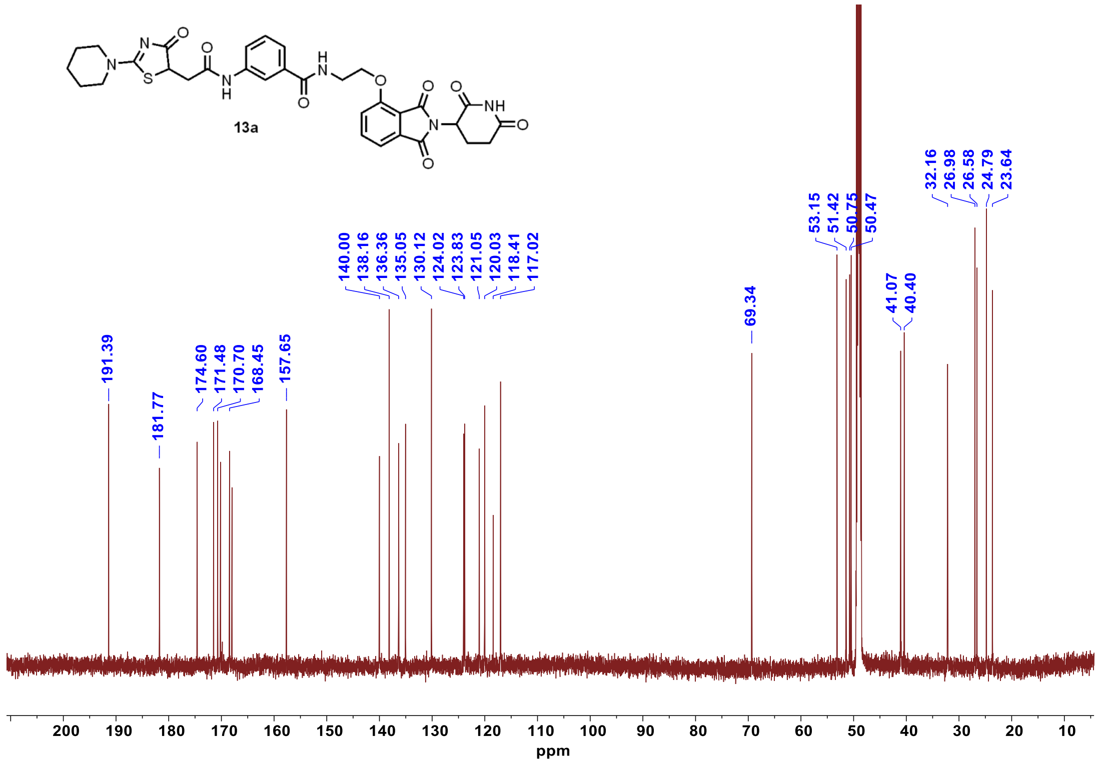

[1]H NMR **13b**

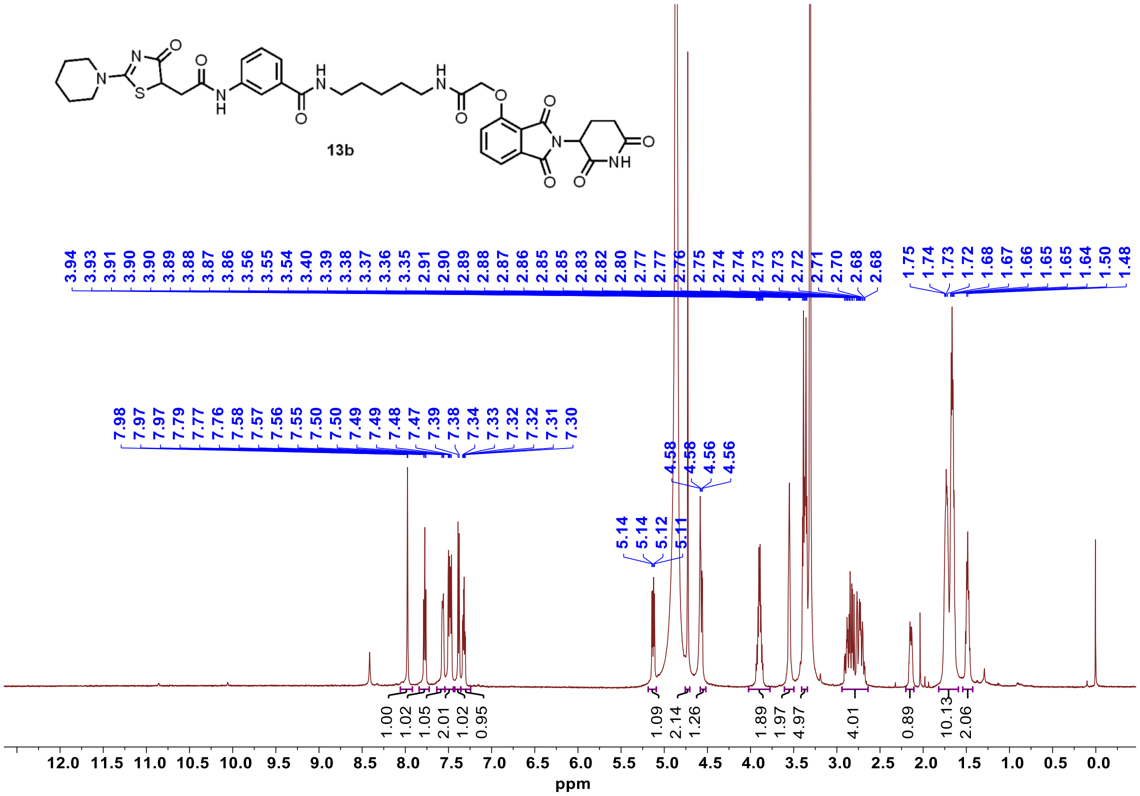


[13]C NMR **13b**

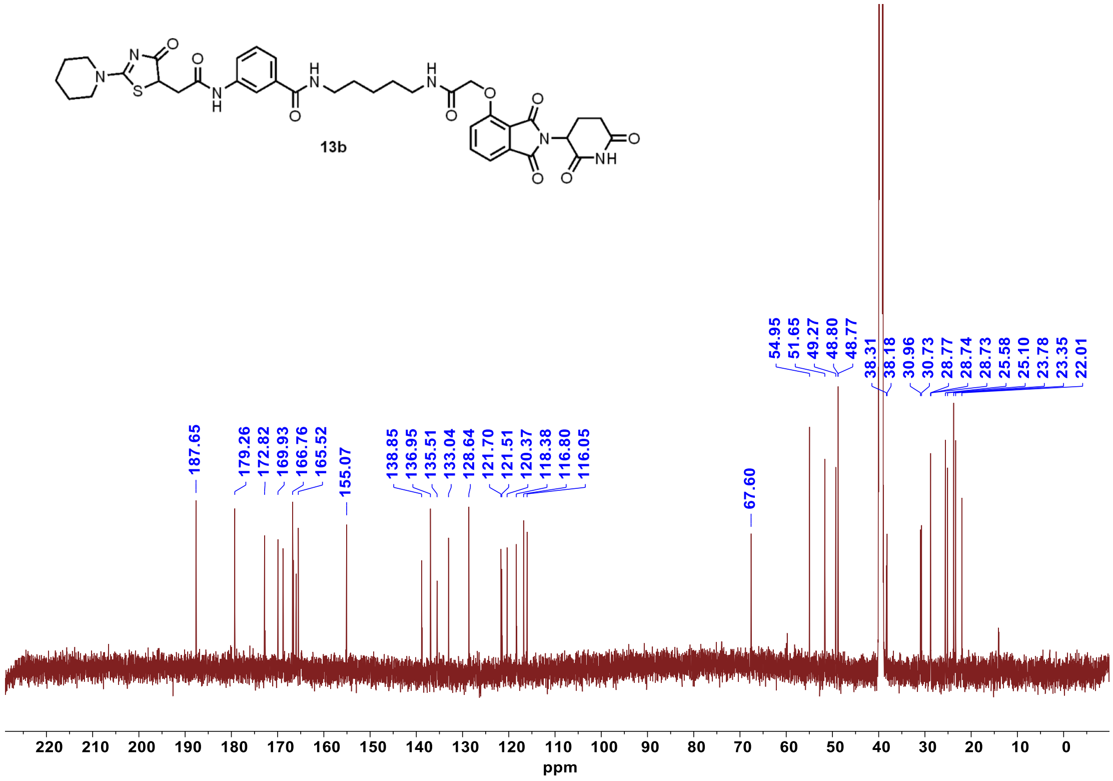

$^{1}H$ NMR **13c**

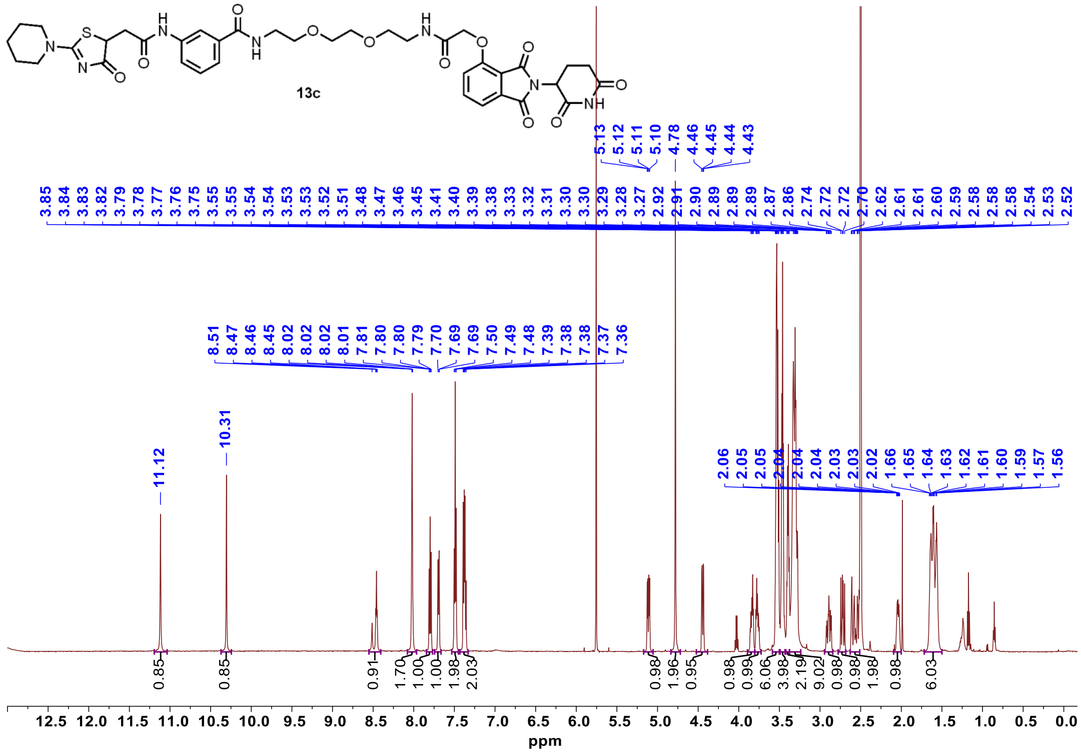


$^{13}C$ NMR **13c**

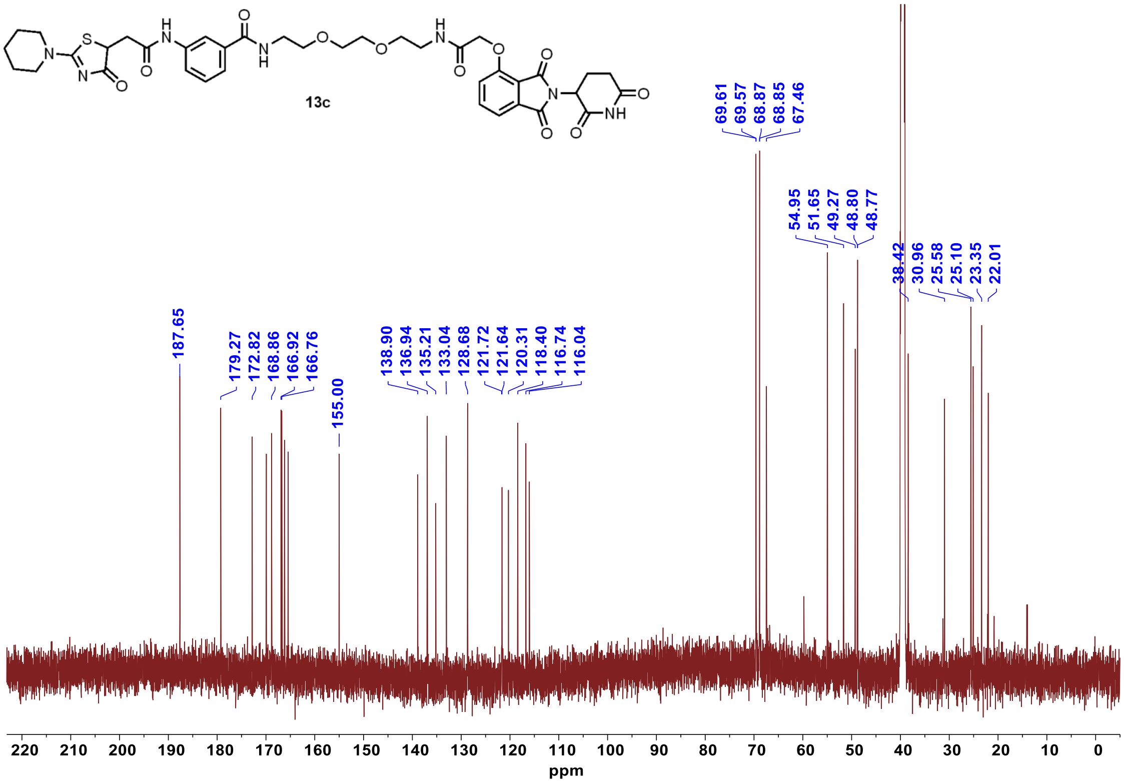

$^1$H NMR **13d**

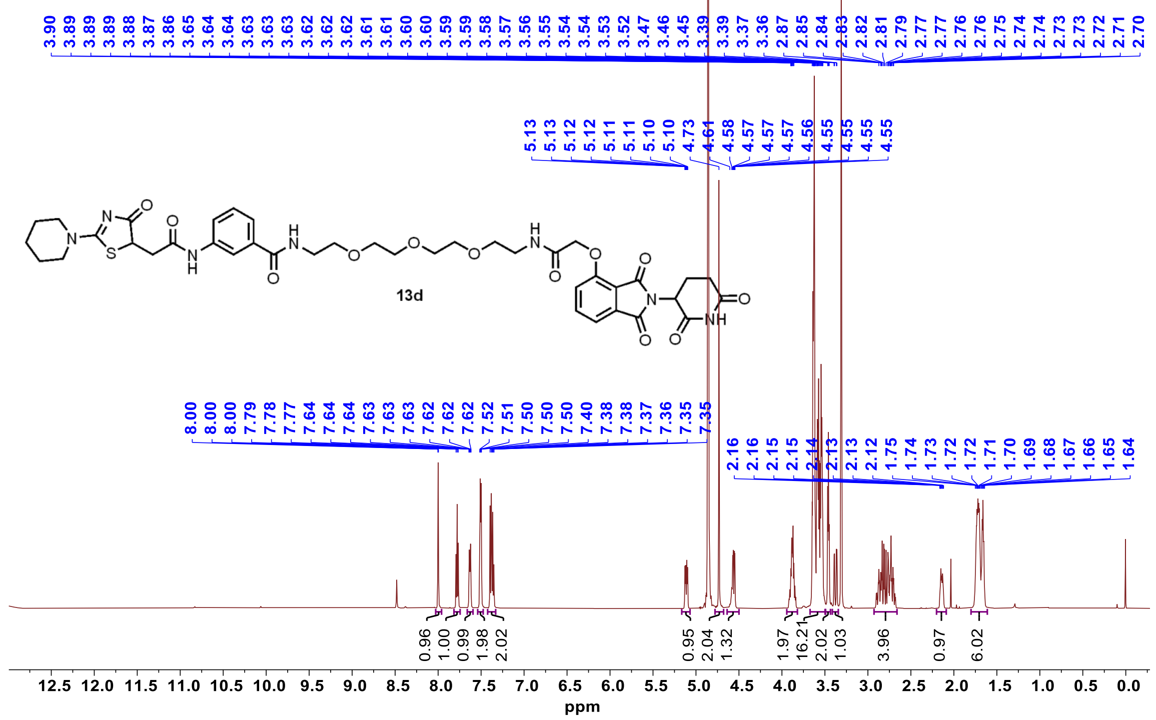


$^{13}$C NMR **13d**

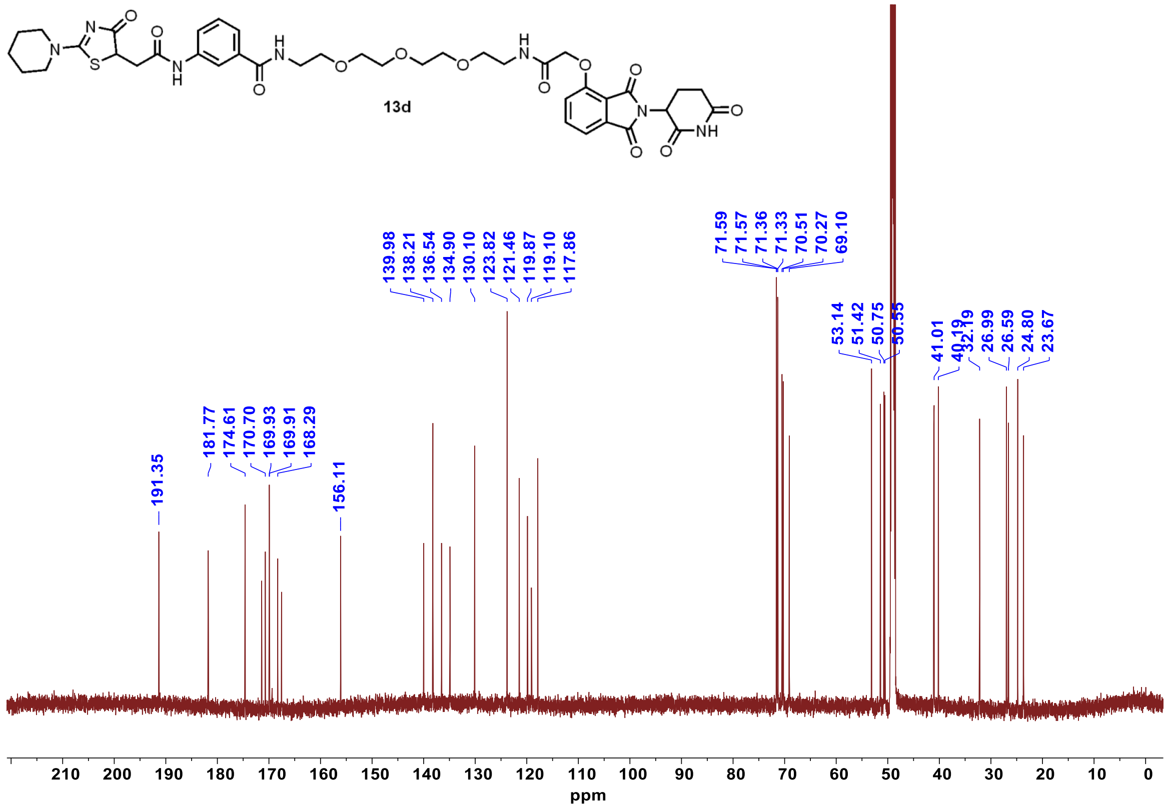

$^{1}$H NMR **14a**

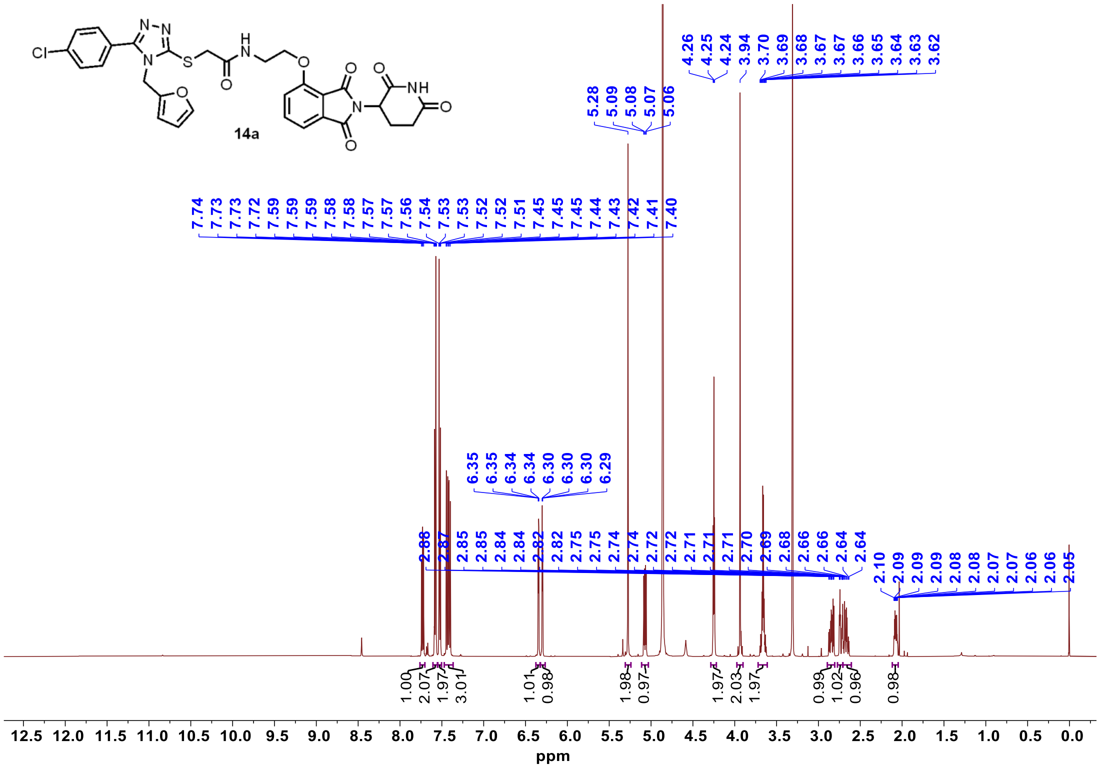

$^{13}$C NMR **14a**

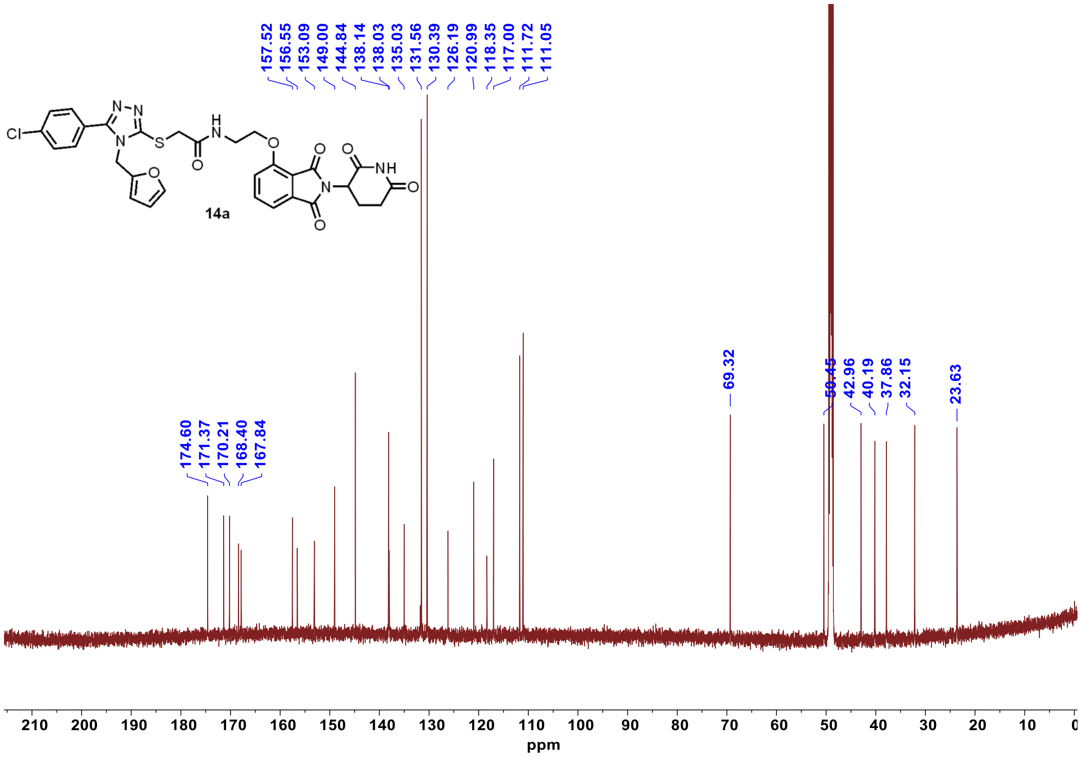

$^{1}$H NMR **14b**

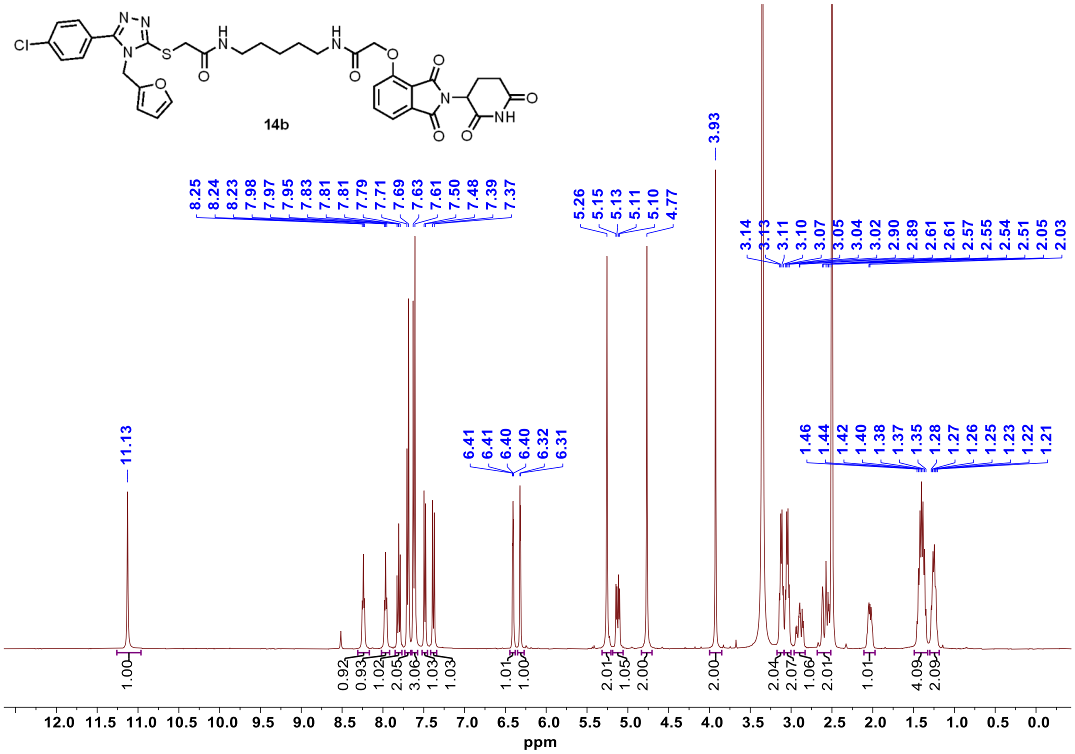


$^{13}$C NMR **14b**

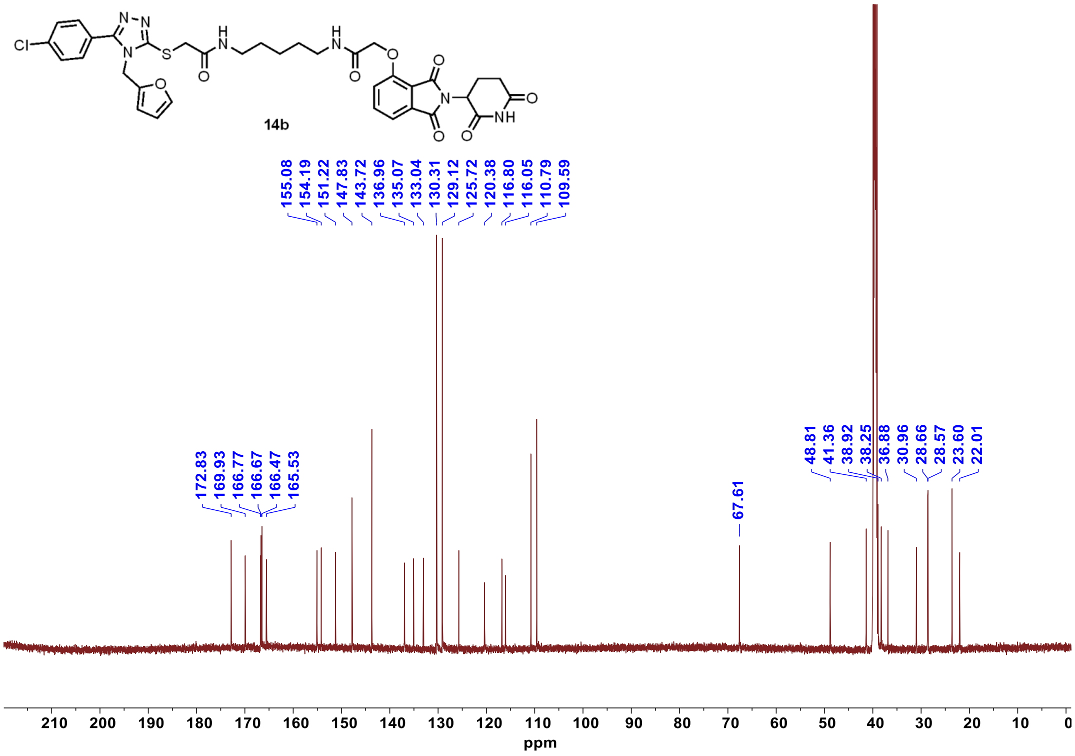

[1]H NMR **14c**

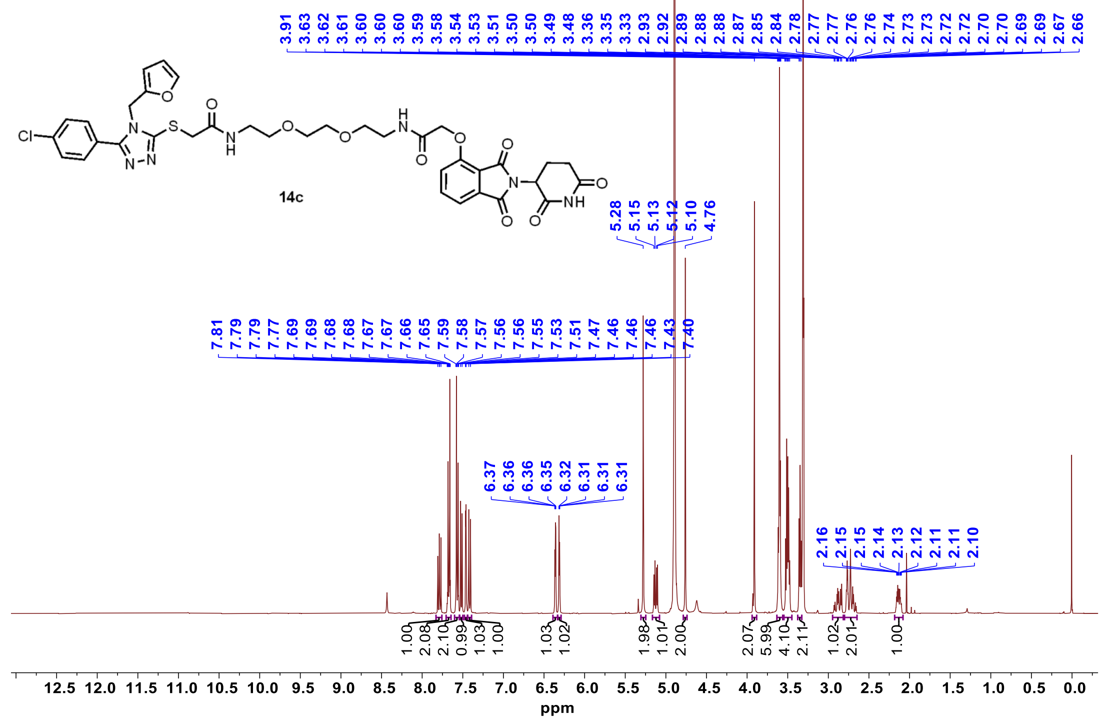


[13]C NMR **14c**

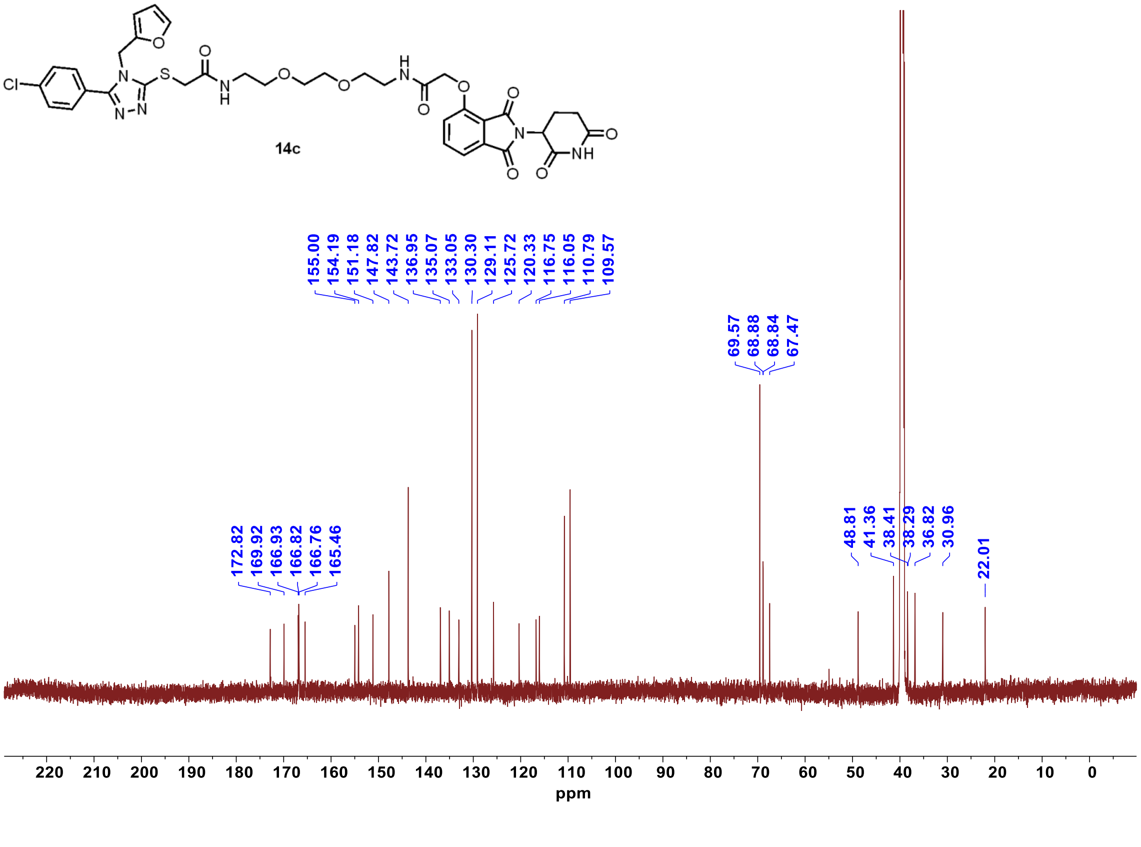

$^1$H NMR **14d**

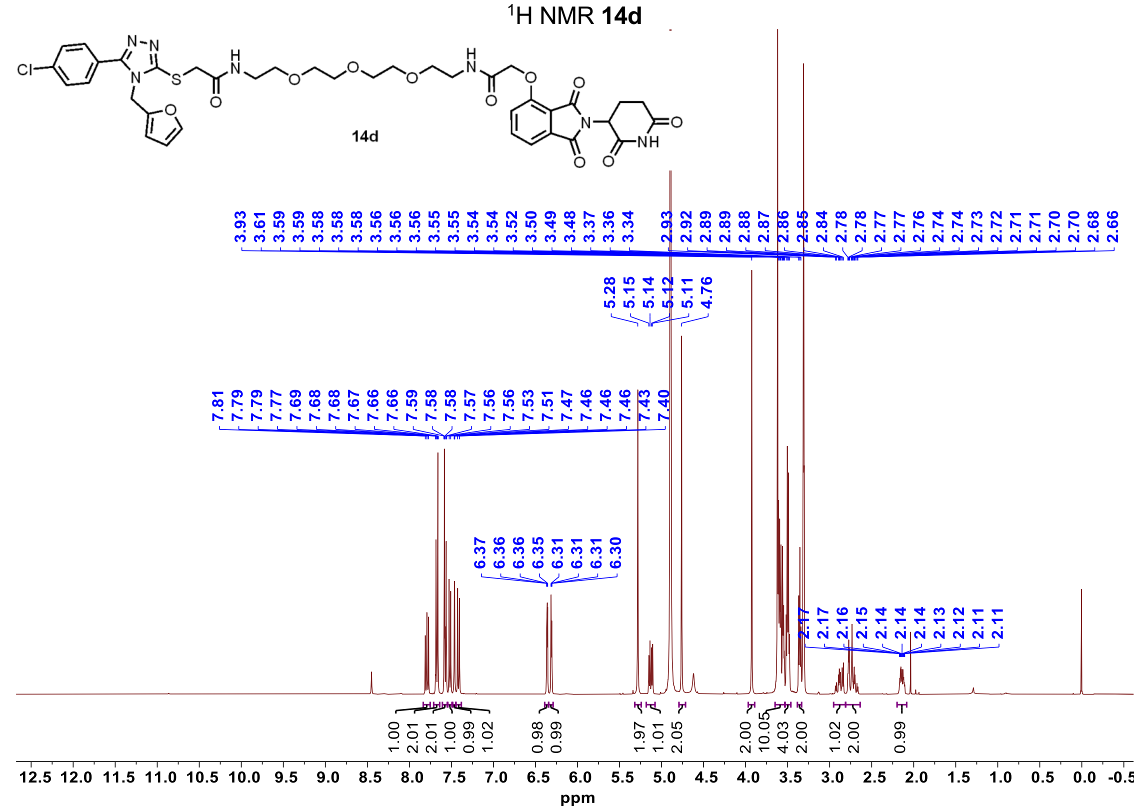


$^{13}$C NMR **14d**

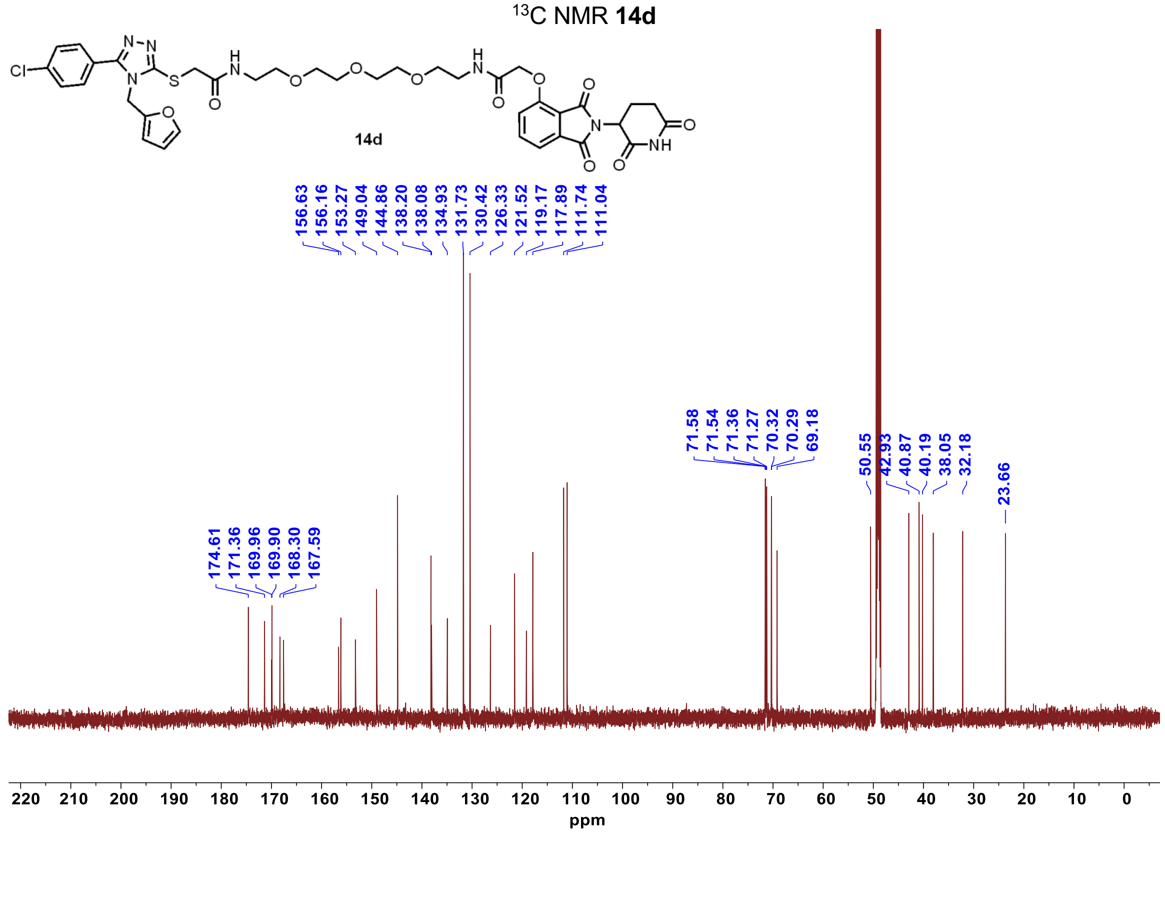

$^{1}$H NMR **15a**

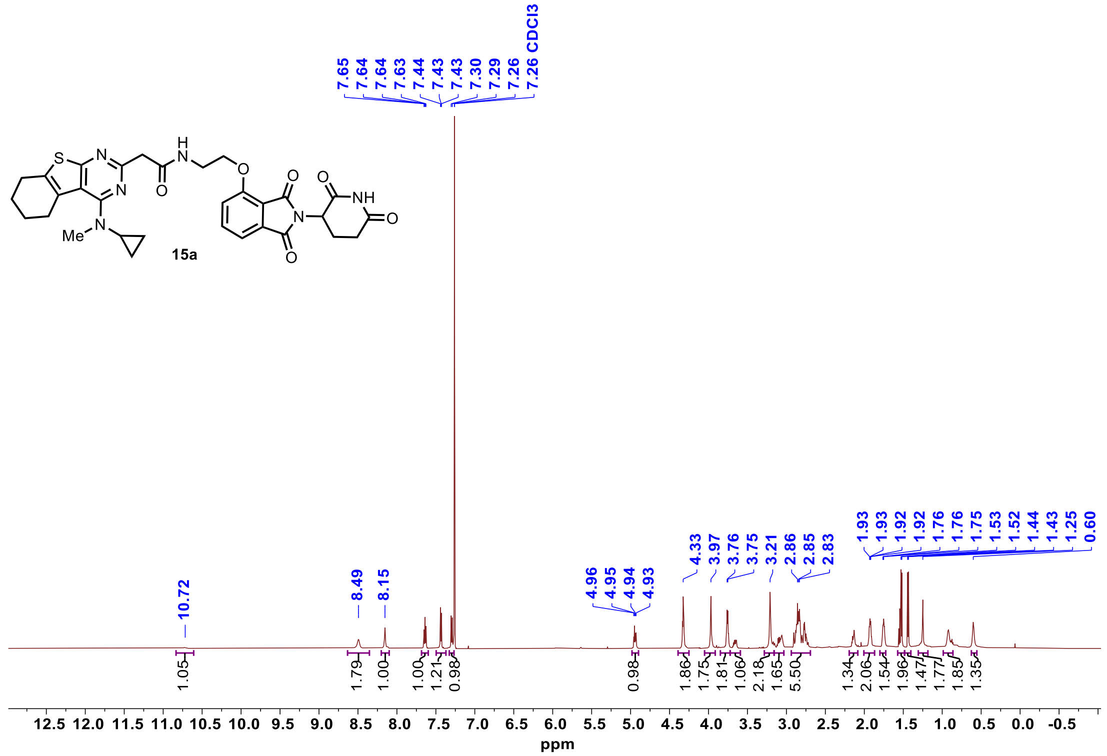


$^{13}$C NMR **15a**

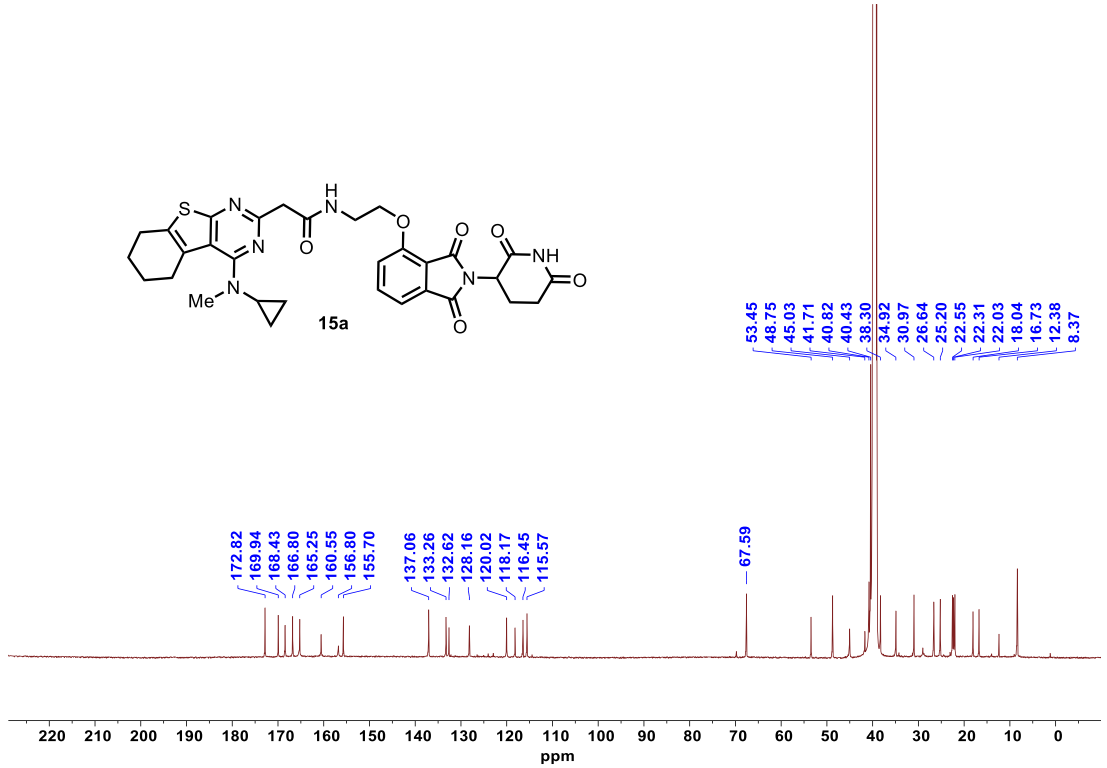

[1]H NMR **15b**

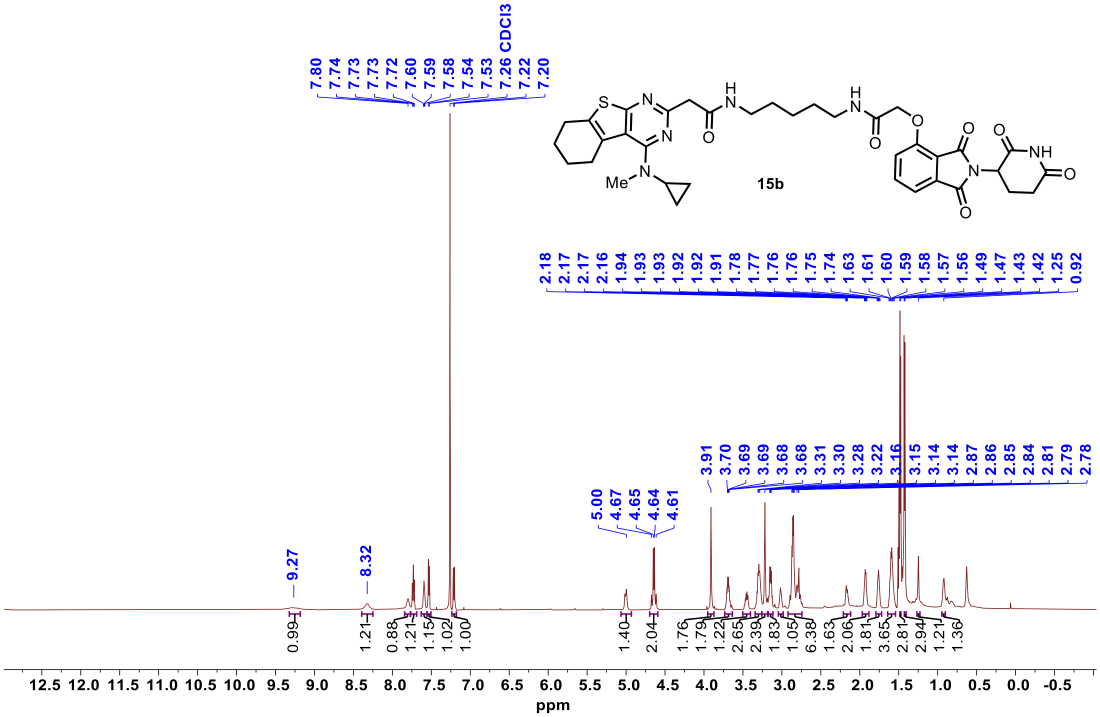


[13]C NMR **15b**

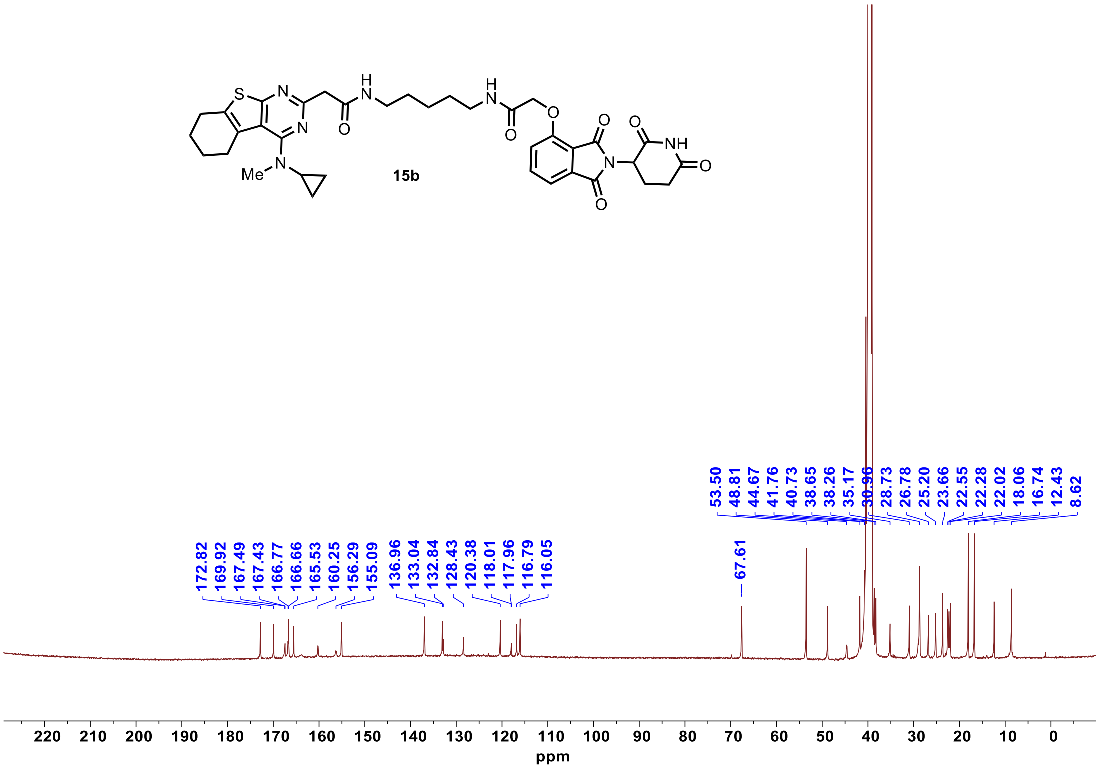

#### $^{1}$H NMR **15c**

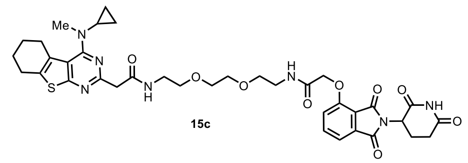


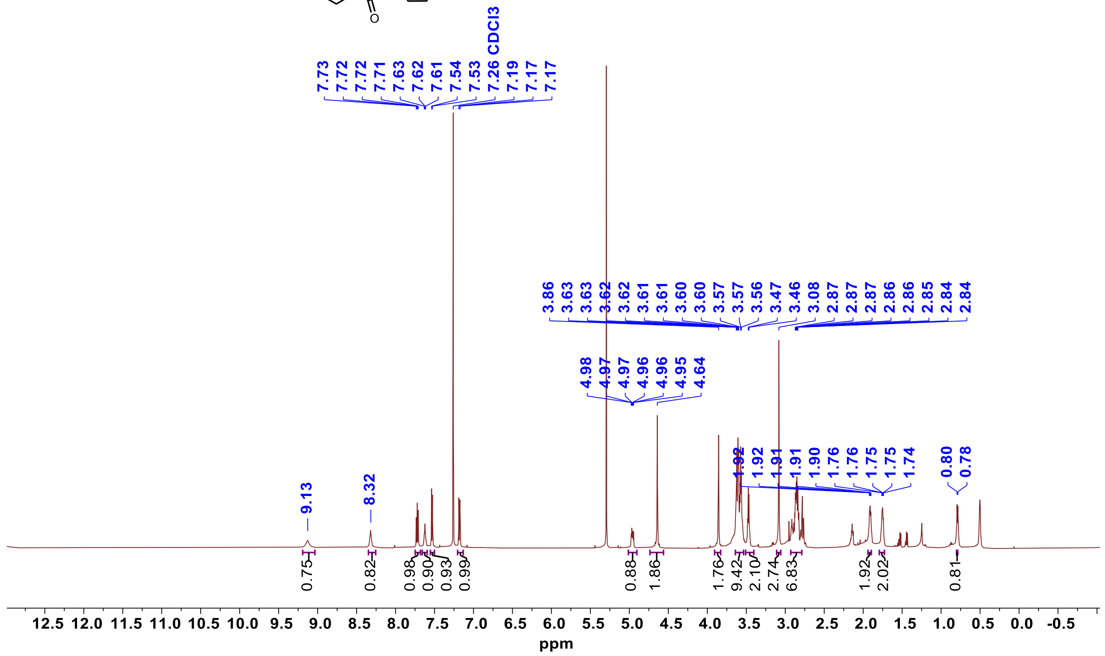


#### $^{13}$C NMR **15c**

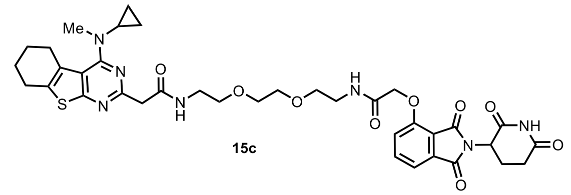


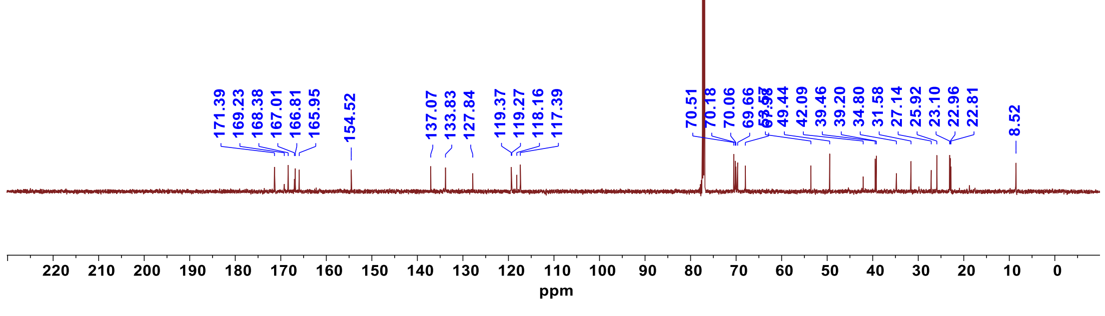

[1]H NMR **15d**

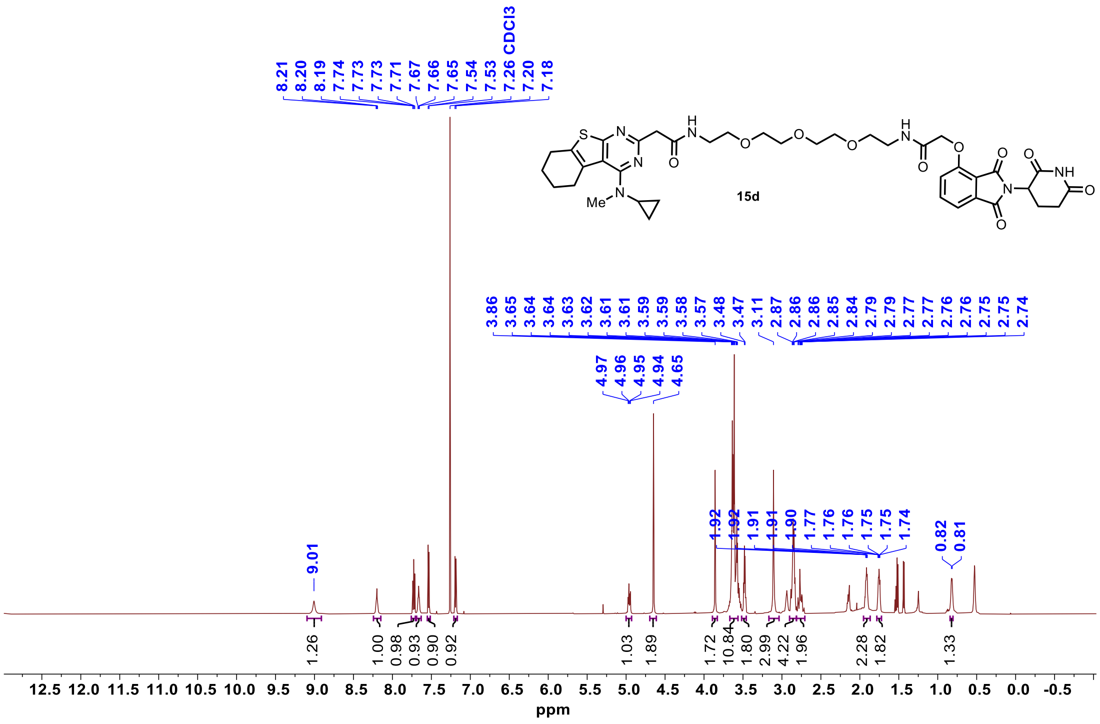


[13]C NMR **15d**

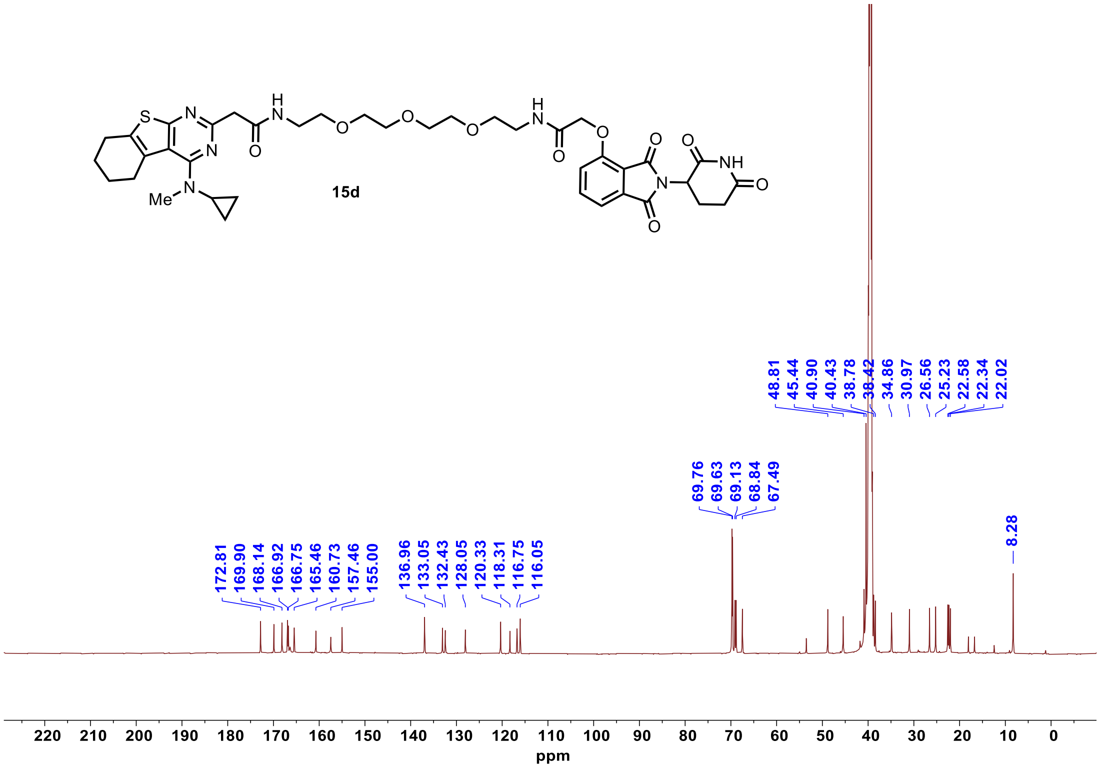

$^{1}$H NMR **16a**

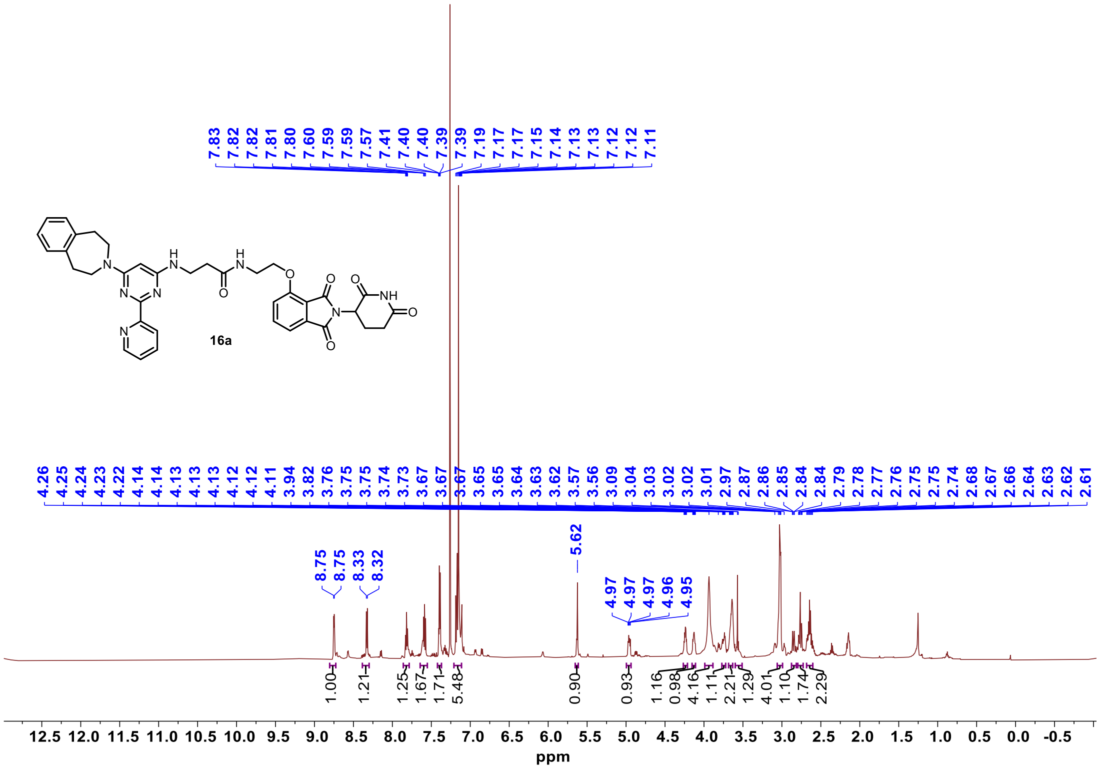

$^{13}$C NMR **16a**

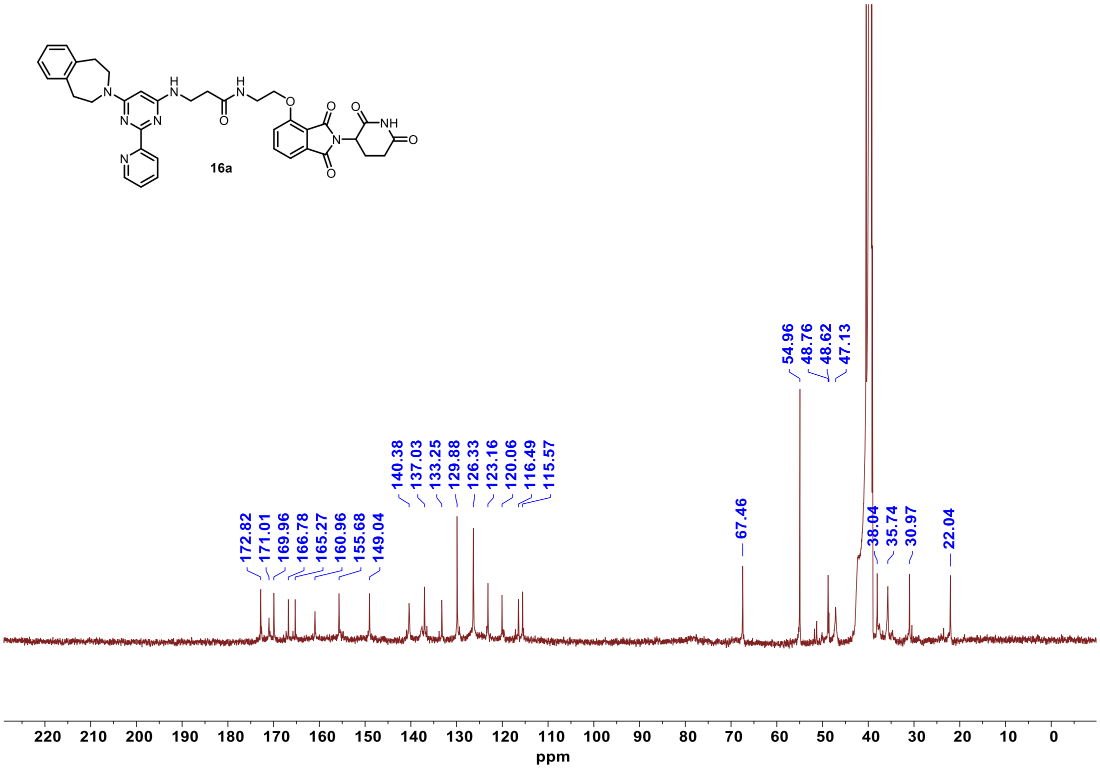

<sup></sup>

$^{1}$H NMR **16b**

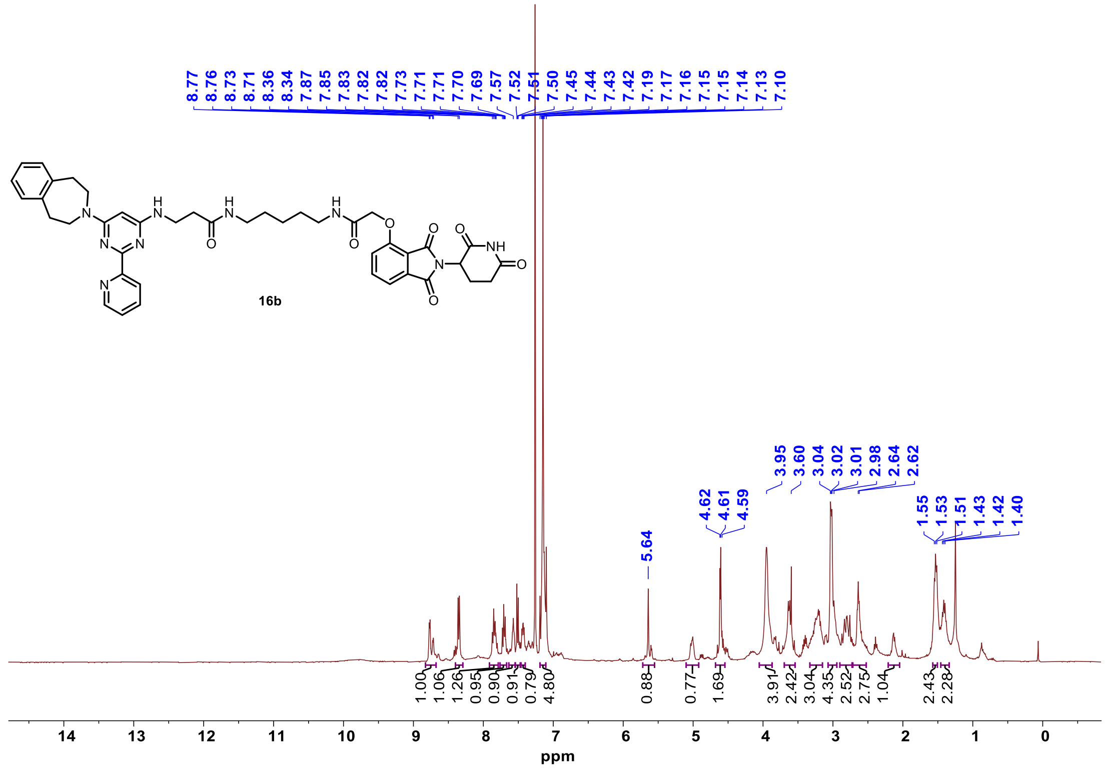

$^{13}$C NMR **16b**

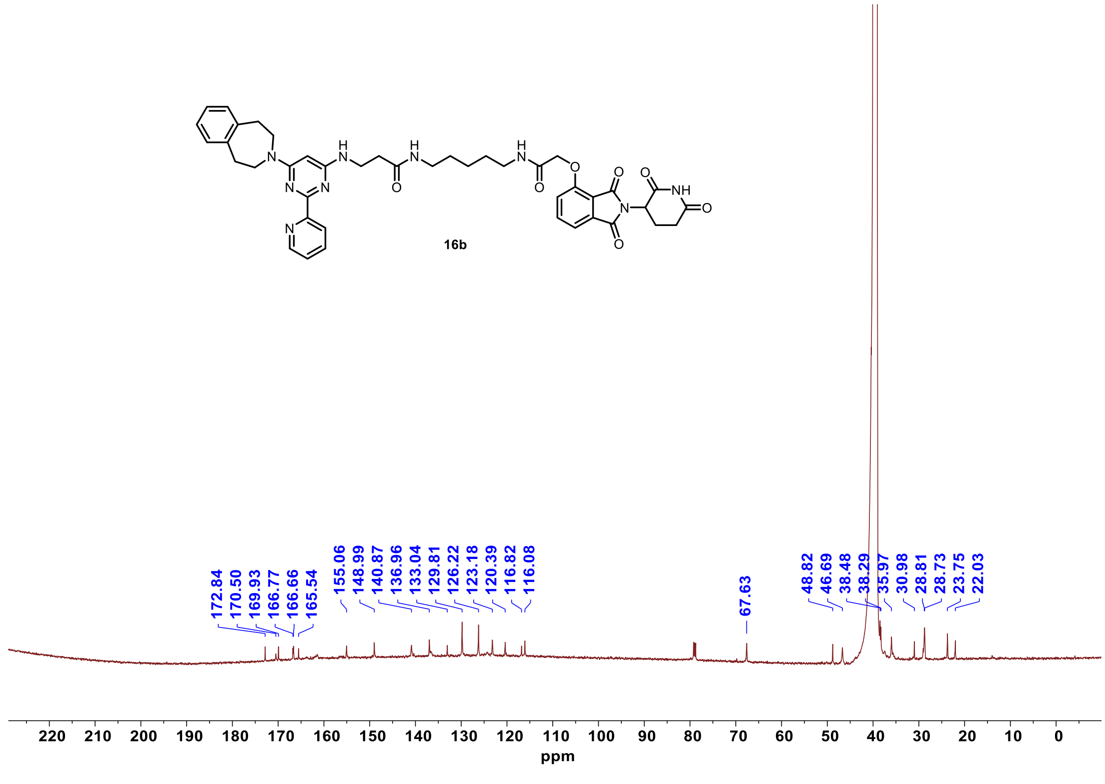

$^{1}$H NMR **16c**

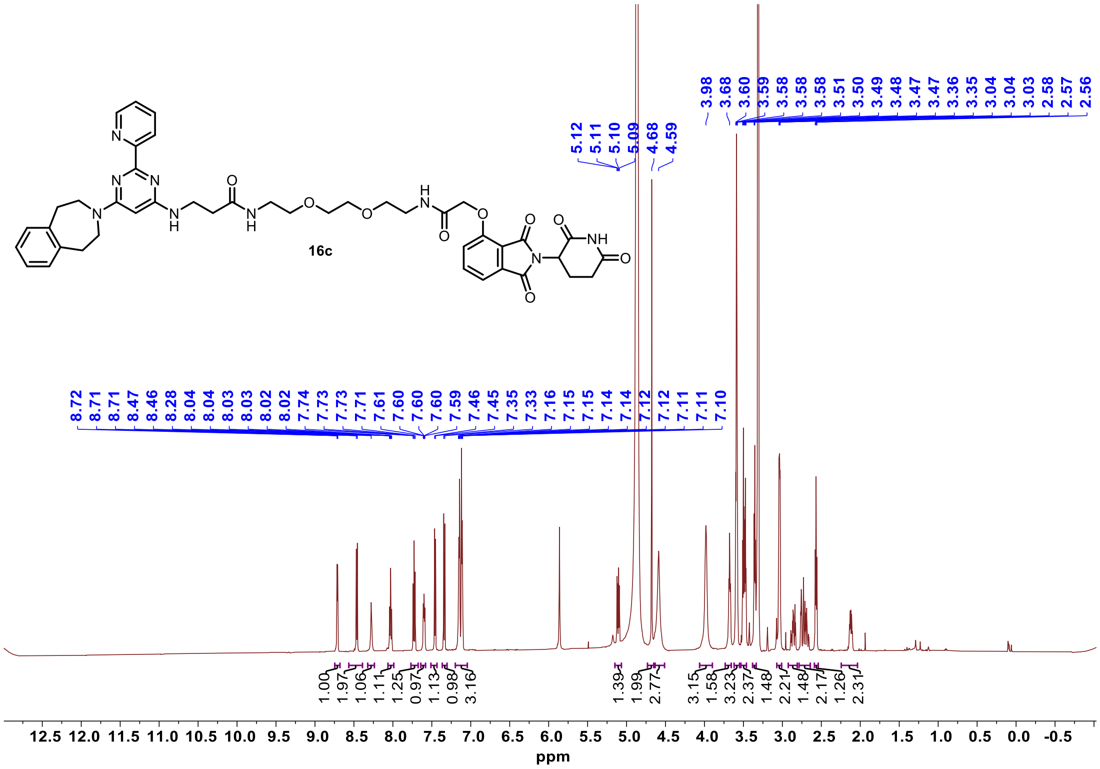

$^{13}$C NMR **16c**

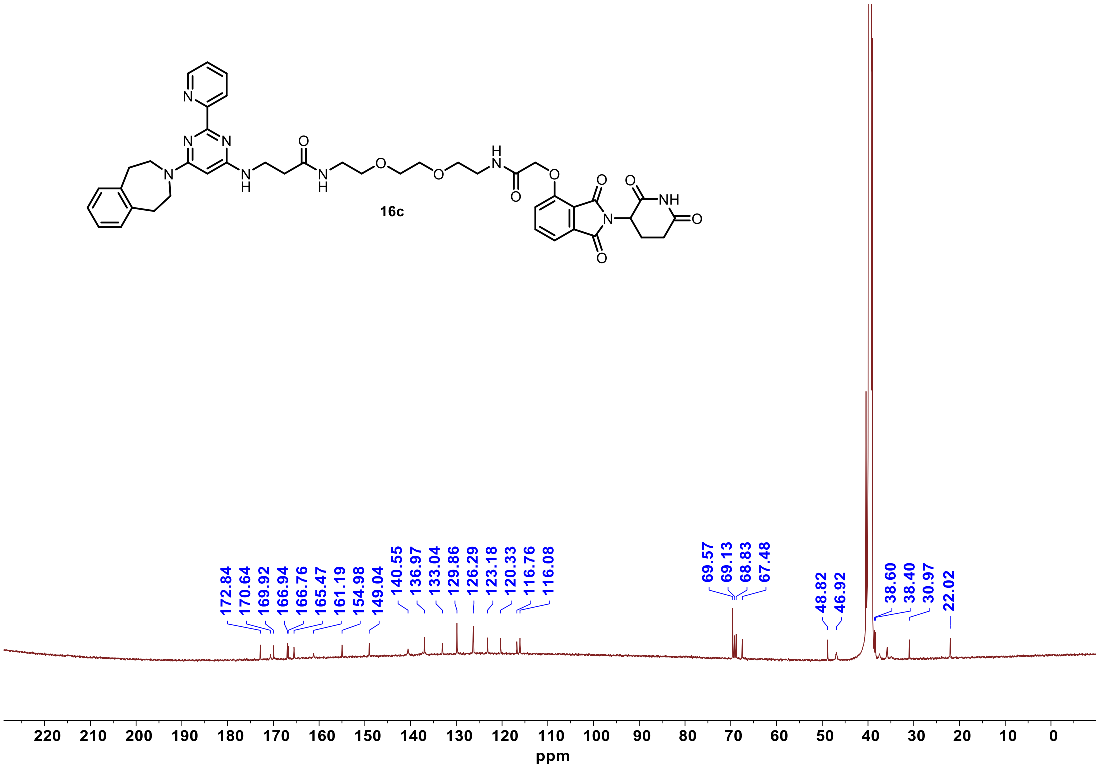

$^{1}$H NMR **16d**

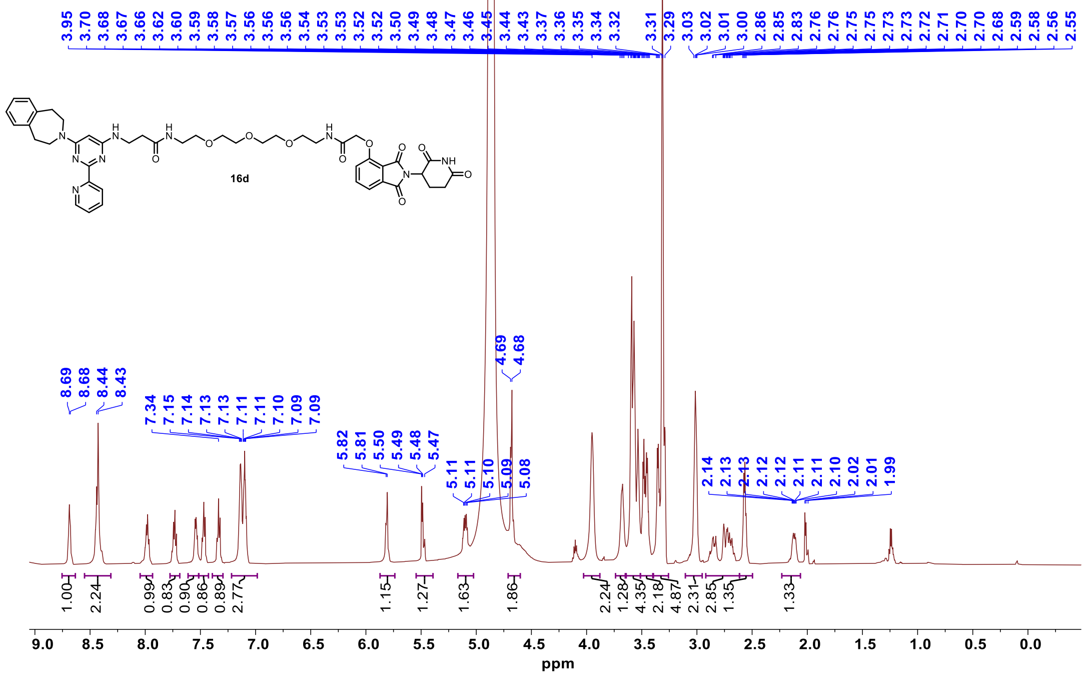


$^{13}$C NMR **16d**

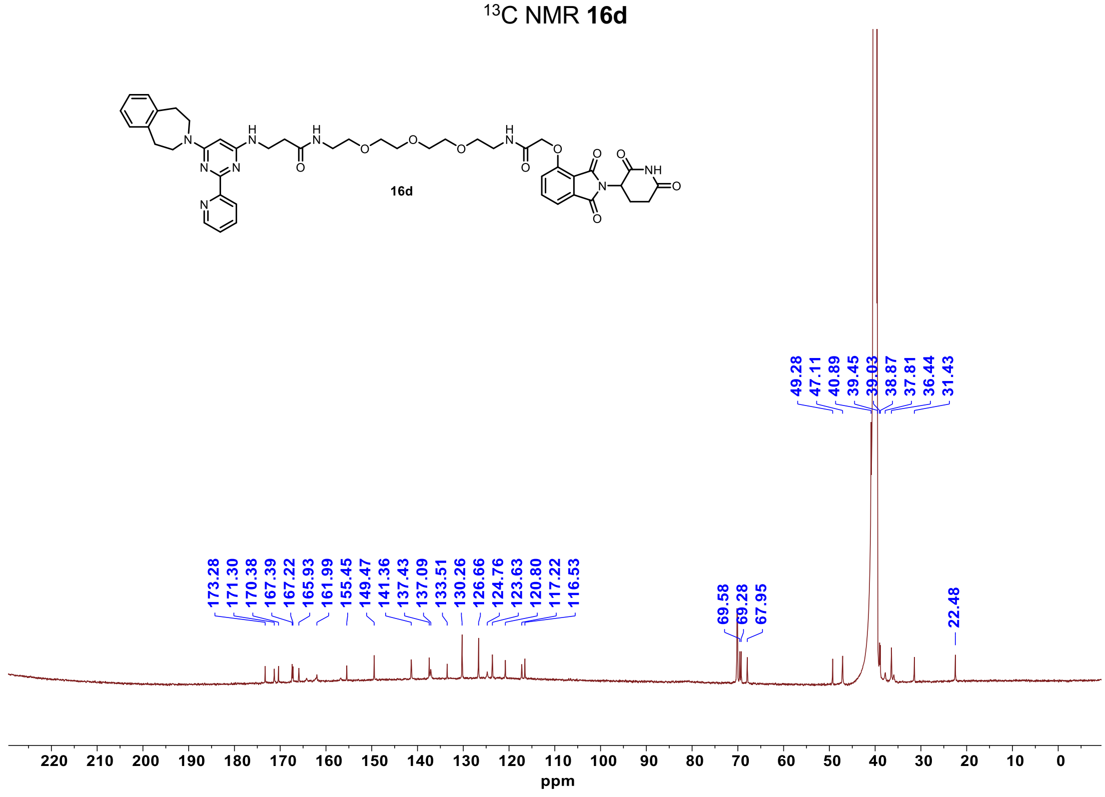

[1]H NMR **17a**

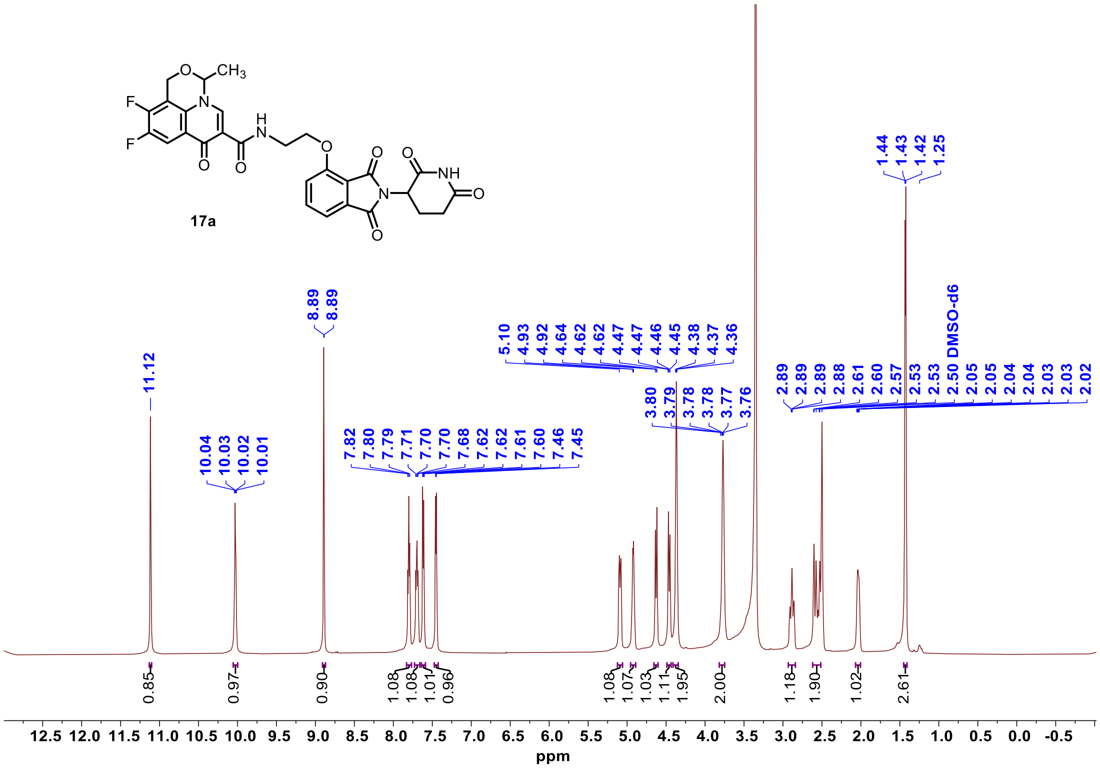


[13]C NMR **17a**

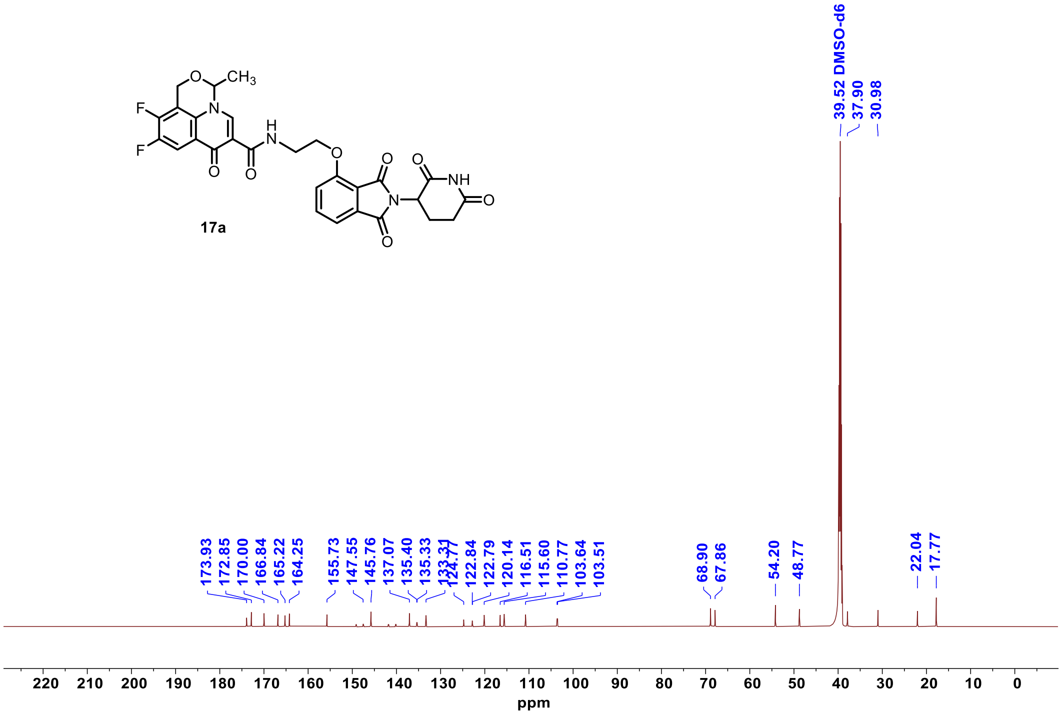

$^{1}H$ NMR **17b**

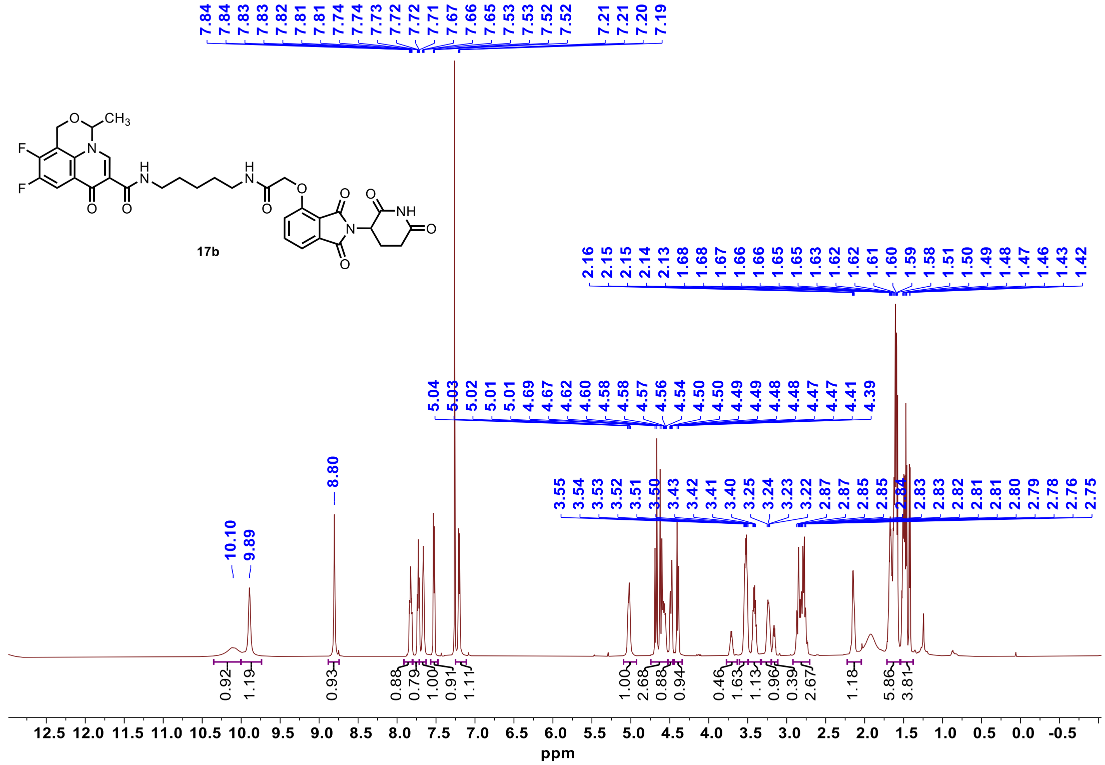


$^{13}C$ NMR **17b**

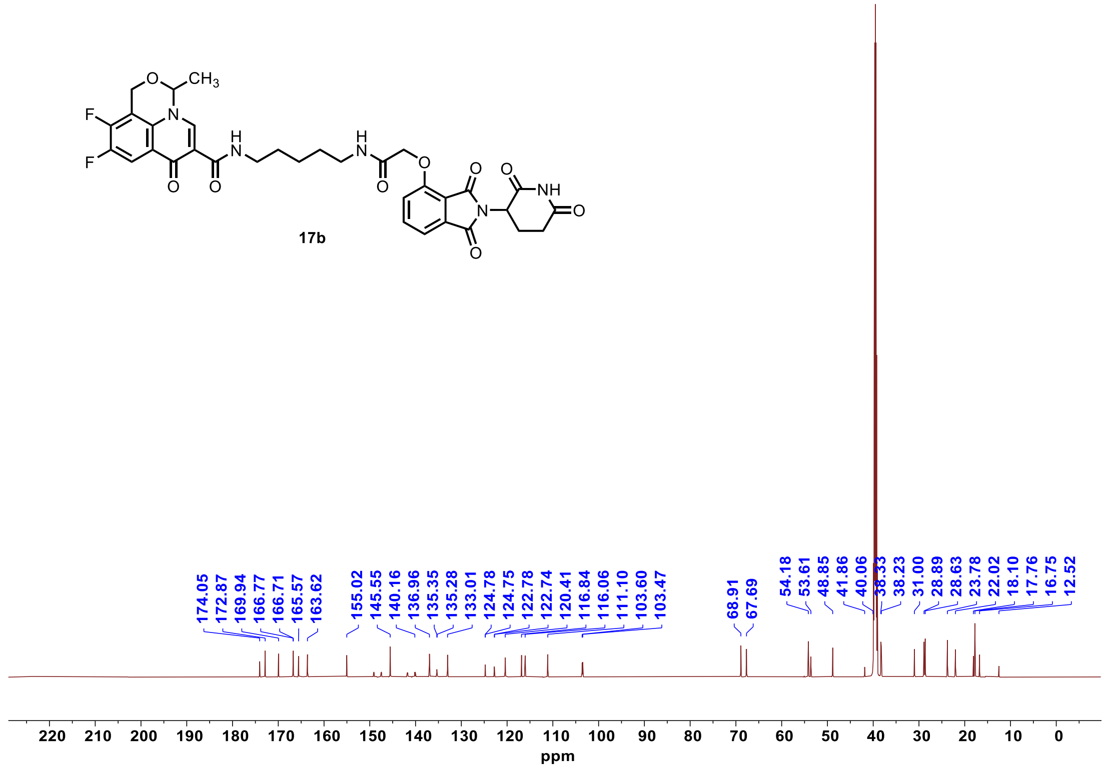

$^1H$ NMR **17c**

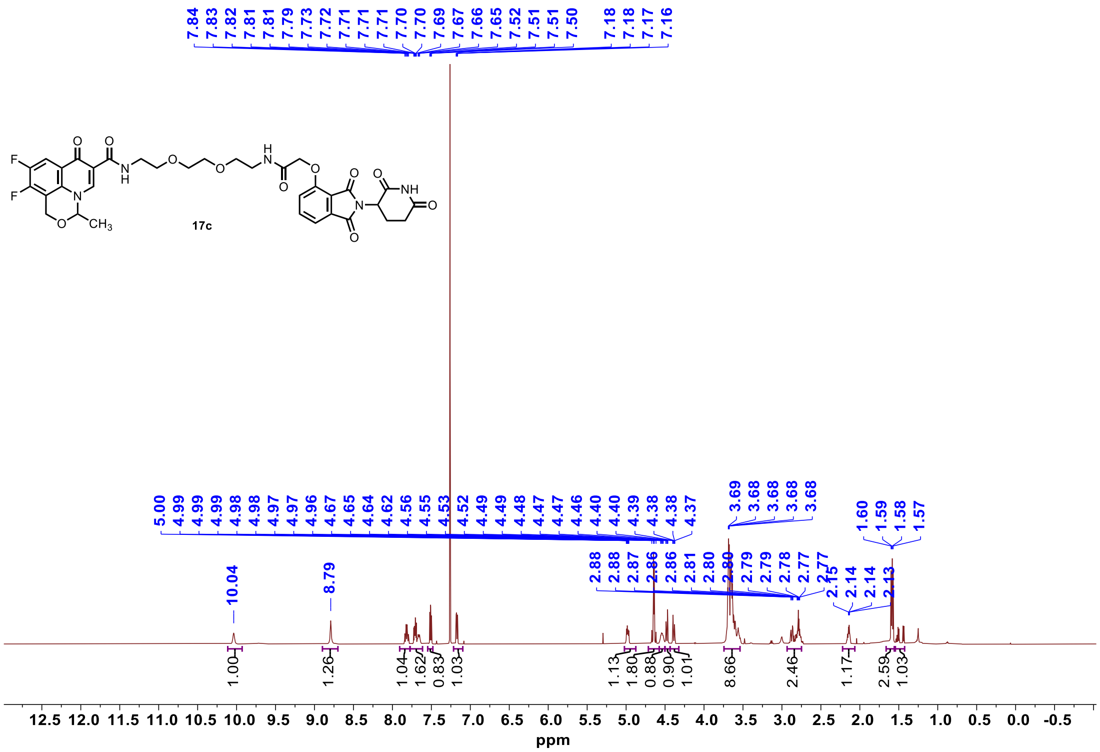

$^{13}C$ NMR **17c**

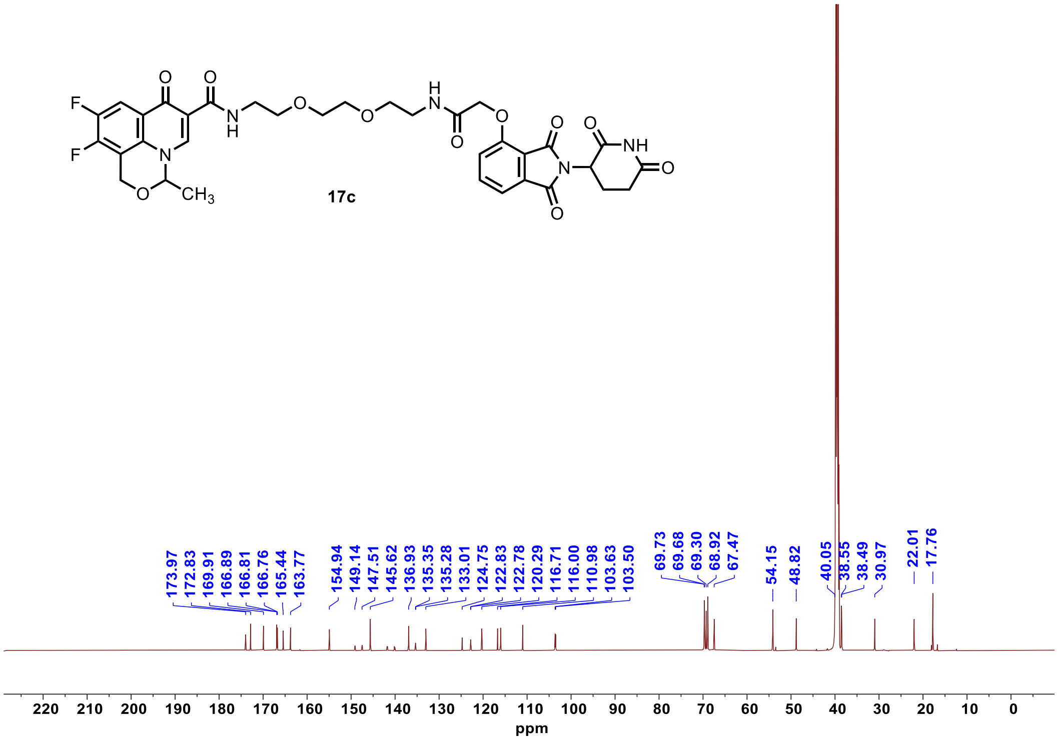

[1]H NMR **17d**

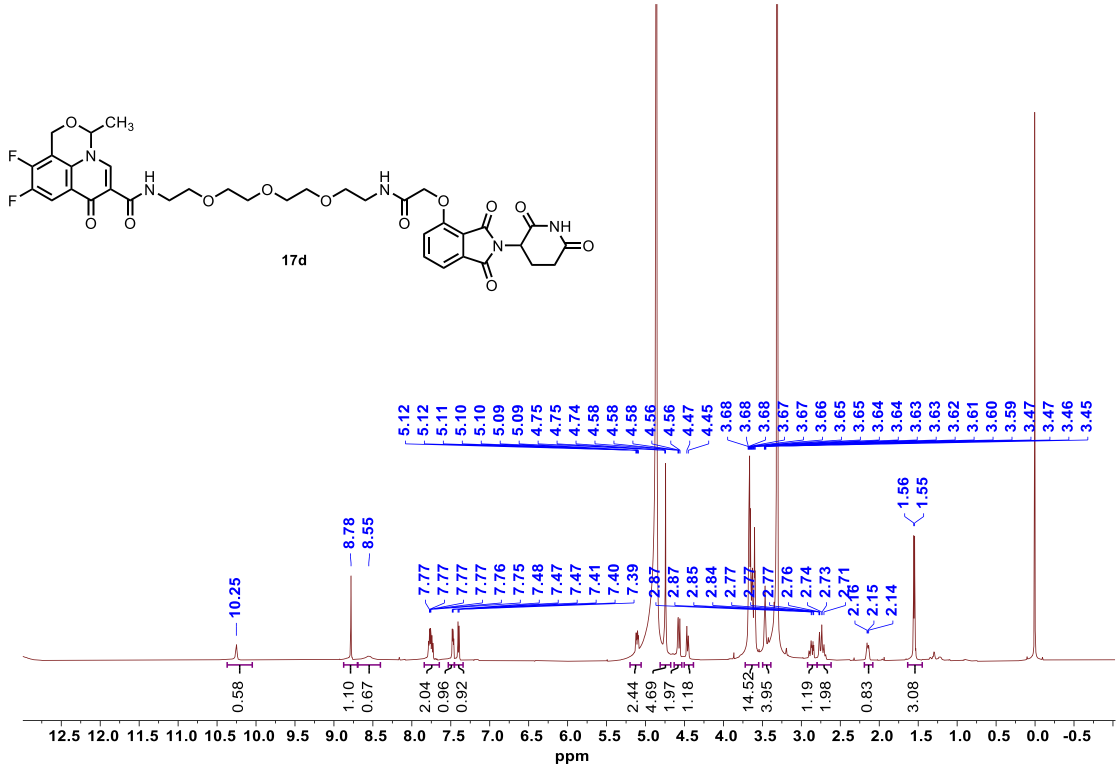

[13]C NMR **17d**

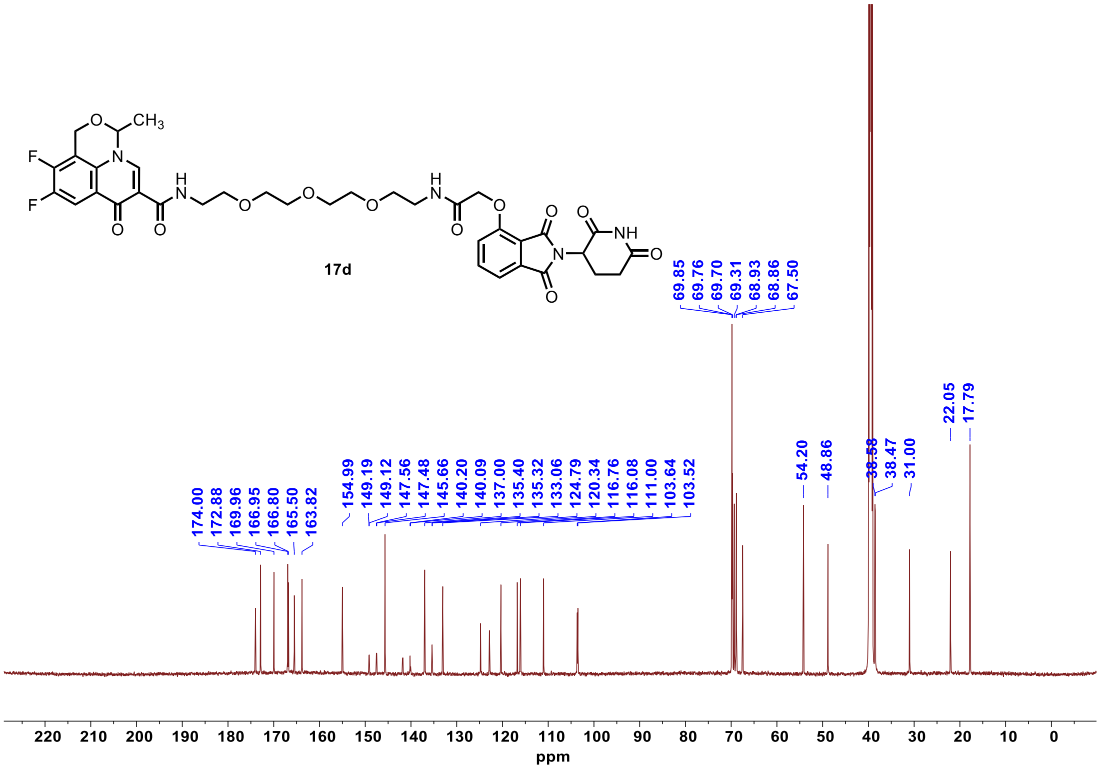

#### $^{1}$H NMR **18a**

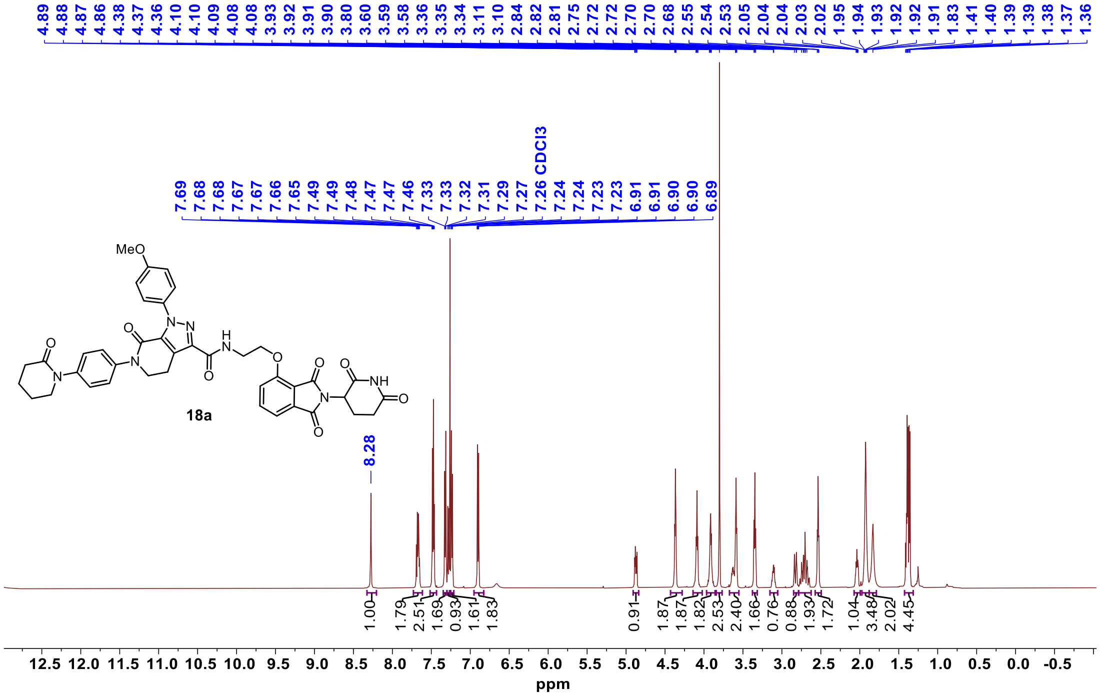


#### $^{13}$C NMR **18a**

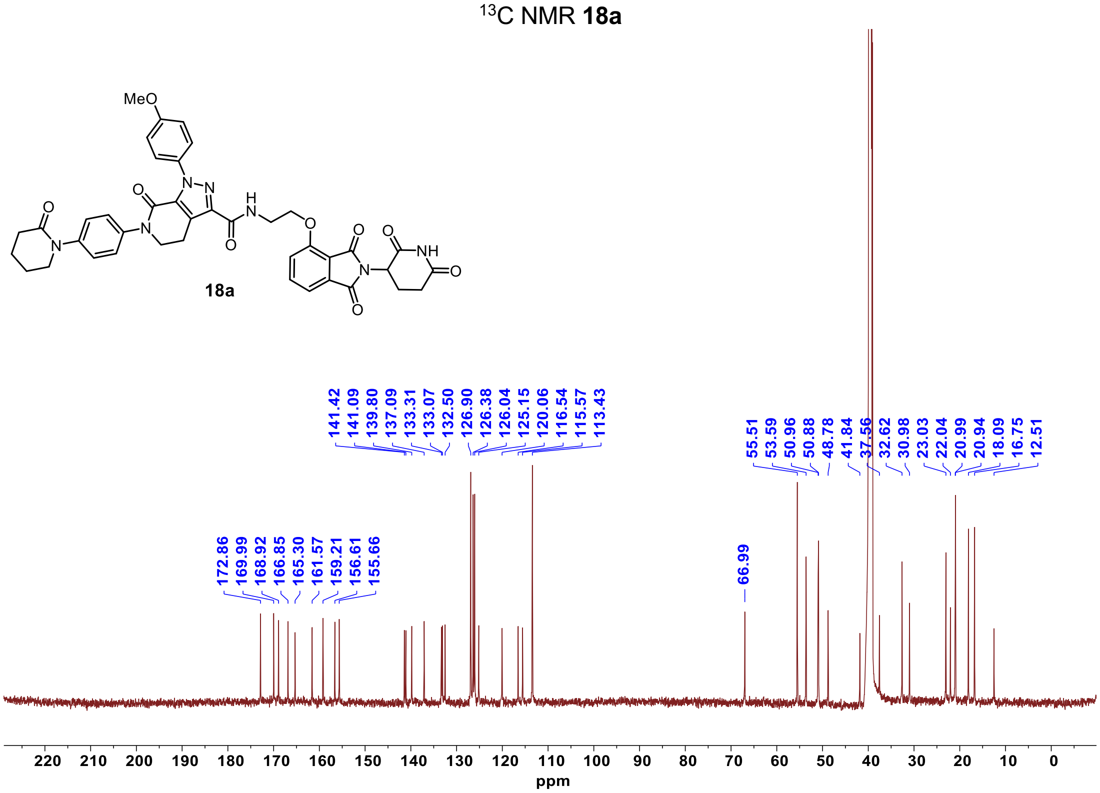

[1]H NMR **18b**

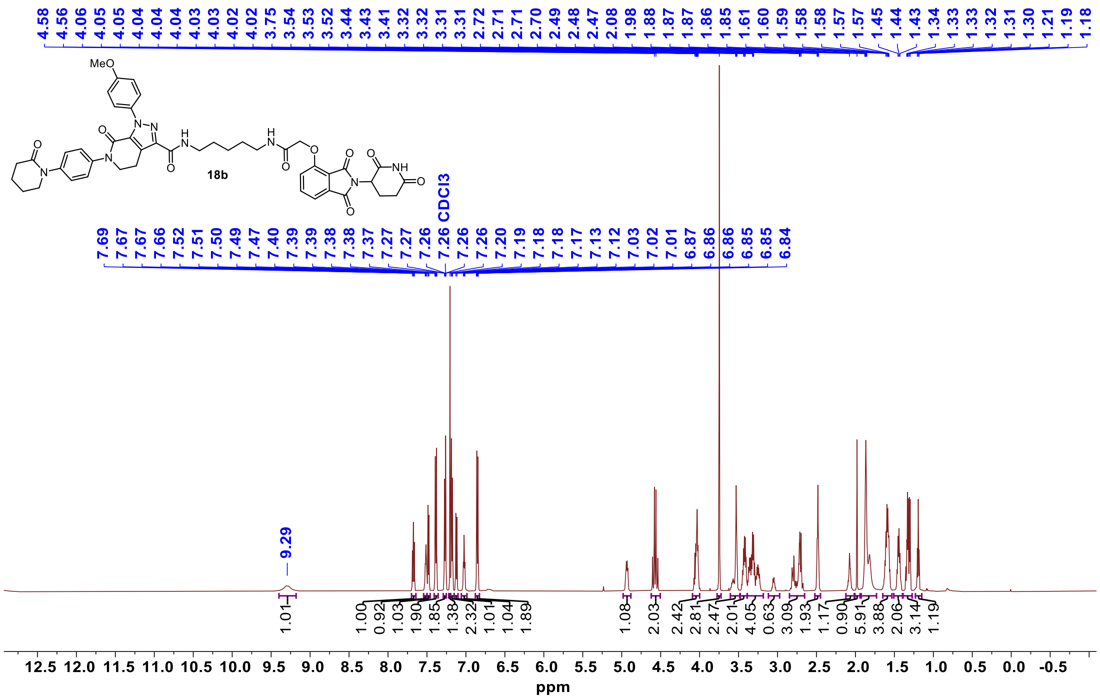


[13]C NMR **18b**

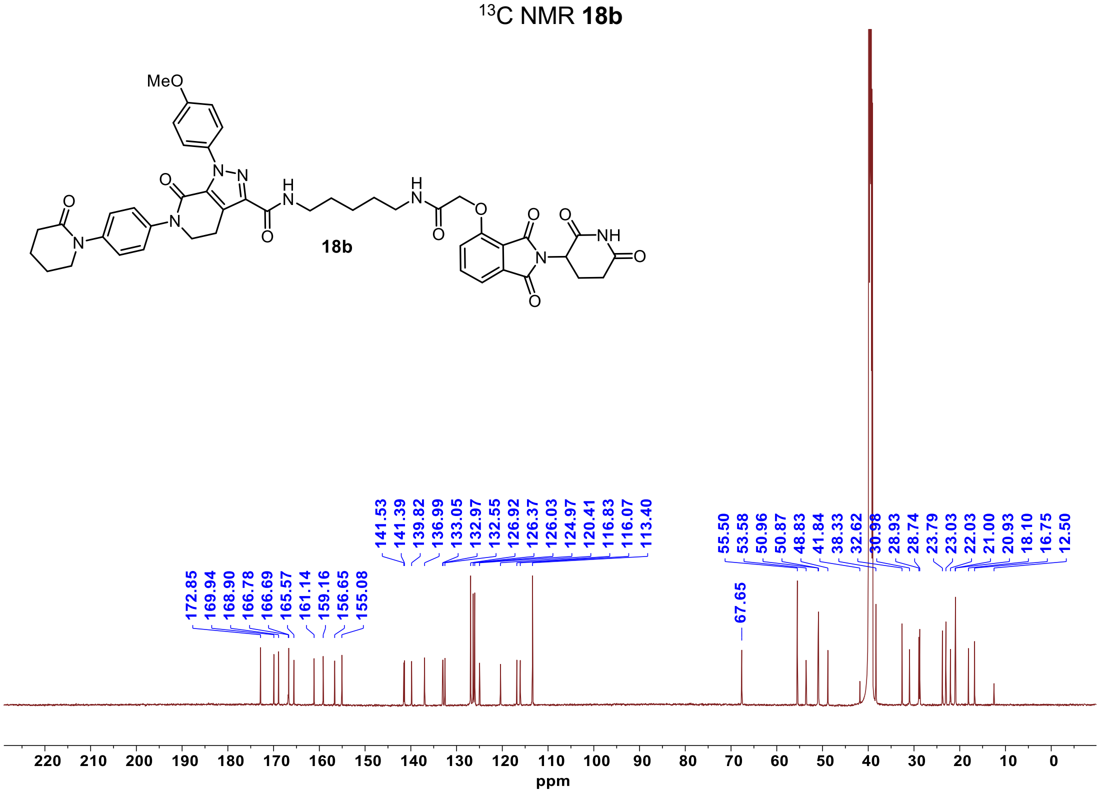

#### [1]H NMR **18c**

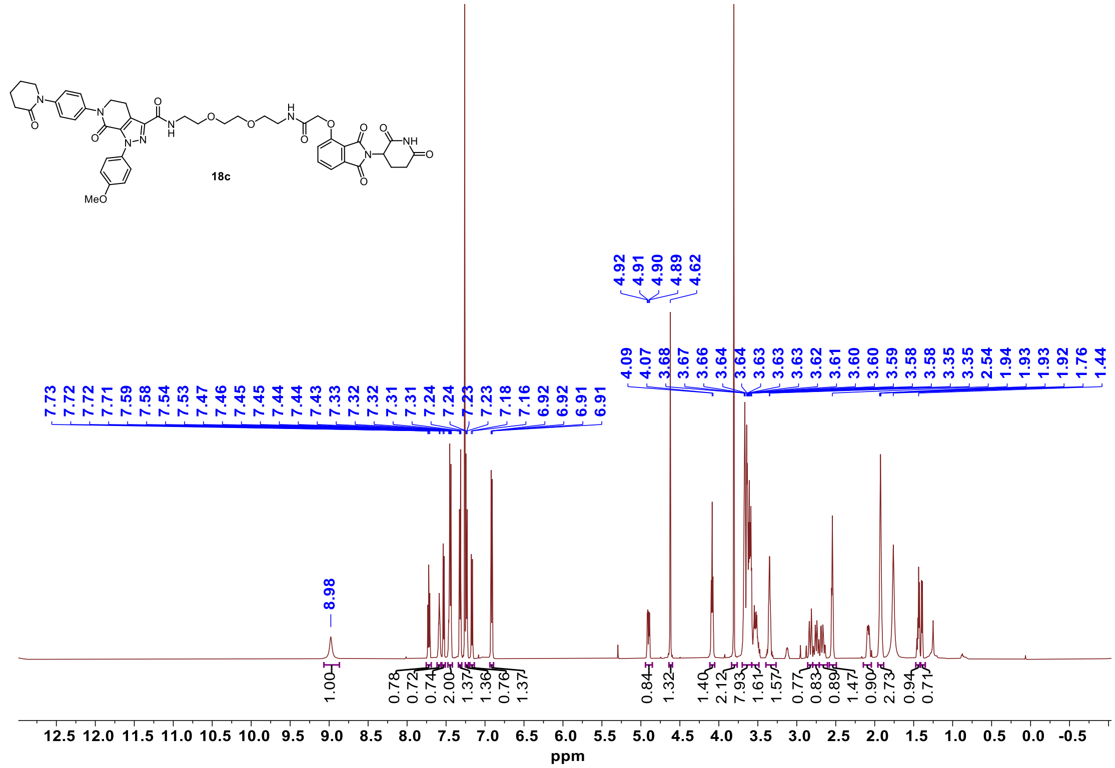


#### [13]C NMR **18c**

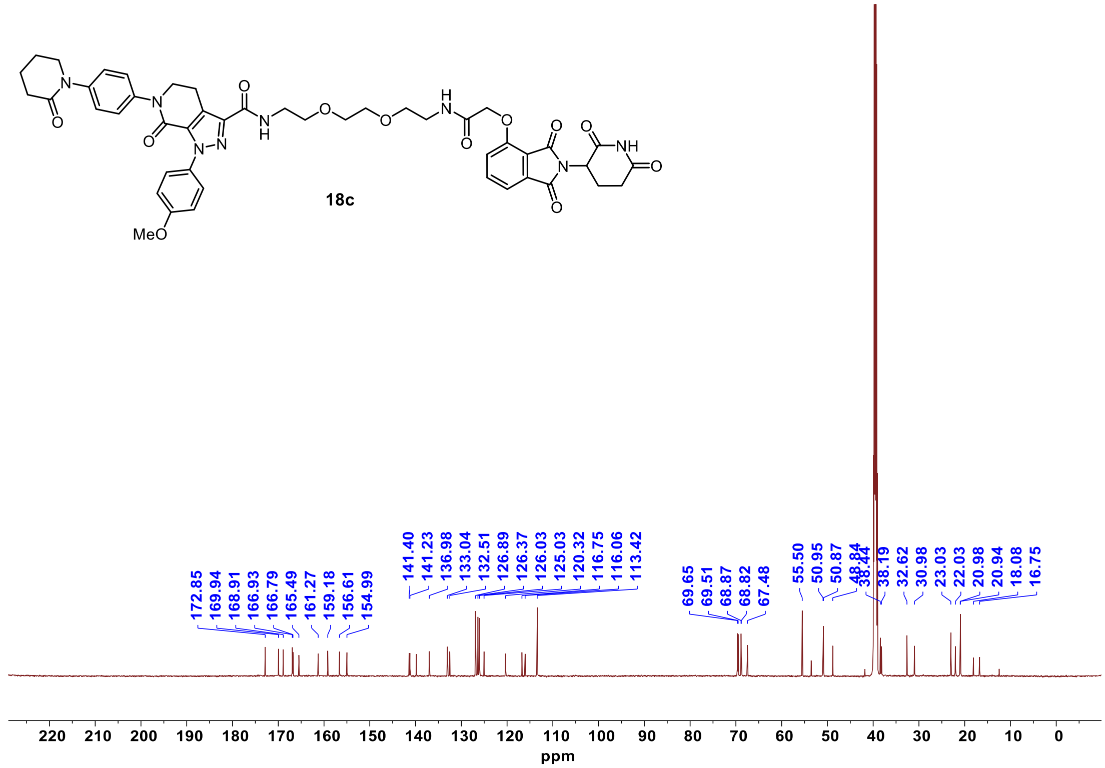

[1]H NMR **18d**

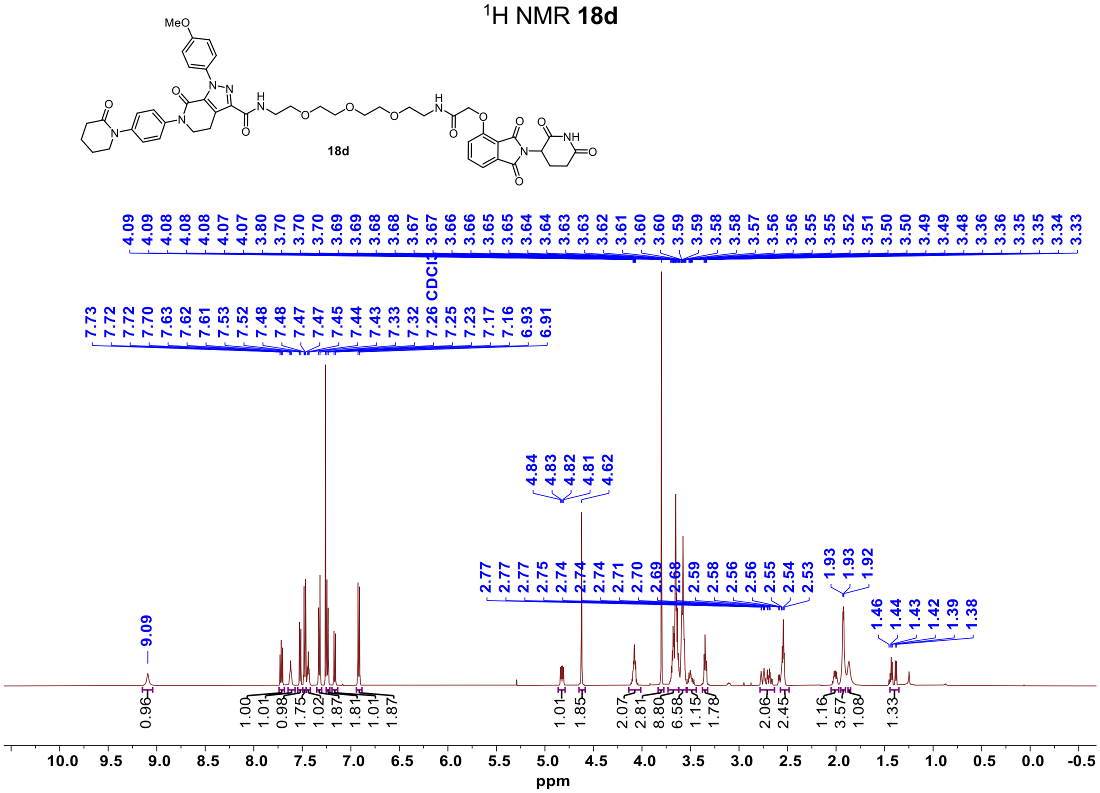


[13]C NMR **18d**

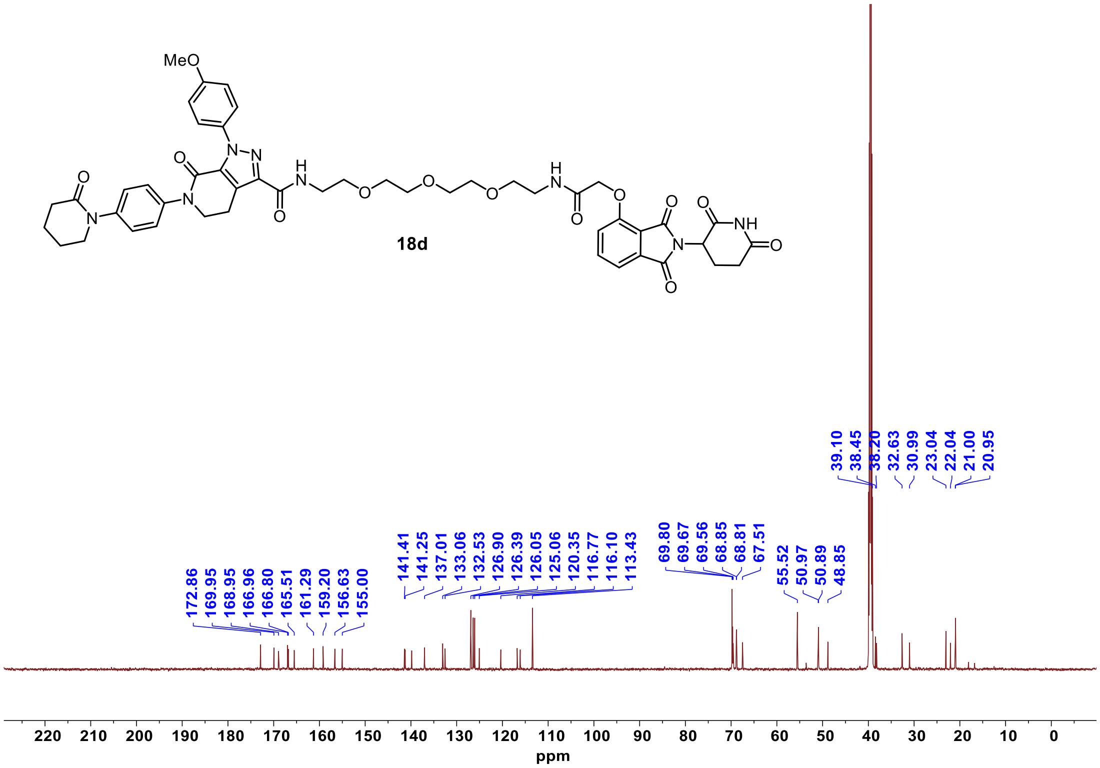
